# Quantitative Principles of *cis*-translational control by general mRNA sequence features in eukaryotes

**DOI:** 10.1101/587584

**Authors:** Jingyi Jessica Li, Guo-Liang Chew, Mark D. Biggin

## Abstract

**BACKGROUND:** General translational *cis*-elements are present in the mRNAs of all genes and affect the recruitment, assembly, and progress of preinitiation complexes and the ribosome under many physiological states. These elements are: mRNA folding, upstream open reading frames, specific nucleotides flanking the initiating AUG codon, protein coding sequence length, and codon usage. The quantitative contributions of these sequence features and how and why they coordinate together to control translation rates are not well understood.

**RESULTS:** Here we show that these sequence features specify 42%–81% of the variance in translation rates in *S*. *cerevisiae, S*. *pombe, Arabidopsis thaliana, M*. *musculus*, and *H*. *Sapiens*. We establish that control by RNA secondary structure is chiefly mediated by highly folded 25–60 nucleotide segments within mRNA 5’ regions; that changes in tri-nucleotide frequencies between highly and poorly translated 5’ regions are correlated between all species; and that control by distinct biochemical processes is extensively correlated as is regulation by a single process acting in different parts of the same mRNA.

**CONCLUSIONS:** Our work shows that the general features control a much larger fraction of the variance in translation rates than previously realized. We provide a more detailed and accurate understanding of the aspects of RNA structure that direct translation in diverse eukaryotes. In addition, we note that the strongly correlated regulation between and within *cis*-control features will cause more even densities of translational complexes along each mRNA and therefore more efficient use of the translation machinery by the cell.

## BACKGROUND

It is a major challenge to determine from nucleotide sequence data the rates at which eukaryotic mRNAs are translated into protein. There are two classes of *cis*-acting elements that determine these rates: general sequence features and gene/condition specific elements [1-11]. The general features are secondary structure in the 5’ portion of the mRNA; upstream open reading frames (uORFs), which lie 5’ of the protein coding sequence (CDS); specific nucleotides immediately flanking the initiating AUG codon (iAUG) at the 5’ of the CDS; CDS length; and codon usage. These five features are each present in all or many mRNAs and function by affecting the formation and progress of preinitiation complexes or the ribosome. As such they act in a wide array of physiological states and tissues. Gene or condition specific elements, in contrast, are short sequences recognized by trans-acting factors such as micro RNAs or regulatory proteins. A given type of specific element is present in only a subset of genes, and its cognate trans-acting factor only functional in a subset of cells or conditions.

Here we focus on the general elements. Our major goal is to estimate the contributions of each of these sequence features—and their associated biochemical mechanisms—to the total variance in translation rates, singly and in combination. We argue that from this one can, in addition, approximate from the unexplained variance the contribution of specific elements, or at least set an upper limit on that contribution. By optimizing models that predict the contribution of each general feature, we also aim to better understand the nucleic acid sequences that encode them. Further, to establish common principles we have analyzed in parallel data from five model eukaryotes that represent yeasts, plants, and animals.

The general sequence features have long been known to affect the translation rates of individual genes. The quantitative contributions of these elements to the variance in genome wide translation rates, however, have not been well characterized. The correlation of subsets of these sequence features with ribosome profiling translation rate data has been estimated in *S*. *cerevisiae* and several animal models [4, 10, 12-16] (Additional file 1: Table S1). In addition, the effect of large numbers of artificial sequence mutations on translation has been determined for 10–50 nucleotide segments proximal to the iAUG codon in *S*. *cerevisiae* and *H*. *sapiens* [12, 17-21] (Additional file 1: Table S2). The role of RNA secondary structure in 5’ untranslated regions (5’UTRs) has also been addressed *in vitro* for *S*. *cerevisae* [22] (Additional file 1: Table S2). A prior study that we conducted was the only one to assess the contributions of all five features [10]. This work established that mRNA folding, uORFs, iAUG proximal sequence elements (APEs), and CDS length explain 58% of the variance in translation rates among *S*. *cerevisiae* genes, presumably due to their impact on the initiation rate. When the effect of codon usage on elongation by the ribosome is also taken into account, 80% of the variance can be explained.

Here we have developed new models that characterize in detail the mRNA secondary structures and sequence motifs that regulate translation rates. By applying our models to five well studied eukaryotes and by examining correlations in control between general features and between different parts of the same mRNA, we identify key principles common to all five species as well as phyla specific differences.

## RESULTS

### Datasets

We chose five model eukaryotes for study: *S*. *cerevisiae, S*. *pombe, Arabidopsis thaliana, M*. *musculus*, and *H*. *Sapiens*. Translation rates were determined from ribosome profiling data as prior work has shown that the density of ribosomes per mRNA (i.e. the translational efficiency) is a useful estimate of the rate [13, 23]. For each species, we chose an example dataset to analyze in most detail [4, 13, 24-26]. In addition, key analyses were repeated on further datasets of three species for other tissues or biological replicas [4, 14, 27, 28]. Figure 1 presents the distributions of translation rates, 5’UTR length, and CDS lengths for the five example datasets. Additional file 1: Figure S1 shows the variation in translation rates for all ten datasets examined. The lengths of 5’UTRs vary among the species, with the average for *S*. *cerevisiae* being the shortest and that for the two mammals and *S*. *pombe* the longest. By contrast, the ranges of CDS lengths are more consistent among the five eukaryotes. The variance in translation rates is relatively narrow for all species and conditions. The mRNA sequences, translation rates, and other primary information for each dataset are given in Additional file 2. Table 1 provides a glossary of abbreviations used.

**Figure 1.**
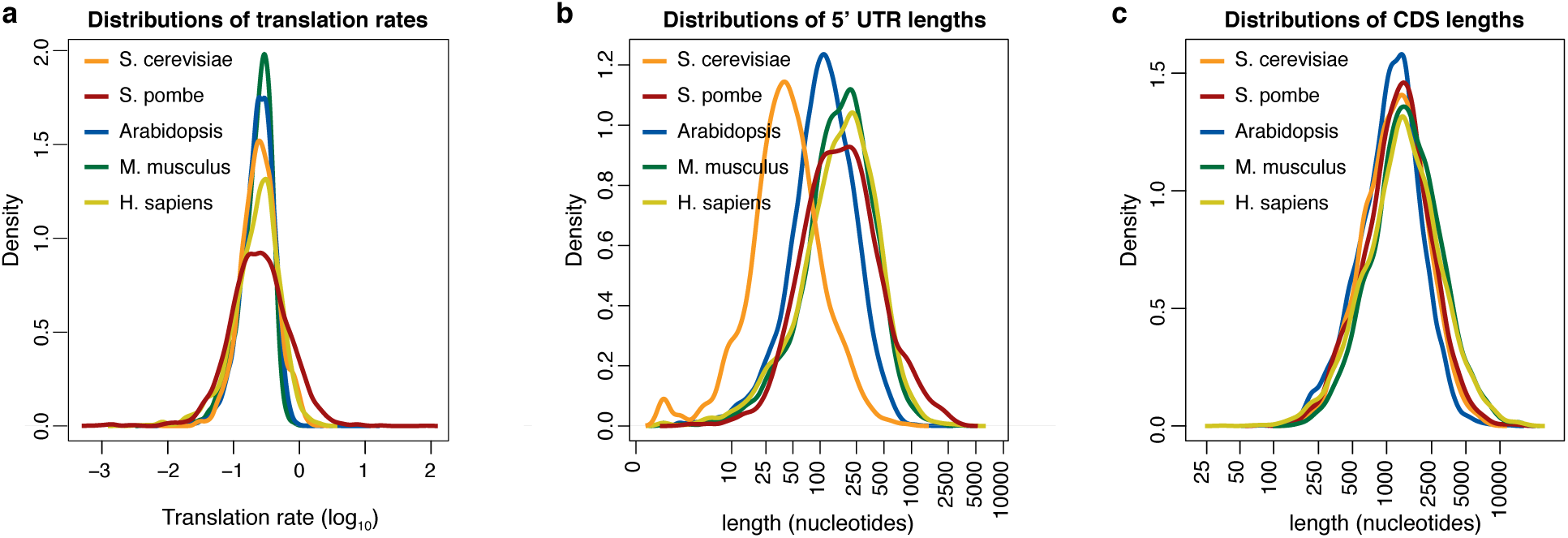
Example datasets for five eukaryotes. (a) The distribution of translation rates for example datasets representing five eukaryotes. Translation rates are defined by the density of ribosomes per mRNA molecule. The log_10_ transformed data have been scaled to have the same median while retaining their original variance. (b) The distributions of lengths of 5’ untranslated regions (UTRs). (c) The distributions of protein coding sequence (CDS) lengths.

**Table 1.**

| **Abbreviation** | **Explanation** |
| --- | --- |
| **APE** | AUG Proximal Element: control sequences that flank the iAUG |
| **dAPE** | downstream APE: that part of the APE located downstream (3') of the iAUG |
| **uAPE** | upstream APE: that part of the APE located upstream (5') of the iAUG |
| **5'ofAPE** | the 5'UTR sequences that lie 5' of the APE |
| **iAUG** | initiating AUG: the AUG codon at the 5' end of the CDS |
| **uAUG** | upstream AUG: an AUG codon in the 5' UTR, i.e. an AUG at the 5' end of an uORF |
| **BIC** | Bayesian Information Criteria: a method to select an efficient set of model features |
| **5' cap** | the 5' most nucleotide of the mRNA |
| **CDS** | coding sequence: the protein coding portion of the mRNA |
| **CDS 3'ofAPE** | that part of the CDS that lies 3' of the APE |
| **MFE** | minimum free energy: the Gibbs free energy of folding for a defined segment of RNA |
| **uORF** | upstream open reading frame: an uAUG containing open reading frame in the 5'UTR |
| **PWM** | Position Weight Matrix: the fraction of A, U, G and Cs at each nucleotide position |
| **5'region** | the region spanning the 5'UTR and the 5' most ~30 nucleotides of the CDS |
| **5'UTR** | 5' untranslated region: all mRNA nucleotides 5' of the CDS |
| **TR** | translation rate: rate of translation per mRNA molecule at steady state |
| **high TR** | the 10% of genes with the highest translation rates |
| **low TR** | the 10% of genes with the lowest translation rates |

### Control by 5’ mRNA secondary structures

Biophysical measurements of short RNA oligonucleotides *in vitro* have allowed the melting temperatures and Gibbs free energies of stem/loop structures to be accurately calculated for any short sequence based on a set of rules for the energies of base pairing and adjacent base stacking as well as the destabilizing entropic effect of loop length [29-32]. For example, the coefficient of determination correlation coefficient between measured and predicted Gibbs energies for 109 4– 14 bp double stranded RNA oligonucleotides from [33] is R^2^=0.98 (Additional file 3). Computational predictions of mRNA secondary structure that incorporate these and other information are quite effective: the algorithm we employed having 74% recall (% of true positive base pairs identified) and 79% precision (% of base pairs identified that are true positives) [32]. Previous work has shown that the free energies of RNA folding from this or similar algorithms positively correlate with translation rates (Additional file 1: Tables S1 and S2), supporting the view that the preinitiation complex and ribosome must unfold RNA structures in order to progress down the mRNA. Our preliminary analyses, however, showed that these prior studies had greatly underestimated the contribution of mRNA folding to translation. We therefore systematically assessed which aspects of secondary structure are most important in determining translational efficiency and used this information to improve predictive models of rates.

Figure 2 shows the mean and the [25^th^ percentile—75^th^ percentile] interval for the free folding energies of 35 nucleotide windows for two cohorts of genes: the 10% of genes with the highest translation rates (high TR) and the 10% with the lowest (low TR). Within the 5’UTR, the low TR cohort have smaller folding energies (i.e. are more folded) than the high TR cohort, consistent with repression of preinitiation complexes by RNA structure. For *S*. *cerevisiae, S*. *pombe*, and *Arabidopsis* the differences between the two cohorts are largest from −35 to +35, −35 to +1, and - 120 to +35 respectively. For mouse and human, the differences are largest towards the 5’ cap. In the CDS region 3’ of +30, the relationship of RNA folding energy and translation rate is not consistent, with either a strong negative correlation in *S*. *pombe*; a weak negative correlation in *S*. *cerevisiae, Arabidopsis*, and *M*. *musculus*; or a positive correlation in *H*. *sapiens*. Prior analysis in *S*. *cerevisiae* implied that strongly folded RNA structures in the CDS are associated with more stable mRNAs and that the negative correlation of CDS folding energy with translation is indirect, being due to the positive correlation between translation and mRNA abundance and the negative effect of RNA turnover on mRNA abundance [10]. Given this and a lack of evidence that RNA structure affects the elongating ribosome, we have limited our models to the 5’UTR and the 5’ most part of CDS, where folding energy values and translation rates correlate positively. We term the combination of the 5’UTR and the short 5’ CDS segment the 5’ region.

**Figure 2.**
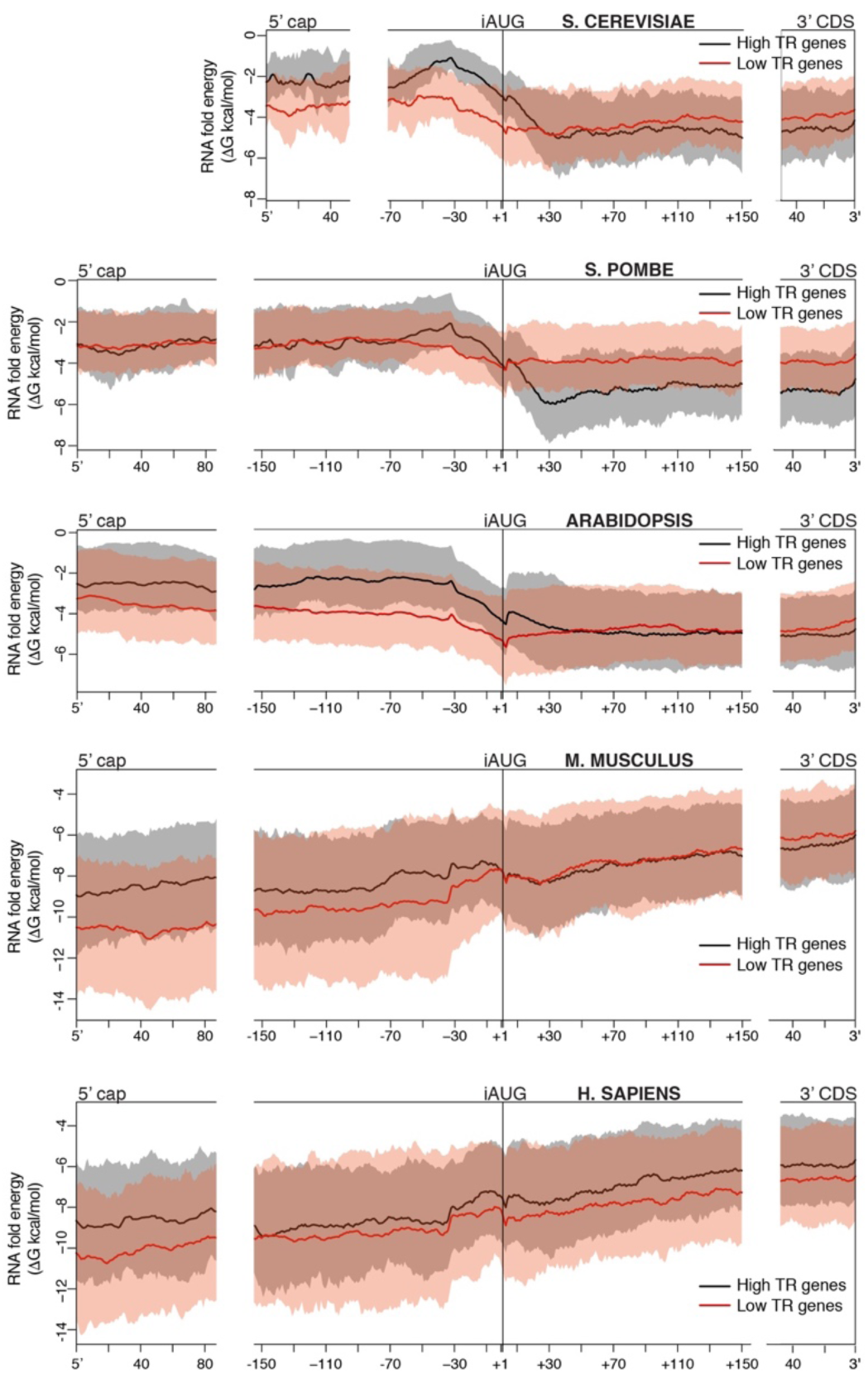
Different distributions of secondary structures in the 5’ portions of mRNAs. The predicted RNA folding energy (ΔG kcal/mol) of 35 nucleotide windows (*y*-axis) are plotted for the 10% most highly translated mRNAs (high TR, black) and the 10% most poorly translated (low TR, red). The *x*-axis shows the position of the 5’ most nucleotide of each window. Windows for every one nucleotide offset were calculated. At each location the free energies for the mean (continuous lines) and the interval between the 25^th^ and 75^th^ percentile (shading) are shown. mRNAs were aligned at their 5’ cap (left), at the iAUG (center), or at the 3’ of the CDS (right). In *S*. *cerevisiae, S*. *pombe* and *Arabidopsis*, the differences between the two cohorts are greatest proximal to the iAUG. In the two mammals, the differences are greatest towards the 5’ cap.

Along the 5’ regions of individual mRNAs free energy values vary dramatically (Additional file 1: Figure S2), reflecting mRNA stem/loop structures at some locations and unfolded regions at others. Despite the mean tendencies shown in Figure 2, Additional file 1: Figure S2 reveals that individual genes can have folded or unfolded regions at almost any location.

To capture and exploit these complex distributions, we devised a number of features that each score every gene using some aspect of predicted RNA folding energy. We also constructed three multivariate linear models that combine multiple features to form feature-sets. The coefficient of determination correlation coefficient (R^2^) was then calculated between each feature or feature-set and the translation rates (Figure 3; Additional file 4). One feature was defined as the folding energy for the contiguous sequence from the 5’ cap to +35 (“whole”). The other features were determined for each of 19 window lengths varying from 6 to 100 nucleotides, the 5’ ends of the windows tiling from the 5’ cap to −1 (Figure 3). Our most accurate prediction of translation derived from a multivariate linear model that was selected by Bayesian Information Criteria (BIC) and that contains 9–33 features of whichever window length(s) provided the most useful information (“RNAfold”, Figure 3; Additional file 5). These “RNAfold” feature-sets explain between 11%–33% of the variance in rates and as such have over twice the predictive power (R^2^) of models used by other groups (Additional file 1: Table S1).

**Figure 3.**
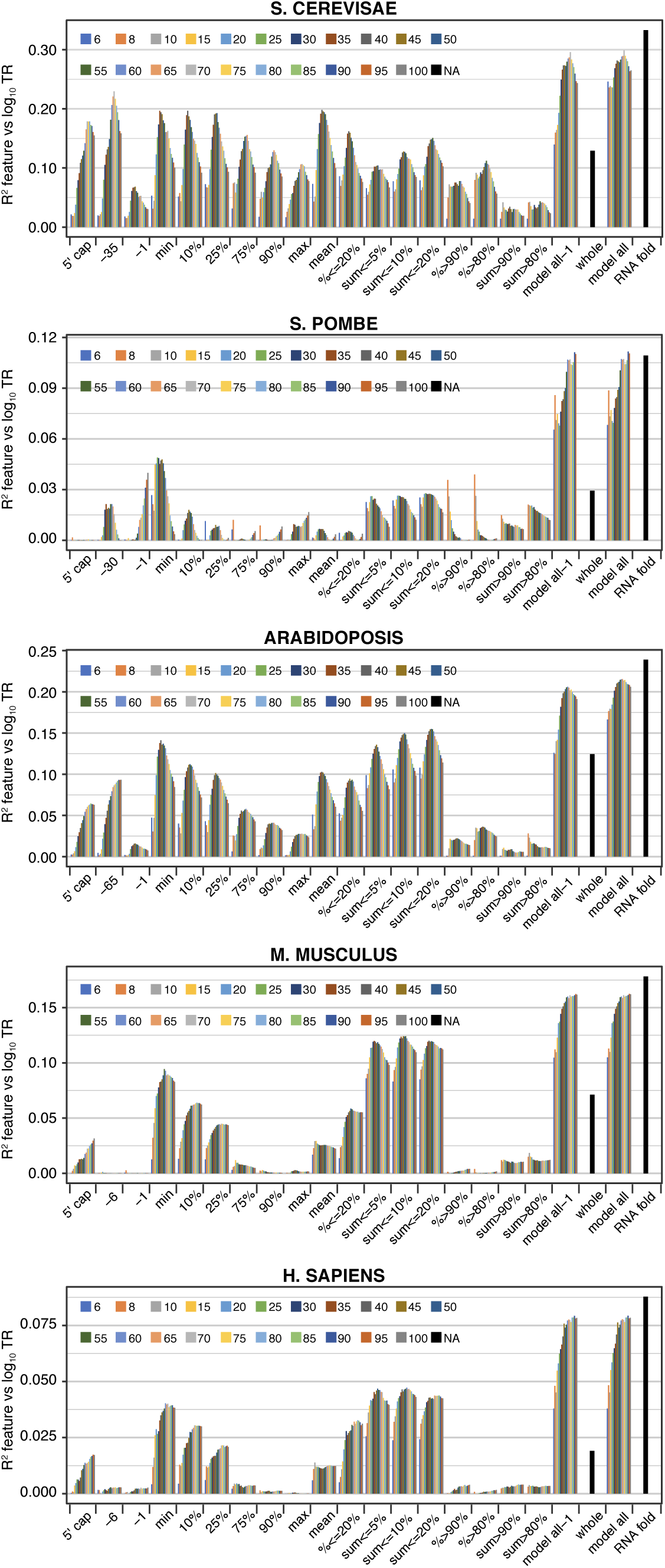
Translation rates are optimally determined by a combination of RNA folding features. The R^2^ coefficients of determination between log_10_ translation rates (TR) and a variety of single features and feature-sets based on RNA folding energies. Nine features use the free energy value of a single window per gene (left): windows “5’ cap” thru “-1” were identified by the location of their 5’ most nucleotide, while windows “min” thru “max” were identified by the rank of their free energy. Nine additional features were each calculated using multiple windows per gene (center): “mean”, the mean of all windows in a gene; “%≤20%”, “%≥=90%”, and “% ≥80%”, the percent of windows in a given gene that respect a threshold on the free energy of all windows for all genes; and “sum≤5%”, “sum≤10%”, “sum≤20%”, “sum≥80%”, and “sum≥90%”, the sum of free energies of windows in a given gene that respect a threshold on the free energy of all windows for all genes. 19 window lengths from 6 to 100 nucleotides were employed for each of the above features. In addition, a final feature, “whole” (right), was calculated from the free energy of folding of the contiguous sequence from the 5’cap to +35 for each mRNA. Three multivariate –feature-sets were also determined (right): “model all-1” combined all window based features for a given window length; “model all” included in addition the “whole” feature; while “RNAfold” used Bayesian Information Criteria to select features from the complete set of features, using whichever window length variant(s) provided the most useful information. “RNAfold” was additionally constrained for *S*. *pombe* by removal of all windows that extend 3’ of +30 nucleotides to avoid sequences showing a strong negative correlation of free energy values and TR. Additional file 4 provides the R^2^ values for all features. Additional file 5 provides details of the features selected for the “RNAfold” model.

Most features employed windows identified based on their free energy values. A substantial proportion of control by RNA secondary structure—often greater than a half—can be explained by the single window that has the minimum free energy within each mRNA (i.e. by the window that has the most folded structure) (“min”, Figures 3 and 4). Less folded windows each have progressively reduced explanatory power compared to the “min” window (“10%”, “25%”, “75%”, “90%”, and “max”, Figure 4). Likewise, features that sum Gibbs energies for the most folded window with the energies of other strongly folded segments (“sum≤5%”, “sum≤10%”, or “sum≤20%”) have more explanatory power than features that sum the energies of less well folded segments (“sum>90%” and “sum>80%”) (Figure 3). A similar relationship is also seen for features that determine the percent of windows in each mRNA that pass some threshold on the energies of all windows in the dataset (“%≤20%” vs “%>80%” and “%>90%”) (Figure 3). These results collectively indicate that translation rates are much more strongly influenced by differences among the more folded segment(s) within 5’ regions than by differences among their less folded sequences.

**Figure 4.**
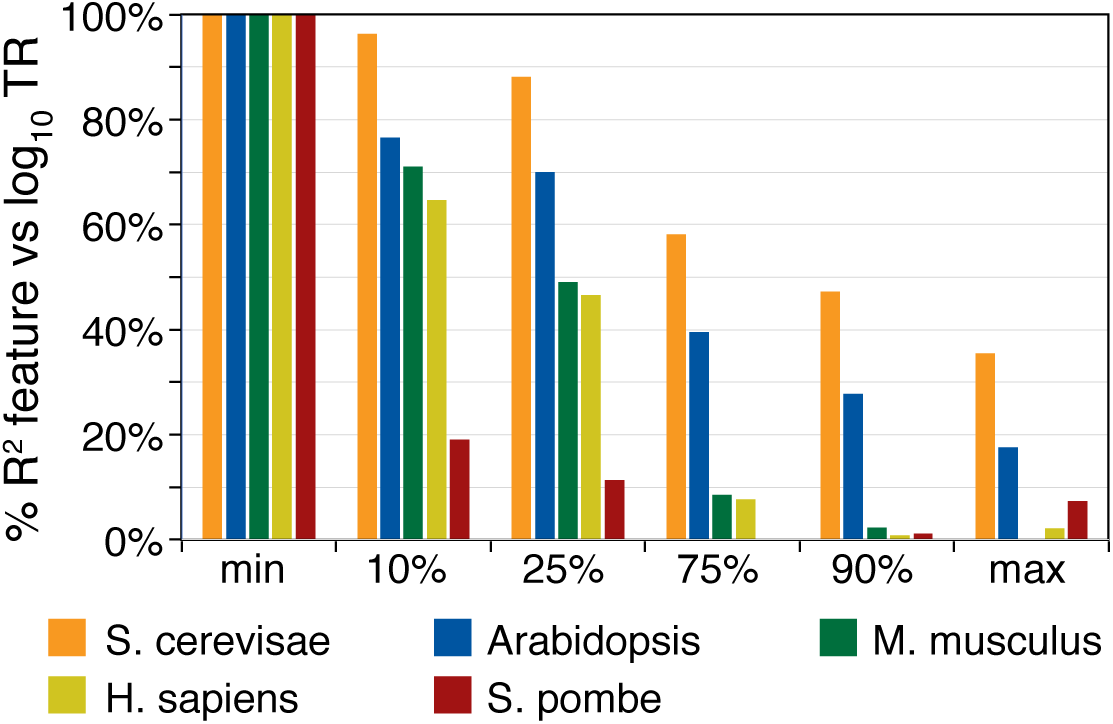
The most folded window controls translation more strongly than less folded windows. The correlation between log_10_ translation rates (TR) and a single windows within mRNA 5’ regions. Six windows were selected based on the rank of their free energy: “min”, “10%”, “25%”, “75%”, “90%” and “max”, where “min” is the most folded (smallest free energy) and “max” the least folded. The values plotted are the R^2^ coefficients of determination expressed as a percent of the R^2^ coefficient for the “min” window. The window length was the optimum for each species: *S*. *cerevisiae*, 35; *Arabidopsis*, 40; and *M*. *musculus*, 55; *H*. *sapiens*, 60; and *S*. *pombe*, 25.

Additional results imply that base-pairing between sequences separated by more than 60 nucleotides does not play a dominant role in regulating translation. Because the “whole” feature includes the full 5’ region, it accounts for all long range interactions. Models that contain all features, however, perform little better than ones lacking “whole” (Figure 3, compare “models all” to “models all-1”).

To further define the secondary structures that control translation, we focused on the most folded window as these are strongly predictive in all species. The optimum length of “min” window ranges from 25–35 nucleotides in the two yeasts to 55–60 in the two mammals (Figure 3). Windows shorter than the optimum are less predictive presumably because they exclude important secondary structures; whereas windows longer than the optimum include unfolded sequences that “dilute” the control information. The optimum-length “min” windows generally contain either one or two stem/loops of varying sizes, some of which include mismatched or single nucleotide bulges within their stems (Additional file 6). For all species, the longest contiguous run of paired nucleotides within a stem contains much of the regulatory information (Figure 5). With the exception of *S*. *pombe*, however, the total number of nucleotide pairs in the window is more predictive of translation than the number of nucleotide pairs in the longest contiguous run or in the longest stem (Figure 5). This suggests that most nucleotide pairs in “min” windows are controlling, including those within smaller secondary-stem/loops. The different result for *S*. *pombe* may well be because its “min” windows have shorter optimum lengths than those of other species (25 vs 35–60 nucleotides) and tend to have only one major stem/loop (Additional file 6). Finally, It is striking that between cohorts of highly translated and lowly translated genes, while free energies and loop sizes vary significantly, the number of nucleotide pairs in contiguous runs, longest stems, or full windows are similar for all five species (Figure 6).

**Figure 5.**
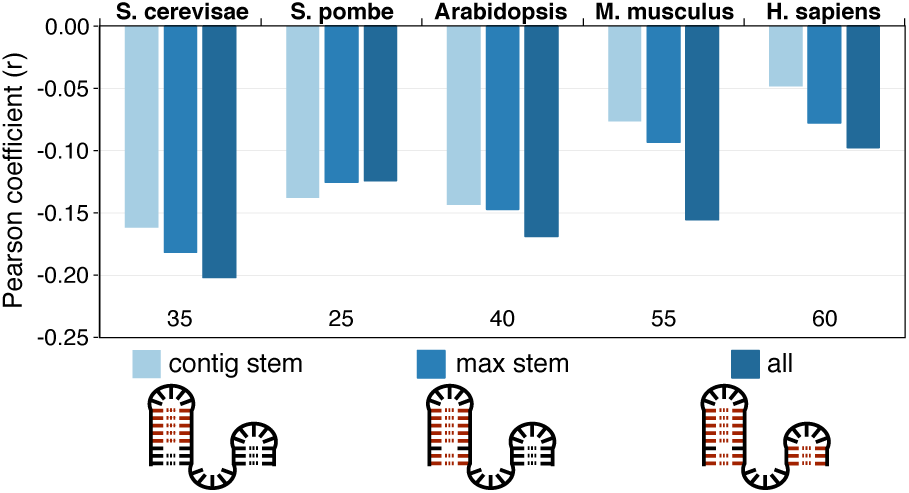
Number of nucleotide pairs describe the regulatory potential of the most folded window. The number of paired nucleotides was calculated in the most folded (“min”) window of each gene for the longest contiguous stem without mismatches or single nucleotide bulges (contig stem, light blue); the longest stem (max stem, mid blue); and for all pairs within the window, including those not in the longest stem (all, dark blue). The Pearson correlation coefficients between each of these three measures and log_10_ translation rates are plotted on the *y*-axis for the five example datasets. The lengths of the most folded widows are also given. For all but *S*. *pombe*, the correlation coefficients are more negative for measures that include more nucleotide pairs. The primary data are provided in Additional file 6.

**Figure 6.**
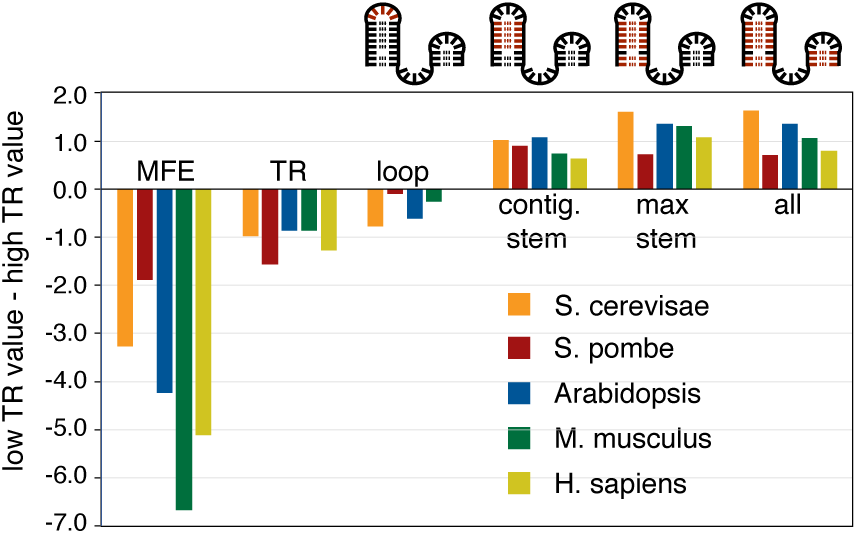
The number of nucleotide pairs controlling translation are similar across diverse eukaryotes. Metrics for “min” windows in the 10% most highly translated mRNAs (high TR) were subtracted from metrics for “min” windows in the 10% most poorly translated mRNAs (low TR), *y*- axis. The metrics are the means of minimum free energy (MFE); log_10_ translation rate (TR); number of unpaired nucleotides linking the longest stem (loop); number of nucleotide pairs in the part of the longest stem that contains no mismatches or single nucleotide bulges (contig. stem); number of all nucleotide pairs in the longest stem (max stem); the total number of nucleotide pairs in the “min” window (all). See Additional file 6 for the primary data. The distributions of the total number of “min”-window nucleotide pairs in high and low TR cohorts are shown in Additional file 1: Figure S3.

We also defined a location-specific group of features that use the fold energy of a single window whose 5’ end is at a defined location: either at the 5’ of the mRNA (“5’cap”) or at a location relative to the iAUG (“-65” to “-1”) (Figure 3). These features give results consistent with the distributions of RNA folding energies shown in Figure 2. For example, in *M*. *Musculus* and *H*. *sapiens* features using windows located at the 5’ cap have more predictive power than those at or near the iAUG. While most of features selected by BIC for the “RNAfold” feature-set used windows identified by their free energies, several others belonged to the location-specific group (Additional file 5). By combining these two feature classes, we are able to more fully capture how mRNA secondary structure directs translation.

### Sequence motifs in 5’ regions

One feature we used previously to explain translation rates in *S*. *cerevisiae* exploited sequence differences between highly and poorly translated mRNAs without regard to the biochemical mechanism(s) of control [10]. This approach proved powerful, showing that nucleotides flanking the iAUG from −35 to +28—a region termed the AUG proximal element (APE)—explain a third of the variance in rates [10]. To extend this strategy to the other four species, we first compared Position Weight Matrices (PWMs) for sequences −80 to +35 for the most highly translated (high TR) and most poorly translated (low TR) 10% of genes (Figure 7a; Additional file 7). The *S*. *pombe* and *Arabidopsis* PWMs resemble those of *S*. *cerevisiae*: for example, A nucleotides are enriched 5’ of the iAUG in high TR genes vs low TR genes, while Gs are depleted. *M*. *musculus* and *H*. *sapiens* mRNAs, by contrast, are GC rich, and sequence differences between high and low TR cohorts are less readily apparent. Following the strategy used earlier for *S*. *cerevisiae* (see Materials and Methods), we assigned a score to each gene based on PWMs of varying lengths in high TR genes and defined APE boundaries by maximizing the R^2^ between the PWM scores and translation rates (Figure 7b; Additional file 1: Figure S4). *Arabidopsis* has, like *S*. *cerevisiae*, an extended APE, spanning nucleotides −65 to +33, whereas *S*. *pombe* and the two mammals have shorter APEs that span −6 to +13 or less.

**Figure 7.**
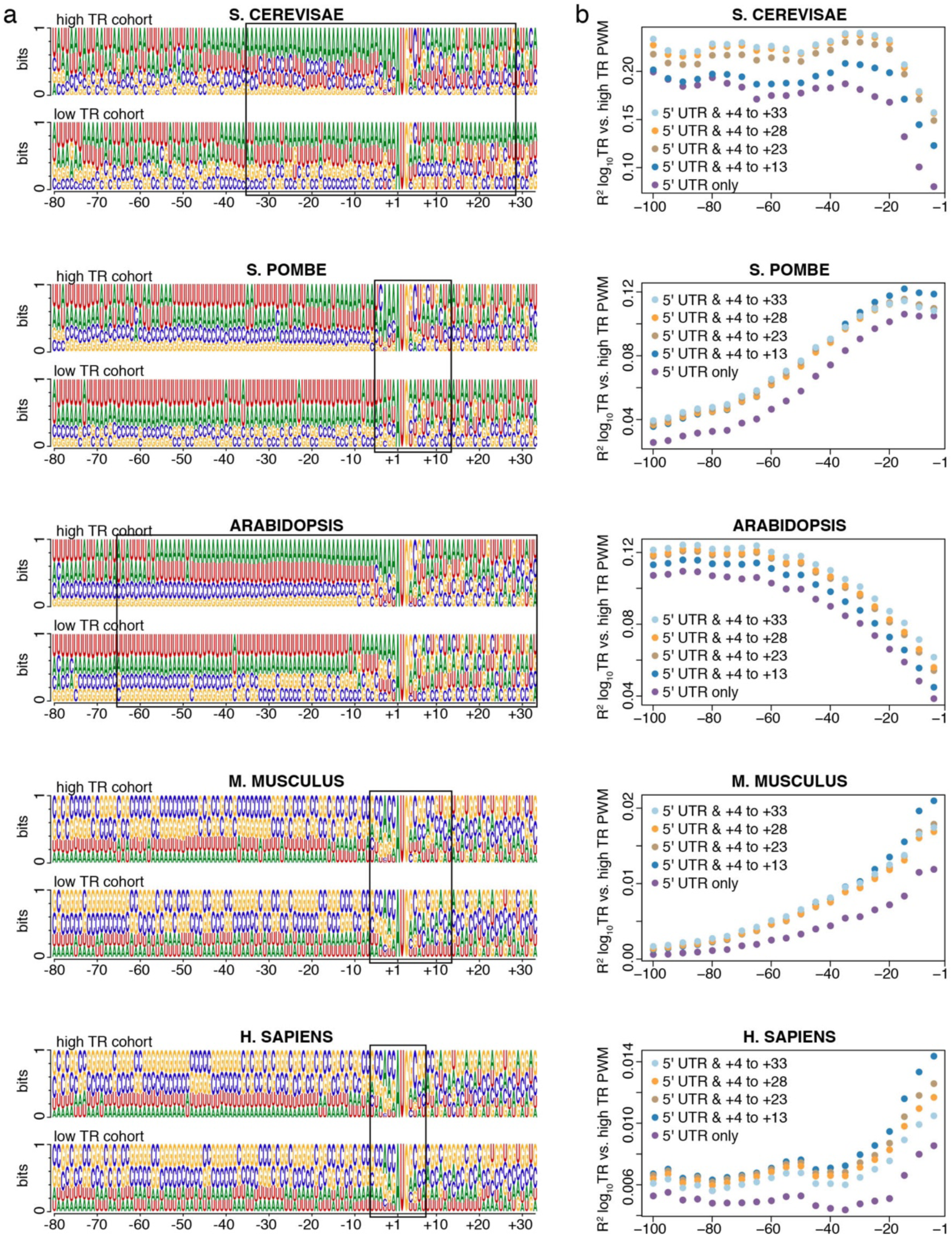
The 5’ and 3’ boundaries of AUG proximal elements. (a) Position Weight Matrices (PWMs) for the 10% of mRNAs with the highest translation rate (high TR cohort) and the 10% with the lowest rate (low TR cohort). Sequence logos show the frequency of each nucleotide at each position relative to the first nucleotide of the iAUG. The location of the AUG proximal element (APE) is indicated with a black box. (b) The R^2^ coefficients of determination between log_10_ translation rates (TR) and PWM scores. PWMs of varying lengths were built from the sequences of the high TR cohort, the PWMs extending 5’ from −1 in 5 nucleotide increments and extending 3’ from +4 in 5 or 10 nucleotide increments. Log odds scores were then calculated for all mRNAs that completely contained a given PWM. Additional file 1: Figure S4 shows more detailed mapping of the 5’ and 3’ boundaries for *M*. *musculus* and *H*. *sapiens*.

To increase these PWM based models’ predictive power, we used BIC to select subsets of di- and tri-nucleotides whose frequencies in each mRNA most strongly explain translation. Di- and tri-nucleotide frequency features were selected for the portion of APE upstream of the iAUG (uAPE) and separately for the portion downstream of the iAUG (dAPE), 4–16 di- and tri-nucleotides being selected per region (Additional file 1: Figure S5, Additional file 5). In the resulting multivariate models, *S*. *cerevisiae, S*. *pombe* and *Arabidopsis* APEs exert stronger control than their mammalian counterparts, the former explaining 16%–33% of the variance in translation, the latter only 3–4% (Additional file 1: Figure S5).

Because the above models capture only part of the 5’ region, we also examined motifs in the area that extends from just 5’ of the APE to the 5’ cap (5’ofAPE). We found that an effective model for this 5’ofAPE region included a five nucleotide length PWM at the 5’ cap and a subset of di- and tri-nucleotide frequencies for the full region that was selected using BIC (Additional file 5; Additional file 1: Figure S6). In non-mammalian species 5’ofAPE sequences are much less predictive of translation than the APE, whereas in mammals the reverse is found (Figure 8). Multivariate models that combine 5’ofAPE and APE motifs provide our most accurate prediction of translation rates using 5’ region sequence motifs (feature-set “5’motifs”, Figure 8).

**Figure 8.**
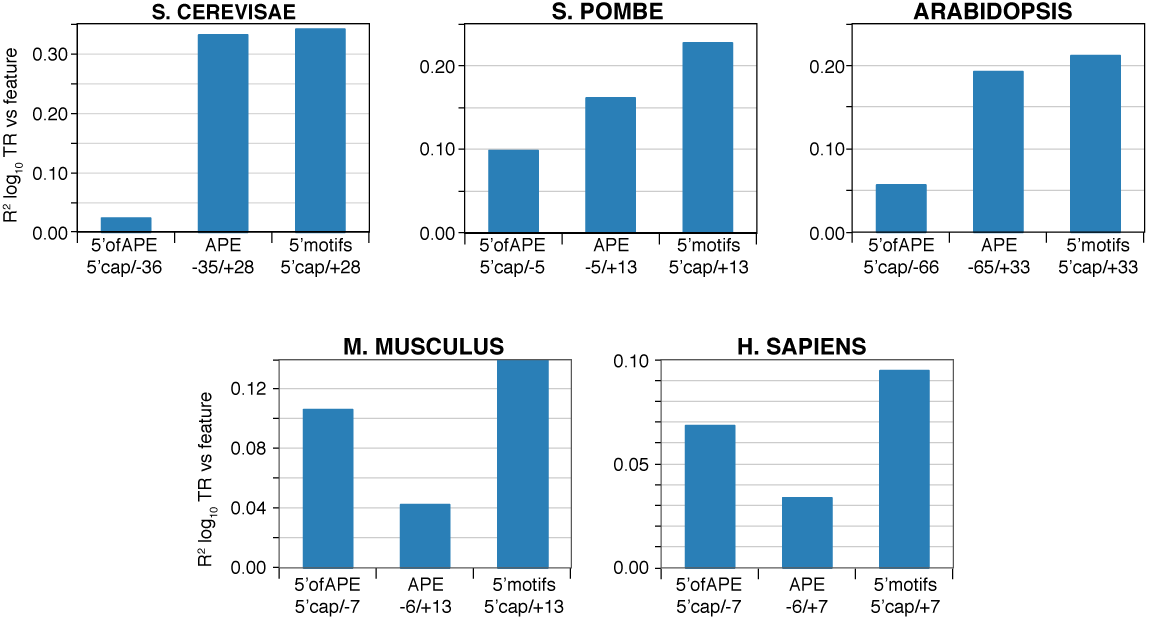
AUG proximal elements (APEs) and the sequences 5’ of these elements (5’ofAPEs) are differently important in mammals and non-mammalian eukaryotes. The R^2^ coefficients of determination between log_10_ translation rate (TR) and models describing the APE; 5’ofAPE; and the combination of these two models (5’motifs). The models are described in Additional file 5.

To determine if control sequences are similar within and between species, we divided each mRNA into four parts: 5’ofAPE, uAPE, dAPE, and the CDS 3’ of the APE (CDS 3’ofAPE). We then calculated the frequencies of each tri-nucleotide in each region of each gene for the high TR and low TR cohorts separately and took the ratio of the means of these values, yielding “high TR / low TR ratios” (Additional file 8). Six results stand out:

i. The (high TR / low TR) ratios in 5’ UTRs vary from 0 to 3.6, with most ratios differing markedly from one (Figures 9 and 11; Additional file 1: Figure S7; Additional file 8). This suggests that a high proportion of all nucleotides in these regions contribute to translational regulation and that the tri-nucleotides whose ratios differ most from one make the largest contributions.
ii. In 5’UTRs, AUG has one of the smallest (high TR / low TR) ratios of any tri-nucleotide (mean (high TR / low TR ratio) = 0.31), showing that our motifs detect the inhibition of initiation at the CDS that results from translation of uORFs (Figure 9; Additional file 1: Figure S7 and Additional file 8).
iii. Within each of the two species that have extended APE sequences—*S*. *cerevisiae* and *Arabidopsis*—the uAPE and dAPE share strong sequence similarities, as shown by the positive correlation between their (high TR / low TR) tri-nucleotide ratios (r=+0.41 and +0.70; Figure 10).
iv. As expected, the CDS 3’ofAPEs show little similarity to 5’UTRs (r<0.1; Figure 10).
v. Some of the strongest (high TR /low TR) ratio correlations are between the uAPEs of the three non-mammals (r>0.65) and separately between the 5’ofAPEs of *M*. *musculus* and *H*. *sapiens* (r=0.8) (Figure 11). There are also, however, lower but clear correlations between the 5’ regions of mammalian and non-mammalian species (Figure 11). Hence, while the mRNA GC contents of these two groups of eukaryotes differ greatly, the variations in 5’ region sequences that direct translation are related across the five species.
vi. The CDS3’ofAPE shows a positive correlation in (high TR /low TR) tri-nucleotide ratios between all species except *H*. *sapiens*, which shows a negative correlation with the other species (Figure 11). This probably reflects a previously characterized negative correlation in tRNA abundances—and as a consequence in optimum codon usage—between cancerous cell lines and other cell types [34] rather than a difference between species as the *H*. *sapiens* dataset was derived from cancerous HeLa cells.

### A model for control by 5’ regions

mRNA 5’ regions control translation in part through their secondary structures and uORFs, both of which reduce the frequency of preinitiation complexes that reach the iAUG. Our motif models for 5’ regions likely include sequence information for these two general features as well as capturing general machinery contacts close to the iAUG and, potentially, other *cis*-elements also. It is interesting, therefore, to quantitate the overlap of regulatory information in models for RNA structure, for uORFs and for 5’ sequence motifs.

**Figure 9.**
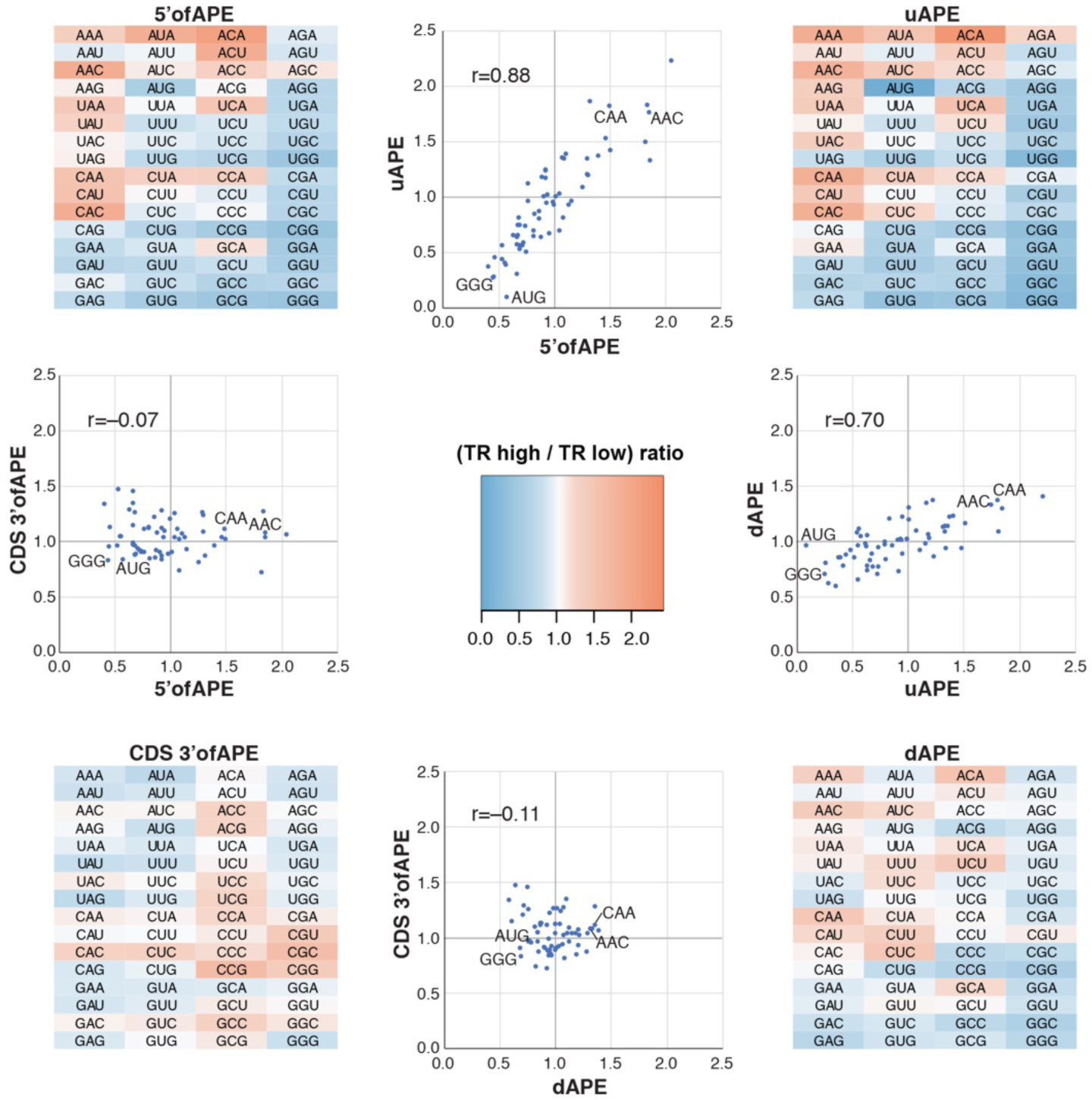
Correlations between regulatory regions in Arabidopsis mRNAs. The four heat maps show the frequency of each tri-nucleotide in the most highly translated 10% of genes divided by its frequency in the most poorly translated 10% of genes ((TR high / TR low) ratio). Ratios were calculated for four separate parts of the mRNA: 5’ofAPE; uAPE; dAPE; and the CDS 3’ofAPE. Scatter plots show the correlation in high/low ratios between selected regions. The Pearson correlation is given. Data points for four tri-nucleotides are indicated (GGG, AUG, CAA, AAC). Strong positive correlates are seen between 5’ofAPE and uAPE and between uAPE and dAPE. CDS 3’ofAPE only correlate weakly with the other regions, consistent with its different function.

**Figure 10.**
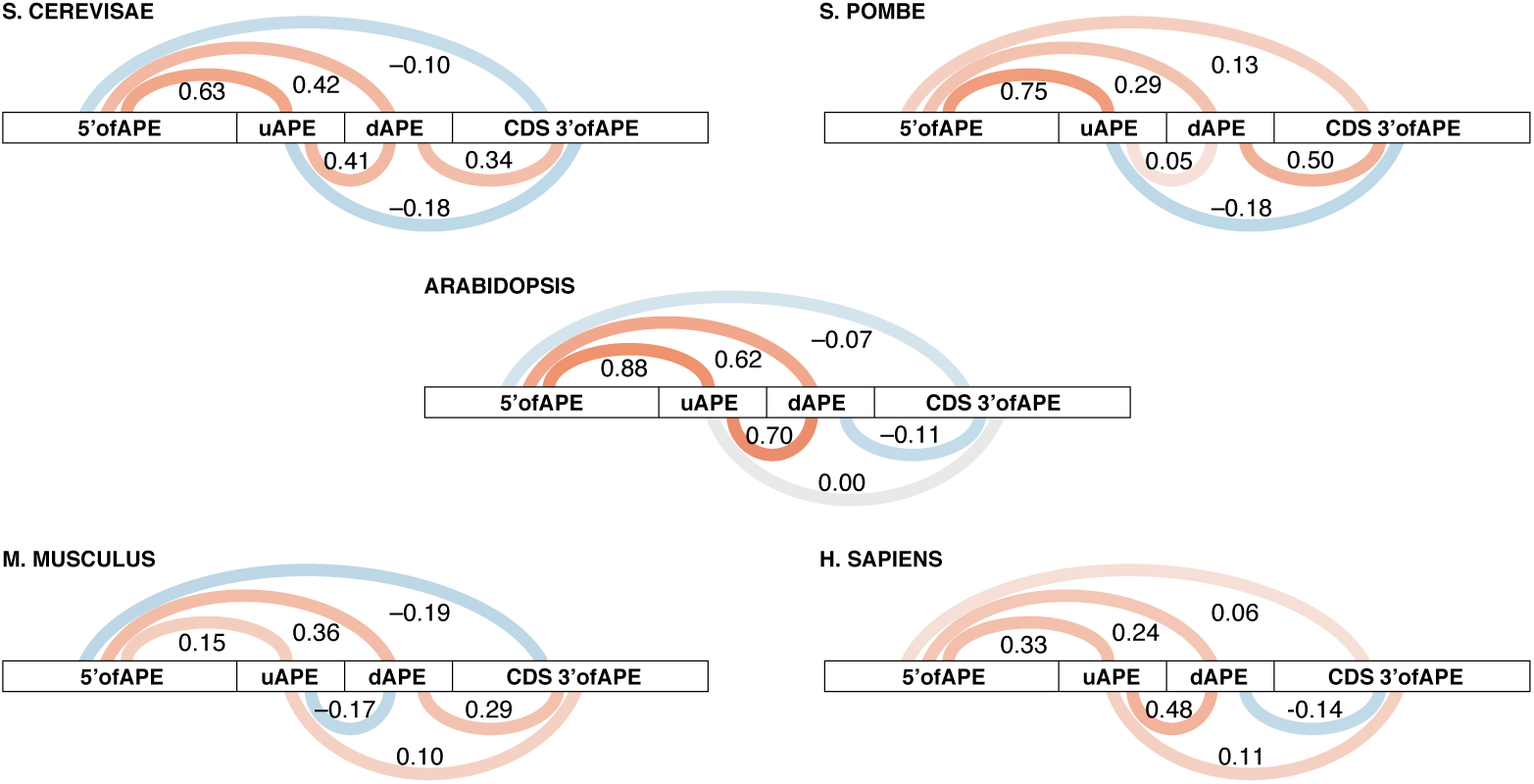
Correlations between translational *cis*-regulatory elements within a species. The Pearson correlation coefficients between the (TR high / TR low) tri-nucleotide ratios for different portions of mRNAs are shown. The correlations were calculated from pairwise comparisons such as those shown in Figure 9. The color intensities are scaled to the correlation coefficient.

**Figure 11.**
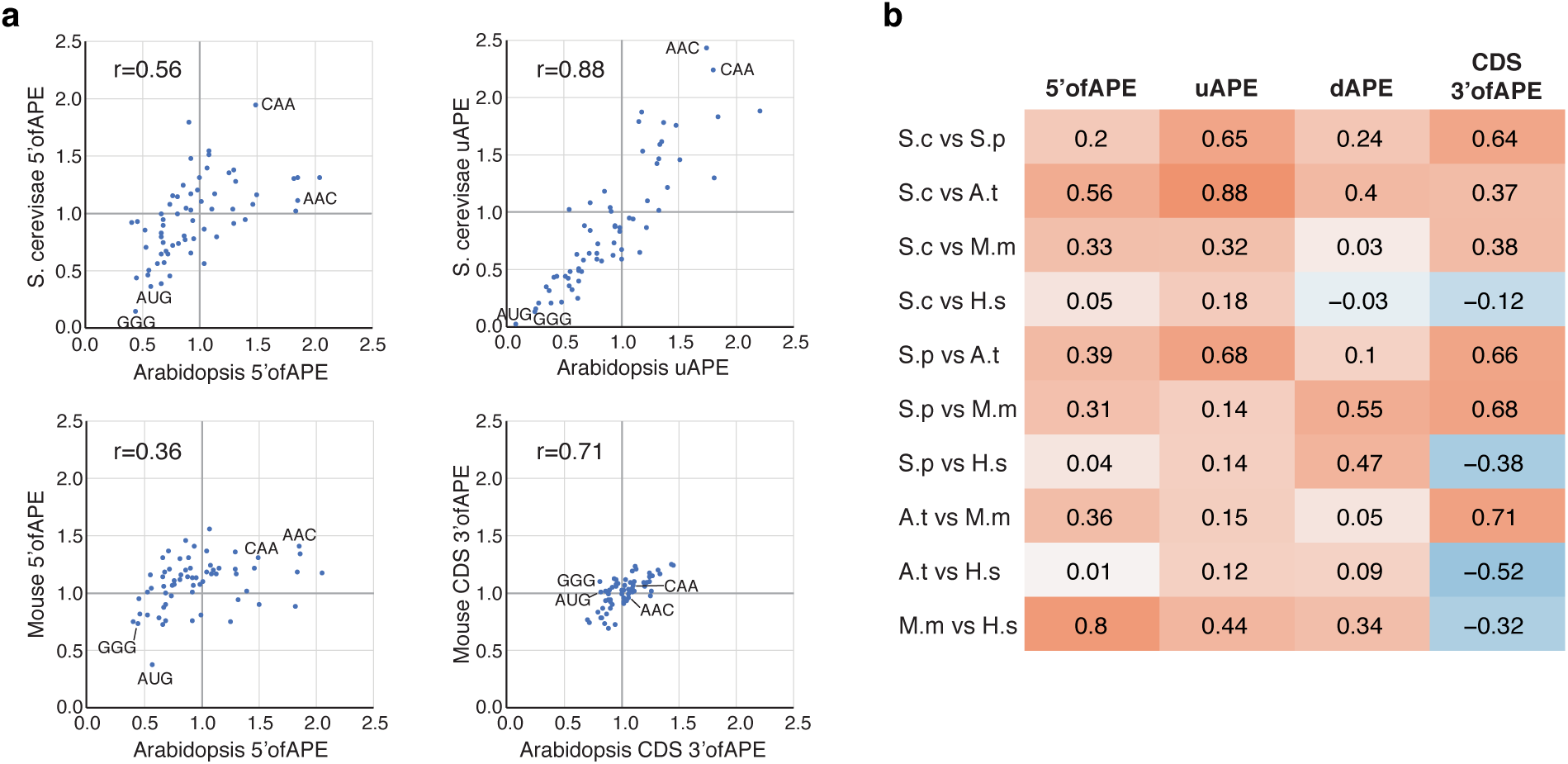
Correlations between translational *cis*-regulatory sequences in different species. (a) Scatter plots showing strong correlations between (TR high / TR low) tri-nucleotide ratios for selected genomic regions from different species. The Pearson correlations (r) are indicated. (b) Pearson correlation coefficients between each of four mRNA regions among the five species. The color intensities are scaled to the correlation coefficient. Clear correlations are observed in many cases.

For this purpose, we represented regulation by RNA structure using the “RNAfold” feature-set; repression by uORFs using the number of upstream AUGs (uAUGs) present in 5’UTRs (“uAUG”); and the combination of these two using a further multivariate model: “5’biochem”. To calculate the overlap of the three resulting biochemical feature-sets with “5’motifs”, three further models were constructed that paired “5’motifs” with each biochemical feature-set (Figure 12). Regulatory overlap is given by the percent of the variance in translation that is collinear (i.e. shared) between feature-sets. Control information in “5’motifs” overlaps strongly with that in “uAUG”, “RNAfold”, and “5’biochem” and *vice versa* (Figure 12). For example, 39%–78% of the explanatory power of “5’motifs” is collinear with “5’biochem”. At the same time, “uAUG”, “RNAfold”, and “5’motifs” each contain unique *cis*-regulatory information not captured by the other. For instance, 48%–72% of the information in the model combining “5’biochem” and “5’motifs” is uniquely obtained from models that employ only one of these two feature-sets, making the combined model our most complete for explaining 5’ region control.

**Figure 12.**
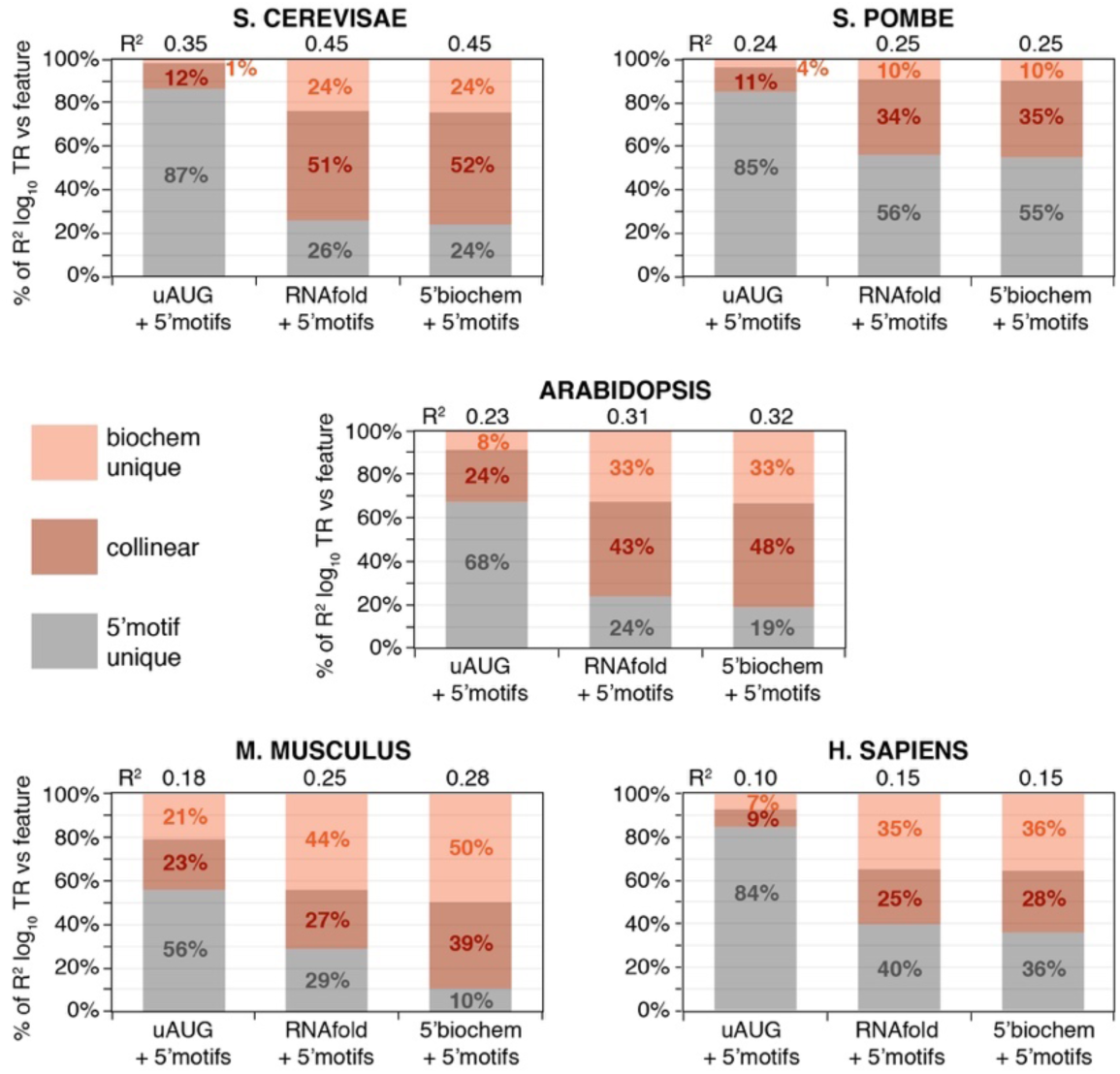
Control by “RNAfold” and “uAUG” is collinear with that by “5’motifs”. Two 5’ features each partially capture control by a biochemical process: “uAUG”, repression due to translation of uORFs; and “RNAfold”, inhibition by RNA secondary structure. A third feature, “5’motifs”, captures 5’ sequences that correlate with translation, without regard to biochemical mechanism. To determine how similar information captured by “5’motifs” is to that in “uAUG” and “RNAfold”, three multivariate models were constructed that combined “5’motifs” either with “uAUG”, with “RNAfold”, or with a third feature (“5’biochem”) that combines both “uAUG” and “RNAfold”. The R^2^ coefficients of these feature-sets vs TR are shown (top). The percent of the R^2^ values contributed by “5’motifs” (grey) or the biochemical process(es) (orange) is shown by length along the *y*-axis. Because some of this information is collinear, the R^2^ coefficient of the combined models are less than that of the sum of the individual contributions. This collinear portion is shown by the darker orange shading, while the lighter orange and the grey shading show the unique contributions.

### The contributions of general features to translation rates

In addition to the 5’ region general features, two others control translation rates: CDS length and codon frequency. CDS lengths are inversely correlated with translation because ribosomal subunits released from mRNAs are recycled from the stop codon to the 5’ cap (40S) or iAUG (60S) within single mRNA molecules, and this process is more efficient for a shorter CDS [9]. Codon frequencies in highly expressed mRNAs more closely match the concentrations of amino acylated tRNAs than do those in poorly expressed mRNAs, which leads to more efficient use of the pool of translational machinery in the cell [10, 13, 34-36].

Therefore, we calculated features-sets that capture regulation by codon frequency and CDS length (features “codon” and “CDS length”) and built a model that combines them with the feature-sets that describe control by the 5’ region. Results were calculated for the five example datasets, for a second *S*. *pombe* dataset, and for data from two additional tissues each for *Arabidopsis* and *M*. *musculus*. The combined models explain 37%–81% of the variance of translation, depending on the dataset, and at least 42% for the best predicted dataset from each species (Figure 13b; Additional file 9; Additional file 1: Figure S8). These results are robust to variations in data as shown by the narrow 95% confidence limits from a 1,000 x bootstrap (Figure 13b). Our results thus suggest a considerably larger role for the general elements than previous models implied (Additional file 1: Table S1). We have also compared the individual contributions of the general features as the percent of either the sum of their five R^2^ values vs translation (Figure 13a) or the R^2^ value of a model combining all five features (Additional file 1: Figure S9). By either approach, the contributions of each feature vary more between the species than between different datasets from the same organism, suggesting that aspects of translational regulation alter quantitatively between species.

**Figure 13.**
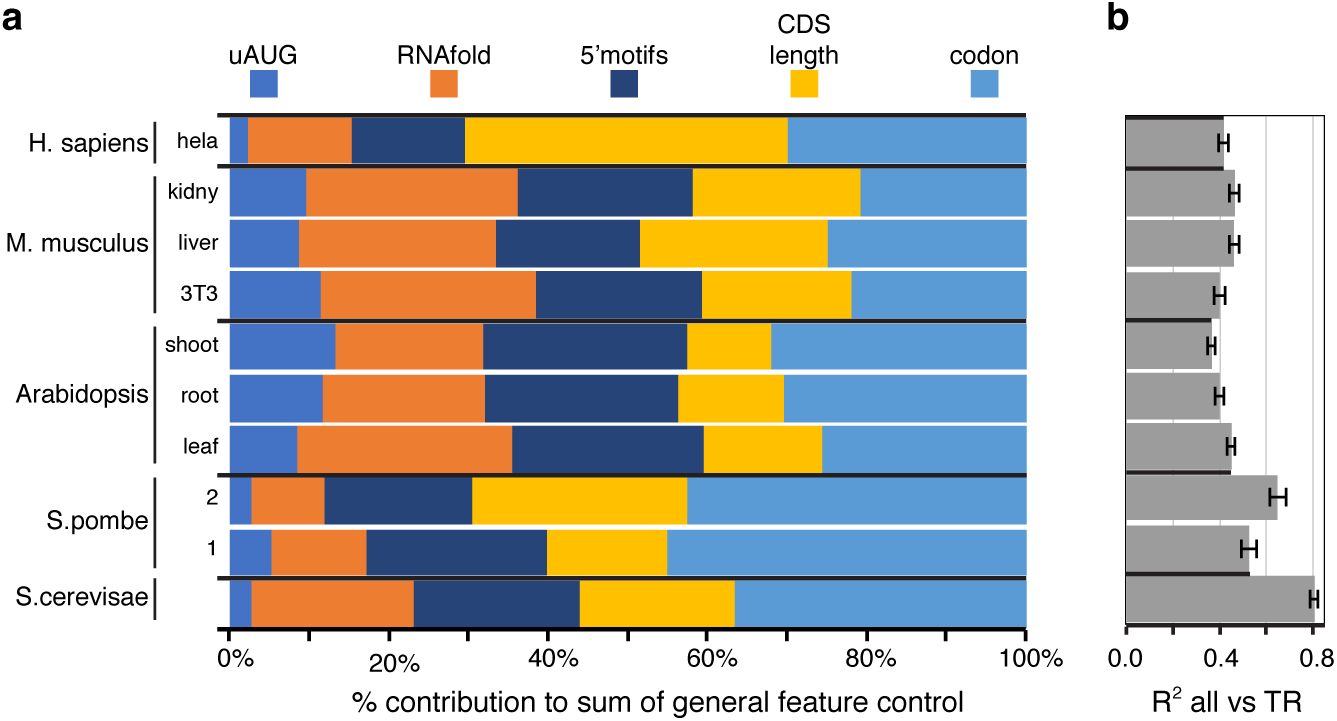
The contributions of general *cis*-control features in five species and several tissues. (a) For each feature or feature-set separately, its R^2^ coefficient of determination vs log_10_ translation rate (TR) is given as a percent of the sum of the R^2^ coefficients for all five features. (b) The R^2^ coefficients between log_10_ TR and models that combine the five features. The 95% confidence limits are shown for 1,000x bootstraps with replacement. The R^2^ values and the values plotted for each feature and the combined model are given in Additional file 9.

### Collinear control

The sum of the variances in translation rates predicted by each of the general features separately is 157%–203% of the variance explained by the model that combines all five features (Additional file 1: Figure S9). Four of the features each represent a separate biochemical process (the exception being “5’motifs”, which captures information redundant with “RNAfold” and “uAUG”, see Figure 12). The *cis*-elements that direct these four biochemical processes must, therefore, have coevolved to work in a correlated manner.

To quantitate the overlap in control, we calculated collinearity between pair-wise combinations of general features within each species. “5’motifs” was excluded, though, to avoid its redundancy with “RNAfold” and “uAUG” (Figure 14). All feature-pairs tested show collinearity. This collinearity is due to inherent correlations among the general mRNA features because if we randomly permute each feature’s observed values independent of other features, the collinearity among features is expected to go down to zero. The overlap in control is largest in the two yeasts, where five out of the six pairs have a remarkable 47%–100% overlap, and smallest in the two mammals, where five of the six pairs have 22%–59% overlap. Some feature-pairs are strongly collinear in all species: for example, “CDS length” and “codon”, which show 45%–62% overlap. Other pairs show dramatic differences: for example, “uAUG” is ≥70% collinear with “RNAfold” in *S*. *cerevisiae, S*. *pombe*, and *Arabidopsis*, but only 7% in *M*. *musculus*.

**Figure 14.**
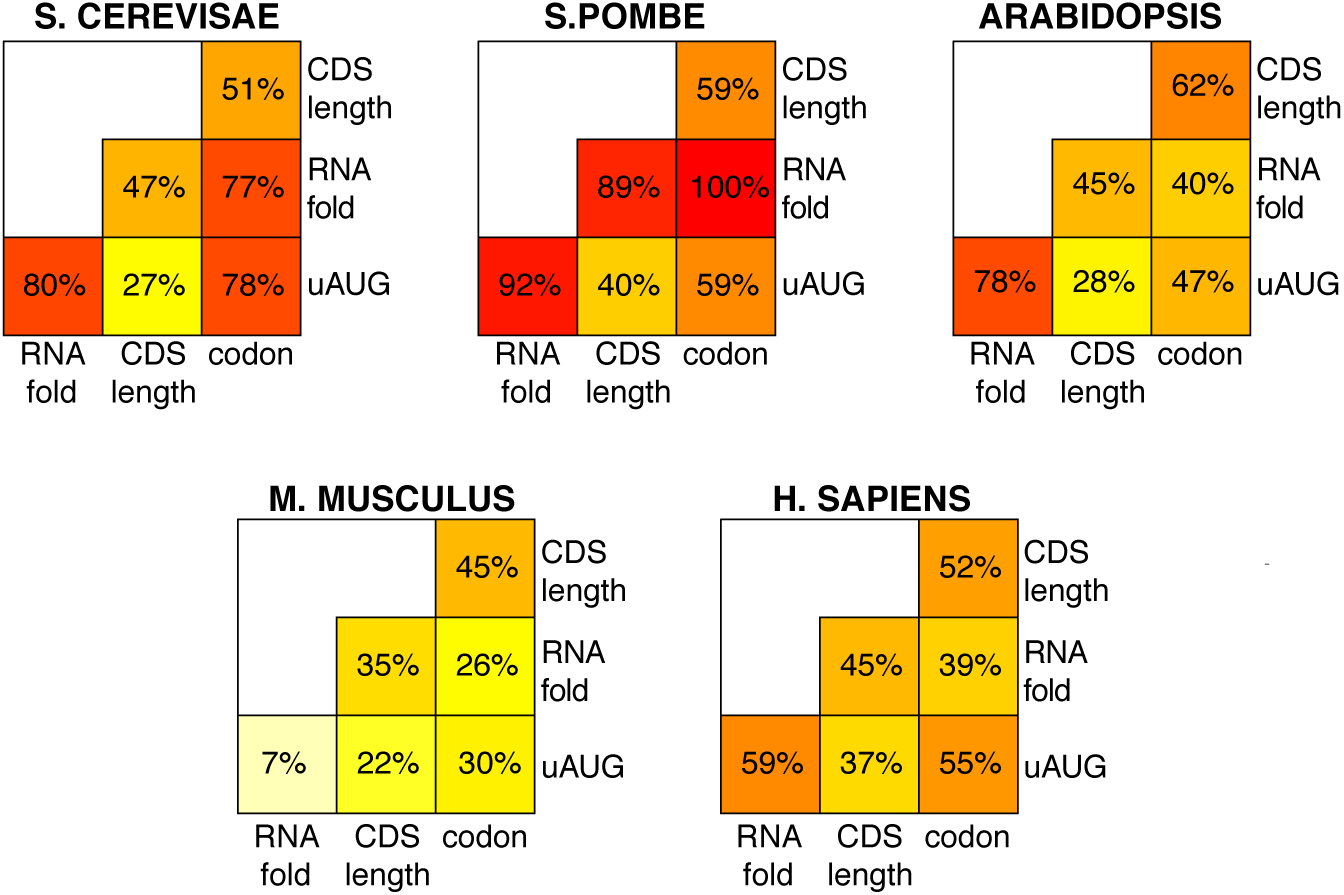
Distinct biochemical processes control translation collinearly. For two features A and B, the R^2^ coefficient of a model that combines both features vs log_10_ translation rate (TR) was defined as R^2^AB, while the R^2^ coefficients of each feature separately vs log_10_ TR were defined as R^2^A and R^2^B. The larger of (R^2^A+R^2^B-R^2^AB)/R^2^A or (R^2^A+R^2^B-R^2^AB)/R^2^B is given and represents the percent in overlap of regulatory control. The overlap in regulatory information varies widely between different pairs, both within and between species.

The collinearity between features in the 5’ region and either “codon” or “CDS length” occurs between physically distinct parts of the mRNA. Our earlier analysis of 5’ region motifs provides additional evidence of correlated regulation between distinct mRNA segments: for example, in the two species with long APEs, tri-nucleotide frequencies correlate strongly between the 5’ofAPE, the uAPE, and the dAPE (Figures 9 and 10). To characterize the collinearity between physically distinct parts of mRNAs further, we identified the second and third most folded windows within 5’ regions that do not overlap with each other or with the most folded window. 63–98% of control exerted by the second most folded window is correlated with that by the most folded window (Figure 15a). Similarly, 67%–92% of control by the third most folded window is collinear with that of the most of folded window (Figure 15b). We also tested for collinearity within the CDS by calculating codon frequencies in N-terminal halves and separately in C-terminal halves. Control by the C-terminal half is 49%–85% collinear with that by the N-terminal half (Figure 15c).

**Figure 15.**
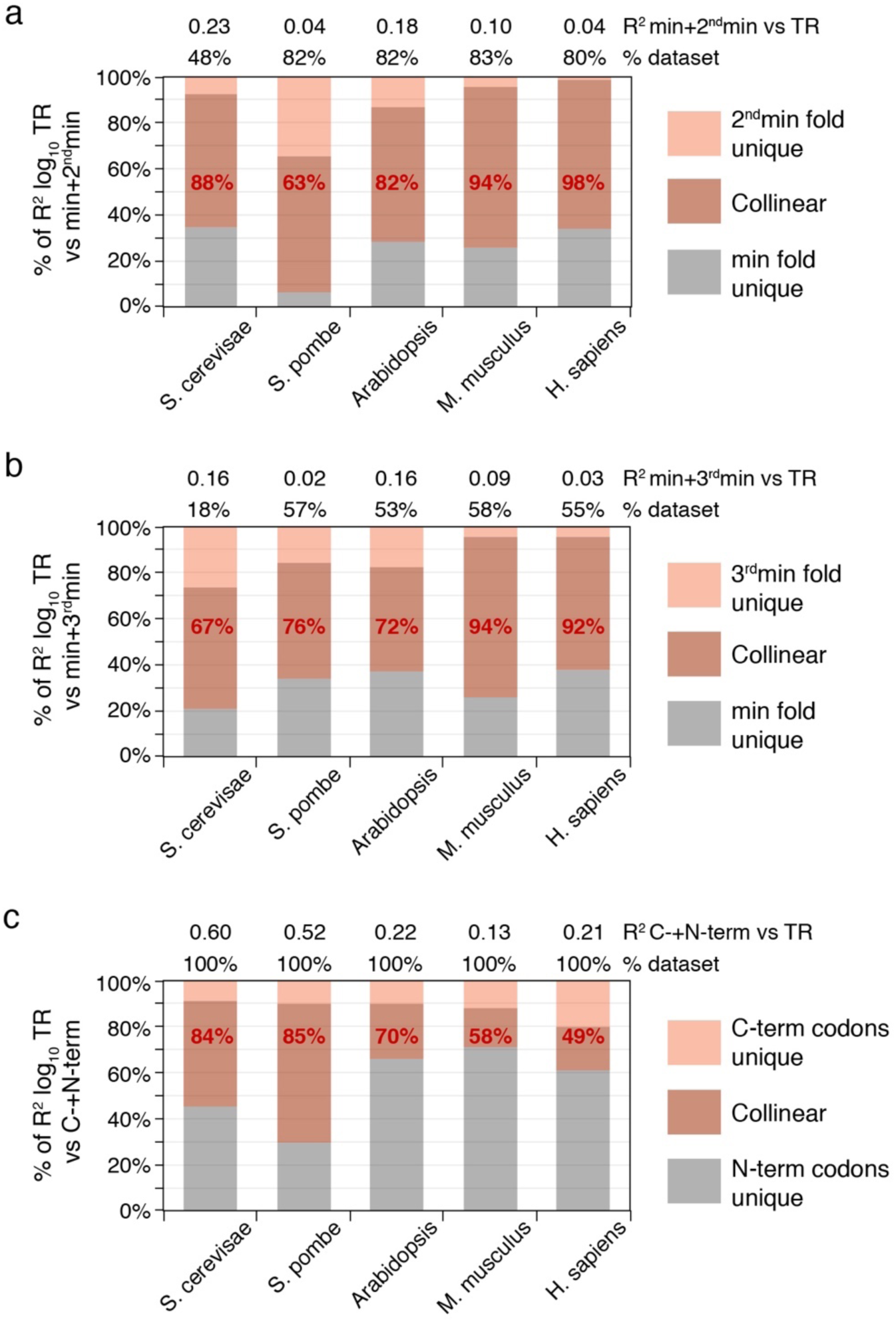
Collinear control between differently located RNA segments. (a) A model was built for the combined control of translation rate (TR) by both the most highly folded window (“min”) and the non-overlapping second most highly folded window (“2^nd^min”). The R^2^ coefficients for these combined features vs log_10_ TR are given (top). The percent of the unique and collinear contributions of “min” and “2^nd^min” to the total variance in translation explained by the model for both features is shown by length on the *y-*axis. The percent of control of TR by “2^nd^min” that is collinear with control by “min” is shown in red text. The percent of genes in the dataset whose 5’ regions are long enough to contain the two non-overlapping windows used in the analysis are indicated (top). (b) The collinearity between the most highly folded window (“min”) and the non-overlapping third most highly folded window (“3^nd^min”) is displayed as described in panel “a”. (c) The collinearity between control of translation by codon frequency in the N-terminal half of each protein (“N-codon”) and C-terminal half of each protein (“C-codon”) is shown as described in panel “a”, except that the percent of control of TR by “C-codon” that is collinear with control by “N-codon” is shown (red text), and 100% of genes in each dataset were used (top).

The correlation of codon frequency with translation rates is determined both by the frequencies of amino acid in proteins and by preference for some synonymous codons versus others [10, 37]. Synonymous codon preferences and amino acid content are logically independent of each other, i.e. in the absence of empirical data one cannot assume *a priori* that their variation between genes is related (Figure 16a). It is possible, then, that their effect on translation could be either collinear or non-collinear. To test for any collinearity, we calculated for each gene both the frequency of each amino acid (“AA”, 20 features in total) and the synonymous preference ratio for each codon (“syn.codon”, 61 features in total) (Figure 16a). As expected, control by both “AA” and “syn.codon” correlates with tRNA abundances (Additional file 1: Figure S10) [10, 13, 34-36], and, importantly, regulation by these two feature-sets is strongly collinear in all species (Figure 16b).

**Figure 16.**
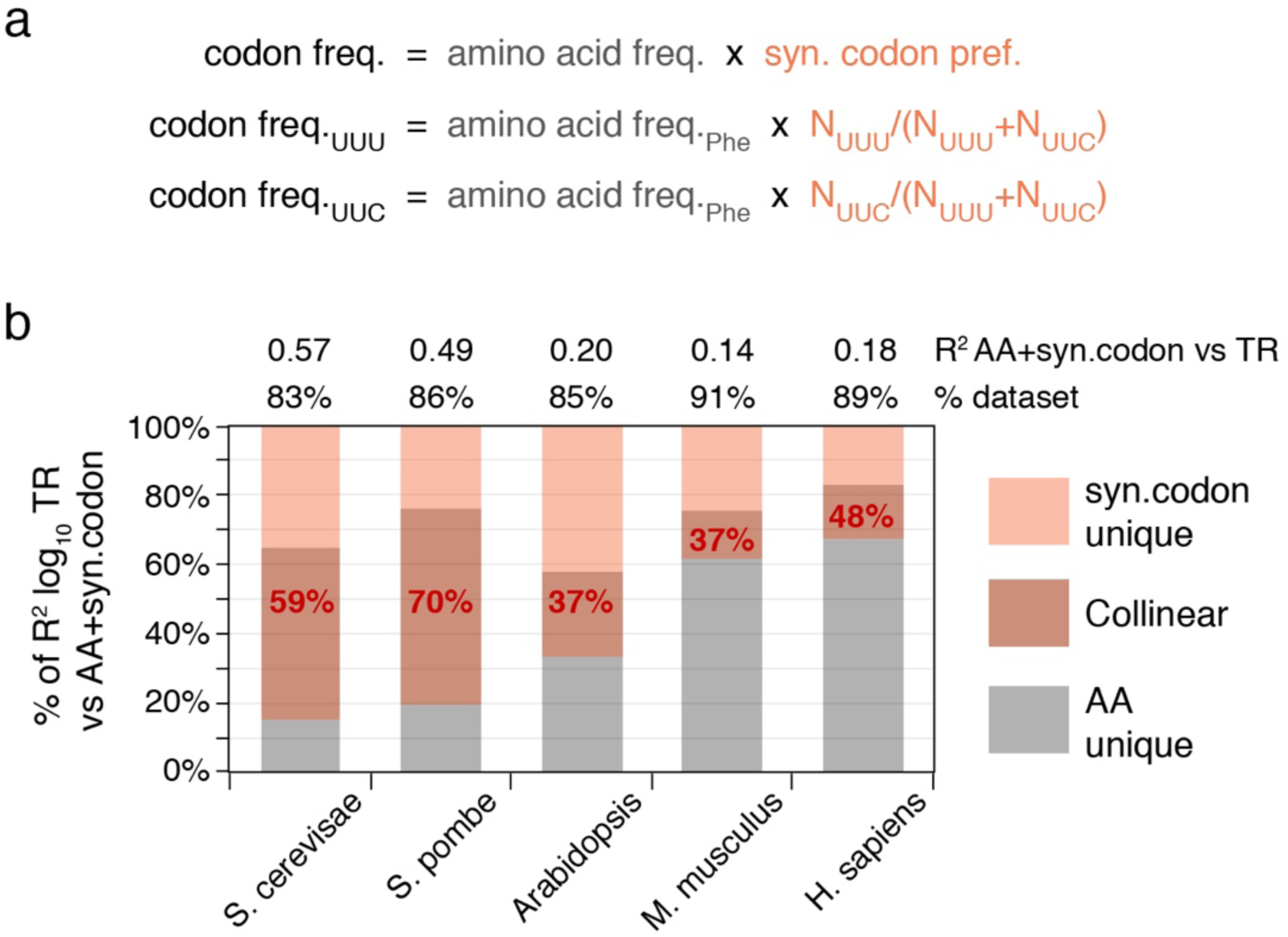
Collinear control by amino acid frequency and synonymous codon preference. (a) The frequency of a codon in a gene is the product of its cognate amino acid frequency (AA) and its synonymous codon preference (syn.codon), where the latter = (number of occurrences of the codon) / (number of occurrences all codons for the cognate amino acid). Examples for the two codons for Phenylalanine (Phe) are shown. *A priori*, there is no necessity for “AA” and “syn.codon” to be correlated. (b) A model was built for control of translation rates (TR) by the combination of “syn.codon” (61 features) and “AA” (20 features). The R^2^ coefficients for this model are given (top). The percent of the unique and collinear contributions of “AA” and “syn.codon” to the total variance in log_10_ TR explained by the combined model is shown by length on the *y-*axis. The percent control of TR by “syn.codon” that is collinear with that by “AA” is shown in red text. The percent of the dataset that contains all 20 amino acids and thus could be used to calculate “syn.codon” is also shown (top). This set of genes was used for all analyses.

Our various analyses thus reveal surprising, strong and complex co-evolution of control between distinct biochemical processes and between multiple segments of mRNA.

## DISCUSSION AND CONCLUSIONS

Here we have sought to better understand how and why differences in translation rates among genes are regulated at steady state by focusing on general mRNA sequence features. We have established common quantitative principles, discussed below, by which these control elements act across diverse eukaryotes. Through optimizing methods to predict translation using mRNA structures, sequence motifs, uORFs, CDS length, and codon usage, we find that the general features together control 81% and 65% of the variance in rates in *S*. *cerevisiae* and *S*. *pombe* respectively and 42%–46% in *Arabidopsis, M*. *musculus* and *H*. *sapiens* (Figure 13). This is significantly higher than earlier estimates (Additional file 1: Table S1). For example, prior work suggested that general features explained 39% of the variance in *S*. *cerevisiae* [13] and 14% in *M*. *musculus* [16].

The part of the variance in measured translation rates that is unexplained by our models for general features is likely due to a combination of:

i. Gene/condition specific regulation by miRNAs and sequence specific RNA binding proteins, which our models do not detect.
ii. Failure of our models to fully capture control by the general features due to, for example, uncertainty in fully predicting RNA structures.
iii. Measurement error in the translation rate data.

It is also possible that our models have exaggerated control by the general features due to “overfitting”, which would lead to aberrant capture of some part(s) of i. and/or iii. Our conservative approach of fitting linear models to features that score known biochemical processes or simple motifs should limit this, however. The likelihood of overfitting is reduced further by our use of Bayesian Information Criterion for feature selection, which we found typically selected fewer features than other approaches, such as ten-fold cross validation (unpublished data). Our models, thus, probably have underestimated true control by the general *cis*-elements by the—unknown—percent of the variance in the translation rate data that is due to ii. and iii.

It is noteworthy that our general feature models predict translation rates more effectively in *S*. *cerevisiae* and *S*. *pombe* than in multicellular eukaryotes. miRNAs are not present in the two yeasts, whereas *Arabidopsis* encodes ~420 miRNAs and *M*. *musculus* and *H*. *sapiens* >1,800 each [38, 39]. Likewise, the multicellular eukaryotes encode over three times more sequence specific RNA binding proteins than *S*. *cerevisiae* and *S*. *pombe* (>350 vs ~100; http://cisbp-rna.ccbr.utoronto.ca [40, 41]). Thus the general features may well play a larger role in the two yeasts than in the multi-cellular eukaryotes as the latter are expected to suffer greater gene and condition specific regulation.

### 5’ mRNA secondary structure

Our analysis provides a more precise understanding of the mRNA secondary structures that control translation within 5’ regions. We have found that the most folded 25–60 nucleotide segments contain far more regulatory information than the less folded parts of 5’ regions in all five species (Figures 3 and 4). Previous studies, by contrast, relied on the mean of the folding energies of short windows that tile the 5’ region, on the folding of a single large window encompassing the entire region, or on windows at fixed locations (Additional file 1: Table S1). Our analysis indicates that the most folded windows alone are more predictive than these prior approaches and that models that combine highly folded windows with location specific ones are even more effective (Figure 3; Additional file 5). Control by the most folded segment is dominated by the longest contiguous run of paired nucleotides within a stem (Figure 5). This control is enhanced by additional nucleotide pairs that lie beyond any mismatched nucleotides or single nucleotide bulges to form a longer stem and by smaller satellite stem loops that reside within the segment (Figure 5). Pairings between nucleotides that are separated by >25–60 nucleotides have only a minimal effect on translation rates (Figure 3). The differences in structures that distinguish translation rates are surprisingly small: only 0.7–1.6 nucleotide pairs on average differentiate the most folded segments in highly translated mRNAs from those in poorly translated mRNAs across all species (Figure 6).

### Conserved 5’ sequence motifs

One of the approaches that we have used to define general *cis*-control elements is an unbiased method that simply categorizes differences in 5’ region sequence motifs between highly and poorly translated mRNAs (Figures 7–11). Non-mammalian species have A/T rich 5’ regions, whereas those of mammals are G/C rich. Despite these differences, we find that when judged by the ratios of the frequencies of each tri-nucleotide in highly and in poorly translated mRNAs (high TR / low TR ratios), there are marked correlations between the *cis*-control sequences in the 5’ regions of mammalian and non-mammalian eukaryotes (Figure 11). The fact that most tri-nucleotide’s (high TR / low TR) ratios ≠1 and are similar across many species (Figures 9 and 11; Additional file 1 Figure S7) strongly implies that a large percent of nucleotides in 5’ regions are involved in regulating translation, rather than having some other function or no function. Those tri-nucleotides whose (high TR / low TR) ratios differ the most from one are presumably those most important for control.

The correlation in 5’ motifs between distant species occurs in part because our motif model captures repression by uORFs via the higher density of uAUGs in poorly translated mRNA (Figures 11 and 12). It also results because our 5’ motifs model captures control by evolutionarily conserved nucleotides that are adjacent to the iAUG and contact the ribosome (Figure 7) [42-45]. Additionally, the correlation is probably due to the fact that di- and tri-nucleotides have differing propensities to form mRNA secondary structures because to their respective base pairing and stacking energies [29, 32, 46, 47]. The important role of RNA structure in directing translation rates will thus drive similar changes in nucleotide content between highly and poorly translated mRNAs in all species. Indeed, 38%–66% of translational control explained by our 5’ motifs model is collinear with control by our model for RNA folding (Figure 12), and those mRNA regions that show the greatest conservation of tri-nucleotide ratios between highly diverged eukaryotes also have the largest differences in RNA folding energies between highly and poorly translated mRNAs (Figures 2, 3, and 11).

### Collinear control between processes and between mRNA segments

Perhaps the most striking observation is that there is a strong correlation in control (i.e. collinearity) between general features that act through distinct biochemical processes and also between different parts of the mRNA. For example, all pair-wise combinations of biochemical process show collinear regulation of translation, many by >50% (Figure 14). Also, the Gibbs energies of the second and third most folded mRNA segment within 5’ regions share a remarkable 63%–98% control with those of the most folded segment; and codon frequencies in the C-terminal half of CDSs share 49%–85% control with those in the N-terminal half (Figure 15). Finally, regulation by synonymous codon preferences is 37%–70% collinear with that by amino acid content, even though the two determinants of codon usage are logically independent of each other (Figure 16). This extensive collinearity reveals a form of strongly repeated information within mRNAs that went undetected by approaches such as testing for nucleotide sequence similarities, illustrating the power of the Analysis of Variance (ANOVA) based approach we have used.

Collinearity between different sections of the mRNA keeps the rate of progress of the translation machinery more constant and thus the density of translational complexes more uniform along any given mRNA (Figure 17a). The translation rate within each mRNA is not determined by a single rate limiting step in this case. Instead, the flux of complexes down a particular mRNA is controlled by a series of steps that are each set to allow a similar rate. The steps share regulation much in the way that the enzymes in a metabolic pathway share fractional control of metabolic flux, as defined by their flux control coefficients [48, 49]. Uncorrelated control, by contrast, would lead to a more uneven distribution of complexes. In this alternate case, the overall flux of complexes for a given mRNA would often be defined by a single rate limiting step, with the important proviso that different steps would limit on different genes to account for the fact that all general features contribute to the overall variance in rates (Figure 17b).

**Figure 17.**
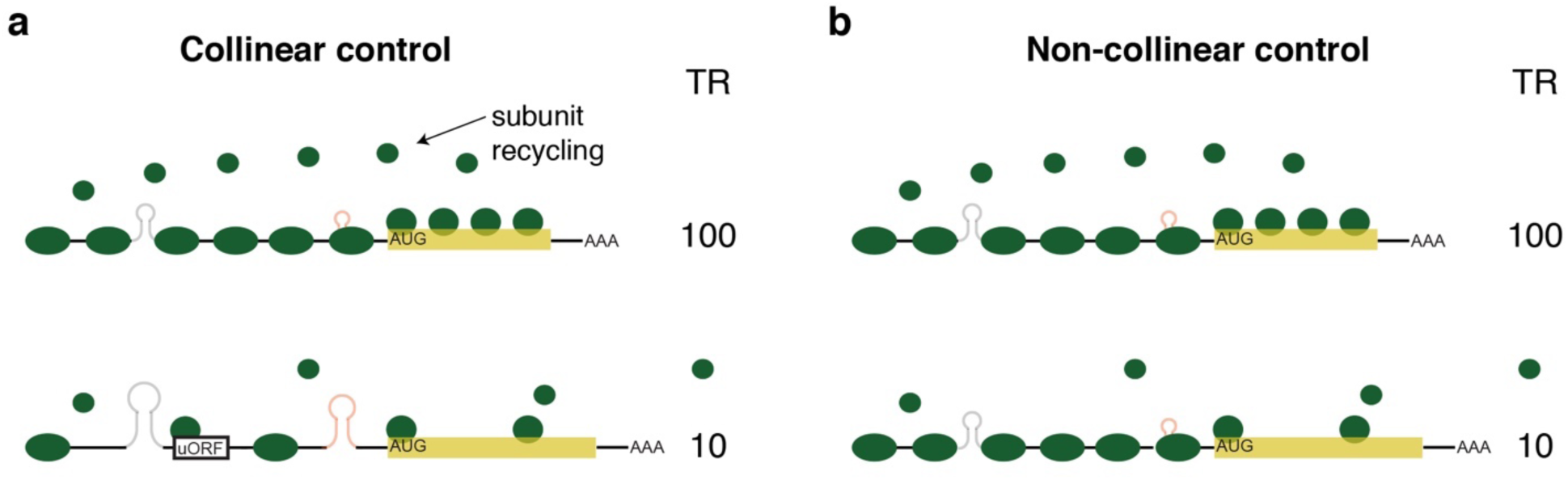
Collinear control results in more uniform particle densities than non-collinear control. (a) Collinear control. The upper, high translation rate (TR) mRNA has multiple small mRNA stem/loops, no uORFs, a short CDS, and optimal codon frequency. The bottom, low TR mRNA has multiple larger mRNA stem/loops, uORFs, a long CDS, and non-optimal codon frequency. As a result both mRNAs have relatively even densities of translational complexes along their length. The CDS length dependent rate of ribosome subunit recycling also matches their respective translation rates. (b) Non-collinear control. Both mRNAs have multiple small mRNA stem/loops and no uORFs resulting in a high density of pre-initiation complexes on the 5’UTR. The top mRNA has a short CDS and optimal codon frequency, which results in a high density of ribosomes along the CDS and efficient ribosome subunit recycling. The bottom mRNA has a suboptimal ribosomal contacts at the iAUG, which reduces the efficiency with which pre-initiation complexes convert to active ribosomes. As a result, the density of ribosomes along the CDS and in the recycling step is lower than that directed by the 5’UTR. The density of translational complexes along this mRNA is thus uneven. The rates of translation of mRNAs in scenarios (a) and (b) are the same. The total number of translation machinery complexes needed in a cell, however, is greater in the non-collinear scenario than in a cell employing collinear control.

There are two known selective pressures that favor—or force—uniform densities of complexes and thus collinear control over non-collinear control.

i. Non-collinear control would increase the frequency of particle collisions when pre-initiation complexes or ribosomes that are moving rapidly down an mRNA enter a region whose regulatory sequences signal a slower rate of progress (Figure 17b, bottom). Ribosomal collisions lead to translational termination and No-Go RNA decay or simply slow the mean elongation rate [50-52]. Collinear control between mRNA segments thus reduces the frequency of these deleterious events. Earlier work has shown that the rate of translation is slower near the N-terminus of the CDS, then increases moderately further 3’ [13, 35]. This arrangement has been proposed to reduce collision frequency [35]. Our results extend this prior model to suggest that it is important to keep a relatively even flux of translational complexes—avoiding strong stops—from the assembly of the preinitiation complex at the 5’ cap all the way to the termination of translation at the stop codon.
ii. Collinear control requires fewer ribosomal subunits per cell than non-collinear control to achieve a given set of translation rates (compare Figure 17 a and b). This is important as the translational machinery is limiting for protein synthesis and cell growth [53-55]: for example, a twofold reduction in the abundances of any one of a number of ribosomal proteins greatly reduces cell division rates in *Drosophila* [53], and under many conditions ≥90% of ribosomal subunits are bound to mRNA and active at any instance [56, 57] and/or are present at locally high concentrations in the vicinity of each mRNA due to CDS-length dependent recycling [9]. There is thus a strong selective pressure to use the set of ribosomal subunits in the cell most efficiently. As Figure 17 shows, this requires collinear control of the complete translation cycle, including the recycling step.

In addition to the two above explanations, a general pressure to increase or decrease protein production from a given gene could lead to multiple, independent mutations affecting different processes or the same process in different mRNA regions. This too would result in collinear control. However, we suggest instead that the discovery of strong collinear control supports other evidence that gene specific translation rates are not determined for the most part by the need to achieve a certain protein abundance. First, translation rates only correlate poorly with protein abundance (R^2^ ~0.08 in *S*. *cerevisiae* and *M*. *musculu*s) [10, 58]. Second, while CDS length is an important determinant of translation rates (Figure 13), the length of a protein is unlikely to be selected only to affect translation. Protein length is presumably set largely by protein function. Third, codon frequency—and thus its correlation with translation—is determined by the amino acid content of proteins as well as by preferences for codons synonymous for the same amino acid (Figure 17). Amino acid contents probably result from selection for protein function rather than for translational regulation. Synonymous codon frequencies in highly abundant mRNAs are selected to optimize use of the total pool of amino acylated tRNAs in the cell [10, 13, 34-36] (Additional file 1: Figure S10); the effect that codon usage has on ribosomal elongation rates could be a secondary consequence, one that is, besides, rather small [13].

Thus both CDS length and codon frequencies are likely driven in part by forces external to the need to determine protein abundances, yet combined the two constitute a significant fraction of control by the general features (Figures 13). The proposed need for collinearity to ensure efficient use of ribosomes by the cell will then force the other three general features to coordinate their activities with CDS length and codon frequency. This may explain why translation is not well correlated with protein abundance. Translation rates are determined by multiple selective pressures. In contrast, the selective pressure on protein abundance dominantly affects transcription [10, 59].

*S*. *cerevisae* and other yeasts have larger effective population sizes than *Arabidopsis, M*. *musculus* and *H*. *sapiens*. As a result, the former have lower mutational loads and have been able to better optimize their genome sequences for energy efficient cell growth than have the latter [60, 61]. These results from population genetics provide an additional explanation for our observation that the general features play a more prominent role in *S*. *cerevisae* and *S*. *pombe* than in the three multicellular eukaryotes: the general elements should be more important in organisms with larger effective population sizes, an idea that has been suggested previously for synonymous codon preference [60, 62] and which we now extend to the other general features.

### High throughput mutant assays

A separate approach to identify translational *cis*-elements has tested large numbers of heterologous reporter genes bearing random mutations in 10 to 50 nucleotide segments proximal to the iAUG [12, 17-21]. In some studies models have been developed that explain ~70%–90% of the variance in translation resulting from these mutations (Additional file 1: Table S2).

It is difficult to relate the results of these reporter gene experiments to the regulation of endogenous mRNAs. The regions mutated in the heterologous reporter assays are smaller than or lie outside of the regions where our analysis indicate that regulatory secondary structures are prevalent. Thus the reporter assays have not captured normal control by RNA folding, let alone that by CDS length and codon usage, which are invariant in a reporter assay. In addition, the consensus sequences for uAUGs associated with repression in a heterologous assay differ dramatically from those identified by studies of uORFs in intact natural transcripts. The high throughput mutational approach identified strong enrichment of A or G at nucleotide −3 relative to uAUGs, similar to the Kozak consensus for the iAUG [20]. In endogenous mRNAs, by contrast, no specific nucleotides are strongly associated with repressive uAUGs at location −3 [16, 25, 63]. Thus the sequences of the *cis*-elements driving translation in the heterologous assays differ strongly from those acting in endogenous mRNAs.

One *S*. *cerevisiae* study did measure protein production from natural 5’UTRs ≤ 50 nucleotides in length fused upstream of a reporter gene [21]. Although this study successfully modelled 60% of the variance in measured output, we find that this output correlates poorly with ribosome profiling data for the corresponding endogenous genes (R^2^ = 0.09, Additional file 10). Thus at least (60-9)/60 = 85% of the variance in reporter expression predicted by the model cannot be explained by measured endogenous translation rates.

Our analysis establishes that there is extensive coordination (collinearity) between control by distinct parts of the mRNA and between multiple steps in the translation cycle. Heterologous reporter constructs that include only short segments of mRNAs thus fail to correctly capture *cis*-elements in part because they miss the normal interactions between steps and between differently located *cis*-elements. It will be necessary to design alternative mutant series to understand *cis*-translational control of endogenous transcripts.

## MATERIALS AND METHODS

### Data and code

An example dataset was chosen from each of the five species to present figures for all analyses: *S*. *cerevisiae* [13]; *S*. *pombe* 2 [26]; *Arabidopsis* leaf (dark) [25]; *M*. *musculus* NIH3T3 cell line [4]; and *H*. *sapiens* HeLa cell line [24]. Results from five additional datasets are shown for a subset of analyses: *S*. *pombe* 1 [4]; *Arabidopsis* root and *Arabidopsis* shoot [27]; and *M*. *musculus* liver and *M*. *musculus* kidney [14, 28].

Where replica data was available from a study, genes without data in all replicas were removed; for each gene, its data were then averaged across all replicas; and if the data in the original publication had not be thresholded, genes with mRNA abundances <1 RPKM were excluded. For all datasets, genes for which data on poly-A tail length were not available from Subtelny et al, 2014 were removed. Additional file 2 provides per gene ribosome profiling data and mRNA abundances as well as scores for each of the general features for the set of genes analyzed for each dataset.

The R code used in this analysis is provided in Additional file 11. We also provide a URL for the complete code files for reproducing the results presented. These files include the R code, input data, processed files, preliminary figures, etc.

https://drive.google.com/file/d/1Zjxg-DbSptfSo7xEKR1vTJEf_ebOregV/view?usp=sharing

### Common aspects of models

Translation rate (TR) is defined as Translation Efficiency (TE) values from ribosome profiling data or in the case of *S*. *cerevisiae* as Initiation Efficiency (IE) values, which are corrected TE values that take into account codon specific elongation rates [13]. Log_10_ transformed TR values were used in all regressions.

All models employed single part, multivariate linear regressions for *S*. *pombe, M*. *musculus* and *H*. *sapiens*; two part, multivariate linear regressions of *Arabidopsis*; and three part, multivariate linear regressions for *S*. *cerevisiae*. The multi part regressions were employed in the latter two species to allow optimal scoring of their extended APE PWMs, described below. Of necessity, multi part regression then had to be employed for all other features in these species to allow a model that combines all general features. For the multi part regressions, genes were divided into groups based on lengths of 5’ untranslated regions (UTRs), with the number of groups equal to the number of parts. For *Arabidopsis*, genes were grouped into those with 5’UTRs <65 nucleotides and those with 5’UTRs ≥65 nucleotides. For *S*. *cerevisiae*, genes were grouped into those with 5’UTRs <20 nucleotides; those with 5’UTRs ≥20 nucleotides but <35 nucleotides; and those with 5’UTRs ≥35 nucleotides. Each part corresponds to a separate multivariate linear model fitted to the corresponding group of genes, with the intercept and feature coefficients allowed to differ between parts. Single part regressions included all genes in the dataset.

The Coefficient of Determination R^2^ coefficient was based on the Ordinary Least Squares (OLS) in all cases. The R^2^ calculated in this way is equal to the square of the Pearson correlation between the observed response values and the predicted response values, and it measures the goodness of fit or the predictive power of a model.

### Control by RNA folding (RNAfold)

To predict control of TR by RNA folding in the 5’ region, a series of features were developed using predicted Gibbs free energies generated by ViennaRNA RNAfold [16, 32] (Figure 3). Most features were calculated in 19 variants based on one nucleotide offset sliding windows of lengths varying from 6 to 100 nucleotides (Figure 3). For these window based features, the 5’ regions used span from the 5’ cap to that part of the CDS that is covered by the window whose 5’ end maps to position −1. The features are:

#### Single window features

**5’ cap**, **−65**, **−30**, **−35**, **−6**, **−1**: the free energy of the window whose 5’ nucleotide lies at the position specified.

**min**, **10%**, **25%**, **75%**, **90%**, **max**: the free energy of the window with the minimum, the 10^th^ percentile, the 25^th^ percentile, the 75^th^ percentile, the 90^th^ percentile, or the maximum energy of the all windows in the 5’ region of the mRNA.

Note that windows corresponding to different single window features are allowed to overlap.

#### Multi window features

**mean**: the mean free energy of windows of the specified length.

**%≤20%**, **%≥=90%**, **% ≥80%**: the percent of windows in a 5’ region that have free energy ≤ the 20^th^ percentile, ≥ the 90^th^ percentile, or ≥ the 80^th^ percentile of the free energy of all windows of all genes in the dataset. Some genes will thus have scores of 0 as they lack any windows defined by these thresholds.

**sum≤5%**, **sum≤10%**, **sum≤20%**, **sum≥80%**, **sum≥90%**: the sum of free energies defined by these thresholds out all windows of all genes in the datasets. Some genes will thus have scores of 0 as they lack any windows defined by these thresholds.

#### Whole 5’ region

**whole**: free energy of a fold of the entire sequence from the 5’ cap to +35.

#### feature-sets

Feature-sets fit multivariate linear models (possibly multi part, depending on the species) between log_10_ TR and multiple of the above features.

**All – 1**: combines all features of a given window length.

**All**: combines all features of a given window length and in addition includes the “whole” feature.

**RNAfold**: uses forward selection with Bayesian Information Criterion (BIC) to identify an effective subset of features to predict log_10_ TR. The model is allowed to select different length windows for different features. In the case of *S*. *pombe*, prior to BIC selection all windows with sequences that extend 3’ of +30 were removed to avoid elements whose Gibbs energies have a strong negative correlation with TR. “RNAfold” is our most accurate model for control of TR by RNA structures (Figure 3). Its BIC selected features are listed in Additional file 5.

### Control by sequence motifs (5’motifs)

#### Defining an optimum PWM for the iAUG proximal elements (APEs)

Various length position weight matrices (PWMs) were calculated using the 10% of genes with the highest TR scores (Figure 7a, high TR cohorts). The mRNA sequences of these sets of genes were aligned such that nucleotide +1 corresponds to the A of the iAUG. The frequencies of A, U, C, and G at every position from −100 to + 35 were calculated (Additional file 7). Sequences of all high TR genes were used, including those with 5’UTRs shorter than 100 nucleotides, these short 5’UTR genes contributing only to the frequencies at the 5’UTR positions they contained.

A series of PWMs were constructed that all contained position nucleotide −1 and positions 5’ of that in five nucleotide steps to −100: i.e. −5 to −1 PWM, −10 to −1 PWM, −15 to −1 PWM etc. Variants of each of these PMWs were constructed that also included nucleotide +4 and positions 3’ in 5 or 10 nucleotide steps within the protein coding sequence (CDS) to +33. i.e. −5 to +13 PWM, −5 to +23 PWM, −10 to +13 PWM etc. Given each PWM, say a PWM for *m* nucleotides from the 5’UTR and for *n* nucleotides of the CDS, we calculated the PWM scores of all the genes in a dataset as follows: score of gene *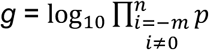*(nucleotide at position *i* of gene *g*).

We then calculated the R^2^ correlation coefficient between the PWM scores based on each PWM boundary and the log_10_ TR values. The results are plotted in Figure 7b. The locations of the approximately optimum APE PWMs are shown boxed (Figure 7a).

#### An APE feature-set

The APE was divided into the part 5’ of +1 (uAPE) and the part 3’ of +3 (dAPE). The optimum length PWMs defined above were divided into two (e.g. *S*. *cerevisiae* uAPE PWM −35 to −1 and dAPE PWM +4 to +28). All genes were scored with the dAPE PWM. In single regression species, all genes were also scored the corresponding uAPE. In *S*. *cerevisiae*, genes with 5’UTRs ≥35 nucleotides were scored with uAPE PWM −35 to −1; genes with 5’UTRs ≥20 nucleotides but <35 nucleotides were scored with uAPE PWM −20 to −1; and genes with 5’UTRs <20 nucleotides with uAPE PWM −5 to −1. In Arabidopsis, genes with 5’UTRs ≥65 nucleotides were scored with uAPE PWM −65 to −1, and genes with 5’UTRs <65 nucleotides with uAPE PWM −5 to −1.

In addition, we calculate the 16 dinucleotide frequencies and 64 trinucleotide frequencies in the uAPE region for every gene as well as the 16 dinucleotide frequencies and 64 trinucleotide frequencies in the dAPE region for every gene. Then we fit a multivariate linear model (possibly multi part, depending on the species) between the log_10_ TR (response) and all features for the uAPE (i.e. uAPE PWM + 16 dinucleotide frequencies + 64 trinucleotide frequencies) and separately all features for the dAPE, and selected features by forward selection with Bayesian Information Criterion. This results in a subset of features that form the uAPE feature-set and the dAPE feature-set (Additional file 1: Figure S5; Additional file 5). In addition, we combined the selected uAPE and dAPE feature-sets to define the APE feature-set (Additional file 1: Figure S5).

#### A 5’ofAPE feature-set

A 5’cap PWM was defined using various length PWMs calculated using the 10% of genes with the highest TR scores in a procedure similar to that used to define the APE (Additional file 1: Figure S6a, high TR cohorts). All 5’cap PWMs share a common 5’ end at the 5’ cap site and differ only in length. The R^2^ correlation coefficients between these PWM scores and log_10_ TR values show a local peak from 3 to 15 nucleotides from the 5’ cap (Additional file 1: Figure S6b). To simplify our model while taking into account the low R^2^ values (<1.2%), a 5’cap PWM was defined as the first 5 nucleotides of the transcript in all species.

In addition, we calculate the 16 dinucleotide frequencies and 64 trinucleotide frequencies in the region 5’ of the APE. Then we fit a multivariate linear model (possibly multi part, depending on the species) between the log_10_ TR (response) and all features for this 5’ofAPE region (i.e. 5’cap PWM + 16 dinucleotide frequencies + 64 trinucleotide frequencies), and selected features by forward selection with Bayesian Information Criterion. The BIC selected features for 5’ofAPE, are listed in Additional file 5 and their R^2^ correlation coefficients vs TR are shown in Figure 8.

#### A 5’motif feature-set

A “5’motifs” feature set was defined by combining the 5’ofAPE, uAPE, and dAPE feature-sets (Figure 8).

### Control by upstream open reading frames (uAUG)

For each gene, the number of AUGs upstream of the iAUG of the CDS was calculated. This feature is designated “uAUG”, and its contribution to the prediction of log_10_ TR is due to the repression of CDS translation by translation of upstream open reading frames (Figure 13; Additional file 1: Figure S9).

### Control by CDS length (CDS length)

For each gene, the log_10_(number of CDS amino acids) was calculated. This feature is designated “CDS length”, and its contribution to the prediction of log_10_ TR is due to the impact of CDS length on the recycling of ribosomal subunits (Figure 13; Additional file 1: Figure S9).

### Control by codon frequencies (codon)

For each gene, we calculate the frequencies of the 61 nonstop codon frequencies in its CDS nucleotide sequence. These 61 features together define the feature-set “codon” (Figure 13; Additional file 1: Figure S9).

### Control by poly-A tail length

For each gene, we took the poly-A tail length defined by Subtelny et al, 2014. For all ten datasets, we find that poly-A length is weakly negatively correlated with translation rates. Prior evidence suggests that this negative correlation does not necessarily reflect a direct control of translation rates, but is instead a result of two phenomenon: the stabilization of mRNAs against degradation by shorter poly-A tails, and the fact that translation rates are positively correlated with mRNA abundance [4, 64]. For this reason, we have not included poly-A length as a feature in our study. Note that there is a much stronger, positive correlation of poly-A tail length with translation rates in pregastrula embryos that likely reflects direct control of translation [4, 15].

### Control by 5’UTR length

5’UTR length has been used as a feature to capture control by 5’ regions in some prior studies [10, 13, 14, 16, 22, 65] (Additional file 1: Tables S1 and S2). We have found, however, that models that combine 5’UTR length with the “5’motifs”, “uAUGs”, and “RNAfold” features only explain an additional 0.03%–0.62% of the variance in translation rates, compared with models that do not include 5’UTR length. This suggests that 5’UTR length is not a direct determinant of translation rates, but is instead largely a co-correlate of other sequence features that do directly affect translation. The fact that control by RNA folding is largely determined by most folded regions, not by less folded regions (Figures 3 and 4), further supports this conclusion. We, therefore, have not included 5’UTR length as a feature in our final models.

### A general feature model using five features

To determine the variance in translation rates explained by “RNAfold”, “5’motifs”, “uAUG”, “CDS length” and “codon”, a multivariate model (possibly multi part, depending on the species) was used to regress log_10_ TR on all five feature-sets together (Figure 13; Additional file 1: Figure S8). The resulting R^2^ correlation coefficient estimates the degree of translational control exerted by these general features.

### Control by amino acid frequencies (AA) and synonymous codon preferences (syn.codon)

For each gene, we calculated the frequencies of the 20 amino acid frequencies in its CDS nucleotide sequence. These 20 features together was defined to be the feature-set “AA” (Figure 16). Then for each of the 61 nonstop codons, we defined a synonymous codon preference as its frequency divided by the sum of the frequencies of synonymous codons that code the same amino acid. For example, AAA, AAT, AAC, and AAG code the same amino acid, so we computed their synonymous codon preferences as AAA/(AAA+AAT+AAG+AAC), AAT/(AAA+AAT+AAG+AAC), AAC/(AAA+AAT+AAG+AAC), and AAG/(AAA+AAT+AAG+AAC). These 61 synonymous codon preferences together define the feature-set “syn.codon” (Figure 16).

## DECLARATIONS

## Acknowledgements

We thank Soile Keranen and Xiaoyong Li for thoughtful critiques of drafts of this manuscript.

## Funding

JJL’s work was supported by the start-up fund of the Department of Statistics at University of California, Los Angeles, a Hellman Fellowship from the Hellman Foundation, the PhRMA Foundation Research Starter Grant in Informatics, the Sloan Research Fellowship, the Johnson & Johnson WiSTEM2D Award, and the NIH/NIGMS grant R01GM120507. Work at Lawrence Berkeley National Laboratory was conducted under U.S. Department of Energy Contract No. DE-AC02-05CH11231.

## Availability of data and materials

The dataset(s) supporting the conclusions of this article is(are) included within the article (and its additional file(s)).”

## Author contributions

JL conducted all statistical analyses. CLC predicted mRNA folding energies and secondary structures. MDB selected the datasets used. MDB, JL and CLC designed the analysis strategies. MDB and JL wrote the manuscript.

## Additional files

Additional file 1. Tables S1–S2 and Figures S—S10. (PDF)

Additional file 2. Translation rate, mRNA sequence and feature score datasets. (XLSX)

Additional file 3: Gibbs energies of dsRNA oligonucleotides from Xia et al 1998. (XLSX)

Additional file 4. RNA feature R^2^ values from Figure 3. (XLSX)

Additional file 5. BIC selected feature sets for “RNAfold” and “5’motifs” features. (XLSX)

Additional file 6. RNA structures for “min” windows. (XLSX)

Additional file 7. APE and 5’cap PWMs. (XLSX)

Additional file 8. Tri-nucleotide high TR / low TR ratios. (XLSX)

Additional file 9. R^2^ values for features and complete model of TR. (XLSX)

Additional file 10. Comparison of Cupress vs Weinberg TR data. (XLSX)

Additional file 11. R source code. (ZIP)

## REFERENCES

1. Hinnebusch AG: Translational regulation of GCN4 and the general amino acid control of yeast. Annu Rev Microbiol 2005, 59:407–450.

2. Gingold H, Pilpel Y: Determinants of translation efficiency and accuracy. Mol Syst Biol 2011, 7:481.

3. Svitkin YV, Yanagiya A, Karetnikov AE, Alain T, Fabian MR, Khoutorsky A, Perreault S, Topisirovic I, Sonenberg N: Control of translation and miRNA-dependent repression by a novel poly(A) binding protein, hnRNP-Q. PLoS Biol 2013, 11:e1001564.

4. Subtelny AO, Eichhorn SW, Chen GR, Sive H, Bartel DP: Poly(A)-tail lengths and a developmental switch in translational control. Nature 2014, 508:66–71.

5. Tuller T, Zur H: Multiple roles of the coding sequence 5’ end in gene expression regulation. Nucleic Acids Res 2015, 43:13–28.

6. Radhakrishnan A, Green R: Connections Underlying Translation and mRNA Stability. J Mol Biol 2016, 428:3558–3564.

7. Hinnebusch AG, Ivanov IP, Sonenberg N: Translational control by 5’-untranslated regions of eukaryotic mRNAs. Science 2016, 352:1413–1416.

8. Thompson MK, Gilbert WV: mRNA length-sensing in eukaryotic translation: reconsidering the “closed loop” and its implications for translational control. Curr Genet 2016.

9. Fernades LD, de Moura APS, Ciandrini L: Gene length as a regulator for ribosome recruitment and protein synthesis: theoretical insights. Scientific Reports 2017, 7:17409.

10. Li JJ, Chew GL, Biggin MD: Quantitating translational control: mRNA abundance-dependent and independent contributions and the mRNA sequences that specify them. Nucleic Acids Res 2017, 45:11821–11836.

11. Cottrell KA, Szczesny P, Djuranovic S: Translation efficiency is a determinant of the magnitude of miRNA-mediated repression. Sci Rep 2017, 7:14884.

12. Shah P, Ding Y, Niemczyk M, Kudla G, Plotkin JB: Rate-limiting steps in yeast protein translation. Cell 2013, 153:1589–1601.

13. Weinberg D, Shah P, Eichhorn S, Hussmann J, Plotkin J, Bartel D: Improved ribosome-footprint and mRNA measurements provide insights into dynamics and regulation of yeast translation. Cell Reports 2016, 14:1787–1799.

14. Janich P, Arpat AB, Castelo-Szekely V, Lopes M, Gatfield D: Ribosome profiling reveals the rhythmic liver translatome and circadian clock regulation by upstream open reading frames. Genome Res 2015, 25:1848–1859.

15. Eichhorn SW, Subtelny AO, Kronja I, Kwasnieski JC, Orr-Weaver TL, Bartel DP: mRNA poly(A)-tail changes specified by deadenylation broadly reshape translation in Drosophila oocytes and early embryos. Elife 2016, 5.

16. Chew GL, Pauli A, Schier AF: Conservation of uORF repressiveness and sequence features in mouse, human and zebrafish. Nat Commun 2016, 7:11663.

17. Dvir S, Velten L, Sharon E, Zeevi D, Carey LB, Weinberger A, Segal E: Deciphering the rules by which 5’-UTR sequences affect protein expression in yeast. Proc Natl Acad Sci U S A 2013, 110:E2792–2801.

18. Noderer WL, Flockhart RJ, Bhaduri A, Diaz de Arce AJ, Zhang J, Khavari PA, Wang CL: Quantitative analysis of mammalian translation initiation sites by FACS-seq. Mol Syst Biol 2014, 10:748.

19. Ben-Yehezkel T, Atar S, Zur H, Diament A, Goz E, Marx T, Cohen R, Dana A, Feldman A, Shapiro E, Tuller T: Rationally designed, heterologous S. cerevisiae transcripts expose novel expression determinants. RNA Biol 2015, 12:972–984.

20. Sample PJ, Wang B, Reid DW, Presnyak V, McFadyen I, Morris DR, Seelig G: Human 5’ UTR design and variant effect prediction from a massively parallel translation assay. BioRxiv 2018, doi:10.1101/310375.

21. Cuperus JT, Groves B, Kuchina A, Rosenberg AB, Jojic N, Fields S, Seelig G: Deep learning of the regulatory grammar of yeast 5’ untranslated regions from 500,000 random sequences. Genome Res 2017, 27:2015–2024.

22. Rojas-Duran MF, Gilbert WV: Alternative transcription start site selection leads to large differences in translation activity in yeast. RNA 2012, 18:2299–2305.

23. Li GW, Burkhardt D, Gross C, Weissman JS: Quantifying absolute protein synthesis rates reveals principles underlying allocation of cellular resources. Cell 2014, 157:624–635.

24. Guo H, Ingolia NT, Weissman JS, Bartel DP: Mammalian microRNAs predominantly act to decrease target mRNA levels. Nature 2010, 466:835–840.

25. Liu MJ, Wu SH, Wu JF, Lin WD, Wu YC, Tsai TY, Tsai HL, Wu SH: Translational landscape of photomorphogenic Arabidopsis. Plant Cell 2013, 25:3699–3710.

26. Duncan CDS, Mata J: Effects of cycloheximide on the interpretation of ribosome profiling experiments in Schizosaccharomyces pombe. Sci Rep 2017, 7:10331.

27. Hsu PY, Calviello L, Wu HL, Li FW, Rothfels CJ, Ohler U, Benfey PN: Super-resolution ribosome profiling reveals unannotated translation events in Arabidopsis. Proc Natl Acad Sci U S A 2016, 113:E7126–E7135.

28. Castelo-Szekely V, Arpat AB, Janich P, Gatfield D: Translational contributions to tissue specificity in rhythmic and constitutive gene expression. Genome Biol 2017, 18:116.

29. Borer PN, Dengler B, Tinoco I, Jr., Uhlenbeck OC: Stability of ribonucleic acid doublestranded helices. J Mol Biol 1974, 86:843–853.

30. Groebe DR, Uhlenbeck OC: Characterization of RNA hairpin loop stability. Nucleic Acids Res 1988, 16:11725–11735.

31. Mathews DH, Sabina J, Zuker M, Turner DH: Expanded sequence dependence of thermodynamic parameters improves prediction of RNA secondary structure. J Mol Biol 1999, 288:911–940.

32. Lorenz R, Bernhart SH, Honer Zu Siederdissen C, Tafer H, Flamm C, Stadler PF, Hofacker IL: ViennaRNA Package 2.0. Algorithms Mol Biol 2011, 6:26.

33. Xia T, SantaLucia J, Jr., Burkard ME, Kierzek R, Schroeder SJ, Jiao X, Cox C, Turner DH: Thermodynamic parameters for an expanded nearest-neighbor model for formation of RNA duplexes with Watson-Crick base pairs. Biochemistry 1998, 37:14719–14735.

34. Gingold H, Tehler D, Christoffersen NR, Nielsen MM, Asmar F, Kooistra SM, Christophersen NS, Christensen LL, Borre M, Sorensen KD, et al: A dual program for translation regulation in cellular proliferation and differentiation. Cell 2014, 158:1281–1292.

35. Tuller T, Carmi A, Vestsigian K, Navon S, Dorfan Y, Zaborske J, Pan T, Dahan O, Furman I, Pilpel Y: An evolutionarily conserved mechanism for controlling the efficiency of protein translation. Cell 2010, 141:344–354.

36. Novoa EM, Ribas de Pouplana L: Speeding with control: codon usage, tRNAs, and ribosomes. Trends Genet 2012, 28:574–581.

37. Presnyak V, Alhusaini N, Chen YH, Martin S, Morris N, Kline N, Olson S, Weinberg D, Baker KE, Graveley BR, Coller J: Codon optimality is a major determinant of mRNA stability. Cell 2015, 160:1111–1124.

38. Kozomara A, Griffiths-Jones S: miRBase: annotating high confidence microRNAs using deep sequencing data. Nucleic Acids Res 2014, 42:D68–73.

39. Jones-Rhoades MW, Bartel DP, Bartel B: MicroRNAS and their regulatory roles in plants. Annu Rev Plant Biol 2006, 57:19–53.

40. Ray D, Kazan H, Cook KB, Weirauch MT, Najafabadi HS, Li X, Gueroussov S, Albu M, Zheng H, Yang A, et al: A compendium of RNA-binding motifs for decoding gene regulation. Nature 2013, 499:172–177.

41. A compendium of RNA-binding motifs for decoding gene regulation. [http://cisbp-rna.ccbr.utoronto.ca]

42. Kozak M: An analysis of 5’-noncoding sequences from 699 vertebrate messenger RNAs. Nucleic Acids Res 1987, 15:8125–8148.

43. Cavener DR, Ray SC: Eukaryotic start and stop translation sites. Nucleic Acids Res 1991, 19:3185–3192.

44. Nakagawa S, Niimura Y, Gojobori T, Tanaka H, Miura K: Diversity of preferred nucleotide sequences around the translation initiation codon in eukaryote genomes. Nucleic Acids Res 2008, 36:861–871.

45. Pisarev AV, Kolupaeva VG, Pisareva VP, Merrick WC, Hellen CU, Pestova TV: Specific functional interactions of nucleotides at key −3 and +4 positions flanking the initiation codon with components of the mammalian 48S translation initiation complex. Genes Dev 2006, 20:624–636.

46. Seol Y, Skinner GM, Visscher K, Buhot A, Halperin A: Stretching of homopolymeric RNA reveals single-stranded helices and base-stacking. Phys Rev Lett 2007, 98:158103.

47. Sponer J, Sponer JE, Mladek A, Jurecka P, Banas P, Otyepka M: Nature and magnitude of aromatic base stacking in DNA and RNA: Quantum chemistry, molecular mechanics, and experiment. Biopolymers 2013, 99:978–988.

48. Kacser H, Burns JA, Fell DA: The control of flux: 21 years on. Biochem Soc Trans 1995, 23:341–366.

49. Fell D: Understanding the control of metabolism. London: Portland Press; 1997.

50. Simms CL, Yan LL, Zaher HS: Ribosome Collision Is Critical for Quality Control during No-Go Decay. Mol Cell 2017, 68:361–373 e365.

51. Shoemaker CJ, Green R: Translation drives mRNA quality control. Nat Struct Mol Biol 2012, 19:594–601.

52. Zarai Y, Margaliot M, Tuller T: On the Ribosomal Density that Maximizes Protein Translation Rate. PLoS One 2016, 11:e0166481.

53. Marygold SJ, Roote J, Reuter G, Lambertsson A, Ashburner M, Millburn GH, Harrison PM, Yu Z, Kenmochi N, Kaufman TC, et al: The ribosomal protein genes and Minute loci of Drosophila melanogaster. Genome Biol 2007, 8:R216.

54. Firczuk H, Kannambath S, Pahle J, Claydon A, Beynon R, Duncan J, Westerhoff H, Mendes P, McCarthy JE: An in vivo control map for the eukaryotic mRNA translation machinery. Mol Syst Biol 2013, 9:635.

55. Sinturel F, Gerber A, Mauvoisin D, Wang J, Gatfield D, Stubblefield JJ, Green CB, Gachon F, Schibler U: Diurnal Oscillations in Liver Mass and Cell Size Accompany Ribosome Assembly Cycles. Cell 2017, 169:651–663 e614.

56. Princiotta MF, Finzi D, Qian SB, Gibbs J, Schuchmann S, Buttgereit F, Bennink JR, Yewdell JW: Quantitating protein synthesis, degradation, and endogenous antigen processing. Immunity 2003, 18:343–354.

57. von der Haar T: A quantitative estimation of the global translational activity in logarithmically growing yeast cells. BMC Syst Biol 2008, 2:87.

58. Li JJ, Bickel PJ, Biggin MD: System wide analyses have underestimated protein abundances and the importance of transcription in mammals. PeerJ 2014, 2:e270.

59. Li JJ, Biggin MD: Gene expression. Statistics requantitates the central dogma. Science 2015, 347:1066–1067.

60. dos Reis M, Wernisch L: Estimating translational selection in eukaryotic genomes. Mol Biol Evol 2009, 26:451–461.

61. Lynch M: The frailty of adaptive hypotheses for the origins of organismal complexity. Proc Natl Acad Sci U S A 2007, 104 Suppl 1:8597–8604.

62. Dana A, Tuller T: Mean of the typical decoding rates: a new translation efficiency index based on the analysis of ribosome profiling data. G3 (Bethesda) 2014, 5:73–80.

63. Johnstone TG, Bazzini AA, Giraldez AJ: Upstream ORFs are prevalent translational repressors in vertebrates. EMBO J 2016, 35:706–723.

64. Lima SA, Chipman LB, Nicholson AL, Chen YH, Yee BA, Yeo GW, Coller J, Pasquinelli AE: Short poly(A) tails are a conserved feature of highly expressed genes. Nat Struct Mol Biol 2017, 24:1057–1063.

65. Sen ND, Zhou F, Harris MS, Ingolia NT, Hinnebusch AG: eIF4B stimulates translation of long mRNAs with structured 5’ UTRs and low closed-loop potential but weak dependence on eIF4G. Proc Natl Acad Sci U S A 2016, 113:10464–10472.

