## Additional file 1 for "Quantitative Principles of *cis*-translational control by general mRNA sequence features in eukaryotes"

**Table S1. Contributions of sequence features to the variance in translation rates among endogenous mRNAs**

| Sequence feature(s) | $R^2$ feature vs TR |
|---|---|
| ***S. cerevisiae, correlation to initiation rate modelled from RP data (Shah et al., 2013)*** | |
| -4 to + 37 folding energy | 0.02 |
| CDS length | 0.35 |
| ***S. cerevisiae, correlation to initiation rate modelled from RP data (Weinberg et al., 2016)*** | |
| 5' most 70 nucleotides folding energy | 0.14 |
| CDS length | 0.24 |
| 5' UTR folding energy, 5' UTR %GC, 5' UTR #uAUGs, 5'UTR length, CDS length | 0.39 |
| ***S. cerevisiae, correlation to initiation rate modelled from RP data (Li et al., 2017)*** | |
| 5' UTR length | 0.05 |
| 5' UTR #uORFs | 0.14 |
| 5' UTR folding energy | 0.19 |
| -35/+28 motif | 0.33 |
| CDS length | 0.32 |
| codon frequency | 0.60 |
| all 5' UTR features, -35/+28 motif, CDS length | 0.58 |
| all 5' UTR features, -35/+28 motif, CDS length, codon frequency | 0.80 |
| ***mouse ES cell line (Chew et al., 2016)*** | |
| 5' UTR density of uAUGs | 0.04 |
| 5' UTR length | 0.01 |
| 5' UTR mean folding energy | 0.01 |
| -25 to +10 folding energy | 0.01 |
| +1 to + 35 folding energy | 0.01 |
| CDS mean folding energy | 0.06 |
| -10 to +13 motif | 0.02 |
| all above seven features | 0.14 |
| ***mouse liver (Janich et al., 2015)*** | |
| 5' UTR length | 0.05 |
| CDS length | 0.16 |
| 3' UTR length | 0.02 |


Chew, G.L., Pauli, A., and Schier, A.F. (2016). Conservation of uORF repressiveness and sequence features in mouse, human and zebrafish. Nat Commun *7*, 11663.

Janich, P., Arpat, A.B., Castelo-Szekely, V., Lopes, M., and Gatfield, D. (2015). Ribosome profiling reveals the rhythmic liver translatome and circadian clock regulation by upstream open reading frames. Genome Res. *25*, 1848-1859.

Li, J.J., Chew, G.L., and Biggin, M.D. (2017). Quantitating translational control: mRNA abundance-dependent and independent contributions and the mRNA sequences that specify them. Nucleic Acids Res. *45*, 11821-11836.

Shah, P., Ding, Y., Niemczyk, M., Kudla, G., and Plotkin, J.B. (2013). Rate-limiting steps in yeast protein translation. Cell *153*, 1589-1601.

Weinberg, D., Shah, P., Eichhorn, S., Hussmann, J., Plotkin, J., and Bartel, D. (2016). Improved ribosome-footprint and mRNA measurements provide insights into dynamics and regulation of yeast translation. Cell Reports *14*, 1787-1799.

**Table S2. Contributions of sequence features/motifs to the variance in translation rates across a set of heterologous mRNAs that have varying 5' UTRs fused upstream of a reporter CDS**

| Sequence feature(s) | $R^2$ feature vs TR |
|---|---|
| ***S. cerevisiae, in vitro, 18 natural 5' UTRs (Rojas-Duran and Gilbert, 2012)*** | |
| 5' UTR length | 0.18 |
| 5' UTR folding energy | 0.25 |
| ***S. cerevisiae, in vivo, 141 random mutations in region –4 to +37 (Shah et al., 2013)*** | |
| -4 to +37 folding energy | 0.22 |
| tRNA index | 0.02 |
| ***S. cerevisiae, in vivo, 2,041 random mutations in region -10 to -1 (Dvir et al., 2013)*** | |
| -15 to +50 folding energy | 0.18 |
| -10 to -1 uAUGs | 0.06 |
| -3 to -1 sequence motif | 0.29 |
| -10 to -1 kmer frequency motif | 0.19 |
| folding energy and sequence motifs (13 features) | 0.68 |
| ***S. cerevisiae, in vivo, 383 random mutations in -14 to -1 or in the CDS (Ben-Yehezkel et al., 2015)*** | |
| -14 to +39 mean folding energy | 0.12* |
| -7 to +33 folding energy | 0.20* |
| A at -3 | 0.12* |
| T at -3 | 0.12* |
| AUG context motif | 0.21* |
| ***S. cerevisae, in vivo, 500,000 random mutations in region -50 to -1 (Cuperus et al., 2017)**** | |
| -50 to -1 folding energy | 0.08 |
| -50 to -1 >100 13 mer PWMs / Neural network model | 0.47-0.62 |
| ***S. cerevisae, in vivo, 11,856 50 nucleotide segments of natural 5' UTRs (Cuperus et al., 2017)**** | |
| >100 13 mer PWMs / Neural network model trained on 500,000 random mutants | 0.60 |
| ***H. sapiens, in vivo, 300,000 random mutations in region -50 to -1 (Sample et al., 2018)*** | |
| -50 to -1 folding energy | 0.19 |
| -50 to -1 >100 8 mer PWMs / Neural network model | 0.93 |
| ***H. sapiens, in vivo, sequences -50 to -1 for 35,212 natural 5' UTRs (Sample et al., 2018)*** | |
| -50 to -1 >100 8 mer PWMs / Neural network model | 0.81 |
| ***H. sapiens, in vivo, 65,536 random mutations in region -6 to +5 (Noderer et al., 2014)*** | |
| PWM and di-nucleotide motif | 0.83 |

* $R^2$ coefficient of determination estimated as the square of a spearman rank correlation coefficient.
** An indirect measure of translation based on competative growth selection which could also capture effects on mRNA stability and transcription.


Ben-Yehezkel, T., Atar, S., Zur, H., Diament, A., Goz, E., Marx, T., Cohen, R., Dana, A., Feldman, A., Shapiro, E., *et al.* (2015). Rationally designed, heterologous S. cerevisiae transcripts expose novel expression determinants. RNA Biol *12*, 972-984.

Cuperus, J.T., Groves, B., Kuchina, A., Rosenberg, A.B., Jojic, N., Fields, S., and Seelig, G. (2017). Deep learning of the regulatory grammar of yeast 5' untranslated regions from 500,000 random sequences. Genome Res. *27*, 2015-2024.

Dvir, S., Velten, L., Sharon, E., Zeevi, D., Carey, L.B., Weinberger, A., and Segal, E. (2013). Deciphering the rules by which 5'-UTR sequences affect protein expression in yeast. Proc. Natl. Acad. Sci. USA *110*, E2792-2801.

Noderer, W.L., Flockhart, R.J., Bhaduri, A., Diaz de Arce, A.J., Zhang, J., Khavari, P.A., and Wang, C.L. (2014). Quantitative analysis of mammalian translation initiation sites by FACS-seq. Mol Syst Biol *10*, 748.

Rojas-Duran, M.F., and Gilbert, W.V. (2012). Alternative transcription start site selection leads to large differences in translation activity in yeast. RNA *18*, 2299-2305.

Sample, P.J., Wang, B., Reid, D.W., Presnyak, V., McFadyen, I., Morris, D.R., and Seelig, G. (2018). Human 5' UTR design and variant effect prediction from a massively parallel translation assay. BioRxiv.

Shah, P., Ding, Y., Niemczyk, M., Kudla, G., and Plotkin, J.B. (2013). Rate-limiting steps in yeast protein translation. Cell *153*, 1589-1601.

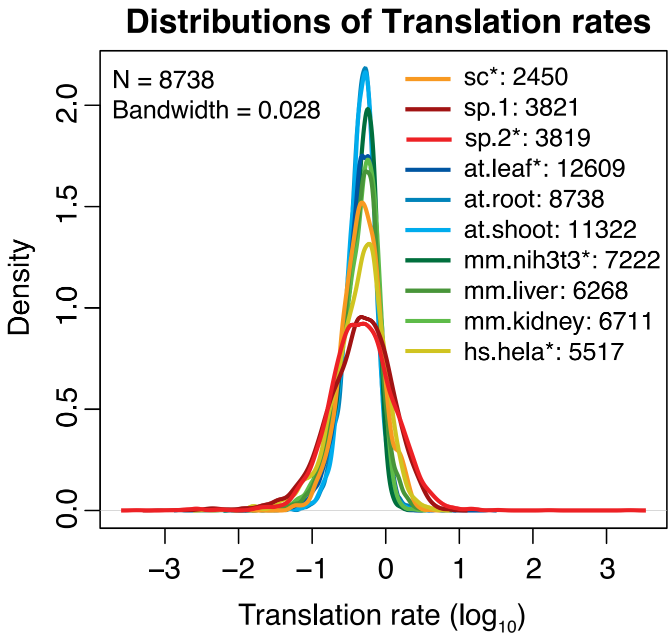


**Figure S1. The distributions of translation rates.** The $\log_{10}$ transformed ribosome profiling data have been scaled to have the same median while retaining their original variance. The species is denoted by a two letter code. The number of genes in each dataset is indicated. The example datasets are sc; sp.2; at.leaf; mm.nih3t3; and hs.hela (*).

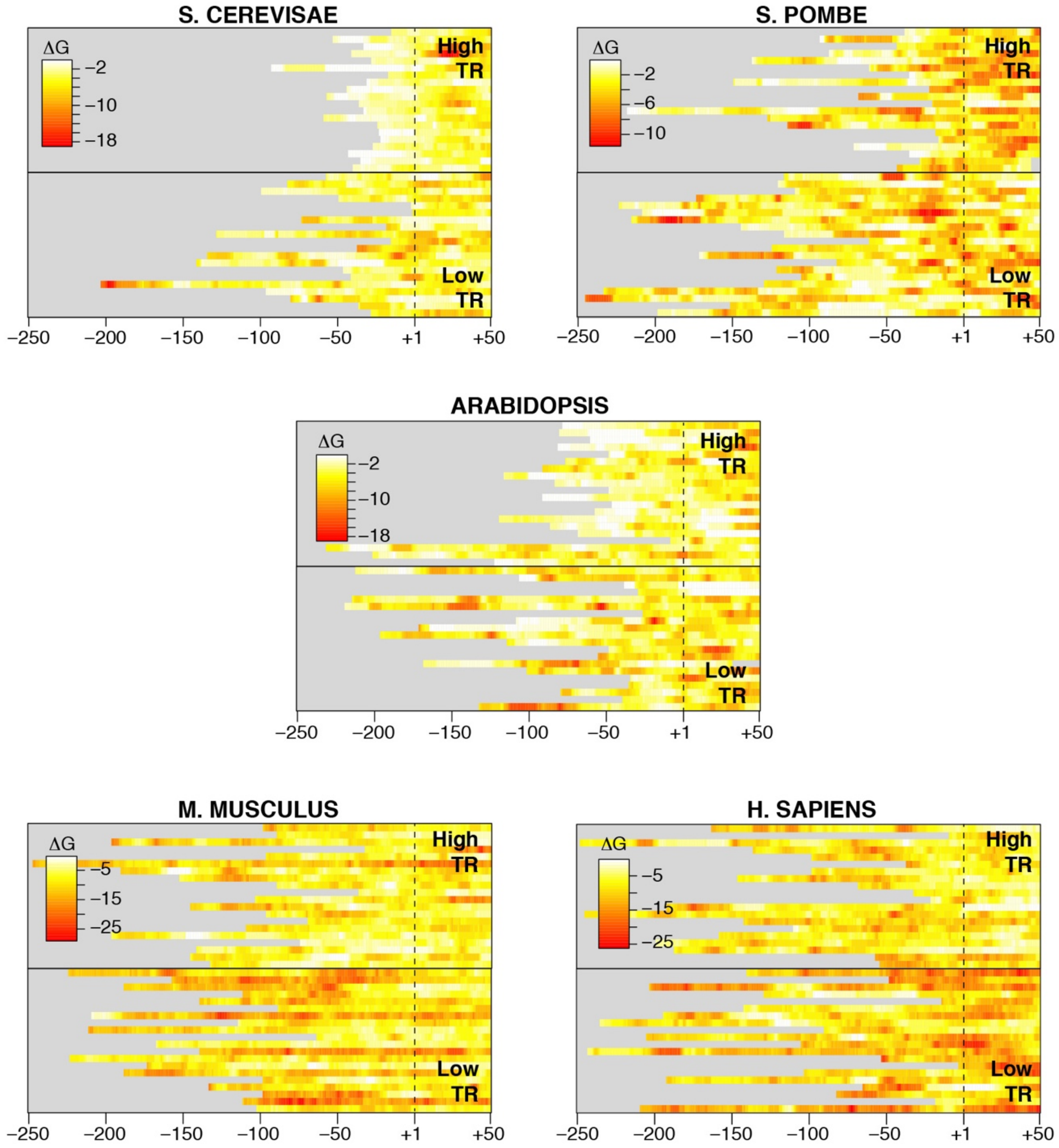


**Figure S2. The locations of secondary structures in individual mRNA are highly variable.** The heat maps show the free energy of RNA folding (ΔG kcal/mol) of 35 nucleotide windows for the 20 most highly translated mRNAs with 5'UTRs ≤250 nucleotides (high TR) and for the 20 most poorly translated mRNAs with the same length constraint (low TR). Windows representing every one nucleotide offset were calculated (*x*-axis). The mRNAs are aligned at the iAUG (dashed line). While the most strongly folded regions tend to be found in low TR mRNAs, the locations of the most folded regions within individual mRNAs are highly variable.

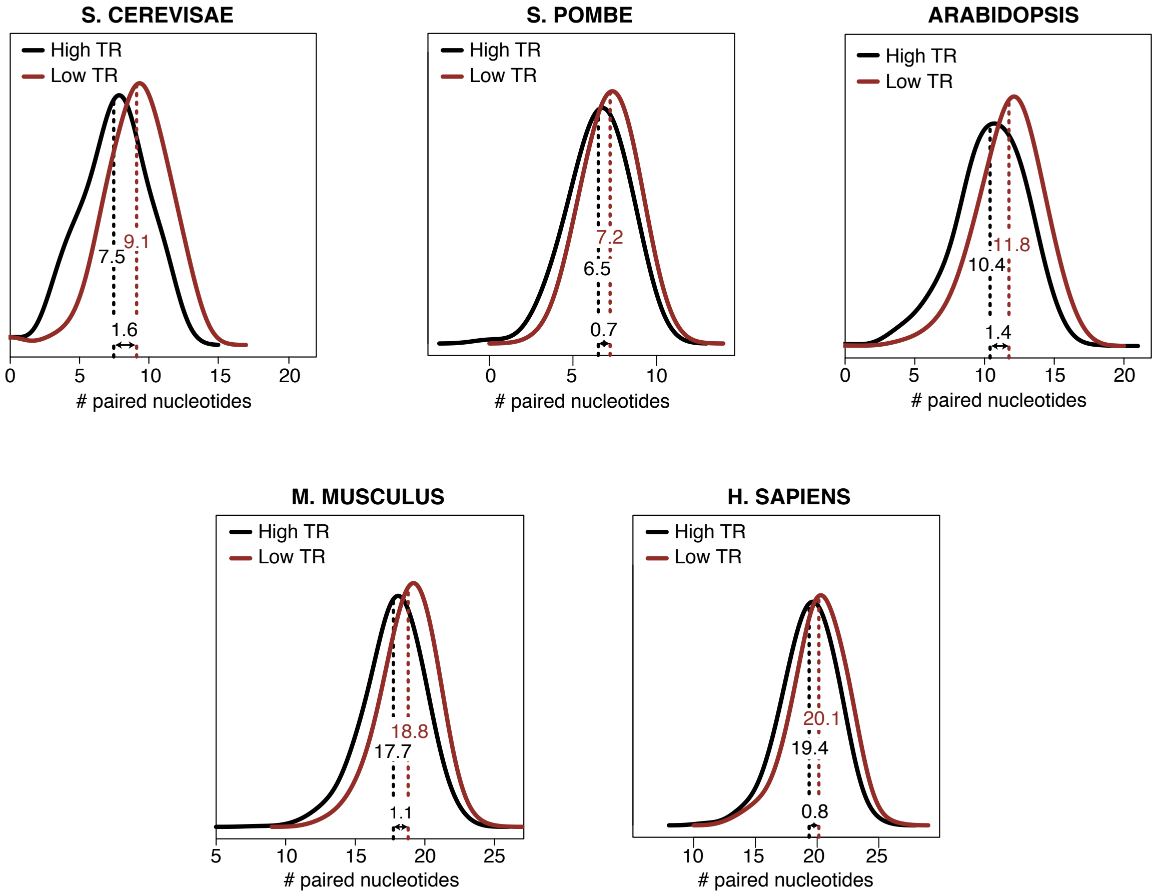


**Figure S3. The distributions of paired nucleotide number in the most folded windows of highly and poorly translated mRNAs.** The most folded ("min") windows were ranked based on the translation rate (TR) of the mRNA, and the 1$^{st}$ (high) and 10$^{th}$ (low) deciles identified. The distributions of the number of paired nucleotides in the most folded windows in each of the two cohorts are plotted. The vertical dotted line indicates the mean for each cohort. The result shows while the low TR cohort contains on average more paired nucleotides than the high TR cohort, there is a considerable overlap in structures between the two cohorts.

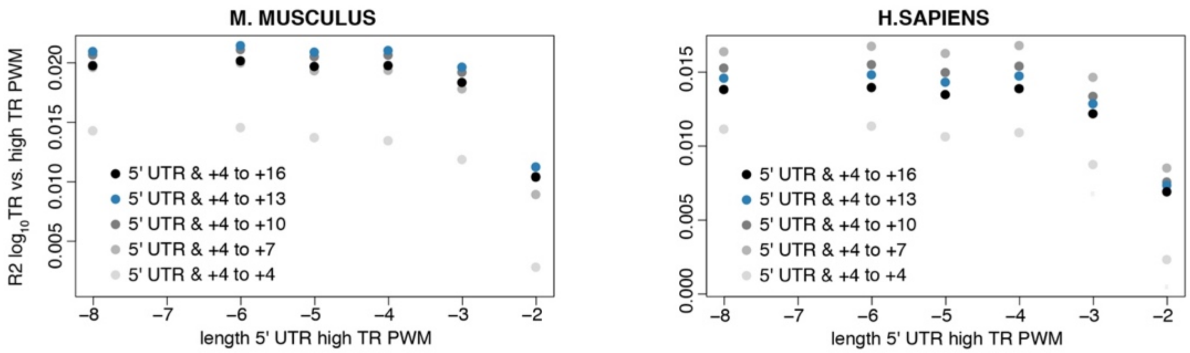


**Figure S4. Fine mapping the 5’ and 3’ boundaries of APEs in *M. musculus* and *H. sapiens*.** The $R^2$ coefficients of determination between $\log_{10}$ translation rates (TR) and PWM scores. The figure is as described in Figure 7 except that PWMs were chosen to more precisely map APE boundaries. The results show the APEs extend from -6 to +13 in *M. musculus* and -6 to +7 in *H. sapiens*.

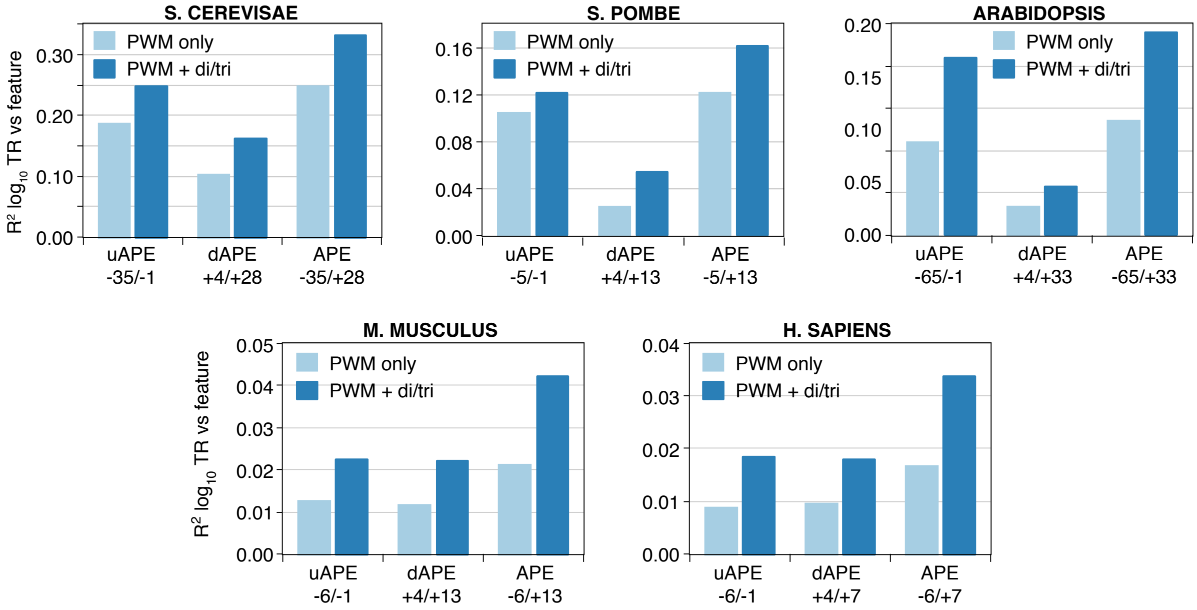


**Figure S5. AUG proximal elements (APEs) comprise sequences upstream and downstream of the iAUG and are best described by Position Weight Matrices (PWMs) and di- and tri-nucleotide frequencies.** The $R^2$ coefficients of determination between $\log_{10}$ translation rate (TR) and feature(s) describing the iAUG upstream portion of the APE (uAPE); the iAUG downstream portion of the APE (dAPE); and the complete APE. Results are shown for a model employing only the PWM score and for a multivariate model combining the PWM score with a BIC selected subset of di and tri-nucleotide frequencies. The models are described in Additional file 5.

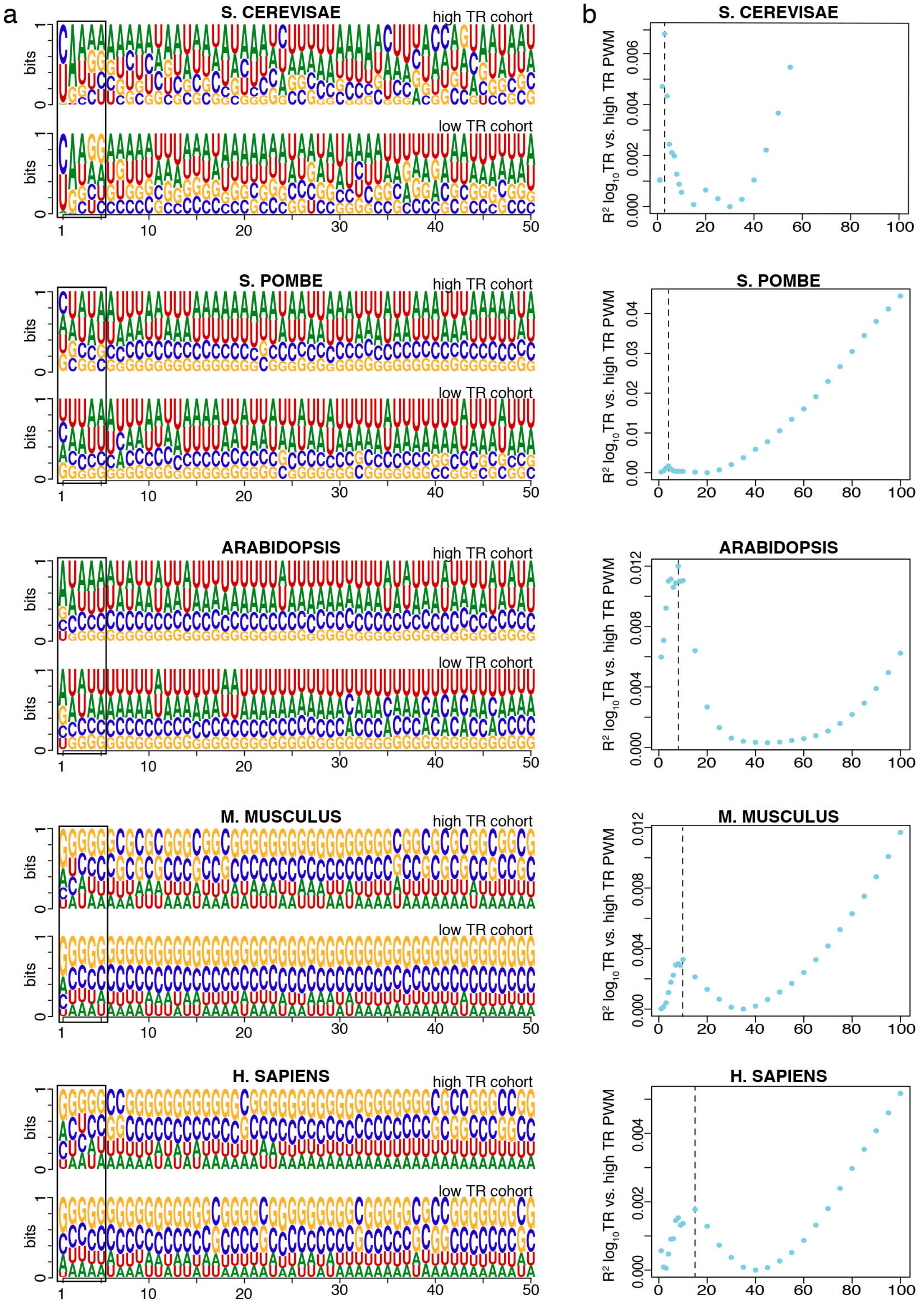


**Figure S6. A 5' cap element.** (a) PWMs for the 10% of mRNAs with the highest translation rate (high TR cohort) and the 10% with the lowest rate (low TR cohort). Sequence logos show the frequency of each nucleotide at each position relative to the first nucleotide of the transcript (i.e. the 5' cap). Only 5'UTR sequences that lie 5' of the APE were included in the analysis. (b) The $R^2$ coefficients of determination between $\log_{10}$ TR and PWM scores. PWMs of varying lengths were built from the sequences of the high TR cohort. Log odds scores were then calculated for all mRNAs that completely contained a given PWM. PWMs extending 3’ from the 5' cap in 1 nucleotide or 5 nucleotide increments were tested (x-axis, right to left). A local maxima in $R^2$ values is seen between 3 to 15 nucleotides from the 5' cap, depending on the species. Because this 5'cap element only controls less than 1.2% of the variance in TR and to simplify our models, the PWM score of the first 5 nucleotides was used in a model for the 5'ofAPE region for all species, together with a BIC selected subset of di and tri-nucleotides from the entire 5'ofAPE region.

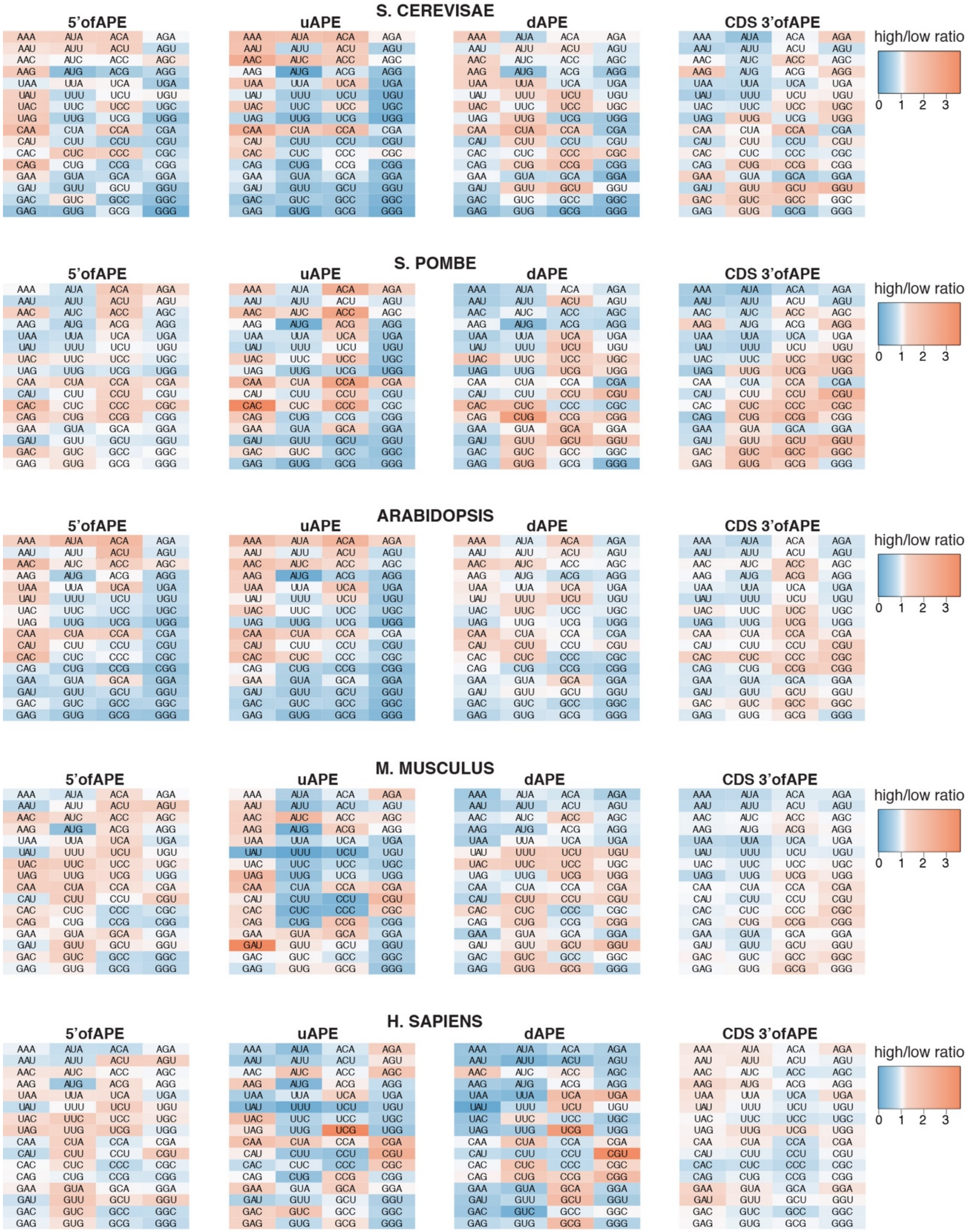


**Figure S7. Tri-nucleotides regulating translation.** The heat maps show the frequency of each tri-nucleotide in the most highly translated 10% of genes divided by its frequency in the most poorly translated 10% of genes ((TR high / TR low) ratios). Results are presented for four mRNA regions of each species.

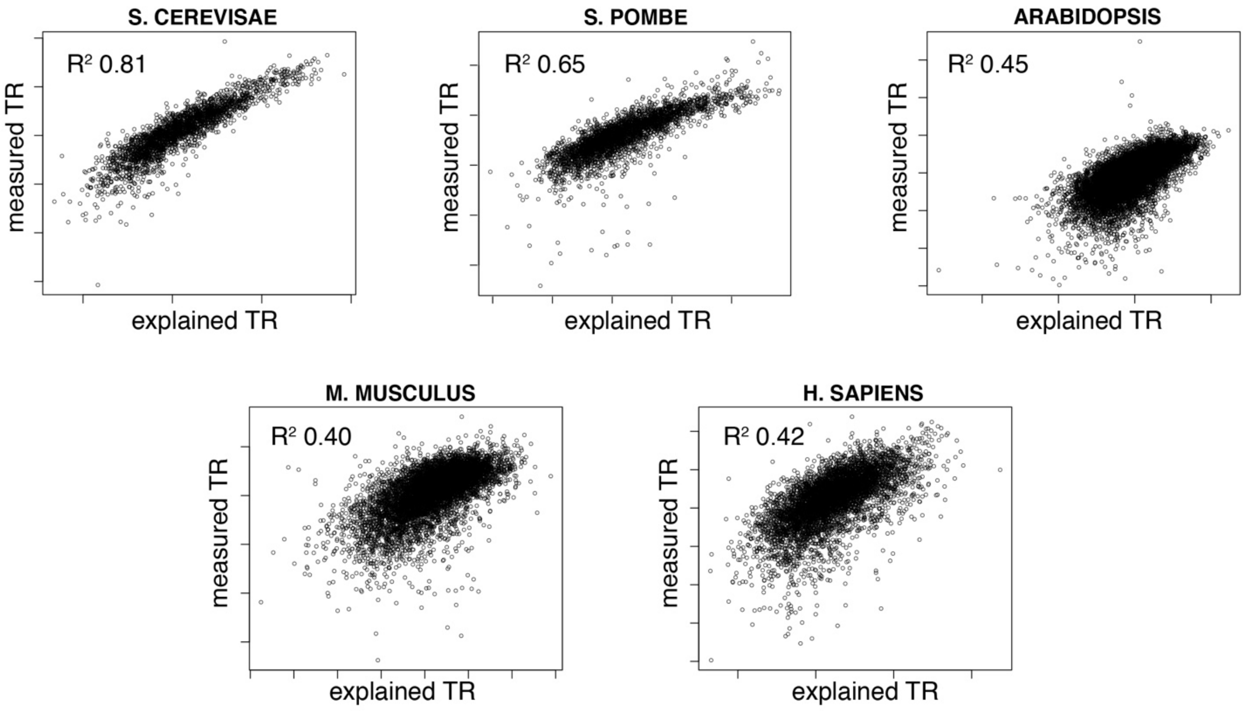


**Figure S8. Multivariate models for the five general features.** Scatter plots show the relationship between measured translation rates (*y*-axis) and translation rates explained by multivariate models for the five general features (*x*-axis). The R2 coefficients of determination are given.

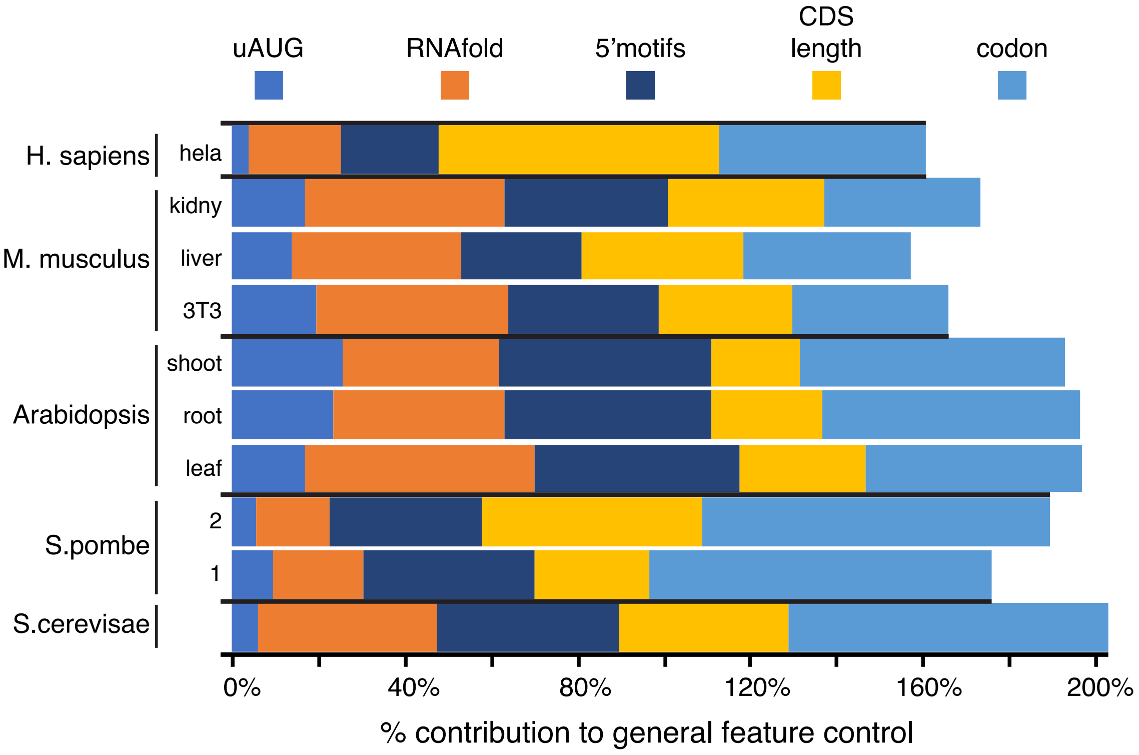


**Figure S9.** (a) For each feature separately, its $R^2$ coefficient of determination vs $\log_{10}$ translation rate is given as a percent the $R^2$ coefficient for a linear multivariate model for all five features. The result shows that the sum of these percent contributions is much greater than the variance in translation explained by the multivariate five-feature model. The $R^2$ values and the values plotted for each feature and the combined model are given in Additional file 9.

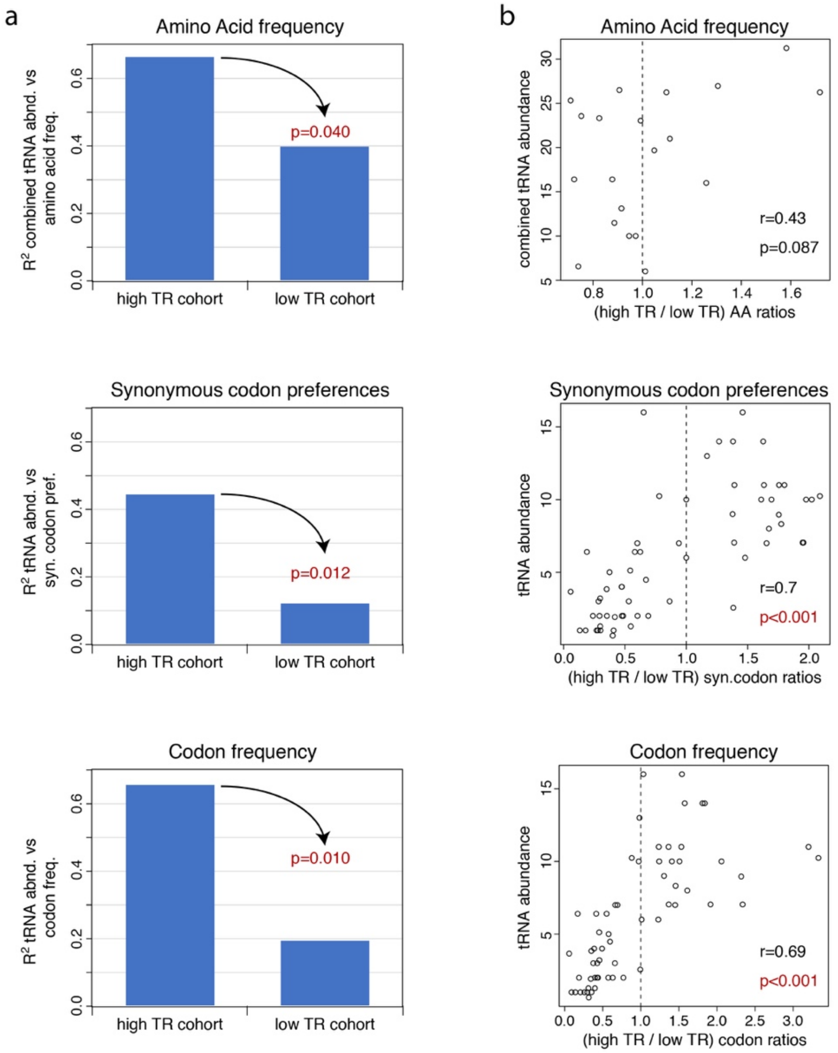


**Figure S10. Control by amino acid frequencies and synonymous codon preferences correlates with tRNA abundances.** The relationship between tRNA abundances and control by amino acid content and synonymous codon preferences was tested in *S. cerevisiae* because estimates of effective tRNA abundances are particularly well established for this species. The frequencies of amino acids (AA), or codons (codon), or the preferences for synonymous codon (syn.codon) were determined separately for the most highly translated 10% of genes (high TR) and for the most poorly translated 10% of genes (low TR). Genes not encoding all 20 amino acids were excluded from the analysis. (a) The coefficient of determination ($R^2$) for the high TR cohort or low TR cohort AA, syn.codon or codon frequencies vs their cognate tRNA abundances. For AA, the frequencies of all cognate tRNAs for each amino acid were summed to give a combined tRNA abundance. *p*-values testing if the correlation of tRNA abundance with the high TR cohort is greater than that with the low TR cohort are given, with significant *p*-values shown in red. The High TR mean frequencies correlate more strongly with tRNA abundances than do the low TR frequencies, indicating that translation of high TR mRNAs uses the cellular population of amino acylated tRNAs more efficiently than translation of low TR mRNAs. (b) The ratios between AA, syn.codon or codon frequencies in the high TR cohort divided by those in the low TR cohort were determined. Ratios > 1 thus indicate a larger frequency in high TR cohorts than in low TR cohorts. Scatter plots are shown between these (high TR /low TR) ratios and tRNA abundance along with the Pearson correlation coefficients (r) and *p*-values testing if the correlations are significant (significant *p*-values in red). Dashed vertical lines indicate a ratio of 1. The Pearson correlation coefficients range from +0.43 – +0.70, establishing that codons for high abundance tRNAs are more prevalent in highly translated mRNAs, whereas codons for low abundance tRNAs are more prevalent in poorly translated mRNAs.
