## Additional file 11 for "Quantitative Principles of *cis*-translational control by general mRNA sequence features in eukaryotes": Code.pdf

### Code for reproducing the results in "Principles of cis-translational control by general mRNA sequence features in six model eukaryotes"

The intermediate files and final output files have been generated and stored in subdirectories. The code chunks that take time to generate intermediate files will not run by default.

```
library(knitr)
opts_chunk$set(tidy.opts=list(width.cutoff=60), tidy=TRUE)
```

#### Required R Packages

```
library(parallel)
library(MASS)
if (!require("readxl")) {
    install.packages("readxl", dependencies = TRUE)
    library(readxl)
}
```

```
## Loading required package: readxl
```

```
if (!require("lmodel2")) {
    install.packages("lmodel2", dependencies = TRUE)
    library(lmodel2)
}
```

```
## Loading required package: lmodel2
```

```
if (!require("stringi")) {
    install.packages("stringi", dependencies = TRUE)
    library(stringi)
}
```

```
## Loading required package: stringi
```

```
if (!require("stringr")) {
    install.packages("stringr", dependencies = TRUE)
    library(stringr)
}
```

```
## Loading required package: stringr
```

```
if (!require("leaps")) {
    install.packages("leaps", dependencies = TRUE)
    library(leaps)
}
```

```
## Loading required package: leaps
```

```
if (!require("Biostrings")) {
    install.packages("Biostrings", dependencies = TRUE)
```

```r
    library(Biostrings)
}
```

```
## Loading required package: Biostrings

## Loading required package: BiocGenerics

##
## Attaching package: 'BiocGenerics'

## The following objects are masked from 'package:parallel':
##
##     clusterApply, clusterApplyLB, clusterCall, clusterEvalQ,
##     clusterExport, clusterMap, parApply, parCapply, parLapply,
##     parLapplyLB, parRapply, parSapply, parSapplyLB

## The following objects are masked from 'package:stats':
##
##     IQR, mad, xtabs

## The following objects are masked from 'package:base':
##
##     anyDuplicated, append, as.data.frame, as.vector, cbind,
##     colnames, do.call, duplicated, eval, evalq, Filter, Find, get,
##     grep, grepl, intersect, is.unsorted, lapply, lengths, Map,
##     mapply, match, mget, order, paste, pmax, pmax.int, pmin,
##     pmin.int, Position, rank, rbind, Reduce, rownames, sapply,
##     setdiff, sort, table, tapply, union, unique, unlist, unsplit

## Loading required package: S4Vectors

## Loading required package: stats4

## Loading required package: IRanges

## Loading required package: XVector
```

```r
if (!require("cocor")) {
    install.packages("cocor", dependencies = TRUE)
    library(cocor)
}
```

```
## Loading required package: cocor
```

```r
if (!require("motifStack")) {
    install.packages("motifStack", dependencies = TRUE)
    library(motifStack)
}
```

```
## Loading required package: motifStack

## Loading required package: grImport

## Loading required package: grid

## Loading required package: XML

## Loading required package: MotIV

##
## Attaching package: 'MotIV'
```

```
## The following object is masked from 'package:stats':
##
##     filter
```

```
## Loading required package: ade4
```

```
##
## Attaching package: 'ade4'
```

```
## The following object is masked from 'package:Biostrings':
##
##     score
```

```
## The following object is masked from 'package:IRanges':
##
##     score
```

```
## The following object is masked from 'package:BiocGenerics':
##
##     score
```

```
if (!require("gplots")) {
    install.packages("gplots", dependencies = TRUE)
    library(gplots)
}
```

```
## Loading required package: gplots
```

```
##
## Attaching package: 'gplots'
```

```
## The following object is masked from 'package:IRanges':
##
##     space
```

```
## The following object is masked from 'package:stats':
##
##     lowess
```

```
if (!require("RColorBrewer")) {
    install.packages("RColorBrewer", dependencies = TRUE)
    library(RColorBrewer)
}
```

```
## Loading required package: RColorBrewer
```

```
if (!require("reshape2")) {
    install.packages("reshape2", dependencies = TRUE)
    library(reshape2)
}
```

```
## Loading required package: reshape2
```

```
if (!require("ggplot2")) {
    install.packages("ggplot2", dependencies = TRUE)
    library(ggplot2)
}
```

```
## Loading required package: ggplot2
```

```
if (!require("zoo")) {
    install.packages("zoo", dependencies = TRUE)
```

```r
    library(zoo)
}
```

```
## Loading required package: zoo

##
## Attaching package: 'zoo'

## The following objects are masked from 'package:base':
##
##     as.Date, as.Date.numeric
```

```r
if (!require("lattice")) {
    install.packages("lattice", dependencies = TRUE)
    library(lattice)
}
```

```
## Loading required package: lattice
```

```r
if (!require("gdata")) {
    install.packages("gdata", dependencies = TRUE)
    library(lattice)
}
```

```
## Loading required package: gdata

## gdata: read.xls support for 'XLS' (Excel 97-2004) files ENABLED.

##

## gdata: read.xls support for 'XLSX' (Excel 2007+) files ENABLED.

##
## Attaching package: 'gdata'

## The following object is masked from 'package:IRanges':
##
##     trim

## The following object is masked from 'package:stats4':
##
##     nobs

## The following object is masked from 'package:BiocGenerics':
##
##     combine

## The following object is masked from 'package:stats':
##
##     nobs

## The following object is masked from 'package:utils':
##
##     object.size
```

```r
if (!require("viridis")) {
    install.packages("viridis", dependencies = TRUE)
    library(lattice)
}
```

```
## Loading required package: viridis
```

```
## Loading required package: viridisLite
```

```
if (!require("xlsx")) {
    install.packages("xlsx", dependencies = TRUE)
    library(lattice)
}
```

```
## Loading required package: xlsx
## Loading required package: rJava
## Loading required package: xlsxjars
```

#### Preprocess data

```
# a function to resolve duplicated short gene names (gene
# IDs) for sequences
resolve_dup_genes <- function(long_seqs, short_names, tissue_long_seqs = NULL) {
    # long_seqs: a character vector of UTR sequences, whose names
    # are long gene names (e.g., transcript IDs) short_names: a
    # character vector of short gene names (e.g., gene IDs);
    # short_names are mathed with long_seqs tissue_long_seqs
    # (optional): a character vector of one tissue's UTR
    # sequences, whose names are long gene names (e.g.,
    # transcript IDs); if not supplied, the longest sequence will
    # be picked for each short name
    long_names <- names(long_seqs)
    dup_idx <- which(duplicated(short_names))
    if (length(dup_idx) > 0) {
        dup_genes <- unique(short_names[dup_idx])
        non_dup_idx <- which(!(short_names %in% dup_genes))
        IDmap <- long_names[non_dup_idx]
        names(IDmap) <- short_names[non_dup_idx]
        if (!is.null(tissue_long_seqs)) {
            checkif <- long_names %in% names(tissue_long_seqs)
        }
        for (dup_gene in dup_genes) {
            if (!is.null(tissue_long_seqs)) {
                tmp_idx <- which(short_names == dup_gene & checkif)
            } else {
                tmp_idx <- which(short_names == dup_gene)
            }
            tmp <- nchar(long_seqs[tmp_idx])
            tmp_idx <- tmp_idx[which.max(tmp)]
            IDmap_add <- long_names[tmp_idx]
            IDmap_name_old <- names(IDmap)
            IDmap_name_add <- short_names[tmp_idx]
            IDmap <- c(IDmap, IDmap_add)
            names(IDmap) <- c(IDmap_name_old, IDmap_name_add)
        }
    } else {
        IDmap <- long_names
        names(IDmap) <- short_names
    }
```

```
    return(IDmap)
    # output: IDmap: a character vector of short names, with
    # names as long names
}

############################## S. cerevisae ##############################

# translational rates and features
sc.data <- read_xlsx("data/Dataset S6 mRNA seq features.xlsx",
    sheet = 1, range = "A23:Q2473")
sc.data <- as.data.frame(sc.data)

# CDS sequences
sc.CDS_seqs <- sc.data$"CDS sequence"
names(sc.CDS_seqs) <- sc.data$ORF

# 5' UTR sequences
sc.UTR_seqs <- sc.data$"5' UTR sequence"
names(sc.UTR_seqs) <- sc.data$ORF

# genes
sc.genes <- sc.data$ORF

# TR data
sc.log10.TR <- sc.data$"log10 TR"
names(sc.log10.TR) <- sc.genes

# mRNA data
sc.log10.mRNA <- sc.data$"log10 RNA"
names(sc.log10.mRNA) <- sc.genes

# UTR sequences
sc.UTR_seqs <- sc.UTR_seqs[sc.genes]

# CDS sequences
sc.CDS_seqs <- sc.CDS_seqs[sc.genes]

# poly A tail length
sc.polyA_lens <- sc.data$"poly-A tail length"
names(sc.polyA_lens) <- sc.genes

# save the processed data to an .Rdata file
save(sc.data, sc.genes, sc.log10.TR, sc.log10.mRNA, sc.UTR_seqs,
    sc.CDS_seqs, sc.polyA_lens, file = "processed_data_w_polyA/sc.Rdata")


############################## S. pombe ##############################

# CDS sequences (downloaded from pombase.org, date 11/1/17)
sp.CDS_fa <- scan("data/s_pombe_cds.fa", what = character(),
    sep = "\n")
name_idx <- which(startsWith(str = sp.CDS_fa, pattern = ">"))
n_seq <- length(name_idx)
```

```r
sp.genes <- gsub(pattern = ">", replacement = "", x = sp.CDS_fa[name_idx])
sp.CDS_seqs <- sapply(1:n_seq, FUN = function(i) {
    start_idx <- name_idx[i] + 1
    if (i < n_seq) {
        stop_idx <- name_idx[i + 1] - 1
    } else {
        stop_idx <- length(sp.CDS_fa)
    }
    paste(sp.CDS_fa[start_idx:stop_idx], collapse = "")
})
names(sp.CDS_seqs) <- sp.genes
rm(sp.CDS_fa, sp.genes)

# 5' UTR sequences (downloaded from pombase.org, date
# 10/18/17)
sp.UTR_fa <- scan("data/s_pombe_5UTR.fa", what = character(),
    sep = "\n")
name_idx <- which(startsWith(str = sp.UTR_fa, pattern = ">"))
n_seq <- length(name_idx)
sp.genes <- sapply(strsplit(gsub(pattern = ">", replacement = "",
    x = sp.UTR_fa[name_idx]), "\\|"), FUN = function(x) x[1])
sp.UTR_seqs <- sapply(1:n_seq, FUN = function(i) {
    start_idx <- name_idx[i] + 1
    if (i < n_seq) {
        stop_idx <- name_idx[i + 1] - 1
    } else {
        stop_idx <- length(sp.UTR_fa)
    }
    paste(sp.UTR_fa[start_idx:stop_idx], collapse = "")
})
names(sp.UTR_seqs) <- sp.genes
rm(sp.UTR_fa, sp.genes)

################## first dataset ##################

# translational rates and features
sp.data <- read_xlsx("data/Subtelny GSE52809_Pombe.xlsx", sheet = 1,
    range = "A5:I3950")
sp.data <- as.data.frame(sp.data)
## the first two columns are exactly the same
sp.data <- sp.data[, -1]

# genes
sp.genes <- sapply(strsplit(sp.data$"Gene name", "\\."), FUN = function(x) paste(x[1:2],
    collapse = "."))

# TR data
sp.log10.TR <- sp.data$"log 10 TE RPKM"
names(sp.log10.TR) <- sp.genes

# mRNA data
sp.log10.mRNA <- sp.data$"log 10 mRNA RPKM"
names(sp.log10.mRNA) <- sp.genes
```

```r
# UTR sequences
sp.UTR_seqs <- sp.UTR_seqs[sp.genes]

# CDS sequences
sp.CDS_seqs <- sp.CDS_seqs[sp.genes]

# poly A tail length
sp.polyA_lens <- sp.data$"Mean poly A tail length"
names(sp.polyA_lens) <- sp.genes

# remove the genes with missing information
sp.rm_idx <- which(is.na(sp.log10.TR) | is.na(sp.log10.mRNA) |
    is.na(sp.UTR_seqs) | is.na(sp.CDS_seqs) | is.na(sp.polyA_lens))
sp.data <- sp.data[-sp.rm_idx, ]
sp.genes <- sp.genes[-sp.rm_idx]
sp.log10.TR <- sp.log10.TR[-sp.rm_idx]
sp.log10.mRNA <- sp.log10.mRNA[-sp.rm_idx]
sp.UTR_seqs <- sp.UTR_seqs[-sp.rm_idx]
sp.CDS_seqs <- sp.CDS_seqs[-sp.rm_idx]
sp.polyA_lens <- sp.polyA_lens[-sp.rm_idx]

# save the processed data to an .Rdata file
save(sp.data, sp.genes, sp.log10.TR, sp.log10.mRNA, sp.UTR_seqs,
    sp.CDS_seqs, sp.polyA_lens, file = "processed_data_w_polyA/sp.Rdata")

################## second dataset ##################

# translational rates (preprocessing)
sp.alt.data <- read_xlsx("data/Pombe Mata 10.23.17 N_present_all_counts.xlsx",
    sheet = 1, range = "A1:E5161")
sp.alt.data <- as.data.frame(sp.alt.data)
for (j in 2:ncol(sp.alt.data)) {
    sp.alt.data[, j] <- sp.alt.data[, j]/(sum(sp.alt.data[, j])/1e+06)  # RPM
}
colnames(sp.alt.data)[1] <- "Systematic_ID"

## annotated ORF lengths
sp.alt.CDS_lens <- read_xlsx("data/Pombe Mata 10.23.17 N_present_all_counts.xlsx",
    sheet = 2, range = "A1:B5124")
sp.alt.CDS_lens <- as.data.frame(sp.alt.CDS_lens)

## make the data frame
sp.alt.data <- merge(sp.alt.data, sp.alt.CDS_lens, by = "Systematic_ID")
for (j in 2:(ncol(sp.alt.data) - 1)) {
    sp.alt.data[, j] <- sp.alt.data[, j]/(sp.alt.data[, ncol(sp.alt.data)]/1000)  # RPKM
}
sp.alt.data$RD_mean <- rowMeans(sp.alt.data[, 2:3])
sp.alt.data$RNA_mean <- rowMeans(sp.alt.data[, 4:5])

## thresholded to >= 1 RPKM for RNA density and > 0 for RD
## density
sp.alt.data <- sp.alt.data[sp.alt.data$RNA_mean >= 1 & sp.alt.data$RD_mean >
    0, ]
```

```r
sp.alt.data$TE <- sp.alt.data$RD_mean/sp.alt.data$RNA_mean

# genes
sp.alt.genes <- sp.alt.data$Systematic_ID

# TR data
sp.alt.log10.TR <- log(sp.alt.data$TE, 10)
names(sp.alt.log10.TR) <- sp.alt.genes

# mRNA data
sp.alt.log10.mRNA <- log(sp.alt.data$RNA_mean, 10)
names(sp.alt.log10.mRNA) <- sp.alt.genes

# UTR sequences
sp.alt.UTR_seqs <- sp.UTR_seqs[sp.alt.genes]

# CDS sequences
sp.alt.CDS_seqs <- sp.CDS_seqs[sp.alt.genes]

# poly A tail length
sp.alt.polyA_lens <- sp.polyA_lens[sp.alt.genes]
names(sp.alt.polyA_lens) <- sp.alt.genes

# remove the genes with missing information
sp.alt.rm_idx <- which(is.na(sp.alt.log10.TR) | is.na(sp.alt.log10.mRNA) |
    is.na(sp.alt.UTR_seqs) | is.na(sp.alt.CDS_seqs) | is.na(sp.alt.polyA_lens))
sp.alt.data <- sp.alt.data[-sp.alt.rm_idx, ]
sp.alt.genes <- sp.alt.genes[-sp.alt.rm_idx]
sp.alt.log10.TR <- sp.alt.log10.TR[-sp.alt.rm_idx]
sp.alt.log10.mRNA <- sp.alt.log10.mRNA[-sp.alt.rm_idx]
sp.alt.UTR_seqs <- sp.alt.UTR_seqs[-sp.alt.rm_idx]
sp.alt.CDS_seqs <- sp.alt.CDS_seqs[-sp.alt.rm_idx]
sp.alt.polyA_lens <- sp.alt.polyA_lens[-sp.alt.rm_idx]

# save the processed data to an .Rdata file
save(sp.alt.genes, sp.alt.log10.TR, sp.alt.log10.mRNA, sp.alt.UTR_seqs,
    sp.alt.CDS_seqs, sp.alt.polyA_lens, file = "processed_data_w_polyA/sp.alt.Rdata")


############################## Arabidopsis ##############################

# CDS sequences
at.CDS_seqs <- readRDS(file = "data/Arabidopsis_CDS.rds")
at.CDS_seqs_short_names <- sapply(strsplit(names(at.CDS_seqs),
    "\\."), FUN = function(x) x[1])

# 5' UTR sequences
at.UTR_seqs <- readRDS(file = "data/Arabidopsis_UTRs.rds")
at.UTR_seqs_short_names <- sapply(strsplit(names(at.UTR_seqs),
    "\\."), FUN = function(x) x[1])

# Hsu data for root and shoot
at.hsu_data <- read_xlsx("data/Hsu et al counts_cds_Hsu.xlsx",
```

```
    sheet = 1, range = "A7:M27213")
at.hsu_data <- as.data.frame(at.hsu_data)
colnames(at.hsu_data)[1] <- "gene_id"
for (j in 2:ncol(at.hsu_data)) {
    at.hsu_data[, j] <- at.hsu_data[, j]/(sum(at.hsu_data[, j])/1e+06)  # RPM
}

################## Leaf ##################

# Leaf translational rates and features (data preprocessing
# in ``030218_mouse_and_arabidopsis_models.Rmd'')
at.leaf.data <- readRDS("data/030218_arab_data.rds")

# genes
at.leaf.genes <- at.leaf.data$GeneID

# TR data
at.leaf.log10.TR <- at.leaf.data$Log10..TE.Dark
names(at.leaf.log10.TR) <- at.leaf.genes

# mRNA data
at.leaf.log10.mRNA <- at.leaf.data$log.10.mRNA.Dark
names(at.leaf.log10.mRNA) <- at.leaf.genes

# UTR sequences
at.leaf.UTR_seqs <- at.UTR_seqs[at.leaf.genes]

# CDS sequences
at.leaf.CDS_seqs <- at.CDS_seqs[at.leaf.genes]

# poly A tail length
at.leaf.polyA_lens <- at.leaf.data$"poly-A tail length"
names(at.leaf.polyA_lens) <- at.leaf.genes

# save the processed data to an .Rdata file
save(at.leaf.data, at.leaf.genes, at.leaf.log10.TR, at.leaf.log10.mRNA,
    at.leaf.UTR_seqs, at.leaf.CDS_seqs, at.leaf.polyA_lens, file = "processed_data_w_polyA/at.leaf.Rdata

############### Root & Shoot ###############

# both tissues have short gene names, so we will resolve the
# duplicated short gene names
at.IDmap <- resolve_dup_genes(long_seqs = at.UTR_seqs, short_names = at.UTR_seqs_short_names,
    tissue_long_seqs = at.leaf.UTR_seqs)

saveRDS(at.IDmap, file = "processed_data_w_polyA/at.IDmap.rds")

# a function to resolve duplicated short gene names (gene
# IDs) for root and shoot data based on the leaf tissue CDS
# sequences
resolve_dup_genes_root_shoot_data <- function(data, leaf_CDS_seqs) {
    # data: a data frame with two key columns: 'gene_id' contains
    # duplicated short gene names; 'Annotated ORF length'
```

```r
    # contains ORF lengths, which are CDS lengths - 3
    # leaf_CDS_seqs: a character vector of leaf tissue's CDS
    # sequences
    leaf_short_names <- sapply(strsplit(names(leaf_CDS_seqs),
        "\\."), FUN = function(x) x[1])
    short_names <- data$gene_id
    dup_idx <- which(duplicated(short_names))
    if (length(dup_idx) > 0) {
        dup_genes <- unique(short_names[dup_idx])
        non_dup_idx <- which(!(short_names %in% dup_genes))
        data_tmp <- data[non_dup_idx, ]
        for (dup_gene in dup_genes) {
            tmp_idx <- which(short_names == dup_gene)
            CDS_len <- nchar(leaf_CDS_seqs[leaf_short_names ==
                dup_gene])
            ORF_lens <- data[tmp_idx, "Annotated ORF length"]
            tmp_idx <- tmp_idx[ORF_lens == CDS_len - 3]
            data_tmp <- rbind(data_tmp, data[tmp_idx[1], ])
        }
    } else {
        data_tmp <- data
    }
    return(data_tmp)
}

################## Root ##################

# Root translational rates (preprocessing) annotated ORF
# lengths
at.root.ORF_lens <- read_xlsx("data/Hsu et al counts_cds_Hsu.xlsx",
    sheet = 2, range = "A3:E20986")
at.root.ORF_lens <- as.data.frame(at.root.ORF_lens)
at.root.ORF_lens <- at.root.ORF_lens[, c("gene_id", "Annoated ORF length")]
## make the data frame
at.root.data <- merge(at.hsu_data, at.root.ORF_lens, by = "gene_id")
for (j in 2:(ncol(at.root.data) - 1)) {
    at.root.data[, j] <- at.root.data[, j]/(at.root.data[, ncol(at.root.data)]/1000)  # RPKM
}
at.root.data$RD_mean <- rowMeans(at.root.data[, 2:4])
at.root.data$RNA_mean <- rowMeans(at.root.data[, 8:10])
## thresholded to >= 1 RPKM for RNA density and > 0 for RD
## density
at.root.data <- at.root.data[at.root.data$RNA_mean >= 1 & at.root.data$RD_mean >
    0, ]
at.root.data$TE <- at.root.data$RD_mean/at.root.data$RNA_mean

# remove duplicate gene names based on the leaf tissue
at.root.data <- resolve_dup_genes_root_shoot_data(data = at.root.data,
    leaf_CDS_seqs = at.leaf.CDS_seqs)

# genes
at.root.genes <- at.root.data$gene_id
```

```r
# TR data
at.root.log10.TR <- log(at.root.data$TE, 10)
names(at.root.log10.TR) <- at.root.genes

# mRNA data
at.root.log10.mRNA <- log(at.root.data$RNA_mean, 10)
names(at.root.log10.mRNA) <- at.root.genes

# UTR sequences
at.root.UTR_seqs <- at.UTR_seqs[at.IDmap[at.root.genes]]
names(at.root.UTR_seqs) <- at.root.genes

# CDS sequences
at.root.CDS_seqs <- at.CDS_seqs[at.IDmap[at.root.genes]]
names(at.root.CDS_seqs) <- at.root.genes

# poly A tail length
at.root.polyA_lens <- at.leaf.polyA_lens[at.IDmap[at.root.genes]]
names(at.root.polyA_lens) <- at.root.genes

# remove the genes with missing information
at.root.rm_idx <- which(is.na(at.root.log10.TR) | is.na(at.root.log10.mRNA) |
    is.na(at.root.UTR_seqs) | is.na(at.root.CDS_seqs) | is.na(at.root.polyA_lens))
at.root.data <- at.root.data[-at.root.rm_idx, ]
at.root.genes <- at.root.genes[-at.root.rm_idx]
at.root.log10.TR <- at.root.log10.TR[-at.root.rm_idx]
at.root.log10.mRNA <- at.root.log10.mRNA[-at.root.rm_idx]
at.root.UTR_seqs <- at.root.UTR_seqs[-at.root.rm_idx]
at.root.CDS_seqs <- at.root.CDS_seqs[-at.root.rm_idx]
at.root.polyA_lens <- at.root.polyA_lens[-at.root.rm_idx]

# save the processed data to an .Rdata file
save(at.root.genes, at.root.log10.TR, at.root.log10.mRNA, at.root.UTR_seqs,
    at.root.CDS_seqs, at.root.polyA_lens, file = "processed_data_w_polyA/at.root.Rdata")

################## Shoot ##################

# Shoot translational rates (preprocessing) annotated ORF
# lengths
at.shoot.ORF_lens <- read_xlsx("data/Hsu et al counts_cds_Hsu.xlsx",
    sheet = 3, range = "A3:E21579")
at.shoot.ORF_lens <- as.data.frame(at.shoot.ORF_lens)
at.shoot.ORF_lens <- at.shoot.ORF_lens[, c("gene_id", "Annotated ORF length")]
## make the data frame
at.shoot.data <- merge(at.hsu_data, at.shoot.ORF_lens, by = "gene_id")
for (j in 2:(ncol(at.shoot.data) - 1)) {
    at.shoot.data[, j] <- at.shoot.data[, j]/(at.shoot.data[,
        ncol(at.shoot.data)]/1000)  # RPKM
}
at.shoot.data$RD_mean <- rowMeans(at.shoot.data[, 5:7])
at.shoot.data$RNA_mean <- rowMeans(at.shoot.data[, 11:13])
## thresholded to >= 1 RPKM for RNA density
at.shoot.data <- at.shoot.data[at.shoot.data$RNA_mean >= 1 &
```

```r
    at.shoot.data$RD_mean > 0, ]
at.shoot.data$TE <- at.shoot.data$RD_mean/at.shoot.data$RNA_mean

# remove duplicate gene names based on the leaf tissue
at.shoot.data <- resolve_dup_genes_root_shoot_data(data = at.shoot.data,
    leaf_CDS_seqs = at.leaf.CDS_seqs)

# genes
at.shoot.genes <- at.shoot.data$gene_id

# TR data
at.shoot.log10.TR <- log(at.shoot.data$TE, 10)
names(at.shoot.log10.TR) <- at.shoot.genes

# mRNA data
at.shoot.log10.mRNA <- log(at.shoot.data$RNA_mean, 10)
names(at.shoot.log10.mRNA) <- at.shoot.genes

# UTR sequences
at.shoot.UTR_seqs <- at.UTR_seqs[at.IDmap[at.shoot.genes]]
names(at.shoot.UTR_seqs) <- at.shoot.genes

# CDS sequences
at.shoot.CDS_seqs <- at.CDS_seqs[at.IDmap[at.shoot.genes]]
names(at.shoot.CDS_seqs) <- at.shoot.genes

# poly A tail length
at.shoot.polyA_lens <- at.leaf.polyA_lens[at.IDmap[at.shoot.genes]]
names(at.shoot.polyA_lens) <- at.shoot.genes

# remove the genes with missing information
at.shoot.rm_idx <- which(is.na(at.shoot.log10.TR) | is.na(at.shoot.log10.mRNA) |
    is.na(at.shoot.UTR_seqs) | is.na(at.shoot.CDS_seqs) | is.na(at.shoot.polyA_lens))
at.shoot.data <- at.shoot.data[-at.shoot.rm_idx, ]
at.shoot.genes <- at.shoot.genes[-at.shoot.rm_idx]
at.shoot.log10.TR <- at.shoot.log10.TR[-at.shoot.rm_idx]
at.shoot.log10.mRNA <- at.shoot.log10.mRNA[-at.shoot.rm_idx]
at.shoot.UTR_seqs <- at.shoot.UTR_seqs[-at.shoot.rm_idx]
at.shoot.CDS_seqs <- at.shoot.CDS_seqs[-at.shoot.rm_idx]
at.shoot.polyA_lens <- at.shoot.polyA_lens[-at.shoot.rm_idx]

# save the processed data to an .Rdata file
save(at.shoot.genes, at.shoot.log10.TR, at.shoot.log10.mRNA,
    at.shoot.UTR_seqs, at.shoot.CDS_seqs, at.shoot.polyA_lens,
    file = "processed_data_w_polyA/at.shoot.Rdata")


############################## Mouse ##############################

# CDS sequences
mm.CDS_seqs <- readRDS(file = "data/Mouse Ensembl CDS.rds")

mm.CDS_fa <- scan("data/Mouse Ensembl CDS.txt", what = character(),
```

```
    sep = "\n")
name_idx <- which(startsWith(str = mm.CDS_fa, pattern = ">"))
mm.CDS_seqs_gene_trpt_names <- sapply(strsplit(gsub(pattern = ">",
    replacement = "", x = mm.CDS_fa[name_idx]), "\\|"), FUN = function(x) x)
mm.CDS_seqs_gene_names <- mm.CDS_seqs_gene_trpt_names[1, ]
names(mm.CDS_seqs_gene_names) <- mm.CDS_seqs_gene_trpt_names[2,
    ]
mm.CDS_seqs_alt_names <- mm.CDS_seqs_gene_names[names(mm.CDS_seqs)]

rm(mm.CDS_fa, name_idx, mm.CDS_seqs_gene_trpt_names, mm.CDS_seqs_gene_names)

# 5' UTR sequences
mm.UTR_seqs <- readRDS(file = "data/Mouse Ensembl 5' UTRs.rds")

mm.UTR_fa <- scan("data/Mouse Ensembl 5' UTRs.txt", what = character(),
    sep = "\n")
name_idx <- which(startsWith(str = mm.UTR_fa, pattern = ">"))
mm.UTR_seqs_gene_trpt_names <- sapply(strsplit(gsub(pattern = ">",
    replacement = "", x = mm.UTR_fa[name_idx]), "\\|"), FUN = function(x) x)
mm.UTR_seqs_gene_names <- mm.UTR_seqs_gene_trpt_names[1, ]
names(mm.UTR_seqs_gene_names) <- mm.UTR_seqs_gene_trpt_names[2,
    ]
mm.UTR_seqs_alt_names <- mm.UTR_seqs_gene_names[names(mm.UTR_seqs)]

rm(mm.UTR_fa, name_idx, mm.UTR_seqs_gene_trpt_names, mm.UTR_seqs_gene_names)

################## NIH3T3 ##################

# NIH3T3 translational rates and features (data preprocessing
# in ``030218_mouse_and_arabidopsis_models.Rmd'')
mm.nih3t3.data <- readRDS(file = "data/mouse_data_Subtelny_NIH3T3.rds")
mm.nih3t3.data <- mm.nih3t3.data[complete.cases(mm.nih3t3.data),
    ]

# genes
mm.nih3t3.genes <- mm.nih3t3.data$Ensembl

# TR data
mm.nih3t3.log10.TR <- mm.nih3t3.data$"log10 TE"
names(mm.nih3t3.log10.TR) <- mm.nih3t3.genes

# mRNA data
mm.nih3t3.log10.mRNA <- mm.nih3t3.data$"log10 mRNA"
names(mm.nih3t3.log10.mRNA) <- mm.nih3t3.genes

# UTR sequences
mm.nih3t3.UTR_seqs <- mm.UTR_seqs[mm.nih3t3.genes]

# CDS sequences
mm.nih3t3.CDS_seqs <- mm.CDS_seqs[mm.nih3t3.genes]

# poly A tail length
mm.nih3t3.polyA_lens <- mm.nih3t3.data$"poly-A tail length"
```

```r
names(mm.nih3t3.polyA_lens) <- mm.nih3t3.genes

# save the processed data to an .Rdata file
save(mm.nih3t3.data, mm.nih3t3.genes, mm.nih3t3.log10.TR, mm.nih3t3.log10.mRNA,
    mm.nih3t3.UTR_seqs, mm.nih3t3.CDS_seqs, mm.nih3t3.polyA_lens,
    file = "processed_data_w_polyA/mm.nih3t3.Rdata")

############### Liver & Kidney ###############

# both tissues have short gene names, so we will resolve the
# duplicated short gene names
mm.IDmap <- resolve_dup_genes(long_seqs = mm.UTR_seqs, short_names = mm.UTR_seqs_alt_names,
    tissue_long_seqs = mm.nih3t3.UTR_seqs)

saveRDS(mm.IDmap, file = "processed_data_w_polyA/mm.IDmap.rds")

################## Liver ##################

# translational rates
mm.liver.data <- read_xlsx("data/GSM1644076 liver 1 RPKM RNA.rpkm copy.xlsx",
    sheet = 3, range = "A2:F10097")
mm.liver.data <- as.data.frame(mm.liver.data)

# genes
mm.liver.genes <- mm.liver.data[, 1]

# TR data
mm.liver.log10.TR <- mm.liver.data$"log TE"
names(mm.liver.log10.TR) <- mm.liver.genes

# mRNA data
mm.liver.log10.mRNA <- mm.liver.data$"log RNA"
names(mm.liver.log10.mRNA) <- mm.liver.genes

# UTR sequences
mm.liver.UTR_seqs <- mm.UTR_seqs[mm.IDmap[mm.liver.genes]]
names(mm.liver.UTR_seqs) <- mm.liver.genes

# CDS sequences
mm.liver.CDS_seqs <- mm.CDS_seqs[mm.IDmap[mm.liver.genes]]
names(mm.liver.CDS_seqs) <- mm.liver.genes

# poly A tail length
mm.liver.polyA_lens <- mm.nih3t3.polyA_lens[mm.IDmap[mm.liver.genes]]
names(mm.liver.polyA_lens) <- mm.liver.genes

# remove the genes with missing information
mm.liver.rm_idx <- which(is.na(mm.liver.log10.TR) | is.na(mm.liver.log10.mRNA) |
    is.na(mm.liver.UTR_seqs) | is.na(mm.liver.CDS_seqs) | is.na(mm.liver.polyA_lens))
mm.liver.data <- mm.liver.data[-mm.liver.rm_idx, ]
mm.liver.genes <- mm.liver.genes[-mm.liver.rm_idx]
mm.liver.log10.TR <- mm.liver.log10.TR[-mm.liver.rm_idx]
mm.liver.log10.mRNA <- mm.liver.log10.mRNA[-mm.liver.rm_idx]
```

```
mm.liver.UTR_seqs <- mm.liver.UTR_seqs[-mm.liver.rm_idx]
mm.liver.CDS_seqs <- mm.liver.CDS_seqs[-mm.liver.rm_idx]
mm.liver.polyA_lens <- mm.liver.polyA_lens[-mm.liver.rm_idx]

# save the processed data to an .Rdata file
save(mm.liver.genes, mm.liver.log10.TR, mm.liver.log10.mRNA,
    mm.liver.UTR_seqs, mm.liver.CDS_seqs, mm.liver.polyA_lens,
    file = "processed_data_w_polyA/mm.liver.Rdata")

################## Kidney ##################

# translational rates
mm.kidney.data <- read_xlsx("data/GSM2149102_Kidney 1 RPKM RNA copy 2.xlsx",
    sheet = 3, range = "A2:F11654")
mm.kidney.data <- as.data.frame(mm.kidney.data)

# genes
mm.kidney.genes <- mm.kidney.data[, 1]

# TR data
mm.kidney.log10.TR <- mm.kidney.data$"log TE"
names(mm.kidney.log10.TR) <- mm.kidney.genes

# mRNA data
mm.kidney.log10.mRNA <- mm.kidney.data$"log RNA"
names(mm.kidney.log10.mRNA) <- mm.kidney.genes

# UTR sequences
mm.kidney.UTR_seqs <- mm.UTR_seqs[mm.IDmap[mm.kidney.genes]]
names(mm.kidney.UTR_seqs) <- mm.kidney.genes

# CDS sequences
mm.kidney.CDS_seqs <- mm.CDS_seqs[mm.IDmap[mm.kidney.genes]]
names(mm.kidney.CDS_seqs) <- mm.kidney.genes

# poly A tail length
mm.kidney.polyA_lens <- mm.nih3t3.polyA_lens[mm.IDmap[mm.kidney.genes]]
names(mm.kidney.polyA_lens) <- mm.kidney.genes

# remove the genes with missing information
mm.kidney.rm_idx <- which(is.na(mm.kidney.log10.TR) | is.na(mm.kidney.log10.mRNA) |
    is.na(mm.kidney.UTR_seqs) | is.na(mm.kidney.CDS_seqs) | is.na(mm.kidney.polyA_lens))
mm.kidney.data <- mm.kidney.data[-mm.kidney.rm_idx, ]
mm.kidney.genes <- mm.kidney.genes[-mm.kidney.rm_idx]
mm.kidney.log10.TR <- mm.kidney.log10.TR[-mm.kidney.rm_idx]
mm.kidney.log10.mRNA <- mm.kidney.log10.mRNA[-mm.kidney.rm_idx]
mm.kidney.UTR_seqs <- mm.kidney.UTR_seqs[-mm.kidney.rm_idx]
mm.kidney.CDS_seqs <- mm.kidney.CDS_seqs[-mm.kidney.rm_idx]
mm.kidney.polyA_lens <- mm.kidney.polyA_lens[-mm.kidney.rm_idx]

# save the processed data to an .Rdata file
save(mm.kidney.genes, mm.kidney.log10.TR, mm.kidney.log10.mRNA,
    mm.kidney.UTR_seqs, mm.kidney.CDS_seqs, mm.kidney.polyA_lens,
```

```
    file = "processed_data_w_polyA/mm.kidney.Rdata")


############################## Human ##############################

# Gene ID conversion: RefSeq vs. Ensembl (downloaded from
# ensembl.org BioMart, GRCh38.p12)
hs.RefSeq_Ensembl <- read.table("data/GRCh38_RefSeq_to_Ensembl.txt",
    header = T, sep = "\t", stringsAsFactors = F)
hs.RefSeq_Ensembl <- hs.RefSeq_Ensembl[hs.RefSeq_Ensembl[, 1] !=
    "" & hs.RefSeq_Ensembl[, 2] != "", ]
hs.RefSeq <- hs.RefSeq_Ensembl[, 2]
hs.Ensembl <- hs.RefSeq_Ensembl[, 1]
names(hs.RefSeq) <- hs.Ensembl
names(hs.Ensembl) <- hs.RefSeq
rm(hs.RefSeq_Ensembl)

# CDS sequences (downloaded from ensembl.org BioMart,
# GRCh38.p12)
hs.CDS_fa <- scan("data/GRCh38_Ensembl_CDS.txt", what = character(),
    sep = "\n")
name_idx <- which(startsWith(str = hs.CDS_fa, pattern = ">"))
n_seq <- length(name_idx)
hs.genes <- gsub(pattern = ">", replacement = "", x = hs.CDS_fa[name_idx])
hs.CDS_seqs <- sapply(1:n_seq, FUN = function(i) {
    start_idx <- name_idx[i] + 1
    if (i < n_seq) {
        stop_idx <- name_idx[i + 1] - 1
    } else {
        stop_idx <- length(hs.CDS_fa)
    }
    paste(hs.CDS_fa[start_idx:stop_idx], collapse = "")
})
names(hs.CDS_seqs) <- hs.genes
hs.CDS_seqs <- hs.CDS_seqs[hs.CDS_seqs != "Sequence unavailable"]
rm(hs.CDS_fa)

# UTR sequences (downloaded from ensembl.org BioMart,
# GRCh38.p12)
hs.UTR_fa <- scan("data/GRCh38_Ensembl_5UTR.txt", what = character(),
    sep = "\n")
name_idx <- which(startsWith(str = hs.UTR_fa, pattern = ">"))
n_seq <- length(name_idx)
hs.genes <- gsub(pattern = ">", replacement = "", x = hs.UTR_fa[name_idx])
hs.UTR_seqs <- sapply(1:n_seq, FUN = function(i) {
    start_idx <- name_idx[i] + 1
    if (i < n_seq) {
        stop_idx <- name_idx[i + 1] - 1
    } else {
        stop_idx <- length(hs.UTR_fa)
    }
    paste(hs.UTR_fa[start_idx:stop_idx], collapse = "")
})
```

```r
names(hs.UTR_seqs) <- hs.genes
hs.UTR_seqs <- hs.UTR_seqs[hs.UTR_seqs != "Sequence unavailable"]
rm(hs.UTR_fa)

################## Hela2 (Guo et al.) ##################

# hela2 translational rates
hs.hela2.data <- read_xlsx("data/Guo et al HeLa.xlsx", sheet = 1,
    range = "A9:G10131")
hs.hela2.data <- as.data.frame(hs.hela2.data)

# genes
hs.hela2.genes <- hs.hela2.data$RefSeq_Accession

# TR data
hs.hela2.log10.TR <- hs.hela2.data$"Log10 TE"
names(hs.hela2.log10.TR) <- hs.hela2.genes

# mRNA data
hs.hela2.log10.mRNA <- hs.hela2.data$"Log10 mRNA"
names(hs.hela2.log10.mRNA) <- hs.hela2.genes

# UTR sequences
hs.hela2.UTR_seqs <- hs.UTR_seqs[hs.Ensembl[hs.hela2.genes]]
names(hs.hela2.UTR_seqs) <- hs.hela2.genes

# CDS sequences
hs.hela2.CDS_seqs <- hs.CDS_seqs[hs.Ensembl[hs.hela2.genes]]
names(hs.hela2.CDS_seqs) <- hs.hela2.genes

# poly A tail length borrow the information from another hela
# data
hs.hela.data <- read_xlsx("data/Subtelny GSE52809_HeLa.xlsx",
    sheet = 1, range = "A5:K6923")
hs.hela.data <- as.data.frame(hs.hela.data)
hs.hela.genes <- hs.hela.data$"Transcript ID"
hs.hela.polyA_lens <- as.numeric(hs.hela.data$"Median poly A length")
names(hs.hela.polyA_lens) <- hs.hela.genes
hs.hela2.polyA_lens <- hs.hela.polyA_lens[hs.hela2.genes]

# remove the genes with missing information or containing 'N'
# in sequences
hs.hela2.rm_idx <- which(is.na(hs.hela2.log10.TR) | is.na(hs.hela2.log10.mRNA) |
    is.na(hs.hela2.UTR_seqs) | is.na(hs.hela2.CDS_seqs) | is.na(hs.hela2.polyA_lens) |
    grepl("N", hs.hela2.UTR_seqs) | grepl("N", hs.hela2.CDS_seqs))
hs.hela2.data <- hs.hela2.data[-hs.hela2.rm_idx, ]
hs.hela2.genes <- hs.hela2.genes[-hs.hela2.rm_idx]
hs.hela2.log10.TR <- hs.hela2.log10.TR[-hs.hela2.rm_idx]
hs.hela2.log10.mRNA <- hs.hela2.log10.mRNA[-hs.hela2.rm_idx]
hs.hela2.UTR_seqs <- hs.hela2.UTR_seqs[-hs.hela2.rm_idx]
hs.hela2.CDS_seqs <- hs.hela2.CDS_seqs[-hs.hela2.rm_idx]
hs.hela2.polyA_lens <- hs.hela2.polyA_lens[-hs.hela2.rm_idx]
```

```
# save the processed data to an .Rdata file
save(hs.hela2.data, hs.hela2.genes, hs.hela2.log10.TR, hs.hela2.log10.mRNA,
    hs.hela2.UTR_seqs, hs.hela2.CDS_seqs, hs.hela2.polyA_lens,
    file = "processed_data_w_polyA/hs.hela2.Rdata")
```

### Read in the preprocessed data

```
# a list of species and tissues
species_tissue_list <- list(sc = "", sp = c("", "alt"), at = c("leaf",
    "root", "shoot"), mm = c("nih3t3", "liver", "kidney"), hs = c("hela2"))

# a list of species and exemplar tissues
species_exemplar_tissue_list <- list(sc = "", sp = "alt", at = "leaf",
    mm = "nih3t3", hs = "hela2")

# a vector of species and examplar tissue names
species_exemplar_tissue_names <- sapply(1:length(species_exemplar_tissue_list),
    FUN = function(i) {
        if (species_exemplar_tissue_list[[i]] != "") {
            paste(names(species_exemplar_tissue_list)[i], species_exemplar_tissue_list[[i]],
                sep = ".")
        } else {
            names(species_exemplar_tissue_list)[i]
        }
    })

# a vector of colors for species
species_cols <- c("#EB9D41", "#94241D", "#225392", "#2F6E42",
    "#CEC448")
names(species_cols) <- species_exemplar_tissue_names

for (species in names(species_tissue_list)) {
    for (species_tissue in species_tissue_list[[species]]) {
        if (species_tissue != "") {
            tissue_dot <- paste0(".", species_tissue)
            tissue_us <- paste0("_", species_tissue)
        } else {
            tissue_dot <- ""
            tissue_us <- ""
        }
        load(file = paste0("processed_data_w_polyA/", species,
            tissue_dot, ".Rdata"))
    }
}

## mouse ID map
mm.IDmap <- readRDS(file = "processed_data_w_polyA/mm.IDmap.rds")

## Arabidopsis ID map
at.IDmap <- readRDS(file = "processed_data_w_polyA/at.IDmap.rds")
```

### Generate and select RNA folding energy features

#### Settings

```
# folding energy window lengths
window_lens <- paste(c(6, 8, seq(from=10, to=100, by=5)))

# uTICE length
species_uTICE_len_list <- list(
    sc=35,
    sp=30, # only for calculating the 5'mostTICE feature
    at=65,
    mm=5,
    hs=list("hela2"=6)
)

# dTICE length
species_dTICE_len_list <- list(
    sc=28,
    sp=13,
    at=33,
    mm=13,
    hs=list("hela2"=7)
)

# UTR length breaks to obtain gene groups
species_len_break_list <- list(
    sc=c(20,35),
    sp=NULL, # one-part regression
    at=65,
    mm=NULL,
    hs=NULL # one-part regression
)
```

#### The following code only needs to be run once

```
# UTR folding energy of different window lengths
for (species_short in names(species_tissue_list)) {
    for (species_tissue in species_tissue_list[[species_short]]) {
        if (species_tissue != "") {
            species_genes <- get(paste(species_short, species_tissue,
                "genes", sep = "."))
        } else {
            species_genes <- get(paste(species_short, "genes",
                sep = "."))
        }
        ## read in the whole UTR folding energy feature (one value per
        ## gene)
        if (species_tissue != "") {
            whole_5UTR_folding_energy <- readRDS(file = paste0("processed_data_w_polyA/whole_UTR_folding
                species_short, "_", species_tissue, "_whole_UTR_folding_energy_feature.rds"))
```

```
    } else {
        whole_5UTR_folding_energy <- readRDS(file = paste0("processed_data_w_polyA/whole_UTR_folding
            species_short, "_whole_UTR_folding_energy_features.rds"))
    }

    if (species_short != "hs") {
        uTICE_len <- species_uTICE_len_list[[species_short]]
    } else {
        uTICE_len <- species_uTICE_len_list[[species_short]][[species_tissue]]
    }

    # generate folding energy features
    for (window_len in window_lens) {
        if (species_tissue != "") {
            UTR_folding_energy_all <- readRDS(file = paste0("processed_data_w_polyA/var_win_fold_UTR
              species_short, "_", species_tissue, "_5UTR_CDS_folding_energy_",
              window_len, ".rds"))
        } else {
            UTR_folding_energy_all <- readRDS(file = paste0("processed_data_w_polyA/var_win_fold_UTR
              species_short, "_5UTR_CDS_folding_energy_",
              window_len, ".rds"))
        }
        UTR_folding_energy_all <- UTR_folding_energy_all$UTR_folding_energy_all[species_genes]
        UTR_folding_energy_percentiles <- quantile(unlist(UTR_folding_energy_all),
            probs = c(0.05, 0.1, 0.2, 0.8, 0.9))
        UTR_folding_energy_features <- t(sapply(UTR_folding_energy_all,
            FUN = function(x) {
              if (is.null(x)) {
                return(rep(NA, 18))
              } else {
                x1 <- x[1]
                x2 <- rev(x)[1]
                x3 <- rev(x)[min(uTICE_len, length(x))]
                x4 <- mean(x)
                x5 <- min(x)
                x6 <- max(x)
                tmp <- quantile(x, probs = c(0.9, 0.1, 0.75,
                  0.25))
                n <- length(x)
                x7 <- tmp[1]
                x8 <- tmp[2]
                x9 <- tmp[3]
                x10 <- tmp[4]
                x11 <- sum(x <= UTR_folding_energy_percentiles[3])/n
                x16 <- sum(x > UTR_folding_energy_percentiles[5])/n
                x17 <- sum(x > UTR_folding_energy_percentiles[4])/n
                x13 <- sum(x[x <= UTR_folding_energy_percentiles[1]])
                x14 <- sum(x[x <= UTR_folding_energy_percentiles[2]])
                x15 <- sum(x[x <= UTR_folding_energy_percentiles[3]])
                x18 <- sum(x[x > UTR_folding_energy_percentiles[5]])
                x19 <- sum(x[x > UTR_folding_energy_percentiles[4]])
                return(c(x1, x2, x3, x4, x5, x6, x7, x8,
                  x9, x10, x11, x13, x14, x15, x16, x17,
```

```
                    x18, x19))
                }
            }))
        UTR_folding_energy_features <- cbind(UTR_folding_energy_features,
            whole_5UTR_folding_energy)

        # deal with the missing data
        if (any(is.na(UTR_folding_energy_features[, 1]))) {
            missing_idx <- which(is.na(UTR_folding_energy_features[,
              1]))
            for (i in missing_idx) {
              for (j in 1:(ncol(UTR_folding_energy_features) -
                1)) {
                quantile_ij <- sum(UTR_folding_energy_features_old[,
                  j] <= UTR_folding_energy_features_old[i,
                  j])/nrow(UTR_folding_energy_features)
                UTR_folding_energy_features[i, j] <- quantile(UTR_folding_energy_features[,
                  j], probs = quantile_ij, na.rm = T)
              }
            }
        }

        colnames(UTR_folding_energy_features) <- c("5'most",
            "-1window", "5'mostTICE", "mean", "min", "max",
            "90%", "10%", "75%", "25%", "%<=20%", "sum<=5%",
            "sum<=10%", "sum<=20%", "%>90%", "%>80%", "sum>90%",
            "sum>80%", "whole")

        ## save the folding energy features to an RDS file
        if (species_tissue != "") {
            saveRDS(UTR_folding_energy_features, file = paste0("processed_data_w_polyA/folding_energ
              species_short, "_", species_tissue, "_UTR_folding_energy_features_",
              window_len, ".rds"))
        } else {
            saveRDS(UTR_folding_energy_features, file = paste0("processed_data_w_polyA/folding_energ
              species_short, "_UTR_folding_energy_features_",
              window_len, ".rds"))
        }

        ## store the features for possible use in the next iteration
        UTR_folding_energy_features_old <- UTR_folding_energy_features
      }
    }
}

## specially create a second version of the features for
## 'spom': don't include any folding energy windows that
## extend into 3' of +30
for (species_short in c("sp")) {
    for (species_tissue in species_tissue_list[[species_short]]) {
        if (species_tissue != "") {
            species_genes <- get(paste(species_short, species_tissue,
                "genes", sep = "."))
```

```
    } else {
        species_genes <- get(paste(species_short, "genes",
            sep = "."))
    }
    ## read in the whole UTR folding energy feature (one value per
    ## gene)
    if (species_tissue != "") {
        whole_5UTR_folding_energy <- readRDS(file = paste0("processed_data_w_polyA/whole_UTR_folding
            species_short, "_", species_tissue, "_whole_UTR_folding_energy_feature.rds"))
    } else {
        whole_5UTR_folding_energy <- readRDS(file = paste0("processed_data_w_polyA/whole_UTR_folding
            species_short, "_whole_UTR_folding_energy_features.rds"))
    }

    if (species_short != "hs") {
        uTICE_len <- species_uTICE_len_list[[species_short]]
    } else {
        uTICE_len <- species_uTICE_len_list[[species_short]][[species_tissue]]
    }

    # generate folding energy features
    for (window_len in window_lens) {
        if (species_tissue != "") {
            UTR_folding_energy_all <- readRDS(file = paste0("processed_data_w_polyA/var_win_fold_UTR
              species_short, "_", species_tissue, "_5UTR_CDS_folding_energy_",
              window_len, ".rds"))
        } else {
            UTR_folding_energy_all <- readRDS(file = paste0("processed_data_w_polyA/var_win_fold_UTR
              species_short, "_5UTR_CDS_folding_energy_",
              window_len, ".rds"))
        }
        UTR_folding_energy_all <- UTR_folding_energy_all$UTR_folding_energy_all[species_genes]
        # remove the long windows that extend beyond +30 before
        # calculating the percentiles

        trun_pos <- as.numeric(window_len) - 30 + 1  # the position (-trun_pos) relative to AUG; wi
        UTR_folding_energy_all_trun <- lapply(UTR_folding_energy_all,
            FUN = function(x) {
              if (trun_pos > length(x)) {
                x <- NULL
              } else if (trun_pos > 0) {
                x <- rev(rev(x)[trun_pos:length(x)])
              }
              return(x)
            })
        UTR_folding_energy_percentiles <- quantile(unlist(UTR_folding_energy_all_trun),
            probs = c(0.05, 0.1, 0.2, 0.8, 0.9))

        UTR_folding_energy_features <- t(sapply(1:length(UTR_folding_energy_all_trun),
            FUN = function(i) {
              x <- UTR_folding_energy_all_trun[[i]]
              x_orig <- UTR_folding_energy_all[[i]]
              if (is.null(x)) {
```

```
        return(rep(NA, 18))
      } else {
        x1 <- x[1]
        if (trun_pos > 1) {
          x2 <- NA
        } else {
          x2 <- rev(x_orig)[1]
        }
        if (trun_pos > uTICE_len) {
          x3 <- NA
        } else {
          x3 <- rev(x_orig)[uTICE_len]
        }
        x4 <- mean(x)
        x5 <- min(x)
        x6 <- max(x)
        tmp <- quantile(x, probs = c(0.9, 0.1, 0.75,
          0.25))
        n <- length(x)
        x7 <- tmp[1]
        x8 <- tmp[2]
        x9 <- tmp[3]
        x10 <- tmp[4]
        x11 <- sum(x <= UTR_folding_energy_percentiles[3])/n
        x16 <- sum(x > UTR_folding_energy_percentiles[5])/n
        x17 <- sum(x > UTR_folding_energy_percentiles[4])/n
        x13 <- sum(x[x <= UTR_folding_energy_percentiles[1]])
        x14 <- sum(x[x <= UTR_folding_energy_percentiles[2]])
        x15 <- sum(x[x <= UTR_folding_energy_percentiles[3]])
        x18 <- sum(x[x > UTR_folding_energy_percentiles[5]])
        x19 <- sum(x[x > UTR_folding_energy_percentiles[4]])
        return(c(x1, x2, x3, x4, x5, x6, x7, x8,
          x9, x10, x11, x13, x14, x15, x16, x17,
          x18, x19))
      }
    }))
nonNA_idx <- apply(UTR_folding_energy_features, 2,
    FUN = function(y) !all(is.na(y)))
UTR_folding_energy_features <- cbind(UTR_folding_energy_features[,
    nonNA_idx], whole_5UTR_folding_energy)

# deal with the missing data
if (any(is.na(UTR_folding_energy_features[, 1]))) {
    missing_idx <- which(is.na(UTR_folding_energy_features[,
      1]))
    for (i in missing_idx) {
      for (j in 1:(ncol(UTR_folding_energy_features) -
        1)) {
        quantile_ij <- sum(UTR_folding_energy_features_old[,
          j] <= UTR_folding_energy_features_old[i,
          j])/nrow(UTR_folding_energy_features)
        UTR_folding_energy_features[i, j] <- quantile(UTR_folding_energy_features[,
          j], probs = quantile_ij, na.rm = T)
```

```
                  }
                }
            }

            colnames(UTR_folding_energy_features) <- c(c("5'most",
                "-1window", "5'mostTICE", "mean", "min", "max",
                "90%", "10%", "75%", "25%", "%<=20%", "sum<=5%",
                "sum<=10%", "sum<=20%", "%>90%", "%>80%", "sum>90%",
                "sum>80%")[nonNA_idx], "whole")

            ## save the folding energy features to an RDS file
            if (species_tissue != "") {
                saveRDS(UTR_folding_energy_features, file = paste0("processed_data_w_polyA/folding_energ
                  species_short, "_", species_tissue, "_UTR_folding_energy_features_",
                  window_len, "_trun.rds"))
            } else {
                saveRDS(UTR_folding_energy_features, file = paste0("processed_data_w_polyA/folding_energ
                  species_short, "_UTR_folding_energy_features_",
                  window_len, "_trun.rds"))
            }

            ## store the features for possible use in the next iteration
            UTR_folding_energy_features_old <- UTR_folding_energy_features
        }
    }
}

# select folding energy features for every species function
fold_energy_feature_select <- function(species_prefix, tissue_prefix,
    len_breaks, ifexemplar, selected_feature_names = NULL, iftrun = F) {
    # species_prefix = 'sc', 'mm', 'at', 'sp', 'dm', 'hs'
    # tissue_prefix = '' ('sc'), c('nih3t3', 'liver', 'kidney')
    # ('mm'), c('leaf', 'root', 'shoot') ('at'), c('', 'alt')
    # ('sp'), c('0to1hr', '3to4hr') ('dm'), c('hela', 'hek293t')
    # ('hs') len_breaks = c(20, 35) ('sc'), NULL ('mm'), 65
    # ('at'), ...  ifexemplar = TRUE/FLASE; if TRUE, perform
    # feature selection; otherwise, use selected_feature_names
    # selected_feature_names = a character vector of the selected
    # feature names from the exemplar tissue

    ## combine folding energy features from previous calculation
    ## in
    ## '040418_check_folding_energy_window_length_effect(add_a_feature_to_arab).R'
    fold_energy_features <- NULL
    cors <- NULL
    if (tissue_prefix != "") {
        UTR_seqs <- get(paste(species_prefix, tissue_prefix,
            "UTR_seqs", sep = "."))
        log10.TR <- get(paste(species_prefix, tissue_prefix,
            "log10.TR", sep = "."))
    } else {
        UTR_seqs <- get(paste(species_prefix, "UTR_seqs", sep = "."))
        log10.TR <- get(paste(species_prefix, "log10.TR", sep = "."))
```

```
    }
    log10.UTR_lens <- log(nchar(UTR_seqs), 10)
    n <- length(log10.UTR_lens)

    for (window_len in window_lens) {
        if (!iftrun) {
            if (tissue_prefix != "") {
                tmp_features <- readRDS(paste0("processed_data_w_polyA/folding_energy_features/",
                  species_prefix, "_", tissue_prefix, "_UTR_folding_energy_features_",
                  window_len, ".rds"))
            } else {
                tmp_features <- readRDS(paste0("processed_data_w_polyA/folding_energy_features/",
                  species_prefix, "_UTR_folding_energy_features_",
                  window_len, ".rds"))
            }
        } else {
            if (tissue_prefix != "") {
                tmp_features <- readRDS(paste0("processed_data_w_polyA/folding_energy_features/",
                  species_prefix, "_", tissue_prefix, "_UTR_folding_energy_features_",
                  window_len, "_trun.rds"))
            } else {
                tmp_features <- readRDS(paste0("processed_data_w_polyA/folding_energy_features/",
                  species_prefix, "_UTR_folding_energy_features_",
                  window_len, "_trun.rds"))
            }
        }
        if (window_len != rev(window_lens)[1]) {
            # handle the 'whole' feature, which doesn't need a window
            # length
            tmp_features <- tmp_features[, -ncol(tmp_features)]
            tmp_feature_names <- paste(colnames(tmp_features),
                window_len, sep = "_")
        } else {
            tmp_feature_names <- paste(colnames(tmp_features)[-ncol(tmp_features)],
                window_len, sep = "_")
            tmp_feature_names <- c(tmp_feature_names, colnames(tmp_features)[ncol(tmp_features)])
        }
        tmp_cors <- apply(tmp_features, 2, FUN = function(x) cor(log10.TR,
            x, use = "complete.obs"))
        if (length(len_breaks) > 0) {
            tmp_features_by_length <- NULL
            tmp_cors_by_group <- NULL  # Pearson correlation between log10.TR and every feature within
            num_groups <- length(len_breaks) + 1  # number of gene groups divided by UTR lengths
            for (i in 1:num_groups) {
                if (i == 1) {
                  tmp_features_one_group <- as.numeric(log10.UTR_lens <
                    log(len_breaks[1], 10)) * tmp_features
                  colnames(tmp_features_one_group) <- paste(tmp_feature_names,
                    paste0("<", len_breaks[1]), sep = "_")
                  idx_one_group <- which(log10.UTR_lens < log(len_breaks[1],
                    10))
                } else if (i == num_groups) {
                  tmp_features_one_group <- as.numeric(log10.UTR_lens >=
```

```r
                    log(len_breaks[num_groups - 1], 10)) * cbind(tmp_features,
                    rep(1, n))
                  colnames(tmp_features_one_group) <- paste(c(tmp_feature_names,
                    "Intercept"), paste0(">=", len_breaks[num_groups -
                    1]), sep = "_")
                  idx_one_group <- which(log10.UTR_lens >= log(len_breaks[num_groups -
                    1], 10))
                } else {
                  tmp_features_one_group <- as.numeric(log10.UTR_lens >=
                    log(len_breaks[i - 1], 10) & log10.UTR_lens <
                    log(len_breaks[i], 10)) * cbind(tmp_features,
                    rep(1, n))
                  colnames(tmp_features_one_group) <- paste(c(tmp_feature_names,
                    "Intercept"), paste0(">=", len_breaks[i -
                    1], "&<", len_breaks[i]), sep = "_")
                  idx_one_group <- which(log10.UTR_lens >= log(len_breaks[i -
                    1], 10) & log10.UTR_lens < log(len_breaks[i],
                    10))
                }
                tmp_features_by_length <- cbind(tmp_features_by_length,
                  tmp_features_one_group)
                tmp_cors_one_group <- apply(tmp_features_one_group,
                  2, FUN = function(x) cor(log10.TR[idx_one_group],
                    x[idx_one_group], use = "complete.obs"))
                tmp_cors_by_group <- c(tmp_cors_by_group, tmp_cors_one_group)
            }
            tmp_features <- tmp_features_by_length
            tmp_cors <- tmp_cors_by_group
        } else {
            colnames(tmp_features) <- tmp_feature_names
        }
        fold_energy_features <- cbind(fold_energy_features, tmp_features)
        cors <- c(cors, tmp_cors)
    }

    ## remove duplicated column names
    keep_idx <- !duplicated(colnames(fold_energy_features))
    fold_energy_features <- fold_energy_features[, keep_idx]
    cors <- cors[keep_idx]

    fold_energy_features_all <- as.data.frame(cbind(log10.TR,
        fold_energy_features))
    if (ifexemplar) {
        ## use forward selection + Bayesian Information Criterion
        ## (BIC) to select features
        reg_fwd <- regsubsets(log10.TR ~ ., data = fold_energy_features_all,
            nvmax = 70, method = "forward")
        reg_summary <- summary(reg_fwd)
        idx_bic <- which.min(reg_summary$bic)
        selected_feature_idx <- which(reg_summary$outmat[idx_bic,
            ] == "*")
    } else {
        ## use 'selected_feature_names'
```

```
        selected_feature_idx <- which(colnames(fold_energy_features) %in%
            selected_feature_names)
    }

    selected_feature_coef <- summary(lm(log10.TR ~ ., data = fold_energy_features_all[,
        c(1, selected_feature_idx + 1)]))$coefficients[-1, ]
    selected_feature_coef <- cbind(selected_feature_coef, cors[selected_feature_idx])
    colnames(selected_feature_coef)[5] <- "Pearson Cor vs TR"
    ## order the selected features by the p-values of their
    ## coefficients in a multiple linear model
    selected_feature_summary <- selected_feature_coef[order(selected_feature_coef[,
        4]), ]

    ## selected features
    fold_energy_features_selected <- fold_energy_features_all[,
        c(1, selected_feature_idx + 1)]
    ## R2 of the selected features
    selected_feature_summary_R2 <- summary(lm(log10.TR ~ ., data = fold_energy_features_selected))$r.squ[illegible]

    return(list(selected_features = fold_energy_features_selected,
        selected_feature_summary = selected_feature_summary,
        selected_feature_R2 = selected_feature_summary_R2, all_features = fold_energy_features_all))
}

for (species_prefix in names(species_tissue_list)) {
    len_breaks <- species_len_break_list[[species_prefix]]
    for (i in 1:length(species_tissue_list[[species_prefix]])) {
        tissue_prefix <- species_tissue_list[[species_prefix]][i]
        fold_energy_features_selected <- fold_energy_feature_select(species_prefix,
            tissue_prefix, len_breaks, ifexemplar = T)
        if (tissue_prefix != "") {
            saveRDS(fold_energy_features_selected, file = paste0("processed_data_w_polyA/folding_energy[illegible]
                species_prefix, "_", tissue_prefix, "_UTR_folding_energy_features_selected.rds"))
        } else {
            saveRDS(fold_energy_features_selected, file = paste0("processed_data_w_polyA/folding_energy[illegible]
                species_prefix, "_UTR_folding_energy_features_selected.rds"))
        }
        # deal with the truncated features of sp
        if (species_prefix == "sp") {
            fold_energy_features_selected <- fold_energy_feature_select(species_prefix,
                tissue_prefix, len_breaks, ifexemplar = T, iftrun = T)
            if (tissue_prefix != "") {
                saveRDS(fold_energy_features_selected, file = paste0("processed_data_w_polyA/folding_ene[illegible]
                  species_prefix, "_", tissue_prefix, "_UTR_folding_energy_features_selected_trun.rds")[illegible]
            } else {
                saveRDS(fold_energy_features_selected, file = paste0("processed_data_w_polyA/folding_ene[illegible]
                  species_prefix, "_UTR_folding_energy_features_selected_trun.rds"))
            }
        }
    }
}
```

#### Construct features for the full model

##### Settings

```
##################### species-independent information ##################### all
##################### the 4 nucleotides
nts <- c("A", "T", "C", "G")
# all the 16 di-nucleotides
di_nts <- as.character(sapply(nts, FUN = function(nt1) sapply(nts,
    FUN = function(nt2) paste0(nt1, nt2))))
# all the 64 tri-nucleotides
tri_nts <- as.character(sapply(nts, FUN = function(nt1) sapply(nts,
    FUN = function(nt2) sapply(nts, FUN = function(nt3) paste0(nt1,
        nt2, nt3)))))
## translating the 64 tri nucleotides to amino acides
tri_nts_to_aas <- sapply(tri_nts, FUN = function(nuc) {
    as.character(translate(DNAString(nuc)))
})
## 20 amino acids
aas <- unique(tri_nts_to_aas)
aas <- aas[aas != "*"]
## nonstopping 61 codon frequencies (trinucleotide in the open
## reading frame)
nonstop_codons <- tri_nts[tri_nts_to_aas != "*"]

# a function to construct the PWM based on a set of 5' UTR
# sequences (right aligned at the TSS)
UTR_pwm <- function(UTR_seqs, n_left) {
    # n_left: -n_left:-1 is the region to construct the PWM
    tmp_seq <- t(sapply(UTR_seqs, FUN = function(x) {
        tmp <- strsplit(x, "")[[1]]
        rev(tmp)[1:n_left]
    }))
    count_A <- colSums(tmp_seq == "A", na.rm = T)
    count_T <- colSums(tmp_seq == "T", na.rm = T)
    count_C <- colSums(tmp_seq == "C", na.rm = T)
    count_G <- colSums(tmp_seq == "G", na.rm = T)
    count_tot <- count_A + count_T + count_C + count_G
    freq <- cbind(count_A/count_tot, count_T/count_tot, count_C/count_tot,
        count_G/count_tot)
    colnames(freq) <- c("A", "T", "C", "G")
    rownames(freq) <- (-1):(-n_left)
    return(freq[n_left:1, ])
}

# a function to calculate 5' UTR PWM scores for a set of
# genes (aligned at TSS)
UTR_pwm_score <- function(PWM, UTR_seqs) {
    len <- nrow(PWM)
    PWM <- PWM[len:1, ]  # reverse the order, so -1 is the first row
    sapply(UTR_seqs, FUN = function(x) {
        tmp <- rev(strsplit(x, "")[[1]])  # reverse the sequence, so -1 is the first position
        min_len <- min(c(len, length(tmp)))
```

```
        log(prod(sapply(1:min_len, FUN = function(i) {
            PWM[i, tmp[i]]
        })), 10)
    })
}

# a function to construct the PWM based on a set of CDS
# sequences truncated in the +4:+n_right region
CDS_pwm <- function(CDS_seqs_n_right, n_right) {
    tmp_seq <- t(sapply(CDS_seqs_n_right, FUN = function(x) {
        tmp <- strsplit(x, "")[[1]]
    }))
    freq <- t(sapply(1:ncol(tmp_seq), FUN = function(i) {
        tmp_count <- c(sum(tmp_seq[, i] == "A"), sum(tmp_seq[,
            i] == "T"), sum(tmp_seq[, i] == "C"), sum(tmp_seq[,
            i] == "G"))
        return(tmp_count/sum(tmp_count))
    }))
    colnames(freq) <- c("A", "T", "C", "G")
    rownames(freq) <- 4:n_right
    return(freq)
}

# a function to calculate CDS PWM scores for a set of genes
# truncated in the +4:+n_right region
CDS_pwm_score <- function(PWM_n_right, CDS_seqs_n_right) {
    UTR_pwm_score(PWM_n_right, CDS_seqs_n_right)
}

# a function to construct the PWM based on a set of 5' UTR
# sequences and ORF sequences of the same genes (aligned at
# TSS)
UTR_CDS_pwm <- function(UTR_seqs, CDS_seqs, n_UTR, n_CDS) {
    tmp_seq <- t(sapply(UTR_seqs, FUN = function(x) {
        tmp <- strsplit(x, "")[[1]]
        rev(tmp)[1:n_UTR]
    }))
    count_A <- colSums(tmp_seq == "A", na.rm = T)
    count_T <- colSums(tmp_seq == "T", na.rm = T)
    count_C <- colSums(tmp_seq == "C", na.rm = T)
    count_G <- colSums(tmp_seq == "G", na.rm = T)
    count_tot <- count_A + count_T + count_C + count_G
    freq_UTR <- cbind(count_A/count_tot, count_T/count_tot, count_C/count_tot,
        count_G/count_tot)

    tmp_seq <- t(sapply(CDS_seqs, FUN = function(x) {
        tmp <- strsplit(x, "")[[1]]
        rev(tmp[4:n_CDS])  # do not include +1 to +3 (often ATG)
    }))
    if (nrow(tmp_seq) == 1)
        tmp_seq <- t(tmp_seq)
    count_A <- colSums(tmp_seq == "A", na.rm = T)
    count_T <- colSums(tmp_seq == "T", na.rm = T)
```

```
    count_C <- colSums(tmp_seq == "C", na.rm = T)
    count_G <- colSums(tmp_seq == "G", na.rm = T)
    count_tot <- count_A + count_T + count_C + count_G
    freq_CDS <- cbind(count_A/count_tot, count_T/count_tot, count_C/count_tot,
        count_G/count_tot)

    freq <- rbind(freq_CDS, freq_UTR)

    colnames(freq) <- c("A", "U", "C", "G")
    rownames(freq) <- c((n_CDS:4), ((-1):(-n_UTR)))  # do not include +1 to +3 (often ATG)
    return(freq[nrow(freq):1, c("A", "C", "G", "U")])
}

# a function to construct the 5ofTICE PWM based on a set of
# 5' UTR sequences in the region of length n_5ofTICE (left
# aligned at the 5' end)
UTR_pwm_5_aligned <- function(UTR_seqs, n_5ofTICE) {
    tmp_seq <- t(sapply(UTR_seqs, FUN = function(x) {
        tmp <- strsplit(x, "")[[1]]
        tmp[1:n_5ofTICE]
    }))
    if (n_5ofTICE == 1) {
        tmp_seq <- t(tmp_seq)
    }
    count_A <- colSums(tmp_seq == "A", na.rm = T)
    count_T <- colSums(tmp_seq == "T", na.rm = T)
    count_C <- colSums(tmp_seq == "C", na.rm = T)
    count_G <- colSums(tmp_seq == "G", na.rm = T)
    count_tot <- count_A + count_T + count_C + count_G
    freq <- cbind(count_A/count_tot, count_T/count_tot, count_C/count_tot,
        count_G/count_tot)
    colnames(freq) <- c("A", "T", "C", "G")
    rownames(freq) <- 1:n_5ofTICE
    return(freq)
}

# a function to calculate 5ofTICE PWM scores for a set of
# genes (left aligned at the 5' end)
UTR_pwm_5_aligned_score <- function(PWM, UTR_seqs) {
    # 5' end is the first row
    len <- nrow(PWM)
    sapply(UTR_seqs, FUN = function(x) {
        tmp <- strsplit(x, "")[[1]]
        min_len <- min(c(len, length(tmp)))
        log(prod(sapply(1:min_len, FUN = function(i) {
            PWM[i, tmp[i]]
        })), 10)
    })
}

##################### parameters for each species ##################### S.
##################### cerevisae
sc.n_left <- 35  # the minimum UTR length of 'long' genes
```

```
sc.n_right <- 28  # the CDS PWM length
sc.TR_top_pctl <- 0.9  # the percentile of TR top genes
sc.TR_bottom_pctl <- 0.1  # the percentile of TR top genes
sc.PWM_lens <- c(5, 20, 35)  # lengths of 5' UTR PWMs
sc.len_breaks <- c(20, 35)  # UTR length breaks to divide genes into groups
sc.n_5ofTICE <- 5  # the 5ofTICE PWM length (from the 5' end)

# S. pombe
sp.n_left <- 5  # the minimum UTR length of 'long' genes
sp.n_right <- 13  # the CDS PWM length
sp.TR_top_pctl <- 0.9  # the percentile of TR top genes
sp.TR_bottom_pctl <- 0.1  # the percentile of TR top genes
sp.PWM_lens <- 5  # lengths of 5' UTR PWMs
sp.len_breaks <- NULL  # UTR length breaks to divide genes into groups
sp.n_5ofTICE <- 5  # the 5ofTICE PWM length (from the 5' end)

# Arabidopsis
at.n_left <- 65  # the minimum UTR length of 'long' genes
at.n_right <- 33  # the CDS PWM length
at.TR_top_pctl <- 0.9  # the percentile of TR top genes
at.TR_bottom_pctl <- 0.1  # the percentile of TR top genes
at.PWM_lens <- c(5, 65)  # lengths of 5' UTR PWMs
at.len_breaks <- c(65)  # UTR length breaks to divide genes into groups
at.n_5ofTICE <- 5  # the 5ofTICE PWM length (from the 5' end)

# Mouse
mm.n_left <- 6  # the minimum UTR length of 'long' genes
mm.n_right <- 13  # the CDS PWM length
mm.TR_top_pctl <- 0.9  # the percentile of TR top genes
mm.TR_bottom_pctl <- 0.1  # the percentile of TR top genes
mm.PWM_lens <- c(6)  # lengths of 5' UTR PWMs
mm.len_breaks <- NULL  # UTR length breaks to divide genes into groups
mm.n_5ofTICE <- 5  # the 5ofTICE PWM length (from the 5' end)

# Human hela 2
hs.hela2.n_left <- 6  # the minimum UTR length of 'long' genes
hs.hela2.n_right <- 7  # the CDS PWM length
hs.hela2.TR_top_pctl <- 0.9  # the percentile of TR top genes
hs.hela2.TR_bottom_pctl <- 0.1  # the percentile of TR top genes
hs.hela2.PWM_lens <- 6  # lengths of 5' UTR PWMs
hs.hela2.len_breaks <- NULL  # UTR length breaks to divide genes into groups
hs.hela2.n_5ofTICE <- 5  # the 5ofTICE PWM length (from the 5' end)
```

##### Read in the already generated features & find the TR top and bottom genes

```
for (species in names(species_tissue_list)) {
    for (species_tissue in species_tissue_list[[species]]) {
        if (species_tissue != "") {
            tissue_dot <- paste0(".", species_tissue)
            tissue_us <- paste0("_", species_tissue)
        } else {
            tissue_dot <- ""
```

```
        tissue_us <- ""
    }
    ## 5' UTR log10 lengths (a feature)
    assign(x = paste0(species, tissue_dot, ".log10.UTR_lens"),
        value = log(nchar(get(paste0(species, tissue_dot,
            ".UTR_seqs"))[get(paste0(species, tissue_dot,
            ".genes"))]), 10))
    ## CDS log10 lengths (a feature)
    assign(x = paste0(species, tissue_dot, ".log10.CDS_lens"),
        value = log(nchar(get(paste0(species, tissue_dot,
            ".CDS_seqs"))[get(paste0(species, tissue_dot,
            ".genes"))]), 10))
    ## uAUG counts (a feature)
    assign(x = paste0(species, tissue_dot, ".uAUG_counts"),
        value = stri_count_regex(get(paste0(species, tissue_dot,
            ".UTR_seqs")), "(?=ATG)"))
    ## selected folding energy features (a mega-feature)
    if (species != "sp") {
        assign(x = paste0(species, tissue_dot, ".fold_energy_features_selected"),
            value = readRDS(file = paste0("processed_data_w_polyA/folding_energy_features_selected/
              species, tissue_us, "_UTR_folding_energy_features_selected.rds"))[[1]][,
              -1])  ## the first column is log10.TR
    } else {
        assign(x = paste0(species, tissue_dot, ".fold_energy_features_selected"),
            value = readRDS(file = paste0("processed_data_w_polyA/folding_energy_features_selected/
              species, tissue_us, "_UTR_folding_energy_features_selected_trun.rds"))[[1]][,
              -1])  ## the first column is log10.TR
    }
    ## all folding energy features (multiple individual features)
    assign(x = paste0(species, tissue_dot, ".fold_energy_features_all"),
        value = readRDS(file = paste0("processed_data_w_polyA/folding_energy_features_selected/",
            species, tissue_us, "_UTR_folding_energy_features_selected.rds"))$all_features[,
            -1])  ## the first column is log10.TR
    ## whole CDS folding energy (a feature)
    assign(x = paste0(species, tissue_dot, ".CDS_fold_energy"),
        value = readRDS(file = paste0("processed_data_w_polyA/whole_CDS_folding_energy_feature/",
            species, tissue_us, "_whole_CDS_folding_energy_feature.rds")))
    ## TR top and bottom genes
    if (species != "hs") {
        TR_quantiles <- quantile(get(paste0(species, tissue_dot,
            ".log10.TR")), prob = c(get(paste0(species, ".TR_bottom_pctl")),
            get(paste0(species, ".TR_top_pctl"))))
    } else {
        TR_quantiles <- quantile(get(paste0(species, tissue_dot,
            ".log10.TR")), prob = c(get(paste0(species, tissue_dot,
            ".TR_bottom_pctl")), get(paste0(species, tissue_dot,
            ".TR_top_pctl"))))
    }
    assign(x = paste0(species, tissue_dot, ".TR_bottom_genes"),
        value = get(paste0(species, tissue_dot, ".genes"))[get(paste0(species,
            tissue_dot, ".log10.TR")) <= TR_quantiles[1]])
    assign(x = paste0(species, tissue_dot, ".TR_top_genes"),
        value = get(paste0(species, tissue_dot, ".genes"))[get(paste0(species,
```

```
                tissue_dot, ".log10.TR")) >= TR_quantiles[2]])
    }
}
```

##### The following code only needs to be run once

```
##### uTICE features (a mega-feature)
make_uTICE_features <- function(UTR_seqs, log10.TR, TR_top_pctl,
    n_left, PWM_lens, len_breaks, ifexemplar, selected_feature_names = NULL) {
    # use forward + BIC to select features among the following
    # features (1) 16 dinucleotide frequencies in the -n_left to
    # -1 region (2) 64 trinucleotide frequencies in the -n_left
    # to -1 region (3) PWM scores based on the PWM constructed
    # from the top 10% TR genes' sequences

    # UTR_seqs: a character vector of UTR sequences, with its
    # names as gene names log10.TR: a numeric vector of log10.TR
    # values, with its names as gene names TR_top_pctl: the
    # percentile to define TR top genes, e.g., 0.9 n_left: the
    # minimum UTR length of 'long' genes PWM_lens: a numeric
    # vector containing the possible PWM lengths len_breaks: a
    # numeric vector containing the UTR length breaks, which
    # divide genes into groups ifexemplar = TRUE/FLASE; if TRUE,
    # perform feature selection; otherwise, use
    # selected_feature_names selected_feature_names = a character
    # vector of the selected feature names from the exemplar
    # tissue

    n <- length(UTR_seqs)
    UTR_lens <- nchar(UTR_seqs)
    TR_top_UTR_seqs <- UTR_seqs[log10.TR >= quantile(log10.TR,
        prob = TR_top_pctl)]

    # cut each gene's 5' UTR sequence to the -n_left:-1 region
    UTR_seqs_n_left <- substr(x = UTR_seqs, start = nchar(UTR_seqs) -
        n_left + 1, stop = nchar(UTR_seqs))
    TR_top_UTR_seqs_n_left <- substr(x = TR_top_UTR_seqs, start = nchar(TR_top_UTR_seqs) -
        n_left + 1, stop = nchar(TR_top_UTR_seqs))

    # count the dinucleotide frequencies in the -n_left:-1 region
    # of each gene
    UTR_di_counts_n_left <- sapply(di_nts, FUN = function(nuc) {
        stri_count_regex(UTR_seqs_n_left, paste0("(?=", nuc,
            ")"))
    })
    rownames(UTR_di_counts_n_left) <- names(UTR_seqs_n_left)
    UTR_di_freqs_n_left <- UTR_di_counts_n_left/rowSums(UTR_di_counts_n_left)
    UTR_di_freqs_n_left[is.na(UTR_di_freqs_n_left)] <- 0

    # count the trinucleotide frequencies in the -n_left:-1
    # region of each gene
    UTR_tri_counts_n_left <- sapply(tri_nts, FUN = function(nuc) {
```

```
    stri_count_regex(UTR_seqs_n_left, paste0("(?=", nuc,
        ")"))
})
rownames(UTR_tri_counts_n_left) <- names(UTR_seqs_n_left)
UTR_tri_freqs_n_left <- UTR_tri_counts_n_left/rowSums(UTR_tri_counts_n_left)
UTR_tri_freqs_n_left[is.na(UTR_tri_freqs_n_left)] <- 0

# calculate PWM scores for multiple PWM lengths for each gene
PWM_UTR_scores <- sapply(PWM_lens, FUN = function(len) {
    TR_top_PWM_UTR <- UTR_pwm(TR_top_UTR_seqs_n_left, len)
    UTR_pwm_score(TR_top_PWM_UTR, UTR_seqs_n_left)
})

# combine the features into a design matrix
features <- cbind(UTR_di_freqs_n_left, UTR_tri_freqs_n_left)
feature_names <- colnames(features)
if (length(len_breaks) > 0) {
    features_by_group <- NULL
    cors_by_group <- NULL  # Pearson correlation between log10.TR and every feature within a group
    num_groups <- length(len_breaks) + 1  # number of gene groups divided by UTR lengths
    for (i in 1:num_groups) {
        if (i == 1) {
            features_one_group <- as.numeric(UTR_lens < len_breaks[1]) *
              cbind(features, PWM_UTR_scores[, 1])
            colnames(features_one_group) <- c(paste(feature_names,
              paste0("<", len_breaks[1]), sep = "_"), paste0("PWM",
              PWM_lens[1]))
            idx_one_group <- which(UTR_lens < len_breaks[1])
        } else if (i == num_groups) {
            features_one_group <- as.numeric(UTR_lens >=
              len_breaks[num_groups - 1]) * cbind(features,
              PWM_UTR_scores[, num_groups], rep(1, n))
            colnames(features_one_group) <- c(paste(feature_names,
              paste0(">=", len_breaks[num_groups - 1]), sep = "_"),
              paste0("PWM", PWM_lens[num_groups]), paste0("Intercept_>=",
                len_breaks[num_groups - 1]))
            idx_one_group <- which(UTR_lens >= len_breaks[num_groups -
              1])
        } else {
            features_one_group <- as.numeric(UTR_lens >=
              len_breaks[i - 1] & UTR_lens < len_breaks[i]) *
              cbind(features, PWM_UTR_scores[, i], rep(1,
                n))
            colnames(features_one_group) <- c(paste(feature_names,
              paste0(">=", len_breaks[i - 1], "&<", len_breaks[i]),
              sep = "_"), paste0("PWM", PWM_lens[i]), paste0("Intercept_>=",
              len_breaks[i - 1], "&<", len_breaks[i]))
            idx_one_group <- which(UTR_lens >= len_breaks[i -
              1] & UTR_lens < len_breaks[i])
        }
        features_by_group <- cbind(features_by_group, features_one_group)
        cors_one_group <- apply(features_one_group, 2, FUN = function(x) cor(log10.TR[idx_one_group]
            x[idx_one_group], use = "complete.obs"))
```

```
        cors_by_group <- c(cors_by_group, cors_one_group)
    }
    features <- features_by_group
    cors <- cors_by_group
} else {
    features <- cbind(features, PWM_UTR_scores)
    colnames(features) <- c(feature_names, paste0("PWM",
        PWM_lens))
    cors <- apply(features, 2, FUN = function(x) cor(log10.TR,
        x, use = "complete.obs"))
}

# combine the response and the design matrix into a data
# frame
PWM_idx <- which(grepl("PWM", colnames(features)))
data_frame <- cbind.data.frame(log10.TR = log10.TR, features[,
    PWM_idx], features[, -PWM_idx])
colnames(data_frame) <- c("log10.TR", colnames(features)[PWM_idx],
    colnames(features)[-PWM_idx])
cors <- c(cors[PWM_idx], cors[-PWM_idx])

# R2 with the PWM scores and intercepts only
R2_PWM <- summary(lm(log10.TR ~ features[, which(grepl("PWM",
    colnames(features)) | grepl("Intercept", colnames(features)))]))$r.squared

if (ifexemplar) {
    # use forward selection + Bayesian Information Criterion
    # (BIC) to select features
    regfit.fwd <- regsubsets(log10.TR ~ ., data = data_frame,
        nvmax = 70, force.in = 1:length(PWM_idx), method = "forward")

    reg.summary <- summary(regfit.fwd)
    idx_bic <- which.min(reg.summary$bic)
    features_selected_idx <- which(reg.summary$outmat[idx_bic,
        ] == "*")
} else {
    features_selected_idx <- which(colnames(data_frame) %in%
        selected_feature_names) - 1
}

lm_selected <- summary(lm(log10.TR ~ ., data = data_frame[,
    c(1, features_selected_idx + 1)]))
coefs_selected <- lm_selected$coefficients[-1, ]
R2_selected <- lm_selected$r.squared
coefs_selected <- cbind(coefs_selected, cors[features_selected_idx])
colnames(coefs_selected)[5] <- "Pearson Cor vs TR"
## order the selected features by the p-values of their
## coefficients in a multiple linear model
coefs_selected <- coefs_selected[order(coefs_selected[, 4]),
    ]

uTICE_features <- data_frame[, features_selected_idx + 1]
return(list(uTICE_features = uTICE_features, uTICE_feature_coefs = coefs_selected,
```

```
        R2 = R2_selected, R2_PWM = R2_PWM))
}

# generate uTICE features
for (species in names(species_tissue_list)) {
    for (i in 1:length(species_tissue_list[[species]])) {
        species_tissue <- species_tissue_list[[species]][i]
        if (species_tissue != "") {
            tissue_dot <- paste0(".", species_tissue)
            tissue_us <- paste0("_", species_tissue)
        } else {
            tissue_dot <- ""
            tissue_us <- ""
        }
        if (species != "hs") {
            assign(x = paste0(species, tissue_dot, ".uTICE_features"),
                value = make_uTICE_features(get(paste0(species,
                  tissue_dot, ".UTR_seqs")), get(paste0(species,
                  tissue_dot, ".log10.TR")), get(paste0(species,
                  ".TR_top_pctl")), get(paste0(species, ".n_left")),
                  get(paste0(species, ".PWM_lens")), get(paste0(species,
                    ".len_breaks")), ifexemplar = T))
        } else {
            assign(x = paste0(species, tissue_dot, ".uTICE_features"),
                value = make_uTICE_features(get(paste0(species,
                  tissue_dot, ".UTR_seqs")), get(paste0(species,
                  tissue_dot, ".log10.TR")), get(paste0(species,
                  tissue_dot, ".TR_top_pctl")), get(paste0(species,
                  tissue_dot, ".n_left")), get(paste0(species,
                  tissue_dot, ".PWM_lens")), get(paste0(species,
                  tissue_dot, ".len_breaks")), ifexemplar = T))
        }
        saveRDS(get(paste0(species, tissue_dot, ".uTICE_features")),
            file = paste0("processed_data_w_polyA/uTICE_features/",
                species, tissue_dot, ".uTICE_features.rds"))
    }
}

##### dTICE features (a mega-feature)
make_dTICE_features <- function(CDS_seqs, UTR_seqs, log10.TR,
    TR_top_pctl, n_right, len_breaks, ifexemplar, selected_feature_names = NULL) {
    # use forward + BIC to select features among the following
    # features (1) 16 dinucleotide frequencies in the +4 to
    # +n_right region (2) 64 trinucleotide frequencies in the +4
    # to +n_right region (3) PWM scores based on the PWM
    # constructed from the top 10% TR genes' sequences

    # CDS_seqs: a character vector of CDS sequences, with its
    # names as gene names UTR_seqs: a character vector of UTR
    # sequences, with its names as gene names log10.TR: a numeric
    # vector of log10.TR values, with its names as gene names
    # TR_top_pctl: the percentile to define TR top genes, e.g.,
    # 0.9 n_right: the CDS PWM length len_breaks: a numeric
```

```
# vector containing the UTR length breaks, which divide genes
# into groups ifexemplar = TRUE/FLASE; if TRUE, perform
# feature selection; otherwise, use selected_feature_names
# selected_feature_names = a character vector of the selected
# feature names from the exemplar tissue

n <- length(UTR_seqs)
UTR_lens <- nchar(UTR_seqs)
TR_top_CDS_seqs <- CDS_seqs[log10.TR >= quantile(log10.TR,
    prob = TR_top_pctl)]

# cut each gene's CDS sequence to the +4:+n_right region
CDS_seqs_n_right <- substr(x = CDS_seqs, start = 4, stop = n_right)
TR_top_CDS_seqs_n_right <- substr(x = TR_top_CDS_seqs, start = 4,
    stop = n_right)

# count the dinucleotide frequencies in the +4:+n_right
# region of each gene
CDS_di_counts_n_right <- sapply(di_nts, FUN = function(nuc) {
    stri_count_regex(CDS_seqs_n_right, paste0("(?=", nuc,
        ")"))
})
rownames(CDS_di_counts_n_right) <- names(CDS_seqs_n_right)
CDS_di_freqs_n_right <- CDS_di_counts_n_right/rowSums(CDS_di_counts_n_right)
CDS_di_freqs_n_right[is.na(CDS_di_freqs_n_right)] <- 0

# count the trinucleotide frequencies in the +4:+n_right
# region of each gene
CDS_tri_counts_n_right <- sapply(tri_nts, FUN = function(nuc) {
    stri_count_regex(CDS_seqs_n_right, paste0("(?=", nuc,
        ")"))
})
rownames(CDS_tri_counts_n_right) <- names(CDS_seqs_n_right)
CDS_tri_freqs_n_right <- CDS_tri_counts_n_right/rowSums(CDS_tri_counts_n_right)
CDS_tri_freqs_n_right[is.na(CDS_tri_freqs_n_right)] <- 0

# calculate PWM scores based on the TR top PWM in the
# +4:+n_right region for each gene
PWM_CDS_scores <- sapply(n_right, FUN = function(len) {
    TR_top_PWM_CDS <- CDS_pwm(TR_top_CDS_seqs_n_right, n_right)
    CDS_pwm_score(TR_top_PWM_CDS, CDS_seqs_n_right)
})
colnames(PWM_CDS_scores) <- paste0("PWM", n_right)

# combine the features into a design matrix
features <- cbind(CDS_di_freqs_n_right, CDS_tri_freqs_n_right,
    PWM_CDS_scores)
feature_names <- colnames(features)
cors <- apply(features, 2, FUN = function(x) cor(log10.TR,
    x, "complete.obs"))
if (length(len_breaks) > 0) {
    features_by_group <- NULL
    cors_by_group <- NULL  # Pearson correlation between log10.TR and every feature within a group
```

```
    num_groups <- length(len_breaks) + 1  # number of gene groups divided by UTR lengths
    for (i in 1:num_groups) {
        if (i == 1) {
            features_one_group <- as.numeric(UTR_lens < len_breaks[1]) *
              features
            colnames(features_one_group) <- paste(feature_names,
              paste0("<", len_breaks[1]), sep = "_")
            idx_one_group <- which(UTR_lens < len_breaks[1])
        } else if (i == num_groups) {
            features_one_group <- as.numeric(UTR_lens >=
              len_breaks[num_groups - 1]) * cbind(features,
              rep(1, n))
            colnames(features_one_group) <- paste(c(feature_names,
              "Intercept"), paste0(">=", len_breaks[num_groups -
              1]), sep = "_")
            idx_one_group <- which(UTR_lens >= len_breaks[num_groups -
              1])
        } else {
            features_one_group <- as.numeric(UTR_lens >=
              len_breaks[i - 1] & UTR_lens < len_breaks[i]) *
              cbind(features, rep(1, n))
            colnames(features_one_group) <- paste(c(feature_names,
              "Intercept"), paste0(">=", len_breaks[i - 1],
              "&<", len_breaks[i]), sep = "_")
            idx_one_group <- which(UTR_lens >= len_breaks[i -
              1] & UTR_lens < len_breaks[i])
        }
        features_by_group <- cbind(features_by_group, features_one_group)
        cors_one_group <- apply(features_one_group, 2, FUN = function(x) cor(log10.TR[idx_one_group
            x[idx_one_group], use = "complete.obs"))
        cors_by_group <- c(cors_by_group, cors_one_group)
    }
    features <- features_by_group
    cors <- cors_by_group
}

# combine the response and the design matrix into a data
# frame
PWM_idx <- which(grepl("PWM", colnames(features)))
data_frame <- cbind.data.frame(log10.TR = log10.TR, features[,
    PWM_idx], features[, -PWM_idx])
colnames(data_frame) <- c("log10.TR", colnames(features)[PWM_idx],
    colnames(features)[-PWM_idx])
cors <- c(cors[PWM_idx], cors[-PWM_idx])

# R2 with the PWM scores and intercepts only
R2_PWM <- summary(lm(log10.TR ~ features[, which(grepl("PWM",
    colnames(features)) | grepl("Intercept", colnames(features)))]))$r.squared

if (ifexemplar) {
    # use forward selection + Bayesian Information Criterion
    # (BIC) to select features
    failure <- try(regfit.fwd <- regsubsets(log10.TR ~ .,
```

```
            data = data_frame, nvmax = 70, force.in = 1:length(PWM_idx),
            method = "forward"))

        if (class(failure) == "try-error") {
            regfit.fwd <- regsubsets(log10.TR ~ ., data = data_frame[,
                -(length(PWM_idx) + 1)], nvmax = 70, force.in = 1:length(PWM_idx),
                method = "forward")
        }

        reg.summary <- summary(regfit.fwd)
        idx_bic <- which.min(reg.summary$bic)
        features_selected_idx <- which(reg.summary$outmat[idx_bic,
            ] == "*")
    } else {
        features_selected_idx <- which(colnames(data_frame) %in%
            selected_feature_names) - 1
    }

    lm_selected <- summary(lm(log10.TR ~ ., data = data_frame[,
        c(1, features_selected_idx + 1)]))
    coefs_selected <- lm_selected$coefficients[-1, ]
    R2_selected <- lm_selected$r.squared
    coefs_selected <- cbind(coefs_selected, cors[features_selected_idx])
    colnames(coefs_selected)[5] <- "Pearson Cor vs TR"
    ## order the selected features by the p-values of their
    ## coefficients in a multiple linear model
    coefs_selected <- coefs_selected[order(coefs_selected[, 4]),
        ]

    dTICE_features <- data_frame[, features_selected_idx + 1]
    return(list(dTICE_features = dTICE_features, dTICE_feature_coefs = coefs_selected,
        R2 = R2_selected, R2_PWM = R2_PWM))
}

# generate dTICE features
for (species in names(species_tissue_list)) {
    for (i in 1:length(species_tissue_list[[species]])) {
        species_tissue <- species_tissue_list[[species]][i]
        if (species_tissue != "") {
            tissue_dot <- paste0(".", species_tissue)
            tissue_us <- paste0("_", species_tissue)
        } else {
            tissue_dot <- ""
            tissue_us <- ""
        }
        if (species != "hs") {
            assign(x = paste0(species, tissue_dot, ".dTICE_features"),
                value = make_dTICE_features(get(paste0(species,
                  tissue_dot, ".CDS_seqs")), get(paste0(species,
                  tissue_dot, ".UTR_seqs")), get(paste0(species,
                  tissue_dot, ".log10.TR")), get(paste0(species,
                  ".TR_top_pctl")), get(paste0(species, ".n_right")),
                  get(paste0(species, ".len_breaks")), ifexemplar = T))
```

```
        } else {
            assign(x = paste0(species, tissue_dot, ".dTICE_features"),
                value = make_dTICE_features(get(paste0(species,
                  tissue_dot, ".CDS_seqs")), get(paste0(species,
                  tissue_dot, ".UTR_seqs")), get(paste0(species,
                  tissue_dot, ".log10.TR")), get(paste0(species,
                  tissue_dot, ".TR_top_pctl")), get(paste0(species,
                  tissue_dot, ".n_right")), get(paste0(species,
                  tissue_dot, ".len_breaks")), ifexemplar = T))
        }
        saveRDS(get(paste0(species, tissue_dot, ".dTICE_features")),
            file = paste0("processed_data_w_polyA/dTICE_features/",
                species, tissue_dot, ".dTICE_features.rds"))
    }
}

##### 5' of TICE features (a mega-feature)
make_5ofTICE_features <- function(UTR_seqs, log10.TR, TR_top_pctl,
    n_left, n_5ofTICE, len_breaks, ifexemplar, selected_feature_names = NULL) {
    # use forward + BIC to select features among the following
    # features (1) 16 dinucleotide frequencies in the 5' end to
    # -(n_left+1) region (2) 64 trinucleotide frequencies in the
    # 5' end to -(n_left+1) region (3) PWM scores based on the
    # PWM constructed from the top 10% TR genes' sequences

    # UTR_seqs: a character vector of UTR sequences, with its
    # names as gene names log10.TR: a numeric vector of log10.TR
    # values, with its names as gene names TR_top_pctl: the
    # percentile to define TR top genes, e.g., 0.9 n_left: the
    # minimum UTR length of 'long' genes n_5ofTICE: the 5ofTICE
    # PWM length (from the 5' end) len_breaks: a numeric vector
    # containing the UTR length breaks, which divide genes into
    # groups ifexemplar = TRUE/FLASE; if TRUE, perform feature
    # selection; otherwise, use selected_feature_names
    # selected_feature_names = a character vector of the selected
    # feature names from the exemplar tissue

    n <- length(UTR_seqs)
    UTR_lens <- nchar(UTR_seqs)
    TR_top_UTR_seqs <- UTR_seqs[log10.TR >= quantile(log10.TR,
        prob = TR_top_pctl)]

    # cut each gene's 5' UTR sequence to the (5' end):-(n_left+1)
    # region
    UTR_seqs_5ofTICE <- substr(x = UTR_seqs, start = 1, stop = nchar(UTR_seqs) -
        n_left)
    TR_top_UTR_seqs_5ofTICE <- substr(x = TR_top_UTR_seqs, start = 1,
        stop = nchar(TR_top_UTR_seqs) - n_left)

    # count the dinucleotide frequencies in the (5'
    # end):-(n_left+1) region of each gene
    UTR_di_counts_5ofTICE <- sapply(di_nts, FUN = function(nuc) {
        stri_count_regex(UTR_seqs_5ofTICE, paste0("(?=", nuc,
```

```
        ")"))
})
rownames(UTR_di_counts_5ofTICE) <- names(UTR_seqs_5ofTICE)
UTR_di_freqs_5ofTICE <- UTR_di_counts_5ofTICE/rowSums(UTR_di_counts_5ofTICE)
UTR_di_freqs_5ofTICE[is.na(UTR_di_freqs_5ofTICE)] <- 0

# count the trinucleotide frequencies in the (5'
# end):-(n_left+1) region of each gene
UTR_tri_counts_5ofTICE <- sapply(tri_nts, FUN = function(nuc) {
    stri_count_regex(UTR_seqs_5ofTICE, paste0("(?=", nuc,
        ")"))
})
rownames(UTR_tri_counts_5ofTICE) <- names(UTR_seqs_5ofTICE)
UTR_tri_freqs_5ofTICE <- UTR_tri_counts_5ofTICE/rowSums(UTR_tri_counts_5ofTICE)
UTR_tri_freqs_5ofTICE[is.na(UTR_tri_freqs_5ofTICE)] <- 0

# calculate PWM scores based on the TR top PWM in the 5' most
# region of length n_5ofTICE for each gene the 5ofTICE PWM
# scores will be used
if (n_5ofTICE > 0) {
    PWM_5ofTICE_scores <- sapply(n_5ofTICE, FUN = function(len) {
        TR_top_PWM_5ofTICE <- UTR_pwm_5_aligned(TR_top_UTR_seqs_5ofTICE,
            len)
        UTR_pwm_5_aligned_score(TR_top_PWM_5ofTICE, UTR_seqs)
    })
    colnames(PWM_5ofTICE_scores) <- paste0("PWM", n_5ofTICE)
    features <- cbind(UTR_di_freqs_5ofTICE, UTR_tri_freqs_5ofTICE,
        PWM_5ofTICE_scores)
} else {
    # the 5ofTICE PWM scores will not be used
    features <- cbind(UTR_di_freqs_5ofTICE, UTR_tri_freqs_5ofTICE)
}

# combine the features into a design matrix
feature_names <- colnames(features)
cors <- apply(features, 2, FUN = function(x) cor(log10.TR,
    x, use = "complete.obs"))
if (length(len_breaks) > 0) {
    features_by_group <- NULL
    cors_by_group <- NULL  # Pearson correlation between log10.TR and every feature within a group
    num_groups <- length(len_breaks) + 1  # number of gene groups divided by UTR lengths
    for (i in 1:num_groups) {
        if (i == 1) {
            features_one_group <- as.numeric(UTR_lens < len_breaks[1]) *
              features
            colnames(features_one_group) <- paste(feature_names,
              paste0("<", len_breaks[1]), sep = "_")
            idx_one_group <- which(UTR_lens < len_breaks[1])
        } else if (i == num_groups) {
            features_one_group <- as.numeric(UTR_lens >=
              len_breaks[num_groups - 1]) * cbind(features,
              rep(1, n))
            colnames(features_one_group) <- paste(c(feature_names,
```

```
                  "Intercept"), paste0(">=", len_breaks[num_groups -
                  1]), sep = "_")
                idx_one_group <- which(UTR_lens >= len_breaks[num_groups -
                  1])
            } else {
                features_one_group <- as.numeric(UTR_lens >=
                  len_breaks[i - 1] & UTR_lens < len_breaks[i]) *
                  cbind(features, rep(1, n))
                colnames(features_one_group) <- paste(c(feature_names,
                  "Intercept"), paste0(">=", len_breaks[i - 1],
                  "&<", len_breaks[i]), sep = "_")
                idx_one_group <- which(UTR_lens >= len_breaks[i -
                  1] & UTR_lens < len_breaks[i])
            }
            features_by_group <- cbind(features_by_group, features_one_group)
            cors_one_group <- apply(features_one_group, 2, FUN = function(x) cor(log10.TR[idx_one_group]
                x[idx_one_group], use = "complete.obs"))
            cors_by_group <- c(cors_by_group, cors_one_group)
        }
        features <- features_by_group
        cors <- cors_by_group
    }

    # combine the response and the design matrix into a data
    # frame use forward selection + Bayesian Information
    # Criterion (BIC) to select features the 5ofTICE PWM scores
    # will be used
    if (n_5ofTICE > 0) {
        PWM_idx <- which(grepl("PWM", colnames(features)))
        data_frame <- cbind.data.frame(log10.TR = log10.TR, features[,
            PWM_idx], features[, -PWM_idx])
        colnames(data_frame) <- c("log10.TR", colnames(features)[PWM_idx],
            colnames(features)[-PWM_idx])
        cors <- c(cors[PWM_idx], cors[-PWM_idx])
        if (ifexemplar) {
            regfit.fwd <- regsubsets(log10.TR ~ ., data = data_frame,
                nvmax = 70, force.in = 1:length(PWM_idx), method = "forward")
        }
    } else {
        # the 5ofTICE PWM scores will not be used
        data_frame <- cbind.data.frame(log10.TR = log10.TR, features)
        if (ifexemplar) {
            regfit.fwd <- regsubsets(log10.TR ~ ., data = data_frame,
                nvmax = 70, method = "forward")
        }
    }

    if (ifexemplar) {
        reg.summary <- summary(regfit.fwd)
        idx_bic <- which.min(reg.summary$bic)
        features_selected_idx <- which(reg.summary$outmat[idx_bic,
            ] == "*")
    } else {
```

```
        features_selected_idx <- which(colnames(data_frame) %in%
            selected_feature_names) - 1
    }

    lm_selected <- summary(lm(log10.TR ~ ., data = data_frame[,
        c(1, features_selected_idx + 1)]))
    coefs_selected <- lm_selected$coefficients[-1, ]
    R2_selected <- lm_selected$r.squared
    coefs_selected <- cbind(coefs_selected, cors[features_selected_idx])
    colnames(coefs_selected)[5] <- "Pearson Cor vs TR"
    ## order the selected features by the p-values of their
    ## coefficients in a multiple linear model
    coefs_selected <- coefs_selected[order(coefs_selected[, 4]),
        ]

    fiveofTICE_features <- data_frame[, features_selected_idx +
        1]
    return(list(fiveofTICE_features = fiveofTICE_features, fiveofTICE_feature_coefs = coefs_selected,
        R2 = R2_selected))
}

# generate 5' of TICE features
for (species in names(species_tissue_list)) {
    for (i in 1:length(species_tissue_list[[species]])) {
        species_tissue <- species_tissue_list[[species]][i]
        if (species_tissue != "") {
            tissue_dot <- paste0(".", species_tissue)
            tissue_us <- paste0("_", species_tissue)
        } else {
            tissue_dot <- ""
            tissue_us <- ""
        }
        if (species != "hs") {
            assign(x = paste0(species, tissue_dot, ".5ofTICE_features"),
                value = make_5ofTICE_features(get(paste0(species,
                  tissue_dot, ".UTR_seqs")), get(paste0(species,
                  tissue_dot, ".log10.TR")), get(paste0(species,
                  ".TR_top_pctl")), get(paste0(species, ".n_left")),
                  get(paste0(species, ".n_5ofTICE")), get(paste0(species,
                    ".len_breaks")), ifexemplar = T))
        } else {
            assign(x = paste0(species, tissue_dot, ".5ofTICE_features"),
                value = make_5ofTICE_features(get(paste0(species,
                  tissue_dot, ".UTR_seqs")), get(paste0(species,
                  tissue_dot, ".log10.TR")), get(paste0(species,
                  tissue_dot, ".TR_top_pctl")), get(paste0(species,
                  tissue_dot, ".n_left")), get(paste0(species,
                  tissue_dot, ".n_5ofTICE")), get(paste0(species,
                  tissue_dot, ".len_breaks")), ifexemplar = T))
        }
        saveRDS(get(paste0(species, tissue_dot, ".5ofTICE_features")),
            file = paste0("processed_data_w_polyA/5ofTICE_features/",
                species, tissue_dot, ".5ofTICE_features.rds"))
```

```
    }
}

##### 20 amino acid frequencies in the CDS (a mega-feature) 'K'
##### is the reference level
make_20aa_features <- function(aa_counts, log10.TR, log10.UTR_lens,
    len_breaks) {
    n <- length(log10.UTR_lens)
    UTR_lens <- 10^log10.UTR_lens
    features <- aa_counts/rowSums(aa_counts)
    features <- features[, -1]  # use 'K' as the reference level
    cors <- apply(features, 2, FUN = function(x) cor(log10.TR,
        x, use = "complete.obs"))
    feature_names <- colnames(features)
    if (length(len_breaks) > 0) {
        features_by_group <- NULL
        cors_by_group <- NULL  # Pearson correlation between log10.TR and every feature within a group
        num_groups <- length(len_breaks) + 1  # number of gene groups divided by UTR lengths
        for (i in 1:num_groups) {
            if (i == 1) {
                features_one_group <- as.numeric(UTR_lens < len_breaks[1]) *
                  features
                colnames(features_one_group) <- paste(feature_names,
                  paste0("<", len_breaks[1]), sep = "_")
                idx_one_group <- which(UTR_lens < len_breaks[1])
            } else if (i == num_groups) {
                features_one_group <- as.numeric(UTR_lens >=
                  len_breaks[num_groups - 1]) * cbind(features,
                  rep(1, n))
                colnames(features_one_group) <- paste(c(feature_names,
                  "Intercept"), paste0(">=", len_breaks[num_groups -
                  1]), sep = "_")
                idx_one_group <- which(UTR_lens >= len_breaks[num_groups -
                  1])
            } else {
                features_one_group <- as.numeric(UTR_lens >=
                  len_breaks[i - 1] & UTR_lens < len_breaks[i]) *
                  cbind(features, rep(1, n))
                colnames(features_one_group) <- paste(c(feature_names,
                  "Intercept"), paste0(">=", len_breaks[i - 1],
                  "&<", len_breaks[i]), sep = "_")
                idx_one_group <- which(UTR_lens >= len_breaks[i -
                  1] & UTR_lens < len_breaks[i])
            }
            features_by_group <- cbind(features_by_group, features_one_group)
            cors_one_group <- apply(features_one_group, 2, FUN = function(x) cor(log10.TR[idx_one_group]
                x[idx_one_group], use = "complete.obs"))
            cors_by_group <- c(cors_by_group, cors_one_group)
        }
        features <- features_by_group
        cors <- cors_by_group
    }
    # combine the response and the design matrix into a data
```

```
    # frame
    data_frame <- cbind.data.frame(log10.TR = log10.TR, features)
    lm_selected <- summary(lm(log10.TR ~ ., data = data_frame))
    coefs_selected <- lm_selected$coefficients[-1, ]
    R2_selected <- lm_selected$r.squared
    coefs_selected <- cbind(coefs_selected, cors)
    colnames(coefs_selected)[5] <- "Pearson Cor vs TR"
    ## order the features by the p-values of their coefficients in
    ## a multiple linear model
    coefs_selected <- coefs_selected[order(coefs_selected[, 4]),
        ]
    return(list(aa_features = features, aa_feature_coefs = coefs_selected,
        R2 = R2_selected))
}

# generate aa features
for (species in names(species_tissue_list)) {
    for (i in 1:length(species_tissue_list[[species]])) {
        species_tissue <- species_tissue_list[[species]][i]
        if (species_tissue != "") {
            tissue_dot <- paste0(".", species_tissue)
            tissue_us <- paste0("_", species_tissue)
        } else {
            tissue_dot <- ""
            tissue_us <- ""
        }
        ## count the frequency of 64 codons (including 3 stop codons)
        ## in every gene
        CDS_seqs <- get(paste0(species, tissue_dot, ".CDS_seqs"))
        codon_counts <- t(sapply(CDS_seqs, FUN = function(seq) {
            sst <- strsplit(seq, "")[[1]]
            n <- length(sst)
            out <- paste0(sst[seq(from = 1, to = n, by = 3)],
                sst[seq(from = 2, to = n, by = 3)], sst[seq(from = 3,
                  to = n, by = 3)])
            table(out)[tri_nts]
        }))
        codon_counts[is.na(codon_counts)] <- 0
        rownames(codon_counts) <- names(CDS_seqs)
        colnames(codon_counts) <- tri_nts
        ## count the frequency of 20 amino acids (AAs) in every gene
        aa_counts <- sapply(aas, FUN = function(aa) {
            idx <- which(tri_nts_to_aas == aa)
            if (length(idx) == 1) {
                codon_counts[, idx]
            } else {
                rowSums(codon_counts[, idx])
            }
        })

        saveRDS(aa_counts/rowSums(aa_counts), file = paste0("processed_data_w_polyA/aa_features/",
            species, tissue_dot, ".aa_freqs.rds"))
```

```
        # features divided by 5'UTR length groups
        if (species != "hs") {
            aa_features <- make_20aa_features(aa_counts, get(paste0(species,
                tissue_dot, ".log10.TR")), get(paste0(species,
                tissue_dot, ".log10.UTR_lens")), get(paste0(species,
                ".len_breaks")))
        } else {
            aa_features <- make_20aa_features(aa_counts, get(paste0(species,
                tissue_dot, ".log10.TR")), get(paste0(species,
                tissue_dot, ".log10.UTR_lens")), get(paste0(species,
                tissue_dot, ".len_breaks")))
        }

        saveRDS(aa_features, file = paste0("processed_data_w_polyA/aa_features/",
            species, tissue_dot, ".aa_features.rds"))
    }
}

##### 61 nonstop codon frequencies in the CDS (a mega-feature)
##### 'AAA' is the reference level
make_61codon_features <- function(codon_counts, log10.TR, log10.UTR_lens,
    len_breaks) {
    n <- length(log10.UTR_lens)
    UTR_lens <- 10^log10.UTR_lens
    features <- codon_counts/rowSums(codon_counts)
    features <- features[, -1]  # use 'AAA' as the reference level
    cors <- apply(features, 2, FUN = function(x) cor(log10.TR,
        x, use = "complete.obs"))
    feature_names <- colnames(features)
    if (length(len_breaks) > 0) {
        features_by_group <- NULL
        cors_by_group <- NULL  # Pearson correlation between log10.TR and every feature within a group
        num_groups <- length(len_breaks) + 1  # number of gene groups divided by UTR lengths
        for (i in 1:num_groups) {
            if (i == 1) {
                features_one_group <- as.numeric(UTR_lens < len_breaks[1]) *
                  features
                colnames(features_one_group) <- paste(feature_names,
                  paste0("<", len_breaks[1]), sep = "_")
                idx_one_group <- which(UTR_lens < len_breaks[1])
            } else if (i == num_groups) {
                features_one_group <- as.numeric(UTR_lens >=
                  len_breaks[num_groups - 1]) * cbind(features,
                  rep(1, n))
                colnames(features_one_group) <- paste(c(feature_names,
                  "Intercept"), paste0(">=", len_breaks[num_groups -
                  1]), sep = "_")
                idx_one_group <- which(UTR_lens >= len_breaks[num_groups -
                  1])
            } else {
                features_one_group <- as.numeric(UTR_lens >=
                  len_breaks[i - 1] & UTR_lens < len_breaks[i]) *
                  cbind(features, rep(1, n))
```

```
                colnames(features_one_group) <- paste(c(feature_names,
                  "Intercept"), paste0(">=", len_breaks[i - 1],
                  "&<", len_breaks[i]), sep = "_")
                idx_one_group <- which(UTR_lens >= len_breaks[i -
                  1] & UTR_lens < len_breaks[i])
            }
            features_by_group <- cbind(features_by_group, features_one_group)
            cors_one_group <- apply(features_one_group, 2, FUN = function(x) cor(log10.TR[idx_one_group]
                x[idx_one_group], use = "complete.obs"))
            cors_by_group <- c(cors_by_group, cors_one_group)
        }
        features <- features_by_group
        cors <- cors_by_group
    }
    # combine the response and the design matrix into a data
    # frame
    data_frame <- cbind.data.frame(log10.TR = log10.TR, features)
    lm_selected <- summary(lm(log10.TR ~ ., data = data_frame))
    coefs_selected <- lm_selected$coefficients[-1, ]
    R2_selected <- lm_selected$r.squared
    coefs_selected <- cbind(coefs_selected, cors)
    colnames(coefs_selected)[5] <- "Pearson Cor vs TR"
    ## order the features by the p-values of their coefficients in
    ## a multiple linear model
    coefs_selected <- coefs_selected[order(coefs_selected[, 4]),
        ]
    return(list(codon_features = features, codon_feature_coefs = coefs_selected,
        R2 = R2_selected))
}

# generate codon features
for (species in names(species_tissue_list)) {
    for (i in 1:length(species_tissue_list[[species]])) {
        species_tissue <- species_tissue_list[[species]][i]
        if (species_tissue != "") {
            tissue_dot <- paste0(".", species_tissue)
            tissue_us <- paste0("_", species_tissue)
        } else {
            tissue_dot <- ""
            tissue_us <- ""
        }
        ## count the frequency of 64 codons (including 3 stop codons)
        ## in every gene
        CDS_seqs <- get(paste0(species, tissue_dot, ".CDS_seqs"))
        codon_counts <- t(sapply(CDS_seqs, FUN = function(seq) {
            sst <- strsplit(seq, "")[[1]]
            n <- length(sst)
            out <- paste0(sst[seq(from = 1, to = n, by = 3)],
                sst[seq(from = 2, to = n, by = 3)], sst[seq(from = 3,
                  to = n, by = 3)])
            table(out)[nonstop_codons]
        }))
        codon_counts[is.na(codon_counts)] <- 0
```

```
        rownames(codon_counts) <- names(CDS_seqs)
        colnames(codon_counts) <- nonstop_codons

        saveRDS(codon_counts/rowSums(codon_counts), file = paste0("processed_data_w_polyA/codon_features
            species, tissue_dot, ".codon_freqs.rds"))

        # features divided by 5'UTR length groups
        if (species != "hs") {
            codon_features <- make_61codon_features(codon_counts,
                get(paste0(species, tissue_dot, ".log10.TR")),
                get(paste0(species, tissue_dot, ".log10.UTR_lens")),
                get(paste0(species, ".len_breaks")))
        } else {
            codon_features <- make_61codon_features(codon_counts,
                get(paste0(species, tissue_dot, ".log10.TR")),
                get(paste0(species, tissue_dot, ".log10.UTR_lens")),
                get(paste0(species, tissue_dot, ".len_breaks")))
        }

        saveRDS(codon_features, file = paste0("processed_data_w_polyA/codon_features/",
            species, tissue_dot, ".codon_features.rds"))
    }
}
#####

##### 61 nonstop codon frequencies in the first and second halves
##### of CDS
for (species in names(species_tissue_list)) {
    for (i in 1:length(species_tissue_list[[species]])) {
        species_tissue <- species_tissue_list[[species]][i]
        if (species_tissue != "") {
            tissue_dot <- paste0(".", species_tissue)
            tissue_us <- paste0("_", species_tissue)
        } else {
            tissue_dot <- ""
            tissue_us <- ""
        }
        ## count the frequency of 64 codons (including 3 stop codons)
        ## in every gene's first and second halves of CDS
        CDS_seqs <- get(paste0(species, tissue_dot, ".CDS_seqs"))
        CDS_seqs_1 <- substr(x = CDS_seqs, start = 1, stop = round(nchar(CDS_seqs)/2))
        CDS_seqs_2 <- substr(x = CDS_seqs, start = round(nchar(CDS_seqs)/2) +
            1, stop = nchar(CDS_seqs))

        codon_counts_1 <- t(sapply(CDS_seqs_1, FUN = function(seq) {
            sst <- strsplit(seq, "")[[1]]
            n <- length(sst)
            out <- paste0(sst[seq(from = 1, to = n, by = 3)],
                sst[seq(from = 2, to = n, by = 3)], sst[seq(from = 3,
                  to = n, by = 3)])
            table(out)[nonstop_codons]
        }))
        codon_counts_1[is.na(codon_counts_1)] <- 0
```

```
        rownames(codon_counts_1) <- names(CDS_seqs_1)
        colnames(codon_counts_1) <- nonstop_codons

        codon_counts_2 <- t(sapply(CDS_seqs_2, FUN = function(seq) {
            sst <- strsplit(seq, "")[[1]]
            n <- length(sst)
            out <- paste0(sst[seq(from = 1, to = n, by = 3)],
                sst[seq(from = 2, to = n, by = 3)], sst[seq(from = 3,
                  to = n, by = 3)])
            table(out)[nonstop_codons]
        }))
        codon_counts_2[is.na(codon_counts_2)] <- 0
        rownames(codon_counts_2) <- names(CDS_seqs_2)
        colnames(codon_counts_2) <- nonstop_codons

        if (species != "hs") {
            codon_features_1 <- make_61codon_features(codon_counts_1,
                get(paste0(species, tissue_dot, ".log10.TR")),
                get(paste0(species, tissue_dot, ".log10.UTR_lens")),
                get(paste0(species, ".len_breaks")))
            codon_features_2 <- make_61codon_features(codon_counts_2,
                get(paste0(species, tissue_dot, ".log10.TR")),
                get(paste0(species, tissue_dot, ".log10.UTR_lens")),
                get(paste0(species, ".len_breaks")))
        } else {
            codon_features_1 <- make_61codon_features(codon_counts_1,
                get(paste0(species, tissue_dot, ".log10.TR")),
                get(paste0(species, tissue_dot, ".log10.UTR_lens")),
                get(paste0(species, tissue_dot, ".len_breaks")))
            codon_features_2 <- make_61codon_features(codon_counts_2,
                get(paste0(species, tissue_dot, ".log10.TR")),
                get(paste0(species, tissue_dot, ".log10.UTR_lens")),
                get(paste0(species, tissue_dot, ".len_breaks")))
        }

        saveRDS(codon_features_1, file = paste0("processed_data_w_polyA/codon_features/",
            species, tissue_dot, ".codon_features_1.rds"))
        saveRDS(codon_features_2, file = paste0("processed_data_w_polyA/codon_features/",
            species, tissue_dot, ".codon_features_2.rds"))
    }
}

##### 61 synonymous codon frequencies in the CDS (a mega-feature)
##### For example, AAA, AAT, AAC and AAG code the same AA, so we
##### compute their synonymous codon frequencies by normalizing
##### AA frequencies as follows AAA / (AAA+AAT+AAG+AAC) AAT /
##### (AAA+AAT+AAG+AAC) AAC / (AAA+AAT+AAG+AAC) AAG /
##### (AAA+AAT+AAG+AAC)
make_61syncodon_features <- function(syncodon_ratios, log10.TR,
    log10.UTR_lens, len_breaks) {
    n <- length(log10.UTR_lens)
    UTR_lens <- 10^log10.UTR_lens
    features <- syncodon_ratios
```

```
    cors <- apply(features, 2, FUN = function(x) cor(log10.TR,
        x, use = "complete.obs"))
    feature_names <- colnames(features)
    if (length(len_breaks) > 0) {
        features_by_group <- NULL
        cors_by_group <- NULL  # Pearson correlation between log10.TR and every feature within a group
        num_groups <- length(len_breaks) + 1  # number of gene groups divided by UTR lengths
        for (i in 1:num_groups) {
            if (i == 1) {
                features_one_group <- as.numeric(UTR_lens < len_breaks[1]) *
                  features
                colnames(features_one_group) <- paste(feature_names,
                  paste0("<", len_breaks[1]), sep = "_")
                idx_one_group <- which(UTR_lens < len_breaks[1])
            } else if (i == num_groups) {
                features_one_group <- as.numeric(UTR_lens >=
                  len_breaks[num_groups - 1]) * cbind(features,
                  rep(1, n))
                colnames(features_one_group) <- paste(c(feature_names,
                  "Intercept"), paste0(">=", len_breaks[num_groups -
                  1]), sep = "_")
                idx_one_group <- which(UTR_lens >= len_breaks[num_groups -
                  1])
            } else {
                features_one_group <- as.numeric(UTR_lens >=
                  len_breaks[i - 1] & UTR_lens < len_breaks[i]) *
                  cbind(features, rep(1, n))
                colnames(features_one_group) <- paste(c(feature_names,
                  "Intercept"), paste0(">=", len_breaks[i - 1],
                  "&<", len_breaks[i]), sep = "_")
                idx_one_group <- which(UTR_lens >= len_breaks[i -
                  1] & UTR_lens < len_breaks[i])
            }
            features_by_group <- cbind(features_by_group, features_one_group)
            cors_one_group <- apply(features_one_group, 2, FUN = function(x) cor(log10.TR[idx_one_group]
                x[idx_one_group], use = "complete.obs"))
            cors_by_group <- c(cors_by_group, cors_one_group)
        }
        features <- features_by_group
        cors <- cors_by_group
    }
    # combine the response and the design matrix into a data
    # frame
    data_frame <- cbind.data.frame(log10.TR = log10.TR, features)
    lm_selected <- summary(lm(log10.TR ~ ., data = data_frame))
    coefs_selected <- lm_selected$coefficients[-1, ]
    R2_selected <- lm_selected$r.squared
    coefs_selected <- cbind(coefs_selected, cors)
    colnames(coefs_selected)[5] <- "Pearson Cor vs TR"
    ## order the features by the p-values of their coefficients in
    ## a multiple linear model
    coefs_selected <- coefs_selected[order(coefs_selected[, 4]),
        ]
```

```r
    return(list(codon_features = features, codon_feature_coefs = coefs_selected,
        R2 = R2_selected))
}

# generate synonymous codon features
for (species in names(species_tissue_list)) {
    for (i in 1:length(species_tissue_list[[species]])) {
        species_tissue <- species_tissue_list[[species]][i]
        if (species_tissue != "") {
            tissue_dot <- paste0(".", species_tissue)
            tissue_us <- paste0("_", species_tissue)
        } else {
            tissue_dot <- ""
            tissue_us <- ""
        }
        ## count the frequency of 64 codons (including 3 stop codons)
        ## in every gene
        CDS_seqs <- get(paste0(species, tissue_dot, ".CDS_seqs"))
        codon_counts <- t(sapply(CDS_seqs, FUN = function(seq) {
            sst <- strsplit(seq, "")[[1]]
            n <- length(sst)
            out <- paste0(sst[seq(from = 1, to = n, by = 3)],
                sst[seq(from = 2, to = n, by = 3)], sst[seq(from = 3,
                  to = n, by = 3)])
            table(out)[nonstop_codons]
        }))
        codon_counts[is.na(codon_counts)] <- 0
        rownames(codon_counts) <- names(CDS_seqs)
        colnames(codon_counts) <- nonstop_codons

        ## normalize the codon counts by their corresponding AA count
        syncodon_ratios <- codon_counts
        for (aa in aas) {
            tmp_codons <- names(tri_nts_to_aas[which(tri_nts_to_aas ==
                aa)])
            syncodon_ratios[, tmp_codons] <- codon_counts[, tmp_codons]/rowSums(codon_counts[,
                tmp_codons, drop = F])
        }

        saveRDS(syncodon_ratios, file = paste0("processed_data_w_polyA/syncodon_features/",
            species, tissue_dot, ".syncodon_ratios.rds"))

        # features divided by 5'UTR length groups
        if (species != "hs") {
            syncodon_features <- make_61syncodon_features(syncodon_ratios,
                get(paste0(species, tissue_dot, ".log10.TR")),
                get(paste0(species, tissue_dot, ".log10.UTR_lens")),
                get(paste0(species, ".len_breaks")))
        } else {
            syncodon_features <- make_61syncodon_features(syncodon_ratios,
                get(paste0(species, tissue_dot, ".log10.TR")),
                get(paste0(species, tissue_dot, ".log10.UTR_lens")),
                get(paste0(species, tissue_dot, ".len_breaks")))
```

```
        }

        saveRDS(syncodon_features, file = paste0("processed_data_w_polyA/syncodon_features/",
            species, tissue_dot, ".syncodon_features.rds"))
    }
}

##### tRNA abundances for sc
for (species in "sc") {
    for (i in 1:length(species_tissue_list[[species]])) {
        species_tissue <- species_tissue_list[[species]][i]
        if (species_tissue != "") {
            tissue_dot <- paste0(".", species_tissue)
            tissue_us <- paste0("_", species_tissue)
        } else {
            tissue_dot <- ""
            tissue_us <- ""
        }

        # read in combined tRNAs abundances (one per AA)
        aa_tRNA_abun <- read.table("../data/Weinberg tRNA abundances (aa).txt",
            header = T, sep = "\t")
        aa_tRNA_abun <- sapply(aas, FUN = function(aa) {
            aa_tRNA_abun[aa_tRNA_abun[, 1] == aa, 2]
        })
        names(aa_tRNA_abun) <- aas

        # read in tRNAs abundances (one per codon)
        codon_tRNA_abun <- read.table("../data/Weinberg tRNA abundances (codon).txt",
            header = F, skip = 1, sep = "\t")
        codon_tRNA_abun <- sapply(nonstop_codons, FUN = function(codon) {
            codon_tRNA_abun[codon_tRNA_abun[, 2] == codon, 5]
        })

        saveRDS(aa_tRNA_abun, file = paste0("processed_data_w_polyA/tRNA_abun/",
            species, tissue_dot, ".aa_tRNA_abun.rds"))

        saveRDS(codon_tRNA_abun, file = paste0("processed_data_w_polyA/tRNA_abun/",
            species, tissue_dot, ".codon_tRNA_abun.rds"))
    }
}
```

##### Read in the already generated mega-features

```
# read in uTICE features
for (species in names(species_tissue_list)) {
    for (i in 1:length(species_tissue_list[[species]])) {
        species_tissue <- species_tissue_list[[species]][i]
        if (species_tissue != "") {
            tissue_dot <- paste0(".", species_tissue)
            tissue_us <- paste0("_", species_tissue)
        } else {
```

```
            tissue_dot <- ""
            tissue_us <- ""
        }
        assign(x = paste0(species, tissue_dot, ".uTICE_features"),
            value = readRDS(file = paste0("processed_data_w_polyA/uTICE_features/",
                species, tissue_dot, ".uTICE_features.rds")))
    }
}

# read in dTICE features
for (species in names(species_tissue_list)) {
    for (i in 1:length(species_tissue_list[[species]])) {
        species_tissue <- species_tissue_list[[species]][i]
        if (species_tissue != "") {
            tissue_dot <- paste0(".", species_tissue)
            tissue_us <- paste0("_", species_tissue)
        } else {
            tissue_dot <- ""
            tissue_us <- ""
        }
        assign(x = paste0(species, tissue_dot, ".dTICE_features"),
            value = readRDS(file = paste0("processed_data_w_polyA/dTICE_features/",
                species, tissue_dot, ".dTICE_features.rds")))
    }
}

# read in 5' of TICE features
for (species in names(species_tissue_list)) {
    for (i in 1:length(species_tissue_list[[species]])) {
        species_tissue <- species_tissue_list[[species]][i]
        if (species_tissue != "") {
            tissue_dot <- paste0(".", species_tissue)
            tissue_us <- paste0("_", species_tissue)
        } else {
            tissue_dot <- ""
            tissue_us <- ""
        }
        assign(x = paste0(species, tissue_dot, ".5ofTICE_features"),
            value = readRDS(file = paste0("processed_data_w_polyA/5ofTICE_features/",
                species, tissue_dot, ".5ofTICE_features.rds")))
    }
}

# read in aa features
for (species in names(species_tissue_list)) {
    for (i in 1:length(species_tissue_list[[species]])) {
        species_tissue <- species_tissue_list[[species]][i]
        if (species_tissue != "") {
            tissue_dot <- paste0(".", species_tissue)
            tissue_us <- paste0("_", species_tissue)
        } else {
            tissue_dot <- ""
            tissue_us <- ""
```

```
        }
        assign(x = paste0(species, tissue_dot, ".aa_features"),
            value = readRDS(file = paste0("processed_data_w_polyA/aa_features/",
                species, tissue_dot, ".aa_features.rds")))
    }
}

# read in codon features
for (species in names(species_tissue_list)) {
    for (i in 1:length(species_tissue_list[[species]])) {
        species_tissue <- species_tissue_list[[species]][i]
        if (species_tissue != "") {
            tissue_dot <- paste0(".", species_tissue)
            tissue_us <- paste0("_", species_tissue)
        } else {
            tissue_dot <- ""
            tissue_us <- ""
        }
        assign(x = paste0(species, tissue_dot, ".codon_features"),
            value = readRDS(file = paste0("processed_data_w_polyA/codon_features/",
                species, tissue_dot, ".codon_features.rds")))
    }
}
```

##### Divide original single features into by-group features

```
### a function
div_original_features <- function(features, feature_names, log10.UTR_lens,
    len_breaks) {
    # for features that have not been divided into groups
    # (original features)
    n <- length(log10.UTR_lens)
    UTR_lens <- 10^log10.UTR_lens

    features <- matrix(features, ncol = 1)
    colnames(features) <- feature_names

    if (length(len_breaks) > 0) {
        features_by_group <- NULL
        num_groups <- length(len_breaks) + 1  # number of gene groups divided by UTR lengths
        for (i in 1:num_groups) {
            if (i == 1) {
                features_one_group <- as.numeric(UTR_lens < len_breaks[1]) *
                  features
                colnames(features_one_group) <- paste(feature_names,
                  paste0("<", len_breaks[1]), sep = "_")
            } else if (i == num_groups) {
                features_one_group <- as.numeric(UTR_lens >=
                  len_breaks[num_groups - 1]) * cbind(features,
                  rep(1, n))
                colnames(features_one_group) <- paste(c(feature_names,
                  "Intercept"), paste0(">=", len_breaks[num_groups -
```

```
                1]), sep = "_")
            } else {
                features_one_group <- as.numeric(UTR_lens >=
                  len_breaks[i - 1] & UTR_lens < len_breaks[i]) *
                  cbind(features, rep(1, n))
                colnames(features_one_group) <- paste(c(feature_names,
                  "Intercept"), paste0(">=", len_breaks[i - 1],
                  "&<", len_breaks[i]), sep = "_")
            }
            features_by_group <- cbind(features_by_group, features_one_group)
        }
        features <- features_by_group
    }

    return(features)
}

### log10.UTR_lens
for (species in names(species_tissue_list)) {
    for (i in 1:length(species_tissue_list[[species]])) {
        species_tissue <- species_tissue_list[[species]][i]
        if (species_tissue != "") {
            tissue_dot <- paste0(".", species_tissue)
            tissue_us <- paste0("_", species_tissue)
        } else {
            tissue_dot <- ""
            tissue_us <- ""
        }
        if (species != "hs") {
            assign(x = paste0(species, tissue_dot, ".log10.UTR_lens.div"),
                value = div_original_features(get(paste0(species,
                  tissue_dot, ".log10.UTR_lens")), "log10.UTR_lens",
                  get(paste0(species, tissue_dot, ".log10.UTR_lens")),
                  get(paste0(species, ".len_breaks"))))
        } else {
            assign(x = paste0(species, tissue_dot, ".log10.UTR_lens.div"),
                value = div_original_features(get(paste0(species,
                  tissue_dot, ".log10.UTR_lens")), "log10.UTR_lens",
                  get(paste0(species, tissue_dot, ".log10.UTR_lens")),
                  get(paste0(species, tissue_dot, ".len_breaks"))))
        }
    }
}

### log10.CDS_lens
for (species in names(species_tissue_list)) {
    for (i in 1:length(species_tissue_list[[species]])) {
        species_tissue <- species_tissue_list[[species]][i]
        if (species_tissue != "") {
            tissue_dot <- paste0(".", species_tissue)
            tissue_us <- paste0("_", species_tissue)
        } else {
            tissue_dot <- ""
```

```
            tissue_us <- ""
        }
        if (species != "hs") {
            assign(x = paste0(species, tissue_dot, ".log10.CDS_lens.div"),
                value = div_original_features(get(paste0(species,
                  tissue_dot, ".log10.CDS_lens")), "log10.CDS_lens",
                  get(paste0(species, tissue_dot, ".log10.UTR_lens")),
                  get(paste0(species, ".len_breaks"))))
        } else {
            assign(x = paste0(species, tissue_dot, ".log10.CDS_lens.div"),
                value = div_original_features(get(paste0(species,
                  tissue_dot, ".log10.CDS_lens")), "log10.CDS_lens",
                  get(paste0(species, tissue_dot, ".log10.UTR_lens")),
                  get(paste0(species, tissue_dot, ".len_breaks"))))
        }
    }
}

### uAUG_counts
for (species in names(species_tissue_list)) {
    for (i in 1:length(species_tissue_list[[species]])) {
        species_tissue <- species_tissue_list[[species]][i]
        if (species_tissue != "") {
            tissue_dot <- paste0(".", species_tissue)
            tissue_us <- paste0("_", species_tissue)
        } else {
            tissue_dot <- ""
            tissue_us <- ""
        }
        if (species != "hs") {
            assign(x = paste0(species, tissue_dot, ".uAUG_counts.div"),
                value = div_original_features(get(paste0(species,
                  tissue_dot, ".uAUG_counts")), "uAUG_counts",
                  get(paste0(species, tissue_dot, ".log10.UTR_lens")),
                  get(paste0(species, ".len_breaks"))))
        } else {
            assign(x = paste0(species, tissue_dot, ".uAUG_counts.div"),
                value = div_original_features(get(paste0(species,
                  tissue_dot, ".uAUG_counts")), "uAUG_counts",
                  get(paste0(species, tissue_dot, ".log10.UTR_lens")),
                  get(paste0(species, tissue_dot, ".len_breaks"))))
        }
    }
}

### CDS_fold_energy
for (species in names(species_tissue_list)) {
    for (i in 1:length(species_tissue_list[[species]])) {
        species_tissue <- species_tissue_list[[species]][i]
        if (species_tissue != "") {
            tissue_dot <- paste0(".", species_tissue)
            tissue_us <- paste0("_", species_tissue)
        } else {
```

```
            tissue_dot <- ""
            tissue_us <- ""
        }
        if (species != "hs") {
            assign(x = paste0(species, tissue_dot, ".CDS_fold_energy.div"),
                value = div_original_features(get(paste0(species,
                  tissue_dot, ".CDS_fold_energy")), "CDS_fold_energy",
                  get(paste0(species, tissue_dot, ".log10.UTR_lens")),
                  get(paste0(species, ".len_breaks"))))
        } else {
            assign(x = paste0(species, tissue_dot, ".CDS_fold_energy.div"),
                value = div_original_features(get(paste0(species,
                  tissue_dot, ".CDS_fold_energy")), "CDS_fold_energy",
                  get(paste0(species, tissue_dot, ".log10.UTR_lens")),
                  get(paste0(species, tissue_dot, ".len_breaks"))))
        }
    }
}

### polyA_lens
for (species in names(species_tissue_list)) {
    for (i in 1:length(species_tissue_list[[species]])) {
        species_tissue <- species_tissue_list[[species]][i]
        if (species_tissue != "") {
            tissue_dot <- paste0(".", species_tissue)
            tissue_us <- paste0("_", species_tissue)
        } else {
            tissue_dot <- ""
            tissue_us <- ""
        }
        if (species != "hs") {
            assign(x = paste0(species, tissue_dot, ".polyA_lens.div"),
                value = div_original_features(get(paste0(species,
                  tissue_dot, ".polyA_lens")), "polyA_lens",
                  get(paste0(species, tissue_dot, ".log10.UTR_lens")),
                  get(paste0(species, ".len_breaks"))))
        } else {
            assign(x = paste0(species, tissue_dot, ".polyA_lens.div"),
                value = div_original_features(get(paste0(species,
                  tissue_dot, ".polyA_lens")), "polyA_lens",
                  get(paste0(species, tissue_dot, ".log10.UTR_lens")),
                  get(paste0(species, tissue_dot, ".len_breaks"))))
        }
    }
}
```

#### Save the selected RNA folding features to excel files

```
for (species in names(species_tissue_list)) {
    for (species_tissue in species_tissue_list[[species]]) {
        if (species_tissue != "") {
```

```
            tissue_dot <- paste0(".", species_tissue)
            tissue_us <- paste0("_", species_tissue)
        } else {
            tissue_dot <- ""
            tissue_us <- ""
        }
        if (species != "sp") {
            output <- readRDS(file = paste0("processed_data_w_polyA/folding_energy_features_selected/",
                species, tissue_us, "_UTR_folding_energy_features_selected.rds"))[[2]]
        } else {
            output <- readRDS(file = paste0("processed_data_w_polyA/folding_energy_features_selected/",
                species, tissue_us, "_UTR_folding_energy_features_selected_trun.rds"))[[2]]
        }
        write.csv(output, file = paste0("tables/folding_energy_features_selected/",
            species, tissue_dot, ".fold_energy_features_selected.csv"),
            quote = F)
    }
}
```

#### Save the selected 5ofTICE, uTICE and dTICE features to excel files

```
for (species in names(species_tissue_list)) {
    for (species_tissue in species_tissue_list[[species]]) {
        if (species_tissue != "") {
            tissue_dot <- paste0(".", species_tissue)
            tissue_us <- paste0("_", species_tissue)
        } else {
            tissue_dot <- ""
            tissue_us <- ""
        }

        output_5ofTICE <- readRDS(file = paste0("processed_data_w_polyA/5ofTICE_features/",
            species, tissue_dot, ".5ofTICE_features.rds"))[[2]]
        output_uTICE <- readRDS(file = paste0("processed_data_w_polyA/uTICE_features/",
            species, tissue_dot, ".uTICE_features.rds"))[[2]]
        output_dTICE <- readRDS(file = paste0("processed_data_w_polyA/dTICE_features/",
            species, tissue_dot, ".dTICE_features.rds"))[[2]]

        write.csv(output_5ofTICE, file = paste0("tables/5ofTICE_features/",
            species, tissue_dot, ".5ofTICE_features.csv"), quote = F)
        write.csv(output_uTICE, file = paste0("tables/uTICE_features/",
            species, tissue_dot, ".uTICE_features.csv"), quote = F)
        write.csv(output_dTICE, file = paste0("tables/dTICE_features/",
            species, tissue_dot, ".dTICE_features.csv"), quote = F)
    }
}
```

#### Pearson correlation(s) between the amino acid frequency feature(s) or codon frequency feature(s) and log10(TR)

##### save the values to excel files (the following code only needs to be run once)

```
# calculate the Pearson correlation between each of the 20
# amino acid frequencies and the 61 codon frequencies in the
# CDS (don't divide every feature into multiple groups) and
# log10(TR)
calc_aa_codon_features.logTR.cors <- function(species, tissue_dot) {
    ## log10.TR
    log10.TR <- get(paste0(species, tissue_dot, ".log10.TR"))
    ## count the frequency of 64 codons (including 3 stop codons)
    ## in every gene
    CDS_seqs <- get(paste0(species, tissue_dot, ".CDS_seqs"))
    codon_counts <- t(sapply(CDS_seqs, FUN = function(seq) {
        sst <- strsplit(seq, "")[[1]]
        n <- length(sst)
        out <- paste0(sst[seq(from = 1, to = n, by = 3)], sst[seq(from = 2,
            to = n, by = 3)], sst[seq(from = 3, to = n, by = 3)])
        table(out)[tri_nts]
    }))
    codon_counts[is.na(codon_counts)] <- 0
    rownames(codon_counts) <- names(CDS_seqs)
    colnames(codon_counts) <- tri_nts

    ## count the frequency of 20 amino acids (AAs) in every gene
    aa_counts <- sapply(aas, FUN = function(aa) {
        idx <- which(tri_nts_to_aas == aa)
        if (length(idx) == 1) {
            codon_counts[, idx]
        } else {
            rowSums(codon_counts[, idx])
        }
    })

    aa_features <- aa_counts/rowSums(aa_counts)
    aa_cors <- apply(aa_features, 2, FUN = function(x) cor(log10.TR,
        x, use = "complete.obs"))
    names(aa_cors) <- colnames(aa_features)

    codon_features <- codon_counts/rowSums(codon_counts)
    codon_cors <- apply(codon_features, 2, FUN = function(x) cor(log10.TR,
        x, use = "complete.obs"))
    names(codon_cors) <- colnames(codon_features)

    return(list(aa_cors = aa_cors, codon_cors = codon_cors))
}


aa_features.logTR.cors <- NULL
rownames_aa_features.logTR.cors <- NULL
```

```
codon_features.logTR.cors <- NULL
rownames_codon_features.logTR.cors <- NULL

for (species in names(species_tissue_list)) {
    for (i in 1:length(species_tissue_list[[species]])) {
        species_tissue <- species_tissue_list[[species]][i]
        if (species_tissue != "") {
            tissue_dot <- paste0(".", species_tissue)
            tissue_us <- paste0("_", species_tissue)
        } else {
            tissue_dot <- ""
            tissue_us <- ""
        }

        results <- calc_aa_codon_features.logTR.cors(species,
            tissue_dot)

        aa_features.logTR.cors <- rbind(aa_features.logTR.cors,
            results$aa_cors)
        rownames_aa_features.logTR.cors <- c(rownames_aa_features.logTR.cors,
            paste0(species, tissue_dot))

        codon_features.logTR.cors <- rbind(codon_features.logTR.cors,
            results$codon_cors)
        rownames_codon_features.logTR.cors <- c(rownames_codon_features.logTR.cors,
            paste0(species, tissue_dot))
    }
}

rownames(aa_features.logTR.cors) <- rownames_aa_features.logTR.cors
rownames(codon_features.logTR.cors) <- rownames_codon_features.logTR.cors

write.csv(aa_features.logTR.cors, file = "tables/aa_features.logTR.cors.csv",
    quote = F)
write.csv(codon_features.logTR.cors, file = "tables/codon_features.logTR.cors.csv",
    quote = F)
```

##### plot the results (the following code only needs to be run once)

```
aa_features.logTR.cors <- read.csv(file = "tables/aa_features.logTR.cors.csv")
colnames(aa_features.logTR.cors)[1] <- "species.tissue"

codon_features.logTR.cors <- read.csv(file = "tables/codon_features.logTR.cors.csv")
colnames(codon_features.logTR.cors)[1] <- "species.tissue"

aa_features.logTR.cors.for_plot <- setNames(melt(aa_features.logTR.cors),
    c("species.tissue", "aa.freq", "cor"))
aa_features.logTR.cors.for_plot$species.tissue <- factor(aa_features.logTR.cors.for_plot$species.tissue
    levels = as.character(aa_features.logTR.cors$species.tissue))

codon_features.logTR.cors.for_plot <- setNames(melt(codon_features.logTR.cors),
```

```
    c("species.tissue", "codon.freq", "cor"))
codon_features.logTR.cors.for_plot$species.tissue <- factor(codon_features.logTR.cors.for_plot$species.[illegible]
    levels = as.character(codon_features.logTR.cors$species.tissue))

pdf("tables/aa_features.logTR.cors.pdf", width = 10, height = 5)
p <- ggplot(aa_features.logTR.cors.for_plot, aes(x = aa.freq,
    y = species.tissue)) + geom_tile(aes(fill = cor)) + scale_fill_viridis() +
    ggtitle("Pearson cor with log10(TR)")
print(p)
dev.off()

pdf("tables/codon_features.logTR.cors.pdf", width = 10, height = 5)
p <- ggplot(codon_features.logTR.cors.for_plot, aes(x = codon.freq,
    y = species.tissue)) + geom_tile(aes(fill = cor)) + scale_fill_viridis() +
    theme(axis.text.x = element_text(size = 6, angle = 45, vjust = 1,
        hjust = 1)) + ggtitle("Pearson cor with log10(TR)")
print(p)
dev.off()
```

##### R2 of the amino acid frequency feature(s) or codon frequency feature(s) in a multi-variate multi-part linear model

```
for (species in names(species_tissue_list)) {
    for (species_tissue in species_tissue_list[[species]]) {
        if (species_tissue != "") {
            tissue_dot <- paste0(".", species_tissue)
            tissue_us <- paste0("_", species_tissue)
        } else {
            tissue_dot <- ""
            tissue_us <- ""
        }

        aa_features <- readRDS(paste0("processed_data_w_polyA/aa_features/",
            species, tissue_dot, ".aa_features.rds"))[[1]]
        cat(paste0(species, tissue_dot, " R2 of aa features: "))
        cat(summary(lm(get(paste0(species, tissue_dot, ".log10.TR")) ~
            as.matrix(aa_features)))$r.squared)
        cat("\n")

        codon_features <- readRDS(paste0("processed_data_w_polyA/codon_features/",
            species, tissue_dot, ".codon_features.rds"))[[1]]
        cat(paste0(species, tissue_dot, " R2 of codon features: "))
        cat(summary(lm(get(paste0(species, tissue_dot, ".log10.TR")) ~
            as.matrix(codon_features)))$r.squared)
        cat("\n")

        cat(paste0(species, tissue_dot, " R2 of aa + codon features: "))
        cat(summary(lm(get(paste0(species, tissue_dot, ".log10.TR")) ~
            as.matrix(cbind(aa_features, codon_features))))$r.squared)
        cat("\n\n")
    }
}
```

```
## sc R2 of aa features: 0.4049897
## sc R2 of codon features: 0.5955437
## sc R2 of aa + codon features: 0.5955437
##
## sp R2 of aa features: 0.2897104
## sp R2 of codon features: 0.4167586
## sp R2 of aa + codon features: 0.4167586
##
## sp.alt R2 of aa features: 0.3883752
## sp.alt R2 of codon features: 0.520032
## sp.alt R2 of aa + codon features: 0.520032
##
## at.leaf R2 of aa features: 0.1292097
## at.leaf R2 of codon features: 0.2252499
## at.leaf R2 of aa + codon features: 0.2252499
##
## at.root R2 of aa features: 0.1678659
## at.root R2 of codon features: 0.238489
## at.root R2 of aa + codon features: 0.238489
##
## at.shoot R2 of aa features: 0.1617223
## at.shoot R2 of codon features: 0.2257369
## at.shoot R2 of aa + codon features: 0.2257369
##
## mm.nih3t3 R2 of aa features: 0.1005773
## mm.nih3t3 R2 of codon features: 0.1455983
## mm.nih3t3 R2 of aa + codon features: 0.1455983
##
## mm.liver R2 of aa features: 0.1033705
## mm.liver R2 of codon features: 0.1808625
## mm.liver R2 of aa + codon features: 0.1808625
##
## mm.kidney R2 of aa features: 0.1360198
## mm.kidney R2 of codon features: 0.1673889
## mm.kidney R2 of aa + codon features: 0.1673889
##
## hs.hela2 R2 of aa features: 0.1605634
## hs.hela2 R2 of codon features: 0.2006944
## hs.hela2 R2 of aa + codon features: 0.2006944
```

### Generate tri-nucleotide frequencies in 5'ofTICE, uTICE, dTICE, and 3'ofTICE regions for all genes in each tissue of each species

```
for (species in names(species_tissue_list)) {
    for (species_tissue in species_tissue_list[[species]]) {
        if (species_tissue != "") {
            species.UTR_seqs <- get(paste(species, species_tissue,
                "UTR_seqs", sep = "."))
            species.CDS_seqs <- get(paste(species, species_tissue,
                "CDS_seqs", sep = "."))
            connector <- paste0(".", species_tissue)
```

```
} else {
    species.UTR_seqs <- get(paste(species, "UTR_seqs",
        sep = "."))
    species.CDS_seqs <- get(paste(species, "CDS_seqs",
        sep = "."))
    connector <- ""
}

if (species != "hs") {
    species.uTICE_len <- species_uTICE_len_list[[species]]
    species.dTICE_len <- species_dTICE_len_list[[species]]
} else {
    species.uTICE_len <- species_uTICE_len_list[[species]][[species_tissue]]
    species.dTICE_len <- species_dTICE_len_list[[species]][[species_tissue]]
}

# 5' of TICE
species.UTR_seqs_5ofTICE <- substr(x = species.UTR_seqs,
    start = 1, stop = nchar(species.UTR_seqs) - species.uTICE_len)
species.UTR_tri_counts_5ofTICE <- sapply(tri_nts, FUN = function(nuc) {
    stri_count_regex(species.UTR_seqs_5ofTICE, paste0("(?=",
        nuc, ")"))
})
rownames(species.UTR_tri_counts_5ofTICE) <- names(species.UTR_seqs_5ofTICE)
species.UTR_tri_freqs_5ofTICE <- species.UTR_tri_counts_5ofTICE/rowSums(species.UTR_tri_counts_[illegible]
species.UTR_tri_freqs_5ofTICE[is.na(species.UTR_tri_freqs_5ofTICE)] <- 0

saveRDS(species.UTR_tri_freqs_5ofTICE, file = paste0("processed_data_w_polyA/tri_freqs/",
    species, connector, ".UTR_tri_freqs_5ofTICE.rds"))

# uTICE
species.UTR_seqs_uTICE <- substr(x = species.UTR_seqs,
    start = nchar(species.UTR_seqs) - species.uTICE_len +
        1, stop = nchar(species.UTR_seqs))
species.UTR_tri_counts_uTICE <- sapply(tri_nts, FUN = function(nuc) {
    stri_count_regex(species.UTR_seqs_uTICE, paste0("(?=",
        nuc, ")"))
})
rownames(species.UTR_tri_counts_uTICE) <- names(species.UTR_seqs_uTICE)
species.UTR_tri_freqs_uTICE <- species.UTR_tri_counts_uTICE/rowSums(species.UTR_tri_counts_uTIC[illegible]
species.UTR_tri_freqs_uTICE[is.na(species.UTR_tri_freqs_uTICE)] <- 0

saveRDS(species.UTR_tri_freqs_uTICE, file = paste0("processed_data_w_polyA/tri_freqs/",
    species, connector, ".UTR_tri_freqs_uTICE.rds"))

# dTICE
species.CDS_seqs_dTICE <- substr(x = species.CDS_seqs,
    start = 4, stop = species.dTICE_len)
species.CDS_tri_counts_dTICE <- sapply(tri_nts, FUN = function(nuc) {
    stri_count_regex(species.CDS_seqs_dTICE, paste0("(?=",
        nuc, ")"))
})
rownames(species.CDS_tri_counts_dTICE) <- names(species.CDS_seqs_dTICE)
```

```
        species.CDS_tri_freqs_dTICE <- species.CDS_tri_counts_dTICE/rowSums(species.CDS_tri_counts_dTIC
        species.CDS_tri_freqs_dTICE[is.na(species.CDS_tri_freqs_dTICE)] <- 0

        saveRDS(species.CDS_tri_freqs_dTICE, file = paste0("processed_data_w_polyA/tri_freqs/",
            species, connector, ".CDS_tri_freqs_dTICE.rds"))

        # 3' of TICE
        species.CDS_seqs_3ofTICE <- substr(x = species.CDS_seqs,
            start = species.dTICE_len + 1, stop = nchar(species.CDS_seqs))
        species.CDS_tri_counts_3ofTICE <- sapply(tri_nts, FUN = function(nuc) {
            stri_count_regex(species.CDS_seqs_3ofTICE, paste0("(?=",
                nuc, ")"))
        })
        rownames(species.CDS_tri_counts_3ofTICE) <- names(species.CDS_seqs_3ofTICE)
        species.CDS_tri_freqs_3ofTICE <- species.CDS_tri_counts_3ofTICE/rowSums(species.CDS_tri_counts_
        species.CDS_tri_freqs_3ofTICE[is.na(species.CDS_tri_freqs_3ofTICE)] <- 0

        saveRDS(species.CDS_tri_freqs_3ofTICE, file = paste0("processed_data_w_polyA/tri_freqs/",
            species, connector, ".CDS_tri_freqs_3ofTICE.rds"))
    }
}
```

#### information needed to plot a trinucleotide ratio heatmap

```
## trinucleotide table
tri_nts_table <- rbind(matrix(tri_nts[1:16], nrow = 4), matrix(tri_nts[17:32],
    nrow = 4), matrix(tri_nts[33:48], nrow = 4), matrix(tri_nts[49:64],
    nrow = 4))

## creates a own color palette from blue to red
my_palette <- colorRampPalette(c(rgb(103, 169, 207, maxColorValue = 255),
    rgb(247, 247, 247, maxColorValue = 255), rgb(239, 138, 98,
        maxColorValue = 255)))(n = 299)
```

#### Find the minimum, the 2nd minimum, and the 3rd minimum folding energy windows in the 5'UTR for every gene; make the windows non-overlapping

```
# find the optimal window length based on the R2 of the
# minimum folding energy and log10(TR)
for (species in names(species_tissue_list)) {
    for (species_tissue in species_tissue_list[[species]]) {
        if (species_tissue != "") {
            tissue_dot <- paste0(".", species_tissue)
            tissue_us <- paste0("_", species_tissue)
        } else {
            tissue_dot <- ""
            tissue_us <- ""
```

```
    }
    if (species == "sp") {
        fold_energy_features_all <- readRDS(paste0("processed_data_w_polyA/folding_energy_features_
            species, tissue_us, "_UTR_folding_energy_features_selected_trun.rds"))[[4]]
    } else {
        fold_energy_features_all <- readRDS(paste0("processed_data_w_polyA/folding_energy_features_
            species, tissue_us, "_UTR_folding_energy_features_selected.rds"))[[4]]
    }

    R2s_min_fold_energy_features <- sapply(window_lens, FUN = function(window_len) {
        min_fold_energy_features <- fold_energy_features_all[,
            c(which(grepl(paste0("^min", "_", window_len,
              "$"), colnames(fold_energy_features_all))),
              which(grepl(paste0("^min", "_", window_len,
                "_"), colnames(fold_energy_features_all))),
              which(grepl(paste0("^Intercept", "_", window_len),
                colnames(fold_energy_features_all))))]
        summary(lm(get(paste0(species, tissue_dot, ".log10.TR")) ~
            as.matrix(min_fold_energy_features)))$r.squared
    })

    min_fold_energy_window_len <- as.numeric(window_lens[which.max(R2s_min_fold_energy_features)])

    min_fold_energy_features <- fold_energy_features_all[,
        c(which(grepl(paste0("^min", "_", min_fold_energy_window_len,
            "$"), colnames(fold_energy_features_all))), which(grepl(paste0("^min",
            "_", min_fold_energy_window_len, "_"), colnames(fold_energy_features_all))),
            which(grepl(paste0("^Intercept", "_", min_fold_energy_window_len),
              colnames(fold_energy_features_all))))]

    UTR_folding_energy_all <- readRDS(file = paste0("processed_data_w_polyA/var_win_fold_UTR5_CDSsh
        species, tissue_us, "_5UTR_CDS_folding_energy_",
        min_fold_energy_window_len, ".rds"))
    species_genes <- get(paste0(species, tissue_dot, ".genes"))
    UTR_folding_energy_all <- UTR_folding_energy_all$UTR_folding_energy_all[species_genes]

    if (species != "sp") {
        min_2ndmin_3rdmin_fold_energy_features <- t(sapply(UTR_folding_energy_all,
            FUN = function(x) {
              if (is.null(x)) {
                return(rep(NA, 3))
              } else {
                n <- length(x)
                x1 <- min(x)
                x1_loc <- which.min(x)
                rm_idx <- max(1, x1_loc - min_fold_energy_window_len +
                  1):min(n, x1_loc + min_fold_energy_window_len)
                x_kept <- x[-rm_idx]
                if (length(x_kept) > 0) {
                  x2 <- min(x_kept)
                  x2_loc <- which(x == x2)
                  rm_idx <- c(rm_idx, max(1, x2_loc - min_fold_energy_window_len +
                    1):min(n, x2_loc + min_fold_energy_window_len))
```

```
                x_kept <- x[-rm_idx]
                if (length(x_kept) > 0) {
                  x3 <- min(x_kept)
                } else {
                  x3 <- NA
                }
              } else {
                x2 <- NA
                x3 <- NA
              }
            }
            return(c(x1, x2, x3))
          }))
    } else {
        # sp, needs truncation to remove long windows that extend
        # beyond +30
        trun_pos <- as.numeric(window_len) - 30 + 1  # the position (-trun_pos) relative to AUG; wi[illegible]
        UTR_folding_energy_all_trun <- lapply(UTR_folding_energy_all,
            FUN = function(x) {
              if (trun_pos > length(x)) {
                x <- NULL
              } else if (trun_pos > 0) {
                x <- rev(rev(x)[trun_pos:length(x)])
              }
              return(x)
            })
        min_2ndmin_3rdmin_fold_energy_features <- t(sapply(UTR_folding_energy_all_trun,
            FUN = function(x) {
              if (is.null(x)) {
                return(rep(NA, 3))
              } else {
                n <- length(x)
                x1 <- min(x)
                x1_loc <- which.min(x)
                rm_idx <- max(1, x1_loc - min_fold_energy_window_len +
                  1):min(n, x1_loc + min_fold_energy_window_len)
                x_kept <- x[-rm_idx]
                if (length(x_kept) > 0) {
                  x2 <- min(x_kept)
                  x2_loc <- which(x == x2)
                  rm_idx <- c(rm_idx, max(1, x2_loc - min_fold_energy_window_len +
                    1):min(n, x2_loc + min_fold_energy_window_len))
                  x_kept <- x[-rm_idx]
                  if (length(x_kept) > 0) {
                    x3 <- min(x_kept)
                  } else {
                    x3 <- NA
                  }
                } else {
                  x2 <- NA
                  x3 <- NA
                }
              }
```

```
                return(c(x1, x2, x3))
            }))
        }

        if (species != "hs") {
            min_fold_energy_features.div <- div_original_features(min_2ndmin_3rdmin_fold_energy_features
                1], paste("min_fold_energy", min_fold_energy_window_len,
                sep = "_"), get(paste0(species, tissue_dot, ".log10.UTR_lens")),
                get(paste0(species, ".len_breaks")))

            second_min_fold_energy_features.div <- div_original_features(min_2ndmin_3rdmin_fold_energy_
                2], paste("2ndmin_fold_energy", min_fold_energy_window_len,
                sep = "_"), get(paste0(species, tissue_dot, ".log10.UTR_lens")),
                get(paste0(species, ".len_breaks")))

            third_min_fold_energy_features.div <- div_original_features(min_2ndmin_3rdmin_fold_energy_f
                3], paste("3rdmin_fold_energy", min_fold_energy_window_len,
                sep = "_"), get(paste0(species, tissue_dot, ".log10.UTR_lens")),
                get(paste0(species, ".len_breaks")))
        } else {
            min_fold_energy_features.div <- div_original_features(min_2ndmin_3rdmin_fold_energy_features
                1], paste("min_fold_energy", min_fold_energy_window_len,
                sep = "_"), get(paste0(species, tissue_dot, ".log10.UTR_lens")),
                get(paste0(species, tissue_dot, ".len_breaks")))

            second_min_fold_energy_features.div <- div_original_features(min_2ndmin_3rdmin_fold_energy_
                2], paste("2ndmin_fold_energy", min_fold_energy_window_len,
                sep = "_"), get(paste0(species, tissue_dot, ".log10.UTR_lens")),
                get(paste0(species, tissue_dot, ".len_breaks")))

            third_min_fold_energy_features.div <- div_original_features(min_2ndmin_3rdmin_fold_energy_f
                3], paste("3rdmin_fold_energy", min_fold_energy_window_len,
                sep = "_"), get(paste0(species, tissue_dot, ".log10.UTR_lens")),
                get(paste0(species, tissue_dot, ".len_breaks")))
        }

        saveRDS(min_fold_energy_features.div, file = paste0("processed_data_w_polyA/min_2ndmin_3rdmin_f
            species, tissue_us, "_min_fold_energy_features.div.rds"))

        saveRDS(second_min_fold_energy_features.div, file = paste0("processed_data_w_polyA/min_2ndmin_3
            species, tissue_us, "_second_min_fold_energy_features.div.rds"))

        saveRDS(third_min_fold_energy_features.div, file = paste0("processed_data_w_polyA/min_2ndmin_3r
            species, tissue_us, "_third_min_fold_energy_features.div.rds"))

        saveRDS(min_2ndmin_3rdmin_fold_energy_features, file = paste0("processed_data_w_polyA/min_2ndmi
            species, tissue_us, "_min_2ndmin_3rdmin_fold_energy_features"))
    }
}
```

#### Figure 1. Distributions of log10(TR), log10(5'UTR length), and log10(CDS length)

```
# a list of species and colors
species_colors <- list(sc = "#EB9D41", sp = "#94241D", at = "#22549C",
    mm = "#2F6E42", hs = "#CEC448")

# number of species
m <- length(species_colors)

# number of genes in each exemplar tissue of each species
num_genes <- NULL
# distributions of the log10(TR), log10(UTR_lens) and
# log10(CDS_lens) values of each exemplar tissue in each
# species after aligning their median
center.log10.TR <- NULL
for (species in names(species_tissue_list)) {
    for (i in 1:length(species_exemplar_tissue_list[[species]])) {
        species_tissue <- species_exemplar_tissue_list[[species]][i]
        if (species_tissue != "") {
            tissue_dot <- paste0(".", species_tissue)
        } else {
            tissue_dot <- ""
        }
        num_genes <- c(num_genes, sum(!is.na(get(paste0(species,
            tissue_dot, ".log10.TR")))))
        center.log10.TR <- c(center.log10.TR, median(get(paste0(species,
            tissue_dot, ".log10.TR"))))
    }
}
center.log10.TR <- mean(center.log10.TR)
den.log10.TR <- vector("list", 0)
ymax.log10.TR <- NULL
xmin.log10.TR <- NULL
xmax.log10.TR <- NULL
den.log10.UTR_lens <- vector("list", 0)
ymax.log10.UTR_lens <- NULL
xmin.log10.UTR_lens <- NULL
xmax.log10.UTR_lens <- NULL
den.log10.CDS_lens <- vector("list", 0)
ymax.log10.CDS_lens <- NULL
xmin.log10.CDS_lens <- NULL
xmax.log10.CDS_lens <- NULL
for (species in names(species_tissue_list)) {
    for (i in 1:length(species_exemplar_tissue_list[[species]])) {
        species_tissue <- species_exemplar_tissue_list[[species]][i]
        if (species_tissue != "") {
            tissue_dot <- paste0(".", species_tissue)
        } else {
            tissue_dot <- ""
        }
        x <- get(paste0(species, tissue_dot, ".log10.TR"))
```

```
        den.log10.TR[[species]] <- density(x - median(x) + center.log10.TR)
        ymax.log10.TR <- c(ymax.log10.TR, max(den.log10.TR[[species]]$y))
        xmin.log10.TR <- c(xmin.log10.TR, min(den.log10.TR[[species]]$x))
        xmax.log10.TR <- c(xmax.log10.TR, max(den.log10.TR[[species]]$x))
        x <- log(nchar(get(paste0(species, tissue_dot, ".UTR_seqs"))) +
            1, 10)
        den.log10.UTR_lens[[species]] <- density(x)
        ymax.log10.UTR_lens <- c(ymax.log10.UTR_lens, max(den.log10.UTR_lens[[species]]$y))
        xmin.log10.UTR_lens <- c(xmin.log10.UTR_lens, min(den.log10.UTR_lens[[species]]$x))
        xmax.log10.UTR_lens <- c(xmax.log10.UTR_lens, max(den.log10.UTR_lens[[species]]$x))
        x <- log(nchar(get(paste0(species, tissue_dot, ".CDS_seqs"))) +
            1, 10)
        den.log10.CDS_lens[[species]] <- density(x)
        ymax.log10.CDS_lens <- c(ymax.log10.CDS_lens, max(den.log10.CDS_lens[[species]]$y))
        xmin.log10.CDS_lens <- c(xmin.log10.CDS_lens, min(den.log10.CDS_lens[[species]]$x))
        xmax.log10.CDS_lens <- c(xmax.log10.CDS_lens, max(den.log10.CDS_lens[[species]]$x))
    }
}
max_idx.log10.TR <- which(ymax.log10.TR == max(ymax.log10.TR))
max_idx.log10.UTR_lens <- which(ymax.log10.UTR_lens == max(ymax.log10.UTR_lens))
max_idx.log10.CDS_lens <- which(ymax.log10.CDS_lens == max(ymax.log10.CDS_lens))

# plot the distributions
plot(den.log10.TR[[max_idx.log10.TR]], lwd = 2, col = species_colors[[max_idx.log10.TR]],
    xlim = c(min(xmin.log10.TR), max(xmax.log10.TR)), ylim = c(0,
        max(ymax.log10.TR)), main = "Distributions of log10(TR) values")
for (i in 1:m) {
    if (i != max_idx.log10.TR) {
        lines(den.log10.TR[[i]], lwd = 2, col = species_colors[[i]])
    }
}
legend("topleft", paste(paste(names(species_exemplar_tissue_list),
    unlist(species_exemplar_tissue_list)), paste(num_genes, "genes"),
    sep = ": "), lwd = 2, col = unlist(species_colors), bty = "n")
```

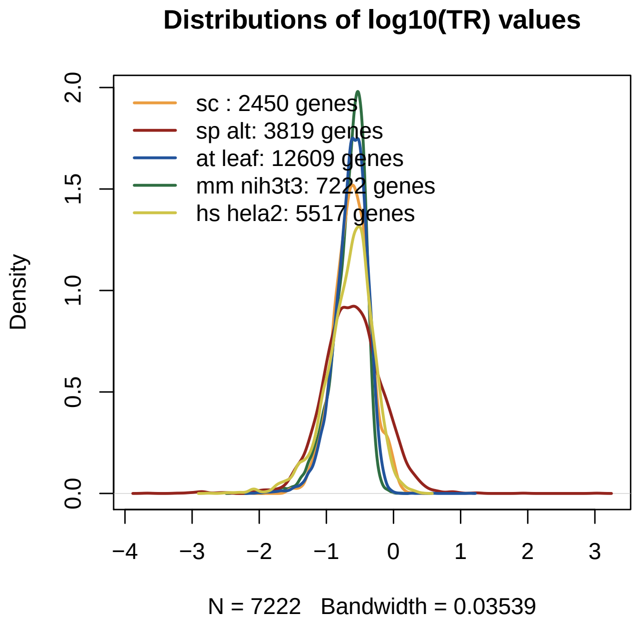


```
plot(den.log10.UTR_lens[[max_idx.log10.UTR_lens]], lwd = 2, col = species_colors[[max_idx.log10.UTR_lens
    xlim = c(min(c(xmin.log10.UTR_lens, xmin.log10.CDS_lens)),
        max(c(xmax.log10.UTR_lens, xmax.log10.CDS_lens))), ylim = c(0,
        max(ymax.log10.UTR_lens)), main = "Distributions of log10(UTR len + 1) values",
    axes = F, xlab = "length (bp)")
axis(1, log(c(0, 10, 25, 50, 100, 250, 500, 1000, 2500, 5000,
    10000) + 1, 10), paste(c(0, 10, 25, 50, 100, 250, 500, 1000,
    2500, 5000, 10000)), las = 2)
axis(2)
box()
for (i in 1:m) {
    if (i != max_idx.log10.UTR_lens) {
        lines(den.log10.UTR_lens[[i]], lwd = 2, col = species_colors[[i]])
    }
}
legend("topleft", paste(paste(names(species_exemplar_tissue_list),
    unlist(species_exemplar_tissue_list)), paste(num_genes, "genes"),
    sep = ": "), lwd = 2, col = unlist(species_colors), bty = "n")
```

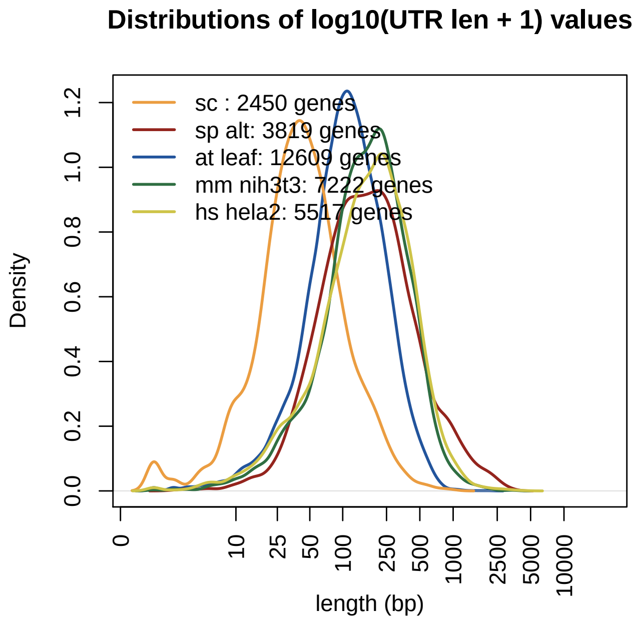


```
plot(den.log10.CDS_lens[[max_idx.log10.CDS_lens]], lwd = 2, col = species_colors[[max_idx.log10.CDS_lens
    xlim = c(min(c(xmin.log10.UTR_lens, xmin.log10.CDS_lens)),
        max(c(xmax.log10.UTR_lens, xmax.log10.CDS_lens))), ylim = c(0,
        max(ymax.log10.CDS_lens)), main = "Distributions of log10(CDS len + 1) values",
    axes = F, xlab = "length (bp)")
axis(1, log(c(0, 10, 25, 50, 100, 250, 500, 1000, 2500, 5000,
    10000) + 1, 10), paste(c(0, 10, 25, 50, 100, 250, 500, 1000,
    2500, 5000, 10000)), las = 2)
axis(2)
box()
for (i in 1:m) {
    if (i != max_idx.log10.CDS_lens) {
        lines(den.log10.CDS_lens[[i]], lwd = 2, col = species_colors[[i]])
    }
}
legend("topleft", paste(paste(names(species_exemplar_tissue_list),
    unlist(species_exemplar_tissue_list)), paste(num_genes, "genes"),
    sep = ": "), lwd = 2, col = unlist(species_colors), bty = "n")
```

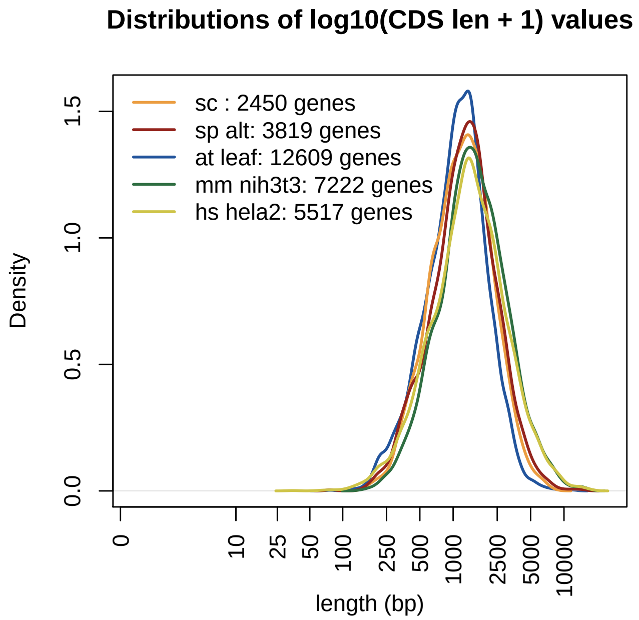


#### Figure 1 - figure supplement 1. Distributions of log10(TR) in all the tissues

```
# a list of species and colors
species_tissue_colors <- list(sc = "#EB9D41", sp = list("#94241D",
    alt = "#DA3A32"), at = list(leaf = "#22549C", root = "#3881B4",
    shoot = "#4CAAE7"), mm = list(nih3t3 = "#2F6E42", liver = "#5C9345",
    kidney = "#76B958"), hs = "#CEC448")

# number of species
m <- length(unlist(species_tissue_colors))

# number of genes in each tissue of each species
num_genes <- NULL
# distributions of the log10(TR) of each tissue in each
# species after aligning their median
center.log10.TR <- NULL
for (species in names(species_tissue_list)) {
    for (i in 1:length(species_tissue_list[[species]])) {
        species_tissue <- species_tissue_list[[species]][i]
        if (species_tissue != "") {
            tissue_dot <- paste0(".", species_tissue)
        } else {
            tissue_dot <- ""
        }
        num_genes <- c(num_genes, sum(!is.na(get(paste0(species,
            tissue_dot, ".log10.TR")))))
        center.log10.TR <- c(center.log10.TR, median(get(paste0(species,
            tissue_dot, ".log10.TR"))))
    }
}
```

```r
center.log10.TR <- mean(center.log10.TR)
den.log10.TR <- vector("list", 0)
ymax.log10.TR <- NULL
xmin.log10.TR <- NULL
xmax.log10.TR <- NULL
for (species in names(species_tissue_list)) {
    for (i in 1:length(species_tissue_list[[species]])) {
        species_tissue <- species_tissue_list[[species]][i]
        if (species_tissue != "") {
            tissue_dot <- paste0(".", species_tissue)
        } else {
            tissue_dot <- ""
        }
        name <- paste0(species, tissue_dot)
        x <- get(paste0(species, tissue_dot, ".log10.TR"))
        den.log10.TR[[name]] <- density(x - median(x) + center.log10.TR)
        ymax.log10.TR <- c(ymax.log10.TR, max(den.log10.TR[[name]]$y))
        xmin.log10.TR <- c(xmin.log10.TR, min(den.log10.TR[[name]]$x))
        xmax.log10.TR <- c(xmax.log10.TR, max(den.log10.TR[[name]]$x))
    }
}
max_idx.log10.TR <- which(ymax.log10.TR == max(ymax.log10.TR))

# plot the distributions
plot(den.log10.TR[[max_idx.log10.TR]], lwd = 2, col = unlist(species_tissue_colors)[max_idx.log10.TR],
    xlim = c(min(xmin.log10.TR), max(xmax.log10.TR)), ylim = c(0,
        max(ymax.log10.TR)), main = "Distributions of log10(TR) values")
for (i in 1:m) {
    if (i != max_idx.log10.TR) {
        lines(den.log10.TR[[i]], lwd = 2, col = unlist(species_tissue_colors)[i])
    }
}
legend("topleft", paste(names(unlist(species_tissue_colors)),
    paste(num_genes, "genes"), sep = ": "), lwd = 2, col = unlist(species_tissue_colors),
    bty = "n")
```

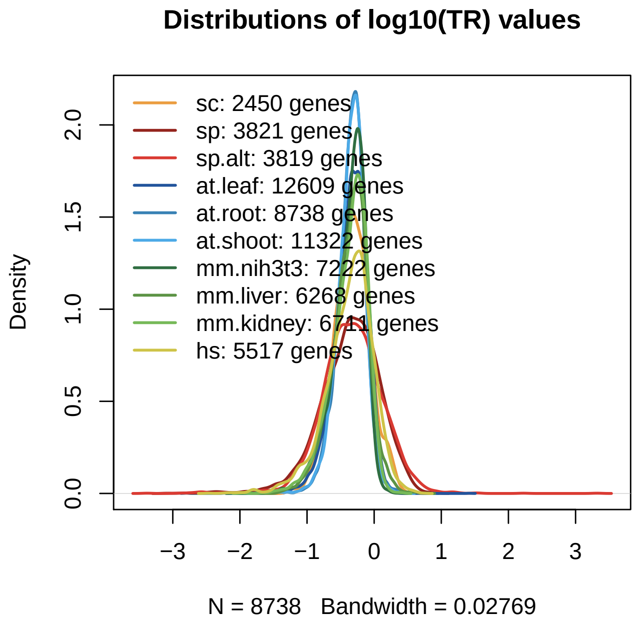


#### Figure 2 - figure supplement 1. Folding energy of the top and bottom 20 translated genes

##### the following code only needs to be run once

```
# a function to generate the top and bottom genes' folding
# energy values aligned at AUG
top_bottom_TR_genes_fold_engy_AUG_aligned <- function(n_left,
    n_right, top_genes, bottom_genes, UTR_folding_energy_all,
    CDS_folding_energy_all) {
    # n_left: the number of positions on the left of AUG n_right:
    # the number of positions on the right of AUG
    top_results <- sapply(top_genes, FUN = function(x) {
        c(rev(rev(UTR_folding_energy_all[[x]])[1:n_left]), CDS_folding_energy_all[[x]][1:n_right])
    })

    bottom_results <- sapply(bottom_genes, FUN = function(x) {
        c(rev(rev(UTR_folding_energy_all[[x]])[1:n_left]), CDS_folding_energy_all[[x]][1:n_right])
    })

    return(list(top = top_results, bottom = bottom_results))
}

n_left <- 250
n_right <- 50
for (species in names(species_tissue_list)) {
    for (species_tissue in species_tissue_list[[species]]) {
        if (species_tissue != "") {
            tissue_dot <- paste0(".", species_tissue)
            tissue_us <- paste0("_", species_tissue)
        } else {
```

```
            tissue_dot <- ""
            tissue_us <- ""
        }
        UTR_seqs <- get(paste0(species, tissue_dot, ".UTR_seqs"))
        log10.UTR_len <- log(nchar(UTR_seqs), 10)

        folding_energy_all <- readRDS(paste0("processed_data_w_polyA/35_win_fold_UTR5_CDSfull_all/",
            species, tissue_us, "_UTR5_full_CDS_window_35.rds"))
        UTR_folding_energy_all <- folding_energy_all$UTR_folding_energy_all
        CDS_folding_energy_all <- folding_energy_all$CDS_folding_energy_all
        rm(folding_energy_all)
        gc()

        log10.TR <- get(paste0(species, tissue_dot, ".log10.TR"))
        log10.tmp <- sort(log10.TR[log10.UTR_len <= log(n_left,
            10)])
        TR_top_20_genes <- names(rev(log10.tmp)[1:20])
        TR_bottom_20_genes <- names(log10.tmp[1:20])

        fold_engy_AUG_aligned <- top_bottom_TR_genes_fold_engy_AUG_aligned(n_left = n_left,
            n_right = n_right, top_genes = TR_top_20_genes, bottom_genes = TR_bottom_20_genes,
            UTR_folding_energy_all, CDS_folding_energy_all)
        TR_top_20_heatmap <- fold_engy_AUG_aligned$top
        TR_bottom_20_heatmap <- fold_engy_AUG_aligned$bottom
        rm(fold_engy_AUG_aligned)

        labels <- as.character(c((-n_left):(-1), 1:n_right))
        breakpoints <- c(seq(1, n_left, by = 10), n_left, n_left +
            1, rev(seq(n_left + n_right, n_left + 1, by = -10)))
        labels[-breakpoints] <- ""

        rownames(TR_top_20_heatmap) <- labels
        rownames(TR_bottom_20_heatmap) <- labels

        saveRDS(TR_top_20_heatmap, file = paste0("processed_data_w_polyA/TR_top_bottom_20_heatmaps/",
            species, tissue_dot, ".TR_top_20_heatmap.rds"))
        saveRDS(TR_bottom_20_heatmap, file = paste0("processed_data_w_polyA/TR_top_bottom_20_heatmaps/"
            species, tissue_dot, ".TR_bottom_20_heatmap.rds"))
    }
}
```

#### plot

```
n_left <- 250
n_right <- 50
breakpoints <- c(seq(1, n_left, by = 10), n_left, n_left + 1,
    rev(seq(n_left + n_right, n_left + 1, by = -10)))

for (species in names(species_tissue_list)) {
    for (species_tissue in species_tissue_list[[species]]) {
        if (species_tissue != "") {
            tissue_dot <- paste0(".", species_tissue)
```

```
            tissue_us <- paste0("_", species_tissue)
        } else {
            tissue_dot <- ""
            tissue_us <- ""
        }
        TR_top_20_heatmap <- readRDS(file = paste0("processed_data_w_polyA/TR_top_bottom_20_heatmaps/",
            species, tissue_dot, ".TR_top_20_heatmap.rds"))
        TR_bottom_20_heatmap <- readRDS(file = paste0("processed_data_w_polyA/TR_top_bottom_20_heatmaps
            species, tissue_dot, ".TR_bottom_20_heatmap.rds"))
        heatmap.range <- range(c(TR_top_20_heatmap, TR_bottom_20_heatmap),
            na.rm = T)
        print(levelplot(TR_top_20_heatmap, scales = list(x = list(at = breakpoints,
            rot = 90, cex = 0.3)), at = seq(heatmap.range[1],
            heatmap.range[2], length.out = 31), main = paste0(species,
            tissue_dot, " TR top 20 genes"), col.regions = heat.colors(120)))
        print(levelplot(TR_bottom_20_heatmap, scales = list(x = list(at = breakpoints,
            rot = 90, cex = 0.3)), at = seq(heatmap.range[1],
            heatmap.range[2], length.out = 31), main = paste0(species,
            tissue_dot, " TR bottom 20 genes"), col.regions = heat.colors(120)))
    }
}
```

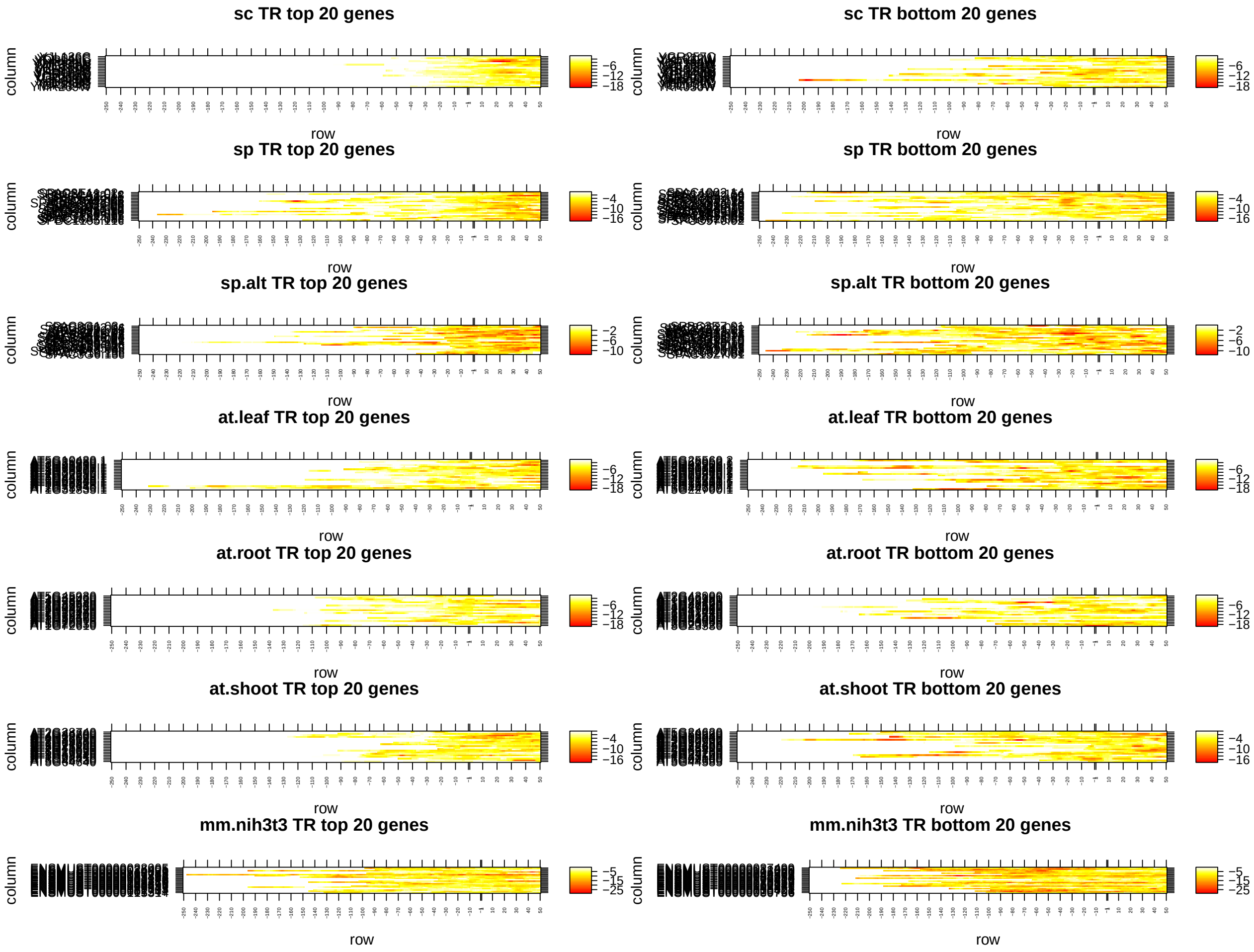

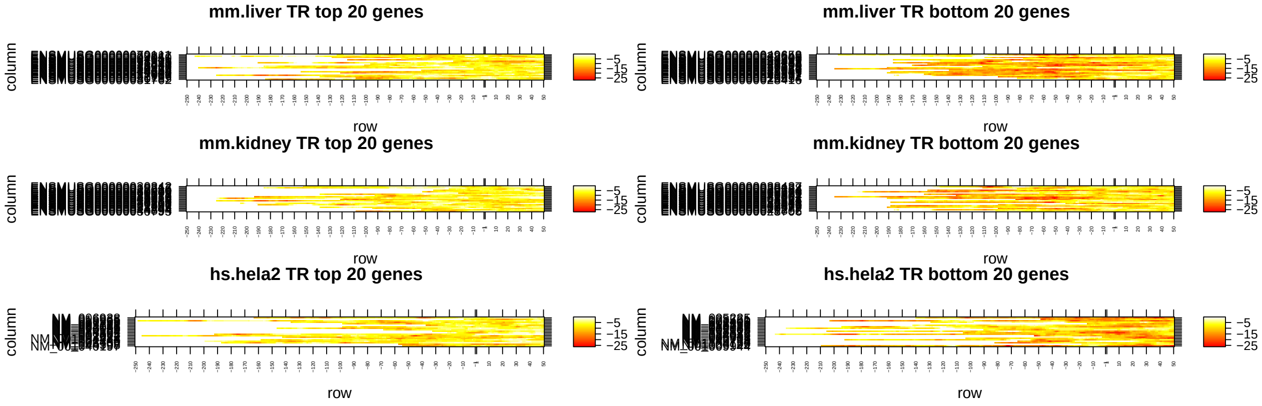


### Figure 3. R2s of RNA folding energy features

#### the following code only needs to be run once

```
for (species in names(species_tissue_list)) {
    for (species_tissue in species_tissue_list[[species]]) {
        if (species_tissue != "") {
            tissue_dot <- paste0(".", species_tissue)
            tissue_us <- paste0("_", species_tissue)
        } else {
            tissue_dot <- ""
            tissue_us <- ""
        }

        load(file = paste0("processed_data_w_polyA/", species,
            tissue_dot, ".Rdata"))
        fold_energy_features_selected <- readRDS(file = paste0("processed_data_w_polyA/folding_energy_f
            species, tissue_us, "_UTR_folding_energy_features_selected.rds"))

        all_features <- fold_energy_features_selected$all_features
        if (species != "sp") {
            selected_feature_R2 <- fold_energy_features_selected$selected_feature_R2
        } else {
            selected_feature_R2 <- readRDS(file = paste0("processed_data_w_polyA/folding_energy_feature
                species, tissue_us, "_UTR_folding_energy_features_selected_trun.rds"))$selected_feature_
        }


        feature_names <- c("5'most", "5'mostTICE", "-1window",
            "min", "10%", "25%", "75%", "90%", "max", "mean",
            "%<=20%", "sum<=5%", "sum<=10%", "sum<=20%", "%>90%",
            "%>80%", "sum>90%", "sum>80%", "model all-1", "whole",
            "model all", "RNA fold")
        names(feature_names) <- c("5' cap", "5'mostTICE", "-1",
            "min", "10%", "25%", "75%", "90%", "max", "mean",
            "%<=20%", "sum<=5%", "sum<=10%", "sum<=20%", "%>90%",
            "%>80%", "sum>90%", "sum>80%", "model all-1", "whole",
            "model all", "RNA fold")
```

```
feature_window_R2s <- matrix(NA, nrow = length(window_lens),
    ncol = length(feature_names))  # a matrix to store the results

rownames(feature_window_R2s) <- window_lens
colnames(feature_window_R2s) <- names(feature_names)

for (j in 1:length(feature_names)) {
    feature_name <- feature_names[j]
    if (!(feature_name %in% c("whole", "RNA fold"))) {
        # these two features do not vary with window lengths
        for (i in 1:length(window_lens)) {
          window_len <- window_lens[i]
          if (!(feature_name %in% c("model all-1", "model all"))) {
            # single features
            tmp1 <- which(colnames(all_features) == paste0(feature_name,
              "_", window_len))
            tmp2 <- which(grepl(paste0("^", feature_name,
              "_", window_len, "_"), colnames(all_features)))
            if (length(tmp1) > 0 & length(tmp2) == 0) {
              idx <- c(tmp1, which(grepl("Intercept",
                colnames(all_features))))
            } else if (length(tmp1) == 0 & length(tmp2) >
              0) {
              idx <- c(tmp2, which(grepl("Intercept",
                colnames(all_features))))
            } else {
              idx <- NULL
            }
          } else if (feature_name == "model all-1") {
            if (length(tmp1) > 0 & length(tmp2) == 0) {
              idx <- c(which(grepl(paste0("_", window_len,
                "$"), colnames(all_features))), which(grepl("Intercept",
                colnames(all_features))))
            } else if (length(tmp1) == 0 & length(tmp2) >
              0) {
              idx <- c(which(grepl(paste0("_", window_len,
                "_"), colnames(all_features))), which(grepl("Intercept",
                colnames(all_features))))
            } else {
              idx <- NULL
            }
          } else if (feature_name == "model all") {
            if (length(tmp1) > 0 & length(tmp2) == 0) {
              idx <- c(which(grepl(paste0("_", window_len,
                "$"), colnames(all_features))), which(grepl("Intercept",
                colnames(all_features))), which(grepl("whole",
                colnames(all_features))))
            } else if (length(tmp1) == 0 & length(tmp2) >
              0) {
              idx <- c(which(grepl(paste0("_", window_len,
                "_"), colnames(all_features))), which(grepl("Intercept",
                colnames(all_features))), which(grepl("whole",
                colnames(all_features))))
```

```
                    } else {
                      idx <- NULL
                    }
                  }
                  features <- all_features[, idx]
                  if (length(idx) > 0) {
                    feature_window_R2s[i, j] <- summary(lm(get(paste0(species,
                      tissue_dot, ".log10.TR")) ~ as.matrix(features)))$r.squared
                  } else {
                    feature_window_R2s[i, j] <- 0
                  }
                }
            } else if (feature_name == "whole") {
                idx <- c(which(grepl("whole", colnames(all_features))),
                  which(grepl("Intercept", colnames(all_features))))
                features <- all_features[, idx]
                feature_window_R2s[, j] <- summary(lm(get(paste0(species,
                  tissue_dot, ".log10.TR")) ~ as.matrix(features)))$r.squared
            } else {
                # feature == 'RNA fold' the BIC selected ones
                feature_window_R2s[, j] <- selected_feature_R2
            }
        }
        write.csv(t(feature_window_R2s), file = paste0("processed_data_w_polyA/folding_energy_feature_R2
            species, ".", species_tissue, ".csv"), quote = F)
        saveRDS(feature_window_R2s, file = paste0("processed_data_w_polyA/folding_energy_feature_R2s/",
            species, ".", species_tissue, ".rds"))
    }
}
```

#### plot

```
# correlation between each RNA folding energy and log10(TR)
for (species in names(species_tissue_list)) {
    for (species_tissue in species_tissue_list[[species]]) {
        feature_window_R2s <- readRDS(file = paste0("processed_data_w_polyA/folding_energy_feature_R2s/
            species, ".", species_tissue, ".rds"))
        # make a histogram
        window_cols <- c("#4F71BE", "#DF8344", "#A5A5A5", "#F6C243",
            "#6A9AD0", "#7EAB55", "#2D4474", "#944D20", "#636363",
            "#937524", "#355D8D", "#4B6733", "#718DCB", "#E59B65",
            "#B7B7B7", "#F7CF55", "#86ADD9", "#97C072", "#3C599C",
            "#C4672C", "#848484", "#000000")
        names(window_cols) <- c(window_lens, "NA")
        feature_window_R2s_for_plot <- setNames(melt(feature_window_R2s),
            c("Window_length", "Feature", "R2"))
        feature_window_R2s_for_plot[, 1] <- as.factor(feature_window_R2s_for_plot[,
            1])

        theme_set(theme_bw())
        print(ggplot(feature_window_R2s_for_plot, aes(x = Feature,
            y = R2, fill = Window_length)) + geom_bar(stat = "identity",
```

```
            position = position_dodge()) + scale_fill_manual(values = window_cols) +
            theme(axis.text.x = element_text(angle = 45, vjust = 1,
                hjust = 1)) + ggtitle(paste0(species, ".", species_tissue)))
    }
}
```

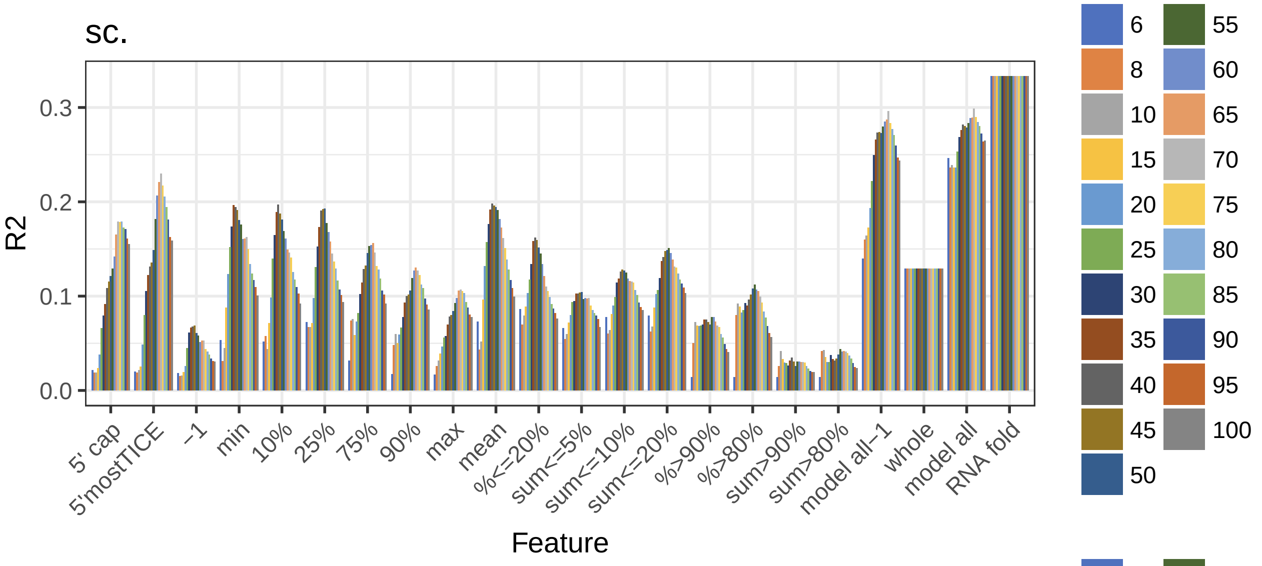


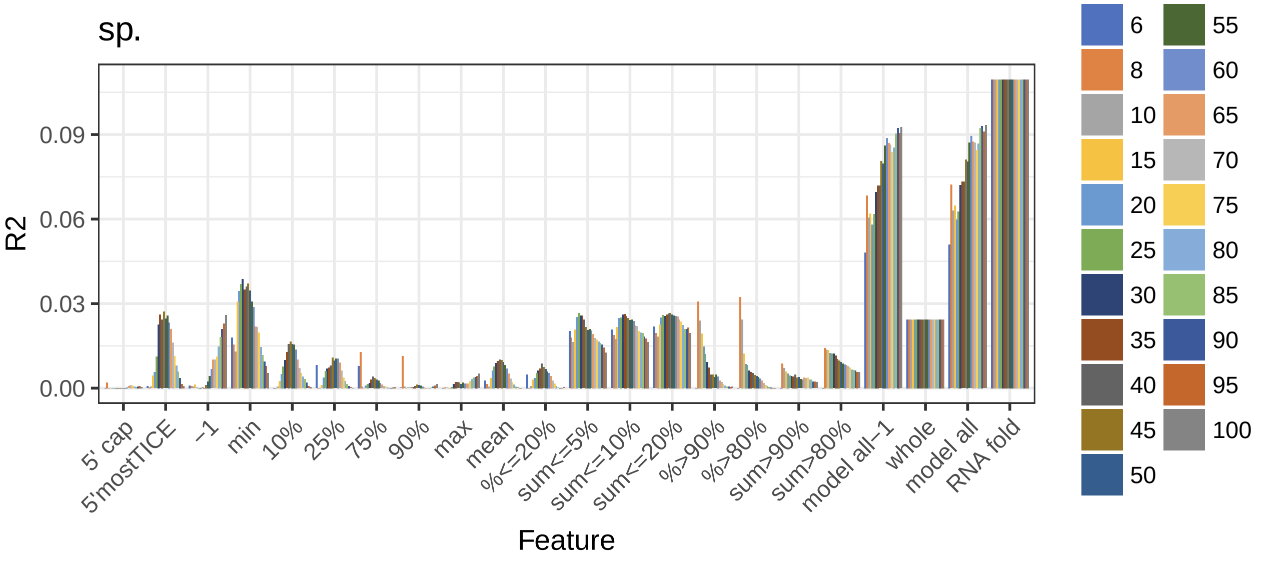

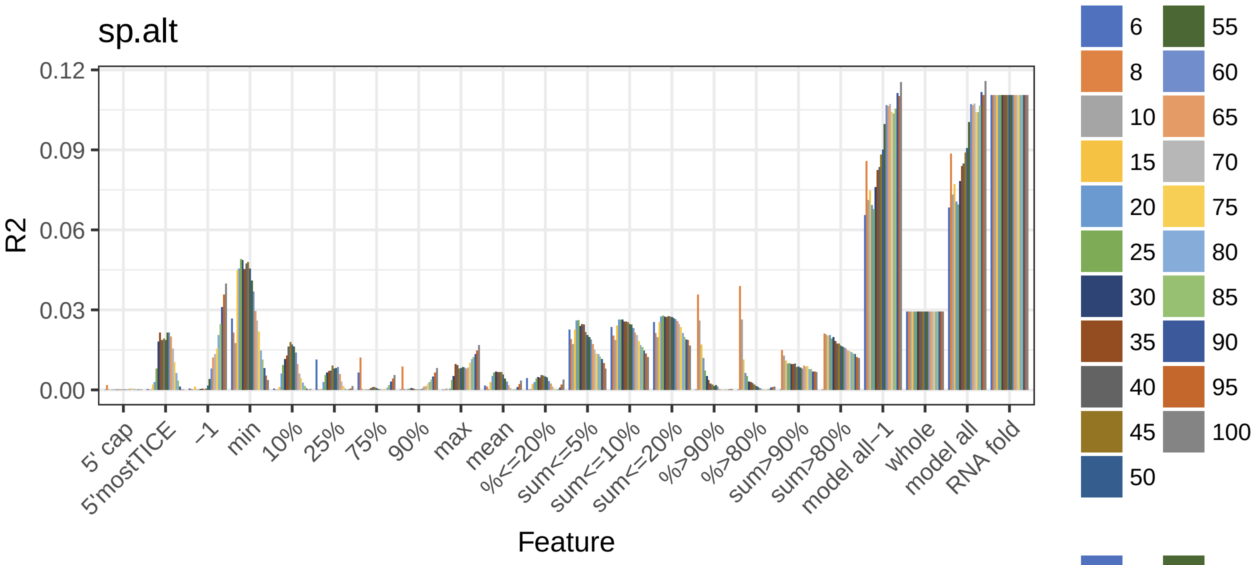
sp.alt
R2
0.12
0.09
0.06
0.03
0.00
5' cap
5'mostTICE
−1
min
10%
25%
75%
90%
max
mean
%<=20%
sum<=5%
sum<=10%
sum<=20%
%>90%
%>80%
sum>90%
sum>80%
model all−1
whole
model all
RNA fold
Feature
6
8
10
15
20
25
30
35
40
45
50
55
60
65
70
75
80
85
90
95
100

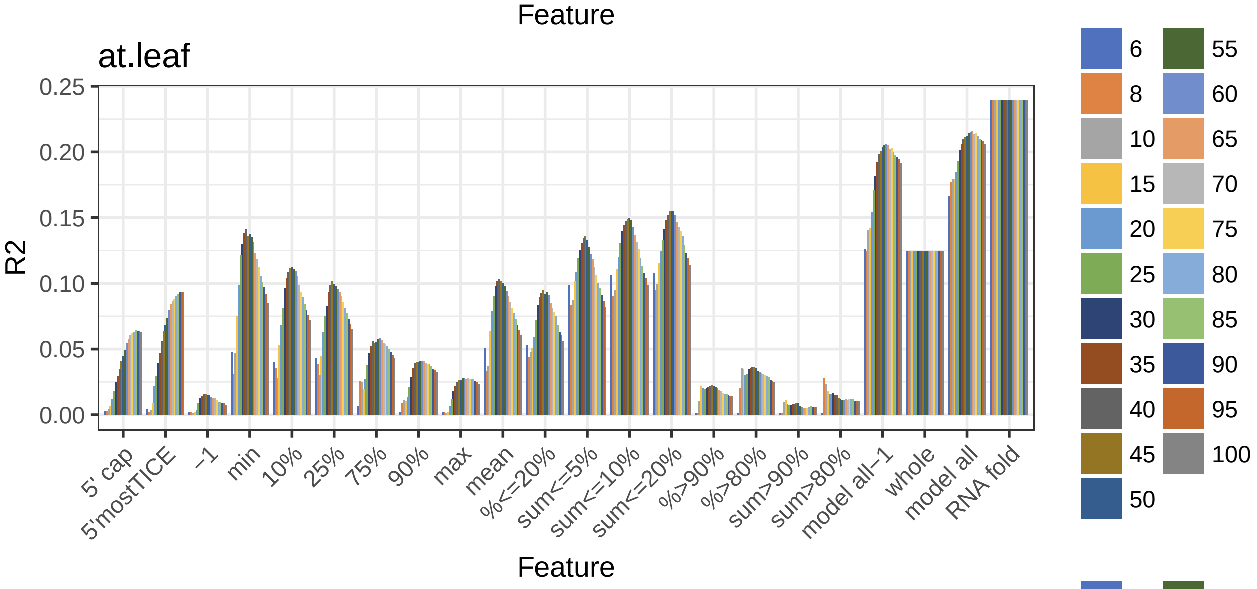
at.leaf
R2
0.25
0.20
0.15
0.10
0.05
0.00
5' cap
5'mostTICE
−1
min
10%
25%
75%
90%
max
mean
%<=20%
sum<=5%
sum<=10%
sum<=20%
%>90%
%>80%
sum>90%
sum>80%
model all−1
whole
model all
RNA fold
Feature
6
8
10
15
20
25
30
35
40
45
50
55
60
65
70
75
80
85
90
95
100

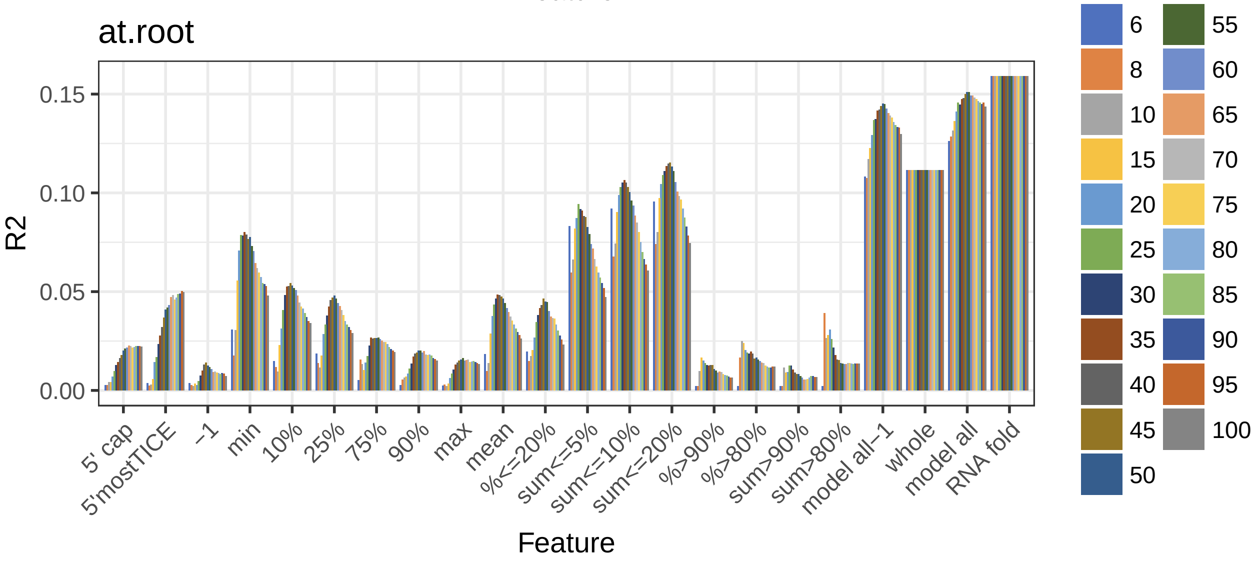
at.root
R2
0.15
0.10
0.05
0.00
5' cap
5'mostTICE
−1
min
10%
25%
75%
90%
max
mean
%<=20%
sum<=5%
sum<=10%
sum<=20%
%>90%
%>80%
sum>90%
sum>80%
model all−1
whole
model all
RNA fold
Feature
6
8
10
15
20
25
30
35
40
45
50
55
60
65
70
75
80
85
90
95
100

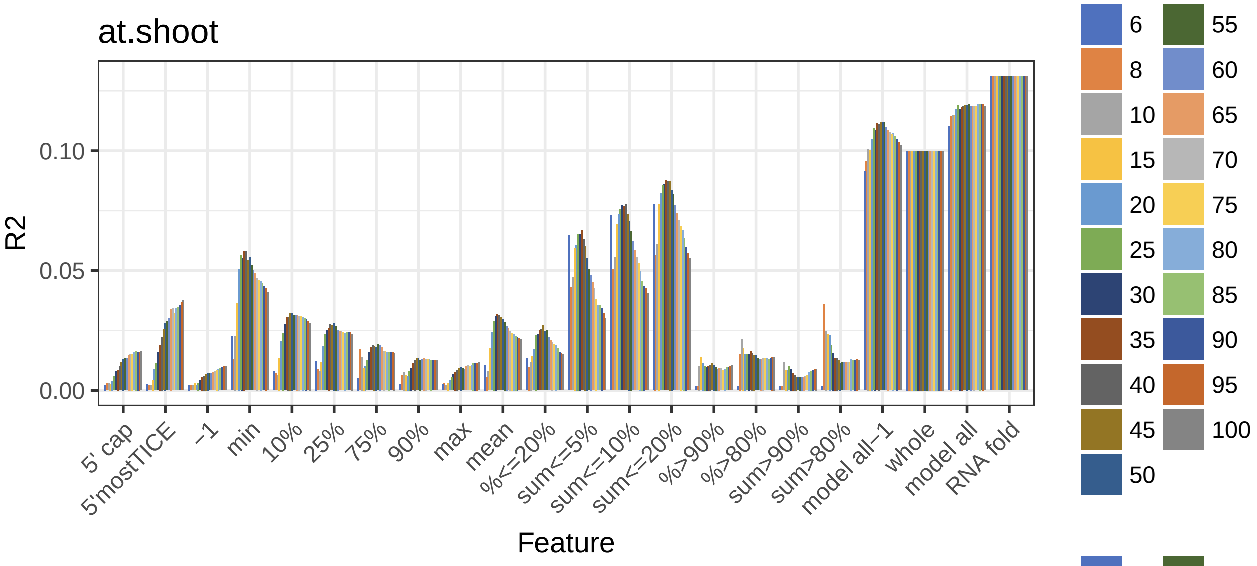
at.shoot
R2
0.00
0.05
0.10
5' cap
5'mostTICE
−1
min
10%
25%
75%
90%
max
mean
%<=20%
sum<=5%
sum<=10%
sum<=20%
%>90%
%>80%
sum>90%
sum>80%
model all−1
whole
model all
RNA fold
Feature
6
8
10
15
20
25
30
35
40
45
50
55
60
65
70
75
80
85
90
95
100

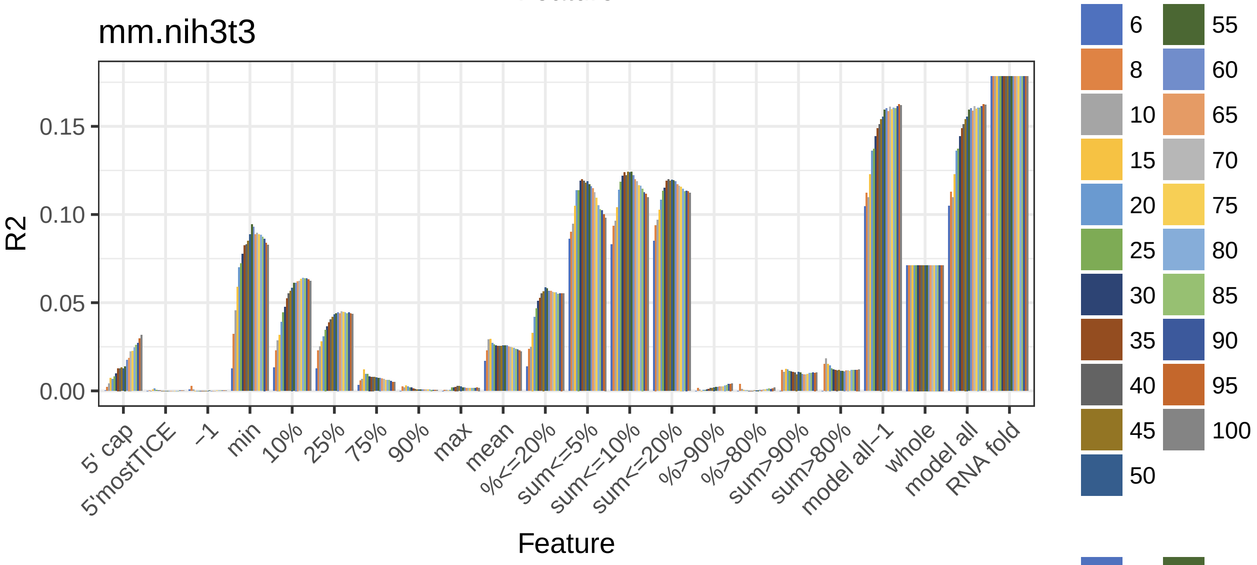
mm.nih3t3
R2
0.00
0.05
0.10
0.15
5' cap
5'mostTICE
−1
min
10%
25%
75%
90%
max
mean
%<=20%
sum<=5%
sum<=10%
sum<=20%
%>90%
%>80%
sum>90%
sum>80%
model all−1
whole
model all
RNA fold
Feature
6
8
10
15
20
25
30
35
40
45
50
55
60
65
70
75
80
85
90
95
100

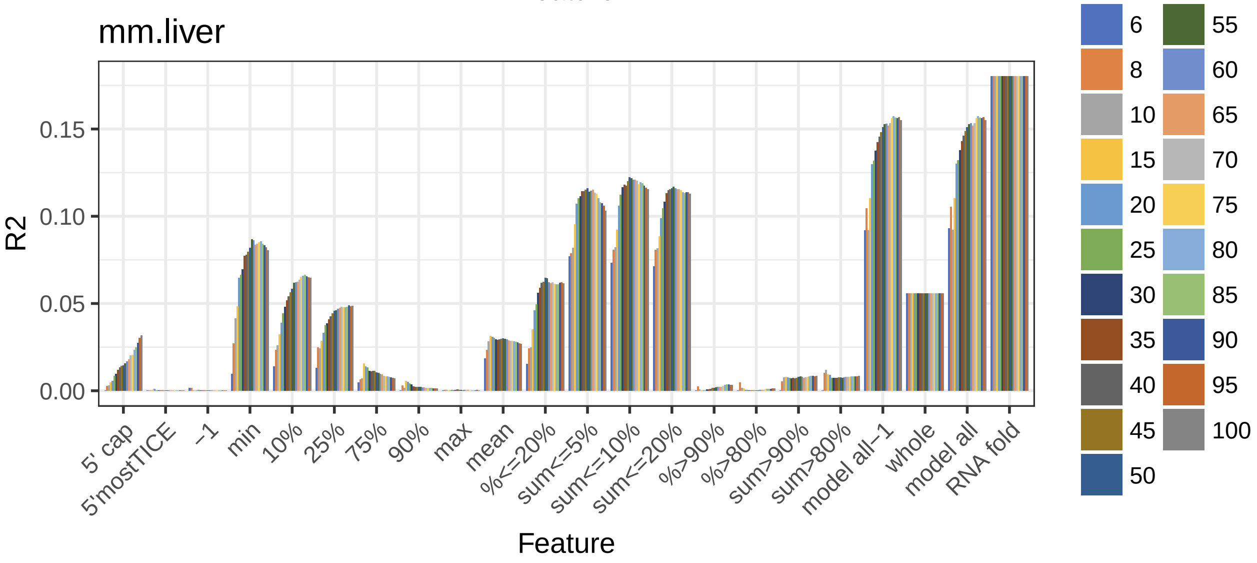
mm.liver
R2
0.00
0.05
0.10
0.15
5' cap
5'mostTICE
−1
min
10%
25%
75%
90%
max
mean
%<=20%
sum<=5%
sum<=10%
sum<=20%
%>90%
%>80%
sum>90%
sum>80%
model all−1
whole
model all
RNA fold
Feature
6
8
10
15
20
25
30
35
40
45
50
55
60
65
70
75
80
85
90
95
100

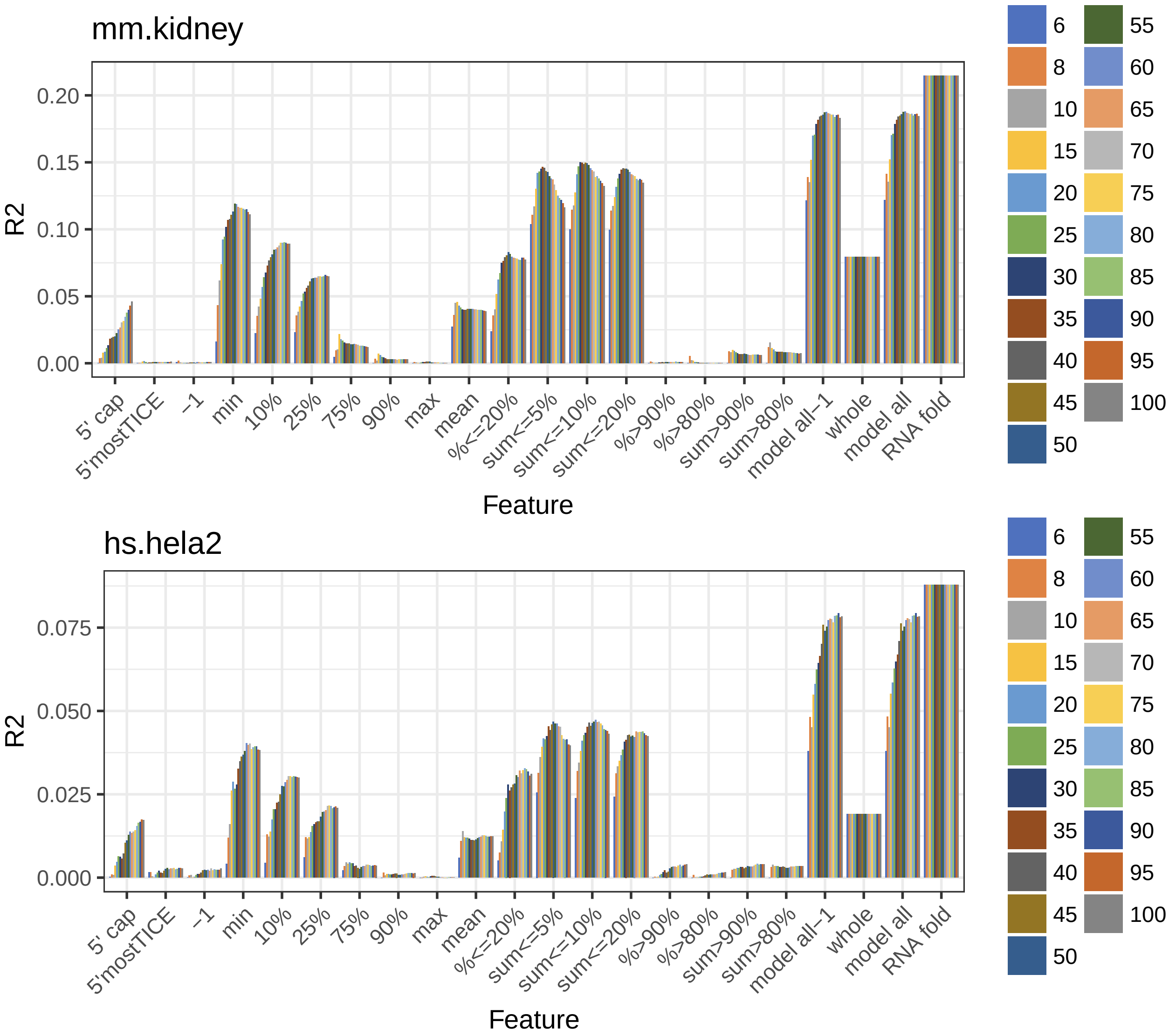


#### Figure 5. Pearson correlation(s) between min fold energy window feature(s) and log(TR)

##### the following code only needs to be run once

```
min_fold_energy_window_feature_cors <- NULL
rownames_min_fold_energy_window_feature_cors <- NULL
colnames_min_fold_energy_window_feature_cors <- c("contig stem",
    "max stem", "all")

for (species in names(species_tissue_list)) {
    for (species_tissue in species_tissue_list[[species]]) {
        if (species_tissue != "") {
            tissue_dot <- paste0(".", species_tissue)
            tissue_us <- paste0("_", species_tissue)
```

```
        } else {
            tissue_dot <- ""
            tissue_us <- ""
        }
        min_fold_energy_window_features <- read.table(paste0("data/min_window_seq_parameters/",
            species, tissue_us, ".tsv"), header = T, sep = "\t",
            stringsAsFactors = FALSE)

        # feature 1
        max_continuous_stem <- min_fold_energy_window_features$max_continuous_stem
        names(max_continuous_stem) <- min_fold_energy_window_features$id
        cor_max_continuous_stem <- cor(get(paste0(species, tissue_dot,
            ".log10.TR")), max_continuous_stem[get(paste0(species,
            tissue_dot, ".genes"))], use = "complete.obs")

        # feature 2
        max_stem_paired <- min_fold_energy_window_features$max_stem_paired
        names(max_stem_paired) <- min_fold_energy_window_features$id
        cor_max_stem_paired <- cor(get(paste0(species, tissue_dot,
            ".log10.TR")), max_stem_paired[get(paste0(species,
            tissue_dot, ".genes"))], use = "complete.obs")

        # feature 3
        paired_nts <- min_fold_energy_window_features$paired_nts
        names(paired_nts) <- min_fold_energy_window_features$id
        cor_paired_nts <- cor(get(paste0(species, tissue_dot,
            ".log10.TR")), paired_nts[get(paste0(species, tissue_dot,
            ".genes"))], use = "complete.obs")

        min_fold_energy_window_feature_cors <- rbind(min_fold_energy_window_feature_cors,
            c(cor_max_continuous_stem, cor_max_stem_paired, cor_paired_nts))
        rownames_min_fold_energy_window_feature_cors <- c(rownames_min_fold_energy_window_feature_cors,
            paste0(species, tissue_dot))
    }
}
rownames(min_fold_energy_window_feature_cors) <- rownames_min_fold_energy_window_feature_cors
colnames(min_fold_energy_window_feature_cors) <- colnames_min_fold_energy_window_feature_cors

write.csv(min_fold_energy_window_feature_cors, file = "tables/min_fold_energy_window_feature_cors.csv",
    quote = F)
```

##### plot

```
min_fold_energy_window_feature_cors <- read.csv(file = "tables/min_fold_energy_window_feature_cors.csv"
colnames(min_fold_energy_window_feature_cors)[1] <- "species.tissue"

min_fold_energy_window_feature_cols <- c("#AECDE1", "#447DB2",
    "#386895")
min_fold_energy_window_feature_cors_for_plot <- setNames(melt(min_fold_energy_window_feature_cors),
    c("Species_Tissue", "Feature", "Pearson_Cor"))
```

```
## Using species.tissue as id variables
```

```r
theme_set(theme_bw())
print(ggplot(min_fold_energy_window_feature_cors_for_plot, aes(x = Species_Tissue,
    y = Pearson_Cor, fill = Feature)) + geom_bar(stat = "identity",
    position = position_dodge()) + scale_fill_manual(values = min_fold_energy_window_feature_cols) +
    theme(axis.text.x = element_text(angle = 45, vjust = 1, hjust = 1)) +
    ggtitle("min fold energy window features"))
```

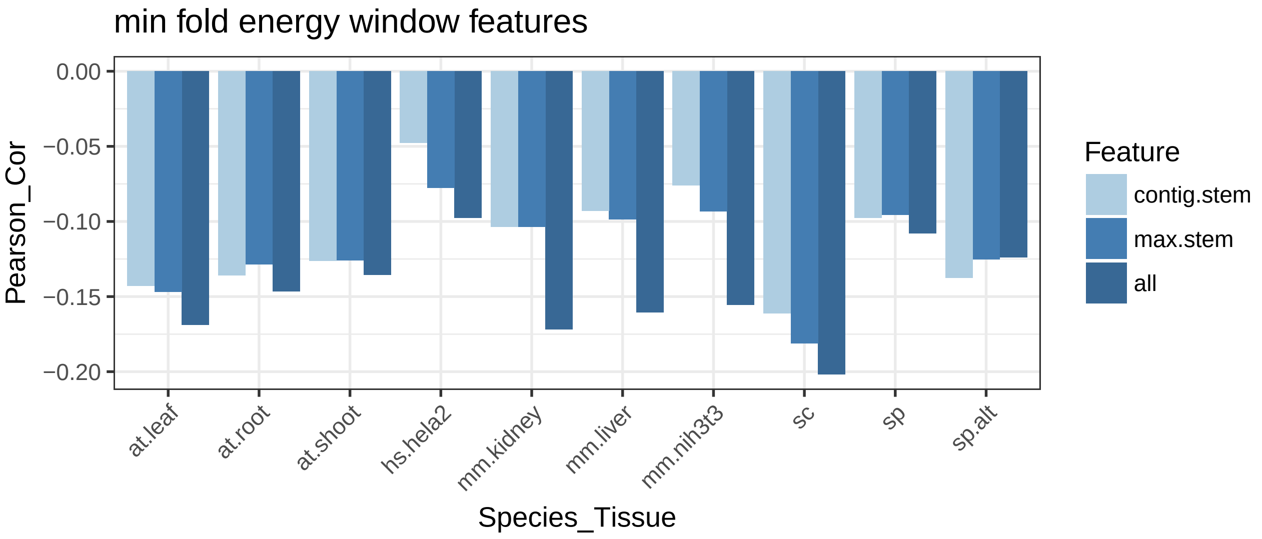


#### Figure 6. Difference(s) of min fold energy window feature(s) between top TR and bottom TR genes

##### the following code only needs to be run once

```r
min_fold_energy_window_feature_top_bottom_diffs <- NULL
rownames_min_fold_energy_window_feature_top_bottom_diffs <- NULL
colnames_min_fold_energy_window_feature_top_bottom_diffs <- c("MFE",
    "TR", "loop", "contig stem", "max stem", "all")

for (species in names(species_tissue_list)) {
    for (species_tissue in species_tissue_list[[species]]) {
        if (species_tissue != "") {
            tissue_dot <- paste0(".", species_tissue)
            tissue_us <- paste0("_", species_tissue)
        } else {
            tissue_dot <- ""
            tissue_us <- ""
        }
        min_fold_energy_window_features <- read.table(paste0("data/min_window_seq_parameters/",
            species, tissue_us, ".tsv"), header = T, sep = "\t",
            stringsAsFactors = FALSE)

        # feature 1
        mfe <- min_fold_energy_window_features$mfe
        names(mfe) <- min_fold_energy_window_features$id
```

```
        diff_mfe <- mean(mfe[get(paste0(species, tissue_dot,
            ".TR_bottom_genes"))], na.rm = T) - mean(mfe[get(paste0(species,
            tissue_dot, ".TR_top_genes"))], na.rm = T)

        # feature 2
        log10.TR <- get(paste0(species, tissue_dot, ".log10.TR"))
        diff_log10.TR <- mean(log10.TR[get(paste0(species, tissue_dot,
            ".TR_bottom_genes"))], na.rm = T) - mean(log10.TR[get(paste0(species,
            tissue_dot, ".TR_top_genes"))], na.rm = T)

        # feature 3
        max_loop <- min_fold_energy_window_features$max_loop
        names(max_loop) <- min_fold_energy_window_features$id
        diff_max_loop <- mean(max_loop[get(paste0(species, tissue_dot,
            ".TR_bottom_genes"))], na.rm = T) - mean(max_loop[get(paste0(species,
            tissue_dot, ".TR_top_genes"))], na.rm = T)

        # feature 4
        max_continuous_stem <- min_fold_energy_window_features$max_continuous_stem
        names(max_continuous_stem) <- min_fold_energy_window_features$id
        diff_max_continuous_stem <- mean(max_continuous_stem[get(paste0(species,
            tissue_dot, ".TR_bottom_genes"))], na.rm = T) - mean(max_continuous_stem[get(paste0(species
            tissue_dot, ".TR_top_genes"))], na.rm = T)

        # feature 5
        max_stem_paired <- min_fold_energy_window_features$max_stem_paired
        names(max_stem_paired) <- min_fold_energy_window_features$id
        diff_max_stem_paired <- mean(max_stem_paired[get(paste0(species,
            tissue_dot, ".TR_bottom_genes"))], na.rm = T) - mean(max_stem_paired[get(paste0(species,
            tissue_dot, ".TR_top_genes"))], na.rm = T)

        # feature 6
        paired_nts <- min_fold_energy_window_features$paired_nts
        names(paired_nts) <- min_fold_energy_window_features$id
        diff_paired_nts <- mean(paired_nts[get(paste0(species,
            tissue_dot, ".TR_bottom_genes"))], na.rm = T) - mean(paired_nts[get(paste0(species,
            tissue_dot, ".TR_top_genes"))], na.rm = T)

        min_fold_energy_window_feature_top_bottom_diffs <- rbind(min_fold_energy_window_feature_top_bot
            c(diff_mfe, diff_log10.TR, diff_max_loop, diff_max_continuous_stem,
                diff_max_stem_paired, diff_paired_nts))
        rownames_min_fold_energy_window_feature_top_bottom_diffs <- c(rownames_min_fold_energy_window_f
            paste0(species, tissue_dot))
    }
}
rownames(min_fold_energy_window_feature_top_bottom_diffs) <- rownames_min_fold_energy_window_feature_top
colnames(min_fold_energy_window_feature_top_bottom_diffs) <- colnames_min_fold_energy_window_feature_top

write.csv(min_fold_energy_window_feature_top_bottom_diffs, file = "tables/min_fold_energy_window_featur
    quote = F)
```

##### plot

```
min_fold_energy_window_feature_top_bottom_diffs <- read.csv(file = "tables/min_fold_energy_window_featu
colnames(min_fold_energy_window_feature_top_bottom_diffs)[1] <- "species.tissue"

# only keep the exemplar tissues
min_fold_energy_window_feature_top_bottom_diffs_for_plot <- setNames(melt(min_fold_energy_window_featur
    species_exemplar_tissue_names, ]), c("Species_Tissue", "Feature",
    "Bottom_minus_Top_Diff"))
```

```
## Using species.tissue as id variables
```

```
min_fold_energy_window_feature_top_bottom_diffs_for_plot$Species_Tissue <- factor(min_fold_energy_windo
    levels = c("sc", "sp.alt", "at.leaf", "mm.nih3t3", "hs.hela2"))

theme_set(theme_bw())
print(ggplot(min_fold_energy_window_feature_top_bottom_diffs_for_plot,
    aes(x = Feature, y = Bottom_minus_Top_Diff, fill = Species_Tissue)) +
    geom_bar(stat = "identity", position = position_dodge()) +
    scale_fill_manual(values = species_cols[c("sc", "sp.alt",
        "at.leaf", "mm.nih3t3", "hs.hela2")]) + theme(axis.text.x = element_text(angle = 45,
    vjust = 1, hjust = 1)) + ggtitle("min fold energy window features (TR bottom - TR top)"))
```

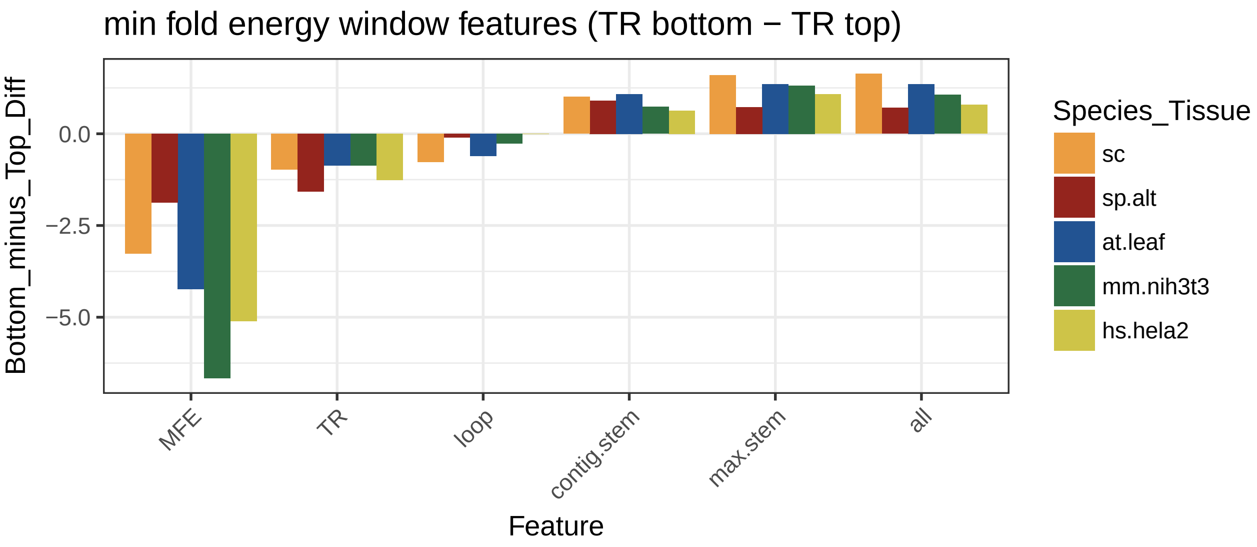


#### Figure 6 - figure supplement 1. Distribution(s) of the number(s) of paired nucleotides in high/low MFE genes or high/low TR genes

```
# my_plot_hook <- function(x, options) paste('\n',
# knitr::hook_plot_tex(x, options), '\n')
# knitr::knit_hooks$set(plot = my_plot_hook)

myred <- "#88342D"

for (species in names(species_tissue_list)) {
    for (species_tissue in species_tissue_list[[species]]) {
```

```r
    if (species_tissue != "") {
        tissue_dot <- paste0(".", species_tissue)
        tissue_us <- paste0("_", species_tissue)
    } else {
        tissue_dot <- ""
        tissue_us <- ""
    }
    min_fold_energy_window_features <- read.table(paste0("data/min_window_seq_parameters/",
        species, tissue_us, ".tsv"), header = T, sep = "\t",
        stringsAsFactors = FALSE)

    mfe_quantiles <- quantile(min_fold_energy_window_features$mfe,
        prob = c(0.1, 0.9))

    for (feature_name in c("paired_nts")) {
        feature <- min_fold_energy_window_features[, feature_name]
        names(feature) <- min_fold_energy_window_features$id

        density1 <- density(feature[min_fold_energy_window_features$mfe >=
            mfe_quantiles[2]], bw = 1, na.rm = TRUE)
        density2 <- density(feature[min_fold_energy_window_features$mfe <=
            mfe_quantiles[1]], bw = 1, na.rm = TRUE)
        density3 <- density(feature[get(paste0(species, tissue_dot,
            ".TR_top_genes"))], bw = 1, na.rm = TRUE)
        density4 <- density(feature[get(paste0(species, tissue_dot,
            ".TR_bottom_genes"))], bw = 1, na.rm = TRUE)

        mean1 <- mean(feature[min_fold_energy_window_features$mfe >=
            mfe_quantiles[2]], na.rm = TRUE)
        mean2 <- mean(feature[min_fold_energy_window_features$mfe <=
            mfe_quantiles[1]], na.rm = TRUE)
        mean3 <- mean(feature[get(paste0(species, tissue_dot,
            ".TR_top_genes"))], na.rm = TRUE)
        mean4 <- mean(feature[get(paste0(species, tissue_dot,
            ".TR_bottom_genes"))], na.rm = TRUE)

        plot(0, 0, type = "n", xlim = range(c(density1$x,
            density2$x, density3$x, density4$x)), ylim = range(c(density1$y,
            density2$y, density3$y, density4$y)), xaxt = "n",
            xlab = feature, ylab = "Density", main = paste0(species,
              tissue_dot))
        axis(1)
        lines(density1, col = 1, lwd = 3)
        abline(v = mean1, lty = 2, col = 1, lwd = 3)
        lines(density2, col = myred, lwd = 3)
        abline(v = mean2, lty = 2, col = myred, lwd = 3)
        legend("topright", c("Top MFE", "Bottom MFE"), col = c(1,
            myred), lty = c(1, 1), bty = "n")
        arrows(x0 = mean1, y0 = 0, x1 = mean2, y1 = 0, col = 1,
            length = 0.05, code = 3)
        text(x = (mean1 + mean2)/2, y = 0.01, labels = round(mean2 -
            mean1, 1))
        text(x = mean1 - 1, y = 0.02, labels = round(mean1,
```

```
                1))
            text(x = mean2 + 1, y = 0.02, labels = round(mean2,
                1), col = myred)

            plot(0, 0, type = "n", xlim = range(c(density1$x,
                density2$x, density3$x, density4$x)), ylim = range(c(density1$y,
                density2$y, density3$y, density4$y)), xaxt = "n",
                xlab = feature, ylab = "Density", main = paste0(species,
                  tissue_dot))
            axis(1)
            lines(density3, col = 1, lwd = 3)
            abline(v = mean3, lty = 2, col = 1, lwd = 3)
            lines(density4, col = myred, lwd = 3)
            abline(v = mean4, lty = 2, col = myred, lwd = 3)
            legend("topright", c("Top TR", "Bottom TR"), col = c(1,
                myred), lty = c(1, 1), bty = "n")
            arrows(x0 = mean3, y0 = 0, x1 = mean4, y1 = 0, col = 1,
                length = 0.05, code = 3)
            text(x = (mean3 + mean4)/2, y = 0.01, labels = round(mean4 -
                mean3, 1))
            text(x = mean3 - 1, y = 0.02, labels = round(mean3,
                1))
            text(x = mean4 + 1, y = 0.02, labels = round(mean4,
                1), col = myred)
        }
    }
}
```

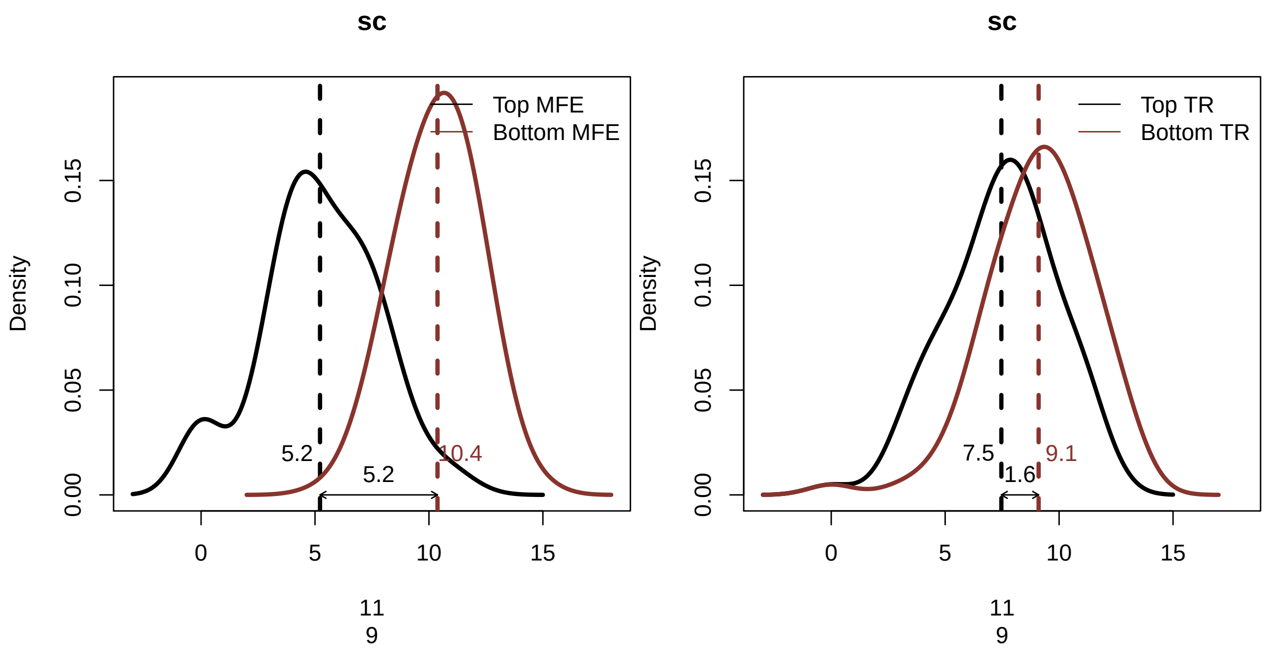

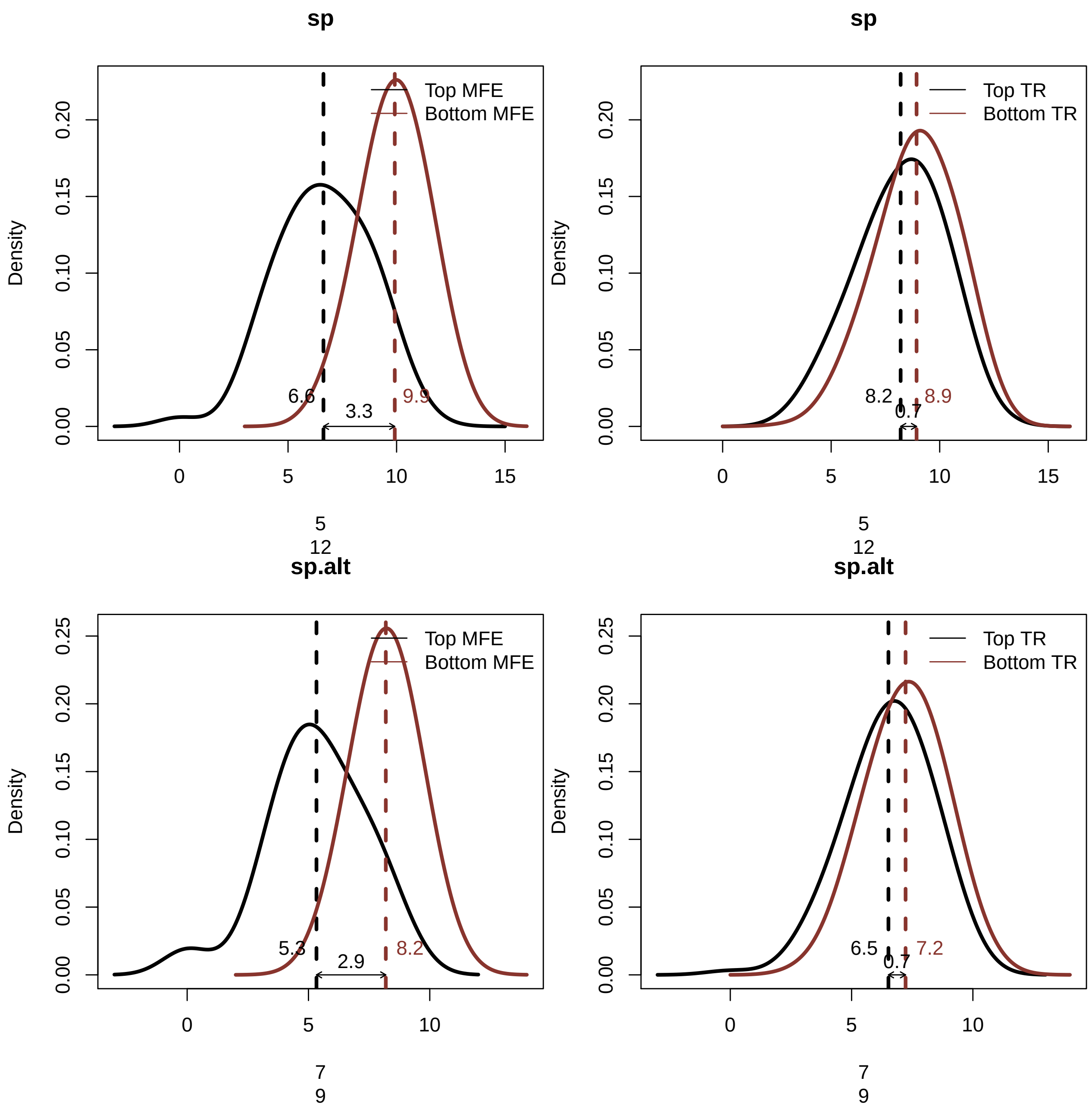
sp
Top MFE
Bottom MFE
Density
6.6
3.3
9.9
5
12
sp
Top TR
Bottom TR
Density
8.2
0.7
8.9
5
12
sp.alt
Top MFE
Bottom MFE
Density
5.3
2.9
8.2
7
9
sp.alt
Top TR
Bottom TR
Density
6.5
0.7
7.2
7
9

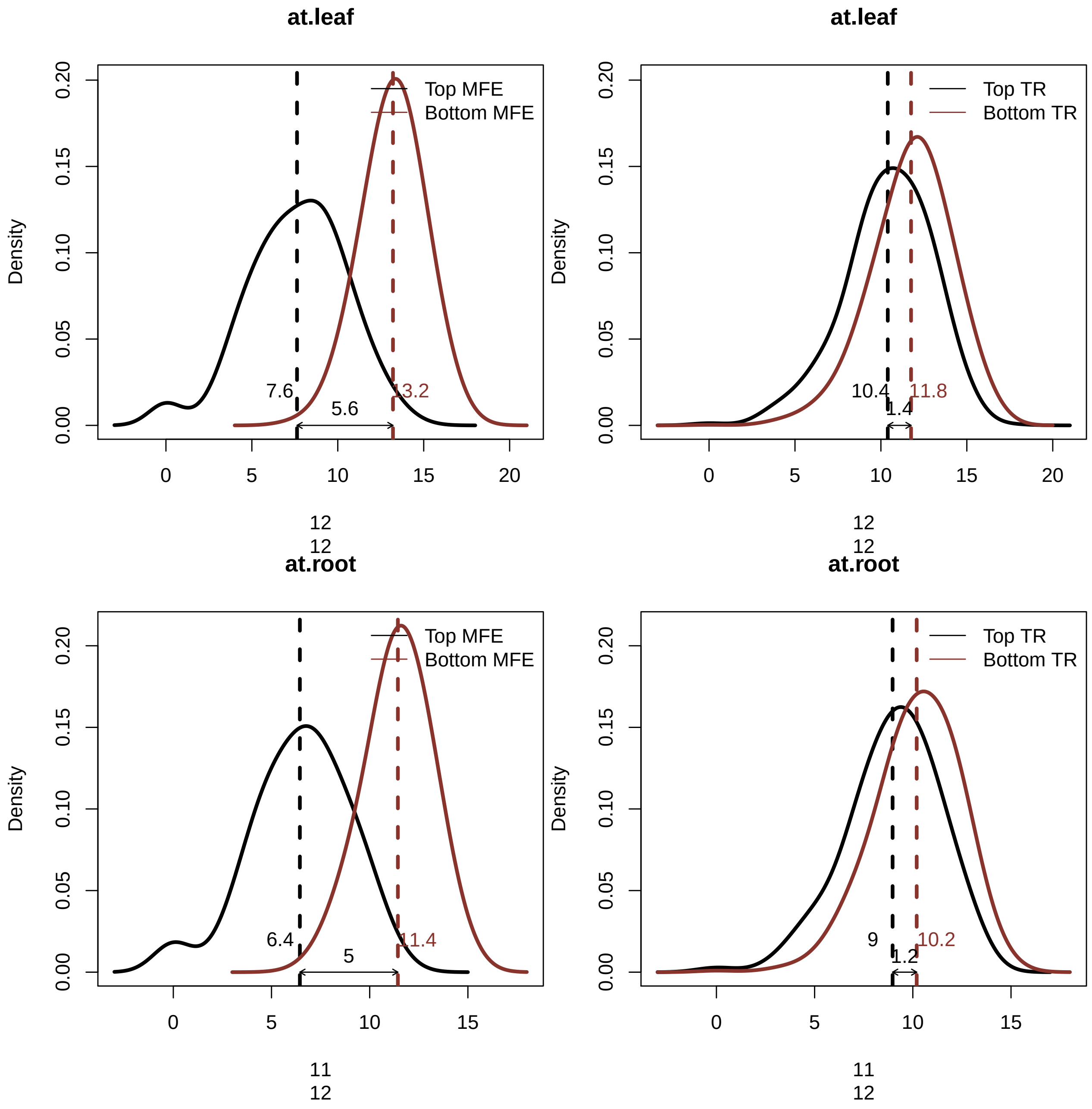
at.leaf
Top MFE
Bottom MFE
Density
7.6
13.2
5.6
12
12
at.leaf
Top TR
Bottom TR
Density
10.4
11.8
1.4
12
12
at.root
Top MFE
Bottom MFE
Density
6.4
11.4
5
11
12
at.root
Top TR
Bottom TR
Density
9
10.2
1.2
11
12

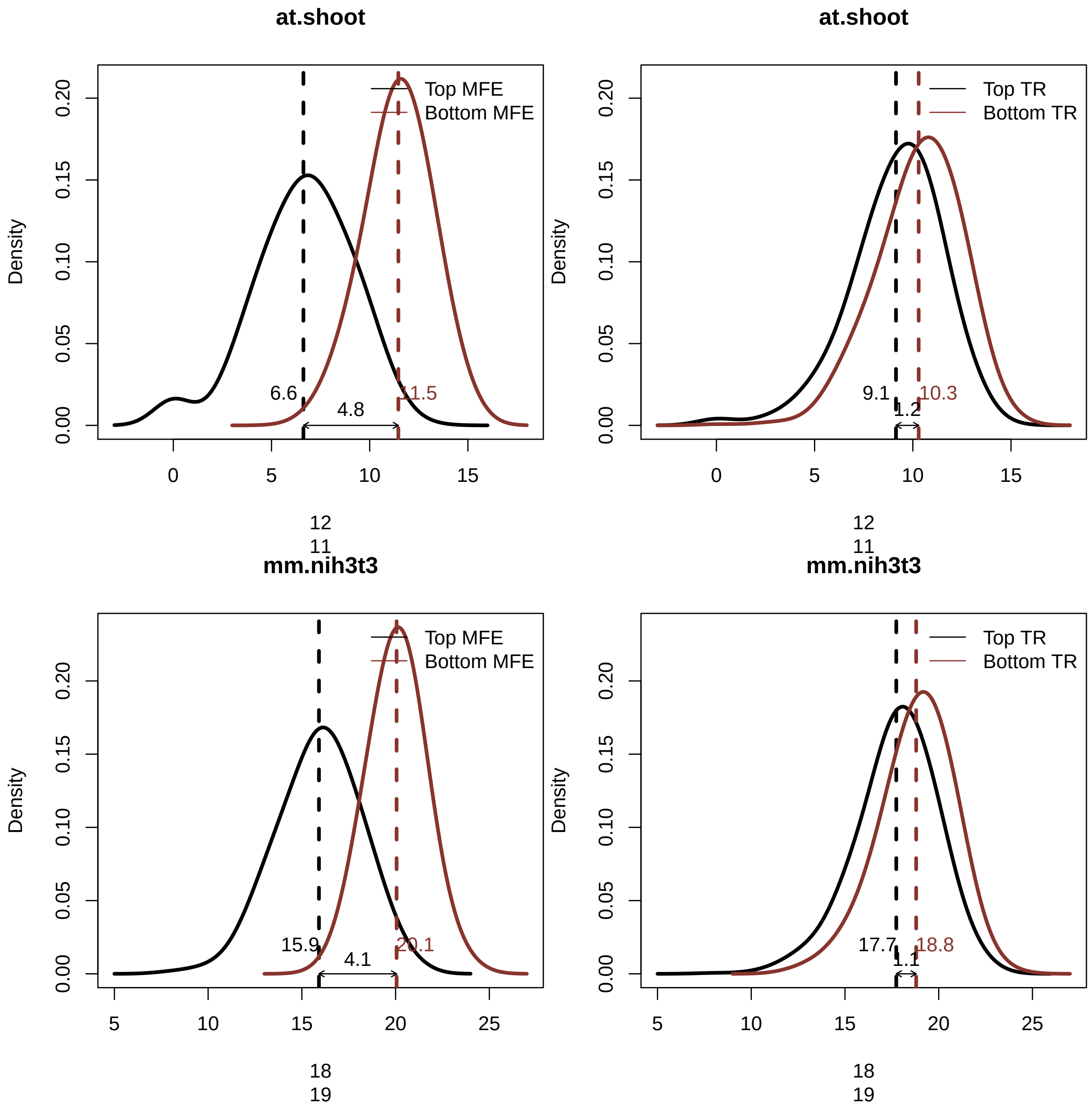
at.shoot
Top MFE
Bottom MFE
6.6
11.5
4.8
Density
12
11
at.shoot
Top TR
Bottom TR
9.1
10.3
1.2
Density
12
11
mm.nih3t3
Top MFE
Bottom MFE
15.9
20.1
4.1
Density
18
19
mm.nih3t3
Top TR
Bottom TR
17.7
18.8
1.1
Density
18
19

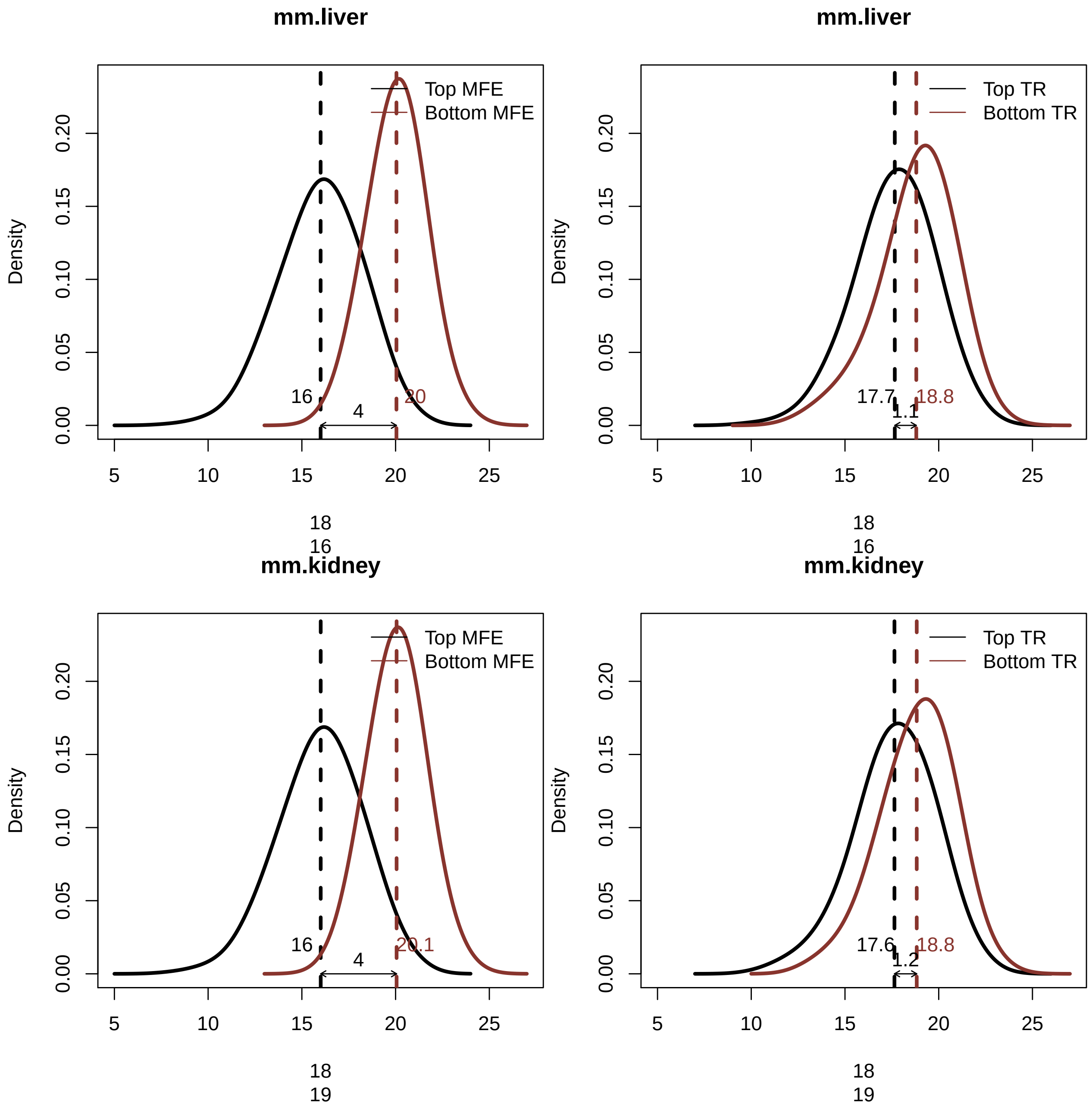
mm.liver
Top MFE
Bottom MFE
Density
16
20
4
18
16
mm.liver
Top TR
Bottom TR
Density
17.7
18.8
1.1
18
16
mm.kidney
Top MFE
Bottom MFE
Density
16
20.1
4
18
19
mm.kidney
Top TR
Bottom TR
Density
17.6
18.8
1.2
18
19

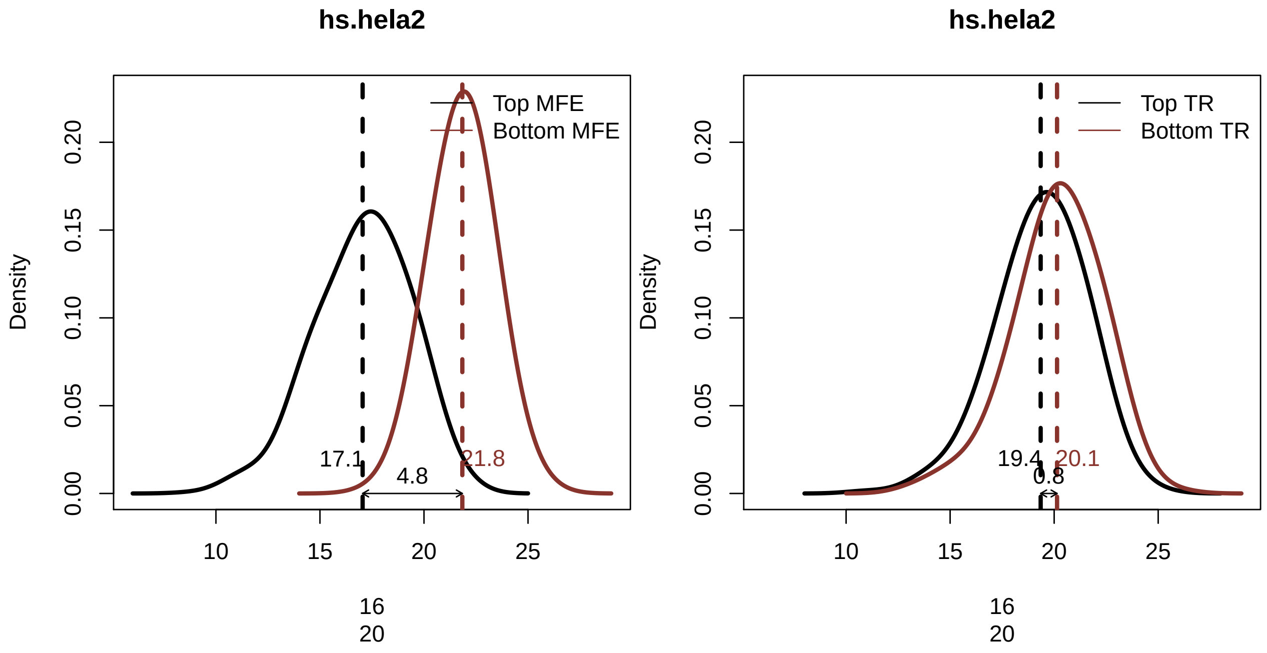


### Figure 7. The 5' and 3' boundaries of TICE.

```
for (species in names(species_exemplar_tissue_list)) {
    for (species_tissue in species_exemplar_tissue_list[[species]]) {
        if (species_tissue != "") {
            species.genes <- get(paste(species, species_tissue,
                "genes", sep = "."))
            species.UTR_seqs <- get(paste(species, species_tissue,
                "UTR_seqs", sep = "."))
            species.CDS_seqs <- get(paste(species, species_tissue,
                "CDS_seqs", sep = "."))
            species.log10.TR <- get(paste(species, species_tissue,
                "log10.TR", sep = "."))
            species.TR_top_genes <- get(paste(species, species_tissue,
                "TR_top_genes", sep = "."))
            species.TR_bottom_genes <- get(paste(species, species_tissue,
                "TR_bottom_genes", sep = "."))
            connector <- paste0(species_tissue, ".")
        } else {
            species.genes <- get(paste(species, "genes", sep = "."))
            species.UTR_seqs <- get(paste(species, "UTR_seqs",
                sep = "."))
            species.CDS_seqs <- get(paste(species, "CDS_seqs",
                sep = "."))
            species.log10.TR <- get(paste(species, "log10.TR",
                sep = "."))
            species.TR_top_genes <- get(paste(species, "TR_top_genes",
                sep = "."))
            species.TR_bottom_genes <- get(paste(species, "TR_bottom_genes",
                sep = "."))
            connector <- ""
```

```
}
species.log10.UTR_len <- log(nchar(species.UTR_seqs),
    10)
if (species != "hs") {
    species.uTICE_len <- species_uTICE_len_list[[species]]
} else {
    species.uTICE_len <- species_uTICE_len_list[[species]][[species_tissue]]
}

## for the genes in the top and bottom cohort, find the 5' UTR
## sequences and CDS sequences
species.TR_top_UTR_seqs <- species.UTR_seqs[species.TR_top_genes]
species.TR_top_CDS_seqs <- species.CDS_seqs[species.TR_top_genes]

species.TR_bottom_UTR_seqs <- species.UTR_seqs[species.TR_bottom_genes]
species.TR_bottom_CDS_seqs <- species.CDS_seqs[species.TR_bottom_genes]

# construct Position Weight Matrices (PWMs) based on the top
# 10% TR genes with different 5' UTR lengths and ORF lengths
max_UTR_len <- 100
len_breaks <- seq(5, max_UTR_len, by = 5)  # 5' UTR lengths vary from 100 bp to 5 bp

## top TR
species.TR_top_PWM_max_UTR_len <- UTR_pwm(UTR_seqs = species.TR_top_UTR_seqs,
    n = max_UTR_len)  # no CDS length

species.TR_top_PWM_max_UTR_len_CDS_13 <- UTR_CDS_pwm(UTR_seqs = species.TR_top_UTR_seqs,
    CDS_seqs = species.TR_top_CDS_seqs, n_UTR = max_UTR_len,
    n_CDS = 13)  # 13 bp CDS
colnames(species.TR_top_PWM_max_UTR_len_CDS_13) <- c("A",
    "C", "G", "T")

species.TR_top_PWM_max_UTR_len_CDS_23 <- UTR_CDS_pwm(UTR_seqs = species.TR_top_UTR_seqs,
    CDS_seqs = species.TR_top_CDS_seqs, n_UTR = max_UTR_len,
    n_CDS = 23)  # 23 bp CDS
colnames(species.TR_top_PWM_max_UTR_len_CDS_23) <- c("A",
    "C", "G", "T")

species.TR_top_PWM_max_UTR_len_CDS_28 <- UTR_CDS_pwm(UTR_seqs = species.TR_top_UTR_seqs,
    CDS_seqs = species.TR_top_CDS_seqs, n_UTR = max_UTR_len,
    n_CDS = 28)  # 28 bp CDS
colnames(species.TR_top_PWM_max_UTR_len_CDS_28) <- c("A",
    "C", "G", "T")

species.TR_top_PWM_max_UTR_len_CDS_33 <- UTR_CDS_pwm(UTR_seqs = species.TR_top_UTR_seqs,
    CDS_seqs = species.TR_top_CDS_seqs, n_UTR = max_UTR_len,
    n_CDS = 33)  # 33 bp CDS
colnames(species.TR_top_PWM_max_UTR_len_CDS_33) <- c("A",
    "C", "G", "T")

saveRDS(species.TR_top_PWM_max_UTR_len_CDS_33, file = paste0("processed_data_w_polyA/AUGPWM_R2s/
    species, ".", connector, "TR_top_PWM_AUG_aligned.rds"))
```

```
## bottom TR
species.TR_bottom_PWM_max_UTR_len <- UTR_pwm(UTR_seqs = species.TR_bottom_UTR_seqs,
    n = max_UTR_len)  # no CDS length

species.TR_bottom_PWM_max_UTR_len_CDS_13 <- UTR_CDS_pwm(UTR_seqs = species.TR_bottom_UTR_seqs,
    CDS_seqs = species.TR_bottom_CDS_seqs, n_UTR = max_UTR_len,
    n_CDS = 13)  # 13 bp CDS
colnames(species.TR_bottom_PWM_max_UTR_len_CDS_13) <- c("A",
    "C", "G", "T")

species.TR_bottom_PWM_max_UTR_len_CDS_23 <- UTR_CDS_pwm(UTR_seqs = species.TR_bottom_UTR_seqs,
    CDS_seqs = species.TR_bottom_CDS_seqs, n_UTR = max_UTR_len,
    n_CDS = 23)  # 23 bp CDS
colnames(species.TR_bottom_PWM_max_UTR_len_CDS_23) <- c("A",
    "C", "G", "T")

species.TR_bottom_PWM_max_UTR_len_CDS_28 <- UTR_CDS_pwm(UTR_seqs = species.TR_bottom_UTR_seqs,
    CDS_seqs = species.TR_bottom_CDS_seqs, n_UTR = max_UTR_len,
    n_CDS = 28)  # 28 bp CDS
colnames(species.TR_bottom_PWM_max_UTR_len_CDS_28) <- c("A",
    "C", "G", "T")

species.TR_bottom_PWM_max_UTR_len_CDS_33 <- UTR_CDS_pwm(UTR_seqs = species.TR_bottom_UTR_seqs,
    CDS_seqs = species.TR_bottom_CDS_seqs, n_UTR = max_UTR_len,
    n_CDS = 33)  # 33 bp CDS
colnames(species.TR_bottom_PWM_max_UTR_len_CDS_33) <- c("A",
    "C", "G", "T")

saveRDS(species.TR_bottom_PWM_max_UTR_len_CDS_33, file = paste0("processed_data_w_polyA/AUGPWM_
    species, ".", connector, "TR_bottom_PWM_AUG_aligned.rds"))


# PWM scores (aligned at AUG)

## top TR varying 5' UTR lengths, 0 bp CDS length
filename <- paste0("processed_data_w_polyA/AUGPWM_R2s/",
    species, ".", connector, "TR_top_PWM_max_UTR_len_score.rds")
if (!file.exists(filename)) {
    species.TR_top_PWM_max_UTR_len_score <- sapply(len_breaks,
        FUN = function(i) {
          UTR_pwm_score(species.TR_top_PWM_max_UTR_len[(i -
            5 + 1):nrow(species.TR_top_PWM_max_UTR_len),
            ], species.UTR_seqs)
        })
    saveRDS(species.TR_top_PWM_max_UTR_len_score, file = filename)
}

### varying 5' UTR lengths, 13 bp CDS length
filename <- paste0("processed_data_w_polyA/AUGPWM_R2s/",
    species, ".", connector, "TR_top_PWM_max_UTR_len_CDS_13_score.rds")
if (!file.exists(filename)) {
    species.UTR_CDS_13_seqs <- sapply(species.genes,
        FUN = function(x) {
```

```
        paste(species.UTR_seqs[x], paste(strsplit(species.CDS_seqs[x],
          "")[[1]][4:13], collapse = ""), sep = "")
      })
    species.TR_top_PWM_max_UTR_len_CDS_13_score <- sapply(len_breaks,
      FUN = function(i) {
        UTR_pwm_score(species.TR_top_PWM_max_UTR_len_CDS_13[(i -
          5 + 1):nrow(species.TR_top_PWM_max_UTR_len_CDS_13),
          ], species.UTR_CDS_13_seqs)
      })
    saveRDS(species.TR_top_PWM_max_UTR_len_CDS_13_score,
      file = filename)
}

### varying 5' UTR lengths, 23 bp CDS length
filename <- paste0("processed_data_w_polyA/AUGPWM_R2s/",
    species, ".", connector, "TR_top_PWM_max_UTR_len_CDS_23_score.rds")
if (!file.exists(filename)) {
    species.UTR_CDS_23_seqs <- sapply(species.genes,
      FUN = function(x) {
        paste(species.UTR_seqs[x], paste(strsplit(species.CDS_seqs[x],
          "")[[1]][4:23], collapse = ""), sep = "")
      })
    species.TR_top_PWM_max_UTR_len_CDS_23_score <- sapply(len_breaks,
      FUN = function(i) {
        UTR_pwm_score(species.TR_top_PWM_max_UTR_len_CDS_23[(i -
          5 + 1):nrow(species.TR_top_PWM_max_UTR_len_CDS_23),
          ], species.UTR_CDS_23_seqs)
      })
    saveRDS(species.TR_top_PWM_max_UTR_len_CDS_23_score,
      file = filename)
}

### varying 5' UTR lengths, 28 bp CDS length
filename <- paste0("processed_data_w_polyA/AUGPWM_R2s/",
    species, ".", connector, "TR_top_PWM_max_UTR_len_CDS_28_score.rds")
if (!file.exists(filename)) {
    species.UTR_CDS_28_seqs <- sapply(species.genes,
      FUN = function(x) {
        paste(species.UTR_seqs[x], paste(strsplit(species.CDS_seqs[x],
          "")[[1]][4:28], collapse = ""), sep = "")
      })
    species.TR_top_PWM_max_UTR_len_CDS_28_score <- sapply(len_breaks,
      FUN = function(i) {
        UTR_pwm_score(species.TR_top_PWM_max_UTR_len_CDS_28[(i -
          5 + 1):nrow(species.TR_top_PWM_max_UTR_len_CDS_28),
          ], species.UTR_CDS_28_seqs)
      })
    saveRDS(species.TR_top_PWM_max_UTR_len_CDS_28_score,
      file = filename)
}

### varying 5' UTR lengths, 33 bp CDS length
filename <- paste0("processed_data_w_polyA/AUGPWM_R2s/",
```

```r
            species, ".", connector, "TR_top_PWM_max_UTR_len_CDS_33_score.rds")
        if (!file.exists(filename)) {
            species.UTR_CDS_33_seqs <- sapply(species.genes,
                FUN = function(x) {
                  paste(species.UTR_seqs[x], paste(strsplit(species.CDS_seqs[x],
                    "")[[1]][4:33], collapse = ""), sep = "")
                })
            species.TR_top_PWM_max_UTR_len_CDS_33_score <- sapply(len_breaks,
                FUN = function(i) {
                  UTR_pwm_score(species.TR_top_PWM_max_UTR_len_CDS_33[(i -
                    5 + 1):nrow(species.TR_top_PWM_max_UTR_len_CDS_33),
                    ], species.UTR_CDS_33_seqs)
                })
            saveRDS(species.TR_top_PWM_max_UTR_len_CDS_33_score,
                file = filename)
        }
    }
}
```

```r
# a function to plot a UTR-CDS PWM
plot_UTR_CDS_pwm <- function(PWM, n_UTR, n_CDS, title) {
    PWM_plot <- rbind(PWM[1:n_UTR, ], c(1, 0, 0, 0), c(0, 0,
        0, 1), c(0, 0, 1, 0), PWM[(n_UTR + 1):nrow(PWM), ])
    colnames(PWM_plot) <- c("A", "C", "G", "U")
    PWM_motif <- new("pcm", mat = t(PWM_plot), name = title)
    plot(PWM_motif, ic.scale = F)
}

max_UTR_len <- 100
len_breaks <- seq(5, max_UTR_len, by = 5)  # 5' UTR lengths vary from 100 bp to 5 bp

for (species in names(species_exemplar_tissue_list)) {
    for (species_tissue in species_exemplar_tissue_list[[species]]) {
        if (species_tissue != "") {
            species.log10.TR <- get(paste(species, species_tissue,
                "log10.TR", sep = "."))
            species.UTR_seqs <- get(paste(species, species_tissue,
                "UTR_seqs", sep = "."))
            connector <- paste0(species_tissue, ".")
        } else {
            species.log10.TR <- get(paste(species, "log10.TR",
                sep = "."))
            species.UTR_seqs <- get(paste(species, "UTR_seqs",
                sep = "."))
            connector <- ""
        }
        species.log10.UTR_len <- log(nchar(species.UTR_seqs),
            10)

        species.TR_top_PWM_AUG_aligned <- readRDS(file = paste0("processed_data_w_polyA/AUGPWM_R2s/",
            species, ".", connector, "TR_top_PWM_AUG_aligned.rds"))
        species.TR_bottom_PWM_AUG_aligned <- readRDS(file = paste0("processed_data_w_polyA/AUGPWM_R2s/"
            species, ".", connector, "TR_bottom_PWM_AUG_aligned.rds"))
        ## plot the PWM matrices
```

```
plot_UTR_CDS_pwm(species.TR_top_PWM_AUG_aligned[21:nrow(species.TR_top_PWM_AUG_aligned),
    ], n_UTR = 80, n_CDS = 33, title = paste0(species,
    ".", connector, "TR_top_PWM"))
plot_UTR_CDS_pwm(species.TR_bottom_PWM_AUG_aligned[21:nrow(species.TR_bottom_PWM_AUG_aligned),
    ], n_UTR = 80, n_CDS = 33, title = paste0(species,
    ".", connector, "TR_bottom_PWM"))

# PWM scores (aligned at AUG)
species.TR_top_PWM_max_UTR_len_score <- readRDS(paste0("processed_data_w_polyA/AUGPWM_R2s/",
    species, ".", connector, "TR_top_PWM_max_UTR_len_score.rds"))

species.TR_top_PWM_max_UTR_len_CDS_13_score <- readRDS(paste0("processed_data_w_polyA/AUGPWM_R2s
    species, ".", connector, "TR_top_PWM_max_UTR_len_CDS_13_score.rds"))

species.TR_top_PWM_max_UTR_len_CDS_23_score <- readRDS(paste0("processed_data_w_polyA/AUGPWM_R2s
    species, ".", connector, "TR_top_PWM_max_UTR_len_CDS_23_score.rds"))

species.TR_top_PWM_max_UTR_len_CDS_28_score <- readRDS(paste0("processed_data_w_polyA/AUGPWM_R2s
    species, ".", connector, "TR_top_PWM_max_UTR_len_CDS_28_score.rds"))

species.TR_top_PWM_max_UTR_len_CDS_33_score <- readRDS(paste0("processed_data_w_polyA/AUGPWM_R2s
    species, ".", connector, "TR_top_PWM_max_UTR_len_CDS_33_score.rds"))

# calculate the R2s for genes with different minimum 5' UTR
# lengths top TR
species.TR_top_R2s_w_TR <- sapply(1:length(len_breaks),
    FUN = function(idx) {
        long_idx <- which(species.log10.UTR_len >= log(100 -
          len_breaks[idx] + 5, 10))
        c(cor(species.log10.TR[long_idx], species.TR_top_PWM_max_UTR_len_score[,
          idx][long_idx])^2, cor(species.log10.TR[long_idx],
          species.TR_top_PWM_max_UTR_len_CDS_13_score[,
            idx][long_idx])^2, cor(species.log10.TR[long_idx],
          species.TR_top_PWM_max_UTR_len_CDS_23_score[,
            idx][long_idx])^2, cor(species.log10.TR[long_idx],
          species.TR_top_PWM_max_UTR_len_CDS_28_score[,
            idx][long_idx])^2, cor(species.log10.TR[long_idx],
          species.TR_top_PWM_max_UTR_len_CDS_33_score[,
            idx][long_idx])^2)
    })
# plot the R2's

## top TR
plot(x = seq(from = -100, to = -5, by = 5), y = species.TR_top_R2s_w_TR[1,
    ], ylim = range(species.TR_top_R2s_w_TR), main = paste0(species,
    ".", connector), xlab = "length 5' UTR TR top PWM",
    ylab = "R2 logTR vs. TR top PWM score", pch = 20,
    axes = T, col = gray(0.9))
points(x = seq(from = -100, to = -5, by = 5), y = species.TR_top_R2s_w_TR[2,
    ], pch = 20, col = gray(0.7))
points(x = seq(from = -100, to = -5, by = 5), y = species.TR_top_R2s_w_TR[3,
    ], pch = 20, col = gray(0.5))
points(x = seq(from = -100, to = -5, by = 5), y = species.TR_top_R2s_w_TR[4,
```

```
        ], pch = 20, col = gray(0.3))
    points(x = seq(from = -100, to = -5, by = 5), y = species.TR_top_R2s_w_TR[5,
        ], pch = 20, col = gray(0.1))
    max_idx <- which(species.TR_top_R2s_w_TR == max(species.TR_top_R2s_w_TR),
        arr.ind = T)
    abline(v = seq(from = -100, to = -5, by = 5)[which.max(species.TR_top_R2s_w_TR[max_idx[1],
        ])], lty = 2)
    legend("bottomleft", c("5' UTR only", "5' UTR & +4 to +13",
        "5' UTR & +4 to +23", "5' UTR & +4 to +28", "5' UTR & +4 to +33"),
        pch = c(20, 20, 20, 20, 20), col = c(gray(0.9), gray(0.7),
            gray(0.5), gray(0.3), gray(0.1)), bty = "n")
    }
}
```

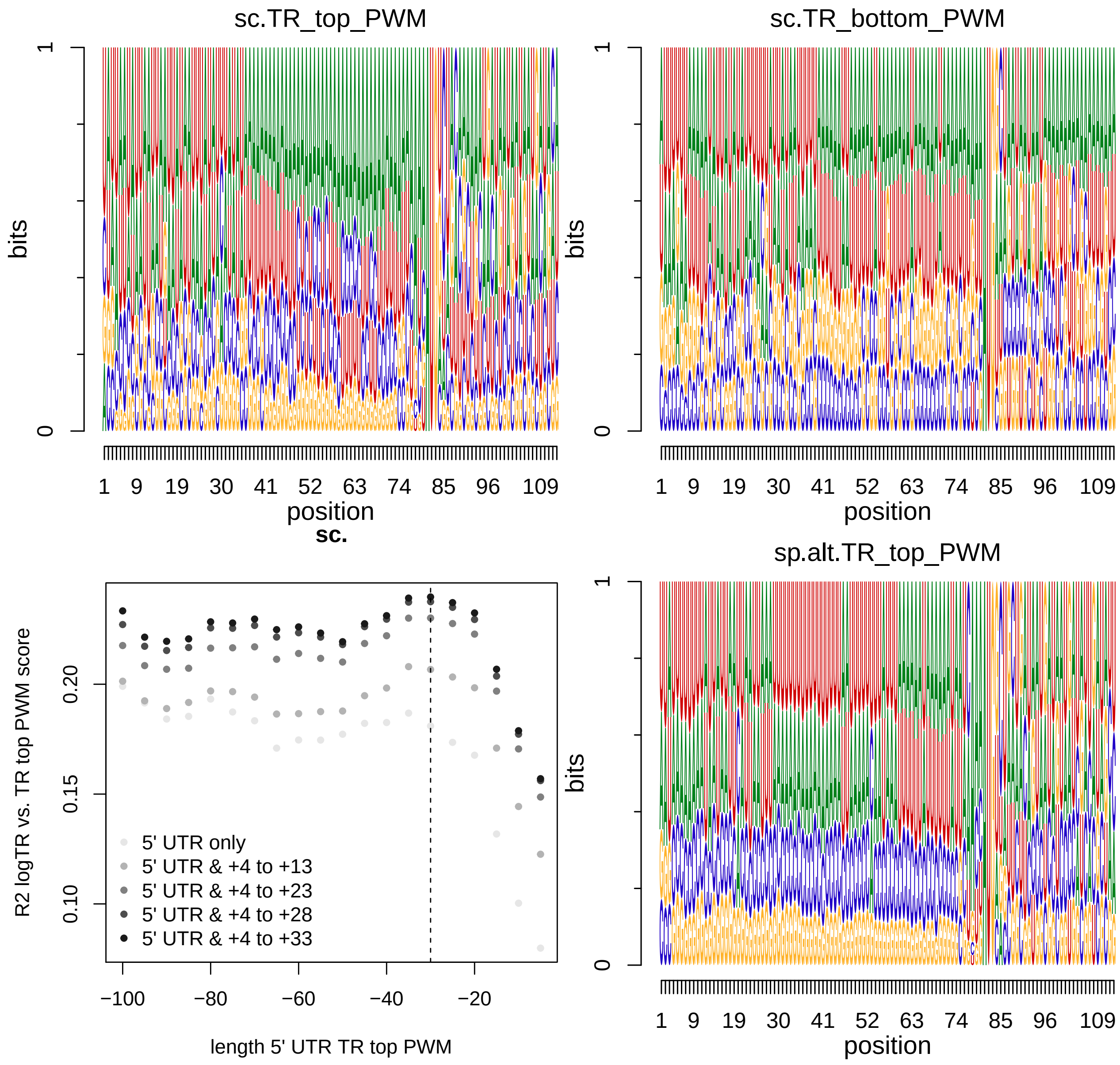

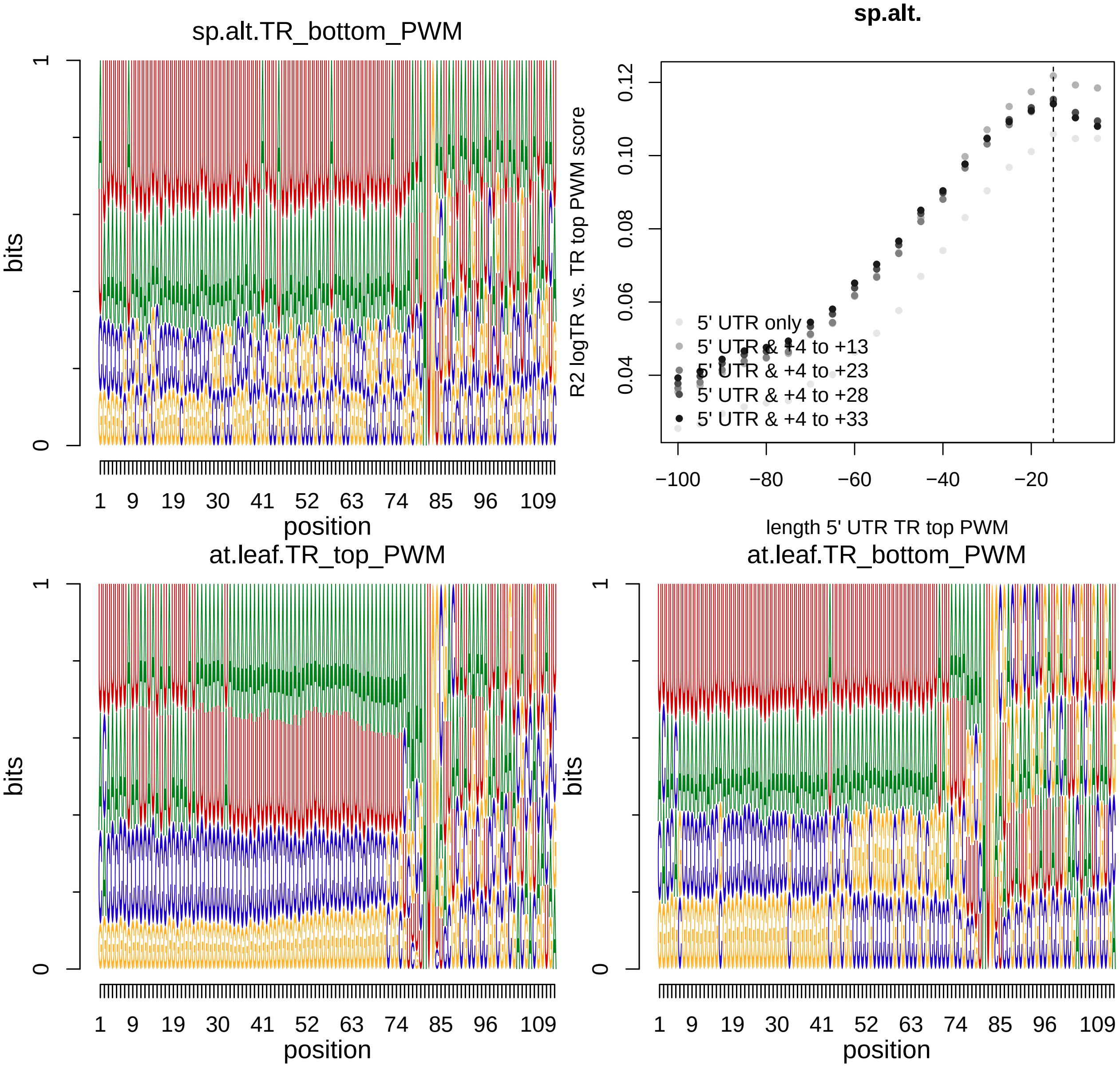
sp.alt.TR_bottom_PWM
bits
position
sp.alt.
R2 logTR vs. TR top PWM score
length 5' UTR TR top PWM
5' UTR only
5' UTR & +4 to +13
5' UTR & +4 to +23
5' UTR & +4 to +28
5' UTR & +4 to +33
at.leaf.TR_top_PWM
bits
position
at.leaf.TR_bottom_PWM
bits
position

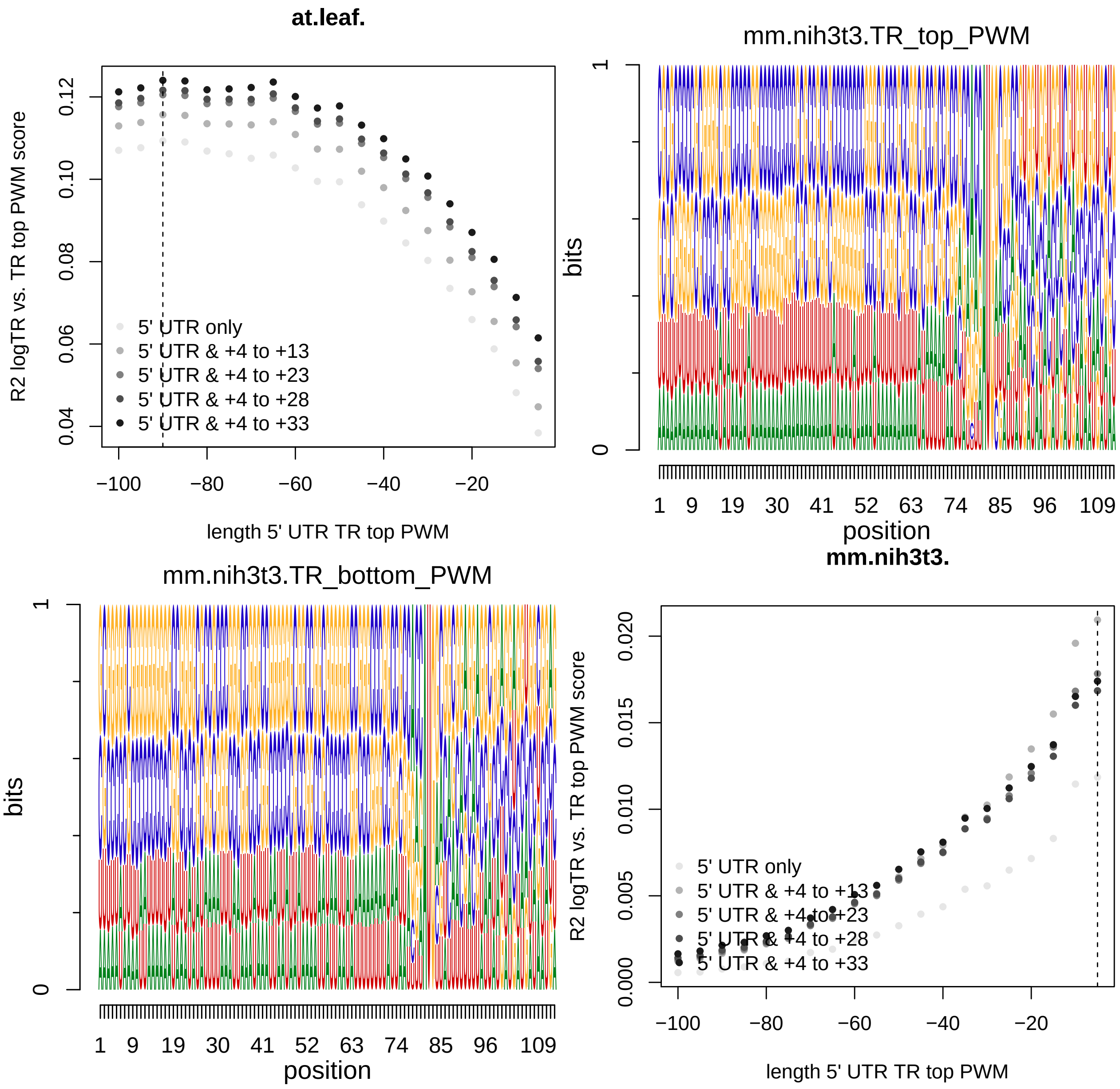
at.leaf.
R2 logTR vs. TR top PWM score
0.04
0.06
0.08
0.10
0.12
−100
−80
−60
−40
−20
length 5' UTR TR top PWM
5' UTR only
5' UTR & +4 to +13
5' UTR & +4 to +23
5' UTR & +4 to +28
5' UTR & +4 to +33
mm.nih3t3.TR_top_PWM
bits
0
1
1 9 19 30 41 52 63 74 85 96 109
position
mm.nih3t3.TR_bottom_PWM
bits
0
1
1 9 19 30 41 52 63 74 85 96 109
position
mm.nih3t3.
R2 logTR vs. TR top PWM score
0.000
0.005
0.010
0.015
0.020
−100
−80
−60
−40
−20
length 5' UTR TR top PWM
5' UTR only
5' UTR & +4 to +13
5' UTR & +4 to +23
5' UTR & +4 to +28
5' UTR & +4 to +33

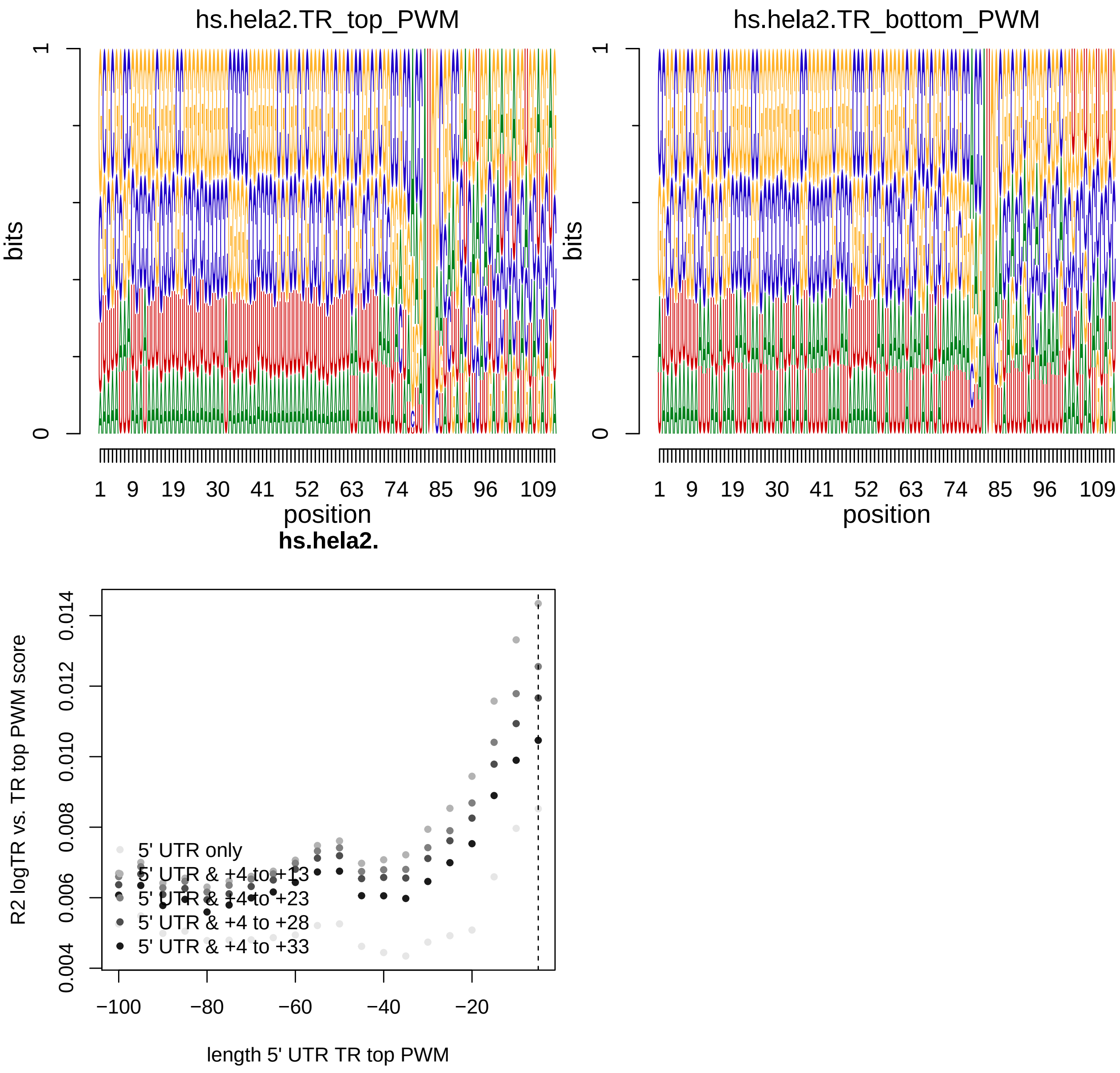


#### Figure 7 - figure supplement 2. R2s between uTICE (PWM only or PWM + selected di/tri nucleotides), dTICE (PWM only or PWM + selected di/tri nucleotides), or TICE (PWM only or PWM + selected di/tri nucleotides)

##### the following code only needs to be run once

```
R2s_uTICE_dTICE_TICE <- NULL
rownames_R2s_uTICE_dTICE_TICE <- NULL
colnames_R2s_uTICE_dTICE_TICE <- c("uTICE_PWM", "uTICE_PWM+di/tri",
    "dTICE_PWM", "dTICE_PWM+di/tri", "TICE_PWM", "TICE_PWM+di/tri")
```

```
for (species in names(species_tissue_list)) {
    for (species_tissue in species_tissue_list[[species]]) {
        if (species_tissue != "") {
            tissue_dot <- paste0(".", species_tissue)
            tissue_us <- paste0("_", species_tissue)
        } else {
            tissue_dot <- ""
            tissue_us <- ""
        }

        uTICE_features <- get(paste0(species, tissue_dot, ".uTICE_features"))[[1]]
        dTICE_features <- get(paste0(species, tissue_dot, ".dTICE_features"))[[1]]

        uTICE_PWM_features <- uTICE_features[, c(which(grepl("^PWM",
            colnames(uTICE_features))), which(grepl("^Intercept",
            colnames(uTICE_features))))]
        dTICE_PWM_features <- dTICE_features[, c(which(grepl("^PWM",
            colnames(dTICE_features))), which(grepl("^Intercept",
            colnames(dTICE_features))))]

        TICE_features <- cbind(uTICE_features, dTICE_features)
        TICE_PWM_features <- cbind(uTICE_PWM_features, dTICE_PWM_features)

        R2s_uTICE_dTICE_TICE <- rbind(R2s_uTICE_dTICE_TICE, c(summary(lm(get(paste0(species,
            tissue_dot, ".log10.TR")) ~ as.matrix(uTICE_PWM_features)))$r.squared,
            summary(lm(get(paste0(species, tissue_dot, ".log10.TR")) ~
                as.matrix(uTICE_features)))$r.squared, summary(lm(get(paste0(species,
                tissue_dot, ".log10.TR")) ~ as.matrix(dTICE_PWM_features)))$r.squared,
            summary(lm(get(paste0(species, tissue_dot, ".log10.TR")) ~
                as.matrix(dTICE_features)))$r.squared, summary(lm(get(paste0(species,
                tissue_dot, ".log10.TR")) ~ as.matrix(TICE_PWM_features)))$r.squared,
            summary(lm(get(paste0(species, tissue_dot, ".log10.TR")) ~
                as.matrix(TICE_features)))$r.squared))
        rownames_R2s_uTICE_dTICE_TICE <- c(rownames_R2s_uTICE_dTICE_TICE,
            paste0(species, tissue_dot))
    }
}
rownames(R2s_uTICE_dTICE_TICE) <- rownames_R2s_uTICE_dTICE_TICE
colnames(R2s_uTICE_dTICE_TICE) <- colnames_R2s_uTICE_dTICE_TICE

write.csv(R2s_uTICE_dTICE_TICE, file = "tables/R2s_uTICE_dTICE_TICE.csv",
    quote = F)
```

#### plot

```
R2s_uTICE_dTICE_TICE <- read.csv(file = "tables/R2s_uTICE_dTICE_TICE.csv")
colnames(R2s_uTICE_dTICE_TICE)[1] <- "species.tissue"

# only keep the exemplar tissues
for (species in names(species_tissue_list)) {
    for (i in 1:length(species_exemplar_tissue_list[[species]])) {
```

```
        species_tissue <- species_exemplar_tissue_list[[species]][i]
        if (species_tissue != "") {
            tissue_dot <- paste0(".", species_tissue)
            tissue_us <- paste0("_", species_tissue)
        } else {
            tissue_dot <- ""
            tissue_us <- ""
        }
        tmp_for_plot <- as.data.frame(matrix(R2s_uTICE_dTICE_TICE[R2s_uTICE_dTICE_TICE$species.tissue ==
            paste0(species, tissue_dot), 2:ncol(R2s_uTICE_dTICE_TICE)],
            ncol = 1))
        colnames(tmp_for_plot) <- "R2"
        tmp_for_plot$Feature <- factor(rep(c("uTICE", "dTICE",
            "TICE"), each = 2), levels = c("uTICE", "dTICE",
            "TICE"))
        tmp_for_plot$ifPWMonly <- factor(rep(c("PWM", "PWM+di/tri"),
            3), levels = c("PWM", "PWM+di/tri"))
        tmp_for_plot$R2 <- as.numeric(tmp_for_plot$R2)

        theme_set(theme_bw())
        print(ggplot(tmp_for_plot, aes(x = Feature, y = R2, fill = ifPWMonly)) +
            geom_bar(stat = "identity", position = position_dodge()) +
            scale_fill_manual(values = c("#AECDE1", "#447DB2")) +
            theme(axis.text.x = element_text(angle = 45, vjust = 1,
                hjust = 1)) + ggtitle(paste0(species, tissue_dot)))
    }
}
```

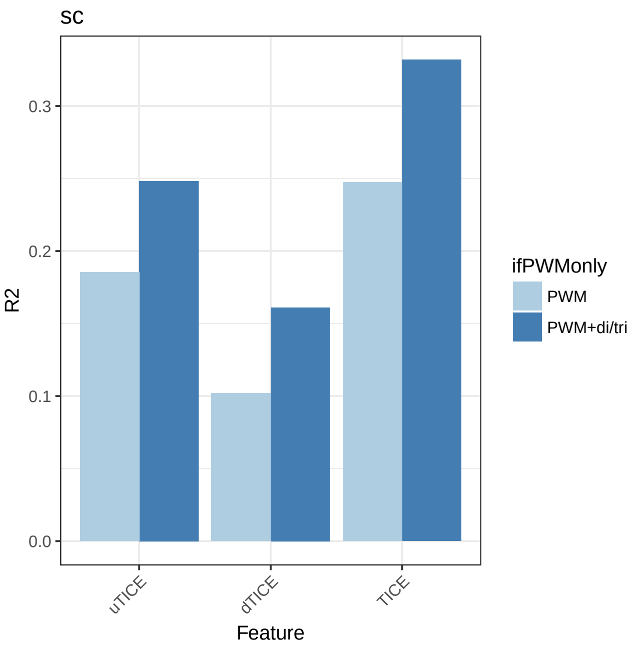


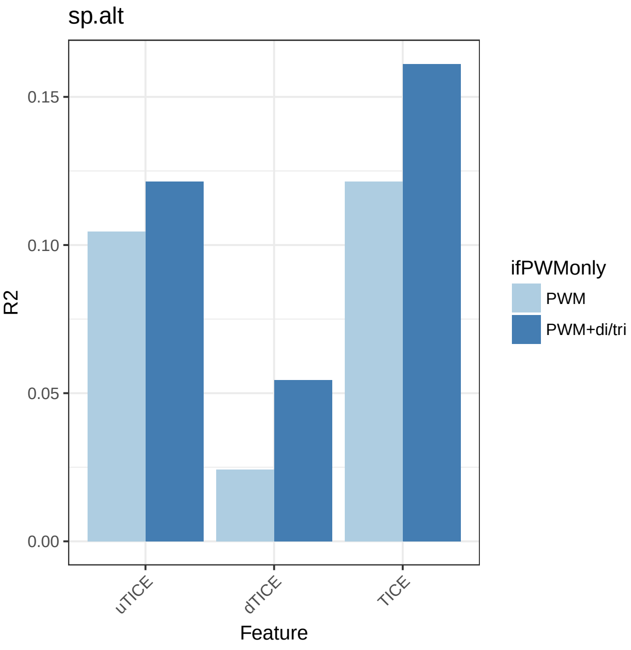

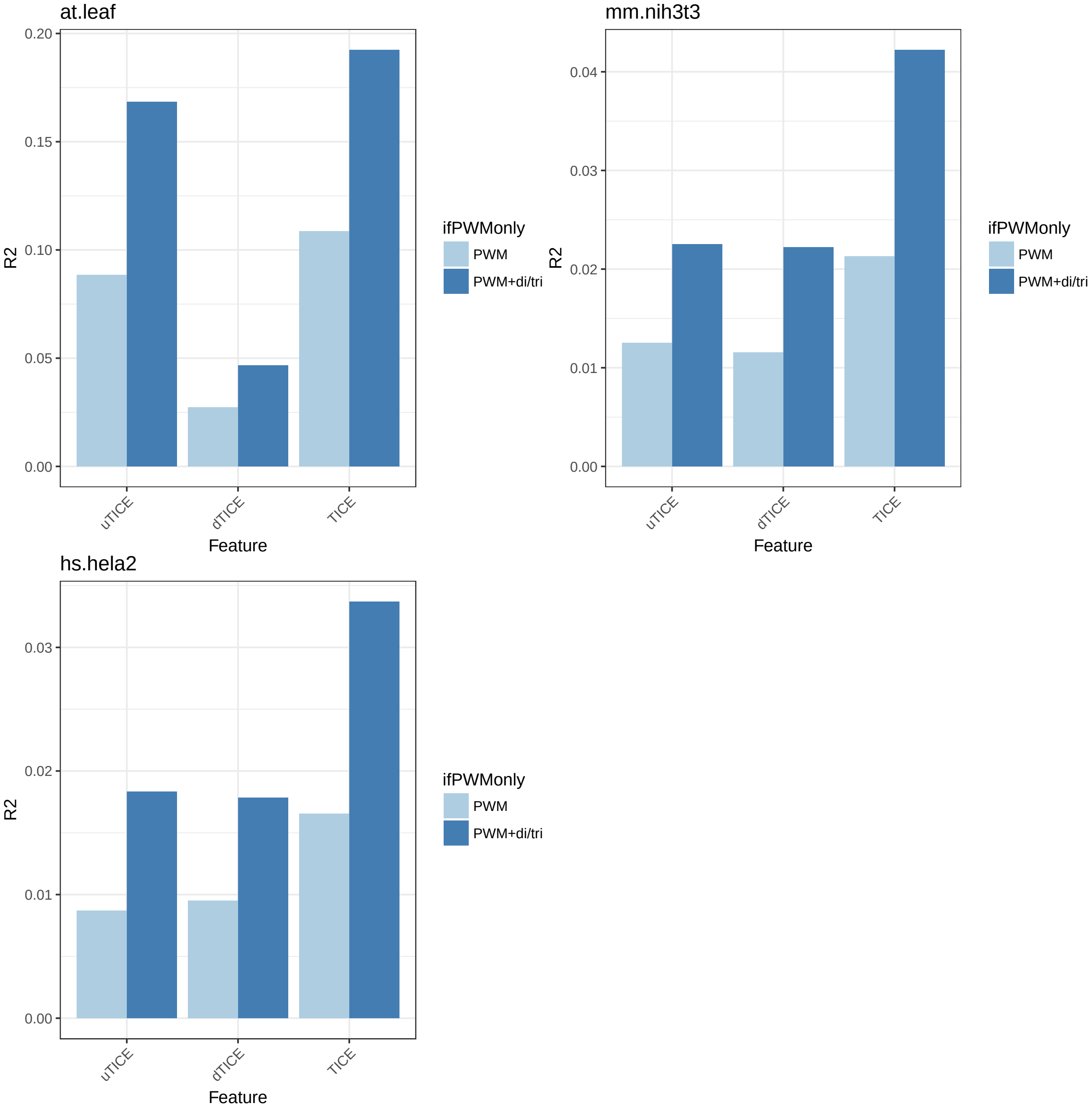


### Figure 8. R2 of 5'ofTICE vs. R2 of TICE vs. R2 of 5'motifs (5'ofTICE + TICE)

#### the following code only needs to be run once

```
R2s_5ofTICE_TICE_5motifs <- NULL
rownames_R2s_5ofTICE_TICE_5motifs <- NULL
colnames_R2s_5ofTICE_TICE_5motifs <- c("5'ofTICE", "TICE", "5'motifs")

for (species in names(species_tissue_list)) {
```

```
    for (species_tissue in species_tissue_list[[species]]) {
        if (species_tissue != "") {
            tissue_dot <- paste0(".", species_tissue)
            tissue_us <- paste0("_", species_tissue)
        } else {
            tissue_dot <- ""
            tissue_us <- ""
        }

        R2s_5ofTICE_TICE_5motifs <- rbind(R2s_5ofTICE_TICE_5motifs,
            c(summary(lm(get(paste0(species, tissue_dot, ".log10.TR")) ~
                as.matrix(get(paste0(species, tissue_dot, ".5ofTICE_features"))[[1]])))$r.squared,
                summary(lm(get(paste0(species, tissue_dot, ".log10.TR")) ~
                  as.matrix(cbind(get(paste0(species, tissue_dot,
                    ".uTICE_features"))[[1]], get(paste0(species,
                    tissue_dot, ".dTICE_features"))[[1]]))))$r.squared,
                summary(lm(get(paste0(species, tissue_dot, ".log10.TR")) ~
                  as.matrix(cbind(get(paste0(species, tissue_dot,
                    ".uTICE_features"))[[1]], get(paste0(species,
                    tissue_dot, ".dTICE_features"))[[1]], get(paste0(species,
                    tissue_dot, ".5ofTICE_features"))[[1]]))))$r.squared))
        rownames_R2s_5ofTICE_TICE_5motifs <- c(rownames_R2s_5ofTICE_TICE_5motifs,
            paste0(species, tissue_dot))
    }
}
rownames(R2s_5ofTICE_TICE_5motifs) <- rownames_R2s_5ofTICE_TICE_5motifs
colnames(R2s_5ofTICE_TICE_5motifs) <- colnames_R2s_5ofTICE_TICE_5motifs

write.csv(R2s_5ofTICE_TICE_5motifs, file = "tables/R2s_5ofTICE_TICE_5motifs.csv",
    quote = F)
```

### plot

```
R2s_5ofTICE_TICE_5motifs <- read.csv(file = "tables/R2s_5ofTICE_TICE_5motifs.csv")
colnames(R2s_5ofTICE_TICE_5motifs)[1] <- "species.tissue"

# only keep the exemplar tissues
R2s_5ofTICE_TICE_5motifs_for_plot <- setNames(melt(R2s_5ofTICE_TICE_5motifs[R2s_5ofTICE_TICE_5motifs$spe[illegible]
    species_exemplar_tissue_names, ]), c("Species_Tissue", "Feature",
    "R2"))
```

```
## Using species.tissue as id variables
```

```
for (species in names(species_tissue_list)) {
    for (i in 1:length(species_exemplar_tissue_list[[species]])) {
        species_tissue <- species_exemplar_tissue_list[[species]][i]
        if (species_tissue != "") {
            tissue_dot <- paste0(".", species_tissue)
            tissue_us <- paste0("_", species_tissue)
        } else {
            tissue_dot <- ""
            tissue_us <- ""
        }
```

```
        tmp_for_plot <- R2s_5ofTICE_TICE_5motifs_for_plot[R2s_5ofTICE_TICE_5motifs_for_plot$Species_Tis
            paste0(species, tissue_dot), ]

        tmp_for_plot$Feature <- factor(tmp_for_plot$Feature,
            levels = c("X5.ofTICE", "TICE", "X5.motifs"))

        theme_set(theme_bw())
        print(ggplot(tmp_for_plot, aes(x = Feature, y = R2, fill = Species_Tissue)) +
            geom_bar(stat = "identity", position = position_dodge()) +
            scale_fill_manual(values = "#447DB2") + theme(axis.text.x = element_text(angle = 45,
            vjust = 1, hjust = 1)) + ggtitle(paste0(species,
            tissue_dot)))
    }
}
```

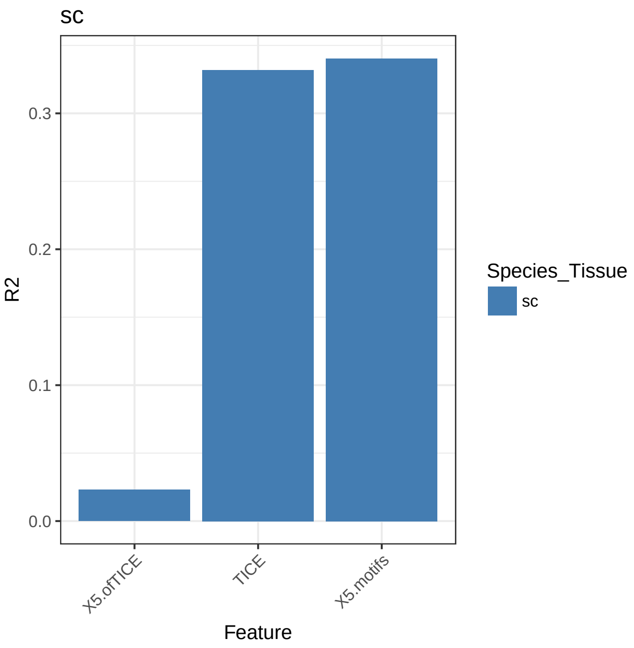


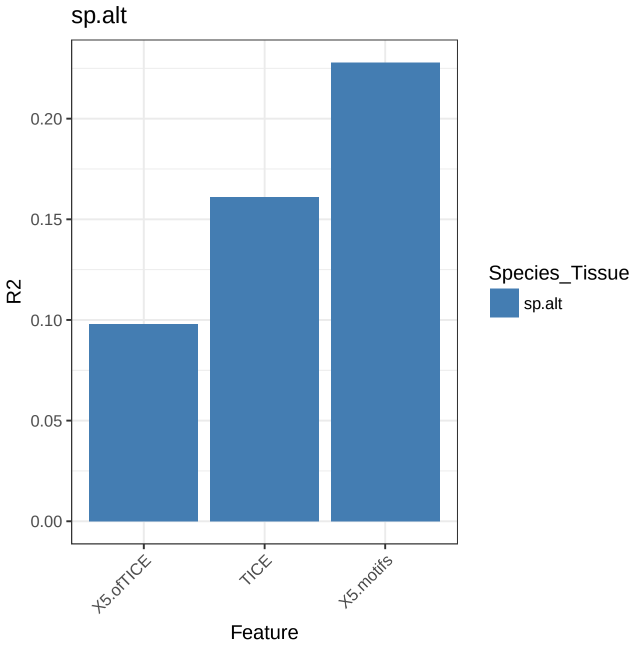

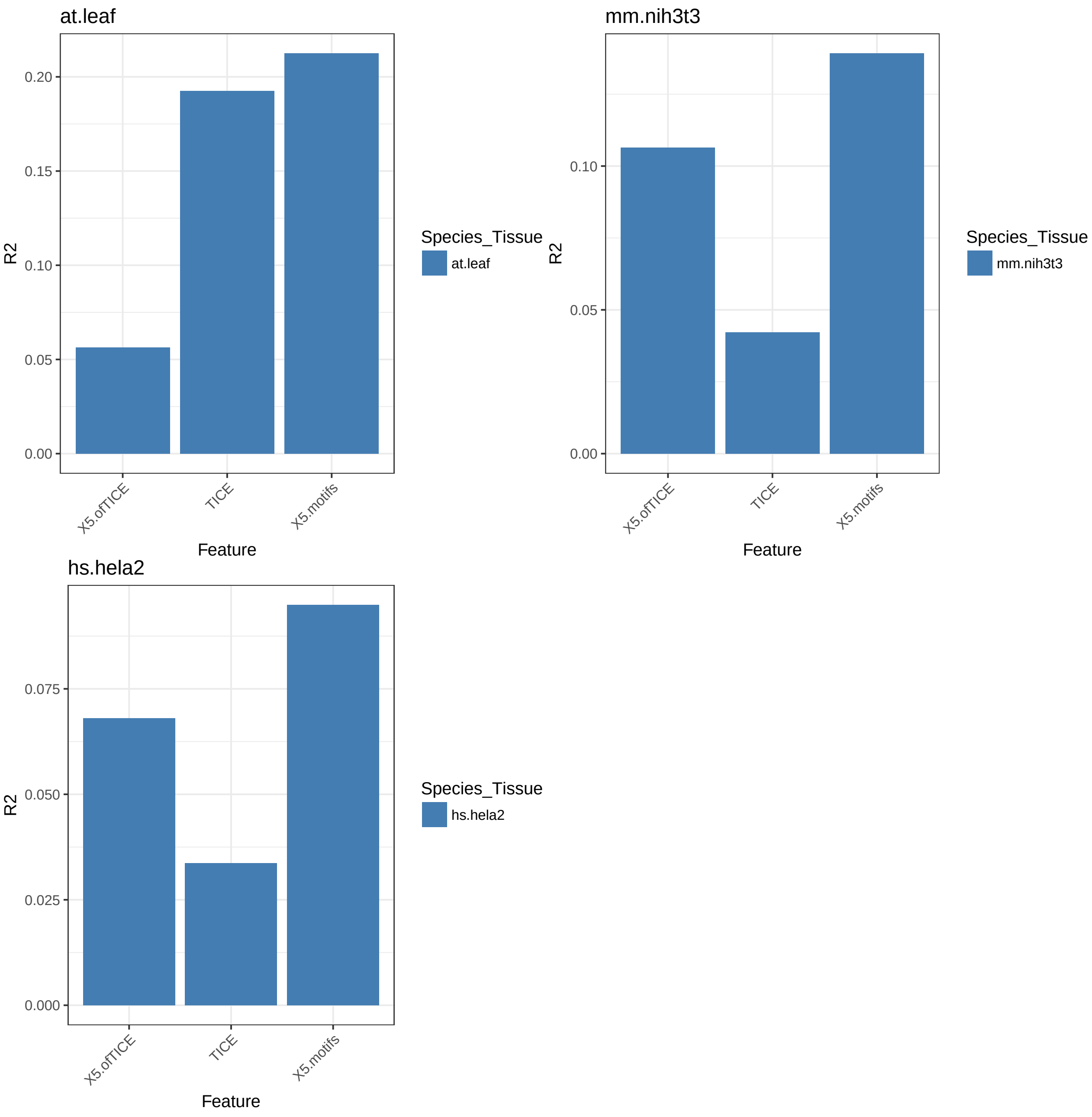


#### Figure 8 - figure supplement 1. 5' cap PWMs and R2s of various PWM boundaries

##### the following code only needs to be run once

```
for (species in names(species_exemplar_tissue_list)) {
    for (species_tissue in species_exemplar_tissue_list[[species]]) {
        if (species_tissue != "") {
            species.genes <- get(paste(species, species_tissue,
                "genes", sep = "."))
```

```
        species.UTR_seqs <- get(paste(species, species_tissue,
            "UTR_seqs", sep = "."))
        species.log10.TR <- get(paste(species, species_tissue,
            "log10.TR", sep = "."))
        species.TR_top_genes <- get(paste(species, species_tissue,
            "TR_top_genes", sep = "."))
        species.TR_bottom_genes <- get(paste(species, species_tissue,
            "TR_bottom_genes", sep = "."))
        connector <- paste0(species_tissue, ".")
    } else {
        species.genes <- get(paste(species, "genes", sep = "."))
        species.UTR_seqs <- get(paste(species, "UTR_seqs",
            sep = "."))
        species.log10.TR <- get(paste(species, "log10.TR",
            sep = "."))
        species.TR_top_genes <- get(paste(species, "TR_top_genes",
            sep = "."))
        species.TR_bottom_genes <- get(paste(species, "TR_bottom_genes",
            sep = "."))
        connector <- ""
    }
    species.log10.UTR_len <- log(nchar(species.UTR_seqs),
        10)
    if (species != "hs") {
        species.uTICE_len <- species_uTICE_len_list[[species]]
    } else {
        species.uTICE_len <- species_uTICE_len_list[[species]][[species_tissue]]
    }

    ## for the genes in the top and bottom cohort, find the 5' UTR
    ## sequences and CDS sequences
    species.TR_top_UTR_seqs <- species.UTR_seqs[species.TR_top_genes]
    species.TR_bottom_UTR_seqs <- species.UTR_seqs[species.TR_bottom_genes]

    # construct Position Weight Matrices (PWMs) based on the top
    # or bottom 10% TR genes from the 5' end with different
    # lengths, excluding the TICE sequence
    len_breaks <- c(1:10, seq(from = 15, to = 100, by = 5))  # varying UTR from the 5' end; do not

    ## top TR
    species.TR_top_PWM_5_aligned <- UTR_pwm_5_aligned(UTR_seqs = substr(x = species.TR_top_UTR_seqs
        start = 1, stop = nchar(species.TR_top_UTR_seqs) -
            species.uTICE_len), n = max(len_breaks))
    colnames(species.TR_top_PWM_5_aligned) <- c("A", "T",
        "C", "G")
    saveRDS(species.TR_top_PWM_5_aligned, file = paste0("processed_data_w_polyA/5'PWM_R2s/",
        species, ".", connector, "TR_top_PWM_5_aligned.rds"))

    ## bottom TR
    species.TR_bottom_PWM_5_aligned <- UTR_pwm_5_aligned(UTR_seqs = substr(x = species.TR_bottom_UTI
        start = 1, stop = nchar(species.TR_bottom_UTR_seqs) -
            species.uTICE_len), n = max(len_breaks))
    colnames(species.TR_bottom_PWM_5_aligned) <- c("A", "T",
```

```
            "C", "G")
        saveRDS(species.TR_bottom_PWM_5_aligned, file = paste0("processed_data_w_polyA/5'PWM_R2s/",
            species, ".", connector, "TR_bottom_PWM_5_aligned.rds"))

        # PWM scores (aligned at the 5' end)

        ## top TR
        species.TR_top_PWM_5_aligned_score <- sapply(len_breaks,
            FUN = function(i) {
                if (i == 1) {
                  PWM_tmp <- matrix(species.TR_top_PWM_5_aligned[1:i,
                    ], nrow = 1)
                  colnames(PWM_tmp) <- c("A", "T", "C", "G")
                } else {
                  PWM_tmp <- species.TR_top_PWM_5_aligned[1:i,
                    ]
                }
                UTR_pwm_5_aligned_score(PWM_tmp, species.UTR_seqs)
            })
        saveRDS(species.TR_top_PWM_5_aligned_score, file = paste0("processed_data_w_polyA/5'PWM_R2s/",
            species, ".", connector, "TR_top_PWM_5_aligned_score.rds"))
    }
}
```

##### plot

```
# a function to plot a 5' cap PWM
plot_pwm_5_aligned <- function(PWM, title) {
    colnames(PWM) <- c("A", "U", "C", "G")
    PWM_motif <- new("pcm", mat = t(PWM), name = title)
    plot(PWM_motif, ic.scale = F)
}

len_breaks <- c(1:10, seq(from = 15, to = 100, by = 5))

for (species in names(species_exemplar_tissue_list)) {
    for (species_tissue in species_exemplar_tissue_list[[species]]) {
        if (species_tissue != "") {
            species.log10.TR <- get(paste(species, species_tissue,
                "log10.TR", sep = "."))
            connector <- paste0(species_tissue, ".")
        } else {
            species.log10.TR <- get(paste(species, "log10.TR",
                sep = "."))
            connector <- ""
        }

        species.TR_top_PWM_5_aligned <- readRDS(file = paste0("processed_data_w_polyA/5'PWM_R2s/",
            species, ".", connector, "TR_top_PWM_5_aligned.rds"))
        species.TR_bottom_PWM_5_aligned <- readRDS(file = paste0("processed_data_w_polyA/5'PWM_R2s/",
            species, ".", connector, "TR_bottom_PWM_5_aligned.rds"))
        ## plot the PWM matrices
```

```
        plot_pwm_5_aligned(species.TR_top_PWM_5_aligned[1:50,
            ], title = paste0(species, ".", connector, "TR_top_PWM_5_aligned"))
        plot_pwm_5_aligned(species.TR_bottom_PWM_5_aligned[1:50,
            ], title = paste0(species, ".", connector, "TR_bottom_PWM_5_aligned"))

        # PWM scores (aligned at the 5' end)
        species.TR_top_PWM_5_aligned_score <- readRDS(file = paste0("processed_data_w_polyA/5'PWM_R2s/"
            species, ".", connector, "TR_top_PWM_5_aligned_score.rds"))

        # calculate the R2s for genes with different minimum 5' UTR
        # lengths
        species.TR_top_R2s_w_TR <- sapply(1:length(len_breaks),
            FUN = function(idx) {
                R2 <- cor(species.log10.TR, species.TR_top_PWM_5_aligned_score[,
                  idx])^2
                return(R2)
            })

        # plot the R2's
        plot(x = len_breaks, y = species.TR_top_R2s_w_TR, main = paste0(species,
            ".", connector), xlab = "length of 5' UTR TR top PWM (5' aligned)",
            ylab = "R2 logTR vs. TR top PWM score", pch = 20,
            axes = T, col = 1)
    }
}
```

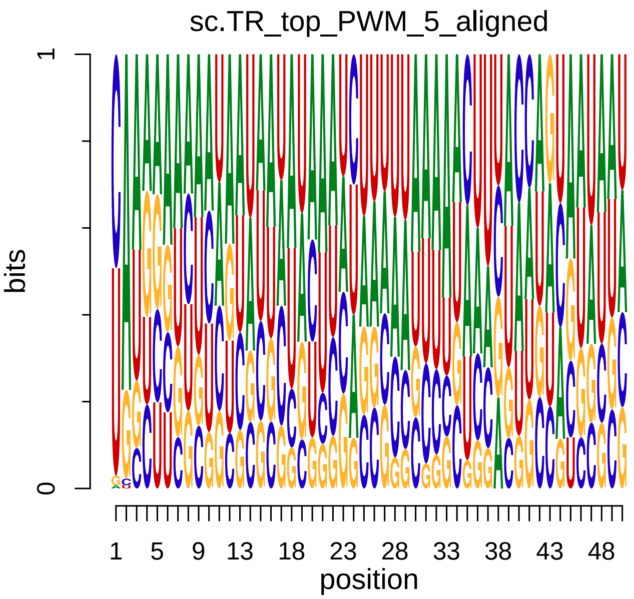


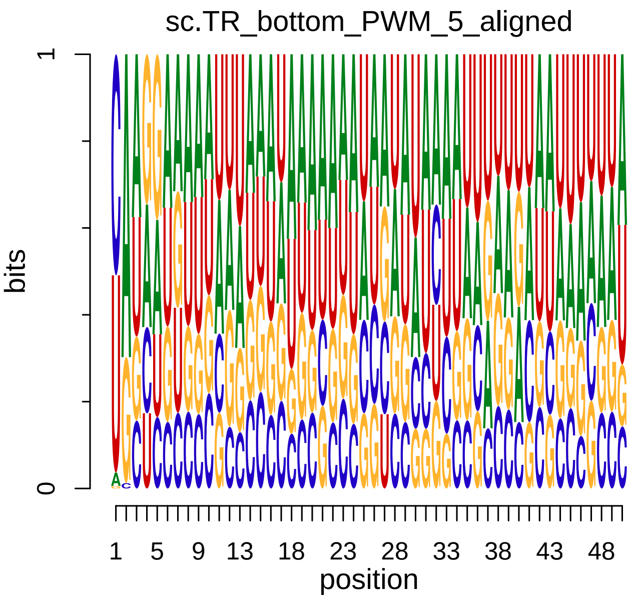

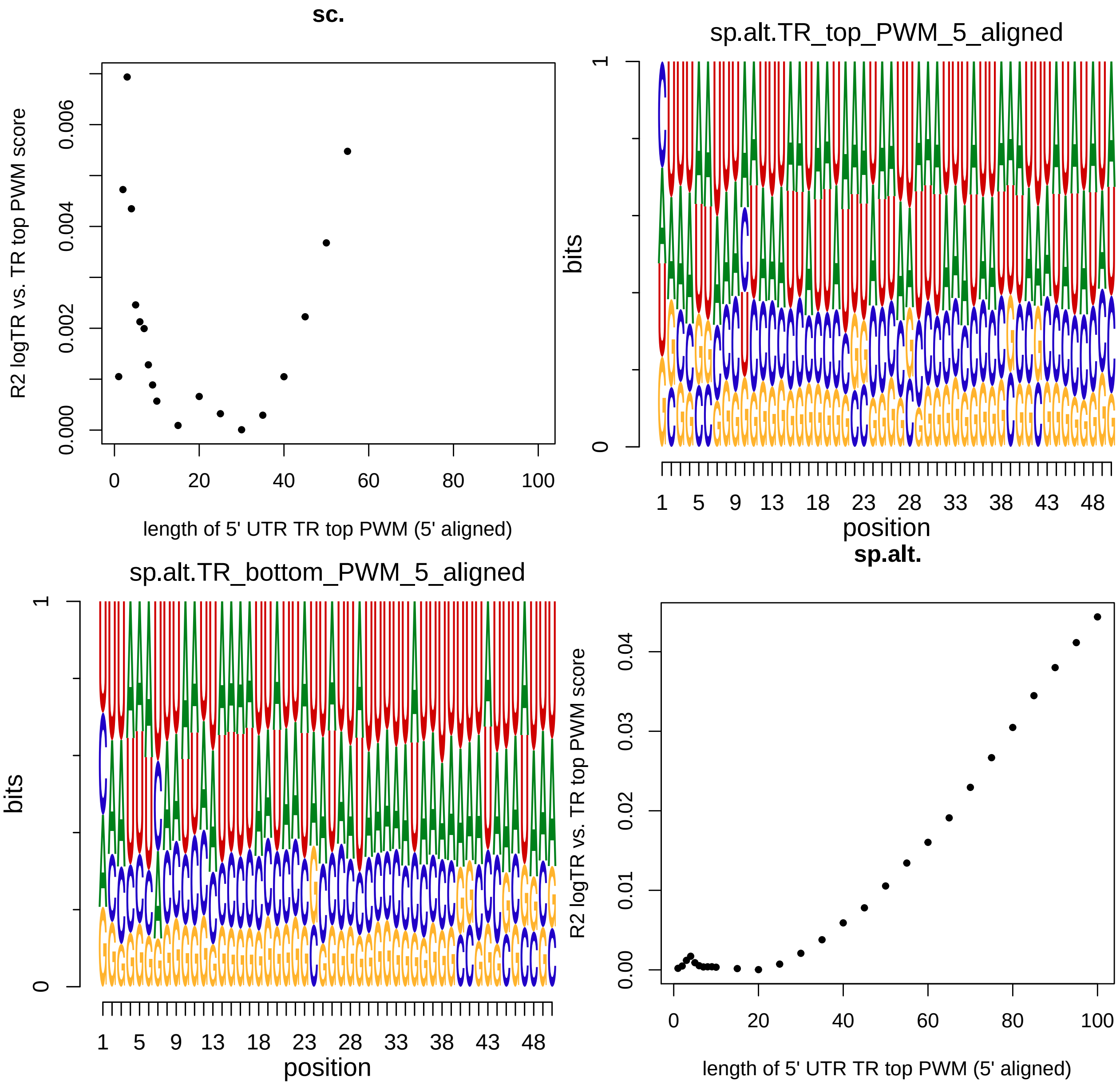
sc.
R2 logTR vs. TR top PWM score
length of 5' UTR TR top PWM (5' aligned)
sp.alt.TR_top_PWM_5_aligned
bits
position
sp.alt.TR_bottom_PWM_5_aligned
bits
position
sp.alt.
R2 logTR vs. TR top PWM score
length of 5' UTR TR top PWM (5' aligned)

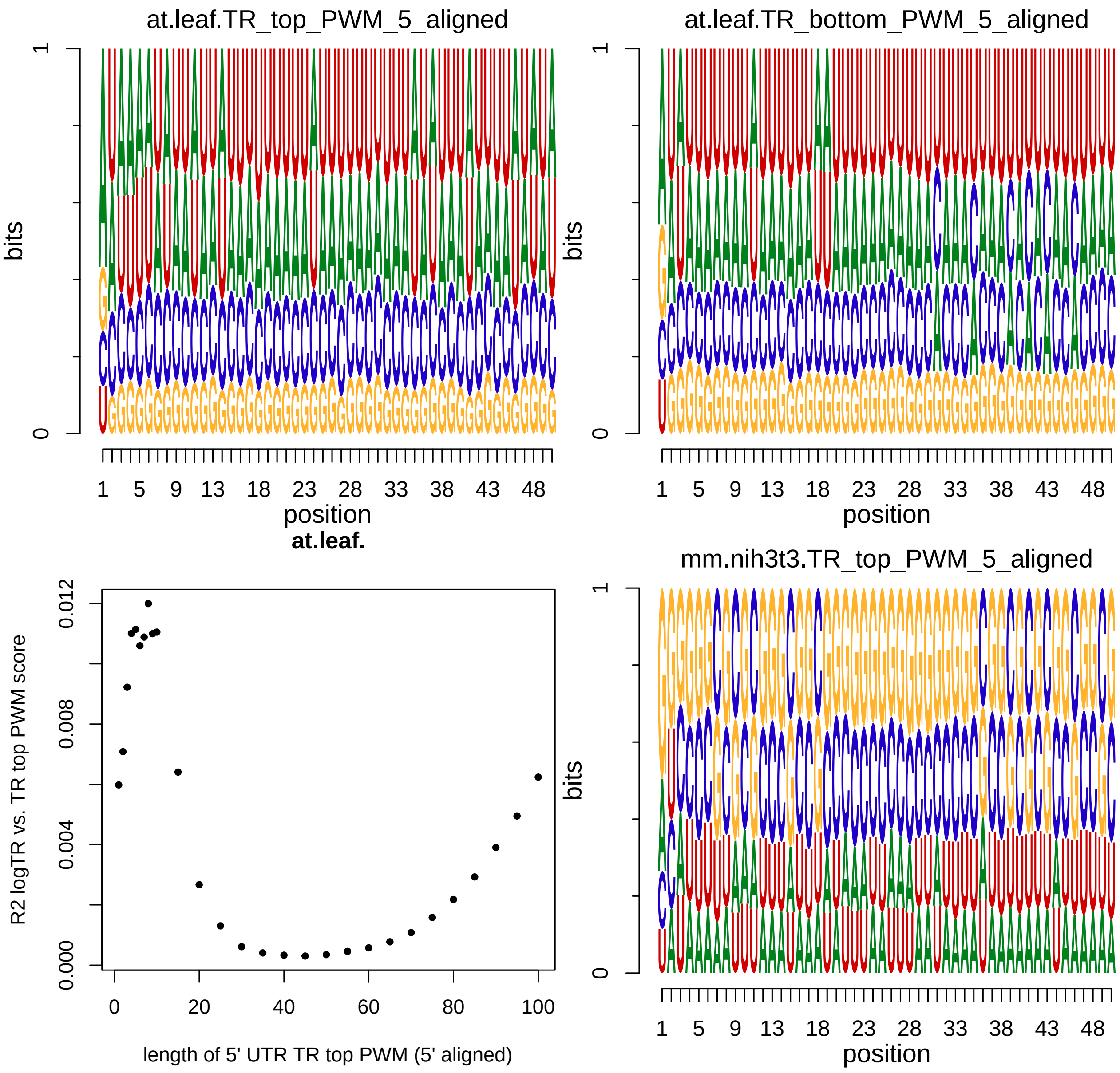
at.leaf.TR_top_PWM_5_aligned
at.leaf.TR_bottom_PWM_5_aligned
bits
position
at.leaf.
R2 logTR vs. TR top PWM score
length of 5' UTR TR top PWM (5' aligned)
mm.nih3t3.TR_top_PWM_5_aligned

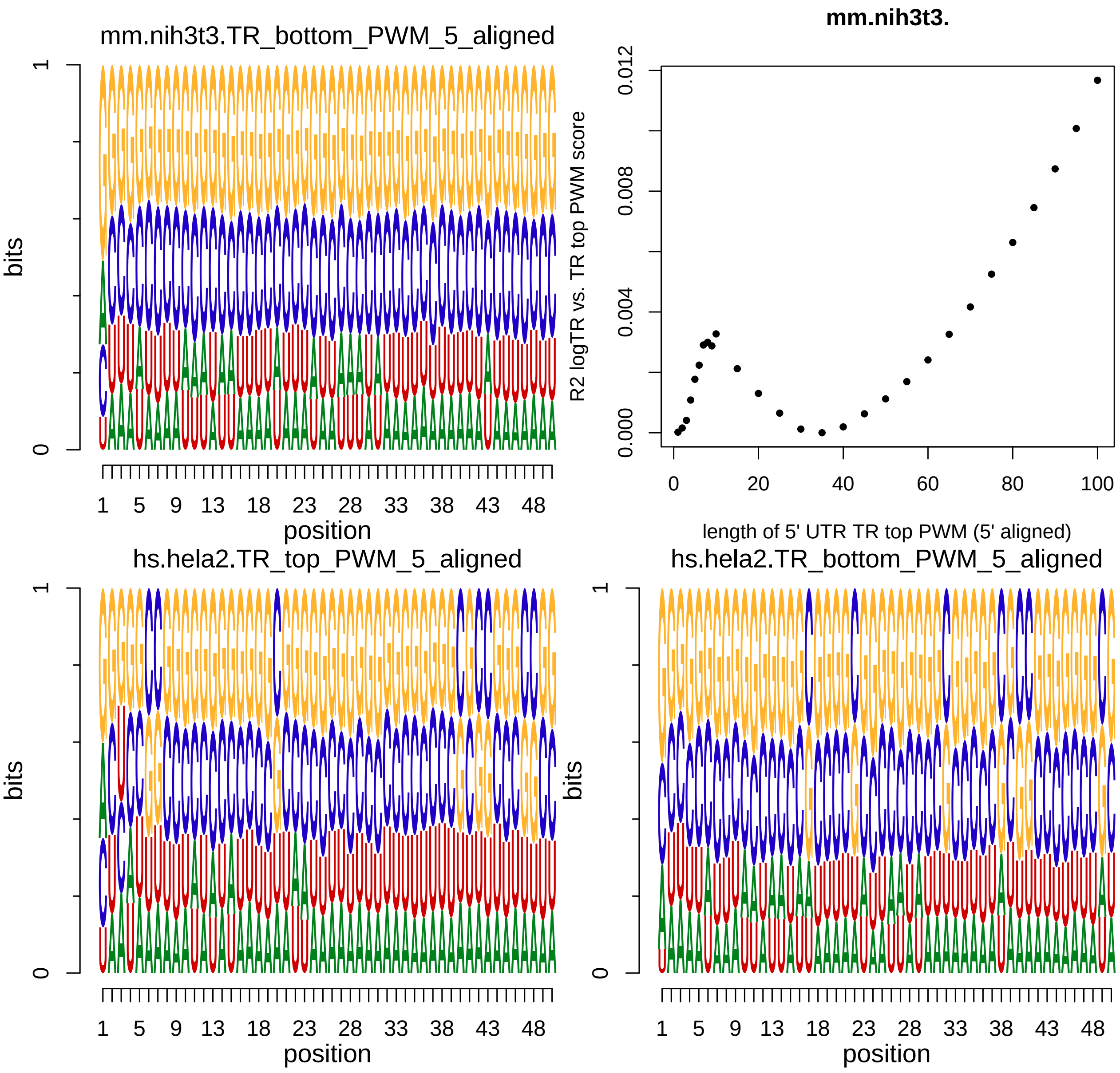
mm.nih3t3.TR_bottom_PWM_5_aligned
bits
0
1
1 5 9 13 18 23 28 33 38 43 48
position
mm.nih3t3.
R2 logTR vs. TR top PWM score
0.000 0.004 0.008 0.012
0 20 40 60 80 100
length of 5' UTR TR top PWM (5' aligned)
hs.hela2.TR_top_PWM_5_aligned
bits
0
1
1 5 9 13 18 23 28 33 38 43 48
position
hs.hela2.TR_bottom_PWM_5_aligned
bits
0
1
1 5 9 13 18 23 28 33 38 43 48
position

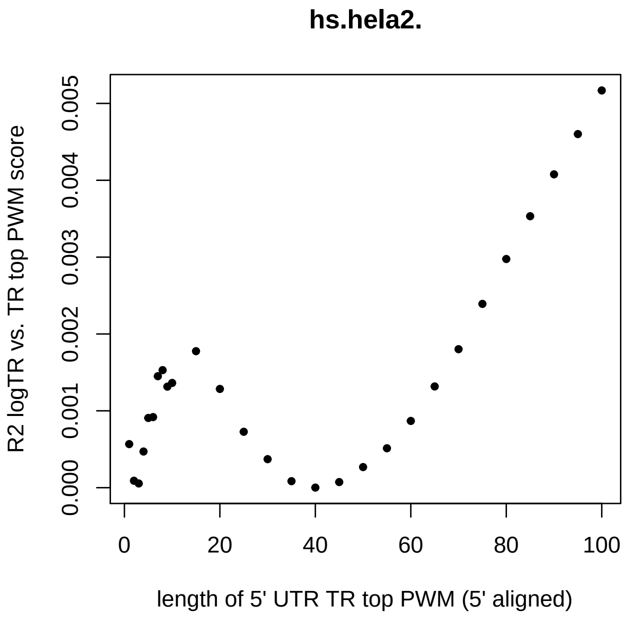


#### Figure 9 - figure supplement 1. Ratios of trinucleotide frequencies between the top and bottom TR genes in the 5' of TICE region, the uTICE region, the dTICE region, and the 3' of TICE region

##### the following code only needs to be run once

```
col_max <- NULL  # record the maximum value to plot in next part

# Remove genes with short 5' UTRs. Keep genes with at least 3
# bp in each region of interest
for (species in names(species_exemplar_tissue_list)) {
    for (species_tissue in species_exemplar_tissue_list[[species]]) {
        if (species_tissue != "") {
            species.genes <- get(paste(species, species_tissue,
                "genes", sep = "."))
            species.UTR_seqs <- get(paste(species, species_tissue,
                "UTR_seqs", sep = "."))
            species.CDS_seqs <- get(paste(species, species_tissue,
                "CDS_seqs", sep = "."))
            species.log10.TR <- get(paste(species, species_tissue,
                "log10.TR", sep = "."))
            species.TR_top_genes <- get(paste(species, species_tissue,
                "TR_top_genes", sep = "."))
            species.TR_bottom_genes <- get(paste(species, species_tissue,
                "TR_bottom_genes", sep = "."))
            connector <- paste0(".", species_tissue)
        } else {
            species.genes <- get(paste(species, "genes", sep = "."))
            species.UTR_seqs <- get(paste(species, "UTR_seqs",
                sep = "."))
```

```
        species.CDS_seqs <- get(paste(species, "CDS_seqs",
            sep = "."))
        species.log10.TR <- get(paste(species, "log10.TR",
            sep = "."))
        species.TR_top_genes <- get(paste(species, "TR_top_genes",
            sep = "."))
        species.TR_bottom_genes <- get(paste(species, "TR_bottom_genes",
            sep = "."))
        connector <- ""
    }

    if (species != "hs") {
        species.uTICE_len <- species_uTICE_len_list[[species]]
        species.dTICE_len <- species_dTICE_len_list[[species]]
    } else {
        species.uTICE_len <- species_uTICE_len_list[[species]][[species_tissue]]
        species.dTICE_len <- species_dTICE_len_list[[species]][[species_tissue]]
    }

    species.UTR_tri_freqs_5ofTICE <- readRDS(file = paste0("processed_data_w_polyA/tri_freqs/",
        species, connector, ".UTR_tri_freqs_5ofTICE.rds"))
    species.UTR_tri_freqs_uTICE <- readRDS(file = paste0("processed_data_w_polyA/tri_freqs/",
        species, connector, ".UTR_tri_freqs_uTICE.rds"))
    species.CDS_tri_freqs_dTICE <- readRDS(file = paste0("processed_data_w_polyA/tri_freqs/",
        species, connector, ".CDS_tri_freqs_dTICE.rds"))
    species.CDS_tri_freqs_3ofTICE <- readRDS(file = paste0("processed_data_w_polyA/tri_freqs/",
        species, connector, ".CDS_tri_freqs_3ofTICE.rds"))

    species.TR_top_UTR_seqs <- species.UTR_seqs[species.TR_top_genes]
    species.TR_bottom_UTR_seqs <- species.UTR_seqs[species.TR_bottom_genes]
    species.TR_top_CDS_seqs <- species.CDS_seqs[species.TR_top_genes]
    species.TR_bottom_CDS_seqs <- species.CDS_seqs[species.TR_bottom_genes]

    # 5' of TICE
    species.UTR_min_len <- species.uTICE_len + 3

    species.UTR_tri_freq_top_bottom_ratios_5ofTICE <- colMeans(species.UTR_tri_freqs_5ofTICE[specie
        species.UTR_min_len], ])/colMeans(species.UTR_tri_freqs_5ofTICE[species.TR_bottom_genes[ncha
        species.UTR_min_len], ])
    species.UTR_tri_freq_top_bottom_ratios_5ofTICE_table <- rbind(matrix(species.UTR_tri_freq_top_bo
        nrow = 4), matrix(species.UTR_tri_freq_top_bottom_ratios_5ofTICE[17:32],
        nrow = 4), matrix(species.UTR_tri_freq_top_bottom_ratios_5ofTICE[33:48],
        nrow = 4), matrix(species.UTR_tri_freq_top_bottom_ratios_5ofTICE[49:64],
        nrow = 4))

    write.csv(species.UTR_tri_freq_top_bottom_ratios_5ofTICE_table,
        file = paste0("tables/tri_freq_tables/", species,
            connector, ".UTR_tri_freq_top_bottom_ratios_5ofTICE_table.csv"),
        quote = F)

    # uTICE
    species.UTR_tri_freq_top_bottom_ratios_uTICE <- colMeans(species.UTR_tri_freqs_uTICE[species.TR_
        3], ])/colMeans(species.UTR_tri_freqs_uTICE[species.TR_bottom_genes[nchar(species.TR_bottom_
```

```
            3], ])
        species.UTR_tri_freq_top_bottom_ratios_uTICE_table <- rbind(matrix(species.UTR_tri_freq_top_bott
            nrow = 4), matrix(species.UTR_tri_freq_top_bottom_ratios_uTICE[17:32],
            nrow = 4), matrix(species.UTR_tri_freq_top_bottom_ratios_uTICE[33:48],
            nrow = 4), matrix(species.UTR_tri_freq_top_bottom_ratios_uTICE[49:64],
            nrow = 4))

        write.csv(species.UTR_tri_freq_top_bottom_ratios_uTICE_table,
            file = paste0("tables/tri_freq_tables/", species,
                connector, ".UTR_tri_freq_top_bottom_ratios_uTICE_table.csv"),
            quote = F)

        # dTICE
        species.CDS_tri_freq_top_bottom_ratios_dTICE <- colMeans(species.CDS_tri_freqs_dTICE[species.TR_
            3], ])/colMeans(species.CDS_tri_freqs_dTICE[species.TR_bottom_genes[nchar(species.TR_bottom_
            3], ])
        species.CDS_tri_freq_top_bottom_ratios_dTICE_table <- rbind(matrix(species.CDS_tri_freq_top_bott
            nrow = 4), matrix(species.CDS_tri_freq_top_bottom_ratios_dTICE[17:32],
            nrow = 4), matrix(species.CDS_tri_freq_top_bottom_ratios_dTICE[33:48],
            nrow = 4), matrix(species.CDS_tri_freq_top_bottom_ratios_dTICE[49:64],
            nrow = 4))
        write.csv(species.CDS_tri_freq_top_bottom_ratios_dTICE_table,
            file = paste0("tables/tri_freq_tables/", species,
                connector, ".CDS_tri_freq_top_bottom_ratios_dTICE_table.csv"),
            quote = F)

        # 3' of TICE
        species.CDS_min_len <- species.dTICE_len + 3

        species.CDS_tri_freq_top_bottom_ratios_3ofTICE <- colMeans(species.CDS_tri_freqs_3ofTICE[species
            species.CDS_min_len], ])/colMeans(species.CDS_tri_freqs_3ofTICE[species.TR_bottom_genes[ncha
            species.CDS_min_len], ])
        species.CDS_tri_freq_top_bottom_ratios_3ofTICE_table <- rbind(matrix(species.CDS_tri_freq_top_bo
            nrow = 4), matrix(species.CDS_tri_freq_top_bottom_ratios_3ofTICE[17:32],
            nrow = 4), matrix(species.CDS_tri_freq_top_bottom_ratios_3ofTICE[33:48],
            nrow = 4), matrix(species.CDS_tri_freq_top_bottom_ratios_3ofTICE[49:64],
            nrow = 4))
        write.csv(species.CDS_tri_freq_top_bottom_ratios_3ofTICE_table,
            file = paste0("tables/tri_freq_tables/", species,
                connector, ".CDS_tri_freq_top_bottom_ratios_3ofTICE_table.csv"),
            quote = F)

        col_max <- max(col_max, max(species.UTR_tri_freq_top_bottom_ratios_5ofTICE_table),
            max(species.UTR_tri_freq_top_bottom_ratios_uTICE_table),
            max(species.CDS_tri_freq_top_bottom_ratios_dTICE_table),
            max(species.CDS_tri_freq_top_bottom_ratios_3ofTICE_table))
    }
}

saveRDS(col_max, file = "processed_data_w_polyA/tri_freq_tables/col_max.rds")
```

#### plot

```
col_max <- readRDS(file="processed_data_w_polyA/tri_freq_tables/col_max.rds")

col_breaks <- c(seq(0, 0.8, length=100),  # for blue
  seq(0.81, 1.25, length=100),           # for white
  seq(1.26, col_max, length=100))       # for red

for (species in names(species_exemplar_tissue_list)) {
    for (species_tissue in species_exemplar_tissue_list[[species]]) {
        if (species_tissue!="") {
            connector <- paste0(".", species_tissue)
        } else {
            connector <- ""
        }

      # 5' of TICE
      species.UTR_tri_freq_top_bottom_ratios_5ofTICE_table <- as.matrix(read.csv(file=paste0("tables/tri

      heatmap.2(species.UTR_tri_freq_top_bottom_ratios_5ofTICE_table,
                cellnote = tri_nts_table,  # same data set for cell labels
                main = paste0(species, connector, " top/bottom ratios\nin the 5ofTICE region"), # heat m
                notecol="black",      # change font color of cell labels to black
                density.info="none",  # turns off density plot inside color legend
                trace="none",         # turns off trace lines inside the heat map
                #margins =c(12,9),     # widens margins around plot
                col=my_palette,       # use on color palette defined earlier
                breaks=col_breaks,    # enable color transition at specified limits
                dendrogram="none",    # only draw a row dendrogram
                Rowv="NA",            # turn off row clustering
                Colv="NA",            # turn off column clustering
                labRow="",            # turn off row labeling
                labCol=""             # turn off column labeling
      )

      # uTICE
      species.UTR_tri_freq_top_bottom_ratios_uTICE_table <- as.matrix(read.csv(file=paste0("tables/tri_

      heatmap.2(species.UTR_tri_freq_top_bottom_ratios_uTICE_table,
                cellnote = tri_nts_table,  # same data set for cell labels
                main = paste0(species, connector, " top/bottom ratios\nin the uTICE region"), # heat ma
                notecol="black",      # change font color of cell labels to black
                density.info="none",  # turns off density plot inside color legend
                trace="none",         # turns off trace lines inside the heat map
                #margins =c(12,9),     # widens margins around plot
                col=my_palette,       # use on color palette defined earlier
                breaks=col_breaks,    # enable color transition at specified limits
                dendrogram="none",    # only draw a row dendrogram
                Rowv="NA",            # turn off row clustering
                Colv="NA",            # turn off column clustering
                labRow="",            # turn off row labeling
                labCol=""             # turn off column labeling
      )
```

```
        # dTICE
        species.CDS_tri_freq_top_bottom_ratios_dTICE_table <- as.matrix(read.csv(file=paste0("tables/tri_

        heatmap.2(species.CDS_tri_freq_top_bottom_ratios_dTICE_table,
                  cellnote = tri_nts_table,  # same data set for cell labels
                  main = paste0(species, connector, " top/bottom ratios\nin the dTICE region"), # heat ma
                  notecol="black",      # change font color of cell labels to black
                  density.info="none",  # turns off density plot inside color legend
                  trace="none",         # turns off trace lines inside the heat map
                  #margins =c(12,9),     # widens margins around plot
                  col=my_palette,       # use on color palette defined earlier
                  breaks=col_breaks,    # enable color transition at specified limits
                  dendrogram="none",    # only draw a row dendrogram
                  Rowv="NA",            # turn off row clustering
                  Colv="NA",            # turn off column clustering
                  labRow="",            # turn off row labeling
                  labCol=""             # turn off column labeling
        )

        # 3' of TICE
        species.CDS_tri_freq_top_bottom_ratios_3ofTICE_table <- as.matrix(read.csv(file=paste0("tables/tri

        heatmap.2(species.CDS_tri_freq_top_bottom_ratios_3ofTICE_table,
                  cellnote = tri_nts_table,  # same data set for cell labels
                  main = paste0(species, connector, " top/bottom ratios\nin the uTICE region"), # heat ma
                  notecol="black",      # change font color of cell labels to black
                  density.info="none",  # turns off density plot inside color legend
                  trace="none",         # turns off trace lines inside the heat map
                  #margins =c(12,9),     # widens margins around plot
                  col=my_palette,       # use on color palette defined earlier
                  breaks=col_breaks,    # enable color transition at specified limits
                  dendrogram="none",    # only draw a row dendrogram
                  Rowv="NA",            # turn off row clustering
                  Colv="NA",            # turn off column clustering
                  labRow="",            # turn off row labeling
                  labCol=""             # turn off column labeling
        )
    }
}
```

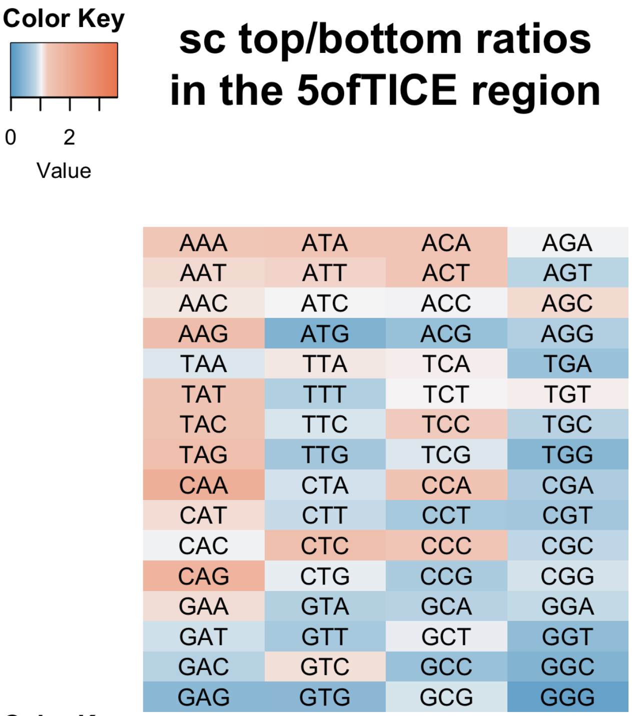
Color Key
0 2
Value
sc top/bottom ratios in the 5ofTICE region
AAA ATA ACA AGA
AAT ATT ACT AGT
AAC ATC ACC AGC
AAG ATG ACG AGG
TAA TTA TCA TGA
TAT TTT TCT TGT
TAC TTC TCC TGC
TAG TTG TCG TGG
CAA CTA CCA CGA
CAT CTT CCT CGT
CAC CTC CCC CGC
CAG CTG CCG CGG
GAA GTA GCA GGA
GAT GTT GCT GGT
GAC GTC GCC GGC
GAG GTG GCG GGG

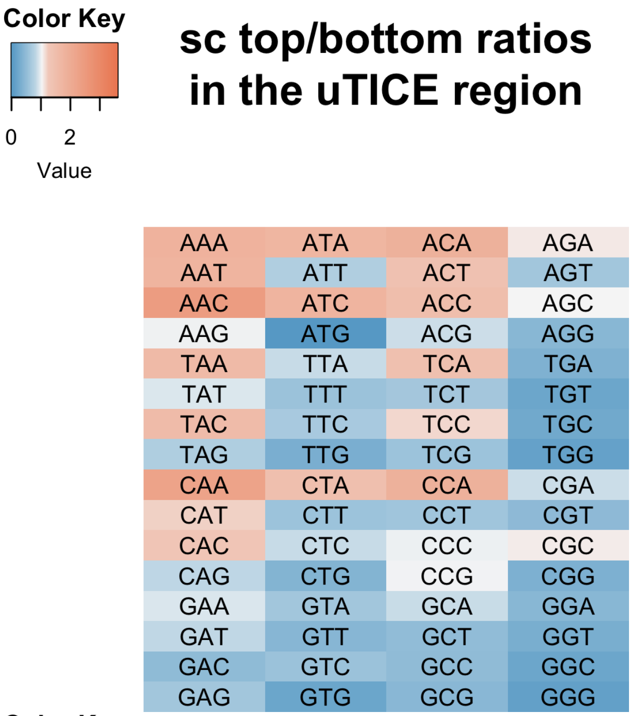
Color Key
0 2
Value
sc top/bottom ratios in the uTICE region
AAA ATA ACA AGA
AAT ATT ACT AGT
AAC ATC ACC AGC
AAG ATG ACG AGG
TAA TTA TCA TGA
TAT TTT TCT TGT
TAC TTC TCC TGC
TAG TTG TCG TGG
CAA CTA CCA CGA
CAT CTT CCT CGT
CAC CTC CCC CGC
CAG CTG CCG CGG
GAA GTA GCA GGA
GAT GTT GCT GGT
GAC GTC GCC GGC
GAG GTG GCG GGG

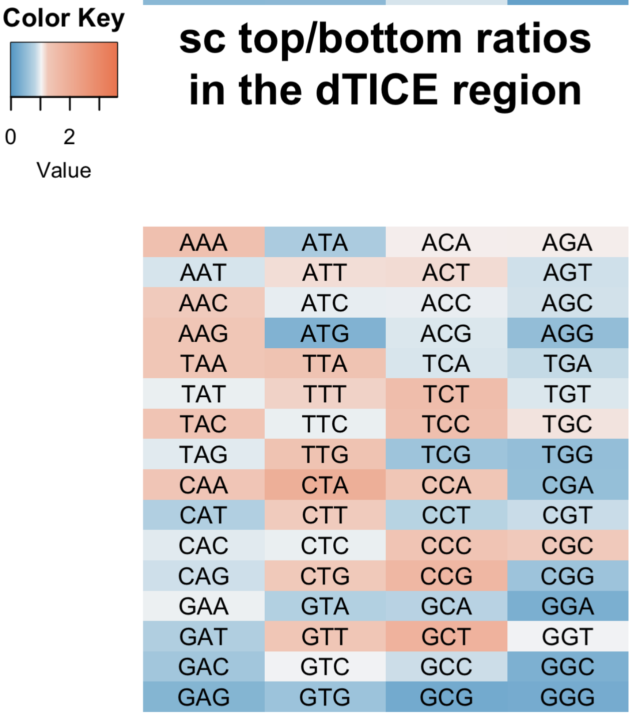
Color Key
0 2
Value
sc top/bottom ratios in the dTICE region
AAA ATA ACA AGA
AAT ATT ACT AGT
AAC ATC ACC AGC
AAG ATG ACG AGG
TAA TTA TCA TGA
TAT TTT TCT TGT
TAC TTC TCC TGC
TAG TTG TCG TGG
CAA CTA CCA CGA
CAT CTT CCT CGT
CAC CTC CCC CGC
CAG CTG CCG CGG
GAA GTA GCA GGA
GAT GTT GCT GGT
GAC GTC GCC GGC
GAG GTG GCG GGG

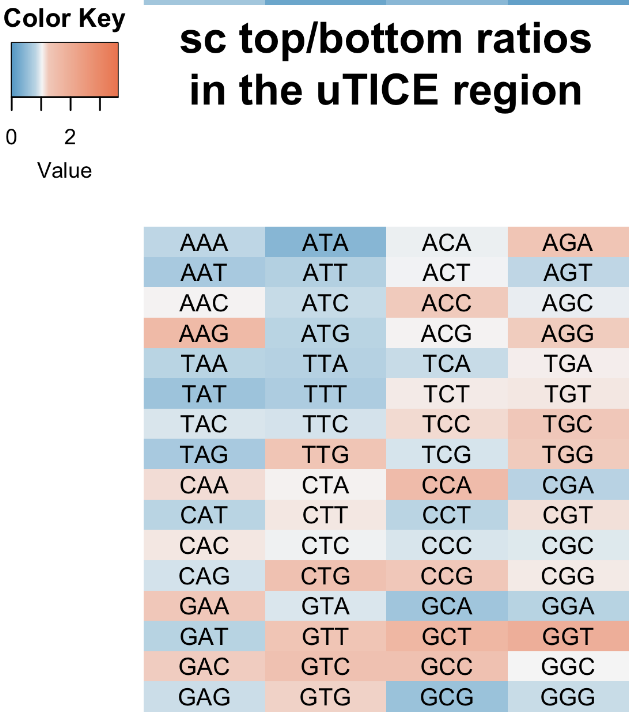
Color Key
0 2
Value
sc top/bottom ratios in the uTICE region
AAA ATA ACA AGA
AAT ATT ACT AGT
AAC ATC ACC AGC
AAG ATG ACG AGG
TAA TTA TCA TGA
TAT TTT TCT TGT
TAC TTC TCC TGC
TAG TTG TCG TGG
CAA CTA CCA CGA
CAT CTT CCT CGT
CAC CTC CCC CGC
CAG CTG CCG CGG
GAA GTA GCA GGA
GAT GTT GCT GGT
GAC GTC GCC GGC
GAG GTG GCG GGG

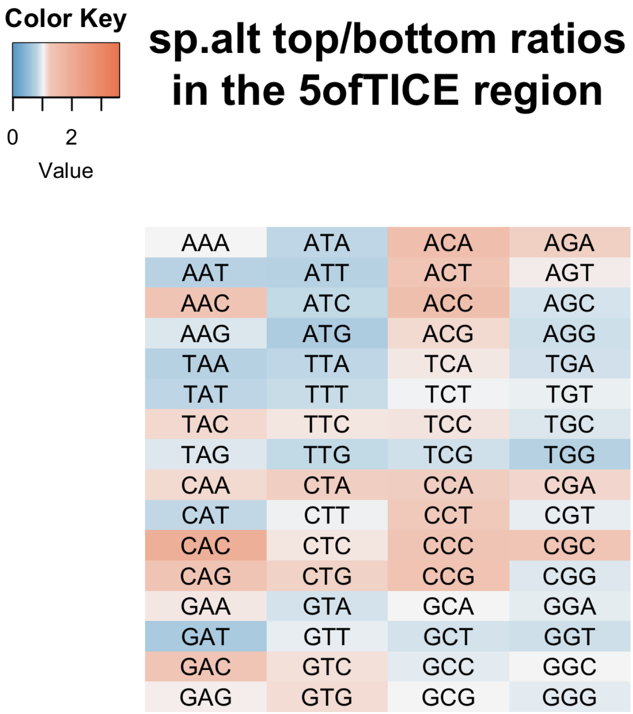
Color Key
0 2
Value
sp.alt top/bottom ratios in the 5ofTICE region
AAA ATA ACA AGA
AAT ATT ACT AGT
AAC ATC ACC AGC
AAG ATG ACG AGG
TAA TTA TCA TGA
TAT TTT TCT TGT
TAC TTC TCC TGC
TAG TTG TCG TGG
CAA CTA CCA CGA
CAT CTT CCT CGT
CAC CTC CCC CGC
CAG CTG CCG CGG
GAA GTA GCA GGA
GAT GTT GCT GGT
GAC GTC GCC GGC
GAG GTG GCG GGG

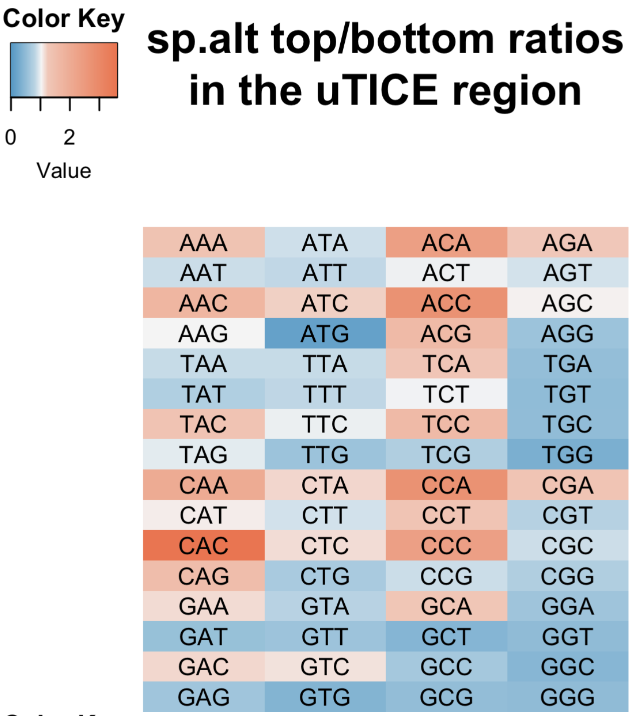
Color Key
0 2
Value
sp.alt top/bottom ratios in the uTICE region
AAA ATA ACA AGA
AAT ATT ACT AGT
AAC ATC ACC AGC
AAG ATG ACG AGG
TAA TTA TCA TGA
TAT TTT TCT TGT
TAC TTC TCC TGC
TAG TTG TCG TGG
CAA CTA CCA CGA
CAT CTT CCT CGT
CAC CTC CCC CGC
CAG CTG CCG CGG
GAA GTA GCA GGA
GAT GTT GCT GGT
GAC GTC GCC GGC
GAG GTG GCG GGG

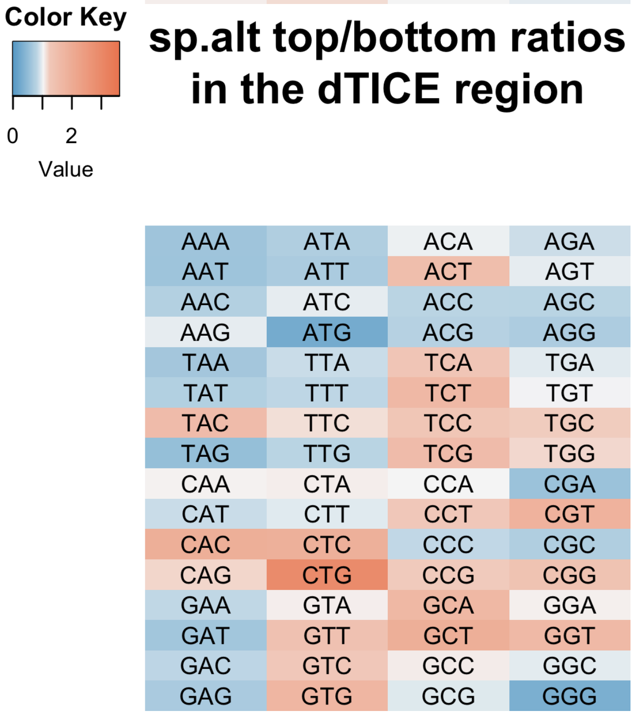
Color Key
0 2
Value
sp.alt top/bottom ratios in the dTICE region
AAA ATA ACA AGA
AAT ATT ACT AGT
AAC ATC ACC AGC
AAG ATG ACG AGG
TAA TTA TCA TGA
TAT TTT TCT TGT
TAC TTC TCC TGC
TAG TTG TCG TGG
CAA CTA CCA CGA
CAT CTT CCT CGT
CAC CTC CCC CGC
CAG CTG CCG CGG
GAA GTA GCA GGA
GAT GTT GCT GGT
GAC GTC GCC GGC
GAG GTG GCG GGG

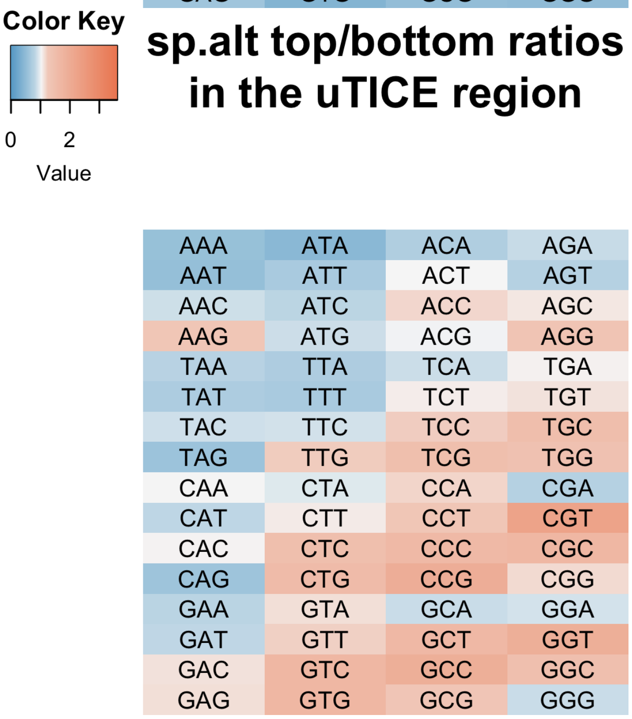
Color Key
0 2
Value
sp.alt top/bottom ratios in the uTICE region
AAA ATA ACA AGA
AAT ATT ACT AGT
AAC ATC ACC AGC
AAG ATG ACG AGG
TAA TTA TCA TGA
TAT TTT TCT TGT
TAC TTC TCC TGC
TAG TTG TCG TGG
CAA CTA CCA CGA
CAT CTT CCT CGT
CAC CTC CCC CGC
CAG CTG CCG CGG
GAA GTA GCA GGA
GAT GTT GCT GGT
GAC GTC GCC GGC
GAG GTG GCG GGG

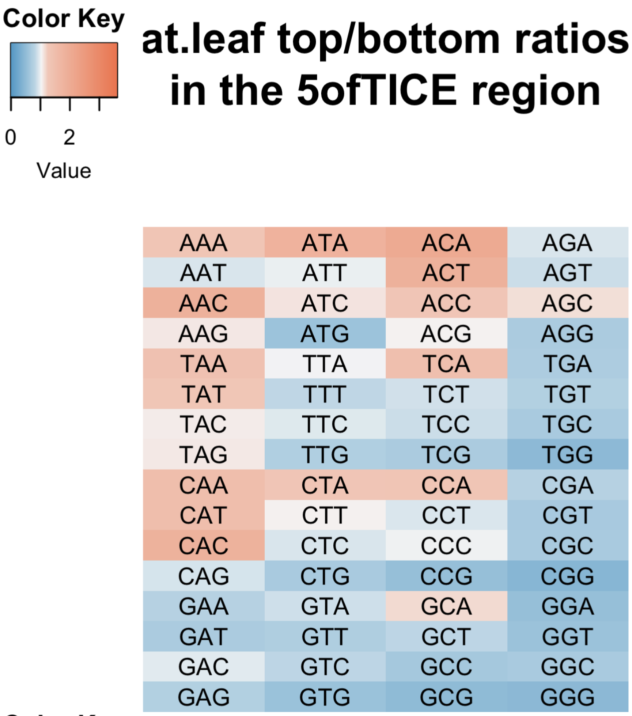
Color Key
0 2
Value
at.leaf top/bottom ratios in the 5ofTICE region
AAA ATA ACA AGA
AAT ATT ACT AGT
AAC ATC ACC AGC
AAG ATG ACG AGG
TAA TTA TCA TGA
TAT TTT TCT TGT
TAC TTC TCC TGC
TAG TTG TCG TGG
CAA CTA CCA CGA
CAT CTT CCT CGT
CAC CTC CCC CGC
CAG CTG CCG CGG
GAA GTA GCA GGA
GAT GTT GCT GGT
GAC GTC GCC GGC
GAG GTG GCG GGG

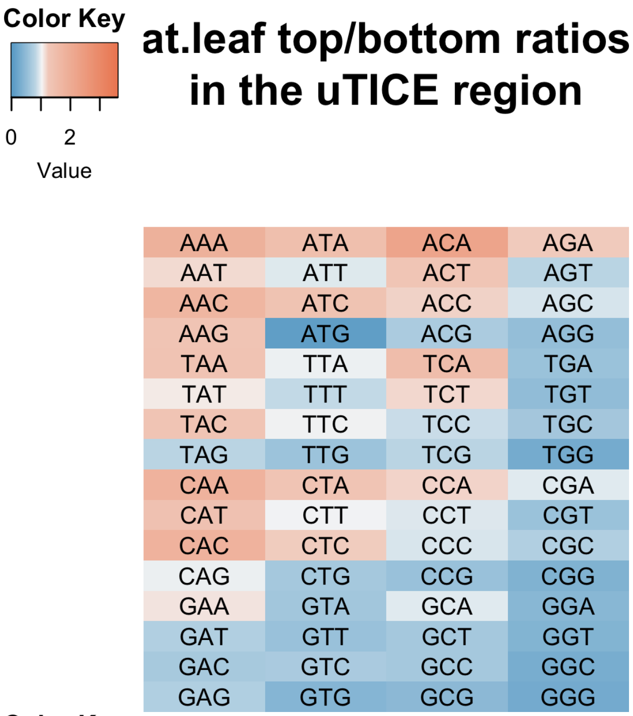
Color Key
0 2
Value
at.leaf top/bottom ratios in the uTICE region
AAA ATA ACA AGA
AAT ATT ACT AGT
AAC ATC ACC AGC
AAG ATG ACG AGG
TAA TTA TCA TGA
TAT TTT TCT TGT
TAC TTC TCC TGC
TAG TTG TCG TGG
CAA CTA CCA CGA
CAT CTT CCT CGT
CAC CTC CCC CGC
CAG CTG CCG CGG
GAA GTA GCA GGA
GAT GTT GCT GGT
GAC GTC GCC GGC
GAG GTG GCG GGG

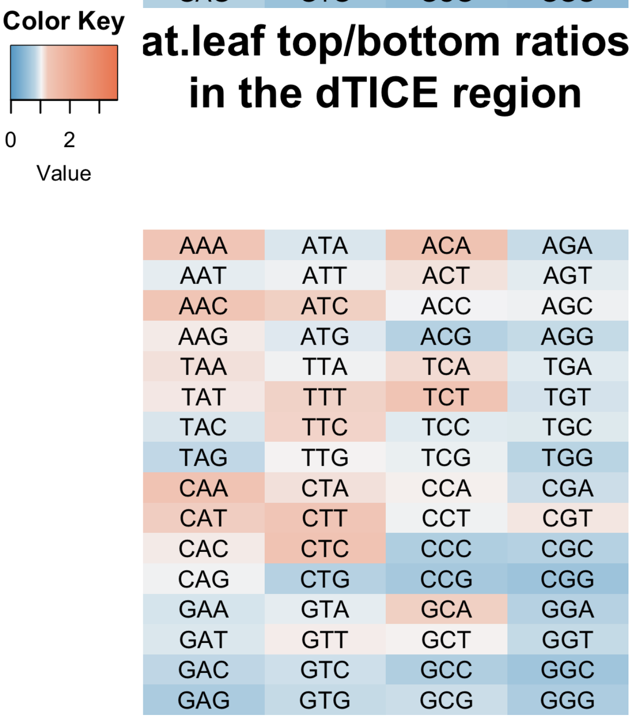
Color Key
0 2
Value
at.leaf top/bottom ratios in the dTICE region
AAA ATA ACA AGA
AAT ATT ACT AGT
AAC ATC ACC AGC
AAG ATG ACG AGG
TAA TTA TCA TGA
TAT TTT TCT TGT
TAC TTC TCC TGC
TAG TTG TCG TGG
CAA CTA CCA CGA
CAT CTT CCT CGT
CAC CTC CCC CGC
CAG CTG CCG CGG
GAA GTA GCA GGA
GAT GTT GCT GGT
GAC GTC GCC GGC
GAG GTG GCG GGG

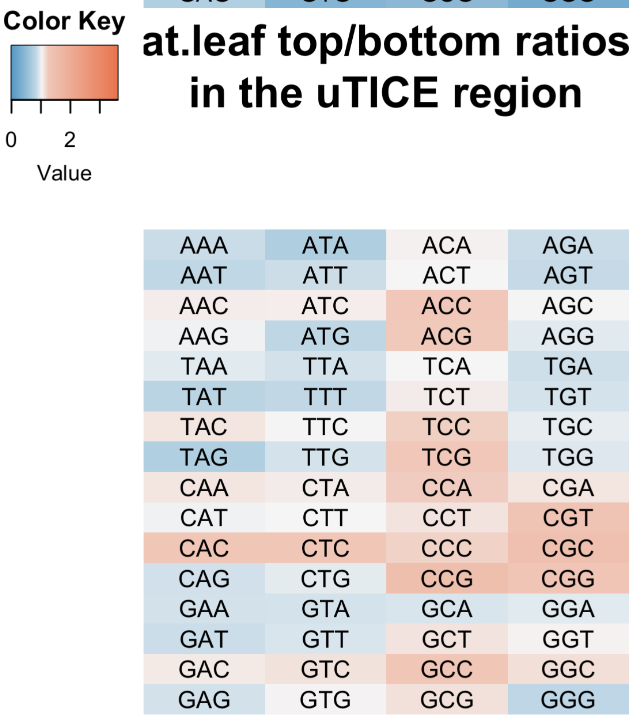
Color Key
0 2
Value
at.leaf top/bottom ratios in the uTICE region
AAA ATA ACA AGA
AAT ATT ACT AGT
AAC ATC ACC AGC
AAG ATG ACG AGG
TAA TTA TCA TGA
TAT TTT TCT TGT
TAC TTC TCC TGC
TAG TTG TCG TGG
CAA CTA CCA CGA
CAT CTT CCT CGT
CAC CTC CCC CGC
CAG CTG CCG CGG
GAA GTA GCA GGA
GAT GTT GCT GGT
GAC GTC GCC GGC
GAG GTG GCG GGG

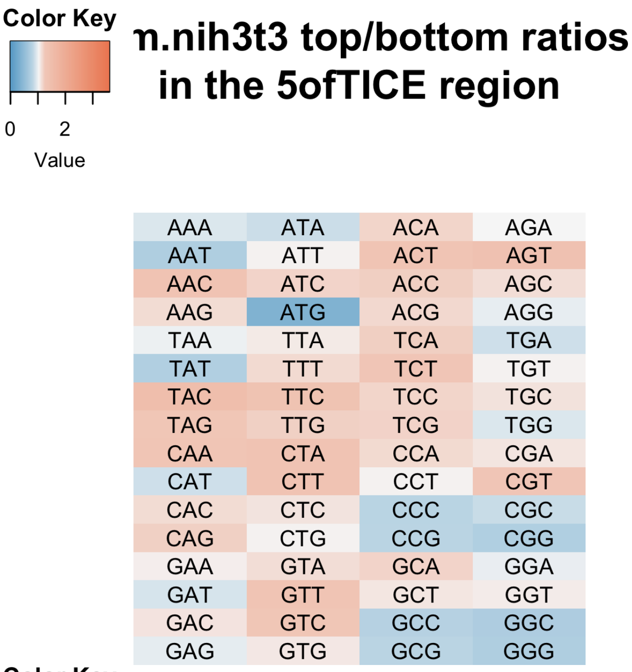
Color Key
0
2
Value
n.nih3t3 top/bottom ratios in the 5ofTICE region
AAA ATA ACA AGA
AAT ATT ACT AGT
AAC ATC ACC AGC
AAG ATG ACG AGG
TAA TTA TCA TGA
TAT TTT TCT TGT
TAC TTC TCC TGC
TAG TTG TCG TGG
CAA CTA CCA CGA
CAT CTT CCT CGT
CAC CTC CCC CGC
CAG CTG CCG CGG
GAA GTA GCA GGA
GAT GTT GCT GGT
GAC GTC GCC GGC
GAG GTG GCG GGG

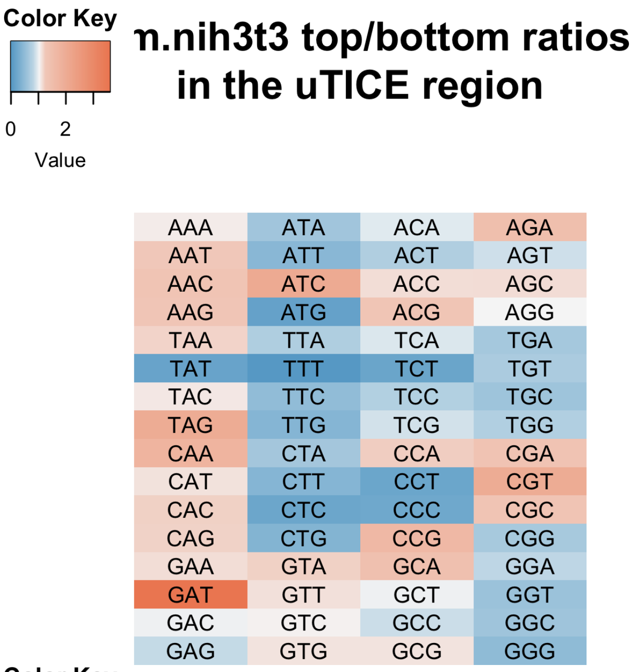
Color Key
0
2
Value
n.nih3t3 top/bottom ratios in the uTICE region
AAA ATA ACA AGA
AAT ATT ACT AGT
AAC ATC ACC AGC
AAG ATG ACG AGG
TAA TTA TCA TGA
TAT TTT TCT TGT
TAC TTC TCC TGC
TAG TTG TCG TGG
CAA CTA CCA CGA
CAT CTT CCT CGT
CAC CTC CCC CGC
CAG CTG CCG CGG
GAA GTA GCA GGA
GAT GTT GCT GGT
GAC GTC GCC GGC
GAG GTG GCG GGG

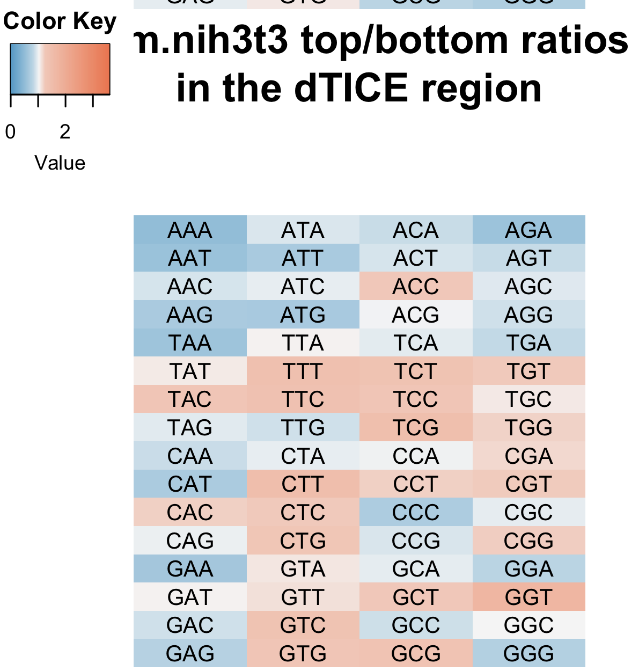
Color Key
0
2
Value
n.nih3t3 top/bottom ratios in the dTICE region
AAA ATA ACA AGA
AAT ATT ACT AGT
AAC ATC ACC AGC
AAG ATG ACG AGG
TAA TTA TCA TGA
TAT TTT TCT TGT
TAC TTC TCC TGC
TAG TTG TCG TGG
CAA CTA CCA CGA
CAT CTT CCT CGT
CAC CTC CCC CGC
CAG CTG CCG CGG
GAA GTA GCA GGA
GAT GTT GCT GGT
GAC GTC GCC GGC
GAG GTG GCG GGG

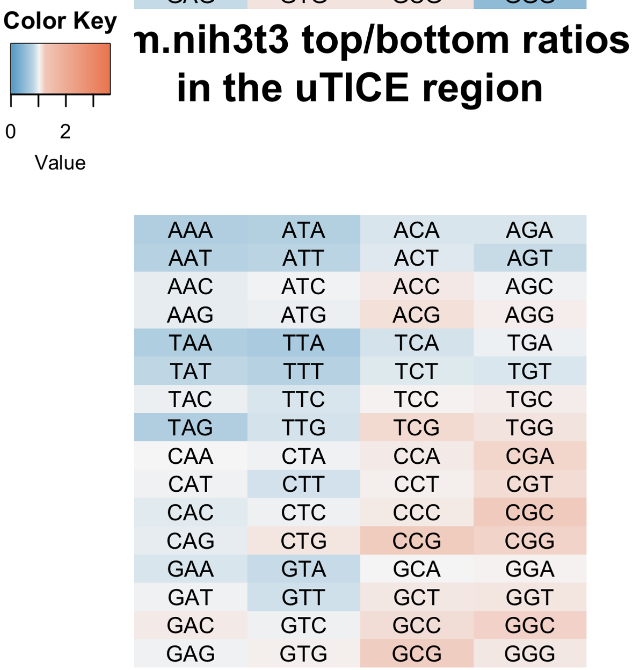
Color Key
0
2
Value
n.nih3t3 top/bottom ratios in the uTICE region
AAA ATA ACA AGA
AAT ATT ACT AGT
AAC ATC ACC AGC
AAG ATG ACG AGG
TAA TTA TCA TGA
TAT TTT TCT TGT
TAC TTC TCC TGC
TAG TTG TCG TGG
CAA CTA CCA CGA
CAT CTT CCT CGT
CAC CTC CCC CGC
CAG CTG CCG CGG
GAA GTA GCA GGA
GAT GTT GCT GGT
GAC GTC GCC GGC
GAG GTG GCG GGG

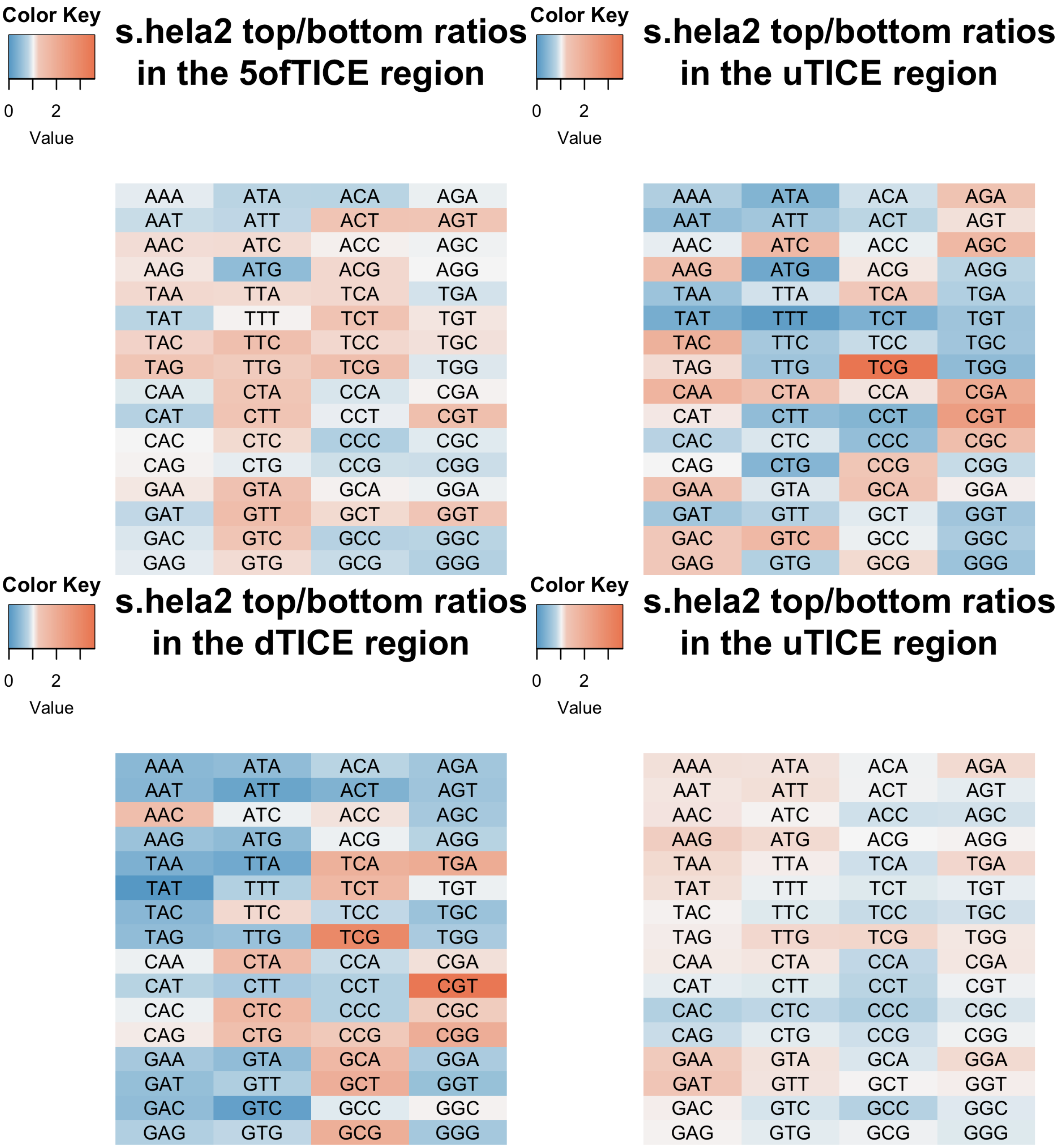


**Figure 10. Pearson correlations between the top/bottom trinucleotide ratios of two regions within a species**

```
for (species in names(species_exemplar_tissue_list)) {
    for (species_tissue in species_exemplar_tissue_list[[species]]) {
        if (species_tissue!="") {
            connector <- paste0(".", species_tissue)
        } else {
```

```
            connector <- ""
        }

      # 5' of TICE
      species.UTR_tri_freq_top_bottom_ratios_5ofTICE_table <- as.numeric(unlist(read.csv(file=paste0("ta

      # uTICE
      species.UTR_tri_freq_top_bottom_ratios_uTICE_table <- as.numeric(unlist(read.csv(file=paste0("tab

      # dTICE
      species.CDS_tri_freq_top_bottom_ratios_dTICE_table <- as.numeric(unlist(read.csv(file=paste0("tab

      # 3' of TICE
      species.CDS_tri_freq_top_bottom_ratios_3ofTICE_table <- as.numeric(unlist(read.csv(file=paste0("ta

      # the correlation table
      species.cors <- matrix(NA, nrow=4, ncol=4)
      rownames(species.cors) <- c("5ofTICE", "uTICE", "dTICE", "3ofTICE")
      colnames(species.cors) <- c("5ofTICE", "uTICE", "dTICE", "3ofTICE")
      species.cors["uTICE", "5ofTICE"] <- cor(species.UTR_tri_freq_top_bottom_ratios_uTICE_table, specie
      species.cors["dTICE", "5ofTICE"] <- cor(species.CDS_tri_freq_top_bottom_ratios_dTICE_table, specie
      species.cors["dTICE", "uTICE"] <- cor(species.CDS_tri_freq_top_bottom_ratios_dTICE_table, species
      species.cors["3ofTICE", "5ofTICE"] <- cor(species.CDS_tri_freq_top_bottom_ratios_3ofTICE_table, sp
      species.cors["3ofTICE", "uTICE"] <- cor(species.CDS_tri_freq_top_bottom_ratios_3ofTICE_table, spec
      species.cors["3ofTICE", "dTICE"] <- cor(species.CDS_tri_freq_top_bottom_ratios_3ofTICE_table, spec

      col_breaks <- c(seq(-1,-0.1,length=100),  # for blue
                      seq(-0.09,0.09,length=100),  # for white
                      seq(0.1,1,length=100)  # for red
                      )

      # make the within-species correlation heatmap
      heatmap.2(species.cors,
                main = paste0(species, connector, " cor of\ntrinucleotide ratios"), # heat map title
                cellnote = round(species.cors, 2),  # same data set for cell labels
                notecol="black",      # change font color of cell labels to black
                density.info="none",  # turns off density plot inside color legend
                trace="none",         # turns off trace lines inside the heat map
                margins =c(15,15),     # widens margins around plot
                col=my_palette,       # use on color palette defined earlier
                breaks=col_breaks,    # enable color transition at specified limits
                dendrogram="none",    # only draw a row dendrogram
                Rowv="NA",            # turn off row clustering
                Colv="NA"            # turn off column clustering
                )
    }
}
```

```
## Warning in image.default(z = matrix(z, ncol = 1), col = col, breaks =
## tmpbreaks, : unsorted 'breaks' will be sorted before use
```

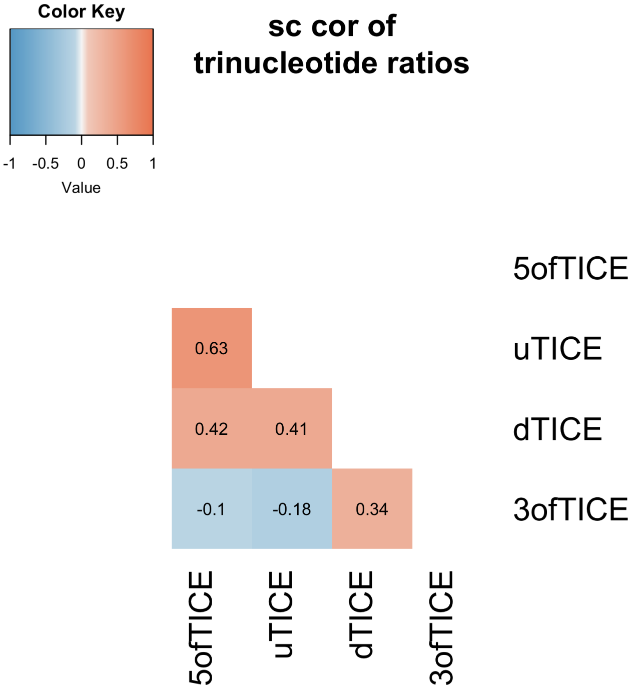


```
## Warning in image.default(z = matrix(z, ncol = 1), col = col, breaks =
## tmpbreaks, : unsorted 'breaks' will be sorted before use
```

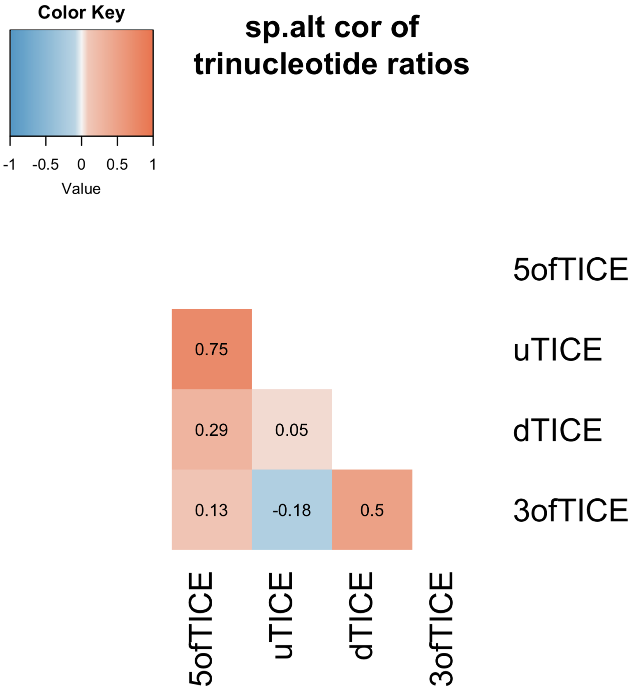


```
## Warning in image.default(z = matrix(z, ncol = 1), col = col, breaks =
## tmpbreaks, : unsorted 'breaks' will be sorted before use
```

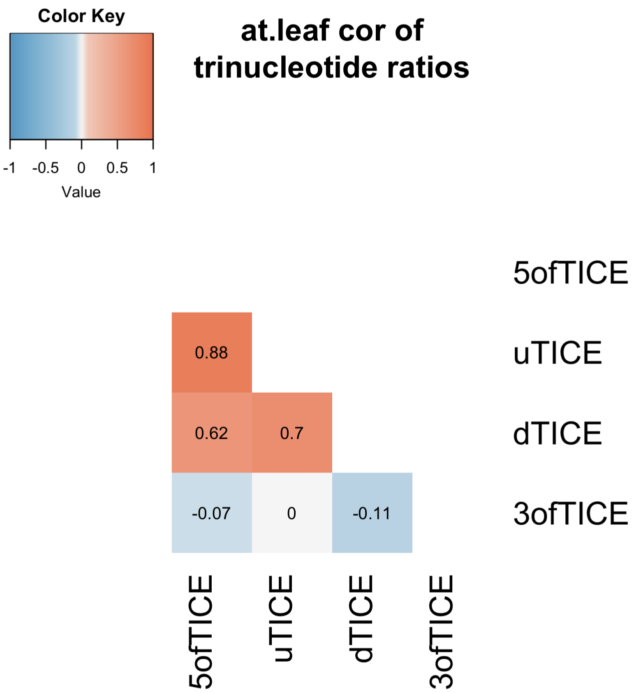


```
## Warning in image.default(z = matrix(z, ncol = 1), col = col, breaks =
## tmpbreaks, : unsorted 'breaks' will be sorted before use
```

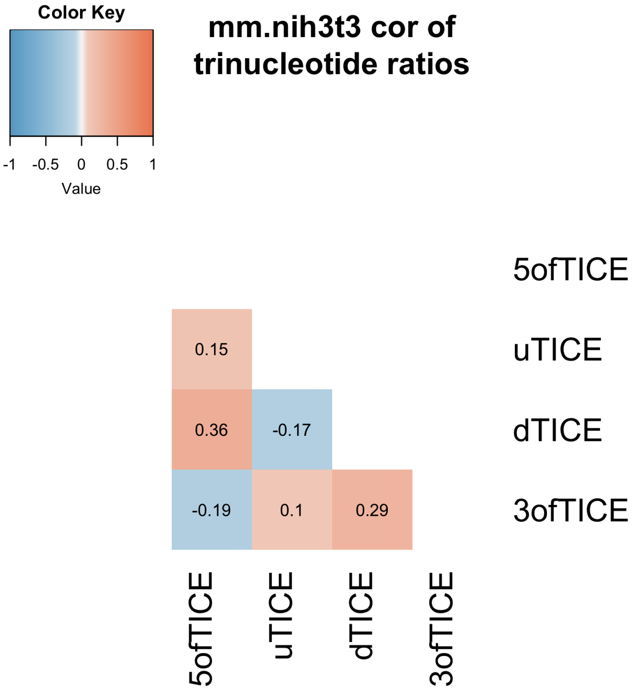


```
## Warning in image.default(z = matrix(z, ncol = 1), col = col, breaks =
## tmpbreaks, : unsorted 'breaks' will be sorted before use
```

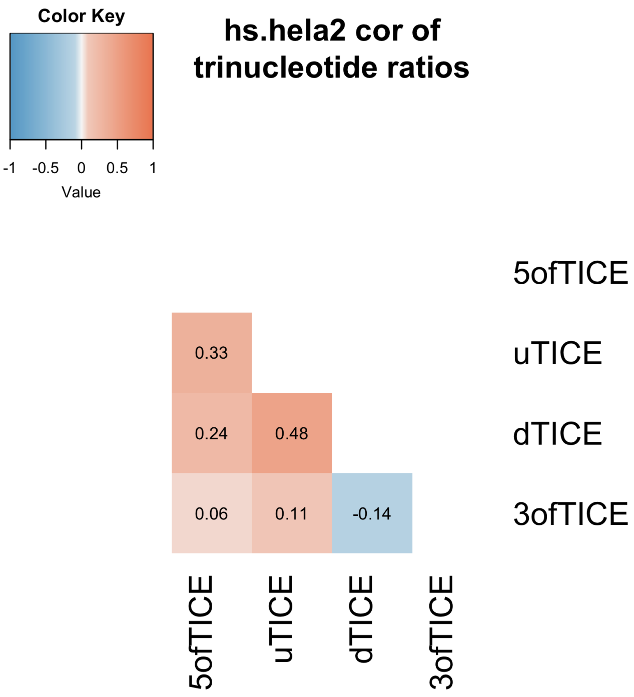


**Figure 11. Pearson correlations between the top/bottom trinucleotide ratios of the region of two species**

```
btw_species_cors <- NULL
rownames_btw_species_cors <- NULL
colnames_btw_species_cors <- c("5ofTICE vs 5ofTICE", "uTICE vs uTICE", "dTICE vs dTICE", "3ofTICE vs 3o
for (i in 1:(length(species_exemplar_tissue_list)-1)) {
  for (j in (i+1):length(species_exemplar_tissue_list)) {
    species1 <- names(species_exemplar_tissue_list)[i]
    species2 <- names(species_exemplar_tissue_list)[j]
    species1_tissue <- species_exemplar_tissue_list[[species1]]
    species2_tissue <- species_exemplar_tissue_list[[species2]]
    if (species1_tissue!="") {
            connector1 <- paste0(".", species1_tissue)
        } else {
            connector1 <- ""
        }
    if (species2_tissue!="") {
            connector2 <- paste0(".", species2_tissue)
        } else {
            connector2 <- ""
        }

    # 5' of TICE
      species1.UTR_tri_freq_top_bottom_ratios_5ofTICE_table <- as.numeric(unlist(read.csv(file=paste0("
      species2.UTR_tri_freq_top_bottom_ratios_5ofTICE_table <- as.numeric(unlist(read.csv(file=paste0("

      # uTICE
      species1.UTR_tri_freq_top_bottom_ratios_uTICE_table <- as.numeric(unlist(read.csv(file=paste0("ta
```

```
      species2.UTR_tri_freq_top_bottom_ratios_uTICE_table <- as.numeric(unlist(read.csv(file=paste0("ta

      # dTICE
      species1.CDS_tri_freq_top_bottom_ratios_dTICE_table <- as.numeric(unlist(read.csv(file=paste0("ta
      species2.CDS_tri_freq_top_bottom_ratios_dTICE_table <- as.numeric(unlist(read.csv(file=paste0("ta

      # 3' of TICE
      species1.CDS_tri_freq_top_bottom_ratios_3ofTICE_table <- as.numeric(unlist(read.csv(file=paste0("
      species2.CDS_tri_freq_top_bottom_ratios_3ofTICE_table <- as.numeric(unlist(read.csv(file=paste0("

      rownames_btw_species_cors <- c(
        rownames_btw_species_cors,
        paste0(species1, connector1, " vs. ", species2, connector2)
      )

      btw_species_cors <- rbind(
        btw_species_cors,
        c(
          cor(species1.UTR_tri_freq_top_bottom_ratios_5ofTICE_table, species2.UTR_tri_freq_top_bottom_r
          cor(species1.UTR_tri_freq_top_bottom_ratios_uTICE_table, species2.UTR_tri_freq_top_bottom_rat
          cor(species1.CDS_tri_freq_top_bottom_ratios_dTICE_table, species2.CDS_tri_freq_top_bottom_rat
          cor(species1.CDS_tri_freq_top_bottom_ratios_3ofTICE_table, species2.CDS_tri_freq_top_bottom_r
        )
      )
  }
}
rownames(btw_species_cors) <- rownames_btw_species_cors
colnames(btw_species_cors) <- colnames_btw_species_cors

# make the between-species correlation heatmap
heatmap.2(btw_species_cors,
  main = "Cor of trinucleotide\nratios between species", # heat map title
  cellnote = round(btw_species_cors, 2),  # same data set for cell labels
  notecol="black",      # change font color of cell labels to black
  density.info="none",  # turns off density plot inside color legend
  trace="none",         # turns off trace lines inside the heat map
  margins =c(15,15),     # widens margins around plot
  col=my_palette,       # use on color palette defined earlier
  breaks=col_breaks,    # enable color transition at specified limits
  dendrogram="none",    # only draw a row dendrogram
  Rowv="NA",            # turn off row clustering
  Colv="NA"            # turn off column clustering
)
```

```
## Warning in image.default(z = matrix(z, ncol = 1), col = col, breaks =
## tmpbreaks, : unsorted 'breaks' will be sorted before use
```

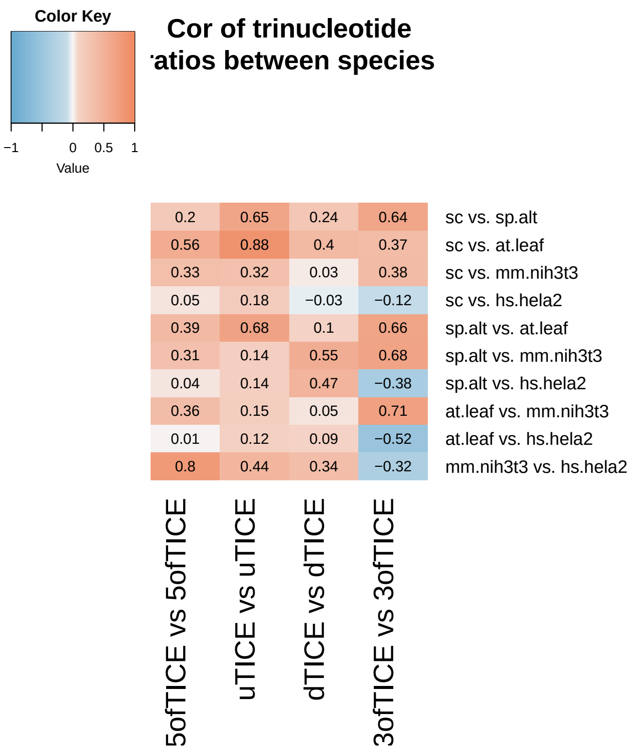


### Figure 12. Collinearity analysis: 5’biochem (i.e. uAUG and RNAfold) and 5’motif (i.e. 5’ofTICE+uTICE+dTICE)

```
for (species in names(species_tissue_list)) {
    for (species_tissue in species_tissue_list[[species]]) {
        if (species_tissue != "") {
            tissue_dot <- paste0(".", species_tissue)
            tissue_us <- paste0("_", species_tissue)
        } else {
            tissue_dot <- ""
            tissue_us <- ""
        }

        features_uAUG <- get(paste0(species, tissue_dot, ".uAUG_counts.div"))

        features_RNAfold <- get(paste0(species, tissue_dot, ".fold_energy_features_selected"))

        features_5biochem <- cbind(features_uAUG, features_RNAfold)

        features_5motif <- cbind(get(paste0(species, tissue_dot,
            ".5ofTICE_features"))[[1]], get(paste0(species, tissue_dot,
            ".uTICE_features"))[[1]], get(paste0(species, tissue_dot,
            ".dTICE_features"))[[1]])

        # uAUG vs. 5motif
        R2_uAUG <- summary(lm(get(paste0(species, tissue_dot,
            ".log10.TR")) ~ as.matrix(features_uAUG)))$r.squared
```

```r
R2_5motif <- summary(lm(get(paste0(species, tissue_dot,
    ".log10.TR")) ~ as.matrix(features_5motif)))$r.squared
R2_uAUG_5motif <- summary(lm(get(paste0(species, tissue_dot,
    ".log10.TR")) ~ as.matrix(cbind(features_uAUG, features_5motif))))$r.squared

cat(paste0(species, tissue_dot, "\nR2 of uAUG+5motif: "))
cat(R2_uAUG_5motif)
cat("\n")
cat("% of uAUG: ")
cat(round(R2_uAUG/R2_uAUG_5motif * 100, 3))
cat("%\n")
cat("% of 5motif: ")
cat(round(R2_5motif/R2_uAUG_5motif * 100, 3))
cat("%\n")
cat("% of collinearity: ")
cat(round((R2_uAUG + R2_5motif - R2_uAUG_5motif)/R2_uAUG_5motif *
    100, 3))
cat("%\n\n")

# RNAfold vs. 5motif
R2_RNAfold <- summary(lm(get(paste0(species, tissue_dot,
    ".log10.TR")) ~ as.matrix(features_RNAfold)))$r.squared
R2_RNAfold_5motif <- summary(lm(get(paste0(species, tissue_dot,
    ".log10.TR")) ~ as.matrix(cbind(features_RNAfold,
    features_5motif))))$r.squared

cat("R2 of RNAfold+5motif: ")
cat(R2_RNAfold_5motif)
cat("\n")
cat("% of RNAfold: ")
cat(round(R2_RNAfold/R2_RNAfold_5motif * 100, 3))
cat("%\n")
cat("% of 5motif: ")
cat(round(R2_5motif/R2_RNAfold_5motif * 100, 3))
cat("%\n")
cat("% of collinearity: ")
cat(round((R2_RNAfold + R2_5motif - R2_RNAfold_5motif)/R2_RNAfold_5motif *
    100, 3))
cat("%\n\n")

# 5biochem vs. 5motif
R2_5biochem <- summary(lm(get(paste0(species, tissue_dot,
    ".log10.TR")) ~ as.matrix(features_5biochem)))$r.squared
R2_5biochem_5motif <- summary(lm(get(paste0(species,
    tissue_dot, ".log10.TR")) ~ as.matrix(cbind(features_5biochem,
    features_5motif))))$r.squared

cat("R2 of 5biochem+5motif: ")
cat(R2_5biochem_5motif)
cat("\n")
cat("% of 5biochem: ")
cat(round(R2_5biochem/R2_5biochem_5motif * 100, 3))
cat("%\n")
```

```r
        cat("% of 5motif: ")
        cat(round(R2_5motif/R2_5biochem_5motif * 100, 3))
        cat("%\n")
        cat("% of collinearity: ")
        cat(round((R2_5biochem + R2_5motif - R2_5biochem_5motif)/R2_5biochem_5motif *
            100, 3))
        cat("%\n\n")
    }
}
```

```
## sc
## R2 of uAUG+5motif: 0.3455477
## % of uAUG: 13.448%
## % of 5motif: 98.506%
## % of collinearity: 11.955%
##
## R2 of RNAfold+5motif: 0.4475022
## % of RNAfold: 74.444%
## % of 5motif: 76.064%
## % of collinearity: 50.508%
##
## R2 of 5biochem+5motif: 0.4494612
## % of 5biochem: 76.194%
## % of 5motif: 75.732%
## % of collinearity: 51.926%
##
## sp
## R2 of uAUG+5motif: 0.2227838
## % of uAUG: 22.207%
## % of 5motif: 93.95%
## % of collinearity: 16.157%
##
## R2 of RNAfold+5motif: 0.2591796
## % of RNAfold: 42.28%
## % of 5motif: 80.757%
## % of collinearity: 23.037%
##
## R2 of 5biochem+5motif: 0.2594661
## % of 5biochem: 47.779%
## % of 5motif: 80.668%
## % of collinearity: 28.447%
##
## sp.alt
## R2 of uAUG+5motif: 0.236603
## % of uAUG: 14.748%
## % of 5motif: 96.332%
## % of collinearity: 11.08%
##
## R2 of RNAfold+5motif: 0.2520633
## % of RNAfold: 43.829%
## % of 5motif: 90.424%
## % of collinearity: 34.253%
##
## R2 of 5biochem+5motif: 0.25252
```

```
## % of 5biochem: 44.851%
## % of 5motif: 90.26%
## % of collinearity: 35.111%
##
## at.leaf
## R2 of uAUG+5motif: 0.2321785
## % of uAUG: 32.411%
## % of 5motif: 91.51%
## % of collinearity: 23.921%
##
## R2 of RNAfold+5motif: 0.3149283
## % of RNAfold: 75.918%
## % of 5motif: 67.465%
## % of collinearity: 43.383%
##
## R2 of 5biochem+5motif: 0.3179388
## % of 5biochem: 80.865%
## % of 5motif: 66.826%
## % of collinearity: 47.691%
##
## at.root
## R2 of uAUG+5motif: 0.2106883
## % of uAUG: 44.04%
## % of 5motif: 90.034%
## % of collinearity: 34.074%
##
## R2 of RNAfold+5motif: 0.2576114
## % of RNAfold: 61.739%
## % of 5motif: 73.634%
## % of collinearity: 35.374%
##
## R2 of 5biochem+5motif: 0.2616652
## % of 5biochem: 70.499%
## % of 5motif: 72.494%
## % of collinearity: 42.992%
##
## at.shoot
## R2 of uAUG+5motif: 0.2008575
## % of uAUG: 46.782%
## % of 5motif: 89.821%
## % of collinearity: 36.603%
##
## R2 of RNAfold+5motif: 0.2255777
## % of RNAfold: 58.13%
## % of 5motif: 79.978%
## % of collinearity: 38.108%
##
## R2 of 5biochem+5motif: 0.2303972
## % of 5biochem: 68.267%
## % of 5motif: 78.305%
## % of collinearity: 46.572%
##
## mm.nih3t3
## R2 of uAUG+5motif: 0.1756989
```

```
## % of uAUG: 43.86%
## % of 5motif: 79.207%
## % of collinearity: 23.068%
##
## R2 of RNAfold+5motif: 0.2501444
## % of RNAfold: 71.265%
## % of 5motif: 55.635%
## % of collinearity: 26.9%
##
## R2 of 5biochem+5motif: 0.2786699
## % of 5biochem: 89.552%
## % of 5motif: 49.94%
## % of collinearity: 39.491%
##
## mm.liver
## R2 of uAUG+5motif: 0.1595938
## % of uAUG: 40.033%
## % of 5motif: 81.79%
## % of collinearity: 21.823%
##
## R2 of RNAfold+5motif: 0.2490616
## % of RNAfold: 72.354%
## % of 5motif: 52.41%
## % of collinearity: 24.764%
##
## R2 of 5biochem+5motif: 0.274304
## % of 5biochem: 89.533%
## % of 5motif: 47.587%
## % of collinearity: 37.12%
##
## mm.kidney
## R2 of uAUG+5motif: 0.2120517
## % of uAUG: 36.346%
## % of 5motif: 83.312%
## % of collinearity: 19.658%
##
## R2 of RNAfold+5motif: 0.3036289
## % of RNAfold: 70.684%
## % of 5motif: 58.184%
## % of collinearity: 28.868%
##
## R2 of 5biochem+5motif: 0.3346106
## % of 5biochem: 88.964%
## % of 5motif: 52.797%
## % of collinearity: 41.761%
##
## hs.hela2
## R2 of uAUG+5motif: 0.1020453
## % of uAUG: 15.726%
## % of 5motif: 92.977%
## % of collinearity: 8.703%
##
## R2 of RNAfold+5motif: 0.1458517
## % of RNAfold: 60.212%
```

```
## % of 5motif: 65.052%
## % of collinearity: 25.263%
##
## R2 of 5biochem+5motif: 0.1474626
## % of 5biochem: 64.002%
## % of 5motif: 64.341%
## % of collinearity: 28.342%
```

#### A comment in the text - Collinearity analysis: 5'UTR length and 5'biochem+5'motif

```
for (species in names(species_tissue_list)) {
    for (species_tissue in species_tissue_list[[species]]) {
        if (species_tissue != "") {
            tissue_dot <- paste0(".", species_tissue)
            tissue_us <- paste0("_", species_tissue)
        } else {
            tissue_dot <- ""
            tissue_us <- ""
        }

        features_UTRlens <- get(paste0(species, tissue_dot, ".log10.UTR_lens.div"))

        features_uAUG <- get(paste0(species, tissue_dot, ".uAUG_counts.div"))

        features_RNAfold <- get(paste0(species, tissue_dot, ".fold_energy_features_selected"))

        features_5biochem <- cbind(features_uAUG, features_RNAfold)

        features_5motif <- cbind(get(paste0(species, tissue_dot,
            ".5ofTICE_features"))[[1]], get(paste0(species, tissue_dot,
            ".uTICE_features"))[[1]], get(paste0(species, tissue_dot,
            ".dTICE_features"))[[1]])

        features_5biochem_5motif <- cbind(features_5biochem,
            features_5motif)

        # UTR_lens vs. 5biochem+5motif
        R2_UTRlens <- summary(lm(get(paste0(species, tissue_dot,
            ".log10.TR")) ~ as.matrix(features_UTRlens)))$r.squared
        R2_5biochem_5motif <- summary(lm(get(paste0(species,
            tissue_dot, ".log10.TR")) ~ as.matrix(features_5biochem_5motif)))$r.squared
        R2_UTRlens_5biochem_5motif <- summary(lm(get(paste0(species,
            tissue_dot, ".log10.TR")) ~ as.matrix(cbind(features_UTRlens,
            features_5biochem_5motif))))$r.squared

        cat(paste0(species, tissue_dot, "\nR2 of UTRlens+5biochem+5motif: "))
        cat(R2_UTRlens_5biochem_5motif)
        cat("\n")
        cat("% of UTRlens: ")
        cat(round(R2_UTRlens/R2_UTRlens_5biochem_5motif * 100,
```

```
            3))
        cat("%\n")
        cat("% of 5motif: ")
        cat(round(R2_5biochem_5motif/R2_UTRlens_5biochem_5motif *
            100, 3))
        cat("%\n")
        cat("% of collinearity: ")
        cat(round((R2_UTRlens + R2_5biochem_5motif - R2_UTRlens_5biochem_5motif)/R2_UTRlens_5biochem_5mo
            100, 3))
        cat("%\n\n")
    }
}
```

```
## sc
## R2 of UTRlens+5biochem+5motif: 0.4556677
## % of UTRlens: 11.034%
## % of 5motif: 98.638%
## % of collinearity: 9.672%
##
## sp
## R2 of UTRlens+5biochem+5motif: 0.2598144
## % of UTRlens: 14.677%
## % of 5motif: 99.866%
## % of collinearity: 14.543%
##
## sp.alt
## R2 of UTRlens+5biochem+5motif: 0.2533873
## % of UTRlens: 24.842%
## % of 5motif: 99.658%
## % of collinearity: 24.5%
##
## at.leaf
## R2 of UTRlens+5biochem+5motif: 0.320601
## % of UTRlens: 22.697%
## % of 5motif: 99.17%
## % of collinearity: 21.866%
##
## at.root
## R2 of UTRlens+5biochem+5motif: 0.2641215
## % of UTRlens: 34.597%
## % of 5motif: 99.07%
## % of collinearity: 33.667%
##
## at.shoot
## R2 of UTRlens+5biochem+5motif: 0.233814
## % of UTRlens: 36.533%
## % of 5motif: 98.539%
## % of collinearity: 35.072%
##
## mm.nih3t3
## R2 of UTRlens+5biochem+5motif: 0.2789754
## % of UTRlens: 17.99%
## % of 5motif: 99.89%
## % of collinearity: 17.881%
```

```
##
## mm.liver
## R2 of UTRlens+5biochem+5motif: 0.2755491
## % of UTRlens: 12.507%
## % of 5motif: 99.548%
## % of collinearity: 12.056%
##
## mm.kidney
## R2 of UTRlens+5biochem+5motif: 0.3347473
## % of UTRlens: 15.8%
## % of 5motif: 99.959%
## % of collinearity: 15.759%
##
## hs.hela2
## R2 of UTRlens+5biochem+5motif: 0.1478139
## % of UTRlens: 10.932%
## % of 5motif: 99.762%
## % of collinearity: 10.694%
```

### Figure 13. R2s of six cis-control elements (an excel file to store the R2 values first)

#### functions

```
# a function to calculate the R2 for a full model with 8
# features (five mega features + three single features)
calc_8feature_full_model_R2 <- function(species.log10.TR, species.uTICE_features,
    species.dTICE_features, species.5ofTICE_features, species.61codon_features,
    species.log10.UTR_lens, species.log10.CDS_lens, species.uAUG_counts,
    species.fold_energy_features_selected, species.polyA_lens,
    len_breaks, ifplot = F, title = "") {
    n <- length(species.log10.UTR_lens)
    UTR_lens <- 10^species.log10.UTR_lens

    features <- cbind(species.uTICE_features, species.dTICE_features,
        species.5ofTICE_features, species.61codon_features, species.log10.CDS_lens,
        species.uAUG_counts, species.fold_energy_features_selected,
        species.polyA_lens)

    lm_selected <- lm(species.log10.TR ~ as.matrix(features),
        y = TRUE)

    if (ifplot) {
        plot(x = lm_selected$fitted, y = lm_selected$y, xlab = "log10 TR (predicted)",
            ylab = "log10 TR", main = paste(title, "R2 =", round(summary(lm_selected)$r.squared,
                2)))
    }
    return(summary(lm_selected)$r.squared)
}

# a function to calculate the R2 for a full model with five
```

```
# mega features + two single features (uORFs, CDS length)
calc_5_2_feature_full_model_R2 <- function(species.log10.TR,
    species.uTICE_features, species.dTICE_features, species.5ofTICE_features,
    species.61codon_features, species.log10.UTR_lens, species.log10.CDS_lens,
    species.uAUG_counts, species.fold_energy_features_selected,
    len_breaks, ifplot = F, title = "") {
    n <- length(species.log10.UTR_lens)
    UTR_lens <- 10^species.log10.UTR_lens

    features <- cbind(species.uTICE_features, species.dTICE_features,
        species.5ofTICE_features, species.61codon_features, species.log10.CDS_lens,
        species.uAUG_counts, species.fold_energy_features_selected)

    lm_selected <- lm(species.log10.TR ~ as.matrix(features),
        y = TRUE)

    if (ifplot) {
        plot(x = lm_selected$fitted, y = lm_selected$y, xlab = "log10 TR (predicted)",
            ylab = "log10 TR", main = paste(title, "R2 =", round(summary(lm_selected)$r.squared,
                2)))
    }
    return(summary(lm_selected)$r.squared)
}
```

##### the following code only needs to be run once

```
six_element_R2s <- NULL
rownames_six_element_R2s <- NULL
colnames_six_element_R2s <- c("uAUGs", "RNA fold", "5' motifs",
    "CDS length", "codon usage", "CDS RNA fold", "poly A", "full model 8 features",
    "full model 5 features")
for (species in names(species_tissue_list)) {
    for (species_tissue in species_tissue_list[[species]]) {
        if (species_tissue != "") {
            tissue_dot <- paste0(".", species_tissue)
            tissue_us <- paste0("_", species_tissue)
        } else {
            tissue_dot <- ""
            tissue_us <- ""
        }
        rownames_six_element_R2s <- c(rownames_six_element_R2s,
            paste0(species, tissue_dot))
        tmp_R2s <- NULL
        ## uAUG_counts
        tmp_R2s <- c(tmp_R2s, summary(lm(get(paste0(species,
            tissue_dot, ".log10.TR")) ~ get(paste0(species, tissue_dot,
            ".uAUG_counts.div"))))$r.squared)
        ## selected UTR folding energy features
        tmp_R2s <- c(tmp_R2s, summary(lm(get(paste0(species,
            tissue_dot, ".log10.TR")) ~ as.matrix(get(paste0(species,
            tissue_dot, ".fold_energy_features_selected")))))$r.squared)
        ## 5' motifs (uTICE + dTICE + 5' of TICE features)
```

```
TICE_features <- cbind(get(paste0(species, tissue_dot,
    ".uTICE_features"))[[1]], get(paste0(species, tissue_dot,
    ".dTICE_features"))[[1]], get(paste0(species, tissue_dot,
    ".5ofTICE_features"))[[1]])
tmp_R2s <- c(tmp_R2s, summary(lm(get(paste0(species,
    tissue_dot, ".log10.TR")) ~ as.matrix(TICE_features)))$r.squared)
## log10.CDS_lens
tmp_R2s <- c(tmp_R2s, summary(lm(get(paste0(species,
    tissue_dot, ".log10.TR")) ~ get(paste0(species, tissue_dot,
    ".log10.CDS_lens.div"))))$r.squared)
## codon features
tmp_R2s <- c(tmp_R2s, summary(lm(get(paste0(species,
    tissue_dot, ".log10.TR")) ~ as.matrix(get(paste0(species,
    tissue_dot, ".codon_features"))[[1]])))$r.squared)
## CDS fold energy
tmp_R2s <- c(tmp_R2s, summary(lm(get(paste0(species,
    tissue_dot, ".log10.TR")) ~ get(paste0(species, tissue_dot,
    ".CDS_fold_energy.div"))))$r.squared)
## polyA_lens
tmp_R2s <- c(tmp_R2s, summary(lm(get(paste0(species,
    tissue_dot, ".log10.TR")) ~ get(paste0(species, tissue_dot,
    ".polyA_lens.div"))))$r.squared)

## 8-feature model
tmp_R2s <- c(tmp_R2s, calc_8feature_full_model_R2(get(paste0(species,
    tissue_dot, ".log10.TR")), get(paste0(species, tissue_dot,
    ".uTICE_features"))[[1]], get(paste0(species, tissue_dot,
    ".dTICE_features"))[[1]], get(paste0(species, tissue_dot,
    ".5ofTICE_features"))[[1]], get(paste0(species, tissue_dot,
    ".codon_features"))[[1]], get(paste0(species, tissue_dot,
    ".log10.UTR_lens.div")), get(paste0(species, tissue_dot,
    ".log10.CDS_lens.div")), get(paste0(species, tissue_dot,
    ".uAUG_counts.div")), get(paste0(species, tissue_dot,
    ".fold_energy_features_selected")), get(paste0(species,
    tissue_dot, ".polyA_lens.div")), get(paste0(species,
    tissue_dot, ".len_breaks")), ifplot = F, title = paste(species,
    species_tissue)))

## 5-feature model
tmp_R2s <- c(tmp_R2s, calc_5_2_feature_full_model_R2(get(paste0(species,
    tissue_dot, ".log10.TR")), get(paste0(species, tissue_dot,
    ".uTICE_features"))[[1]], get(paste0(species, tissue_dot,
    ".dTICE_features"))[[1]], get(paste0(species, tissue_dot,
    ".5ofTICE_features"))[[1]], get(paste0(species, tissue_dot,
    ".codon_features"))[[1]], get(paste0(species, tissue_dot,
    ".log10.UTR_lens.div")), get(paste0(species, tissue_dot,
    ".log10.CDS_lens.div")), get(paste0(species, tissue_dot,
    ".uAUG_counts.div")), get(paste0(species, tissue_dot,
    ".fold_energy_features_selected")), get(paste0(species,
    tissue_dot, ".len_breaks")), ifplot = F, title = paste(species,
    species_tissue)))

six_element_R2s <- rbind(six_element_R2s, tmp_R2s)
```

```
    }
}
rownames(six_element_R2s) <- rownames_six_element_R2s
colnames(six_element_R2s) <- colnames_six_element_R2s

write.csv(six_element_R2s, file = "tables/six_element_R2s.csv",
    quote = F)
```

##### plot

```
for (species in names(species_tissue_list)) {
    for (i in 1:length(species_tissue_list[[species]])) {
        species_tissue <- species_tissue_list[[species]][i]
        if (species_tissue != "") {
            tissue_dot <- paste0(".", species_tissue)
            tissue_us <- paste0("_", species_tissue)
        } else {
            tissue_dot <- ""
            tissue_us <- ""
        }
        calc_5_2_feature_full_model_R2(get(paste0(species, tissue_dot,
            ".log10.TR")), get(paste0(species, tissue_dot, ".uTICE_features"))[[1]],
            get(paste0(species, tissue_dot, ".dTICE_features"))[[1]],
            get(paste0(species, tissue_dot, ".5ofTICE_features"))[[1]],
            get(paste0(species, tissue_dot, ".codon_features"))[[1]],
            get(paste0(species, tissue_dot, ".log10.UTR_lens.div")),
            get(paste0(species, tissue_dot, ".log10.CDS_lens.div")),
            get(paste0(species, tissue_dot, ".uAUG_counts.div")),
            get(paste0(species, tissue_dot, ".fold_energy_features_selected")),
            get(paste0(species, tissue_dot, ".len_breaks")),
            ifplot = T, title = paste(species, species_tissue))
    }
}
```

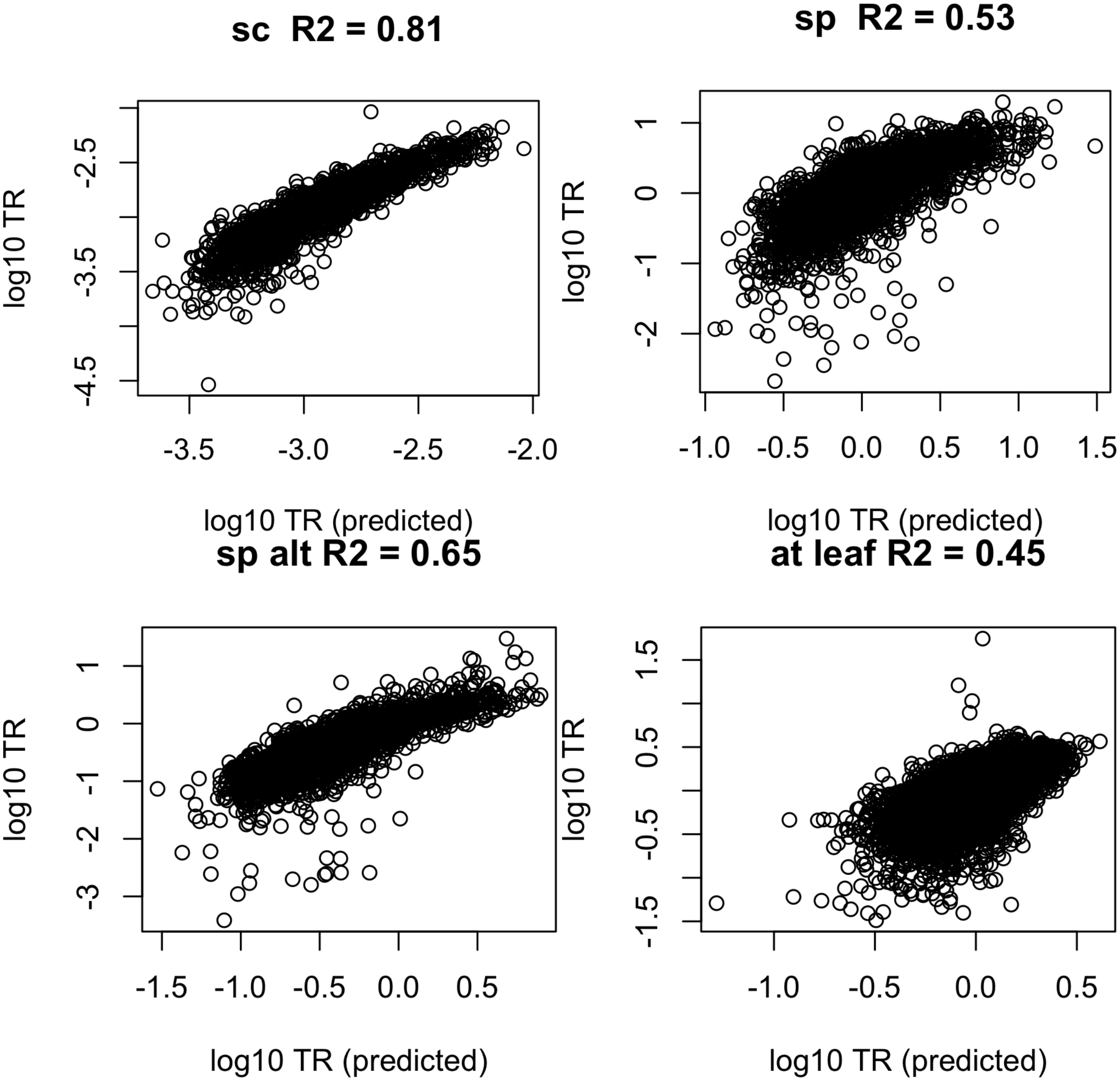
sc R2 = 0.81
sp R2 = 0.53
sp alt R2 = 0.65
at leaf R2 = 0.45
log10 TR
log10 TR (predicted)
-4.5
-3.5
-2.5
-3.5
-3.0
-2.5
-2.0
-2
-1
0
1
-1.0
-0.5
0.0
0.5
1.0
1.5
-3
-2
-1
0
1
-1.5
-1.0
-0.5
0.0
0.5
-1.5
-0.5
0.5
1.5
-1.0
-0.5
0.0
0.5

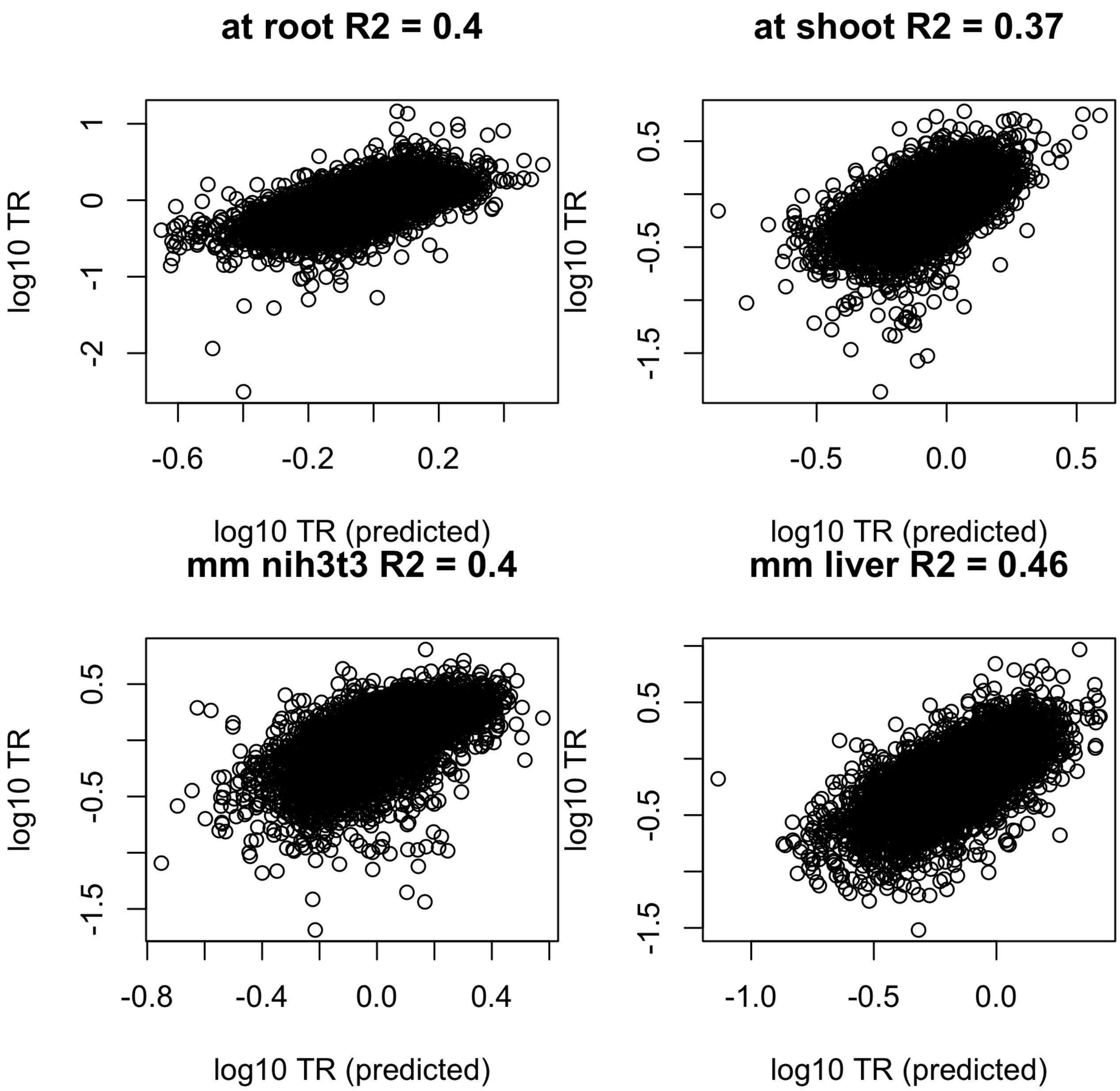
at root R2 = 0.4
at shoot R2 = 0.37
mm nih3t3 R2 = 0.4
mm liver R2 = 0.46
log10 TR
log10 TR (predicted)
1
0
-1
-2
-0.6
-0.2
0.2
0.5
-0.5
-1.5
0.0
-0.8
-0.4
0.4
-1.0

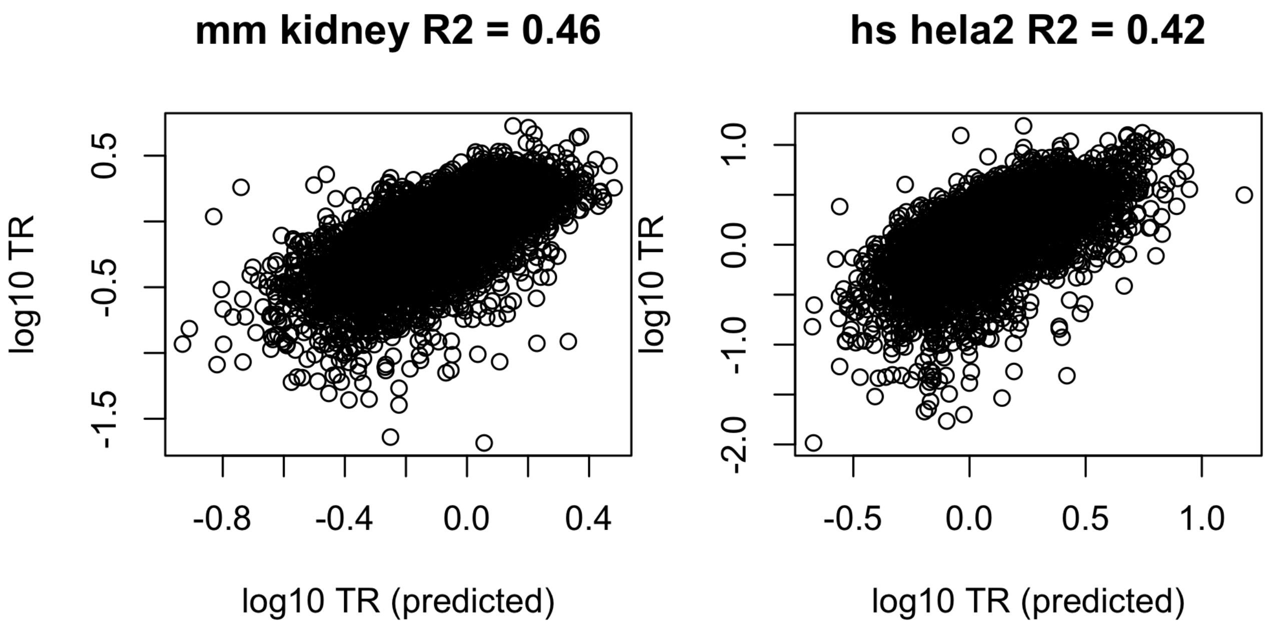


**Figure 14. Collinearity analysis of four features: uAUGs, 5'RNA fold, CDS length, and codon usage**

```
## A function to calculate pairwise collinearity of every two
## features
calc_pariwise_collinearity <- function(features_list, y) {
    p <- length(features_list)
    feature_names <- names(features_list)

    uni_feature_R2s <- sapply(1:p, FUN = function(i) {
        tmp_features_all <- cbind(y, features_list[[i]])
        colnames(tmp_features_all)[1] <- "y"
        ## remove duplicated column names
        tmp_features_all <- tmp_features_all[, !duplicated(colnames(tmp_features_all))]
        tmp_features_all <- as.data.frame(tmp_features_all)
        ## model
        tmp_features_model <- lm(y ~ ., data = tmp_features_all)
        summary(tmp_features_model)$r.squared
    })

    feature_idx_combn <- combn(p, 2)
    bi_feature_R2s <- apply(feature_idx_combn, 2, FUN = function(feature_idx) {
        tmp_features_all <- y
        for (i in feature_idx) {
            tmp_features_all <- cbind(tmp_features_all, features_list[[i]])
        }
        colnames(tmp_features_all)[1] <- "y"
        ## remove duplicated column names
        tmp_features_all <- tmp_features_all[, !duplicated(colnames(tmp_features_all))]
        tmp_features_all <- as.data.frame(tmp_features_all)
        ## model
```

```r
        tmp_features_model <- lm(y ~ ., data = tmp_features_all)
        summary(tmp_features_model)$r.squared
    })

    result <- matrix(NA, p, p)
    colnames(result) <- feature_names
    rownames(result) <- feature_names
    # the order of bi_feature_R2s should correspond to the
    # columns of feature_idx_combn
    for (i in 1:length(bi_feature_R2s)) {
        idx <- feature_idx_combn[, i]
        redundancy <- sum(uni_feature_R2s[idx]) - bi_feature_R2s[i]
        redundancy <- max(0, redundancy)
        result[idx[2], idx[1]] <- max(round(redundancy/uni_feature_R2s[idx[1]] *
            100, 3), round(redundancy/uni_feature_R2s[idx[2]] *
            100, 3))
    }
    result
}

## A function to make levelplot showing values
myPanel <- function(x, y, z, ...) {
    panel.levelplot(x, y, z, ...)
    panel.text(x, y, round(z, 2))
}

for (species in names(species_tissue_list)) {
    for (species_tissue in species_tissue_list[[species]]) {
        if (species_tissue != "") {
            tissue_dot <- paste0(".", species_tissue)
            tissue_us <- paste0("_", species_tissue)
        } else {
            tissue_dot <- ""
            tissue_us <- ""
        }

        features_list <- list(uAUG = get(paste0(species, tissue_dot,
            ".uAUG_counts.div")), RNAfold = get(paste0(species,
            tissue_dot, ".fold_energy_features_selected")), CDSlen = get(paste0(species,
            tissue_dot, ".log10.CDS_lens.div")), CodonUsage = get(paste0(species,
            tissue_dot, ".codon_features"))[[1]])
        tmp <- calc_pariwise_collinearity(features_list, get(paste0(species,
            tissue_dot, ".log10.TR")))
        tmp[tmp > 100] <- 100  # truncate the value at 100
        assign(x = paste0(species, tissue_dot, ".pariwise_collinearity"),
            value = tmp[-1, -4]/100)
        grid <- setNames(melt(get(paste0(species, tissue_dot,
            ".pariwise_collinearity"))), c("Feature_1", "Feature_2",
            "Collinearity"))
        print(levelplot(Collinearity ~ Feature_1 * Feature_2,
            grid, panel = myPanel, scales = list(x = list(rot = 90)),
            main = paste(species, species_tissue), col.regions = rev(heat.colors(120)),
            cuts = 100, at = seq(0, 1, 0.01)))
```

```
        filename <- paste0("tables/pairwise_collinearity/", species,
            tissue_dot, ".pariwise_collinearity.csv")
        if (!file.exists(filename)) {
            write.csv(get(paste0(species, tissue_dot, ".pariwise_collinearity"))/100,
                file = filename, quote = F)
        }
    }
}
```

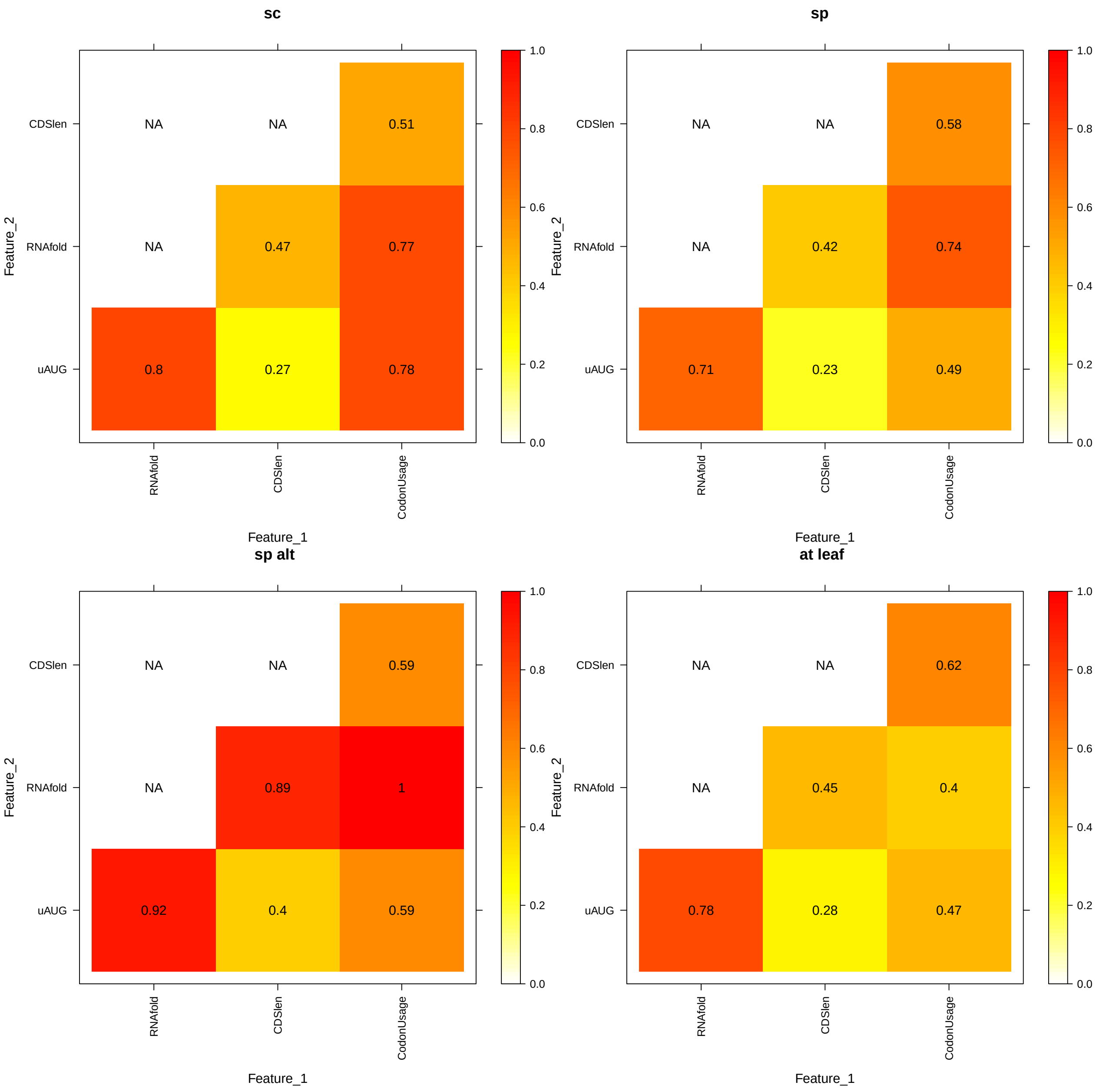

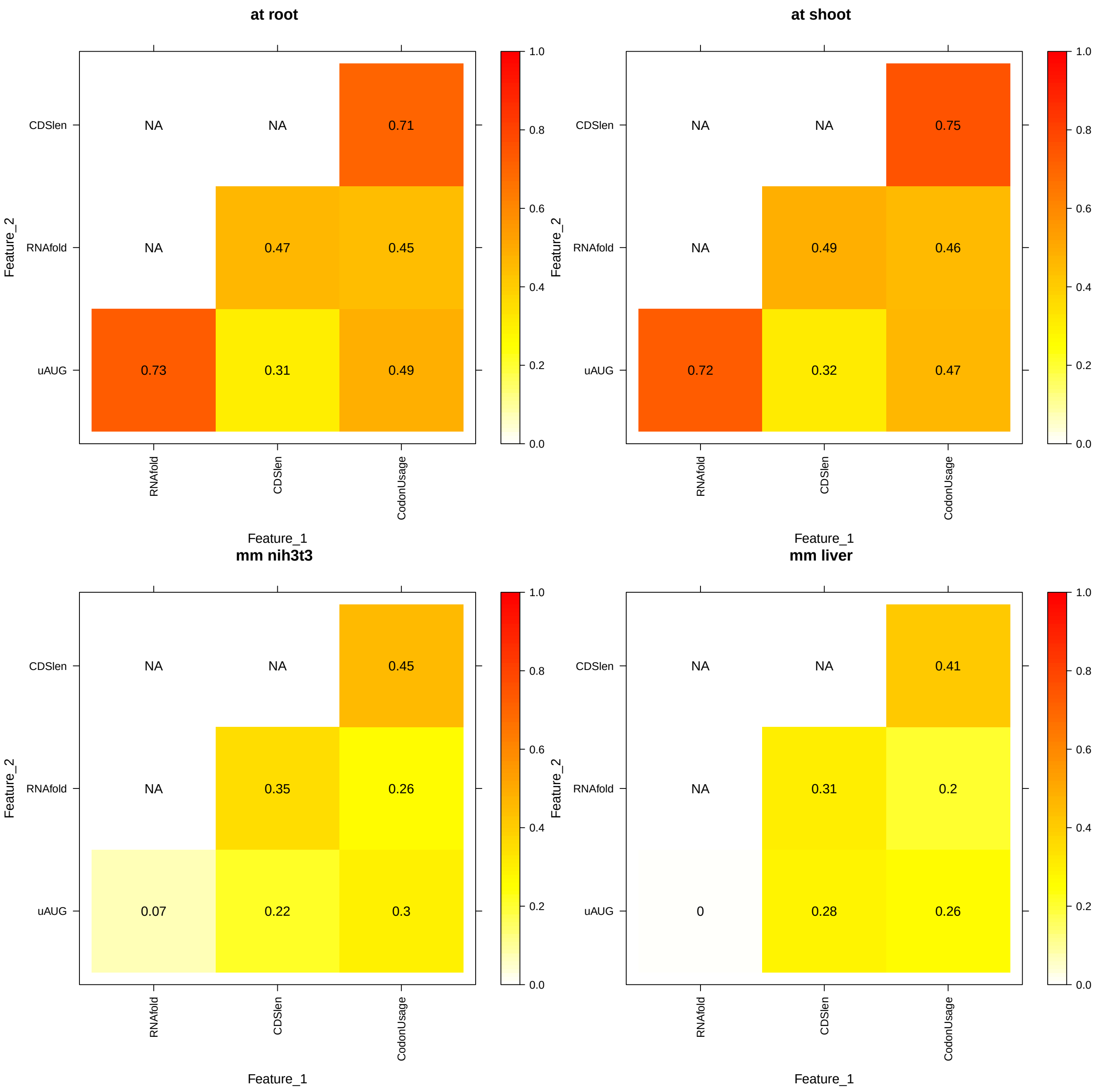
at root
at shoot
mm nih3t3
mm liver
Feature_2
Feature_1
CDSlen
RNAfold
uAUG
CodonUsage
NA
0.71
0.47
0.45
0.73
0.31
0.49
0.75
0.49
0.46
0.72
0.32
0.47
0.45
0.35
0.26
0.07
0.22
0.3
0.41
0.31
0.2
0
0.28
0.26
0.0
0.2
0.4
0.6
0.8
1.0

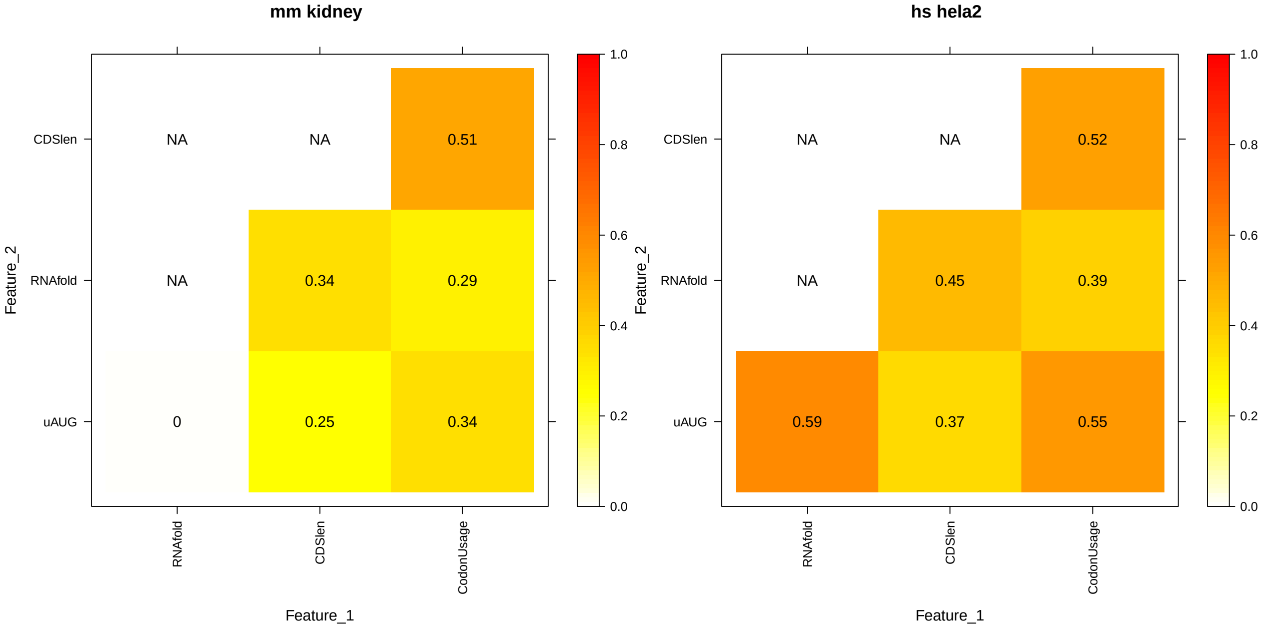


### Collinearity analysis: (codon frequencies in the 1st half of CDS) and (codon frequencies in the 2nd half of CDS)

```
for (species in names(species_tissue_list)) {
    for (species_tissue in species_tissue_list[[species]]) {
        if (species_tissue != "") {
            tissue_dot <- paste0(".", species_tissue)
            tissue_us <- paste0("_", species_tissue)
        } else {
            tissue_dot <- ""
            tissue_us <- ""
        }

        codon_features_1 <- readRDS(paste0("processed_data_w_polyA/codon_features/",
            species, tissue_dot, ".codon_features_1.rds"))[[1]]

        codon_features_2 <- readRDS(paste0("processed_data_w_polyA/codon_features/",
            species, tissue_dot, ".codon_features_2.rds"))[[1]]

        codon_features_1_2 <- cbind(codon_features_1, codon_features_2)

        # codon_features_1 vs. codon_features_2
        R2_features_1 <- summary(lm(get(paste0(species, tissue_dot,
            ".log10.TR")) ~ as.matrix(codon_features_1)))$r.squared
        R2_features_2 <- summary(lm(get(paste0(species, tissue_dot,
            ".log10.TR")) ~ as.matrix(codon_features_2)))$r.squared
        R2_features_1_2 <- summary(lm(get(paste0(species, tissue_dot,
            ".log10.TR")) ~ as.matrix(codon_features_1_2)))$r.squared

        cat(paste0(species, tissue_dot, "\nR2 of codon freq 1st + 2nd halves: "))
        cat(R2_features_1_2)
```

```r
        cat("\n")
        cat("% of codon freq 1st half: ")
        cat(round(R2_features_1/R2_features_1_2 * 100, 3))
        cat("%\n")
        cat("% of codon freq 2nd half: ")
        cat(round(R2_features_2/R2_features_1_2 * 100, 3))
        cat("%\n")
        cat("% of collinearity: ")
        cat(round((R2_features_1 + R2_features_2 - R2_features_1_2)/R2_features_1_2 *
            100, 3))
        cat("%\n\n")
    }
}
```

```
## sc
## R2 of codon freq 1st + 2nd halves: 0.602485
## % of codon freq 1st half: 91.431%
## % of codon freq 2nd half: 54.725%
## % of collinearity: 46.157%
##
## sp
## R2 of codon freq 1st + 2nd halves: 0.4122197
## % of codon freq 1st half: 85.174%
## % of codon freq 2nd half: 76.542%
## % of collinearity: 61.717%
##
## sp.alt
## R2 of codon freq 1st + 2nd halves: 0.5160581
## % of codon freq 1st half: 89.693%
## % of codon freq 2nd half: 70.353%
## % of collinearity: 60.046%
##
## at.leaf
## R2 of codon freq 1st + 2nd halves: 0.2179808
## % of codon freq 1st half: 89.793%
## % of codon freq 2nd half: 33.822%
## % of collinearity: 23.616%
##
## at.root
## R2 of codon freq 1st + 2nd halves: 0.2221062
## % of codon freq 1st half: 78.193%
## % of codon freq 2nd half: 43.456%
## % of collinearity: 21.65%
##
## at.shoot
## R2 of codon freq 1st + 2nd halves: 0.2151379
## % of codon freq 1st half: 74.259%
## % of codon freq 2nd half: 49.958%
## % of collinearity: 24.217%
##
## mm.nih3t3
## R2 of codon freq 1st + 2nd halves: 0.1306327
## % of codon freq 1st half: 87.92%
## % of codon freq 2nd half: 28.934%
```

```
## % of collinearity: 16.854%
##
## mm.liver
## R2 of codon freq 1st + 2nd halves: 0.1694238
## % of codon freq 1st half: 87.715%
## % of codon freq 2nd half: 34.86%
## % of collinearity: 22.575%
##
## mm.kidney
## R2 of codon freq 1st + 2nd halves: 0.1564376
## % of codon freq 1st half: 82.374%
## % of codon freq 2nd half: 36.98%
## % of collinearity: 19.354%
##
## hs.hela2
## R2 of codon freq 1st + 2nd halves: 0.2106567
## % of codon freq 1st half: 79.879%
## % of codon freq 2nd half: 39.31%
## % of collinearity: 19.189%
```

#### Figure 15. Collinearity analysis of min fold energy, 2nd min fold energy, and 3rd min fold energy

```
for (species in names(species_tissue_list)) {
    for (species_tissue in species_tissue_list[[species]]) {
        if (species_tissue != "") {
            tissue_dot <- paste0(".", species_tissue)
            tissue_us <- paste0("_", species_tissue)
        } else {
            tissue_dot <- ""
            tissue_us <- ""
        }

        # min fold energy vs. 2nd min fold energy
        features_1 <- readRDS(file = paste0("processed_data_w_polyA/min_2ndmin_3rdmin_fold_energy_featu[illegible]
            species, tissue_us, "_min_fold_energy_features.div.rds"))

        features_2 <- readRDS(file = paste0("processed_data_w_polyA/min_2ndmin_3rdmin_fold_energy_featu[illegible]
            species, tissue_us, "_second_min_fold_energy_features.div.rds"))

        features_1_2 <- cbind(features_1, features_2)

        nonNA_idx <- which(complete.cases(features_1_2))

        R2_features_1 <- summary(lm(get(paste0(species, tissue_dot,
            ".log10.TR"))[nonNA_idx] ~ as.matrix(features_1[nonNA_idx,
            ])))$r.squared
        R2_features_2 <- summary(lm(get(paste0(species, tissue_dot,
            ".log10.TR"))[nonNA_idx] ~ as.matrix(features_2[nonNA_idx,
            ])))$r.squared
        R2_features_1_2 <- summary(lm(get(paste0(species, tissue_dot,
```

```
    ".log10.TR"))[nonNA_idx] ~ as.matrix(features_1_2[nonNA_idx,
    ])))$r.squared

cat(paste0(species, tissue_dot, "\nR2 of min fold energy + 2nd min fold energy: "))
cat(R2_features_1_2)
cat("\n")
cat("% of min fold energy: ")
cat(round(R2_features_1/R2_features_1_2 * 100, 3))
cat("%\n")
cat("% of 2nd min fold energy: ")
cat(round(R2_features_2/R2_features_1_2 * 100, 3))
cat("%\n")
cat("% of collinearity: ")
cat(round((R2_features_1 + R2_features_2 - R2_features_1_2)/R2_features_1_2 *
    100, 3))
cat("%\n")
cat("# of genes: ")
cat(length(nonNA_idx))
cat("\n\n")

# min fold energy vs. 3rd min fold energy
features_1 <- readRDS(file = paste0("processed_data_w_polyA/min_2ndmin_3rdmin_fold_energy_featu[illegible]
    species, tissue_us, "_min_fold_energy_features.div.rds"))

features_2 <- readRDS(file = paste0("processed_data_w_polyA/min_2ndmin_3rdmin_fold_energy_featu[illegible]
    species, tissue_us, "_third_min_fold_energy_features.div.rds"))

features_1_2 <- cbind(features_1, features_2)

nonNA_idx <- which(complete.cases(features_1_2))

R2_features_1 <- summary(lm(get(paste0(species, tissue_dot,
    ".log10.TR"))[nonNA_idx] ~ as.matrix(features_1[nonNA_idx,
    ])))$r.squared
R2_features_2 <- summary(lm(get(paste0(species, tissue_dot,
    ".log10.TR"))[nonNA_idx] ~ as.matrix(features_2[nonNA_idx,
    ])))$r.squared
R2_features_1_2 <- summary(lm(get(paste0(species, tissue_dot,
    ".log10.TR"))[nonNA_idx] ~ as.matrix(features_1_2[nonNA_idx,
    ])))$r.squared

cat(paste0("R2 of min fold energy + 3rd min fold energy: "))
cat(R2_features_1_2)
cat("\n")
cat("% of min fold energy: ")
cat(round(R2_features_1/R2_features_1_2 * 100, 3))
cat("%\n")
cat("% of 3rd min fold energy: ")
cat(round(R2_features_2/R2_features_1_2 * 100, 3))
cat("%\n")
cat("% of collinearity: ")
cat(round((R2_features_1 + R2_features_2 - R2_features_1_2)/R2_features_1_2 *
    100, 3))
```

```
        cat("%\n")
        cat("# of genes: ")
        cat(length(nonNA_idx))
        cat("\n\n")

        # Pairwise Pearson correlations of min fold energy vs. 2nd
        # min fold energy vs. 3rd min fold energy
        min_2ndmin_3rdmin_fold_energy_features <- readRDS(file = paste0("processed_data_w_polyA/min_2ndm
            species, tissue_us, "_min_2ndmin_3rdmin_fold_energy_features"))[nonNA_idx,
            ]
        cat(paste0("Pearson cor of min fold energy vs. 2nd min fold energy: "))
        cat(cor(min_2ndmin_3rdmin_fold_energy_features[, 1],
            min_2ndmin_3rdmin_fold_energy_features[, 2]))
        cat("\n")
        cat(paste0("Pearson cor of min fold energy vs. 3rd min fold energy: "))
        cat(cor(min_2ndmin_3rdmin_fold_energy_features[, 1],
            min_2ndmin_3rdmin_fold_energy_features[, 3]))
        cat("\n")
        cat(paste0("Pearson cor of 2nd min fold energy vs. 3rd min fold energy: "))
        cat(cor(min_2ndmin_3rdmin_fold_energy_features[, 2],
            min_2ndmin_3rdmin_fold_energy_features[, 3]))
        cat("\n\n")
    }
}
```

```
## sc
## R2 of min fold energy + 2nd min fold energy: 0.231055
## % of min fold energy: 92.441%
## % of 2nd min fold energy: 65.687%
## % of collinearity: 58.128%
## # of genes: 1181
##
## R2 of min fold energy + 3rd min fold energy: 0.161843
## % of min fold energy: 73.857%
## % of 3rd min fold energy: 79.565%
## % of collinearity: 53.422%
## # of genes: 443
##
## Pearson cor of min fold energy vs. 2nd min fold energy: 0.6899256
## Pearson cor of min fold energy vs. 3rd min fold energy: 0.5354447
## Pearson cor of 2nd min fold energy vs. 3rd min fold energy: 0.7557849
##
## sp
## R2 of min fold energy + 2nd min fold energy: 0.02054973
## % of min fold energy: 76.864%
## % of 2nd min fold energy: 81.126%
## % of collinearity: 57.99%
## # of genes: 2784
##
## R2 of min fold energy + 3rd min fold energy: 0.02213057
## % of min fold energy: 75.083%
## % of 3rd min fold energy: 78.04%
## % of collinearity: 53.123%
## # of genes: 1706
```

```
##
## Pearson cor of min fold energy vs. 2nd min fold energy: 0.6814448
## Pearson cor of min fold energy vs. 3rd min fold energy: 0.531558
## Pearson cor of 2nd min fold energy vs. 3rd min fold energy: 0.7565556
##
## sp.alt
## R2 of min fold energy + 2nd min fold energy: 0.03801753
## % of min fold energy: 65.525%
## % of 2nd min fold energy: 94.061%
## % of collinearity: 59.586%
## # of genes: 3142
##
## R2 of min fold energy + 3rd min fold energy: 0.01841481
## % of min fold energy: 84.345%
## % of 3rd min fold energy: 66.27%
## % of collinearity: 50.614%
## # of genes: 2166
##
## Pearson cor of min fold energy vs. 2nd min fold energy: 0.6783187
## Pearson cor of min fold energy vs. 3rd min fold energy: 0.5178331
## Pearson cor of 2nd min fold energy vs. 3rd min fold energy: 0.778292
##
## at.leaf
## R2 of min fold energy + 2nd min fold energy: 0.1798378
## % of min fold energy: 86.978%
## % of 2nd min fold energy: 71.602%
## % of collinearity: 58.58%
## # of genes: 10350
##
## R2 of min fold energy + 3rd min fold energy: 0.1648275
## % of min fold energy: 82.115%
## % of 3rd min fold energy: 62.998%
## % of collinearity: 45.113%
## # of genes: 6740
##
## Pearson cor of min fold energy vs. 2nd min fold energy: 0.6033103
## Pearson cor of min fold energy vs. 3rd min fold energy: 0.4619936
## Pearson cor of 2nd min fold energy vs. 3rd min fold energy: 0.7588875
##
## at.root
## R2 of min fold energy + 2nd min fold energy: 0.120287
## % of min fold energy: 76.857%
## % of 2nd min fold energy: 81.919%
## % of collinearity: 58.775%
## # of genes: 7210
##
## R2 of min fold energy + 3rd min fold energy: 0.1184481
## % of min fold energy: 62.235%
## % of 3rd min fold energy: 81.543%
## % of collinearity: 43.778%
## # of genes: 4638
##
## Pearson cor of min fold energy vs. 2nd min fold energy: 0.5879342
## Pearson cor of min fold energy vs. 3rd min fold energy: 0.4483644
```

```
## Pearson cor of 2nd min fold energy vs. 3rd min fold energy: 0.7564651
##
## at.shoot
## R2 of min fold energy + 2nd min fold energy: 0.09450058
## % of min fold energy: 72.866%
## % of 2nd min fold energy: 86.199%
## % of collinearity: 59.065%
## # of genes: 9410
##
## R2 of min fold energy + 3rd min fold energy: 0.1022632
## % of min fold energy: 55.942%
## % of 3rd min fold energy: 86.429%
## % of collinearity: 42.37%
## # of genes: 6185
##
## Pearson cor of min fold energy vs. 2nd min fold energy: 0.5983952
## Pearson cor of min fold energy vs. 3rd min fold energy: 0.4508112
## Pearson cor of 2nd min fold energy vs. 3rd min fold energy: 0.7514525
##
## mm.nih3t3
## R2 of min fold energy + 2nd min fold energy: 0.09929118
## % of min fold energy: 95.664%
## % of 2nd min fold energy: 73.96%
## % of collinearity: 69.624%
## # of genes: 6002
##
## R2 of min fold energy + 3rd min fold energy: 0.08703582
## % of min fold energy: 96.28%
## % of 3rd min fold energy: 56.276%
## % of collinearity: 52.556%
## # of genes: 4200
##
## Pearson cor of min fold energy vs. 2nd min fold energy: 0.7646191
## Pearson cor of min fold energy vs. 3rd min fold energy: 0.6085559
## Pearson cor of 2nd min fold energy vs. 3rd min fold energy: 0.7971894
##
## mm.liver
## R2 of min fold energy + 2nd min fold energy: 0.09753985
## % of min fold energy: 96.99%
## % of 2nd min fold energy: 70.491%
## % of collinearity: 67.482%
## # of genes: 5181
##
## R2 of min fold energy + 3rd min fold energy: 0.09311743
## % of min fold energy: 97.905%
## % of 3rd min fold energy: 50.463%
## % of collinearity: 48.368%
## # of genes: 3621
##
## Pearson cor of min fold energy vs. 2nd min fold energy: 0.7669807
## Pearson cor of min fold energy vs. 3rd min fold energy: 0.6010318
## Pearson cor of 2nd min fold energy vs. 3rd min fold energy: 0.7904313
##
## mm.kidney
```

```
## R2 of min fold energy + 2nd min fold energy: 0.125633
## % of min fold energy: 95.767%
## % of 2nd min fold energy: 73.422%
## % of collinearity: 69.189%
## # of genes: 5568
##
## R2 of min fold energy + 3rd min fold energy: 0.1111678
## % of min fold energy: 96.434%
## % of 3rd min fold energy: 55.475%
## % of collinearity: 51.909%
## # of genes: 3919
##
## Pearson cor of min fold energy vs. 2nd min fold energy: 0.764149
## Pearson cor of min fold energy vs. 3rd min fold energy: 0.6054119
## Pearson cor of 2nd min fold energy vs. 3rd min fold energy: 0.7959718
##
## hs.hela2
## R2 of min fold energy + 2nd min fold energy: 0.03642137
## % of min fold energy: 98.458%
## % of 2nd min fold energy: 66.216%
## % of collinearity: 64.674%
## # of genes: 4393
##
## R2 of min fold energy + 3rd min fold energy: 0.02758912
## % of min fold energy: 95.291%
## % of 3rd min fold energy: 62.097%
## % of collinearity: 57.388%
## # of genes: 3018
##
## Pearson cor of min fold energy vs. 2nd min fold energy: 0.7792172
## Pearson cor of min fold energy vs. 3rd min fold energy: 0.6356418
## Pearson cor of 2nd min fold energy vs. 3rd min fold energy: 0.8176511
```

### Figure 16A. Collinearity analysis of aa freqs, codon freqs, and synonymous codon ratios

```r
# collinearity between codon freqs, aa freqs, and syn codon
# ratios
for (species in names(species_tissue_list)) {
    for (i in 1:length(species_tissue_list[[species]])) {
        species_tissue <- species_tissue_list[[species]][i]
        if (species_tissue != "") {
            tissue_dot <- paste0(".", species_tissue)
            tissue_us <- paste0("_", species_tissue)
        } else {
            tissue_dot <- ""
            tissue_us <- ""
        }

        aa_features <- readRDS(paste0("processed_data_w_polyA/aa_features/",
            species, tissue_dot, ".aa_features.rds"))[[1]]
```

```r
codon_features <- readRDS(paste0("processed_data_w_polyA/codon_features/",
    species, tissue_dot, ".codon_features.rds"))[[1]]
syncodon_features <- readRDS(paste0("processed_data_w_polyA/syncodon_features/",
    species, tissue_dot, ".syncodon_features.rds"))[[1]]

keep_idx <- which(apply(syncodon_features, 1, FUN = function(x) !(any(x ==
    Inf | is.na(x)))))

log10.CDS_lens <- get(paste0(species, tissue_dot, ".log10.CDS_lens"))

cat(paste0(species, tissue_dot, "\n"))
cat(paste0("# genes with non-Inf syn codon ratios: ",
    length(keep_idx), "\n"))
cat(paste0("median CDS length of included genes: ", median(10^log10.CDS_lens[keep_idx]),
    "\n"))
cat(paste0("median CDS length of excluded genes: ", median(10^log10.CDS_lens[-keep_idx]),
    "\n"))

aa_syncodon_features <- cbind(aa_features, syncodon_features)
codon_syncodon_features <- cbind(codon_features, syncodon_features)

R2_aa_features <- summary(lm(get(paste0(species, tissue_dot,
    ".log10.TR"))[keep_idx] ~ as.matrix(aa_features[keep_idx,
    ])))$r.squared
R2_syncodon_features <- summary(lm(get(paste0(species,
    tissue_dot, ".log10.TR"))[keep_idx] ~ as.matrix(syncodon_features[keep_idx,
    ])))$r.squared
R2_aa_syncodon_features <- summary(lm(get(paste0(species,
    tissue_dot, ".log10.TR"))[keep_idx] ~ as.matrix(aa_syncodon_features[keep_idx,
    ])))$r.squared

cat(paste0("R2 of aa_features + syncodon_features: "))
cat(R2_aa_syncodon_features)
cat("\n")
cat("% of aa_features: ")
cat(round(R2_aa_features/R2_aa_syncodon_features * 100,
    3))
cat("%\n")
cat("% of syncodon_features: ")
cat(round(R2_syncodon_features/R2_aa_syncodon_features *
    100, 3))
cat("%\n")
cat("% of collinearity: ")
cat(round((R2_aa_features + R2_syncodon_features - R2_aa_syncodon_features)/R2_aa_syncodon_featu
    100, 3))
cat("%\n\n")

R2_codon_features <- summary(lm(get(paste0(species, tissue_dot,
    ".log10.TR"))[keep_idx] ~ as.matrix(codon_features[keep_idx,
    ])))$r.squared
R2_codon_syncodon_features <- summary(lm(get(paste0(species,
    tissue_dot, ".log10.TR"))[keep_idx] ~ as.matrix(codon_syncodon_features[keep_idx,
    ])))$r.squared
```

```
        cat(paste0("R2 of codon_features + syncodon_features: "))
        cat(R2_codon_syncodon_features)
        cat("\n")
        cat("% of codon_features: ")
        cat(round(R2_codon_features/R2_codon_syncodon_features *
            100, 3))
        cat("%\n")
        cat("% of syncodon_features: ")
        cat(round(R2_syncodon_features/R2_codon_syncodon_features *
            100, 3))
        cat("%\n")
        cat("% of collinearity: ")
        cat(round((R2_codon_features + R2_syncodon_features -
            R2_codon_syncodon_features)/R2_codon_syncodon_features *
            100, 3))
        cat("%\n\n")
    }
}
```

```
## sc
## # genes with non-Inf syn codon ratios: 2022
## median CDS length of included genes: 1357.5
## median CDS length of excluded genes: 639
## R2 of aa_features + syncodon_features: 0.573553
## % of aa_features: 64.977%
## % of syncodon_features: 84.913%
## % of collinearity: 49.89%
##
## R2 of codon_features + syncodon_features: 0.606262
## % of codon_features: 93.423%
## % of syncodon_features: 80.332%
## % of collinearity: 73.755%
##
## sp
## # genes with non-Inf syn codon ratios: 3268
## median CDS length of included genes: 1401
## median CDS length of excluded genes: 531
## R2 of aa_features + syncodon_features: 0.4057041
## % of aa_features: 68.713%
## % of syncodon_features: 80.684%
## % of collinearity: 49.396%
##
## R2 of codon_features + syncodon_features: 0.4266632
## % of codon_features: 94.752%
## % of syncodon_features: 76.72%
## % of collinearity: 71.472%
##
## sp.alt
## # genes with non-Inf syn codon ratios: 3267
## median CDS length of included genes: 1401
## median CDS length of excluded genes: 529.5
## R2 of aa_features + syncodon_features: 0.4881939
## % of aa_features: 75.795%
```

```
## % of syncodon_features: 80.612%
## % of collinearity: 56.408%
##
## R2 of codon_features + syncodon_features: 0.5047257
## % of codon_features: 96.915%
## % of syncodon_features: 77.972%
## % of collinearity: 74.887%
##
## at.leaf
## # genes with non-Inf syn codon ratios: 10666
## median CDS length of included genes: 1233
## median CDS length of excluded genes: 480
## R2 of aa_features + syncodon_features: 0.2027222
## % of aa_features: 57.599%
## % of syncodon_features: 66.916%
## % of collinearity: 24.514%
##
## R2 of codon_features + syncodon_features: 0.2168145
## % of codon_features: 96.363%
## % of syncodon_features: 62.567%
## % of collinearity: 58.93%
##
## at.root
## # genes with non-Inf syn codon ratios: 7417
## median CDS length of included genes: 1251
## median CDS length of excluded genes: 486
## R2 of aa_features + syncodon_features: 0.2407765
## % of aa_features: 70.505%
## % of syncodon_features: 47.812%
## % of collinearity: 18.318%
##
## R2 of codon_features + syncodon_features: 0.2656062
## % of codon_features: 93.687%
## % of syncodon_features: 43.343%
## % of collinearity: 37.03%
##
## at.shoot
## # genes with non-Inf syn codon ratios: 9690
## median CDS length of included genes: 1263
## median CDS length of excluded genes: 507
## R2 of aa_features + syncodon_features: 0.2234242
## % of aa_features: 72.538%
## % of syncodon_features: 45.056%
## % of collinearity: 17.594%
##
## R2 of codon_features + syncodon_features: 0.2464026
## % of codon_features: 93.97%
## % of syncodon_features: 40.854%
## % of collinearity: 34.824%
##
## mm.nih3t3
## # genes with non-Inf syn codon ratios: 6590
## median CDS length of included genes: 1590
## median CDS length of excluded genes: 591
```

```
## R2 of aa_features + syncodon_features: 0.1370316
## % of aa_features: 75.73%
## % of syncodon_features: 38.593%
## % of collinearity: 14.322%
##
## R2 of codon_features + syncodon_features: 0.157992
## % of codon_features: 92.276%
## % of syncodon_features: 33.473%
## % of collinearity: 25.749%
##
## mm.liver
## # genes with non-Inf syn codon ratios: 5697
## median CDS length of included genes: 1566
## median CDS length of excluded genes: 588
## R2 of aa_features + syncodon_features: 0.1714909
## % of aa_features: 61.73%
## % of syncodon_features: 58.377%
## % of collinearity: 20.106%
##
## R2 of codon_features + syncodon_features: 0.1914157
## % of codon_features: 93.823%
## % of syncodon_features: 52.3%
## % of collinearity: 46.123%
##
## mm.kidney
## # genes with non-Inf syn codon ratios: 6121
## median CDS length of included genes: 1578
## median CDS length of excluded genes: 594
## R2 of aa_features + syncodon_features: 0.1605799
## % of aa_features: 84.525%
## % of syncodon_features: 31.156%
## % of collinearity: 15.681%
##
## R2 of codon_features + syncodon_features: 0.1775161
## % of codon_features: 93.461%
## % of syncodon_features: 28.183%
## % of collinearity: 21.644%
##
## hs.hela2
## # genes with non-Inf syn codon ratios: 4896
## median CDS length of included genes: 1519.5
## median CDS length of excluded genes: 465
## R2 of aa_features + syncodon_features: 0.1801226
## % of aa_features: 82.837%
## % of syncodon_features: 32.888%
## % of collinearity: 15.725%
##
## R2 of codon_features + syncodon_features: 0.2009149
## % of codon_features: 93.589%
## % of syncodon_features: 29.484%
## % of collinearity: 23.074%
```

```
# calculate the correlation between each syn codon's ratio
# and its corresponding AA's freq
```

```
for (species in names(species_tissue_list)) {
    for (i in 1:length(species_tissue_list[[species]])) {
        species_tissue <- species_tissue_list[[species]][i]
        if (species_tissue != "") {
            tissue_dot <- paste0(".", species_tissue)
            tissue_us <- paste0("_", species_tissue)
        } else {
            tissue_dot <- ""
            tissue_us <- ""
        }
        ## count the frequency of 64 codons (including 3 stop codons)
        ## in every gene
        CDS_seqs <- get(paste0(species, tissue_dot, ".CDS_seqs"))
        codon_counts <- t(sapply(CDS_seqs, FUN = function(seq) {
            sst <- strsplit(seq, "")[[1]]
            n <- length(sst)
            out <- paste0(sst[seq(from = 1, to = n, by = 3)],
                sst[seq(from = 2, to = n, by = 3)], sst[seq(from = 3,
                  to = n, by = 3)])
            table(out)[tri_nts]
        }))
        codon_counts[is.na(codon_counts)] <- 0
        codon_freqs <- codon_counts/rowSums(codon_counts)
        rownames(codon_freqs) <- names(CDS_seqs)
        colnames(codon_freqs) <- tri_nts

        ## count the frequency of 20 amino acids (AAs) in every gene
        aa_counts <- sapply(aas, FUN = function(aa) {
            idx <- which(tri_nts_to_aas == aa)
            if (length(idx) == 1) {
                codon_counts[, idx]
            } else {
                rowSums(codon_counts[, idx])
            }
        })
        aa_freqs <- aa_counts/rowSums(aa_counts)
        rownames(aa_freqs) <- names(CDS_seqs)
        colnames(aa_freqs) <- aas

        ## normalize the codon counts by their corresponding AA count
        syncodon_ratios <- codon_freqs[, nonstop_codons]
        for (aa in aas) {
            tmp_codons <- names(tri_nts_to_aas[which(tri_nts_to_aas ==
                aa)])
            syncodon_ratios[, tmp_codons] <- codon_freqs[, tmp_codons]/rowSums(codon_freqs[,
                tmp_codons, drop = F])
        }

        cor_codon_aa <- data.frame(codon = nonstop_codons, aa = tri_nts_to_aas[nonstop_codons],
            cor = sapply(nonstop_codons, FUN = function(codon) {
                x <- syncodon_ratios[, codon]
                y <- aa_freqs[, tri_nts_to_aas[codon]]
                keep_idx <- which(x != Inf & !is.na(x))
```

```
                cor(x[keep_idx], y[keep_idx])
            }))

        cat(paste0("\n", species, tissue_dot, "\n"))
        cat(paste0("# genes with non-Inf syn codon ratios: ",
            sum(apply(syncodon_ratios, 1, FUN = function(z) !any(z ==
                Inf | is.na(z)))), "\n"))
        print(cor_codon_aa)
    }
}
```

### Figure 16B. Collinearity analysis of four features: aa freqs (or codon freqs, or synonymous codon ratios), CDS length, 5'RNA fold, and uAUG

```
## A function to calculate pairwise collinearity of every two
## features
calc_syncodon_vs_features_collinearity <- function(features_list,
    syncodon_features, imp_feature_name, y) {
    p <- length(features_list)
    feature_names <- names(features_list)

    syncodon_R2 <- summary(lm(y ~ as.matrix(syncodon_features)))$r.squared

    uni_feature_R2s <- sapply(1:p, FUN = function(i) {
        tmp_features_all <- cbind(y, features_list[[i]])
        colnames(tmp_features_all)[1] <- "y"
        ## remove duplicated column names
        tmp_features_all <- tmp_features_all[, !duplicated(colnames(tmp_features_all))]
        tmp_features_all <- as.data.frame(tmp_features_all)
        ## model
        tmp_features_model <- lm(y ~ ., data = tmp_features_all)
        summary(tmp_features_model)$r.squared
    })
    names(uni_feature_R2s) <- feature_names

    bi_feature_R2s <- sapply(1:p, FUN = function(i) {
        tmp_features_all <- cbind(y, features_list[[i]], syncodon_features)
        colnames(tmp_features_all)[1] <- "y"
        ## remove duplicated column names
        tmp_features_all <- tmp_features_all[, !duplicated(colnames(tmp_features_all))]
        tmp_features_all <- as.data.frame(tmp_features_all)
        ## model
        tmp_features_model <- lm(y ~ ., data = tmp_features_all)
        summary(tmp_features_model)$r.squared
    })
    names(bi_feature_R2s) <- paste(imp_feature_name, feature_names,
        sep = "-")

    result <- matrix(NA, p, 1)
```

```r
    rownames(result) <- feature_names
    # the order of bi_feature_R2s should correspond to the
    # columns of feature_idx_combn
    for (i in 1:p) {
        redundancy <- syncodon_R2 + uni_feature_R2s[i] - bi_feature_R2s[i]
        redundancy <- max(0, redundancy)
        result[i, 1] <- max(round(redundancy/syncodon_R2 * 100,
            3), round(redundancy/uni_feature_R2s[i] * 100, 3))
    }

    cat(paste0("R2 of ", imp_feature_name, ": "))
    cat(syncodon_R2)
    cat("\n")
    for (i in 1:p) {
        feature_name <- feature_names[i]
        cat(paste0("R2 of ", feature_name, ": "))
        cat(uni_feature_R2s[i])
        cat("\n")
        cat(paste0("R2 of ", imp_feature_name, " + ", feature_name,
            ": "))
        cat(bi_feature_R2s[i])
        cat("\n")
    }
    cat("\n")
    result
}

## A function to make levelplot showing values
myPanel <- function(x, y, z, ...) {
    panel.levelplot(x, y, z, ...)
    panel.text(x, y, round(z, 2))
}

for (species in names(species_tissue_list)) {
    for (species_tissue in species_tissue_list[[species]]) {
        if (species_tissue != "") {
            tissue_dot <- paste0(".", species_tissue)
            tissue_us <- paste0("_", species_tissue)
        } else {
            tissue_dot <- ""
            tissue_us <- ""
        }

        aa_features <- readRDS(paste0("processed_data_w_polyA/aa_features/",
            species, tissue_dot, ".aa_features.rds"))[[1]]
        codon_features <- readRDS(paste0("processed_data_w_polyA/codon_features/",
            species, tissue_dot, ".codon_features.rds"))[[1]]
        syncodon_features <- readRDS(paste0("processed_data_w_polyA/syncodon_features/",
            species, tissue_dot, ".syncodon_features.rds"))[[1]]

        # only keep the genes whose syncodon features are all non-Inf
        # and non-NA
        keep_idx <- which(apply(syncodon_features, 1, FUN = function(x) !(any(x ==
```

```
    Inf | is.na(x)))))
# keep_idx <- 1:nrow(syncodon_features) # all genes keep_idx
# <- which(apply(syncodon_features, 1, FUN=function(x)
# any(x==Inf | is.na(x)))) # removed genes

aa_features <- aa_features[keep_idx, ]
codon_features <- codon_features[keep_idx, ]
syncodon_features <- syncodon_features[keep_idx, ]
features_list <- list(uAUG = get(paste0(species, tissue_dot,
    ".uAUG_counts.div"))[keep_idx, ], RNAfold = get(paste0(species,
    tissue_dot, ".fold_energy_features_selected"))[keep_idx,
    ], CDSlen = get(paste0(species, tissue_dot, ".log10.CDS_lens.div"))[keep_idx,
    ])

cat(paste0(species, tissue_dot, "\n"))

aa_tmp <- calc_syncodon_vs_features_collinearity(features_list,
    aa_features, "aa freqs", get(paste0(species, tissue_dot,
        ".log10.TR"))[keep_idx])

colnames(aa_tmp) <- "AA freqs"
aa_tmp[aa_tmp > 100] <- 100  # truncate the value at 100
assign(x = paste0(species, tissue_dot, ".aa_collinearity"),
    value = aa_tmp/100)

codon_tmp <- calc_syncodon_vs_features_collinearity(features_list,
    codon_features, "codon freqs", get(paste0(species,
        tissue_dot, ".log10.TR"))[keep_idx])
colnames(codon_tmp) <- "codon ratios"
codon_tmp[codon_tmp > 100] <- 100  # truncate the value at 100
assign(x = paste0(species, tissue_dot, ".codon_collinearity"),
    value = codon_tmp/100)

syncodon_tmp <- calc_syncodon_vs_features_collinearity(features_list,
    syncodon_features, "syncodon ratios", get(paste0(species,
        tissue_dot, ".log10.TR"))[keep_idx])
colnames(syncodon_tmp) <- "syncodon ratios"
syncodon_tmp[syncodon_tmp > 100] <- 100  # truncate the value at 100
assign(x = paste0(species, tissue_dot, ".syncodon_collinearity"),
    value = syncodon_tmp/100)

aa_grid <- setNames(melt(get(paste0(species, tissue_dot,
    ".aa_collinearity"))), c("Feature_1", "Feature_2",
    "Collinearity"))
print(levelplot(Collinearity ~ Feature_1 * Feature_2,
    aa_grid, panel = myPanel, scales = list(x = list(rot = 90)),
    main = paste(species, species_tissue, "aa freqs"),
    col.regions = rev(heat.colors(120)), cuts = 100,
    at = seq(0, 1, 0.01)))

codon_grid <- setNames(melt(get(paste0(species, tissue_dot,
    ".codon_collinearity"))), c("Feature_1", "Feature_2",
    "Collinearity"))
```

```
        print(levelplot(Collinearity ~ Feature_1 * Feature_2,
            codon_grid, panel = myPanel, scales = list(x = list(rot = 90)),
            main = paste(species, species_tissue, "codon ratios"),
            col.regions = rev(heat.colors(120)), cuts = 100,
            at = seq(0, 1, 0.01)))

        syncodon_grid <- setNames(melt(get(paste0(species, tissue_dot,
            ".syncodon_collinearity"))), c("Feature_1", "Feature_2",
            "Collinearity"))
        print(levelplot(Collinearity ~ Feature_1 * Feature_2,
            syncodon_grid, panel = myPanel, scales = list(x = list(rot = 90)),
            main = paste(species, species_tissue, "syncodon ratios"),
            col.regions = rev(heat.colors(120)), cuts = 100,
            at = seq(0, 1, 0.01)))

        filename <- paste0("tables/pairwise_collinearity/", species,
            tissue_dot, ".aa_collinearity.csv")
        if (!file.exists(filename)) {
            write.csv(get(paste0(species, tissue_dot, ".aa_collinearity"))/100,
                file = filename, quote = F)
        }

        filename <- paste0("tables/pairwise_collinearity/", species,
            tissue_dot, ".codon_collinearity.csv")
        if (!file.exists(filename)) {
            write.csv(get(paste0(species, tissue_dot, ".codon_collinearity"))/100,
                file = filename, quote = F)
        }

        filename <- paste0("tables/pairwise_collinearity/", species,
            tissue_dot, ".syncodon_collinearity.csv")
        if (!file.exists(filename)) {
            write.csv(get(paste0(species, tissue_dot, ".syncodon_collinearity"))/100,
                file = filename, quote = F)
        }
    }
}
```

```
## sc
## R2 of aa freqs: 0.372675
## R2 of uAUG: 0.06725848
## R2 of aa freqs + uAUG: 0.4055211
## R2 of RNAfold: 0.3040088
## R2 of aa freqs + RNAfold: 0.5233637
## R2 of CDSlen: 0.231087
## R2 of aa freqs + CDSlen: 0.5247433
##
## R2 of codon freqs: 0.5663885
## R2 of uAUG: 0.06725848
## R2 of codon freqs + uAUG: 0.5843258
## R2 of RNAfold: 0.3040088
## R2 of codon freqs + RNAfold: 0.6494725
## R2 of CDSlen: 0.231087
## R2 of codon freqs + CDSlen: 0.7263148
```

```
##
## R2 of syncodon ratios: 0.4870224
## R2 of uAUG: 0.06725848
## R2 of syncodon ratios + uAUG: 0.5060519
## R2 of RNAfold: 0.3040088
## R2 of syncodon ratios + RNAfold: 0.5835885
## R2 of CDSlen: 0.231087
## R2 of syncodon ratios + CDSlen: 0.6591315
```

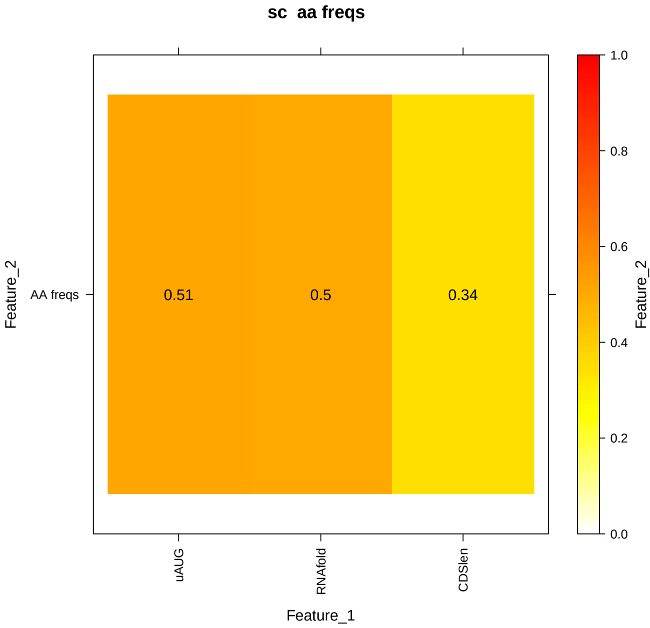


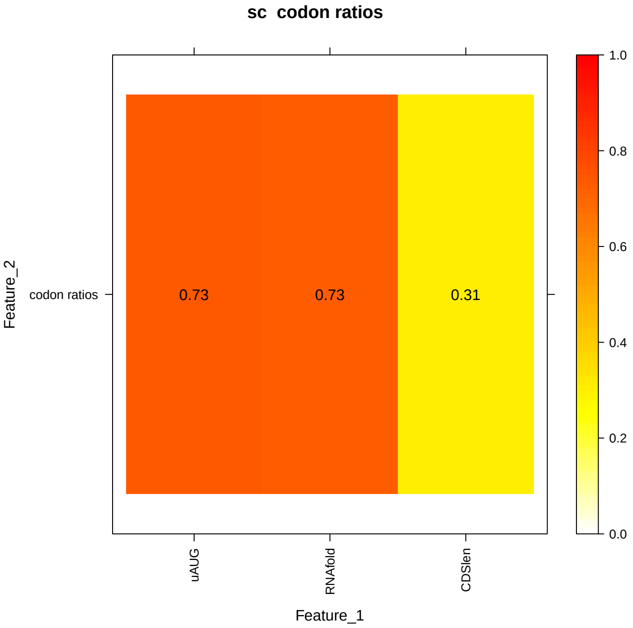


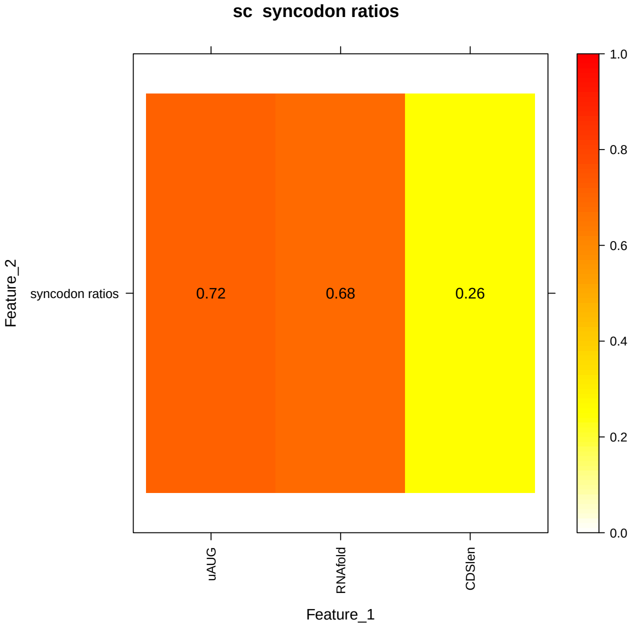


```
## sp
## R2 of aa freqs: 0.2787707
## R2 of uAUG: 0.05094277
## R2 of aa freqs + uAUG: 0.3146855
## R2 of RNAfold: 0.1034295
## R2 of aa freqs + RNAfold: 0.3193937
```

```
## R2 of CDSlen: 0.1260509
## R2 of aa freqs + CDSlen: 0.3625252
##
## R2 of codon freqs: 0.4042725
## R2 of uAUG: 0.05094277
## R2 of codon freqs + uAUG: 0.4318372
## R2 of RNAfold: 0.1034295
## R2 of codon freqs + RNAfold: 0.4303458
## R2 of CDSlen: 0.1260509
## R2 of codon freqs + CDSlen: 0.4782965
##
## R2 of syncodon ratios: 0.3273366
## R2 of uAUG: 0.05094277
## R2 of syncodon ratios + uAUG: 0.3559184
## R2 of RNAfold: 0.1034295
## R2 of syncodon ratios + RNAfold: 0.3623074
## R2 of CDSlen: 0.1260509
## R2 of syncodon ratios + CDSlen: 0.4077895
```

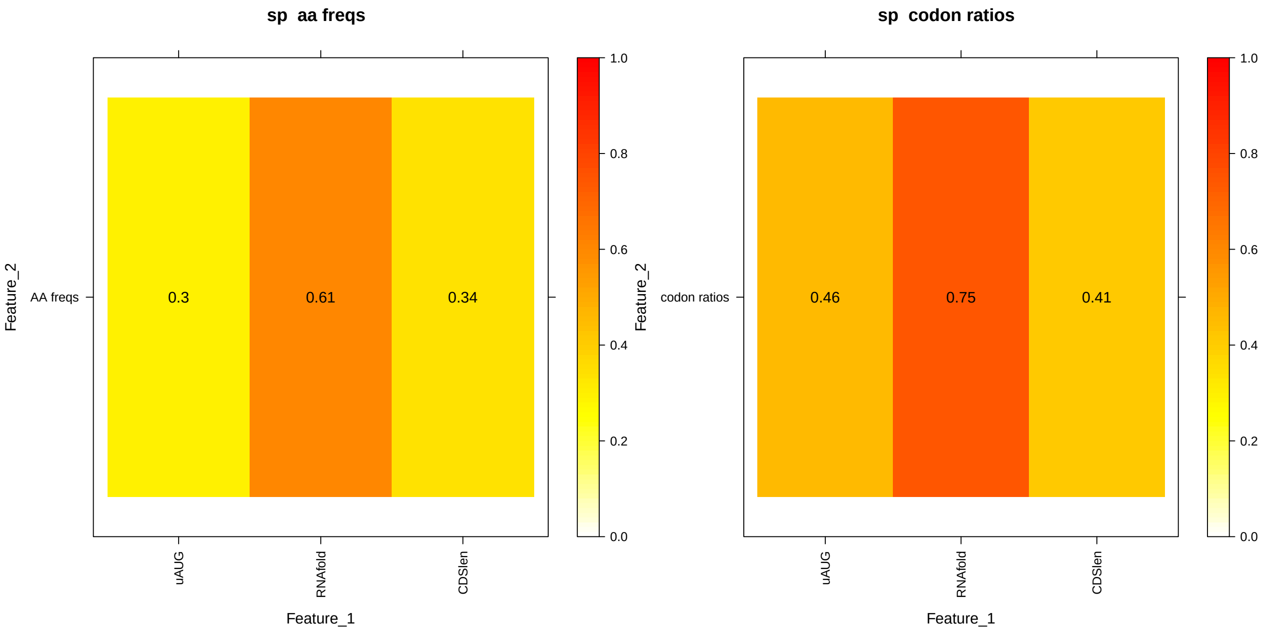

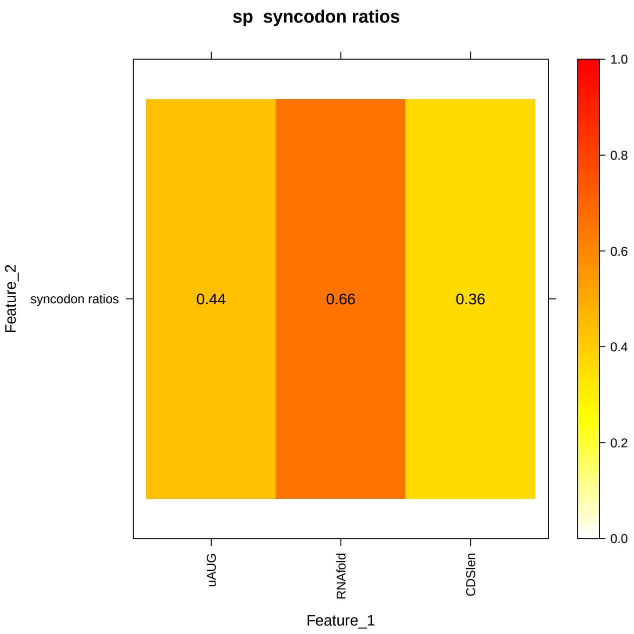


```
## sp.alt
## R2 of aa freqs: 0.370027
## R2 of uAUG: 0.03773724
## R2 of aa freqs + uAUG: 0.3949289
## R2 of RNAfold: 0.09486063
## R2 of aa freqs + RNAfold: 0.3855169
## R2 of CDSlen: 0.2573903
## R2 of aa freqs + CDSlen: 0.5128242
##
## R2 of codon freqs: 0.4891536
## R2 of uAUG: 0.03773724
## R2 of codon freqs + uAUG: 0.50684
## R2 of RNAfold: 0.09486063
## R2 of codon freqs + RNAfold: 0.4863781
## R2 of CDSlen: 0.2573903
## R2 of codon freqs + CDSlen: 0.6231069
##
## R2 of syncodon ratios: 0.393545
## R2 of uAUG: 0.03773724
## R2 of syncodon ratios + uAUG: 0.4113897
## R2 of RNAfold: 0.09486063
## R2 of syncodon ratios + RNAfold: 0.4010344
## R2 of CDSlen: 0.2573903
## R2 of syncodon ratios + CDSlen: 0.5713376
```

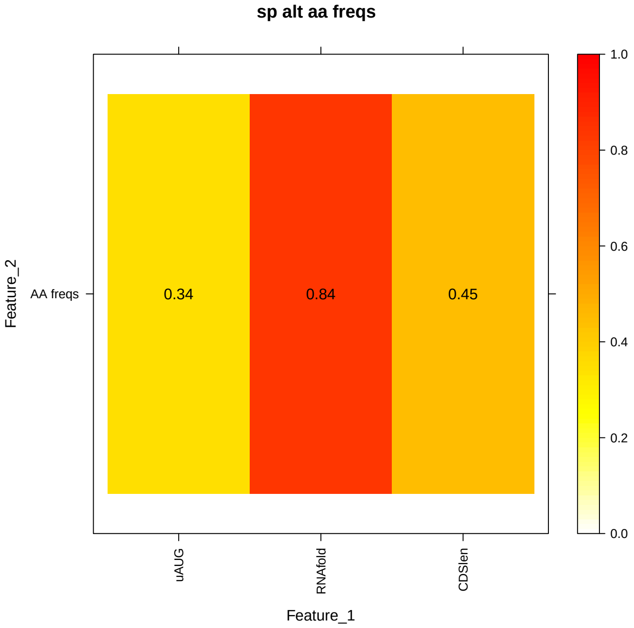


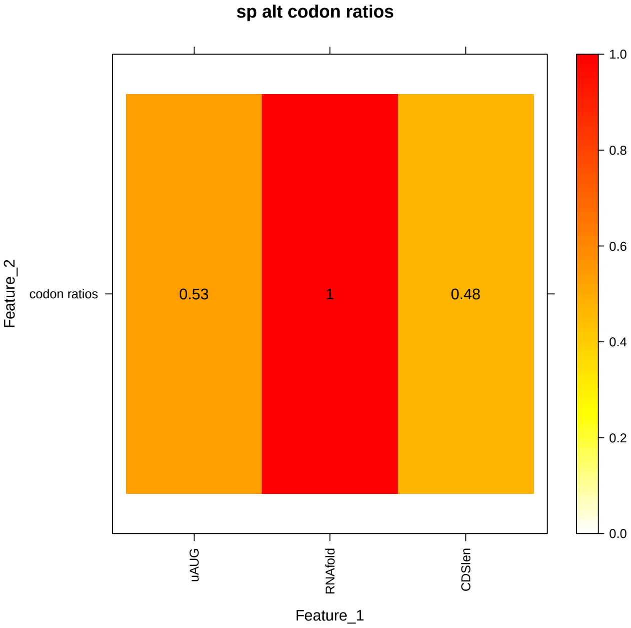


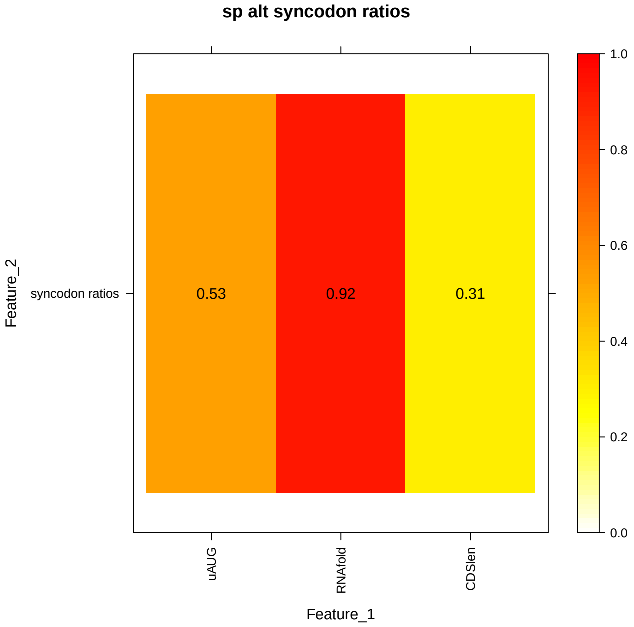


```
## at.leaf
## R2 of aa freqs: 0.1167652
## R2 of uAUG: 0.06804122
## R2 of aa freqs + uAUG: 0.1690409
## R2 of RNAfold: 0.227317
## R2 of aa freqs + RNAfold: 0.3064541
## R2 of CDSlen: 0.08508482
## R2 of aa freqs + CDSlen: 0.1869004
##
## R2 of codon freqs: 0.2089295
## R2 of uAUG: 0.06804122
## R2 of codon freqs + uAUG: 0.2483186
## R2 of RNAfold: 0.227317
## R2 of codon freqs + RNAfold: 0.3573116
```

```
## R2 of CDSlen: 0.08508482
## R2 of codon freqs + CDSlen: 0.2411095
##
## R2 of syncodon ratios: 0.1356532
## R2 of uAUG: 0.06804122
## R2 of syncodon ratios + uAUG: 0.1838037
## R2 of RNAfold: 0.227317
## R2 of syncodon ratios + RNAfold: 0.3024857
## R2 of CDSlen: 0.08508482
## R2 of syncodon ratios + CDSlen: 0.1681209
```

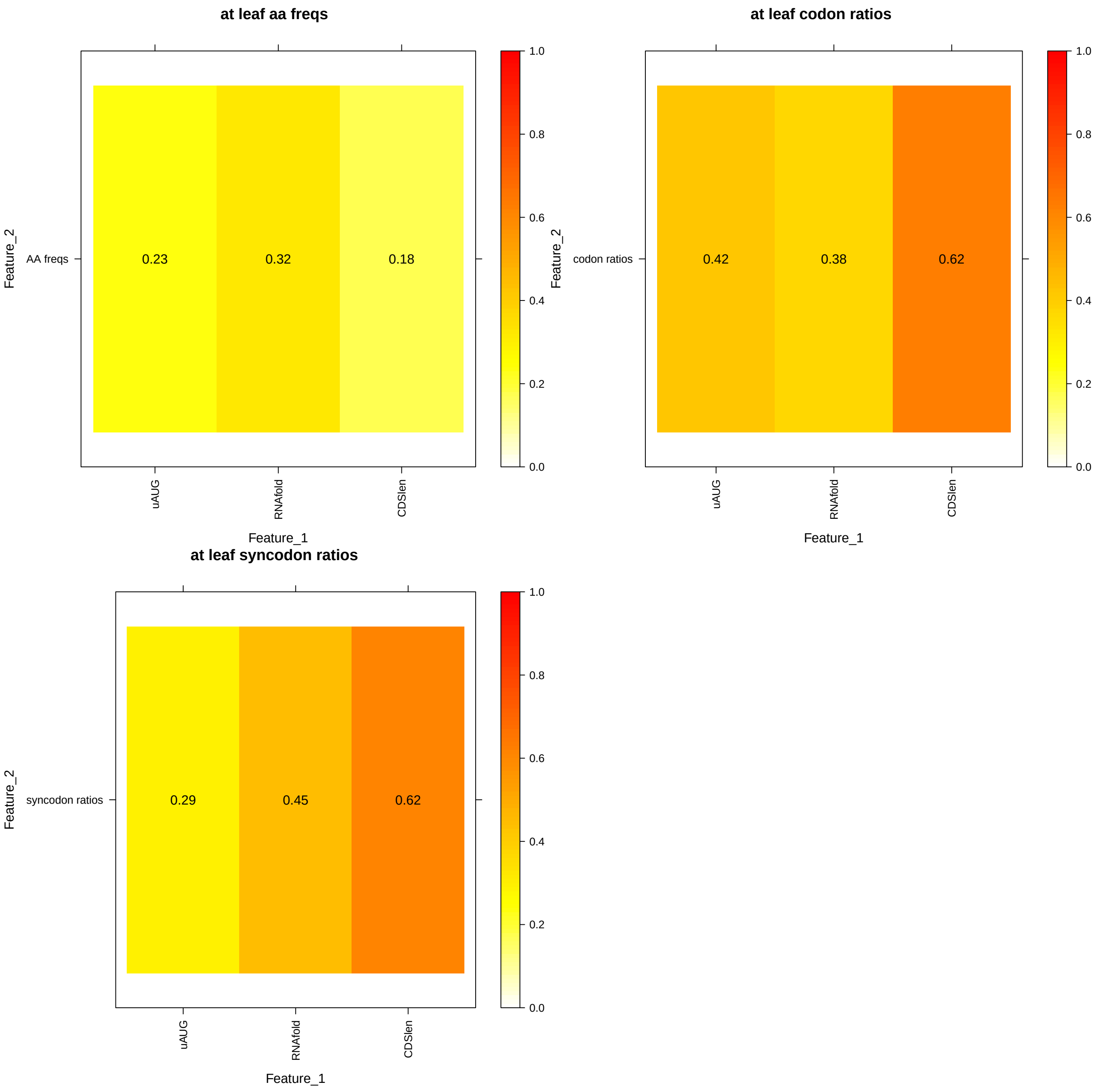


```
## at.root
## R2 of aa freqs: 0.1697607
## R2 of uAUG: 0.0967943
## R2 of aa freqs + uAUG: 0.2326369
```

```
## R2 of RNAfold: 0.1612343
## R2 of aa freqs + RNAfold: 0.2900553
## R2 of CDSlen: 0.08637703
## R2 of aa freqs + CDSlen: 0.2315243
##
## R2 of codon freqs: 0.2488384
## R2 of uAUG: 0.0967943
## R2 of codon freqs + uAUG: 0.2990818
## R2 of RNAfold: 0.1612343
## R2 of codon freqs + RNAfold: 0.3387577
## R2 of CDSlen: 0.08637703
## R2 of codon freqs + CDSlen: 0.2805797
##
## R2 of syncodon ratios: 0.115121
## R2 of uAUG: 0.0967943
## R2 of syncodon ratios + uAUG: 0.1882217
## R2 of RNAfold: 0.1612343
## R2 of syncodon ratios + RNAfold: 0.2343587
## R2 of CDSlen: 0.08637703
## R2 of syncodon ratios + CDSlen: 0.1553111
```

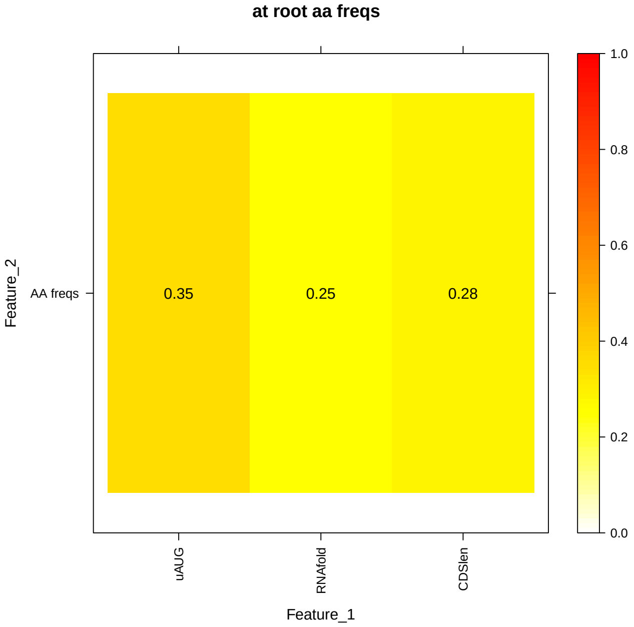


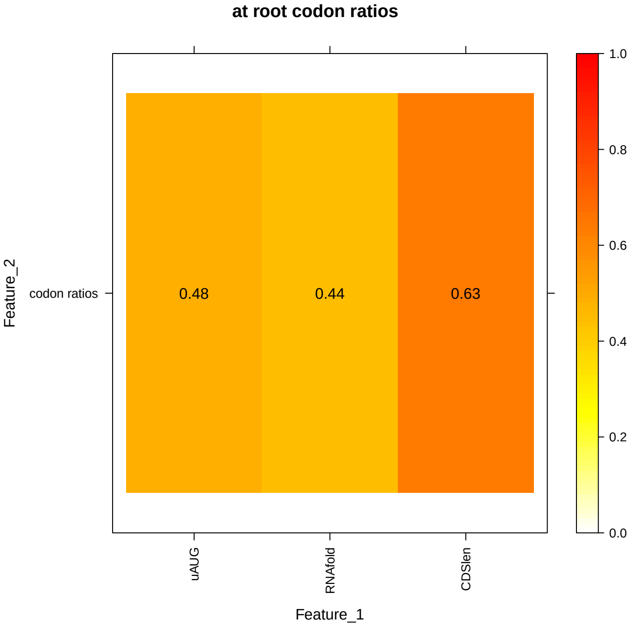

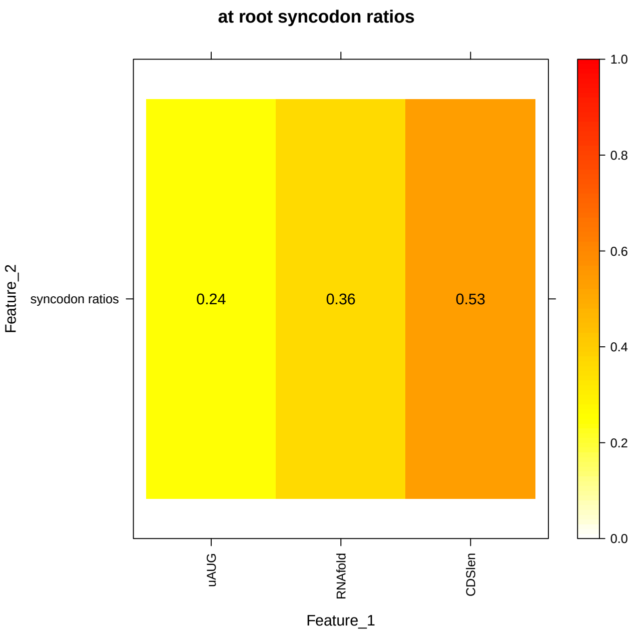


```
## at.shoot
## R2 of aa freqs: 0.1620685
## R2 of uAUG: 0.09685659
## R2 of aa freqs + uAUG: 0.2268717
## R2 of RNAfold: 0.1342398
## R2 of aa freqs + RNAfold: 0.2639618
## R2 of CDSlen: 0.06873281
## R2 of aa freqs + CDSlen: 0.2107153
##
## R2 of codon freqs: 0.231544
## R2 of uAUG: 0.09685659
## R2 of codon freqs + uAUG: 0.2836446
## R2 of RNAfold: 0.1342398
## R2 of codon freqs + RNAfold: 0.3040842
## R2 of CDSlen: 0.06873281
## R2 of codon freqs + CDSlen: 0.2545832
##
## R2 of syncodon ratios: 0.1006659
## R2 of uAUG: 0.09685659
## R2 of syncodon ratios + uAUG: 0.173407
## R2 of RNAfold: 0.1342398
## R2 of syncodon ratios + RNAfold: 0.1951689
## R2 of CDSlen: 0.06873281
## R2 of syncodon ratios + CDSlen: 0.1304149
```

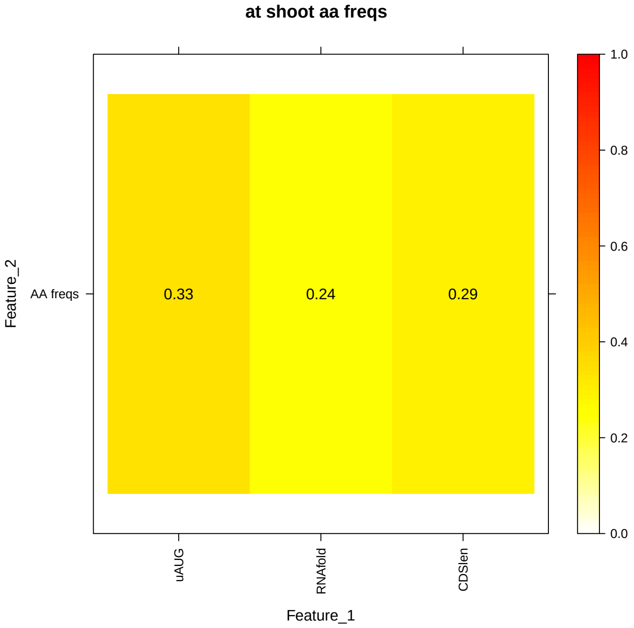


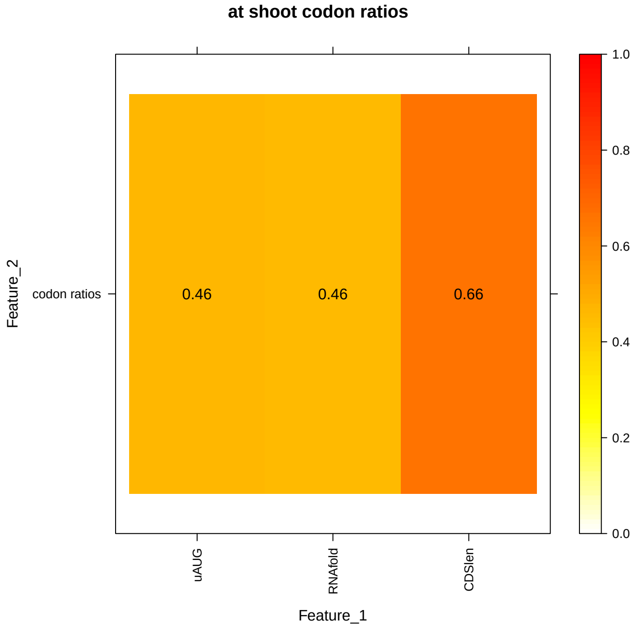


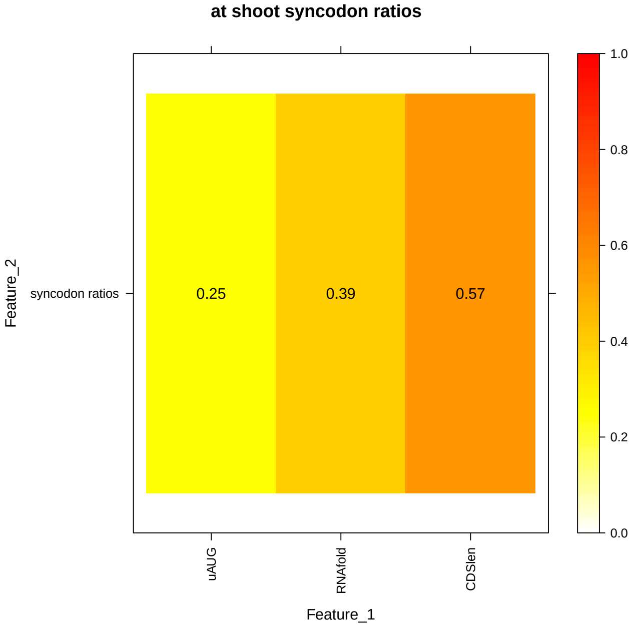


```
## mm.nih3t3
## R2 of aa freqs: 0.1037734
## R2 of uAUG: 0.08337952
## R2 of aa freqs + uAUG: 0.1652441
## R2 of RNAfold: 0.1758413
## R2 of aa freqs + RNAfold: 0.2543673
## R2 of CDSlen: 0.1060529
## R2 of aa freqs + CDSlen: 0.1759301
##
## R2 of codon freqs: 0.1457886
## R2 of uAUG: 0.08337952
## R2 of codon freqs + uAUG: 0.2027361
## R2 of RNAfold: 0.1758413
## R2 of codon freqs + RNAfold: 0.2849187
```

```
## R2 of CDSlen: 0.1060529
## R2 of codon freqs + CDSlen: 0.2101996
##
## R2 of syncodon ratios: 0.05288419
## R2 of uAUG: 0.08337952
## R2 of syncodon ratios + uAUG: 0.1258685
## R2 of RNAfold: 0.1758413
## R2 of syncodon ratios + RNAfold: 0.2114868
## R2 of CDSlen: 0.1060529
## R2 of syncodon ratios + CDSlen: 0.1389477
```

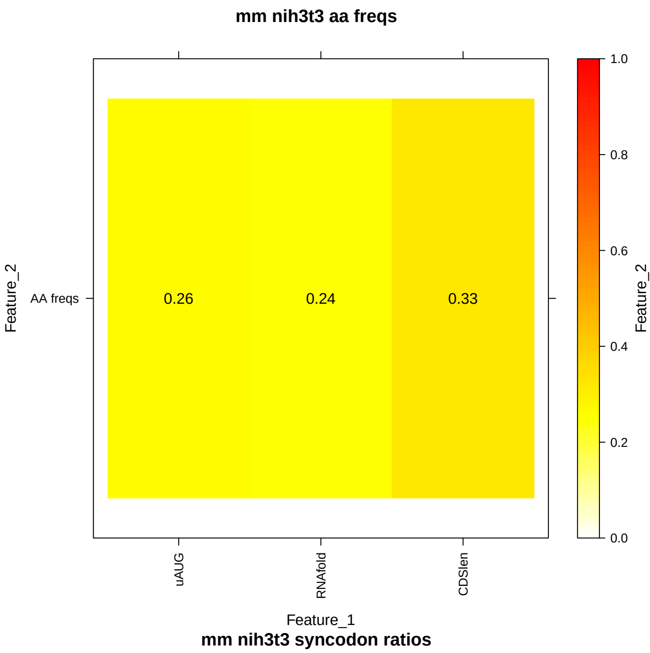


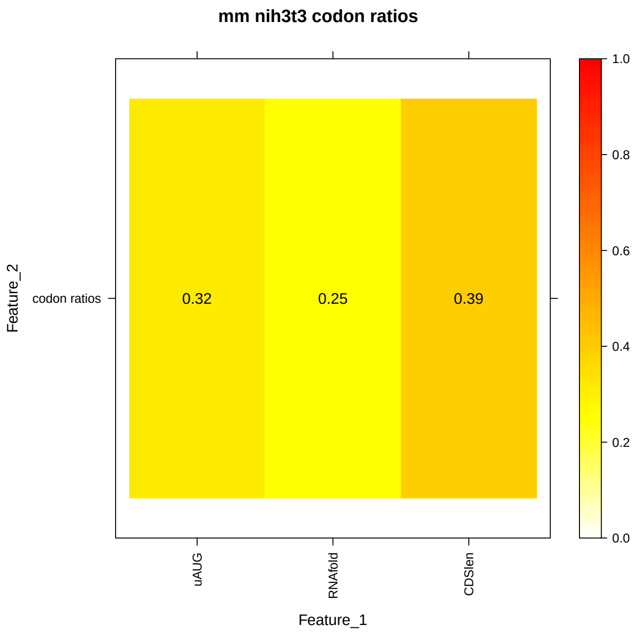


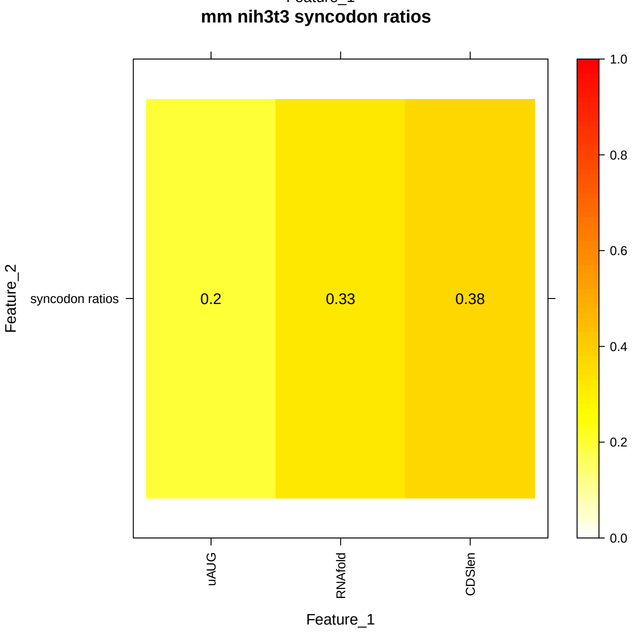


```
## mm.liver
## R2 of aa freqs: 0.1058606
## R2 of uAUG: 0.0727428
## R2 of aa freqs + uAUG: 0.1609637
```

```
## R2 of RNAfold: 0.184825
## R2 of aa freqs + RNAfold: 0.2689322
## R2 of CDSlen: 0.1632853
## R2 of aa freqs + CDSlen: 0.2241327
##
## R2 of codon freqs: 0.1795914
## R2 of uAUG: 0.0727428
## R2 of codon freqs + uAUG: 0.2314827
## R2 of RNAfold: 0.184825
## R2 of codon freqs + RNAfold: 0.3275042
## R2 of CDSlen: 0.1632853
## R2 of codon freqs + CDSlen: 0.2822876
##
## R2 of syncodon ratios: 0.1001104
## R2 of uAUG: 0.0727428
## R2 of syncodon ratios + uAUG: 0.1636221
## R2 of RNAfold: 0.184825
## R2 of syncodon ratios + RNAfold: 0.2602259
## R2 of CDSlen: 0.1632853
## R2 of syncodon ratios + CDSlen: 0.2268437
```

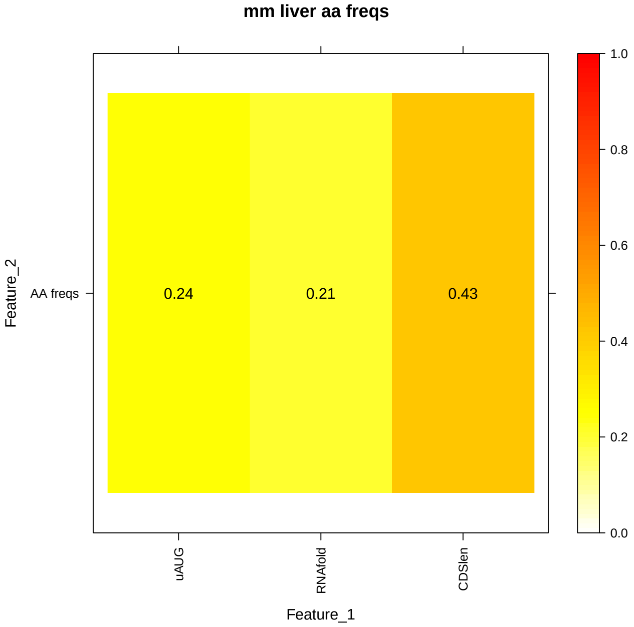


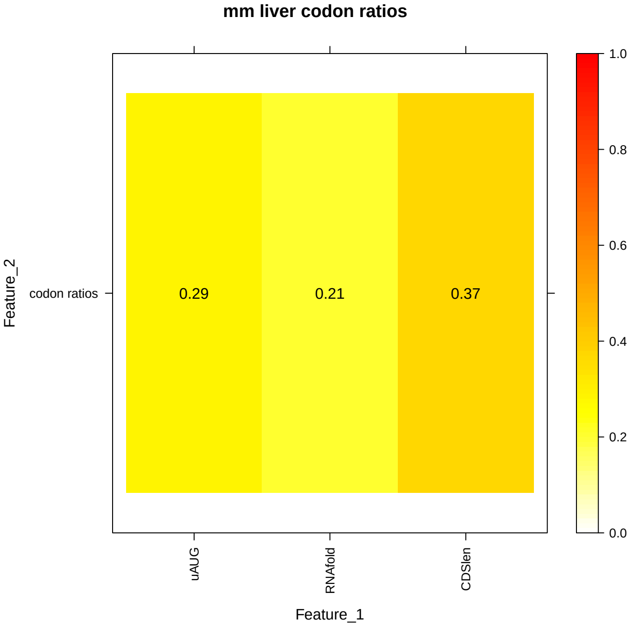

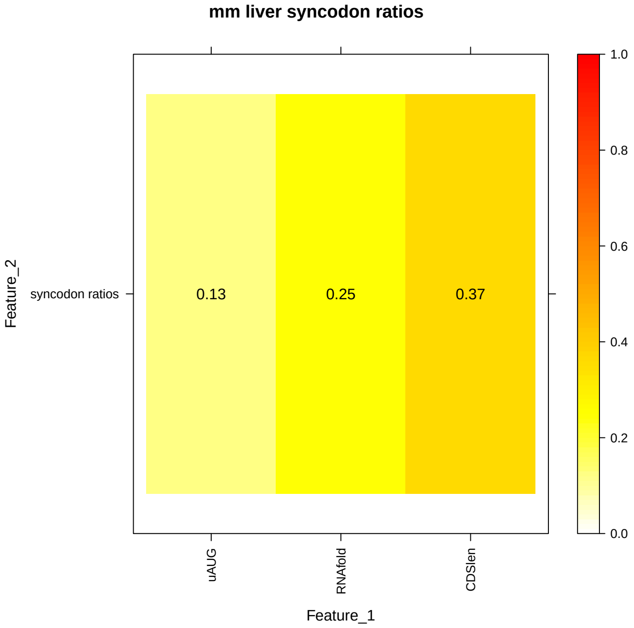


```
## mm.kidney
## R2 of aa freqs: 0.1357304
## R2 of uAUG: 0.08544309
## R2 of aa freqs + uAUG: 0.1942969
## R2 of RNAfold: 0.2129005
## R2 of aa freqs + RNAfold: 0.3112252
## R2 of CDSlen: 0.1536686
## R2 of aa freqs + CDSlen: 0.2220784
##
## R2 of codon freqs: 0.1659077
## R2 of uAUG: 0.08544309
## R2 of codon freqs + uAUG: 0.2203029
## R2 of RNAfold: 0.2129005
## R2 of codon freqs + RNAfold: 0.3313333
## R2 of CDSlen: 0.1536686
## R2 of codon freqs + CDSlen: 0.2464779
##
## R2 of syncodon ratios: 0.0500301
## R2 of uAUG: 0.08544309
## R2 of syncodon ratios + uAUG: 0.1243589
## R2 of RNAfold: 0.2129005
## R2 of syncodon ratios + RNAfold: 0.2472377
## R2 of CDSlen: 0.1536686
## R2 of syncodon ratios + CDSlen: 0.1837945
```

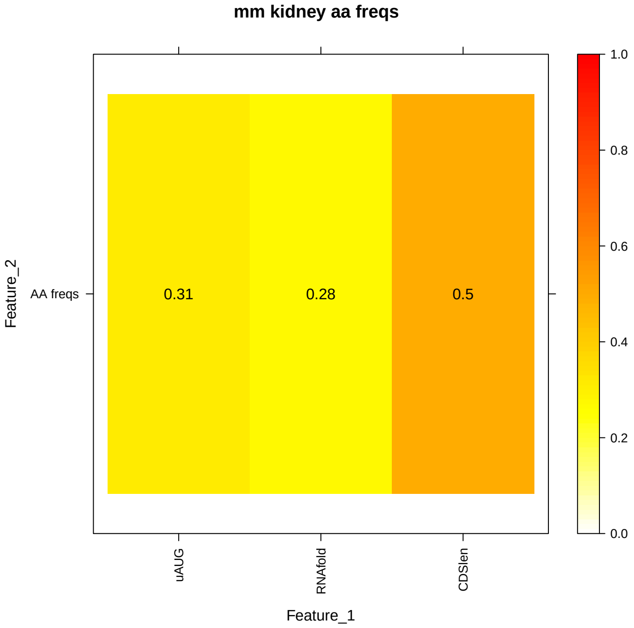


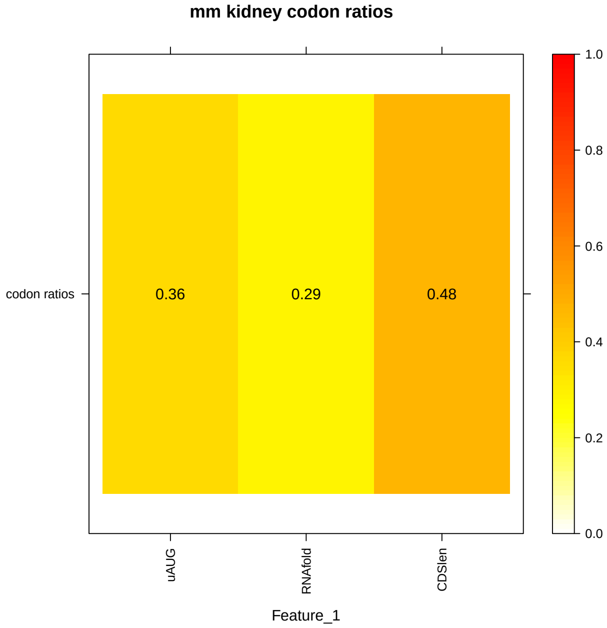


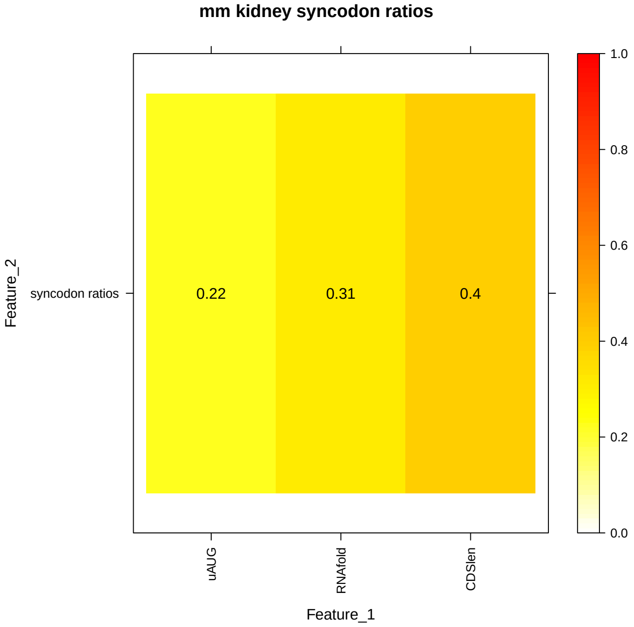


```
## hs.hela2
## R2 of aa freqs: 0.1492081
## R2 of uAUG: 0.02158616
## R2 of aa freqs + uAUG: 0.161095
## R2 of RNAfold: 0.07933165
## R2 of aa freqs + RNAfold: 0.2073126
## R2 of CDSlen: 0.2725774
## R2 of aa freqs + CDSlen: 0.3427748
##
## R2 of codon freqs: 0.1880345
## R2 of uAUG: 0.02158616
## R2 of codon freqs + uAUG: 0.1981765
## R2 of RNAfold: 0.07933165
## R2 of codon freqs + RNAfold: 0.2420245
```

```
## R2 of CDSlen: 0.2725774
## R2 of codon freqs + CDSlen: 0.3690907
##
## R2 of syncodon ratios: 0.0592387
## R2 of uAUG: 0.02158616
## R2 of syncodon ratios + uAUG: 0.07594777
## R2 of RNAfold: 0.07933165
## R2 of syncodon ratios + RNAfold: 0.1329607
## R2 of CDSlen: 0.2725774
## R2 of syncodon ratios + CDSlen: 0.3142431
```

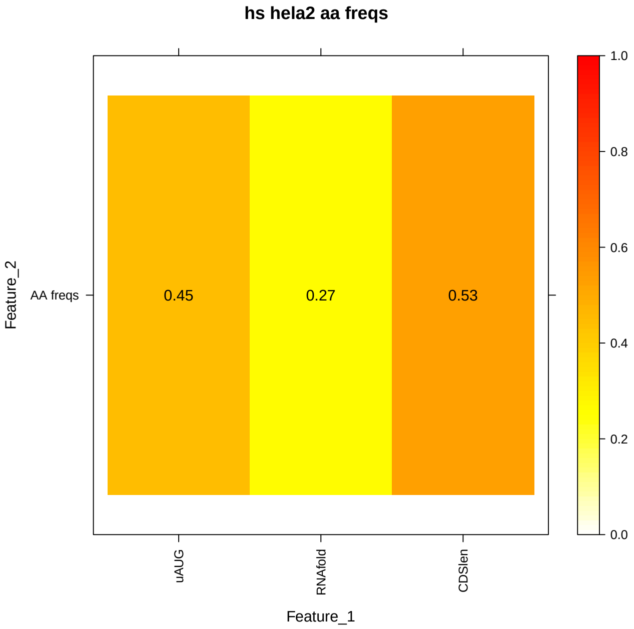


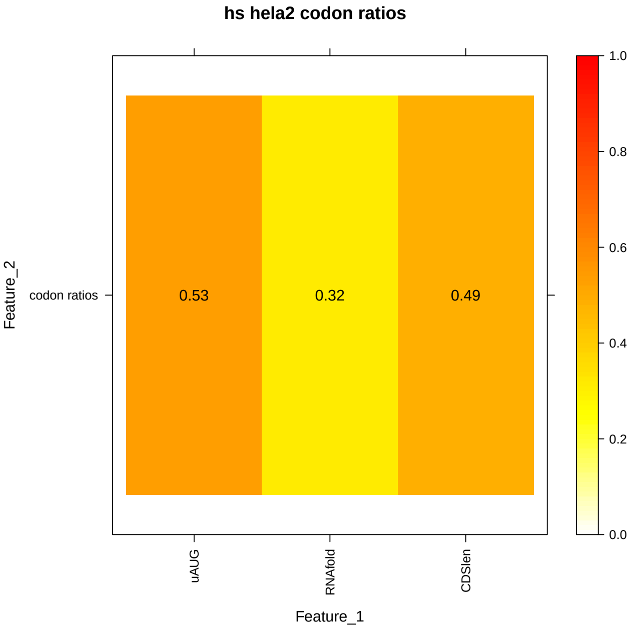


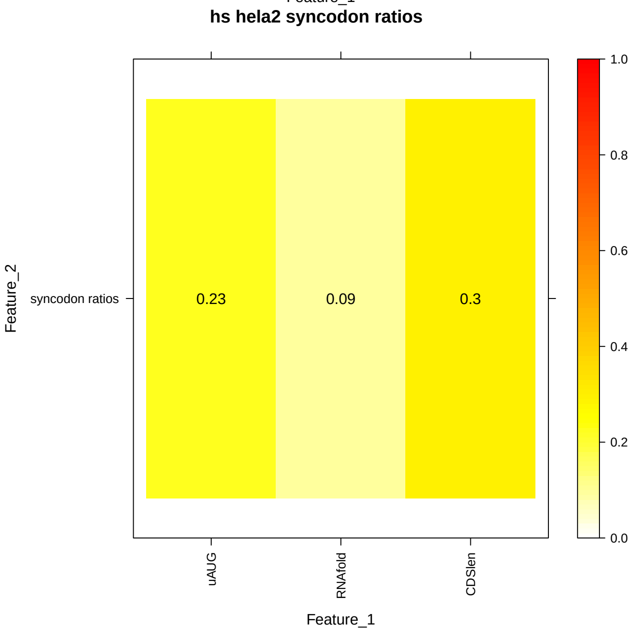

### Figure 17. Collinearity analysis of three features: uAUGs, 5'RNA fold, and CDS length, with AA usage and synonymous codon usage

```
## A function to calculate collinearity between aa usage (or
## synonymous codon usage) and another feature
calc_aa_syncodon_collinearity <- function(features_list, aa_features,
    codon_features, y) {
    p <- length(features_list)
    feature_names <- names(features_list)

    uni_feature_R2s <- sapply(1:p, FUN = function(i) {
        tmp_features_all <- cbind(y, features_list[[i]])
        colnames(tmp_features_all)[1] <- "y"
        ## remove duplicated column names
        tmp_features_all <- tmp_features_all[, !duplicated(colnames(tmp_features_all))]
        tmp_features_all <- as.data.frame(tmp_features_all)
        ## model
        tmp_features_model <- lm(y ~ ., data = tmp_features_all)
        summary(tmp_features_model)$r.squared
    })

    aa_features_R2 <- summary(lm(y ~ as.matrix(aa_features)))$r.squared
    codon_features_R2 <- summary(lm(y ~ as.matrix(codon_features)))$r.squared
    syncodon_features_R2 <- codon_features_R2 - aa_features_R2

    bi_feature_R2s <- sapply(1:p, FUN = function(i) {
        tmp_features_all <- cbind(y, features_list[[i]], aa_features)
        colnames(tmp_features_all)[1] <- "y"
        ## remove duplicated column names
        tmp_features_all <- tmp_features_all[, !duplicated(colnames(tmp_features_all))]
        tmp_features_all <- as.data.frame(tmp_features_all)
        ## model
        tmp_features_model <- lm(y ~ ., data = tmp_features_all)
        aa_bi_feature_R2 <- summary(tmp_features_model)$r.squared

        tmp_features_all <- cbind(y, features_list[[i]], codon_features)
        colnames(tmp_features_all)[1] <- "y"
        ## remove duplicated column names
        tmp_features_all <- tmp_features_all[, !duplicated(colnames(tmp_features_all))]
        tmp_features_all <- as.data.frame(tmp_features_all)
        ## model
        tmp_features_model <- lm(y ~ ., data = tmp_features_all)
        codon_bi_feature_R2 <- summary(tmp_features_model)$r.squared

        return(c(aa_bi_feature_R2, codon_bi_feature_R2 - aa_bi_feature_R2))
    })

    aa_result <- rep(NA, p)
    names(aa_result) <- feature_names

    syncodon_result <- rep(NA, p)
    names(syncodon_result) <- feature_names
```

```
    # the order of bi_feature_R2s should correspond to
    # feature_names
    for (i in 1:p) {
        aa_redundancy <- (uni_feature_R2s[i] + aa_features_R2) -
            bi_feature_R2s[1, i]
        aa_result[i] <- max(round(aa_redundancy/uni_feature_R2s[i] *
            100, 3), round(aa_redundancy/aa_features_R2 * 100,
            3))

        syncodon_redundancy <- (uni_feature_R2s[i] + syncodon_features_R2) -
            bi_feature_R2s[2, i]
        syncodon_result[i] <- max(round(syncodon_redundancy/uni_feature_R2s[i] *
            100, 3), round(syncodon_redundancy/syncodon_features_R2 *
            100, 3))
    }
    return(list(aa_result, syncodon_result))
}

for (species in names(species_tissue_list)) {
    for (species_tissue in species_tissue_list[[species]]) {
        if (species_tissue != "") {
            tissue_dot <- paste0(".", species_tissue)
            tissue_us <- paste0("_", species_tissue)
        } else {
            tissue_dot <- ""
            tissue_us <- ""
        }

        features_list <- list(uAUG = get(paste0(species, tissue_dot,
            ".uAUG_counts.div")), RNAfold = get(paste0(species,
            tissue_dot, ".fold_energy_features_selected")), CDSlen = get(paste0(species,
            tissue_dot, ".log10.CDS_lens.div")))
        aa_features <- get(paste0(species, tissue_dot, ".aa_features"))[[1]]
        codon_features <- get(paste0(species, tissue_dot, ".codon_features"))[[1]]
        tmp <- calc_aa_syncodon_collinearity(features_list, aa_features,
            codon_features, get(paste0(species, tissue_dot, ".log10.TR")))
        aa_result <- tmp[[1]]
        syncodon_result <- tmp[[2]]
        aa_result[aa_result > 100] <- 100  # truncate the value at 100
        syncodon_result[syncodon_result > 100] <- 100  # truncate the value at 100

        aa_result <- aa_result/100
        names(aa_result) <- names(features_list)
        syncodon_result <- syncodon_result/100
        names(syncodon_result) <- names(features_list)

        cat(paste0(species, tissue_dot, "\ncollinearity with AA usage: \n"))
        for (i in 1:length(features_list)) {
            cat(names(features_list)[i])
            cat("\n")
            cat(aa_result[i])
            cat("\n")
        }
```

```r
        cat("\n")

        cat(paste0(species, tissue_dot, "\ncollinearity with synonymous codon usage: \n"))
        for (i in 1:length(features_list)) {
            cat(names(features_list)[i])
            cat("\n")
            cat(syncodon_result[i])
            cat("\n")
        }
        cat("\n")
    }
}
```

```
## sc
## collinearity with AA usage:
## uAUG
## 0.65777
## RNAfold
## 0.54309
## CDSlen
## 0.47887
##
## sc
## collinearity with synonymous codon usage:
## uAUG
## 1
## RNAfold
## 1
## CDSlen
## 1
##
## sp
## collinearity with AA usage:
## uAUG
## 0.37412
## RNAfold
## 0.61339
## CDSlen
## 0.47913
##
## sp
## collinearity with synonymous codon usage:
## uAUG
## 1
## RNAfold
## 1
## CDSlen
## 1
##
## sp.alt
## collinearity with AA usage:
## uAUG
## 0.45983
## RNAfold
```

```
## 0.93162
## CDSlen
## 0.523
##
## sp.alt
## collinearity with synonymous codon usage:
## uAUG
## 1
## RNAfold
## 1
## CDSlen
## 1
##
## at.leaf
## collinearity with AA usage:
## uAUG
## 0.26331
## RNAfold
## 0.35424
## CDSlen
## 0.26378
##
## at.leaf
## collinearity with synonymous codon usage:
## uAUG
## 1
## RNAfold
## 1
## CDSlen
## 1
##
## at.root
## collinearity with AA usage:
## uAUG
## 0.35684
## RNAfold
## 0.27144
## CDSlen
## 0.41684
##
## at.root
## collinearity with synonymous codon usage:
## uAUG
## 1
## RNAfold
## 1
## CDSlen
## 1
##
## at.shoot
## collinearity with AA usage:
## uAUG
## 0.33212
## RNAfold
```

```
## 0.25109
## CDSlen
## 0.42218
##
## at.shoot
## collinearity with synonymous codon usage:
## uAUG
## 1
## RNAfold
## 1
## CDSlen
## 1
##
## mm.nih3t3
## collinearity with AA usage:
## uAUG
## 0.25093
## RNAfold
## 0.26051
## CDSlen
## 0.44254
##
## mm.nih3t3
## collinearity with synonymous codon usage:
## uAUG
## 1
## RNAfold
## 1
## CDSlen
## 1
##
## mm.liver
## collinearity with AA usage:
## uAUG
## 0.2366
## RNAfold
## 0.20597
## CDSlen
## 0.48259
##
## mm.liver
## collinearity with synonymous codon usage:
## uAUG
## 1
## RNAfold
## 1
## CDSlen
## 1
##
## mm.kidney
## collinearity with AA usage:
## uAUG
## 0.30028
## RNAfold
```

```
## 0.28675
## CDSlen
## 0.55993
##
## mm.kidney
## collinearity with synonymous codon usage:
## uAUG
## 1
## RNAfold
## 1
## CDSlen
## 1
##
## hs.hela2
## collinearity with AA usage:
## uAUG
## 0.4724
## RNAfold
## 0.32229
## CDSlen
## 0.58432
##
## hs.hela2
## collinearity with synonymous codon usage:
## uAUG
## 1
## RNAfold
## 1
## CDSlen
## 1
```

### Figure 18. Analysis of aa freqs / codon freqs / synonymous codon ratios vs. tRNA abundance for sc

```
for (species in "sc") {
    for (i in 1:length(species_tissue_list[[species]])) {
        species_tissue <- species_tissue_list[[species]][i]
        if (species_tissue != "") {
            tissue_dot <- paste0(".", species_tissue)
            tissue_us <- paste0("_", species_tissue)
        } else {
            tissue_dot <- ""
            tissue_us <- ""
        }

        aa_freqs <- readRDS(paste0("processed_data_w_polyA/aa_features/",
            species, tissue_dot, ".aa_freqs.rds"))
        codon_freqs <- readRDS(paste0("processed_data_w_polyA/codon_features/",
            species, tissue_dot, ".codon_freqs.rds"))
        syncodon_ratios <- readRDS(paste0("processed_data_w_polyA/syncodon_features/",
            species, tissue_dot, ".syncodon_ratios.rds"))
```

```
genes <- get(paste0(species, tissue_dot, ".genes"))

TR_top_genes <- get(paste0(species, tissue_dot, ".TR_top_genes"))
TR_bottom_genes <- get(paste0(species, tissue_dot, ".TR_bottom_genes"))

keep_idx <- which(apply(syncodon_ratios, 1, FUN = function(x) !(any(x ==
    Inf | is.na(x)))))

aa_freqs <- aa_freqs[keep_idx, ]
codon_freqs <- codon_freqs[keep_idx, ]
syncodon_ratios <- syncodon_ratios[keep_idx, ]
genes <- genes[keep_idx]
TR_top_genes <- TR_top_genes[TR_top_genes %in% genes]
TR_bottom_genes <- TR_bottom_genes[TR_bottom_genes %in%
    genes]

aa_tRNA_abun <- readRDS(paste0("processed_data_w_polyA/tRNA_abun/",
    species, tissue_dot, ".aa_tRNA_abun.rds"))
codon_tRNA_abun <- readRDS(paste0("processed_data_w_polyA/tRNA_abun/",
    species, tissue_dot, ".codon_tRNA_abun.rds"))

TR_top_avg_aa_freqs <- colMeans(aa_freqs[TR_top_genes,
    ])
TR_bottom_avg_aa_freqs <- colMeans(aa_freqs[TR_bottom_genes,
    ])

TR_top_avg_codon_freqs <- colMeans(codon_freqs[TR_top_genes,
    ])
TR_bottom_avg_codon_freqs <- colMeans(codon_freqs[TR_bottom_genes,
    ])

TR_top_avg_syncodon_ratios <- colMeans(syncodon_ratios[TR_top_genes,
    ])
TR_bottom_avg_syncodon_ratios <- colMeans(syncodon_ratios[TR_bottom_genes,
    ])

# output the values

## R2s of tRNA abun ~ AA freqs (TR top vs. bottom)
cat(paste0(species, tissue_dot, "\n"))
cat("Top TR AA freqs. vs tRNAs abundaces: ")
cat(cor(TR_top_avg_aa_freqs, aa_tRNA_abun)^2)
cat("\n")
cat("Bottom TR AA freqs. vs tRNAs abundaces: ")
cat(cor(TR_bottom_avg_aa_freqs, aa_tRNA_abun)^2)
cat("\n")
cat("Test if TR top > TR bottom:\n")
print(cocor.dep.groups.overlap(r.jk = cor(aa_tRNA_abun,
    TR_top_avg_aa_freqs), r.jh = cor(aa_tRNA_abun, TR_bottom_avg_aa_freqs),
    r.kh = cor(TR_top_avg_aa_freqs, TR_bottom_avg_aa_freqs),
    n = 20, alternative = "greater", var.labels = c("tRNA_abun",
        "TR_top_aa_freq", "TR_bottom_aa_freq")))
```

```
## R2s of tRNA abun ~ codon freqs (TR top vs. bottom)
cat(paste0(species, tissue_dot, "\n"))
cat("Top TR codon freqs. vs tRNAs abundaces: ")
cat(cor(TR_top_avg_codon_freqs, codon_tRNA_abun)^2)
cat("\n")
cat("Bottom TR codon freqs. vs tRNAs abundaces: ")
cat(cor(TR_bottom_avg_codon_freqs, codon_tRNA_abun)^2)
cat("\n")
cat("Test if TR top > TR bottom:\n")
print(cocor.dep.groups.overlap(r.jk = cor(codon_tRNA_abun,
    TR_top_avg_codon_freqs), r.jh = cor(codon_tRNA_abun,
    TR_bottom_avg_codon_freqs), r.kh = cor(TR_top_avg_codon_freqs,
    TR_bottom_avg_codon_freqs), n = 20, alternative = "greater",
    var.labels = c("tRNA_abun", "TR_top_aa_freq", "TR_bottom_aa_freq")))

## R2s of tRNA abun ~ syncodon ratios (TR top vs. bottom)
cat(paste0(species, tissue_dot, "\n"))
cat("Top TR syncodon ratios. vs tRNAs abundaces: ")
cat(cor(TR_top_avg_syncodon_ratios, codon_tRNA_abun)^2)
cat("\n")
cat("Bottom TR syncodon ratios. vs tRNAs abundaces: ")
cat(cor(TR_bottom_avg_syncodon_ratios, codon_tRNA_abun)^2)
cat("\n")
cat("Test if TR top > TR bottom:\n")
print(cocor.dep.groups.overlap(r.jk = cor(codon_tRNA_abun,
    TR_top_avg_syncodon_ratios), r.jh = cor(codon_tRNA_abun,
    TR_bottom_avg_syncodon_ratios), r.kh = cor(TR_top_avg_syncodon_ratios,
    TR_bottom_avg_syncodon_ratios), n = 20, alternative = "greater",
    var.labels = c("tRNA_abun", "TR_top_aa_freq", "TR_bottom_aa_freq")))

# scatterplots of tRNA abundance vs. TR top/bottom ratios
TR_top_vs_bottom_avg_aa_freqs_ratios <- TR_top_avg_aa_freqs/TR_bottom_avg_aa_freqs
TR_top_vs_bottom_avg_codon_freqs_ratios <- TR_top_avg_codon_freqs/TR_bottom_avg_codon_freqs
TR_top_vs_bottom_avg_syncodon_ratios_ratios <- TR_top_avg_syncodon_ratios/TR_bottom_avg_syncodon

plot(x = TR_top_vs_bottom_avg_aa_freqs_ratios, y = aa_tRNA_abun,
    xlab = "TR top/bottom AA freq. ratios", ylab = "tRNA abundance")
title(paste("cor =", round(cor(x = TR_top_vs_bottom_avg_aa_freqs_ratios,
    y = aa_tRNA_abun), 2), "p =", round(cor.test(TR_top_vs_bottom_avg_aa_freqs_ratios,
    aa_tRNA_abun, alternative = "greater")$p.value *
    3, 3)))
abline(v = 1, lty = 2)

plot(x = TR_top_vs_bottom_avg_codon_freqs_ratios, y = codon_tRNA_abun,
    xlab = "TR top/bottom codon freq. ratios", ylab = "tRNA abundance")
title(paste("cor =", round(cor(x = TR_top_vs_bottom_avg_codon_freqs_ratios,
    y = codon_tRNA_abun), 2), "p =", round(cor.test(TR_top_vs_bottom_avg_codon_freqs_ratios,
    codon_tRNA_abun, alternative = "greater")$p.value *
    3, 3)))
abline(v = 1, lty = 2)

plot(x = TR_top_vs_bottom_avg_syncodon_ratios_ratios,
    y = codon_tRNA_abun, xlab = "TR top/bottom syncodon ratio ratios",
```

```
            ylab = "tRNA abundance")
        title(paste("cor =", round(cor(x = TR_top_vs_bottom_avg_syncodon_ratios_ratios,
            y = codon_tRNA_abun), 2), "p =", round(cor.test(TR_top_vs_bottom_avg_syncodon_ratios_ratios
            codon_tRNA_abun, alternative = "greater")$p.value *
            3, 3)))
        abline(v = 1, lty = 2)
    }
}
```

```
## sc
## Top TR AA freqs. vs tRNAs abundaces: 0.6619552
## Bottom TR AA freqs. vs tRNAs abundaces: 0.3958626
## Test if TR top > TR bottom:
##
##   Results of a comparison of two overlapping correlations based on dependent groups
##
## Comparison between r.jk (tRNA_abun, TR_top_aa_freq) = 0.8136 and r.jh (tRNA_abun, TR_bottom_aa_freq)
## Difference: r.jk - r.jh = 0.1844
## Related correlation: r.kh = 0.7893
## Data: j = tRNA_abun, k = TR_top_aa_freq, h = TR_bottom_aa_freq
## Group size: n = 20
## Null hypothesis: r.jk is equal to r.jh
## Alternative hypothesis: r.jk is greater than r.jh (one-sided)
## Alpha: 0.05
##
## pearson1898: Pearson and Filon's z (1898)
##   z = 1.7504, p-value = 0.0400
##   Null hypothesis rejected
##
## hotelling1940: Hotelling's t (1940)
##   t = 2.0160, df = 17, p-value = 0.0299
##   Null hypothesis rejected
##
## williams1959: Williams' t (1959)
##   t = 1.9990, df = 17, p-value = 0.0309
##   Null hypothesis rejected
##
## olkin1967: Olkin's z (1967)
##   z = 1.7504, p-value = 0.0400
##   Null hypothesis rejected
##
## dunn1969: Dunn and Clark's z (1969)
##   z = 1.9049, p-value = 0.0284
##   Null hypothesis rejected
##
## hendrickson1970: Hendrickson, Stanley, and Hills' (1970) modification of Williams' t (1959)
##   t = 2.0160, df = 17, p-value = 0.0299
##   Null hypothesis rejected
##
## steiger1980: Steiger's (1980) modification of Dunn and Clark's z (1969) using average correlations
##   z = 1.8581, p-value = 0.0316
##   Null hypothesis rejected
##
## meng1992: Meng, Rosenthal, and Rubin's z (1992)
```

```
##   z = 1.8457, p-value = 0.0325
##   Null hypothesis rejected
##   95% confidence interval for r.jk - r.jh: -0.0246 0.8197
##   Null hypothesis retained (Lower boundary <= 0)
##
## hittner2003: Hittner, May, and Silver's (2003) modification of Dunn and Clark's z (1969) using a bac
##   z = 1.8298, p-value = 0.0336
##   Null hypothesis rejected
##
## zou2007: Zou's (2007) confidence interval
##   95% confidence interval for r.jk - r.jh: -0.0074 0.4975
##   Null hypothesis retained (Lower boundary <= 0)
##
## sc
## Top TR codon freqs. vs tRNAs abundaces: 0.6569623
## Bottom TR codon freqs. vs tRNAs abundaces: 0.1949541
## Test if TR top > TR bottom:
##
##   Results of a comparison of two overlapping correlations based on dependent groups
##
## Comparison between r.jk (tRNA_abun, TR_top_aa_freq) = 0.8105 and r.jh (tRNA_abun, TR_bottom_aa_freq)
## Difference: r.jk - r.jh = 0.369
## Related correlation: r.kh = 0.6046
## Data: j = tRNA_abun, k = TR_top_aa_freq, h = TR_bottom_aa_freq
## Group size: n = 20
## Null hypothesis: r.jk is equal to r.jh
## Alternative hypothesis: r.jk is greater than r.jh (one-sided)
## Alpha: 0.05
##
## pearson1898: Pearson and Filon's z (1898)
##   z = 2.3132, p-value = 0.0104
##   Null hypothesis rejected
##
## hotelling1940: Hotelling's t (1940)
##   t = 2.9368, df = 17, p-value = 0.0046
##   Null hypothesis rejected
##
## williams1959: Williams' t (1959)
##   t = 2.8656, df = 17, p-value = 0.0054
##   Null hypothesis rejected
##
## olkin1967: Olkin's z (1967)
##   z = 2.3132, p-value = 0.0104
##   Null hypothesis rejected
##
## dunn1969: Dunn and Clark's z (1969)
##   z = 2.6087, p-value = 0.0045
##   Null hypothesis rejected
##
## hendrickson1970: Hendrickson, Stanley, and Hills' (1970) modification of Williams' t (1959)
##   t = 2.9364, df = 17, p-value = 0.0046
##   Null hypothesis rejected
##
## steiger1980: Steiger's (1980) modification of Dunn and Clark's z (1969) using average correlations
```

```
##   z = 2.5329, p-value = 0.0057
##   Null hypothesis rejected
##
## meng1992: Meng, Rosenthal, and Rubin's z (1992)
##   z = 2.4886, p-value = 0.0064
##   Null hypothesis rejected
##   95% confidence interval for r.jk - r.jh: 0.1390 1.1699
##   Null hypothesis rejected (Lower boundary > 0)
##
## hittner2003: Hittner, May, and Silver's (2003) modification of Dunn and Clark's z (1969) using a bac
##   z = 2.4681, p-value = 0.0068
##   Null hypothesis rejected
##
## zou2007: Zou's (2007) confidence interval
##   95% confidence interval for r.jk - r.jh: 0.0875 0.7721
##   Null hypothesis rejected (Lower boundary > 0)
##
## sc
## Top TR syncodon ratios. vs tRNAs abundaces: 0.4434027
## Bottom TR syncodon ratios. vs tRNAs abundaces: 0.1222008
## Test if TR top > TR bottom:
##
##   Results of a comparison of two overlapping correlations based on dependent groups
##
## Comparison between r.jk (tRNA_abun, TR_top_aa_freq) = 0.6659 and r.jh (tRNA_abun, TR_bottom_aa_freq)
## Difference: r.jk - r.jh = 0.3163
## Related correlation: r.kh = 0.7589
## Data: j = tRNA_abun, k = TR_top_aa_freq, h = TR_bottom_aa_freq
## Group size: n = 20
## Null hypothesis: r.jk is equal to r.jh
## Alternative hypothesis: r.jk is greater than r.jh (one-sided)
## Alpha: 0.05
##
## pearson1898: Pearson and Filon's z (1898)
##   z = 2.2710, p-value = 0.0116
##   Null hypothesis rejected
##
## hotelling1940: Hotelling's t (1940)
##   t = 2.6581, df = 17, p-value = 0.0083
##   Null hypothesis rejected
##
## williams1959: Williams' t (1959)
##   t = 2.6480, df = 17, p-value = 0.0085
##   Null hypothesis rejected
##
## olkin1967: Olkin's z (1967)
##   z = 2.2710, p-value = 0.0116
##   Null hypothesis rejected
##
## dunn1969: Dunn and Clark's z (1969)
##   z = 2.3672, p-value = 0.0090
##   Null hypothesis rejected
##
## hendrickson1970: Hendrickson, Stanley, and Hills' (1970) modification of Williams' t (1959)
```

```
##   t = 2.6580, df = 17, p-value = 0.0083
##   Null hypothesis rejected
##
## steiger1980: Steiger's (1980) modification of Dunn and Clark's z (1969) using average correlations
##   z = 2.2910, p-value = 0.0110
##   Null hypothesis rejected
##
## meng1992: Meng, Rosenthal, and Rubin's z (1992)
##   z = 2.2588, p-value = 0.0119
##   Null hypothesis rejected
##   95% confidence interval for r.jk - r.jh: 0.0580 0.8187
##   Null hypothesis rejected (Lower boundary > 0)
##
## hittner2003: Hittner, May, and Silver's (2003) modification of Dunn and Clark's z (1969) using a bac
##   z = 2.2672, p-value = 0.0117
##   Null hypothesis rejected
##
## zou2007: Zou's (2007) confidence interval
##   95% confidence interval for r.jk - r.jh: 0.0543 0.6677
##   Null hypothesis rejected (Lower boundary > 0)
```

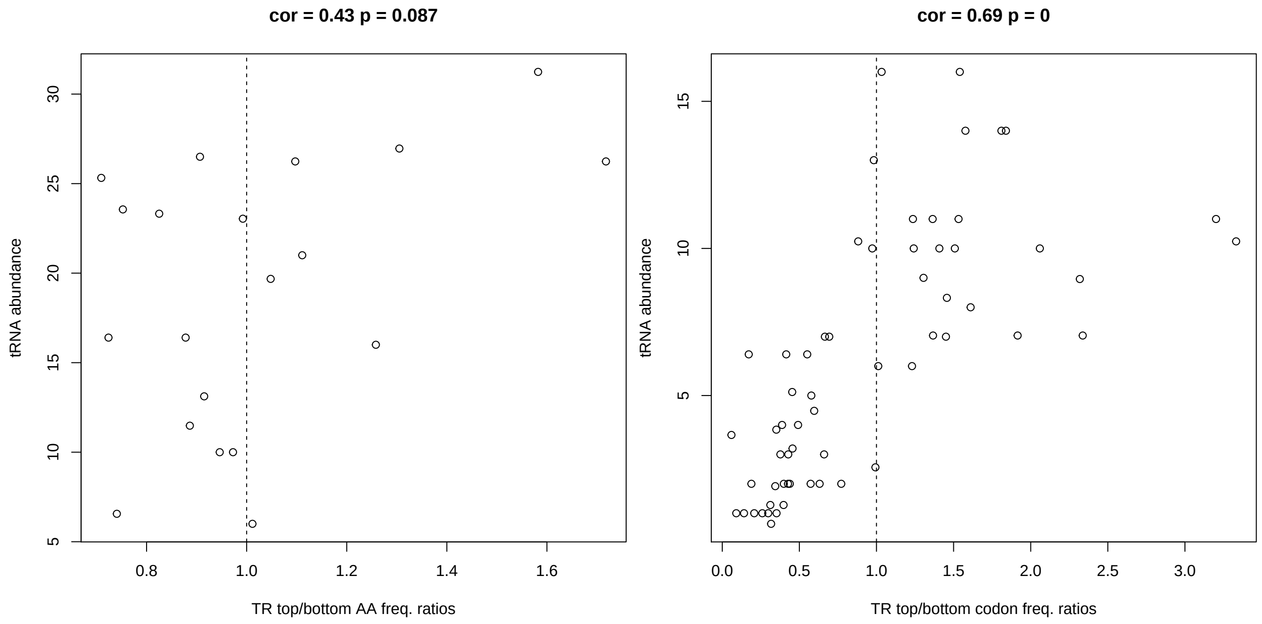

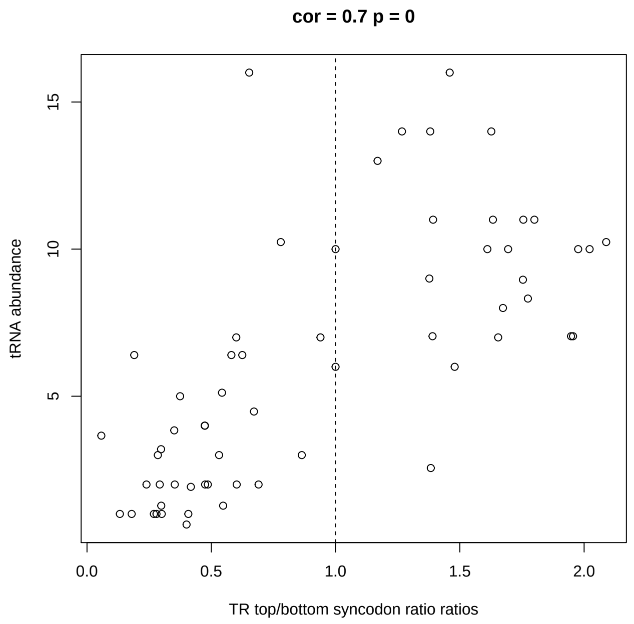


#### Supplementary file 2. Output the processed information to an Excel file, one tab per species & tissue

```
# columns Gene 5' UTR seq CDS seq log10 mRNA (RPKM) log10 TR
# (RPKM) 5' motif score 5' UTR RNA fold engy 5'UTR #uAUGs
# log10 CDS length codon freq.  AA freq.  syn.codon
# preference poly A length log10 5' UTR length

for (species in names(species_tissue_list)) {
    for (species_tissue in species_tissue_list[[species]]) {
        if (species_tissue != "") {
            tissue_dot <- paste0(".", species_tissue)
            tissue_us <- paste0("_", species_tissue)
        } else {
            tissue_dot <- ""
            tissue_us <- ""
        }
        supp2 <- data.frame(Gene = get(paste0(species, tissue_dot,
            ".genes")))
        supp2$"5' UTR seq" <- get(paste0(species, tissue_dot,
            ".UTR_seqs"))
        supp2$"CDS seq" <- get(paste0(species, tissue_dot, ".CDS_seqs"))
        supp2$"log10 mRNA (RPKM)" <- get(paste0(species, tissue_dot,
            ".log10.mRNA"))
        supp2$"log10 TR (RPKM)" <- get(paste0(species, tissue_dot,
            ".log10.TR"))
        supp2$"5' motif score" <- get(paste0(species, tissue_dot,
            ".log10.TR"))

        features_5motif <- cbind(get(paste0(species, tissue_dot,
            ".5ofTICE_features"))[[1]], get(paste0(species, tissue_dot,
```

```
            ".uTICE_features"))[[1]], get(paste0(species, tissue_dot,
            ".dTICE_features"))[[1]])
        supp2$"5' motif score" <- fitted(lm(get(paste0(species,
            tissue_dot, ".log10.TR")) ~ as.matrix(features_5motif),
            na.action = "na.exclude"))

        features_RNAfold <- get(paste0(species, tissue_dot, ".fold_energy_features_selected"))
        supp2$"5' UTR RNA fold engy" <- fitted(lm(get(paste0(species,
            tissue_dot, ".log10.TR")) ~ as.matrix(features_RNAfold),
            na.action = "na.exclude"))

        supp2$"5'UTR #uAUGs" <- get(paste0(species, tissue_dot,
            ".uAUG_counts"))

        features_CDSlen <- get(paste0(species, tissue_dot, ".log10.CDS_lens.div"))
        supp2$"log10 CDS length" <- fitted(lm(get(paste0(species,
            tissue_dot, ".log10.TR")) ~ as.matrix(features_CDSlen),
            na.action = "na.exclude"))

        features_CodonUsage <- get(paste0(species, tissue_dot,
            ".codon_features"))[[1]]
        supp2$"codon freq." <- fitted(lm(get(paste0(species,
            tissue_dot, ".log10.TR")) ~ as.matrix(features_CodonUsage),
            na.action = "na.exclude"))

        features_AAUsage <- get(paste0(species, tissue_dot, ".aa_features"))[[1]]
        supp2$"AA freq." <- fitted(lm(get(paste0(species, tissue_dot,
            ".log10.TR")) ~ as.matrix(features_AAUsage), na.action = "na.exclude"))

        features_SynCodonUsage <- readRDS(paste0("processed_data_w_polyA/syncodon_features/",
            species, tissue_dot, ".syncodon_features.rds"))[[1]]
        supp2$"syn.codon preference" <- fitted(lm(get(paste0(species,
            tissue_dot, ".log10.TR")) ~ as.matrix(features_SynCodonUsage),
            na.action = "na.exclude"))

        supp2$"poly A length" <- get(paste0(species, tissue_dot,
            ".polyA_lens"))
        supp2$"log10 5' UTR length" <- get(paste0(species, tissue_dot,
            ".log10.UTR_lens"))

        write.csv(supp2, file = paste0("supp/SuppFile2.", species,
            tissue_dot, ".csv"), quote = FALSE, row.names = FALSE)
        rm(supp2)
        gc()
    }
}
```

#### Supplementary file 4. Output the RNA secondary structures of the minimum folding energy windows (the most folded windows)

```
# columns id seq mfe mfe_struct max_loop max_continuous_stem
# max_stem all paired nucleotides log TR

for (species in names(species_tissue_list)) {
    for (species_tissue in species_tissue_list[[species]]) {
        if (species_tissue != "") {
            tissue_dot <- paste0(".", species_tissue)
            tissue_us <- paste0("_", species_tissue)
        } else {
            tissue_dot <- ""
            tissue_us <- ""
        }

        min_fold_energy_window_features <- read.table(paste0("data/min_window_seq_parameters/",
            species, tissue_us, ".tsv"), header = T, sep = "\t",
            stringsAsFactors = FALSE)
        rownames(min_fold_energy_window_features) <- min_fold_energy_window_features$id

        supp4 <- data.frame(id = get(paste0(species, tissue_dot,
            ".genes")))
        supp4$seq <- min_fold_energy_window_features$seq[supp4$id]
        supp4$mfe <- min_fold_energy_window_features$mfe[supp4$id]
        supp4$mfe_struct <- min_fold_energy_window_features$mfe_struct[supp4$id]
        supp4$max_loop <- min_fold_energy_window_features$max_loop[supp4$id]
        supp4$max_continuous_stem <- min_fold_energy_window_features$max_continuous_stem[supp4$id]
        supp4$max_stem <- min_fold_energy_window_features$max_stem_paired[supp4$id]
        supp4$"all paired nucleotides" <- min_fold_energy_window_features$paired_nts[supp4$id]
        supp4$"log TR" <- get(paste0(species, tissue_dot, ".log10.TR"))

        write.csv(supp4, file = paste0("supp/SuppFile4.", species,
            tissue_dot, ".csv"), quote = FALSE, row.names = FALSE)
    }
}
```

#### Supplementary file 5. The frequencies of nucleotides at each nucleotide position of PWMs. i. The AUG aligned PWMs for the top and bottom cohorts shown in Figure 7. ie from -80 to +35. PWMs for all 10 datasets. ii. The 5' cap aligned PWMs for the 5' most 50 nucleotides for the top and bottom cohorts shown in Figure 8—figure supplement 1.

```
for (species in names(species_exemplar_tissue_list)) {
    for (species_tissue in species_exemplar_tissue_list[[species]]) {
        if (species_tissue != "") {
            species.log10.TR <- get(paste(species, species_tissue,
```

```
                "log10.TR", sep = "."))
            connector <- paste0(species_tissue, ".")
        } else {
            species.log10.TR <- get(paste(species, "log10.TR",
                sep = "."))
            connector <- ""
        }

        species.TR_top_PWM_AUG_aligned <- readRDS(file = paste0("processed_data_w_polyA/AUGPWM_R2s/",
            species, ".", connector, "TR_top_PWM_AUG_aligned.rds"))
        write.csv(species.TR_top_PWM_AUG_aligned[21:nrow(species.TR_top_PWM_AUG_aligned),
            ], file = paste0("supp/SuppFile5.", species, tissue_dot,
            ".TR_top_PWM_AUG_aligned.csv"), quote = FALSE, row.names = TRUE)

        species.TR_bottom_PWM_AUG_aligned <- readRDS(file = paste0("processed_data_w_polyA/AUGPWM_R2s/"
            species, ".", connector, "TR_bottom_PWM_AUG_aligned.rds"))
        write.csv(species.TR_bottom_PWM_AUG_aligned[21:nrow(species.TR_bottom_PWM_AUG_aligned),
            ], file = paste0("supp/SuppFile5.", species, tissue_dot,
            ".TR_bottom_PWM_AUG_aligned.csv"), quote = FALSE,
            row.names = TRUE)

        species.TR_top_PWM_5_aligned <- readRDS(file = paste0("processed_data_w_polyA/5'PWM_R2s/",
            species, ".", connector, "TR_top_PWM_5_aligned.rds"))
        write.csv(species.TR_top_PWM_5_aligned[1:50, ], file = paste0("supp/SuppFile5.",
            species, tissue_dot, ".TR_top_PWM_5_aligned.csv"),
            quote = FALSE, row.names = TRUE)

        species.TR_bottom_PWM_5_aligned <- readRDS(file = paste0("processed_data_w_polyA/5'PWM_R2s/",
            species, ".", connector, "TR_bottom_PWM_5_aligned.rds"))
        write.csv(species.TR_bottom_PWM_5_aligned[1:50, ], file = paste0("supp/SuppFile5.",
            species, tissue_dot, ".TR_bottom_PWM_5_aligned.csv"),
            quote = FALSE, row.names = TRUE)
    }
}
```

#### Supplementary file 6. Compare the Sc. TR data from Weinberg et al to that of Cuperus et al.

```
# columns gene name cupress 5' UTR seq weinberg 5' UTR seq
# cupress log 10 growth rate weinberg log 10 IE

sc.cuperus.data <- read_xlsx("data/Supplementary file 6 Cuperus vs weinberg.xlsx",
    sheet = 1, range = "A2:N2259")
sc.cuperus.data <- as.data.frame(sc.cuperus.data)

sc.cuperus.genes <- sc.cuperus.data[, 1]
sc.cuperus.log10.TR <- sc.cuperus.data[, 2]
names(sc.cuperus.log10.TR) <- sc.cuperus.genes
sc.cuperus.UTR_lens <- sc.cuperus.data$"5' UTR length"
names(sc.cuperus.UTR_lens) <- sc.cuperus.genes
sc.cuperus.UTR_seqs <- sc.cuperus.data$"UTR sequence"
```

```
names(sc.cuperus.UTR_seqs) <- sc.cuperus.genes

rm(sc.cuperus.data)

sc.common.genes <- intersect(sc.genes, sc.cuperus.genes)
# check the UTR lengths
sc.UTR_lens <- 10^sc.log10.UTR_lens
idx <- which(abs(sc.UTR_lens[sc.common.genes] - sc.cuperus.UTR_lens[sc.common.genes]) <
    2)

cor(sc.log10.TR[sc.common.genes][idx], sc.cuperus.log10.TR[sc.common.genes][idx])
```

```
## [1] 0.3065955
```

```
cor(sc.log10.TR[sc.common.genes][idx], sc.cuperus.log10.TR[sc.common.genes][idx])^2
```

```
## [1] 0.0940008
```

```
supp6 <- data.frame(gene = sc.common.genes[idx])
supp6$"cupress 5' UTR seq" <- sc.cuperus.UTR_seqs[sc.common.genes][idx]
supp6$"weinberg 5' UTR seq" <- sc.UTR_seqs[sc.common.genes][idx]
supp6$"cupress log 10 growth rate" <- sc.cuperus.log10.TR[sc.common.genes][idx]
supp6$"weinberg log 10 IE" <- sc.log10.TR[sc.common.genes][idx]

write.csv(supp6, file = "supp/SuppFile6.csv", quote = FALSE,
    row.names = FALSE)
```
