## Additional file 11 for "Quantitative Principles of *cis*-translational control by general mRNA sequence features in eukaryotes": Code.Rmd

````
---
title: 'Code for reproducing the results in "Principles of cis-translational control by general mRNA sequence features in six model eukaryotes"'
output: 
  pdf_document:
    keep_tex: true
---
The intermediate files and final output files have been generated and stored in subdirectories. The code chunks that take time to generate intermediate files will not run by default.

```{r}
library(knitr)
opts_chunk$set(tidy.opts=list(width.cutoff=60), tidy=TRUE)
```

# Required R Packages
```{r, warning=FALSE}
library(parallel)
library(MASS)
if (!require("readxl")) {
   install.packages("readxl", dependencies = TRUE)
   library(readxl)
}
if (!require("lmodel2")) {
   install.packages("lmodel2", dependencies = TRUE)
   library(lmodel2)
}
if (!require("stringi")) {
   install.packages("stringi", dependencies = TRUE)
   library(stringi)
}
if (!require("stringr")) {
   install.packages("stringr", dependencies = TRUE)
   library(stringr)
}
if (!require("leaps")) {
   install.packages("leaps", dependencies = TRUE)
   library(leaps)
}
if (!require("Biostrings")) {
   install.packages("Biostrings", dependencies = TRUE)
   library(Biostrings)
}
if (!require("cocor")) {
   install.packages("cocor", dependencies = TRUE)
   library(cocor)
}
if (!require("motifStack")) {
   install.packages("motifStack", dependencies = TRUE)
   library(motifStack)
}
if (!require("gplots")) {
   install.packages("gplots", dependencies = TRUE)
   library(gplots)
}
if (!require("RColorBrewer")) {
   install.packages("RColorBrewer", dependencies = TRUE)
   library(RColorBrewer)
}
if (!require("reshape2")) {
   install.packages("reshape2", dependencies = TRUE)
   library(reshape2)
}
if (!require("ggplot2")) {
   install.packages("ggplot2", dependencies = TRUE)
   library(ggplot2)
}
if (!require("zoo")) {
   install.packages("zoo", dependencies = TRUE)
   library(zoo)
}
if (!require("lattice")) {
   install.packages("lattice", dependencies = TRUE)
   library(lattice)
}
if (!require("gdata")) {
   install.packages("gdata", dependencies = TRUE)
   library(lattice)
}
if (!require("viridis")) {
   install.packages("viridis", dependencies = TRUE)
   library(lattice)
}
if (!require("xlsx")) {
   install.packages("xlsx", dependencies = TRUE)
   library(lattice)
}
```

# Preprocess data
```{r, eval = FALSE}
# a function to resolve duplicated short gene names (gene IDs) for sequences
resolve_dup_genes <- function(long_seqs, short_names, tissue_long_seqs=NULL) {
	# long_seqs: a character vector of UTR sequences, whose names are long gene names (e.g., transcript IDs)
	# short_names: a character vector of short gene names (e.g., gene IDs); short_names are mathed with long_seqs
	# tissue_long_seqs (optional): a character vector of one tissue's UTR sequences, whose names are long gene names (e.g., transcript IDs); if not supplied, the longest sequence will be picked for each short name
	long_names <- names(long_seqs)
	dup_idx <- which(duplicated(short_names))
	if (length(dup_idx)>0) {
		dup_genes <- unique(short_names[dup_idx])
		non_dup_idx <- which(!(short_names %in% dup_genes))
		IDmap <- long_names[non_dup_idx]
		names(IDmap) <- short_names[non_dup_idx]
		if (!is.null(tissue_long_seqs)) {
			checkif <- long_names %in% names(tissue_long_seqs)
		}
		for (dup_gene in dup_genes) {
			if (!is.null(tissue_long_seqs)) {
				tmp_idx <- which(short_names == dup_gene & checkif)
			} else {
				tmp_idx <- which(short_names == dup_gene)
			}
			tmp <- nchar(long_seqs[tmp_idx])
			tmp_idx <- tmp_idx[which.max(tmp)]
			IDmap_add <- long_names[tmp_idx]
			IDmap_name_old <- names(IDmap)
			IDmap_name_add <- short_names[tmp_idx]	
			IDmap <- c(IDmap, IDmap_add)
			names(IDmap) <- c(IDmap_name_old, IDmap_name_add)
		}
	} else {
		IDmap <- long_names
		names(IDmap) <- short_names
	}
	return(IDmap)
	# output:
	# IDmap: a character vector of short names, with names as long names
}

############################## S. cerevisae ##############################

# translational rates and features
sc.data <- read_xlsx("data/Dataset S6 mRNA seq features.xlsx", sheet=1, range="A23:Q2473")
sc.data <- as.data.frame(sc.data)

# CDS sequences
sc.CDS_seqs <- sc.data$"CDS sequence"
names(sc.CDS_seqs) <- sc.data$ORF

# 5' UTR sequences
sc.UTR_seqs <- sc.data$"5' UTR sequence"
names(sc.UTR_seqs) <- sc.data$ORF

# genes
sc.genes <- sc.data$ORF

# TR data
sc.log10.TR <- sc.data$"log10 TR"
names(sc.log10.TR) <- sc.genes

# mRNA data
sc.log10.mRNA <- sc.data$"log10 RNA"
names(sc.log10.mRNA) <- sc.genes

# UTR sequences
sc.UTR_seqs <- sc.UTR_seqs[sc.genes]

# CDS sequences
sc.CDS_seqs <- sc.CDS_seqs[sc.genes]

# poly A tail length
sc.polyA_lens <- sc.data$"poly-A tail length"
names(sc.polyA_lens) <- sc.genes

# save the processed data to an .Rdata file
save(sc.data, sc.genes, sc.log10.TR, sc.log10.mRNA, sc.UTR_seqs, sc.CDS_seqs, sc.polyA_lens, file="processed_data_w_polyA/sc.Rdata")


############################## S. pombe ##############################

# CDS sequences (downloaded from pombase.org, date 11/1/17)
sp.CDS_fa <- scan("data/s_pombe_cds.fa", what=character(), sep="\n")
name_idx <- which( startsWith(str=sp.CDS_fa, pattern=">") )
n_seq <- length(name_idx)
sp.genes <- gsub(pattern=">", replacement="", x=sp.CDS_fa[name_idx])
sp.CDS_seqs <- sapply(1:n_seq, FUN=function(i) {
	start_idx <- name_idx[i] + 1
	if (i < n_seq) {
		stop_idx <- name_idx[i+1] - 1
	} else {
		stop_idx <- length(sp.CDS_fa)
	}
	paste(sp.CDS_fa[start_idx:stop_idx], collapse="")
})
names(sp.CDS_seqs) <- sp.genes
rm(sp.CDS_fa, sp.genes)

# 5' UTR sequences (downloaded from pombase.org, date 10/18/17)
sp.UTR_fa <- scan("data/s_pombe_5UTR.fa", what=character(), sep="\n")
name_idx <- which( startsWith(str=sp.UTR_fa, pattern=">") )
n_seq <- length(name_idx)
sp.genes <- sapply( strsplit( gsub(pattern=">", replacement="", x=sp.UTR_fa[name_idx]), "\\|" ), FUN=function(x) x[1])
sp.UTR_seqs <- sapply(1:n_seq, FUN=function(i) {
	start_idx <- name_idx[i] + 1
	if (i < n_seq) {
		stop_idx <- name_idx[i+1] - 1
	} else {
		stop_idx <- length(sp.UTR_fa)
	}
	paste(sp.UTR_fa[start_idx:stop_idx], collapse="")
})
names(sp.UTR_seqs) <- sp.genes
rm(sp.UTR_fa, sp.genes)

################## first dataset ##################

# translational rates and features
sp.data <- read_xlsx("data/Subtelny GSE52809_Pombe.xlsx", sheet=1, range="A5:I3950")
sp.data <- as.data.frame(sp.data)
## the first two columns are exactly the same
sp.data <- sp.data[,-1]

# genes
sp.genes <- sapply(strsplit(sp.data$"Gene name", "\\."), FUN=function(x) paste(x[1:2], collapse="."))

# TR data
sp.log10.TR <- sp.data$"log 10 TE RPKM"
names(sp.log10.TR) <- sp.genes

# mRNA data
sp.log10.mRNA <- sp.data$"log 10 mRNA RPKM"
names(sp.log10.mRNA) <- sp.genes

# UTR sequences
sp.UTR_seqs <- sp.UTR_seqs[sp.genes]

# CDS sequences
sp.CDS_seqs <- sp.CDS_seqs[sp.genes]

# poly A tail length
sp.polyA_lens <- sp.data$"Mean poly A tail length"
names(sp.polyA_lens) <- sp.genes

# remove the genes with missing information
sp.rm_idx <- which(is.na(sp.log10.TR) | is.na(sp.log10.mRNA) | is.na(sp.UTR_seqs) | is.na(sp.CDS_seqs) | is.na(sp.polyA_lens))
sp.data <- sp.data[-sp.rm_idx,]
sp.genes <- sp.genes[-sp.rm_idx]
sp.log10.TR <- sp.log10.TR[-sp.rm_idx]
sp.log10.mRNA <- sp.log10.mRNA[-sp.rm_idx]
sp.UTR_seqs <- sp.UTR_seqs[-sp.rm_idx]
sp.CDS_seqs <- sp.CDS_seqs[-sp.rm_idx]
sp.polyA_lens <- sp.polyA_lens[-sp.rm_idx]

# save the processed data to an .Rdata file
save(sp.data, sp.genes, sp.log10.TR, sp.log10.mRNA, sp.UTR_seqs, sp.CDS_seqs, sp.polyA_lens, file="processed_data_w_polyA/sp.Rdata")

################## second dataset ##################

# translational rates (preprocessing)
sp.alt.data <- read_xlsx("data/Pombe Mata 10.23.17 N_present_all_counts.xlsx", sheet=1, range="A1:E5161")
sp.alt.data <- as.data.frame(sp.alt.data)
for (j in 2:ncol(sp.alt.data)) {
	sp.alt.data[,j] <- sp.alt.data[,j]/(sum(sp.alt.data[,j])/1000000) # RPM
}
colnames(sp.alt.data)[1] <- "Systematic_ID"

## annotated ORF lengths
sp.alt.CDS_lens <- read_xlsx("data/Pombe Mata 10.23.17 N_present_all_counts.xlsx", sheet=2, range="A1:B5124")
sp.alt.CDS_lens <- as.data.frame(sp.alt.CDS_lens)

## make the data frame
sp.alt.data <- merge(sp.alt.data, sp.alt.CDS_lens, by="Systematic_ID")
for (j in 2:(ncol(sp.alt.data)-1)) {
	sp.alt.data[,j] <- sp.alt.data[,j]/(sp.alt.data[,ncol(sp.alt.data)]/1000) # RPKM
}
sp.alt.data$RD_mean <- rowMeans(sp.alt.data[,2:3])
sp.alt.data$RNA_mean <- rowMeans(sp.alt.data[,4:5])

## thresholded to >= 1 RPKM for RNA density and > 0 for RD density
sp.alt.data <- sp.alt.data[sp.alt.data$RNA_mean >= 1 & sp.alt.data$RD_mean > 0,]
sp.alt.data$TE <- sp.alt.data$RD_mean / sp.alt.data$RNA_mean

# genes
sp.alt.genes <- sp.alt.data$Systematic_ID

# TR data
sp.alt.log10.TR <- log(sp.alt.data$TE, 10)
names(sp.alt.log10.TR) <- sp.alt.genes

# mRNA data
sp.alt.log10.mRNA <- log(sp.alt.data$RNA_mean, 10)
names(sp.alt.log10.mRNA) <- sp.alt.genes

# UTR sequences
sp.alt.UTR_seqs <- sp.UTR_seqs[sp.alt.genes]

# CDS sequences
sp.alt.CDS_seqs <- sp.CDS_seqs[sp.alt.genes]

# poly A tail length
sp.alt.polyA_lens <- sp.polyA_lens[sp.alt.genes]
names(sp.alt.polyA_lens) <- sp.alt.genes

# remove the genes with missing information
sp.alt.rm_idx <- which(is.na(sp.alt.log10.TR) | is.na(sp.alt.log10.mRNA) | is.na(sp.alt.UTR_seqs) | is.na(sp.alt.CDS_seqs) | is.na(sp.alt.polyA_lens))
sp.alt.data <- sp.alt.data[-sp.alt.rm_idx,]
sp.alt.genes <- sp.alt.genes[-sp.alt.rm_idx]
sp.alt.log10.TR <- sp.alt.log10.TR[-sp.alt.rm_idx]
sp.alt.log10.mRNA <- sp.alt.log10.mRNA[-sp.alt.rm_idx]
sp.alt.UTR_seqs <- sp.alt.UTR_seqs[-sp.alt.rm_idx]
sp.alt.CDS_seqs <- sp.alt.CDS_seqs[-sp.alt.rm_idx]
sp.alt.polyA_lens <- sp.alt.polyA_lens[-sp.alt.rm_idx]

# save the processed data to an .Rdata file
save(sp.alt.genes, sp.alt.log10.TR, sp.alt.log10.mRNA, sp.alt.UTR_seqs, sp.alt.CDS_seqs, sp.alt.polyA_lens, file="processed_data_w_polyA/sp.alt.Rdata")


############################## Arabidopsis ##############################

# CDS sequences
at.CDS_seqs <- readRDS(file="data/Arabidopsis_CDS.rds")
at.CDS_seqs_short_names <- sapply(strsplit(names(at.CDS_seqs), "\\."), FUN=function(x) x[1])

# 5' UTR sequences
at.UTR_seqs <- readRDS(file="data/Arabidopsis_UTRs.rds")
at.UTR_seqs_short_names <- sapply(strsplit(names(at.UTR_seqs), "\\."), FUN=function(x) x[1])

# Hsu data for root and shoot
at.hsu_data <- read_xlsx("data/Hsu et al counts_cds_Hsu.xlsx", sheet=1, range="A7:M27213")
at.hsu_data <- as.data.frame(at.hsu_data)
colnames(at.hsu_data)[1] <- "gene_id"
for (j in 2:ncol(at.hsu_data)) {
	at.hsu_data[,j] <- at.hsu_data[,j]/(sum(at.hsu_data[,j])/1000000) # RPM
}

################## Leaf ##################

# Leaf translational rates and features (data preprocessing in ``030218_mouse_and_arabidopsis_models.Rmd"")
at.leaf.data <- readRDS("data/030218_arab_data.rds")

# genes
at.leaf.genes <- at.leaf.data$GeneID

# TR data
at.leaf.log10.TR <- at.leaf.data$"Log10..TE.Dark"
names(at.leaf.log10.TR) <- at.leaf.genes

# mRNA data
at.leaf.log10.mRNA <- at.leaf.data$"log.10.mRNA.Dark"
names(at.leaf.log10.mRNA) <- at.leaf.genes

# UTR sequences
at.leaf.UTR_seqs <- at.UTR_seqs[at.leaf.genes]

# CDS sequences
at.leaf.CDS_seqs <- at.CDS_seqs[at.leaf.genes]

# poly A tail length
at.leaf.polyA_lens <- at.leaf.data$"poly-A tail length"
names(at.leaf.polyA_lens) <- at.leaf.genes

# save the processed data to an .Rdata file
save(at.leaf.data, at.leaf.genes, at.leaf.log10.TR, at.leaf.log10.mRNA, at.leaf.UTR_seqs, at.leaf.CDS_seqs, at.leaf.polyA_lens, file="processed_data_w_polyA/at.leaf.Rdata")

############### Root & Shoot ###############

# both tissues have short gene names, so we will resolve the duplicated short gene names 
at.IDmap <- resolve_dup_genes(long_seqs=at.UTR_seqs, short_names=at.UTR_seqs_short_names, tissue_long_seqs=at.leaf.UTR_seqs)

saveRDS(at.IDmap, file="processed_data_w_polyA/at.IDmap.rds")

# a function to resolve duplicated short gene names (gene IDs) for root and shoot data based on the leaf tissue CDS sequences
resolve_dup_genes_root_shoot_data <- function(data, leaf_CDS_seqs) {
	# data: a data frame with two key columns: "gene_id" contains duplicated short gene names; "Annotated ORF length" contains ORF lengths, which are CDS lengths - 3
	# leaf_CDS_seqs: a character vector of leaf tissue's CDS sequences
	leaf_short_names <- sapply(strsplit(names(leaf_CDS_seqs), "\\."), FUN=function(x) x[1])
	short_names <- data$gene_id
	dup_idx <- which(duplicated(short_names))
	if (length(dup_idx)>0) {
		dup_genes <- unique(short_names[dup_idx])
		non_dup_idx <- which(!(short_names %in% dup_genes))
		data_tmp <- data[non_dup_idx, ]
		for (dup_gene in dup_genes) {
			tmp_idx <- which(short_names == dup_gene)
			CDS_len <- nchar(leaf_CDS_seqs[leaf_short_names==dup_gene])
			ORF_lens <- data[tmp_idx, "Annotated ORF length"]
			tmp_idx <- tmp_idx[ORF_lens==CDS_len-3]
			data_tmp <- rbind(data_tmp, data[tmp_idx[1],])
		}
	} else {
		data_tmp <- data
	}
	return(data_tmp)
}

################## Root ##################

# Root translational rates (preprocessing)
## annotated ORF lengths
at.root.ORF_lens <- read_xlsx("data/Hsu et al counts_cds_Hsu.xlsx", sheet=2, range="A3:E20986")
at.root.ORF_lens <- as.data.frame(at.root.ORF_lens)
at.root.ORF_lens <- at.root.ORF_lens[ , c("gene_id", "Annoated ORF length")]
## make the data frame
at.root.data <- merge(at.hsu_data, at.root.ORF_lens, by="gene_id")
for (j in 2:(ncol(at.root.data)-1)) {
	at.root.data[,j] <- at.root.data[,j]/(at.root.data[,ncol(at.root.data)]/1000) # RPKM
}
at.root.data$RD_mean <- rowMeans(at.root.data[,2:4])
at.root.data$RNA_mean <- rowMeans(at.root.data[,8:10])
## thresholded to >= 1 RPKM for RNA density and > 0 for RD density
at.root.data <- at.root.data[at.root.data$RNA_mean >= 1 & at.root.data$RD_mean > 0,]
at.root.data$TE <- at.root.data$RD_mean / at.root.data$RNA_mean

# remove duplicate gene names based on the leaf tissue
at.root.data <- resolve_dup_genes_root_shoot_data(data=at.root.data, leaf_CDS_seqs=at.leaf.CDS_seqs)

# genes
at.root.genes <- at.root.data$gene_id

# TR data
at.root.log10.TR <- log(at.root.data$TE, 10)
names(at.root.log10.TR) <- at.root.genes

# mRNA data
at.root.log10.mRNA <- log(at.root.data$RNA_mean, 10)
names(at.root.log10.mRNA) <- at.root.genes

# UTR sequences
at.root.UTR_seqs <- at.UTR_seqs[at.IDmap[at.root.genes]]
names(at.root.UTR_seqs) <- at.root.genes

# CDS sequences
at.root.CDS_seqs <- at.CDS_seqs[at.IDmap[at.root.genes]]
names(at.root.CDS_seqs) <- at.root.genes

# poly A tail length
at.root.polyA_lens <- at.leaf.polyA_lens[at.IDmap[at.root.genes]]
names(at.root.polyA_lens) <- at.root.genes

# remove the genes with missing information
at.root.rm_idx <- which(is.na(at.root.log10.TR) | is.na(at.root.log10.mRNA) | is.na(at.root.UTR_seqs) | is.na(at.root.CDS_seqs) | is.na(at.root.polyA_lens))
at.root.data <- at.root.data[-at.root.rm_idx,]
at.root.genes <- at.root.genes[-at.root.rm_idx]
at.root.log10.TR <- at.root.log10.TR[-at.root.rm_idx]
at.root.log10.mRNA <- at.root.log10.mRNA[-at.root.rm_idx]
at.root.UTR_seqs <- at.root.UTR_seqs[-at.root.rm_idx]
at.root.CDS_seqs <- at.root.CDS_seqs[-at.root.rm_idx]
at.root.polyA_lens <- at.root.polyA_lens[-at.root.rm_idx]

# save the processed data to an .Rdata file
save(at.root.genes, at.root.log10.TR, at.root.log10.mRNA, at.root.UTR_seqs, at.root.CDS_seqs, at.root.polyA_lens, file="processed_data_w_polyA/at.root.Rdata")

################## Shoot ##################

# Shoot translational rates (preprocessing)
## annotated ORF lengths
at.shoot.ORF_lens <- read_xlsx("data/Hsu et al counts_cds_Hsu.xlsx", sheet=3, range="A3:E21579")
at.shoot.ORF_lens <- as.data.frame(at.shoot.ORF_lens)
at.shoot.ORF_lens <- at.shoot.ORF_lens[ , c("gene_id", "Annotated ORF length")]
## make the data frame
at.shoot.data <- merge(at.hsu_data, at.shoot.ORF_lens, by="gene_id")
for (j in 2:(ncol(at.shoot.data)-1)) {
	at.shoot.data[,j] <- at.shoot.data[,j]/(at.shoot.data[,ncol(at.shoot.data)]/1000) # RPKM
}
at.shoot.data$RD_mean <- rowMeans(at.shoot.data[,5:7])
at.shoot.data$RNA_mean <- rowMeans(at.shoot.data[,11:13])
## thresholded to >= 1 RPKM for RNA density
at.shoot.data <- at.shoot.data[at.shoot.data$RNA_mean >= 1 & at.shoot.data$RD_mean > 0,]
at.shoot.data$TE <- at.shoot.data$RD_mean / at.shoot.data$RNA_mean

# remove duplicate gene names based on the leaf tissue
at.shoot.data <- resolve_dup_genes_root_shoot_data(data=at.shoot.data, leaf_CDS_seqs=at.leaf.CDS_seqs)

# genes
at.shoot.genes <- at.shoot.data$gene_id

# TR data
at.shoot.log10.TR <- log(at.shoot.data$TE, 10)
names(at.shoot.log10.TR) <- at.shoot.genes

# mRNA data
at.shoot.log10.mRNA <- log(at.shoot.data$RNA_mean, 10)
names(at.shoot.log10.mRNA) <- at.shoot.genes

# UTR sequences
at.shoot.UTR_seqs <- at.UTR_seqs[at.IDmap[at.shoot.genes]]
names(at.shoot.UTR_seqs) <- at.shoot.genes

# CDS sequences
at.shoot.CDS_seqs <- at.CDS_seqs[at.IDmap[at.shoot.genes]]
names(at.shoot.CDS_seqs) <- at.shoot.genes

# poly A tail length
at.shoot.polyA_lens <- at.leaf.polyA_lens[at.IDmap[at.shoot.genes]]
names(at.shoot.polyA_lens) <- at.shoot.genes

# remove the genes with missing information
at.shoot.rm_idx <- which(is.na(at.shoot.log10.TR) | is.na(at.shoot.log10.mRNA) | is.na(at.shoot.UTR_seqs) | is.na(at.shoot.CDS_seqs) | is.na(at.shoot.polyA_lens))
at.shoot.data <- at.shoot.data[-at.shoot.rm_idx,]
at.shoot.genes <- at.shoot.genes[-at.shoot.rm_idx]
at.shoot.log10.TR <- at.shoot.log10.TR[-at.shoot.rm_idx]
at.shoot.log10.mRNA <- at.shoot.log10.mRNA[-at.shoot.rm_idx]
at.shoot.UTR_seqs <- at.shoot.UTR_seqs[-at.shoot.rm_idx]
at.shoot.CDS_seqs <- at.shoot.CDS_seqs[-at.shoot.rm_idx]
at.shoot.polyA_lens <- at.shoot.polyA_lens[-at.shoot.rm_idx]

# save the processed data to an .Rdata file
save(at.shoot.genes, at.shoot.log10.TR, at.shoot.log10.mRNA, at.shoot.UTR_seqs, at.shoot.CDS_seqs, at.shoot.polyA_lens, file="processed_data_w_polyA/at.shoot.Rdata")


############################## Mouse ##############################

# CDS sequences
mm.CDS_seqs <- readRDS(file="data/Mouse Ensembl CDS.rds")

mm.CDS_fa <- scan("data/Mouse Ensembl CDS.txt", what=character(), sep="\n")
name_idx <- which( startsWith(str=mm.CDS_fa, pattern=">") )
mm.CDS_seqs_gene_trpt_names <- sapply(strsplit( gsub(pattern=">", replacement="", x=mm.CDS_fa[name_idx]), "\\|" ), FUN=function(x) x)
mm.CDS_seqs_gene_names <- mm.CDS_seqs_gene_trpt_names[1,]
names(mm.CDS_seqs_gene_names) <- mm.CDS_seqs_gene_trpt_names[2,]
mm.CDS_seqs_alt_names <- mm.CDS_seqs_gene_names[names(mm.CDS_seqs)]

rm(mm.CDS_fa, name_idx, mm.CDS_seqs_gene_trpt_names, mm.CDS_seqs_gene_names)

# 5' UTR sequences
mm.UTR_seqs <- readRDS(file="data/Mouse Ensembl 5' UTRs.rds")

mm.UTR_fa <- scan("data/Mouse Ensembl 5' UTRs.txt", what=character(), sep="\n")
name_idx <- which( startsWith(str=mm.UTR_fa, pattern=">") )
mm.UTR_seqs_gene_trpt_names <- sapply(strsplit( gsub(pattern=">", replacement="", x=mm.UTR_fa[name_idx]), "\\|" ), FUN=function(x) x)
mm.UTR_seqs_gene_names <- mm.UTR_seqs_gene_trpt_names[1,]
names(mm.UTR_seqs_gene_names) <- mm.UTR_seqs_gene_trpt_names[2,]
mm.UTR_seqs_alt_names <- mm.UTR_seqs_gene_names[names(mm.UTR_seqs)]

rm(mm.UTR_fa, name_idx, mm.UTR_seqs_gene_trpt_names, mm.UTR_seqs_gene_names)

################## NIH3T3 ##################

# NIH3T3 translational rates and features (data preprocessing in ``030218_mouse_and_arabidopsis_models.Rmd"")
mm.nih3t3.data <- readRDS(file="data/mouse_data_Subtelny_NIH3T3.rds")
mm.nih3t3.data <- mm.nih3t3.data[complete.cases(mm.nih3t3.data),]

# genes
mm.nih3t3.genes <- mm.nih3t3.data$Ensembl

# TR data
mm.nih3t3.log10.TR <- mm.nih3t3.data$"log10 TE"
names(mm.nih3t3.log10.TR) <- mm.nih3t3.genes

# mRNA data
mm.nih3t3.log10.mRNA <- mm.nih3t3.data$"log10 mRNA"
names(mm.nih3t3.log10.mRNA) <- mm.nih3t3.genes

# UTR sequences
mm.nih3t3.UTR_seqs <- mm.UTR_seqs[mm.nih3t3.genes]

# CDS sequences
mm.nih3t3.CDS_seqs <- mm.CDS_seqs[mm.nih3t3.genes]

# poly A tail length
mm.nih3t3.polyA_lens <- mm.nih3t3.data$"poly-A tail length"
names(mm.nih3t3.polyA_lens) <- mm.nih3t3.genes

# save the processed data to an .Rdata file
save(mm.nih3t3.data, mm.nih3t3.genes, mm.nih3t3.log10.TR, mm.nih3t3.log10.mRNA, mm.nih3t3.UTR_seqs, mm.nih3t3.CDS_seqs, mm.nih3t3.polyA_lens, file="processed_data_w_polyA/mm.nih3t3.Rdata")

############### Liver & Kidney ###############

# both tissues have short gene names, so we will resolve the duplicated short gene names 
mm.IDmap <- resolve_dup_genes(long_seqs=mm.UTR_seqs, short_names=mm.UTR_seqs_alt_names, tissue_long_seqs=mm.nih3t3.UTR_seqs)

saveRDS(mm.IDmap, file="processed_data_w_polyA/mm.IDmap.rds")

################## Liver ##################

# translational rates
mm.liver.data <- read_xlsx("data/GSM1644076 liver 1 RPKM RNA.rpkm copy.xlsx", sheet=3, range="A2:F10097")
mm.liver.data <- as.data.frame(mm.liver.data)

# genes
mm.liver.genes <- mm.liver.data[,1]

# TR data
mm.liver.log10.TR <- mm.liver.data$"log TE"
names(mm.liver.log10.TR) <- mm.liver.genes

# mRNA data
mm.liver.log10.mRNA <- mm.liver.data$"log RNA"
names(mm.liver.log10.mRNA) <- mm.liver.genes

# UTR sequences
mm.liver.UTR_seqs <- mm.UTR_seqs[mm.IDmap[mm.liver.genes]]
names(mm.liver.UTR_seqs) <- mm.liver.genes

# CDS sequences
mm.liver.CDS_seqs <- mm.CDS_seqs[mm.IDmap[mm.liver.genes]]
names(mm.liver.CDS_seqs) <- mm.liver.genes

# poly A tail length
mm.liver.polyA_lens <- mm.nih3t3.polyA_lens[mm.IDmap[mm.liver.genes]]
names(mm.liver.polyA_lens) <- mm.liver.genes

# remove the genes with missing information
mm.liver.rm_idx <- which(is.na(mm.liver.log10.TR) | is.na(mm.liver.log10.mRNA) | is.na(mm.liver.UTR_seqs) | is.na(mm.liver.CDS_seqs) | is.na(mm.liver.polyA_lens))
mm.liver.data <- mm.liver.data[-mm.liver.rm_idx,]
mm.liver.genes <- mm.liver.genes[-mm.liver.rm_idx]
mm.liver.log10.TR <- mm.liver.log10.TR[-mm.liver.rm_idx]
mm.liver.log10.mRNA <- mm.liver.log10.mRNA[-mm.liver.rm_idx]
mm.liver.UTR_seqs <- mm.liver.UTR_seqs[-mm.liver.rm_idx]
mm.liver.CDS_seqs <- mm.liver.CDS_seqs[-mm.liver.rm_idx]
mm.liver.polyA_lens <- mm.liver.polyA_lens[-mm.liver.rm_idx]

# save the processed data to an .Rdata file
save(mm.liver.genes, mm.liver.log10.TR, mm.liver.log10.mRNA, mm.liver.UTR_seqs, mm.liver.CDS_seqs, mm.liver.polyA_lens, file="processed_data_w_polyA/mm.liver.Rdata")

################## Kidney ##################

# translational rates
mm.kidney.data <- read_xlsx("data/GSM2149102_Kidney 1 RPKM RNA copy 2.xlsx", sheet=3, range="A2:F11654")
mm.kidney.data <- as.data.frame(mm.kidney.data)

# genes
mm.kidney.genes <- mm.kidney.data[,1]

# TR data
mm.kidney.log10.TR <- mm.kidney.data$"log TE"
names(mm.kidney.log10.TR) <- mm.kidney.genes

# mRNA data
mm.kidney.log10.mRNA <- mm.kidney.data$"log RNA"
names(mm.kidney.log10.mRNA) <- mm.kidney.genes

# UTR sequences
mm.kidney.UTR_seqs <- mm.UTR_seqs[mm.IDmap[mm.kidney.genes]]
names(mm.kidney.UTR_seqs) <- mm.kidney.genes

# CDS sequences
mm.kidney.CDS_seqs <- mm.CDS_seqs[mm.IDmap[mm.kidney.genes]]
names(mm.kidney.CDS_seqs) <- mm.kidney.genes

# poly A tail length
mm.kidney.polyA_lens <- mm.nih3t3.polyA_lens[mm.IDmap[mm.kidney.genes]]
names(mm.kidney.polyA_lens) <- mm.kidney.genes

# remove the genes with missing information
mm.kidney.rm_idx <- which(is.na(mm.kidney.log10.TR) | is.na(mm.kidney.log10.mRNA) | is.na(mm.kidney.UTR_seqs) | is.na(mm.kidney.CDS_seqs) | is.na(mm.kidney.polyA_lens))
mm.kidney.data <- mm.kidney.data[-mm.kidney.rm_idx,]
mm.kidney.genes <- mm.kidney.genes[-mm.kidney.rm_idx]
mm.kidney.log10.TR <- mm.kidney.log10.TR[-mm.kidney.rm_idx]
mm.kidney.log10.mRNA <- mm.kidney.log10.mRNA[-mm.kidney.rm_idx]
mm.kidney.UTR_seqs <- mm.kidney.UTR_seqs[-mm.kidney.rm_idx]
mm.kidney.CDS_seqs <- mm.kidney.CDS_seqs[-mm.kidney.rm_idx]
mm.kidney.polyA_lens <- mm.kidney.polyA_lens[-mm.kidney.rm_idx]

# save the processed data to an .Rdata file
save(mm.kidney.genes, mm.kidney.log10.TR, mm.kidney.log10.mRNA, mm.kidney.UTR_seqs, mm.kidney.CDS_seqs, mm.kidney.polyA_lens, file="processed_data_w_polyA/mm.kidney.Rdata")


############################## Human ##############################

# Gene ID conversion: RefSeq vs. Ensembl (downloaded from ensembl.org BioMart, GRCh38.p12)
hs.RefSeq_Ensembl <- read.table("data/GRCh38_RefSeq_to_Ensembl.txt", header=T, sep="\t", stringsAsFactors=F)
hs.RefSeq_Ensembl <- hs.RefSeq_Ensembl[hs.RefSeq_Ensembl[,1]!="" & hs.RefSeq_Ensembl[,2]!="", ]
hs.RefSeq <- hs.RefSeq_Ensembl[,2]
hs.Ensembl <- hs.RefSeq_Ensembl[,1]
names(hs.RefSeq) <- hs.Ensembl
names(hs.Ensembl) <- hs.RefSeq
rm(hs.RefSeq_Ensembl)

# CDS sequences (downloaded from ensembl.org BioMart, GRCh38.p12)
hs.CDS_fa <- scan("data/GRCh38_Ensembl_CDS.txt", what=character(), sep="\n")
name_idx <- which( startsWith(str=hs.CDS_fa, pattern=">") )
n_seq <- length(name_idx)
hs.genes <- gsub(pattern=">", replacement="", x=hs.CDS_fa[name_idx])
hs.CDS_seqs <- sapply(1:n_seq, FUN=function(i) {
	start_idx <- name_idx[i] + 1
	if (i < n_seq) {
		stop_idx <- name_idx[i+1] - 1
	} else {
		stop_idx <- length(hs.CDS_fa)
	}
	paste(hs.CDS_fa[start_idx:stop_idx], collapse="")
})
names(hs.CDS_seqs) <- hs.genes
hs.CDS_seqs <- hs.CDS_seqs[hs.CDS_seqs!="Sequence unavailable"]
rm(hs.CDS_fa)

# UTR sequences (downloaded from ensembl.org BioMart, GRCh38.p12)
hs.UTR_fa <- scan("data/GRCh38_Ensembl_5UTR.txt", what=character(), sep="\n")
name_idx <- which( startsWith(str=hs.UTR_fa, pattern=">") )
n_seq <- length(name_idx)
hs.genes <- gsub(pattern=">", replacement="", x=hs.UTR_fa[name_idx])
hs.UTR_seqs <- sapply(1:n_seq, FUN=function(i) {
	start_idx <- name_idx[i] + 1
	if (i < n_seq) {
		stop_idx <- name_idx[i+1] - 1
	} else {
		stop_idx <- length(hs.UTR_fa)
	}
	paste(hs.UTR_fa[start_idx:stop_idx], collapse="")
})
names(hs.UTR_seqs) <- hs.genes
hs.UTR_seqs <- hs.UTR_seqs[hs.UTR_seqs!="Sequence unavailable"]
rm(hs.UTR_fa)

################## Hela2 (Guo et al.) ##################

# hela2 translational rates
hs.hela2.data <- read_xlsx("data/Guo et al HeLa.xlsx", sheet=1, range="A9:G10131")
hs.hela2.data <- as.data.frame(hs.hela2.data)

# genes
hs.hela2.genes <- hs.hela2.data$"RefSeq_Accession"

# TR data
hs.hela2.log10.TR <- hs.hela2.data$"Log10 TE"
names(hs.hela2.log10.TR) <- hs.hela2.genes

# mRNA data
hs.hela2.log10.mRNA <- hs.hela2.data$"Log10 mRNA"
names(hs.hela2.log10.mRNA) <- hs.hela2.genes

# UTR sequences
hs.hela2.UTR_seqs <- hs.UTR_seqs[hs.Ensembl[hs.hela2.genes]]
names(hs.hela2.UTR_seqs) <- hs.hela2.genes

# CDS sequences
hs.hela2.CDS_seqs <- hs.CDS_seqs[hs.Ensembl[hs.hela2.genes]]
names(hs.hela2.CDS_seqs) <- hs.hela2.genes

# poly A tail length
## borrow the information from another hela data
hs.hela.data <- read_xlsx("data/Subtelny GSE52809_HeLa.xlsx", sheet=1, range="A5:K6923")
hs.hela.data <- as.data.frame(hs.hela.data)
hs.hela.genes <- hs.hela.data$"Transcript ID"
hs.hela.polyA_lens <- as.numeric(hs.hela.data$"Median poly A length")
names(hs.hela.polyA_lens) <- hs.hela.genes
hs.hela2.polyA_lens <- hs.hela.polyA_lens[hs.hela2.genes]

# remove the genes with missing information or containing "N" in sequences
hs.hela2.rm_idx <- which(is.na(hs.hela2.log10.TR) | is.na(hs.hela2.log10.mRNA) | is.na(hs.hela2.UTR_seqs) | is.na(hs.hela2.CDS_seqs) | is.na(hs.hela2.polyA_lens) | grepl("N", hs.hela2.UTR_seqs) | grepl("N", hs.hela2.CDS_seqs))
hs.hela2.data <- hs.hela2.data[-hs.hela2.rm_idx,]
hs.hela2.genes <- hs.hela2.genes[-hs.hela2.rm_idx]
hs.hela2.log10.TR <- hs.hela2.log10.TR[-hs.hela2.rm_idx]
hs.hela2.log10.mRNA <- hs.hela2.log10.mRNA[-hs.hela2.rm_idx]
hs.hela2.UTR_seqs <- hs.hela2.UTR_seqs[-hs.hela2.rm_idx]
hs.hela2.CDS_seqs <- hs.hela2.CDS_seqs[-hs.hela2.rm_idx]
hs.hela2.polyA_lens <- hs.hela2.polyA_lens[-hs.hela2.rm_idx]

# save the processed data to an .Rdata file
save(hs.hela2.data, hs.hela2.genes, hs.hela2.log10.TR, hs.hela2.log10.mRNA, hs.hela2.UTR_seqs, hs.hela2.CDS_seqs, hs.hela2.polyA_lens, file="processed_data_w_polyA/hs.hela2.Rdata")
```

# Read in the preprocessed data
```{r}
# a list of species and tissues
species_tissue_list <- list(
	sc="", 
	sp=c("", "alt"), 
	at=c("leaf", "root", "shoot"), 
	mm=c("nih3t3", "liver", "kidney"), 
	hs=c("hela2")
)

# a list of species and exemplar tissues
species_exemplar_tissue_list <- list(
	sc="", 
	sp="alt", 
	at="leaf", 
	mm="nih3t3", 
	hs="hela2"
)

# a vector of species and examplar tissue names
species_exemplar_tissue_names <- sapply(1:length(species_exemplar_tissue_list), FUN=function(i) {
  if (species_exemplar_tissue_list[[i]] != "") {
    paste(names(species_exemplar_tissue_list)[i], species_exemplar_tissue_list[[i]], sep=".")
  } else {
    names(species_exemplar_tissue_list)[i]
  }
})

# a vector of colors for species
species_cols <- c("#EB9D41", "#94241D", "#225392", "#2F6E42", "#CEC448")
names(species_cols) <- species_exemplar_tissue_names

for (species in names(species_tissue_list)) {
	for (species_tissue in species_tissue_list[[species]]) {
		if (species_tissue!="") {
			tissue_dot <- paste0(".", species_tissue)
			tissue_us <- paste0("_", species_tissue)
		} else {
			tissue_dot <- ""
			tissue_us <- ""
		}
		load(file=paste0("processed_data_w_polyA/", species, tissue_dot, ".Rdata"))
	}
}

## mouse ID map
mm.IDmap <- readRDS(file="processed_data_w_polyA/mm.IDmap.rds")

## Arabidopsis ID map
at.IDmap <- readRDS(file="processed_data_w_polyA/at.IDmap.rds")
```

# Generate and select RNA folding energy features
## Settings
```{r}
# folding energy window lengths
window_lens <- paste(c(6, 8, seq(from=10, to=100, by=5)))

# uTICE length
species_uTICE_len_list <- list(
	sc=35, 
	sp=30, # only for calculating the 5'mostTICE feature 
	at=65, 
	mm=5, 
	hs=list("hela2"=6)
)
	  
# dTICE length
species_dTICE_len_list <- list(
	sc=28, 
	sp=13,
	at=33, 
	mm=13, 
	hs=list("hela2"=7)
)

# UTR length breaks to obtain gene groups
species_len_break_list <- list(
	sc=c(20,35),
	sp=NULL, # one-part regression
	at=65,
	mm=NULL,
	hs=NULL # one-part regression
)
```
## The following code only needs to be run once
```{r, eval = FALSE}
# UTR folding energy of different window lengths
for (species_short in names(species_tissue_list)) {
	for (species_tissue in species_tissue_list[[species_short]]) {
		if (species_tissue!="") {
			species_genes <- get(paste(species_short, species_tissue, "genes", sep="."))			
		} else {
			species_genes <- get(paste(species_short, "genes", sep="."))
		}
		## read in the whole UTR folding energy feature (one value per gene)
		if (species_tissue!="") {
			whole_5UTR_folding_energy <- readRDS(file=paste0("processed_data_w_polyA/whole_UTR_folding_energy_feature/", species_short, "_", species_tissue, "_whole_UTR_folding_energy_feature.rds"))
		} else {
			whole_5UTR_folding_energy <- readRDS(file=paste0("processed_data_w_polyA/whole_UTR_folding_energy_feature/", species_short, "_whole_UTR_folding_energy_features.rds"))
		}
		
		if (species_short != "hs") {
			uTICE_len <- species_uTICE_len_list[[species_short]]
		} else {
			uTICE_len <- species_uTICE_len_list[[species_short]][[species_tissue]]
		}
		
		# generate folding energy features
		for (window_len in window_lens) {
			if (species_tissue!="") {
				UTR_folding_energy_all <- readRDS(file=paste0("processed_data_w_polyA/var_win_fold_UTR5_CDSshort_all/", species_short, "_", species_tissue, "_5UTR_CDS_folding_energy_", window_len, ".rds"))
			} else {
				UTR_folding_energy_all <- readRDS(file=paste0("processed_data_w_polyA/var_win_fold_UTR5_CDSshort_all/", species_short, "_5UTR_CDS_folding_energy_", window_len, ".rds"))
			}
			UTR_folding_energy_all <- UTR_folding_energy_all$UTR_folding_energy_all[species_genes]
			UTR_folding_energy_percentiles <- quantile(unlist(UTR_folding_energy_all), probs=c(0.05,0.1,0.2,0.8,0.9))
			UTR_folding_energy_features <- t( sapply(UTR_folding_energy_all, FUN=function(x) {
				if (is.null(x)) {
				  return(rep(NA, 18))
				} else {
			  	  x1 <- x[1]
			  	  x2 <- rev(x)[1]
			  	  x3 <- rev(x)[min(uTICE_len,length(x))]
			  	  x4 <- mean(x)
			  	  x5 <- min(x)
			  	  x6 <- max(x)
			  	  tmp <- quantile(x, probs=c(0.9, 0.1, 0.75, 0.25))
			  	  n <- length(x)
			  	  x7 <- tmp[1]
			  	  x8 <- tmp[2]
			  	  x9 <- tmp[3]
			  	  x10 <- tmp[4]
			  	  x11 <- sum(x <= UTR_folding_energy_percentiles[3]) / n
			  	  x16 <- sum(x > UTR_folding_energy_percentiles[5]) / n
			  	  x17 <- sum(x > UTR_folding_energy_percentiles[4]) / n
			  	  x13 <- sum(x[x <= UTR_folding_energy_percentiles[1]])
			  	  x14 <- sum(x[x <= UTR_folding_energy_percentiles[2]])
			  	  x15 <- sum(x[x <= UTR_folding_energy_percentiles[3]])
			  	  x18 <- sum(x[x > UTR_folding_energy_percentiles[5]])
			  	  x19 <- sum(x[x > UTR_folding_energy_percentiles[4]])
			  	  return( c(x1, x2, x3, x4, x5, x6, x7, x8, x9, x10, x11, x13, x14, x15, x16, x17, x18, x19) )
				}
			}) )
			UTR_folding_energy_features <- cbind(UTR_folding_energy_features, whole_5UTR_folding_energy)

			# deal with the missing data
			if (any(is.na(UTR_folding_energy_features[,1]))) {
				missing_idx <- which(is.na(UTR_folding_energy_features[,1]))
				for (i in missing_idx) {
					for (j in 1:(ncol(UTR_folding_energy_features)-1)) {
						quantile_ij <- sum(UTR_folding_energy_features_old[,j] <= UTR_folding_energy_features_old[i,j]) / nrow(UTR_folding_energy_features)
						UTR_folding_energy_features[i,j] <- quantile(UTR_folding_energy_features[,j], probs=quantile_ij, na.rm=T)
					}
				}
			}

			colnames(UTR_folding_energy_features) <- c("5'most", "-1window", "5'mostTICE", "mean", "min", "max", "90%", "10%", "75%", "25%", "%<=20%", "sum<=5%", "sum<=10%", "sum<=20%", "%>90%", "%>80%", "sum>90%", "sum>80%", "whole")

			## save the folding energy features to an RDS file
			if (species_tissue!="") {
				saveRDS(UTR_folding_energy_features, file=paste0("processed_data_w_polyA/folding_energy_features/", species_short, "_", species_tissue, "_UTR_folding_energy_features_", window_len, ".rds"))
			} else {
				saveRDS(UTR_folding_energy_features, file=paste0("processed_data_w_polyA/folding_energy_features/", species_short, "_UTR_folding_energy_features_", window_len, ".rds"))
			}

			## store the features for possible use in the next iteration
			UTR_folding_energy_features_old <- UTR_folding_energy_features
		}
	}
}

## specially create a second version of the features for "spom": don't include any folding energy windows that extend into 3' of +30
for (species_short in c("sp")) {
	for (species_tissue in species_tissue_list[[species_short]]) {
		if (species_tissue!="") {
			species_genes <- get(paste(species_short, species_tissue, "genes", sep="."))			
		} else {
			species_genes <- get(paste(species_short, "genes", sep="."))
		}
		## read in the whole UTR folding energy feature (one value per gene)
		if (species_tissue!="") {
			whole_5UTR_folding_energy <- readRDS(file=paste0("processed_data_w_polyA/whole_UTR_folding_energy_feature/", species_short, "_", species_tissue, "_whole_UTR_folding_energy_feature.rds"))
		} else {
			whole_5UTR_folding_energy <- readRDS(file=paste0("processed_data_w_polyA/whole_UTR_folding_energy_feature/", species_short, "_whole_UTR_folding_energy_features.rds"))
		}
		
		if (species_short != "hs") {
			uTICE_len <- species_uTICE_len_list[[species_short]]
		} else {
			uTICE_len <- species_uTICE_len_list[[species_short]][[species_tissue]]
		}
		
		# generate folding energy features
		for (window_len in window_lens) {
			if (species_tissue!="") {
				UTR_folding_energy_all <- readRDS(file=paste0("processed_data_w_polyA/var_win_fold_UTR5_CDSshort_all/", species_short, "_", species_tissue, "_5UTR_CDS_folding_energy_", window_len, ".rds"))
			} else {
				UTR_folding_energy_all <- readRDS(file=paste0("processed_data_w_polyA/var_win_fold_UTR5_CDSshort_all/", species_short, "_5UTR_CDS_folding_energy_", window_len, ".rds"))
			}
			UTR_folding_energy_all <- UTR_folding_energy_all$UTR_folding_energy_all[species_genes]
			# remove the long windows that extend beyond +30 before calculating the percentiles
			
			trun_pos <- as.numeric(window_len) - 30 + 1 # the position (-trun_pos) relative to AUG; windows whose starting positions to the 3' of trun_pos should be removed
			UTR_folding_energy_all_trun <- lapply(UTR_folding_energy_all, FUN=function(x) {
				if (trun_pos > length(x)) {
					x <- NULL
				} else if (trun_pos > 0) {
					x <- rev(rev(x)[trun_pos:length(x)])
				}
				return(x)
			})
			UTR_folding_energy_percentiles <- quantile(unlist(UTR_folding_energy_all_trun), probs=c(0.05,0.1,0.2,0.8,0.9))
			
			UTR_folding_energy_features <- t( sapply(1:length(UTR_folding_energy_all_trun), FUN=function(i) {
			  x <- UTR_folding_energy_all_trun[[i]]
			  x_orig <- UTR_folding_energy_all[[i]]
				if (is.null(x)) {
				  return(rep(NA, 18))
				} else {
			  	  x1 <- x[1]
				  if (trun_pos > 1) {
				  	x2 <- NA
				  } else {
				  	x2 <- rev(x_orig)[1]
				  }
				  if (trun_pos > uTICE_len) {
				  	x3 <- NA 
				  } else {
				  	x3 <- rev(x_orig)[uTICE_len]
				  }
				  x4 <- mean(x)
				  x5 <- min(x)
				  x6 <- max(x)
				  tmp <- quantile(x, probs=c(0.9, 0.1, 0.75, 0.25))
				  n <- length(x)
				  x7 <- tmp[1]
				  x8 <- tmp[2]
				  x9 <- tmp[3]
				  x10 <- tmp[4]
				  x11 <- sum(x <= UTR_folding_energy_percentiles[3]) / n
				  x16 <- sum(x > UTR_folding_energy_percentiles[5]) / n
				  x17 <- sum(x > UTR_folding_energy_percentiles[4]) / n
				  x13 <- sum(x[x <= UTR_folding_energy_percentiles[1]])
				  x14 <- sum(x[x <= UTR_folding_energy_percentiles[2]])
				  x15 <- sum(x[x <= UTR_folding_energy_percentiles[3]])
				  x18 <- sum(x[x > UTR_folding_energy_percentiles[5]])
				  x19 <- sum(x[x > UTR_folding_energy_percentiles[4]])
				  return( c(x1, x2, x3, x4, x5, x6, x7, x8, x9, x10, x11, x13, x14, x15, x16, x17, x18, x19) )
				}
			}) )
			nonNA_idx <- apply(UTR_folding_energy_features, 2, FUN=function(y) !all(is.na(y)))
			UTR_folding_energy_features <- cbind(UTR_folding_energy_features[,nonNA_idx], whole_5UTR_folding_energy)

			# deal with the missing data
			if (any(is.na(UTR_folding_energy_features[,1]))) {
				missing_idx <- which(is.na(UTR_folding_energy_features[,1]))
				for (i in missing_idx) {
					for (j in 1:(ncol(UTR_folding_energy_features)-1)) {
						quantile_ij <- sum(UTR_folding_energy_features_old[,j] <= UTR_folding_energy_features_old[i,j]) / nrow(UTR_folding_energy_features)
						UTR_folding_energy_features[i,j] <- quantile(UTR_folding_energy_features[,j], probs=quantile_ij, na.rm=T)
					}
				}
			}

			colnames(UTR_folding_energy_features) <- c(c("5'most", "-1window", "5'mostTICE", "mean", "min", "max", "90%", "10%", "75%", "25%", "%<=20%", "sum<=5%", "sum<=10%", "sum<=20%", "%>90%", "%>80%", "sum>90%", "sum>80%")[nonNA_idx], "whole")

			## save the folding energy features to an RDS file
			if (species_tissue!="") {
				saveRDS(UTR_folding_energy_features, file=paste0("processed_data_w_polyA/folding_energy_features/", species_short, "_", species_tissue, "_UTR_folding_energy_features_", window_len, "_trun.rds"))
			} else {
				saveRDS(UTR_folding_energy_features, file=paste0("processed_data_w_polyA/folding_energy_features/", species_short, "_UTR_folding_energy_features_", window_len, "_trun.rds"))
			}

			## store the features for possible use in the next iteration
			UTR_folding_energy_features_old <- UTR_folding_energy_features
		}
	}
}

# select folding energy features for every species
## function
fold_energy_feature_select <- function(species_prefix, tissue_prefix, len_breaks, ifexemplar, selected_feature_names=NULL, iftrun=F) {
	# species_prefix = "sc", "mm", "at", "sp", "dm", "hs"
	# tissue_prefix = "" ("sc"), c("nih3t3", "liver", "kidney") ("mm"), c("leaf", "root", "shoot") ("at"), c("", "alt") ("sp"), c("0to1hr", "3to4hr") ("dm"), c("hela", "hek293t") ("hs")
	# len_breaks = c(20, 35) ("sc"), NULL ("mm"), 65 ("at"), ...
	# ifexemplar = TRUE/FLASE; if TRUE, perform feature selection; otherwise, use selected_feature_names
	# selected_feature_names = a character vector of the selected feature names from the exemplar tissue
	
	## combine folding energy features from previous calculation in "040418_check_folding_energy_window_length_effect(add_a_feature_to_arab).R"
	fold_energy_features <- NULL
	cors <- NULL
	if (tissue_prefix != "") {
		UTR_seqs <- get(paste(species_prefix, tissue_prefix, "UTR_seqs", sep="."))
		log10.TR <- get(paste(species_prefix, tissue_prefix, "log10.TR", sep="."))
	} else {
		UTR_seqs <- get(paste(species_prefix, "UTR_seqs", sep="."))
		log10.TR <- get(paste(species_prefix, "log10.TR", sep="."))
	}	
	log10.UTR_lens <- log(nchar(UTR_seqs), 10)	
	n <- length(log10.UTR_lens)
	
	for (window_len in window_lens) {
		if (!iftrun) {
		  if (tissue_prefix != "") {
			tmp_features <- readRDS(paste0("processed_data_w_polyA/folding_energy_features/", species_prefix, "_", tissue_prefix, "_UTR_folding_energy_features_", window_len, ".rds"))
		  } else {
			tmp_features <- readRDS(paste0("processed_data_w_polyA/folding_energy_features/", species_prefix, "_UTR_folding_energy_features_", window_len, ".rds"))
		  }
		} else {
		  if (tissue_prefix != "") {
			tmp_features <- readRDS(paste0("processed_data_w_polyA/folding_energy_features/", species_prefix, "_", tissue_prefix, "_UTR_folding_energy_features_", window_len, "_trun.rds"))
		  } else {
			tmp_features <- readRDS(paste0("processed_data_w_polyA/folding_energy_features/", species_prefix, "_UTR_folding_energy_features_", window_len, "_trun.rds"))
		  }
		}
		if (window_len != rev(window_lens)[1]) { # handle the "whole" feature, which doesn't need a window length
			tmp_features <- tmp_features[,-ncol(tmp_features)]
			tmp_feature_names <- paste(colnames(tmp_features), window_len, sep="_")
		} else {
			tmp_feature_names <- paste(colnames(tmp_features)[-ncol(tmp_features)], window_len, sep="_")
			tmp_feature_names <- c(tmp_feature_names, colnames(tmp_features)[ncol(tmp_features)])
		}
		tmp_cors <- apply(tmp_features, 2, FUN=function(x) cor(log10.TR, x, use="complete.obs"))
		if (length(len_breaks) > 0) {
			tmp_features_by_length <- NULL		
			tmp_cors_by_group <- NULL # Pearson correlation between log10.TR and every feature within a group	
			num_groups <- length(len_breaks) + 1 # number of gene groups divided by UTR lengths
			for (i in 1:num_groups) {
				if (i == 1) {
					tmp_features_one_group <- as.numeric(log10.UTR_lens<log(len_breaks[1],10)) * tmp_features
					colnames(tmp_features_one_group) <- paste(tmp_feature_names, paste0("<",len_breaks[1]), sep="_")
					idx_one_group <- which(log10.UTR_lens<log(len_breaks[1],10))
				} else if (i == num_groups) {
					tmp_features_one_group <- as.numeric(log10.UTR_lens>=log(len_breaks[num_groups-1],10)) * cbind(tmp_features, rep(1,n))
					colnames(tmp_features_one_group) <- paste(c(tmp_feature_names,"Intercept"), paste0(">=",len_breaks[num_groups-1]), sep="_")
					idx_one_group <- which(log10.UTR_lens>=log(len_breaks[num_groups-1],10))
				} else {
					tmp_features_one_group <- as.numeric(log10.UTR_lens>=log(len_breaks[i-1],10) & log10.UTR_lens<log(len_breaks[i],10)) * cbind(tmp_features, rep(1,n))
					colnames(tmp_features_one_group) <- paste(c(tmp_feature_names,"Intercept"), paste0(">=",len_breaks[i-1],"&<",len_breaks[i]), sep="_")
					idx_one_group <- which(log10.UTR_lens>=log(len_breaks[i-1],10) & log10.UTR_lens<log(len_breaks[i],10))
				}
				tmp_features_by_length <- cbind(tmp_features_by_length, tmp_features_one_group)
				tmp_cors_one_group <- apply(tmp_features_one_group, 2, FUN=function(x) cor(log10.TR[idx_one_group], x[idx_one_group], use="complete.obs"))
				tmp_cors_by_group <- c(tmp_cors_by_group, tmp_cors_one_group)
			}
			tmp_features <- tmp_features_by_length
			tmp_cors <- tmp_cors_by_group			
		} else {
			colnames(tmp_features) <- tmp_feature_names
		}		
		fold_energy_features <- cbind(fold_energy_features, tmp_features)
		cors <- c(cors, tmp_cors)
	}
	
	## remove duplicated column names
	keep_idx <- !duplicated(colnames(fold_energy_features))
	fold_energy_features <- fold_energy_features[,keep_idx]
	cors <- cors[keep_idx]
	
	fold_energy_features_all <- as.data.frame(cbind(log10.TR, fold_energy_features))
	if (ifexemplar) {
		## use forward selection + Bayesian Information Criterion (BIC) to select features		
		reg_fwd <- regsubsets(log10.TR ~. , data=fold_energy_features_all, nvmax=70, method ="forward")
		reg_summary <- summary(reg_fwd)
		idx_bic <- which.min(reg_summary$bic)
		selected_feature_idx <- which(reg_summary$outmat[idx_bic,]=="*")
	} else {
		## use "selected_feature_names"
		selected_feature_idx <- which(colnames(fold_energy_features) %in% selected_feature_names)
	}
		
	selected_feature_coef <- summary(lm(log10.TR ~. , data=fold_energy_features_all[,c(1,selected_feature_idx+1)]))$coefficients[-1,]
	selected_feature_coef <- cbind(selected_feature_coef, cors[selected_feature_idx])
	colnames(selected_feature_coef)[5] <- "Pearson Cor vs TR"
	## order the selected features by the p-values of their coefficients in a multiple linear model
	selected_feature_summary <- selected_feature_coef[order(selected_feature_coef[,4]),]
	
	## selected features
	fold_energy_features_selected <- fold_energy_features_all[,c(1,selected_feature_idx+1)]
	## R2 of the selected features
	selected_feature_summary_R2 <- summary(lm(log10.TR ~. , data=fold_energy_features_selected))$r.squared
	
	return(list(selected_features=fold_energy_features_selected, selected_feature_summary=selected_feature_summary, selected_feature_R2=selected_feature_summary_R2, all_features=fold_energy_features_all))
}

for (species_prefix in names(species_tissue_list)) {
	len_breaks <- species_len_break_list[[species_prefix]]
	for (i in 1:length(species_tissue_list[[species_prefix]])) {
		tissue_prefix <- species_tissue_list[[species_prefix]][i]
		fold_energy_features_selected <- fold_energy_feature_select(species_prefix, tissue_prefix, len_breaks, ifexemplar=T)
		if (tissue_prefix != "") {
			saveRDS(fold_energy_features_selected, file=paste0("processed_data_w_polyA/folding_energy_features_selected/", species_prefix, "_", tissue_prefix, "_UTR_folding_energy_features_selected.rds"))
		} else {
			saveRDS(fold_energy_features_selected, file=paste0("processed_data_w_polyA/folding_energy_features_selected/", species_prefix, "_UTR_folding_energy_features_selected.rds"))
		}
		# deal with the truncated features of sp
		if (species_prefix == "sp") {
		  fold_energy_features_selected <- fold_energy_feature_select(species_prefix, tissue_prefix, len_breaks, ifexemplar=T, iftrun=T)
		  if (tissue_prefix != "") {
			  saveRDS(fold_energy_features_selected, file=paste0("processed_data_w_polyA/folding_energy_features_selected/", species_prefix, "_", tissue_prefix, "_UTR_folding_energy_features_selected_trun.rds"))
		  } else {
			  saveRDS(fold_energy_features_selected, file=paste0("processed_data_w_polyA/folding_energy_features_selected/", species_prefix, "_UTR_folding_energy_features_selected_trun.rds"))
		  }
		}
	}
}
```

# Construct features for the full model
## Settings
```{r}
##################### species-independent information #####################
# all the 4 nucleotides
nts <- c("A", "T", "C", "G")
# all the 16 di-nucleotides
di_nts <- as.character(sapply(nts, FUN=function(nt1) sapply(nts, FUN=function(nt2) paste0(nt1,nt2)))) 
# all the 64 tri-nucleotides
tri_nts <- as.character(sapply(nts, FUN=function(nt1) sapply(nts, FUN=function(nt2) sapply(nts, FUN=function(nt3) paste0(nt1,nt2,nt3)))))
## translating the 64 tri nucleotides to amino acides
tri_nts_to_aas <- sapply(tri_nts, FUN=function(nuc) {
    as.character( translate( DNAString( nuc ) ) )
})
## 20 amino acids
aas <- unique(tri_nts_to_aas)
aas <- aas[aas!="*"]
## nonstopping 61 codon frequencies (trinucleotide in the open reading frame)
nonstop_codons <- tri_nts[tri_nts_to_aas!="*"]

# a function to construct the PWM based on a set of 5' UTR sequences (right aligned at the TSS)
UTR_pwm <- function(UTR_seqs, n_left) {
	# n_left: -n_left:-1 is the region to construct the PWM
    tmp_seq <- t( sapply(UTR_seqs, FUN=function(x) {
        tmp <- strsplit(x, "")[[1]]
        rev(tmp)[1:n_left]
    }) )
    count_A <- colSums( tmp_seq == "A", na.rm=T )
    count_T <- colSums( tmp_seq == "T", na.rm=T )
    count_C <- colSums( tmp_seq == "C", na.rm=T )
    count_G <- colSums( tmp_seq == "G", na.rm=T )
    count_tot <- count_A + count_T + count_C + count_G
    freq <- cbind( count_A / count_tot, count_T / count_tot, count_C/ count_tot, count_G / count_tot )
    colnames(freq) <- c("A", "T", "C", "G")
    rownames(freq) <- (-1):(-n_left)
    return(freq[n_left:1,])
}

# a function to calculate 5' UTR PWM scores for a set of genes (aligned at TSS)
UTR_pwm_score <- function(PWM, UTR_seqs) {
    len <- nrow(PWM)
    PWM <- PWM[len:1,] # reverse the order, so -1 is the first row
    sapply(UTR_seqs, FUN=function(x) {
        tmp <- rev( strsplit(x, "")[[1]] ) # reverse the sequence, so -1 is the first position
        min_len <- min( c(len, length(tmp)) )
        log( prod( sapply( 1:min_len, FUN=function(i) {
            PWM[i, tmp[i]]
        }) ), 10)
    })
}

# a function to construct the PWM based on a set of CDS sequences truncated in the +4:+n_right region
CDS_pwm <- function(CDS_seqs_n_right, n_right) {
    tmp_seq <- t( sapply(CDS_seqs_n_right, FUN=function(x) {
        tmp <- strsplit(x, "")[[1]]
    }) )
	freq <- t( sapply(1:ncol(tmp_seq), FUN=function(i) {
        tmp_count <- c( sum( tmp_seq[,i]=="A" ),
        sum( tmp_seq[,i]=="T" ),
        sum( tmp_seq[,i]=="C" ),
        sum( tmp_seq[,i]=="G" )
        )
        return( tmp_count / sum(tmp_count) )
    }) )
    colnames(freq) <- c("A", "T", "C", "G")
    rownames(freq) <- 4:n_right
    return(freq)
}

# a function to calculate CDS PWM scores for a set of genes truncated in the +4:+n_right region
CDS_pwm_score <- function(PWM_n_right, CDS_seqs_n_right) {
	UTR_pwm_score(PWM_n_right, CDS_seqs_n_right)
}

# a function to construct the PWM based on a set of 5' UTR sequences and ORF sequences of the same genes (aligned at TSS)
UTR_CDS_pwm <- function(UTR_seqs, CDS_seqs, n_UTR, n_CDS) {
    tmp_seq <- t( sapply(UTR_seqs, FUN=function(x) {
        tmp <- strsplit(x, "")[[1]]
        rev(tmp)[1:n_UTR]
    }) )
    count_A <- colSums( tmp_seq == "A", na.rm=T )
    count_T <- colSums( tmp_seq == "T", na.rm=T )
    count_C <- colSums( tmp_seq == "C", na.rm=T )
    count_G <- colSums( tmp_seq == "G", na.rm=T )
    count_tot <- count_A + count_T + count_C + count_G
    freq_UTR <- cbind( count_A / count_tot, count_T / count_tot, count_C/ count_tot, count_G / count_tot )

    tmp_seq <- t( sapply(CDS_seqs, FUN=function(x) {
        tmp <- strsplit(x, "")[[1]]
        rev(tmp[4:n_CDS]) # do not include +1 to +3 (often ATG)
    }) )
    if (nrow(tmp_seq)==1) tmp_seq <- t(tmp_seq)
    count_A <- colSums( tmp_seq == "A", na.rm=T )
    count_T <- colSums( tmp_seq == "T", na.rm=T )
    count_C <- colSums( tmp_seq == "C", na.rm=T )
    count_G <- colSums( tmp_seq == "G", na.rm=T )
    count_tot <- count_A + count_T + count_C + count_G
    freq_CDS <- cbind( count_A / count_tot, count_T / count_tot, count_C/ count_tot, count_G / count_tot )

    freq <- rbind(freq_CDS, freq_UTR)

    colnames(freq) <- c("A", "U", "C", "G")
    rownames(freq) <- c( (n_CDS:4), ((-1):(-n_UTR)) ) # do not include +1 to +3 (often ATG)
    return(freq[nrow(freq):1, c("A", "C", "G", "U")])
}

# a function to construct the 5ofTICE PWM based on a set of 5' UTR sequences in the region of length n_5ofTICE (left aligned at the 5' end)
UTR_pwm_5_aligned <- function(UTR_seqs, n_5ofTICE) {
    tmp_seq <- t( sapply(UTR_seqs, FUN=function(x) {
        tmp <- strsplit(x, "")[[1]]
        tmp[1:n_5ofTICE]
    }) )
	if (n_5ofTICE==1) {
		tmp_seq <- t(tmp_seq)
	}
    count_A <- colSums( tmp_seq == "A", na.rm=T )
    count_T <- colSums( tmp_seq == "T", na.rm=T )
    count_C <- colSums( tmp_seq == "C", na.rm=T )
    count_G <- colSums( tmp_seq == "G", na.rm=T )
    count_tot <- count_A + count_T + count_C + count_G
    freq <- cbind( count_A / count_tot, count_T / count_tot, count_C/ count_tot, count_G / count_tot )
    colnames(freq) <- c("A", "T", "C", "G")
    rownames(freq) <- 1:n_5ofTICE
    return(freq)
}

# a function to calculate 5ofTICE PWM scores for a set of genes (left aligned at the 5' end)
UTR_pwm_5_aligned_score <- function(PWM, UTR_seqs) {
	# 5' end is the first row
    len <- nrow(PWM)
    sapply(UTR_seqs, FUN=function(x) {
        tmp <- strsplit(x, "")[[1]]
        min_len <- min( c(len, length(tmp)) )
        log( prod( sapply( 1:min_len, FUN=function(i) {
            PWM[i, tmp[i]]
        }) ), 10)
    })
}

##################### parameters for each species #####################
# S. cerevisae
sc.n_left <- 35 # the minimum UTR length of "long" genes 
sc.n_right <- 28 # the CDS PWM length
sc.TR_top_pctl <- 0.9 # the percentile of TR top genes
sc.TR_bottom_pctl <- 0.1 # the percentile of TR top genes
sc.PWM_lens <- c(5, 20, 35) # lengths of 5' UTR PWMs
sc.len_breaks <- c(20, 35) # UTR length breaks to divide genes into groups
sc.n_5ofTICE <- 5 # the 5ofTICE PWM length (from the 5' end)

# S. pombe
sp.n_left <- 5 # the minimum UTR length of "long" genes 
sp.n_right <- 13 # the CDS PWM length
sp.TR_top_pctl <- 0.9 # the percentile of TR top genes
sp.TR_bottom_pctl <- 0.1 # the percentile of TR top genes
sp.PWM_lens <- 5 # lengths of 5' UTR PWMs
sp.len_breaks <- NULL # UTR length breaks to divide genes into groups
sp.n_5ofTICE <- 5 # the 5ofTICE PWM length (from the 5' end)

# Arabidopsis
at.n_left <- 65 # the minimum UTR length of "long" genes 
at.n_right <- 33 # the CDS PWM length
at.TR_top_pctl <- 0.9 # the percentile of TR top genes
at.TR_bottom_pctl <- 0.1 # the percentile of TR top genes
at.PWM_lens <- c(5, 65) # lengths of 5' UTR PWMs
at.len_breaks <- c(65) # UTR length breaks to divide genes into groups
at.n_5ofTICE <- 5 # the 5ofTICE PWM length (from the 5' end)

# Mouse
mm.n_left <- 6 # the minimum UTR length of "long" genes 
mm.n_right <- 13 # the CDS PWM length
mm.TR_top_pctl <- 0.9 # the percentile of TR top genes
mm.TR_bottom_pctl <- 0.1 # the percentile of TR top genes
mm.PWM_lens <- c(6) # lengths of 5' UTR PWMs
mm.len_breaks <- NULL # UTR length breaks to divide genes into groups
mm.n_5ofTICE <- 5 # the 5ofTICE PWM length (from the 5' end)

# Human hela 2
hs.hela2.n_left <- 6 # the minimum UTR length of "long" genes 
hs.hela2.n_right <- 7 # the CDS PWM length
hs.hela2.TR_top_pctl <- 0.9 # the percentile of TR top genes
hs.hela2.TR_bottom_pctl <- 0.1 # the percentile of TR top genes
hs.hela2.PWM_lens <- 6 # lengths of 5' UTR PWMs
hs.hela2.len_breaks <- NULL # UTR length breaks to divide genes into groups
hs.hela2.n_5ofTICE <- 5 # the 5ofTICE PWM length (from the 5' end)
```
## Read in the already generated features & find the TR top and bottom genes
```{r}
for (species in names(species_tissue_list)) {
	for (species_tissue in species_tissue_list[[species]]) {
		if (species_tissue!="") {
			tissue_dot <- paste0(".", species_tissue)
			tissue_us <- paste0("_", species_tissue)
		} else {
			tissue_dot <- ""
			tissue_us <- ""
		}
		## 5' UTR log10 lengths (a feature)
		assign(x=paste0(species, tissue_dot, ".log10.UTR_lens"),
			value=log(nchar(get(paste0(species, tissue_dot, ".UTR_seqs"))[get(paste0(species, tissue_dot, ".genes"))]), 10)
		)
		## CDS log10 lengths (a feature)
		assign(x=paste0(species, tissue_dot, ".log10.CDS_lens"),
			value=log(nchar(get(paste0(species, tissue_dot, ".CDS_seqs"))[get(paste0(species, tissue_dot, ".genes"))]), 10)
		)
		## uAUG counts (a feature)
		assign(x=paste0(species, tissue_dot, ".uAUG_counts"),
			value=stri_count_regex(get(paste0(species, tissue_dot, ".UTR_seqs")), '(?=ATG)')
		)
		## selected folding energy features (a mega-feature)
		if (species != "sp") {
		  assign(x=paste0(species, tissue_dot, ".fold_energy_features_selected"),
			value=readRDS(file=paste0("processed_data_w_polyA/folding_energy_features_selected/", species, tissue_us, "_UTR_folding_energy_features_selected.rds"))[[1]][,-1]
		  ) ## the first column is log10.TR
		} else {
		  assign(x=paste0(species, tissue_dot, ".fold_energy_features_selected"),
			value=readRDS(file=paste0("processed_data_w_polyA/folding_energy_features_selected/", species, tissue_us, "_UTR_folding_energy_features_selected_trun.rds"))[[1]][,-1]
		  ) ## the first column is log10.TR
		}
		## all folding energy features (multiple individual features)
		assign(x=paste0(species, tissue_dot, ".fold_energy_features_all"),
			value=readRDS(file=paste0("processed_data_w_polyA/folding_energy_features_selected/", species, tissue_us, "_UTR_folding_energy_features_selected.rds"))$all_features[,-1]
		) ## the first column is log10.TR
		## whole CDS folding energy (a feature)
		assign(x=paste0(species, tissue_dot, ".CDS_fold_energy"),
			value=readRDS(file=paste0("processed_data_w_polyA/whole_CDS_folding_energy_feature/", species, tissue_us, "_whole_CDS_folding_energy_feature.rds"))
		)
		## TR top and bottom genes
		if (species != "hs") {
		  TR_quantiles <- quantile(get(paste0(species, tissue_dot, ".log10.TR")), prob=c(get(paste0(species, ".TR_bottom_pctl")), get(paste0(species, ".TR_top_pctl"))))
		} else {
		  TR_quantiles <- quantile(get(paste0(species, tissue_dot, ".log10.TR")), prob=c(get(paste0(species, tissue_dot, ".TR_bottom_pctl")), get(paste0(species, tissue_dot, ".TR_top_pctl"))))
		}
		assign(x=paste0(species, tissue_dot, ".TR_bottom_genes"),
			value=get(paste0(species, tissue_dot, ".genes"))[get(paste0(species, tissue_dot, ".log10.TR")) <= TR_quantiles[1]]
		)
		assign(x=paste0(species, tissue_dot, ".TR_top_genes"),
			value=get(paste0(species, tissue_dot, ".genes"))[get(paste0(species, tissue_dot, ".log10.TR")) >= TR_quantiles[2]]
		)
	}
}
```
## The following code only needs to be run once
```{r, eval = FALSE}
#####
# uTICE features (a mega-feature)
make_uTICE_features <- function(UTR_seqs, log10.TR, TR_top_pctl, n_left, PWM_lens, len_breaks, ifexemplar, selected_feature_names=NULL) {
	# use forward + BIC to select features among the following features
	#(1) 16 dinucleotide frequencies in the -n_left to -1 region
	#(2) 64 trinucleotide frequencies in the -n_left to -1 region
	#(3) PWM scores based on the PWM constructed from the top 10% TR genes’ sequences
	
	#UTR_seqs: a character vector of UTR sequences, with its names as gene names
	#log10.TR: a numeric vector of log10.TR values, with its names as gene names
	#TR_top_pctl: the percentile to define TR top genes, e.g., 0.9
	#n_left: the minimum UTR length of "long" genes
	#PWM_lens: a numeric vector containing the possible PWM lengths
	#len_breaks: a numeric vector containing the UTR length breaks, which divide genes into groups
	#ifexemplar = TRUE/FLASE; if TRUE, perform feature selection; otherwise, use selected_feature_names
	#selected_feature_names = a character vector of the selected feature names from the exemplar tissue
	
	n <- length(UTR_seqs)
	UTR_lens <- nchar(UTR_seqs)
	TR_top_UTR_seqs <- UTR_seqs[log10.TR >= quantile(log10.TR, prob=TR_top_pctl)]
	
	# cut each gene's 5' UTR sequence to the -n_left:-1 region
	UTR_seqs_n_left <- substr( x=UTR_seqs, start=nchar(UTR_seqs)-n_left+1, stop=nchar(UTR_seqs) )
	TR_top_UTR_seqs_n_left <- substr( x=TR_top_UTR_seqs, start=nchar(TR_top_UTR_seqs)-n_left+1, stop=nchar(TR_top_UTR_seqs) )	
	
	# count the dinucleotide frequencies in the -n_left:-1 region of each gene
	UTR_di_counts_n_left <- sapply(di_nts, FUN=function(nuc) {
	    stri_count_regex(UTR_seqs_n_left, paste0('(?=',nuc,')'))
	})
	rownames(UTR_di_counts_n_left) <- names(UTR_seqs_n_left)
	UTR_di_freqs_n_left <- UTR_di_counts_n_left / rowSums(UTR_di_counts_n_left)
	UTR_di_freqs_n_left[is.na(UTR_di_freqs_n_left)] <- 0
	
	# count the trinucleotide frequencies in the -n_left:-1 region of each gene
	UTR_tri_counts_n_left <- sapply(tri_nts, FUN=function(nuc) {
	    stri_count_regex(UTR_seqs_n_left, paste0('(?=',nuc,')'))
	})
	rownames(UTR_tri_counts_n_left) <- names(UTR_seqs_n_left)
	UTR_tri_freqs_n_left <- UTR_tri_counts_n_left / rowSums(UTR_tri_counts_n_left)
	UTR_tri_freqs_n_left[is.na(UTR_tri_freqs_n_left)] <- 0
	
	# calculate PWM scores for multiple PWM lengths for each gene
	PWM_UTR_scores <- sapply(PWM_lens, FUN=function(len) {
		TR_top_PWM_UTR <- UTR_pwm( TR_top_UTR_seqs_n_left, len )
		UTR_pwm_score( TR_top_PWM_UTR, UTR_seqs_n_left )
	})
	
	# combine the features into a design matrix
	features <- cbind(UTR_di_freqs_n_left, UTR_tri_freqs_n_left)
	feature_names <- colnames(features)
	if (length(len_breaks) > 0) {
		features_by_group <- NULL
		cors_by_group <- NULL # Pearson correlation between log10.TR and every feature within a group
		num_groups <- length(len_breaks) + 1 # number of gene groups divided by UTR lengths
		for (i in 1:num_groups) {
			if (i == 1) {
				features_one_group <- as.numeric(UTR_lens<len_breaks[1]) * cbind(features, PWM_UTR_scores[,1])
				colnames(features_one_group) <- c( paste(feature_names, paste0("<",len_breaks[1]), sep="_"), paste0("PWM", PWM_lens[1]) )
				idx_one_group <- which(UTR_lens<len_breaks[1])								
			} else if (i == num_groups) {
				features_one_group <- as.numeric(UTR_lens>=len_breaks[num_groups-1]) * cbind(features, PWM_UTR_scores[,num_groups], rep(1, n))
				colnames(features_one_group) <- c( paste(feature_names, paste0(">=",len_breaks[num_groups-1]), sep="_"), paste0("PWM", PWM_lens[num_groups]), paste0("Intercept_>=",len_breaks[num_groups-1]) )
				idx_one_group <- which(UTR_lens>=len_breaks[num_groups-1])	
			} else {
				features_one_group <- as.numeric(UTR_lens>=len_breaks[i-1] & UTR_lens<len_breaks[i]) * cbind(features, PWM_UTR_scores[,i], rep(1, n))
				colnames(features_one_group) <- c( paste(feature_names, paste0(">=",len_breaks[i-1],"&<",len_breaks[i]), sep="_"), paste0("PWM", PWM_lens[i]), paste0("Intercept_>=",len_breaks[i-1],"&<",len_breaks[i]) )
				idx_one_group <- which(UTR_lens>=len_breaks[i-1] & UTR_lens<len_breaks[i])	
			}
			features_by_group <- cbind(features_by_group, features_one_group)
			cors_one_group <- apply(features_one_group, 2, FUN=function(x) cor(log10.TR[idx_one_group], x[idx_one_group], use="complete.obs"))
			cors_by_group <- c(cors_by_group, cors_one_group)		
		}
		features <- features_by_group
		cors <- cors_by_group
	} else {
		features <- cbind(features, PWM_UTR_scores)
		colnames(features) <- c(feature_names, paste0("PWM", PWM_lens))
		cors <- apply(features, 2, FUN=function(x) cor(log10.TR, x, use="complete.obs"))
	}
	
	# combine the response and the design matrix into a data frame
	PWM_idx <- which(grepl("PWM", colnames(features)))
	data_frame <- cbind.data.frame(log10.TR=log10.TR, features[,PWM_idx], features[,-PWM_idx])
	colnames(data_frame) <- c("log10.TR", colnames(features)[PWM_idx], colnames(features)[-PWM_idx])
	cors <- c(cors[PWM_idx], cors[-PWM_idx])
	
	# R2 with the PWM scores and intercepts only
	R2_PWM <- summary(lm(log10.TR ~ features[,which(grepl("PWM", colnames(features)) | grepl("Intercept", colnames(features)))]))$r.squared
	
	if (ifexemplar) {
		# use forward selection + Bayesian Information Criterion (BIC) to select features
		regfit.fwd <- regsubsets(log10.TR ~. , data=data_frame, nvmax=70, force.in=1:length(PWM_idx), method ="forward")
	
		reg.summary <- summary(regfit.fwd)
		idx_bic <- which.min(reg.summary$bic )
		features_selected_idx <- which(reg.summary$outmat[idx_bic,]=="*")
	} else {
		features_selected_idx <- which(colnames(data_frame) %in% selected_feature_names) - 1
	}	
	
	lm_selected <- summary(lm(log10.TR ~. , data=data_frame[,c(1,features_selected_idx+1)]))
	coefs_selected <- lm_selected$coefficients[-1,]
	R2_selected <- lm_selected$r.squared
	coefs_selected <- cbind(coefs_selected, cors[features_selected_idx])
	colnames(coefs_selected)[5] <- "Pearson Cor vs TR"
	## order the selected features by the p-values of their coefficients in a multiple linear model
	coefs_selected <- coefs_selected[order(coefs_selected[,4]),]
	
	uTICE_features <- data_frame[,features_selected_idx+1]
	return(list(uTICE_features=uTICE_features, uTICE_feature_coefs=coefs_selected, R2=R2_selected, R2_PWM=R2_PWM))
}

# generate uTICE features
for (species in names(species_tissue_list)) {
	for (i in 1:length(species_tissue_list[[species]])) {
		species_tissue <- species_tissue_list[[species]][i]
		if (species_tissue!="") {
			tissue_dot <- paste0(".", species_tissue)
			tissue_us <- paste0("_", species_tissue)
		} else {
			tissue_dot <- ""
			tissue_us <- ""
		}
		if (species != "hs") {
			assign(x=paste0(species, tissue_dot, ".uTICE_features"),
				value=make_uTICE_features(get(paste0(species, tissue_dot, ".UTR_seqs")), get(paste0(species, tissue_dot, ".log10.TR")), get(paste0(species, ".TR_top_pctl")), get(paste0(species, ".n_left")), get(paste0(species, ".PWM_lens")), get(paste0(species, ".len_breaks")), ifexemplar=T)
			)
		} else {
			assign(x=paste0(species, tissue_dot, ".uTICE_features"),
				value=make_uTICE_features(get(paste0(species, tissue_dot, ".UTR_seqs")), get(paste0(species, tissue_dot, ".log10.TR")), get(paste0(species, tissue_dot, ".TR_top_pctl")), get(paste0(species, tissue_dot, ".n_left")), get(paste0(species, tissue_dot, ".PWM_lens")), get(paste0(species, tissue_dot, ".len_breaks")), ifexemplar=T)
			)
		}
		saveRDS(get(paste0(species, tissue_dot, ".uTICE_features")), file=paste0("processed_data_w_polyA/uTICE_features/", species, tissue_dot, ".uTICE_features.rds"))	
	}
}

#####
# dTICE features (a mega-feature)
make_dTICE_features <- function(CDS_seqs, UTR_seqs, log10.TR, TR_top_pctl, n_right, len_breaks, ifexemplar, selected_feature_names=NULL) {
	# use forward + BIC to select features among the following features
	#(1) 16 dinucleotide frequencies in the +4 to +n_right region
	#(2) 64 trinucleotide frequencies in the +4 to +n_right region
	#(3) PWM scores based on the PWM constructed from the top 10% TR genes’ sequences
	
	#CDS_seqs: a character vector of CDS sequences, with its names as gene names
	#UTR_seqs: a character vector of UTR sequences, with its names as gene names
	#log10.TR: a numeric vector of log10.TR values, with its names as gene names
	#TR_top_pctl: the percentile to define TR top genes, e.g., 0.9
	#n_right: the CDS PWM length
	#len_breaks: a numeric vector containing the UTR length breaks, which divide genes into groups
	#ifexemplar = TRUE/FLASE; if TRUE, perform feature selection; otherwise, use selected_feature_names
	#selected_feature_names = a character vector of the selected feature names from the exemplar tissue	
	
	n <- length(UTR_seqs)
	UTR_lens <- nchar(UTR_seqs)
	TR_top_CDS_seqs <- CDS_seqs[log10.TR >= quantile(log10.TR, prob=TR_top_pctl)]
	
	# cut each gene's CDS sequence to the +4:+n_right region
	CDS_seqs_n_right <- substr( x=CDS_seqs, start=4, stop=n_right )
	TR_top_CDS_seqs_n_right <- substr( x=TR_top_CDS_seqs, start=4, stop=n_right )	
	
	# count the dinucleotide frequencies in the +4:+n_right region of each gene
	CDS_di_counts_n_right <- sapply(di_nts, FUN=function(nuc) {
	    stri_count_regex(CDS_seqs_n_right, paste0('(?=',nuc,')'))
	})
	rownames(CDS_di_counts_n_right) <- names(CDS_seqs_n_right)
	CDS_di_freqs_n_right <- CDS_di_counts_n_right / rowSums(CDS_di_counts_n_right)
	CDS_di_freqs_n_right[is.na(CDS_di_freqs_n_right)] <- 0
	
	# count the trinucleotide frequencies in the +4:+n_right region of each gene
	CDS_tri_counts_n_right <- sapply(tri_nts, FUN=function(nuc) {
	    stri_count_regex(CDS_seqs_n_right, paste0('(?=',nuc,')'))
	})
	rownames(CDS_tri_counts_n_right) <- names(CDS_seqs_n_right)
	CDS_tri_freqs_n_right <- CDS_tri_counts_n_right / rowSums(CDS_tri_counts_n_right)
	CDS_tri_freqs_n_right[is.na(CDS_tri_freqs_n_right)] <- 0
	
	# calculate PWM scores based on the TR top PWM in the +4:+n_right region for each gene
	PWM_CDS_scores <- sapply(n_right, FUN=function(len) {
		TR_top_PWM_CDS <- CDS_pwm( TR_top_CDS_seqs_n_right, n_right )
		CDS_pwm_score( TR_top_PWM_CDS, CDS_seqs_n_right )
	})
	colnames(PWM_CDS_scores) <- paste0("PWM", n_right)
	
	# combine the features into a design matrix
	features <- cbind(CDS_di_freqs_n_right, CDS_tri_freqs_n_right, PWM_CDS_scores)
	feature_names <- colnames(features)
	cors <- apply(features, 2, FUN=function(x) cor(log10.TR, x, "complete.obs"))
	if (length(len_breaks) > 0) {
		features_by_group <- NULL
		cors_by_group <- NULL # Pearson correlation between log10.TR and every feature within a group
		num_groups <- length(len_breaks) + 1 # number of gene groups divided by UTR lengths
		for (i in 1:num_groups) {
			if (i == 1) {
				features_one_group <- as.numeric(UTR_lens<len_breaks[1]) * features
				colnames(features_one_group) <- paste(feature_names, paste0("<",len_breaks[1]), sep="_")
				idx_one_group <- which(UTR_lens<len_breaks[1])
			} else if (i == num_groups) {
				features_one_group <- as.numeric(UTR_lens>=len_breaks[num_groups-1]) * cbind(features, rep(1,n))
				colnames(features_one_group) <- paste(c(feature_names,"Intercept"), paste0(">=",len_breaks[num_groups-1]), sep="_")
				idx_one_group <- which(UTR_lens>=len_breaks[num_groups-1])
			} else {
				features_one_group <- as.numeric(UTR_lens>=len_breaks[i-1] & UTR_lens<len_breaks[i]) * cbind(features, rep(1,n))
				colnames(features_one_group) <- paste(c(feature_names,"Intercept"), paste0(">=",len_breaks[i-1],"&<",len_breaks[i]), sep="_")
				idx_one_group <- which(UTR_lens>=len_breaks[i-1] & UTR_lens<len_breaks[i])
			}
			features_by_group <- cbind(features_by_group, features_one_group)
			cors_one_group <- apply(features_one_group, 2, FUN=function(x) cor(log10.TR[idx_one_group], x[idx_one_group], use="complete.obs"))
			cors_by_group <- c(cors_by_group, cors_one_group)
		}
		features <- features_by_group
		cors <- cors_by_group
	}
	
	# combine the response and the design matrix into a data frame
	PWM_idx <- which(grepl("PWM", colnames(features)))
	data_frame <- cbind.data.frame(log10.TR=log10.TR, features[,PWM_idx], features[,-PWM_idx])
	colnames(data_frame) <- c("log10.TR", colnames(features)[PWM_idx], colnames(features)[-PWM_idx])
	cors <- c(cors[PWM_idx], cors[-PWM_idx])
	
	# R2 with the PWM scores and intercepts only
	R2_PWM <- summary(lm(log10.TR ~ features[,which(grepl("PWM", colnames(features)) | grepl("Intercept", colnames(features)))]))$r.squared
	
	if (ifexemplar) {
		# use forward selection + Bayesian Information Criterion (BIC) to select features
		failure <- try( regfit.fwd <- regsubsets(log10.TR ~. , data=data_frame, nvmax=70, force.in=1:length(PWM_idx), method ="forward") )
		
		if (class(failure)=="try-error") {
			regfit.fwd <- regsubsets(log10.TR ~. , data=data_frame[,-(length(PWM_idx)+1)], nvmax=70, force.in=1:length(PWM_idx), method ="forward")
		}
	
		reg.summary <- summary(regfit.fwd)
		idx_bic <- which.min(reg.summary$bic )
		features_selected_idx <- which(reg.summary$outmat[idx_bic,]=="*")
	} else {
		features_selected_idx <- which(colnames(data_frame) %in% selected_feature_names) - 1
	}
	
	lm_selected <- summary(lm(log10.TR ~. , data=data_frame[,c(1,features_selected_idx+1)]))
	coefs_selected <- lm_selected$coefficients[-1,]
	R2_selected <- lm_selected$r.squared
	coefs_selected <- cbind(coefs_selected, cors[features_selected_idx])
	colnames(coefs_selected)[5] <- "Pearson Cor vs TR"
	## order the selected features by the p-values of their coefficients in a multiple linear model
	coefs_selected <- coefs_selected[order(coefs_selected[,4]),]
	
	dTICE_features <- data_frame[,features_selected_idx+1]
	return(list(dTICE_features=dTICE_features, dTICE_feature_coefs=coefs_selected, R2=R2_selected, R2_PWM=R2_PWM))
}

# generate dTICE features
for (species in names(species_tissue_list)) {
	for (i in 1:length(species_tissue_list[[species]])) {
		species_tissue <- species_tissue_list[[species]][i]
		if (species_tissue!="") {
			tissue_dot <- paste0(".", species_tissue)
			tissue_us <- paste0("_", species_tissue)
		} else {
			tissue_dot <- ""
			tissue_us <- ""
		}
		if (species != "hs") {
			assign(x=paste0(species, tissue_dot, ".dTICE_features"),
				value=make_dTICE_features(get(paste0(species, tissue_dot, ".CDS_seqs")), get(paste0(species, tissue_dot, ".UTR_seqs")), get(paste0(species, tissue_dot, ".log10.TR")), get(paste0(species, ".TR_top_pctl")), get(paste0(species, ".n_right")), get(paste0(species, ".len_breaks")), ifexemplar=T)
			)
		} else {
			assign(x=paste0(species, tissue_dot, ".dTICE_features"),
				value=make_dTICE_features(get(paste0(species, tissue_dot, ".CDS_seqs")), get(paste0(species, tissue_dot, ".UTR_seqs")), get(paste0(species, tissue_dot, ".log10.TR")), get(paste0(species, tissue_dot, ".TR_top_pctl")), get(paste0(species, tissue_dot, ".n_right")), get(paste0(species, tissue_dot, ".len_breaks")), ifexemplar=T)
			)
		}		
		saveRDS(get(paste0(species, tissue_dot, ".dTICE_features")), file=paste0("processed_data_w_polyA/dTICE_features/", species, tissue_dot, ".dTICE_features.rds"))		
	}
}

#####
# 5' of TICE features (a mega-feature)
make_5ofTICE_features <- function(UTR_seqs, log10.TR, TR_top_pctl, n_left, n_5ofTICE, len_breaks, ifexemplar, selected_feature_names=NULL) {
	# use forward + BIC to select features among the following features
	#(1) 16 dinucleotide frequencies in the 5' end to -(n_left+1) region
	#(2) 64 trinucleotide frequencies in the 5' end to -(n_left+1) region
	#(3) PWM scores based on the PWM constructed from the top 10% TR genes’ sequences
	
	#UTR_seqs: a character vector of UTR sequences, with its names as gene names
	#log10.TR: a numeric vector of log10.TR values, with its names as gene names
	#TR_top_pctl: the percentile to define TR top genes, e.g., 0.9
	#n_left: the minimum UTR length of "long" genes
	#n_5ofTICE: the 5ofTICE PWM length (from the 5' end)
	#len_breaks: a numeric vector containing the UTR length breaks, which divide genes into groups
	#ifexemplar = TRUE/FLASE; if TRUE, perform feature selection; otherwise, use selected_feature_names
	#selected_feature_names = a character vector of the selected feature names from the exemplar tissue
	
	n <- length(UTR_seqs)
	UTR_lens <- nchar(UTR_seqs)
	TR_top_UTR_seqs <- UTR_seqs[log10.TR >= quantile(log10.TR, prob=TR_top_pctl)]
	
	# cut each gene's 5' UTR sequence to the (5' end):-(n_left+1) region
	UTR_seqs_5ofTICE <- substr( x=UTR_seqs, start=1, stop=nchar(UTR_seqs)-n_left )
	TR_top_UTR_seqs_5ofTICE <- substr( x=TR_top_UTR_seqs, start=1, stop=nchar(TR_top_UTR_seqs)-n_left )
	
	# count the dinucleotide frequencies in the (5' end):-(n_left+1) region of each gene
	UTR_di_counts_5ofTICE <- sapply(di_nts, FUN=function(nuc) {
	    stri_count_regex(UTR_seqs_5ofTICE, paste0('(?=',nuc,')'))
	})
	rownames(UTR_di_counts_5ofTICE) <- names(UTR_seqs_5ofTICE)
	UTR_di_freqs_5ofTICE <- UTR_di_counts_5ofTICE / rowSums(UTR_di_counts_5ofTICE)
	UTR_di_freqs_5ofTICE[is.na(UTR_di_freqs_5ofTICE)] <- 0
	
	# count the trinucleotide frequencies in the (5' end):-(n_left+1) region of each gene
	UTR_tri_counts_5ofTICE <- sapply(tri_nts, FUN=function(nuc) {
	    stri_count_regex(UTR_seqs_5ofTICE, paste0('(?=',nuc,')'))
	})
	rownames(UTR_tri_counts_5ofTICE) <- names(UTR_seqs_5ofTICE)
	UTR_tri_freqs_5ofTICE <- UTR_tri_counts_5ofTICE / rowSums(UTR_tri_counts_5ofTICE)
	UTR_tri_freqs_5ofTICE[is.na(UTR_tri_freqs_5ofTICE)] <- 0
	
	# calculate PWM scores based on the TR top PWM in the 5' most region of length n_5ofTICE for each gene
	if (n_5ofTICE > 0) { # the 5ofTICE PWM scores will be used
		PWM_5ofTICE_scores <- sapply(n_5ofTICE, FUN=function(len) {
			TR_top_PWM_5ofTICE <- UTR_pwm_5_aligned( TR_top_UTR_seqs_5ofTICE, len )
			UTR_pwm_5_aligned_score( TR_top_PWM_5ofTICE, UTR_seqs )
		})
		colnames(PWM_5ofTICE_scores) <- paste0("PWM", n_5ofTICE)
		features <- cbind(UTR_di_freqs_5ofTICE, UTR_tri_freqs_5ofTICE, PWM_5ofTICE_scores)
	} else { # the 5ofTICE PWM scores will not be used
		features <- cbind(UTR_di_freqs_5ofTICE, UTR_tri_freqs_5ofTICE)
	}
	
	# combine the features into a design matrix
	feature_names <- colnames(features)
	cors <- apply(features, 2, FUN=function(x) cor(log10.TR, x, use="complete.obs"))
	if (length(len_breaks) > 0) {
		features_by_group <- NULL
		cors_by_group <- NULL # Pearson correlation between log10.TR and every feature within a group
		num_groups <- length(len_breaks) + 1 # number of gene groups divided by UTR lengths
		for (i in 1:num_groups) {
			if (i == 1) {
				features_one_group <- as.numeric(UTR_lens<len_breaks[1]) * features
				colnames(features_one_group) <- paste(feature_names, paste0("<",len_breaks[1]), sep="_")
				idx_one_group <- which(UTR_lens<len_breaks[1])
			} else if (i == num_groups) {
				features_one_group <- as.numeric(UTR_lens>=len_breaks[num_groups-1]) * cbind(features, rep(1,n))
				colnames(features_one_group) <- paste(c(feature_names,"Intercept"), paste0(">=",len_breaks[num_groups-1]), sep="_")
				idx_one_group <- which(UTR_lens>=len_breaks[num_groups-1])
			} else {
				features_one_group <- as.numeric(UTR_lens>=len_breaks[i-1] & UTR_lens<len_breaks[i]) * cbind(features, rep(1,n))
				colnames(features_one_group) <- paste(c(feature_names,"Intercept"), paste0(">=",len_breaks[i-1],"&<",len_breaks[i]), sep="_")
				idx_one_group <- which(UTR_lens>=len_breaks[i-1] & UTR_lens<len_breaks[i])
			}
			features_by_group <- cbind(features_by_group, features_one_group)
			cors_one_group <- apply(features_one_group, 2, FUN=function(x) cor(log10.TR[idx_one_group], x[idx_one_group], use="complete.obs"))
			cors_by_group <- c(cors_by_group, cors_one_group)
		}
		features <- features_by_group
		cors <- cors_by_group
	}
	
	# combine the response and the design matrix into a data frame
	# use forward selection + Bayesian Information Criterion (BIC) to select features
	if (n_5ofTICE > 0) { # the 5ofTICE PWM scores will be used
		PWM_idx <- which(grepl("PWM", colnames(features)))
		data_frame <- cbind.data.frame(log10.TR=log10.TR, features[,PWM_idx], features[,-PWM_idx])
		colnames(data_frame) <- c("log10.TR", colnames(features)[PWM_idx], colnames(features)[-PWM_idx])
		cors <- c(cors[PWM_idx], cors[-PWM_idx])
		if (ifexemplar) {
			regfit.fwd <- regsubsets(log10.TR ~. , data=data_frame, nvmax=70, force.in=1:length(PWM_idx), method ="forward")
		}		
	} else { # the 5ofTICE PWM scores will not be used
		data_frame <- cbind.data.frame(log10.TR=log10.TR, features)
		if (ifexemplar) {
			regfit.fwd <- regsubsets(log10.TR ~. , data=data_frame, nvmax=70, method ="forward")
		}		
	}
	
	if (ifexemplar) {
		reg.summary <- summary(regfit.fwd)
		idx_bic <- which.min(reg.summary$bic )
		features_selected_idx <- which(reg.summary$outmat[idx_bic,]=="*")
	} else {
		features_selected_idx <- which(colnames(data_frame) %in% selected_feature_names) - 1
	}
	
	lm_selected <- summary(lm(log10.TR ~. , data=data_frame[,c(1,features_selected_idx+1)]))
	coefs_selected <- lm_selected$coefficients[-1,]
	R2_selected <- lm_selected$r.squared
	coefs_selected <- cbind(coefs_selected, cors[features_selected_idx])
	colnames(coefs_selected)[5] <- "Pearson Cor vs TR"
	## order the selected features by the p-values of their coefficients in a multiple linear model
	coefs_selected <- coefs_selected[order(coefs_selected[,4]),]
	
	fiveofTICE_features <- data_frame[,features_selected_idx+1]
	return(list(fiveofTICE_features=fiveofTICE_features, fiveofTICE_feature_coefs=coefs_selected, R2=R2_selected))
}

# generate 5' of TICE features
for (species in names(species_tissue_list)) {
	for (i in 1:length(species_tissue_list[[species]])) {
		species_tissue <- species_tissue_list[[species]][i]
		if (species_tissue!="") {
			tissue_dot <- paste0(".", species_tissue)
			tissue_us <- paste0("_", species_tissue)
		} else {
			tissue_dot <- ""
			tissue_us <- ""
		}
		if (species != "hs") {
			assign(x=paste0(species, tissue_dot, ".5ofTICE_features"),
				value=make_5ofTICE_features(get(paste0(species, tissue_dot, ".UTR_seqs")), get(paste0(species, tissue_dot, ".log10.TR")), get(paste0(species, ".TR_top_pctl")), get(paste0(species, ".n_left")), get(paste0(species, ".n_5ofTICE")), get(paste0(species, ".len_breaks")), ifexemplar=T)
			)
		} else {
			assign(x=paste0(species, tissue_dot, ".5ofTICE_features"),
				value=make_5ofTICE_features(get(paste0(species, tissue_dot, ".UTR_seqs")), get(paste0(species, tissue_dot, ".log10.TR")), get(paste0(species, tissue_dot, ".TR_top_pctl")), get(paste0(species, tissue_dot, ".n_left")), get(paste0(species, tissue_dot, ".n_5ofTICE")), get(paste0(species, tissue_dot, ".len_breaks")), ifexemplar=T)
			)
		}
		saveRDS(get(paste0(species, tissue_dot, ".5ofTICE_features")), file=paste0("processed_data_w_polyA/5ofTICE_features/", species, tissue_dot, ".5ofTICE_features.rds"))		
	}
}

#####
# 20 amino acid frequencies in the CDS (a mega-feature)
## "K" is the reference level
make_20aa_features <- function(aa_counts, log10.TR, log10.UTR_lens, len_breaks) {
	n <- length(log10.UTR_lens)
	UTR_lens <- 10^log10.UTR_lens
	features <- aa_counts / rowSums(aa_counts)
	features <- features[,-1] # use "K" as the reference level
	cors <- apply(features, 2, FUN=function(x) cor(log10.TR, x, use="complete.obs"))
	feature_names <- colnames(features)
	if (length(len_breaks) > 0) {
		features_by_group <- NULL
		cors_by_group <- NULL # Pearson correlation between log10.TR and every feature within a group
		num_groups <- length(len_breaks) + 1 # number of gene groups divided by UTR lengths
		for (i in 1:num_groups) {
			if (i == 1) {
				features_one_group <- as.numeric(UTR_lens<len_breaks[1]) * features
				colnames(features_one_group) <- paste(feature_names, paste0("<",len_breaks[1]), sep="_")
				idx_one_group <- which(UTR_lens<len_breaks[1])
			} else if (i == num_groups) {
				features_one_group <- as.numeric(UTR_lens>=len_breaks[num_groups-1]) * cbind(features, rep(1,n))
				colnames(features_one_group) <- paste(c(feature_names,"Intercept"), paste0(">=",len_breaks[num_groups-1]), sep="_")
				idx_one_group <- which(UTR_lens>=len_breaks[num_groups-1])
			} else {
				features_one_group <- as.numeric(UTR_lens>=len_breaks[i-1] & UTR_lens<len_breaks[i]) * cbind(features, rep(1,n))
				colnames(features_one_group) <- paste(c(feature_names,"Intercept"), paste0(">=",len_breaks[i-1],"&<",len_breaks[i]), sep="_")
				idx_one_group <- which(UTR_lens>=len_breaks[i-1] & UTR_lens<len_breaks[i])
			}
			features_by_group <- cbind(features_by_group, features_one_group)
			cors_one_group <- apply(features_one_group, 2, FUN=function(x) cor(log10.TR[idx_one_group], x[idx_one_group], use="complete.obs"))
			cors_by_group <- c(cors_by_group, cors_one_group)
		}
		features <- features_by_group
		cors <- cors_by_group
	}
	# combine the response and the design matrix into a data frame
	data_frame <- cbind.data.frame(log10.TR=log10.TR, features)
	lm_selected <- summary(lm(log10.TR ~. , data=data_frame))
	coefs_selected <- lm_selected$coefficients[-1,]
	R2_selected <- lm_selected$r.squared
	coefs_selected <- cbind(coefs_selected, cors)
	colnames(coefs_selected)[5] <- "Pearson Cor vs TR"
	## order the features by the p-values of their coefficients in a multiple linear model
	coefs_selected <- coefs_selected[order(coefs_selected[,4]),]
	return(list(aa_features=features, aa_feature_coefs=coefs_selected, R2=R2_selected))
}

# generate aa features
for (species in names(species_tissue_list)) {
	for (i in 1:length(species_tissue_list[[species]])) {
		species_tissue <- species_tissue_list[[species]][i]
		if (species_tissue!="") {
			tissue_dot <- paste0(".", species_tissue)
			tissue_us <- paste0("_", species_tissue)
		} else {
			tissue_dot <- ""
			tissue_us <- ""
		}
		## count the frequency of 64 codons (including 3 stop codons) in every gene
		CDS_seqs <- get(paste0(species, tissue_dot, ".CDS_seqs"))
		codon_counts <- t( sapply(CDS_seqs, FUN=function(seq) {
		    sst <- strsplit(seq, "")[[1]]
		    n <- length(sst)
		    out <- paste0(sst[seq(from=1, to=n, by=3)], sst[seq(from=2, to=n, by=3)], sst[seq(from=3, to=n, by=3)])
		    table(out)[tri_nts]
		}) )
		codon_counts[is.na(codon_counts)] <- 0
		rownames(codon_counts) <- names(CDS_seqs)
		colnames(codon_counts) <- tri_nts
		## count the frequency of 20 amino acids (AAs) in every gene
		aa_counts <- sapply(aas, FUN=function(aa) {
		    idx <- which(tri_nts_to_aas==aa)
		    if (length(idx)==1) {
		        codon_counts[,idx]
		    } else {
		        rowSums(codon_counts[,idx])
		    }   
		})
		
		saveRDS(aa_counts / rowSums(aa_counts), file=paste0("processed_data_w_polyA/aa_features/", species, tissue_dot, ".aa_freqs.rds"))
		
		# features divided by 5'UTR length groups
		if (species != "hs") {
			aa_features <- make_20aa_features(aa_counts, get(paste0(species, tissue_dot, ".log10.TR")), get(paste0(species, tissue_dot, ".log10.UTR_lens")), get(paste0(species, ".len_breaks")))
		} else {
			aa_features <- make_20aa_features(aa_counts, get(paste0(species, tissue_dot, ".log10.TR")), get(paste0(species, tissue_dot, ".log10.UTR_lens")), get(paste0(species, tissue_dot, ".len_breaks")))
		}
		
		saveRDS(aa_features, file=paste0("processed_data_w_polyA/aa_features/", species, tissue_dot, ".aa_features.rds"))		
	}
}

#####
# 61 nonstop codon frequencies in the CDS (a mega-feature)
## "AAA" is the reference level
make_61codon_features <- function(codon_counts, log10.TR, log10.UTR_lens, len_breaks) {
	n <- length(log10.UTR_lens)
	UTR_lens <- 10^log10.UTR_lens
	features <- codon_counts / rowSums(codon_counts)
	features <- features[,-1] # use "AAA" as the reference level
	cors <- apply(features, 2, FUN=function(x) cor(log10.TR, x, use="complete.obs"))
	feature_names <- colnames(features)
	if (length(len_breaks) > 0) {
		features_by_group <- NULL
		cors_by_group <- NULL # Pearson correlation between log10.TR and every feature within a group
		num_groups <- length(len_breaks) + 1 # number of gene groups divided by UTR lengths
		for (i in 1:num_groups) {
			if (i == 1) {
				features_one_group <- as.numeric(UTR_lens<len_breaks[1]) * features
				colnames(features_one_group) <- paste(feature_names, paste0("<",len_breaks[1]), sep="_")
				idx_one_group <- which(UTR_lens<len_breaks[1])
			} else if (i == num_groups) {
				features_one_group <- as.numeric(UTR_lens>=len_breaks[num_groups-1]) * cbind(features, rep(1,n))
				colnames(features_one_group) <- paste(c(feature_names,"Intercept"), paste0(">=",len_breaks[num_groups-1]), sep="_")
				idx_one_group <- which(UTR_lens>=len_breaks[num_groups-1])
			} else {
				features_one_group <- as.numeric(UTR_lens>=len_breaks[i-1] & UTR_lens<len_breaks[i]) * cbind(features, rep(1,n))
				colnames(features_one_group) <- paste(c(feature_names,"Intercept"), paste0(">=",len_breaks[i-1],"&<",len_breaks[i]), sep="_")
				idx_one_group <- which(UTR_lens>=len_breaks[i-1] & UTR_lens<len_breaks[i])
			}
			features_by_group <- cbind(features_by_group, features_one_group)
			cors_one_group <- apply(features_one_group, 2, FUN=function(x) cor(log10.TR[idx_one_group], x[idx_one_group], use="complete.obs"))
			cors_by_group <- c(cors_by_group, cors_one_group)
		}
		features <- features_by_group
		cors <- cors_by_group
	}
	# combine the response and the design matrix into a data frame
	data_frame <- cbind.data.frame(log10.TR=log10.TR, features)
	lm_selected <- summary(lm(log10.TR ~. , data=data_frame))
	coefs_selected <- lm_selected$coefficients[-1,]
	R2_selected <- lm_selected$r.squared
	coefs_selected <- cbind(coefs_selected, cors)
	colnames(coefs_selected)[5] <- "Pearson Cor vs TR"
	## order the features by the p-values of their coefficients in a multiple linear model
	coefs_selected <- coefs_selected[order(coefs_selected[,4]),]
	return(list(codon_features=features, codon_feature_coefs=coefs_selected, R2=R2_selected))
}

# generate codon features
for (species in names(species_tissue_list)) {
	for (i in 1:length(species_tissue_list[[species]])) {
		species_tissue <- species_tissue_list[[species]][i]
		if (species_tissue!="") {
			tissue_dot <- paste0(".", species_tissue)
			tissue_us <- paste0("_", species_tissue)
		} else {
			tissue_dot <- ""
			tissue_us <- ""
		}
		## count the frequency of 64 codons (including 3 stop codons) in every gene
		CDS_seqs <- get(paste0(species, tissue_dot, ".CDS_seqs"))
		codon_counts <- t( sapply(CDS_seqs, FUN=function(seq) {
		    sst <- strsplit(seq, "")[[1]]
		    n <- length(sst)
		    out <- paste0(sst[seq(from=1, to=n, by=3)], sst[seq(from=2, to=n, by=3)], sst[seq(from=3, to=n, by=3)])
		    table(out)[nonstop_codons]
		}) )
		codon_counts[is.na(codon_counts)] <- 0
		rownames(codon_counts) <- names(CDS_seqs)
		colnames(codon_counts) <- nonstop_codons
		
		saveRDS(codon_counts / rowSums(codon_counts), file=paste0("processed_data_w_polyA/codon_features/", species, tissue_dot, ".codon_freqs.rds"))
		
		# features divided by 5'UTR length groups
		if (species != "hs") {
			codon_features <- make_61codon_features(codon_counts, get(paste0(species, tissue_dot, ".log10.TR")), get(paste0(species, tissue_dot, ".log10.UTR_lens")), get(paste0(species, ".len_breaks")))
		} else {
			codon_features <- make_61codon_features(codon_counts, get(paste0(species, tissue_dot, ".log10.TR")), get(paste0(species, tissue_dot, ".log10.UTR_lens")), get(paste0(species, tissue_dot, ".len_breaks")))
		}
		
		saveRDS(codon_features, file=paste0("processed_data_w_polyA/codon_features/", species, tissue_dot, ".codon_features.rds"))		
	}
}
#####

#####
# 61 nonstop codon frequencies in the first and second halves of CDS
for (species in names(species_tissue_list)) {
	for (i in 1:length(species_tissue_list[[species]])) {
		species_tissue <- species_tissue_list[[species]][i]
		if (species_tissue!="") {
			tissue_dot <- paste0(".", species_tissue)
			tissue_us <- paste0("_", species_tissue)
		} else {
			tissue_dot <- ""
			tissue_us <- ""
		}
		## count the frequency of 64 codons (including 3 stop codons) in every gene's first and second halves of CDS
		CDS_seqs <- get(paste0(species, tissue_dot, ".CDS_seqs"))
		CDS_seqs_1 <- substr( x=CDS_seqs, start=1, stop=round(nchar(CDS_seqs)/2) )
		CDS_seqs_2 <- substr( x=CDS_seqs, start=round(nchar(CDS_seqs)/2)+1, stop=nchar(CDS_seqs) )
		
		codon_counts_1 <- t( sapply(CDS_seqs_1, FUN=function(seq) {
		    sst <- strsplit(seq, "")[[1]]
		    n <- length(sst)
		    out <- paste0(sst[seq(from=1, to=n, by=3)], sst[seq(from=2, to=n, by=3)], sst[seq(from=3, to=n, by=3)])
		    table(out)[nonstop_codons]
		}) )
		codon_counts_1[is.na(codon_counts_1)] <- 0
		rownames(codon_counts_1) <- names(CDS_seqs_1)
		colnames(codon_counts_1) <- nonstop_codons
		
		codon_counts_2 <- t( sapply(CDS_seqs_2, FUN=function(seq) {
		    sst <- strsplit(seq, "")[[1]]
		    n <- length(sst)
		    out <- paste0(sst[seq(from=1, to=n, by=3)], sst[seq(from=2, to=n, by=3)], sst[seq(from=3, to=n, by=3)])
		    table(out)[nonstop_codons]
		}) )
		codon_counts_2[is.na(codon_counts_2)] <- 0
		rownames(codon_counts_2) <- names(CDS_seqs_2)
		colnames(codon_counts_2) <- nonstop_codons
		
		if (species != "hs") {
			codon_features_1 <- make_61codon_features(codon_counts_1, get(paste0(species, tissue_dot, ".log10.TR")), get(paste0(species, tissue_dot, ".log10.UTR_lens")), get(paste0(species, ".len_breaks")))
			codon_features_2 <- make_61codon_features(codon_counts_2, get(paste0(species, tissue_dot, ".log10.TR")), get(paste0(species, tissue_dot, ".log10.UTR_lens")), get(paste0(species, ".len_breaks")))
		} else {
			codon_features_1 <- make_61codon_features(codon_counts_1, get(paste0(species, tissue_dot, ".log10.TR")), get(paste0(species, tissue_dot, ".log10.UTR_lens")), get(paste0(species, tissue_dot, ".len_breaks")))
			codon_features_2 <- make_61codon_features(codon_counts_2, get(paste0(species, tissue_dot, ".log10.TR")), get(paste0(species, tissue_dot, ".log10.UTR_lens")), get(paste0(species, tissue_dot, ".len_breaks")))
		}
		
		saveRDS(codon_features_1, file=paste0("processed_data_w_polyA/codon_features/", species, tissue_dot, ".codon_features_1.rds"))
		saveRDS(codon_features_2, file=paste0("processed_data_w_polyA/codon_features/", species, tissue_dot, ".codon_features_2.rds"))	
	}
}

#####
# 61 synonymous codon frequencies in the CDS (a mega-feature)
## For example, AAA, AAT, AAC and AAG code the same AA, so we compute their synonymous codon frequencies by normalizing AA frequencies as follows
##   AAA / (AAA+AAT+AAG+AAC)
##   AAT / (AAA+AAT+AAG+AAC)
##   AAC / (AAA+AAT+AAG+AAC)
##   AAG / (AAA+AAT+AAG+AAC)
make_61syncodon_features <- function(syncodon_ratios, log10.TR, log10.UTR_lens, len_breaks) {
	n <- length(log10.UTR_lens)
	UTR_lens <- 10^log10.UTR_lens
	features <- syncodon_ratios
	cors <- apply(features, 2, FUN=function(x) cor(log10.TR, x, use="complete.obs"))
	feature_names <- colnames(features)
	if (length(len_breaks) > 0) {
		features_by_group <- NULL
		cors_by_group <- NULL # Pearson correlation between log10.TR and every feature within a group
		num_groups <- length(len_breaks) + 1 # number of gene groups divided by UTR lengths
		for (i in 1:num_groups) {
			if (i == 1) {
				features_one_group <- as.numeric(UTR_lens<len_breaks[1]) * features
				colnames(features_one_group) <- paste(feature_names, paste0("<",len_breaks[1]), sep="_")
				idx_one_group <- which(UTR_lens<len_breaks[1])
			} else if (i == num_groups) {
				features_one_group <- as.numeric(UTR_lens>=len_breaks[num_groups-1]) * cbind(features, rep(1,n))
				colnames(features_one_group) <- paste(c(feature_names,"Intercept"), paste0(">=",len_breaks[num_groups-1]), sep="_")
				idx_one_group <- which(UTR_lens>=len_breaks[num_groups-1])
			} else {
				features_one_group <- as.numeric(UTR_lens>=len_breaks[i-1] & UTR_lens<len_breaks[i]) * cbind(features, rep(1,n))
				colnames(features_one_group) <- paste(c(feature_names,"Intercept"), paste0(">=",len_breaks[i-1],"&<",len_breaks[i]), sep="_")
				idx_one_group <- which(UTR_lens>=len_breaks[i-1] & UTR_lens<len_breaks[i])
			}
			features_by_group <- cbind(features_by_group, features_one_group)
			cors_one_group <- apply(features_one_group, 2, FUN=function(x) cor(log10.TR[idx_one_group], x[idx_one_group], use="complete.obs"))
			cors_by_group <- c(cors_by_group, cors_one_group)
		}
		features <- features_by_group
		cors <- cors_by_group
	}
	# combine the response and the design matrix into a data frame
	data_frame <- cbind.data.frame(log10.TR=log10.TR, features)
	lm_selected <- summary(lm(log10.TR ~. , data=data_frame))
	coefs_selected <- lm_selected$coefficients[-1,]
	R2_selected <- lm_selected$r.squared
	coefs_selected <- cbind(coefs_selected, cors)
	colnames(coefs_selected)[5] <- "Pearson Cor vs TR"
	## order the features by the p-values of their coefficients in a multiple linear model
	coefs_selected <- coefs_selected[order(coefs_selected[,4]),]
	return(list(codon_features=features, codon_feature_coefs=coefs_selected, R2=R2_selected))
}

# generate synonymous codon features
for (species in names(species_tissue_list)) {
	for (i in 1:length(species_tissue_list[[species]])) {
		species_tissue <- species_tissue_list[[species]][i]
		if (species_tissue!="") {
			tissue_dot <- paste0(".", species_tissue)
			tissue_us <- paste0("_", species_tissue)
		} else {
			tissue_dot <- ""
			tissue_us <- ""
		}
		## count the frequency of 64 codons (including 3 stop codons) in every gene
		CDS_seqs <- get(paste0(species, tissue_dot, ".CDS_seqs"))
		codon_counts <- t( sapply(CDS_seqs, FUN=function(seq) {
		    sst <- strsplit(seq, "")[[1]]
		    n <- length(sst)
		    out <- paste0(sst[seq(from=1, to=n, by=3)], sst[seq(from=2, to=n, by=3)], sst[seq(from=3, to=n, by=3)])
		    table(out)[nonstop_codons]
		}) )
		codon_counts[is.na(codon_counts)] <- 0
		rownames(codon_counts) <- names(CDS_seqs)
		colnames(codon_counts) <- nonstop_codons
		
		## normalize the codon counts by their corresponding AA count
		syncodon_ratios <- codon_counts
		for (aa in aas) {
			tmp_codons <- names(tri_nts_to_aas[which(tri_nts_to_aas==aa)])
			syncodon_ratios[,tmp_codons] <- codon_counts[,tmp_codons] / rowSums(codon_counts[,tmp_codons, drop=F])
		}
		
		saveRDS(syncodon_ratios, file=paste0("processed_data_w_polyA/syncodon_features/", species, tissue_dot, ".syncodon_ratios.rds"))
		
		# features divided by 5'UTR length groups
		if (species != "hs") {
			syncodon_features <- make_61syncodon_features(syncodon_ratios, get(paste0(species, tissue_dot, ".log10.TR")), get(paste0(species, tissue_dot, ".log10.UTR_lens")), get(paste0(species, ".len_breaks")))
		} else {
			syncodon_features <- make_61syncodon_features(syncodon_ratios, get(paste0(species, tissue_dot, ".log10.TR")), get(paste0(species, tissue_dot, ".log10.UTR_lens")), get(paste0(species, tissue_dot, ".len_breaks")))
		}
		
		saveRDS(syncodon_features, file=paste0("processed_data_w_polyA/syncodon_features/", species, tissue_dot, ".syncodon_features.rds"))		
	}
}

#####
# tRNA abundances for sc
for (species in "sc") {
	for (i in 1:length(species_tissue_list[[species]])) {
		species_tissue <- species_tissue_list[[species]][i]
		if (species_tissue!="") {
			tissue_dot <- paste0(".", species_tissue)
			tissue_us <- paste0("_", species_tissue)
		} else {
			tissue_dot <- ""
			tissue_us <- ""
		}
		
		# read in combined tRNAs abundances (one per AA)
		aa_tRNA_abun <- read.table("../data/Weinberg tRNA abundances (aa).txt", header=T, sep="\t")
		aa_tRNA_abun <- sapply(aas, FUN=function(aa) {
      aa_tRNA_abun[ aa_tRNA_abun[,1]==aa, 2 ]
		})
		names(aa_tRNA_abun) <- aas

		# read in tRNAs abundances (one per codon)
		codon_tRNA_abun <- read.table("../data/Weinberg tRNA abundances (codon).txt", header=F, skip=1, sep="\t")
		codon_tRNA_abun <- sapply(nonstop_codons, FUN=function(codon) {
			codon_tRNA_abun[ codon_tRNA_abun[,2]==codon, 5 ]
		})
		
		saveRDS(aa_tRNA_abun, file=paste0("processed_data_w_polyA/tRNA_abun/", species, tissue_dot, ".aa_tRNA_abun.rds"))
		
		saveRDS(codon_tRNA_abun, file=paste0("processed_data_w_polyA/tRNA_abun/", species, tissue_dot, ".codon_tRNA_abun.rds"))	
	}
}
```
## Read in the already generated mega-features
```{r}
# read in uTICE features
for (species in names(species_tissue_list)) {
	for (i in 1:length(species_tissue_list[[species]])) {
		species_tissue <- species_tissue_list[[species]][i]
		if (species_tissue!="") {
			tissue_dot <- paste0(".", species_tissue)
			tissue_us <- paste0("_", species_tissue)
		} else {
			tissue_dot <- ""
			tissue_us <- ""
		}
		assign(x=paste0(species, tissue_dot, ".uTICE_features"),
			value=readRDS(file=paste0("processed_data_w_polyA/uTICE_features/", species, tissue_dot, ".uTICE_features.rds"))
		)			
	}
}

# read in dTICE features
for (species in names(species_tissue_list)) {
	for (i in 1:length(species_tissue_list[[species]])) {
		species_tissue <- species_tissue_list[[species]][i]
		if (species_tissue!="") {
			tissue_dot <- paste0(".", species_tissue)
			tissue_us <- paste0("_", species_tissue)
		} else {
			tissue_dot <- ""
			tissue_us <- ""
		}
		assign(x=paste0(species, tissue_dot, ".dTICE_features"),
			value=readRDS(file=paste0("processed_data_w_polyA/dTICE_features/", species, tissue_dot, ".dTICE_features.rds"))
		)			
	}
}

# read in 5' of TICE features
for (species in names(species_tissue_list)) {
	for (i in 1:length(species_tissue_list[[species]])) {
		species_tissue <- species_tissue_list[[species]][i]
		if (species_tissue!="") {
			tissue_dot <- paste0(".", species_tissue)
			tissue_us <- paste0("_", species_tissue)
		} else {
			tissue_dot <- ""
			tissue_us <- ""
		}
		assign(x=paste0(species, tissue_dot, ".5ofTICE_features"),
			value=readRDS(file=paste0("processed_data_w_polyA/5ofTICE_features/", species, tissue_dot, ".5ofTICE_features.rds"))
		)			
	}
}

# read in aa features
for (species in names(species_tissue_list)) {
	for (i in 1:length(species_tissue_list[[species]])) {
		species_tissue <- species_tissue_list[[species]][i]
		if (species_tissue!="") {
			tissue_dot <- paste0(".", species_tissue)
			tissue_us <- paste0("_", species_tissue)
		} else {
			tissue_dot <- ""
			tissue_us <- ""
		}
		assign(x=paste0(species, tissue_dot, ".aa_features"),
			value=readRDS(file=paste0("processed_data_w_polyA/aa_features/", species, tissue_dot, ".aa_features.rds"))
		)			
	}
}

# read in codon features
for (species in names(species_tissue_list)) {
	for (i in 1:length(species_tissue_list[[species]])) {
		species_tissue <- species_tissue_list[[species]][i]
		if (species_tissue!="") {
			tissue_dot <- paste0(".", species_tissue)
			tissue_us <- paste0("_", species_tissue)
		} else {
			tissue_dot <- ""
			tissue_us <- ""
		}
		assign(x=paste0(species, tissue_dot, ".codon_features"),
			value=readRDS(file=paste0("processed_data_w_polyA/codon_features/", species, tissue_dot, ".codon_features.rds"))
		)			
	}
}
```
## Divide original single features into by-group features
```{r}
### a function
div_original_features <- function(features, feature_names, log10.UTR_lens, len_breaks) {
	# for features that have not been divided into groups (original features)
	n <- length(log10.UTR_lens)
	UTR_lens <- 10^log10.UTR_lens
	
	features <- matrix(features, ncol=1)
	colnames(features) <- feature_names
	
	if (length(len_breaks) > 0) {
		features_by_group <- NULL
		num_groups <- length(len_breaks) + 1 # number of gene groups divided by UTR lengths
		for (i in 1:num_groups) {
			if (i == 1) {
				features_one_group <- as.numeric(UTR_lens<len_breaks[1]) * features
				colnames(features_one_group) <- paste(feature_names, paste0("<",len_breaks[1]), sep="_")
			} else if (i == num_groups) {
				features_one_group <- as.numeric(UTR_lens>=len_breaks[num_groups-1]) * cbind(features, rep(1,n))
				colnames(features_one_group) <- paste(c(feature_names,"Intercept"), paste0(">=",len_breaks[num_groups-1]), sep="_")
			} else {
				features_one_group <- as.numeric(UTR_lens>=len_breaks[i-1] & UTR_lens<len_breaks[i]) * cbind(features, rep(1,n))
				colnames(features_one_group) <- paste(c(feature_names,"Intercept"), paste0(">=",len_breaks[i-1],"&<",len_breaks[i]), sep="_")
			}
			features_by_group <- cbind(features_by_group, features_one_group)
		}
		features <- features_by_group
	}
	
	return(features)
}

### log10.UTR_lens
for (species in names(species_tissue_list)) {
	for (i in 1:length(species_tissue_list[[species]])) {
		species_tissue <- species_tissue_list[[species]][i]
		if (species_tissue!="") {
			tissue_dot <- paste0(".", species_tissue)
			tissue_us <- paste0("_", species_tissue)
		} else {
			tissue_dot <- ""
			tissue_us <- ""
		}
		if (species != "hs") {
			assign(x=paste0(species, tissue_dot, ".log10.UTR_lens.div"),
				value=div_original_features(get(paste0(species, tissue_dot, ".log10.UTR_lens")), "log10.UTR_lens", get(paste0(species, tissue_dot, ".log10.UTR_lens")), get(paste0(species, ".len_breaks")))
			)
		} else {
			assign(x=paste0(species, tissue_dot, ".log10.UTR_lens.div"),
				value=div_original_features(get(paste0(species, tissue_dot, ".log10.UTR_lens")), "log10.UTR_lens", get(paste0(species, tissue_dot, ".log10.UTR_lens")), get(paste0(species, tissue_dot, ".len_breaks")))
			)
		}			
	}
}

### log10.CDS_lens
for (species in names(species_tissue_list)) {
	for (i in 1:length(species_tissue_list[[species]])) {
		species_tissue <- species_tissue_list[[species]][i]
		if (species_tissue!="") {
			tissue_dot <- paste0(".", species_tissue)
			tissue_us <- paste0("_", species_tissue)
		} else {
			tissue_dot <- ""
			tissue_us <- ""
		}
		if (species != "hs") {
			assign(x=paste0(species, tissue_dot, ".log10.CDS_lens.div"),
				value=div_original_features(get(paste0(species, tissue_dot, ".log10.CDS_lens")), "log10.CDS_lens", get(paste0(species, tissue_dot, ".log10.UTR_lens")), get(paste0(species, ".len_breaks")))
			)
		} else {
			assign(x=paste0(species, tissue_dot, ".log10.CDS_lens.div"),
				value=div_original_features(get(paste0(species, tissue_dot, ".log10.CDS_lens")), "log10.CDS_lens", get(paste0(species, tissue_dot, ".log10.UTR_lens")), get(paste0(species, tissue_dot, ".len_breaks")))
			)
		}		
	}
}

### uAUG_counts
for (species in names(species_tissue_list)) {
	for (i in 1:length(species_tissue_list[[species]])) {
		species_tissue <- species_tissue_list[[species]][i]
		if (species_tissue!="") {
			tissue_dot <- paste0(".", species_tissue)
			tissue_us <- paste0("_", species_tissue)
		} else {
			tissue_dot <- ""
			tissue_us <- ""
		}
		if (species != "hs") {
			assign(x=paste0(species, tissue_dot, ".uAUG_counts.div"),
				value=div_original_features(get(paste0(species, tissue_dot, ".uAUG_counts")), "uAUG_counts", get(paste0(species, tissue_dot, ".log10.UTR_lens")), get(paste0(species, ".len_breaks")))
			)
		} else {
			assign(x=paste0(species, tissue_dot, ".uAUG_counts.div"),
				value=div_original_features(get(paste0(species, tissue_dot, ".uAUG_counts")), "uAUG_counts", get(paste0(species, tissue_dot, ".log10.UTR_lens")), get(paste0(species, tissue_dot, ".len_breaks")))
			)
		}		
	}
}

### CDS_fold_energy
for (species in names(species_tissue_list)) {
	for (i in 1:length(species_tissue_list[[species]])) {
		species_tissue <- species_tissue_list[[species]][i]
		if (species_tissue!="") {
			tissue_dot <- paste0(".", species_tissue)
			tissue_us <- paste0("_", species_tissue)
		} else {
			tissue_dot <- ""
			tissue_us <- ""
		}
		if (species != "hs") {
			assign(x=paste0(species, tissue_dot, ".CDS_fold_energy.div"),
				value=div_original_features(get(paste0(species, tissue_dot, ".CDS_fold_energy")), "CDS_fold_energy", get(paste0(species, tissue_dot, ".log10.UTR_lens")), get(paste0(species, ".len_breaks")))
			)
		} else {
			assign(x=paste0(species, tissue_dot, ".CDS_fold_energy.div"),
				value=div_original_features(get(paste0(species, tissue_dot, ".CDS_fold_energy")), "CDS_fold_energy", get(paste0(species, tissue_dot, ".log10.UTR_lens")), get(paste0(species, tissue_dot, ".len_breaks")))
			)
		}		
	}
}

### polyA_lens
for (species in names(species_tissue_list)) {
	for (i in 1:length(species_tissue_list[[species]])) {
		species_tissue <- species_tissue_list[[species]][i]
		if (species_tissue!="") {
			tissue_dot <- paste0(".", species_tissue)
			tissue_us <- paste0("_", species_tissue)
		} else {
			tissue_dot <- ""
			tissue_us <- ""
		}
		if (species != "hs") {
			assign(x=paste0(species, tissue_dot, ".polyA_lens.div"),
				value=div_original_features(get(paste0(species, tissue_dot, ".polyA_lens")), "polyA_lens", get(paste0(species, tissue_dot, ".log10.UTR_lens")), get(paste0(species, ".len_breaks")))
			)
		} else {
			assign(x=paste0(species, tissue_dot, ".polyA_lens.div"),
				value=div_original_features(get(paste0(species, tissue_dot, ".polyA_lens")), "polyA_lens", get(paste0(species, tissue_dot, ".log10.UTR_lens")), get(paste0(species, tissue_dot, ".len_breaks")))
			)
		}		
	}
}

```

# Save the selected RNA folding features to excel files
```{r, eval = FALSE}
for (species in names(species_tissue_list)) {
  for (species_tissue in species_tissue_list[[species]]) {
    if (species_tissue!="") {
			tissue_dot <- paste0(".", species_tissue)
			tissue_us <- paste0("_", species_tissue)
		} else {
			tissue_dot <- ""
			tissue_us <- ""
		}
    if (species != "sp") {
      output <- readRDS(file=paste0("processed_data_w_polyA/folding_energy_features_selected/", species, tissue_us, "_UTR_folding_energy_features_selected.rds"))[[2]]
    } else {
      output <- readRDS(file=paste0("processed_data_w_polyA/folding_energy_features_selected/", species, tissue_us, "_UTR_folding_energy_features_selected_trun.rds"))[[2]]
    }
    write.csv(output, file=paste0("tables/folding_energy_features_selected/", species, tissue_dot, ".fold_energy_features_selected.csv"), quote=F)
  }
}
```

# Save the selected 5ofTICE, uTICE and dTICE features to excel files
```{r, eval = FALSE}
for (species in names(species_tissue_list)) {
  for (species_tissue in species_tissue_list[[species]]) {
    if (species_tissue!="") {
			tissue_dot <- paste0(".", species_tissue)
			tissue_us <- paste0("_", species_tissue)
		} else {
			tissue_dot <- ""
			tissue_us <- ""
		}
    
    output_5ofTICE <- readRDS(file=paste0("processed_data_w_polyA/5ofTICE_features/", species, tissue_dot, ".5ofTICE_features.rds"))[[2]]
    output_uTICE <- readRDS(file=paste0("processed_data_w_polyA/uTICE_features/", species, tissue_dot, ".uTICE_features.rds"))[[2]]
    output_dTICE <- readRDS(file=paste0("processed_data_w_polyA/dTICE_features/", species, tissue_dot, ".dTICE_features.rds"))[[2]]
    
    write.csv(output_5ofTICE, file=paste0("tables/5ofTICE_features/", species, tissue_dot, ".5ofTICE_features.csv"), quote=F)
    write.csv(output_uTICE, file=paste0("tables/uTICE_features/", species, tissue_dot, ".uTICE_features.csv"), quote=F)
    write.csv(output_dTICE, file=paste0("tables/dTICE_features/", species, tissue_dot, ".dTICE_features.csv"), quote=F)
  }
}
```

# Pearson correlation(s) between the amino acid frequency feature(s) or codon frequency feature(s) and log10(TR)
## save the values to excel files (the following code only needs to be run once)
```{r, eval = FALSE}
# calculate the Pearson correlation between each of the 20 amino acid frequencies and the 61 codon frequencies in the CDS (don't divide every feature into multiple groups) and log10(TR)
calc_aa_codon_features.logTR.cors <- function(species, tissue_dot) {
  ## log10.TR
  log10.TR <- get(paste0(species, tissue_dot, ".log10.TR"))
	## count the frequency of 64 codons (including 3 stop codons) in every gene
	CDS_seqs <- get(paste0(species, tissue_dot, ".CDS_seqs"))
	codon_counts <- t( sapply(CDS_seqs, FUN=function(seq) {
	    sst <- strsplit(seq, "")[[1]]
	    n <- length(sst)
	    out <- paste0(sst[seq(from=1, to=n, by=3)], sst[seq(from=2, to=n, by=3)], sst[seq(from=3, to=n, by=3)])
	    table(out)[tri_nts]
	}) )
	codon_counts[is.na(codon_counts)] <- 0
	rownames(codon_counts) <- names(CDS_seqs)
	colnames(codon_counts) <- tri_nts
	
	## count the frequency of 20 amino acids (AAs) in every gene
	aa_counts <- sapply(aas, FUN=function(aa) {
	    idx <- which(tri_nts_to_aas==aa)
	    if (length(idx)==1) {
	        codon_counts[,idx]
	    } else {
	        rowSums(codon_counts[,idx])
	    }   
	})
	
	aa_features <- aa_counts / rowSums(aa_counts)
	aa_cors <- apply(aa_features, 2, FUN=function(x) cor(log10.TR, x, use="complete.obs"))
	names(aa_cors) <- colnames(aa_features)
	
	codon_features <- codon_counts / rowSums(codon_counts)
	codon_cors <- apply(codon_features, 2, FUN=function(x) cor(log10.TR, x, use="complete.obs"))
	names(codon_cors) <- colnames(codon_features)
	
	return(list(aa_cors=aa_cors, codon_cors=codon_cors))
}


aa_features.logTR.cors <- NULL
rownames_aa_features.logTR.cors <- NULL

codon_features.logTR.cors <- NULL
rownames_codon_features.logTR.cors <- NULL

for (species in names(species_tissue_list)) {
	for (i in 1:length(species_tissue_list[[species]])) {
		species_tissue <- species_tissue_list[[species]][i]
		if (species_tissue!="") {
			tissue_dot <- paste0(".", species_tissue)
			tissue_us <- paste0("_", species_tissue)
		} else {
			tissue_dot <- ""
			tissue_us <- ""
		}
		
		results <- calc_aa_codon_features.logTR.cors(species, tissue_dot)
				
		aa_features.logTR.cors <- rbind(
			aa_features.logTR.cors,
			results$aa_cors
		)
		rownames_aa_features.logTR.cors <- c(
			rownames_aa_features.logTR.cors,
			paste0(species, tissue_dot)
		)
		
		codon_features.logTR.cors <- rbind(
			codon_features.logTR.cors,
			results$codon_cors
		)
		rownames_codon_features.logTR.cors <- c(
			rownames_codon_features.logTR.cors,
			paste0(species, tissue_dot)
		)		
	}
}

rownames(aa_features.logTR.cors) <- rownames_aa_features.logTR.cors
rownames(codon_features.logTR.cors) <- rownames_codon_features.logTR.cors

write.csv(aa_features.logTR.cors, file="tables/aa_features.logTR.cors.csv", quote=F)
write.csv(codon_features.logTR.cors, file="tables/codon_features.logTR.cors.csv", quote=F)
```
## plot the results (the following code only needs to be run once)
```{r, eval = FALSE}
aa_features.logTR.cors <- read.csv(file="tables/aa_features.logTR.cors.csv")
colnames(aa_features.logTR.cors)[1] <- "species.tissue"

codon_features.logTR.cors <- read.csv(file="tables/codon_features.logTR.cors.csv")
colnames(codon_features.logTR.cors)[1] <- "species.tissue"

aa_features.logTR.cors.for_plot <- setNames(melt(aa_features.logTR.cors), c("species.tissue", "aa.freq", 'cor'))
aa_features.logTR.cors.for_plot$species.tissue <- factor(aa_features.logTR.cors.for_plot$species.tissue, levels=as.character(aa_features.logTR.cors$species.tissue))

codon_features.logTR.cors.for_plot <- setNames(melt(codon_features.logTR.cors), c("species.tissue", "codon.freq", 'cor'))
codon_features.logTR.cors.for_plot$species.tissue <- factor(codon_features.logTR.cors.for_plot$species.tissue, levels=as.character(codon_features.logTR.cors$species.tissue))

pdf("tables/aa_features.logTR.cors.pdf", width=10, height=5)
p <- ggplot(aa_features.logTR.cors.for_plot, aes(x=aa.freq, y=species.tissue)) + 
  geom_tile(aes(fill = cor)) + 
  scale_fill_viridis() + 
  ggtitle("Pearson cor with log10(TR)")
print(p)
dev.off()

pdf("tables/codon_features.logTR.cors.pdf", width=10, height=5)
p <- ggplot(codon_features.logTR.cors.for_plot, aes(x=codon.freq, y=species.tissue)) + 
  geom_tile(aes(fill = cor)) + 
  scale_fill_viridis() +
  theme(axis.text.x = element_text(size = 6, angle = 45, vjust = 1, hjust=1)) +
  ggtitle("Pearson cor with log10(TR)")
print(p)
dev.off()
```
## R2 of the amino acid frequency feature(s) or codon frequency feature(s) in a multi-variate multi-part linear model
```{r}
for (species in names(species_tissue_list)) {
  for (species_tissue in species_tissue_list[[species]]) {
    if (species_tissue!="") {
			tissue_dot <- paste0(".", species_tissue)
			tissue_us <- paste0("_", species_tissue)
		} else {
			tissue_dot <- ""
			tissue_us <- ""
		}
    
    aa_features <- readRDS(paste0("processed_data_w_polyA/aa_features/", species, tissue_dot, ".aa_features.rds"))[[1]]
    cat(paste0(species, tissue_dot, " R2 of aa features: "))
    cat(summary(lm(get(paste0(species, tissue_dot, ".log10.TR")) ~ as.matrix(aa_features)))$r.squared)
    cat("\n")
    
    codon_features <- readRDS(paste0("processed_data_w_polyA/codon_features/", species, tissue_dot, ".codon_features.rds"))[[1]]
    cat(paste0(species, tissue_dot, " R2 of codon features: "))
    cat(summary(lm(get(paste0(species, tissue_dot, ".log10.TR")) ~ as.matrix(codon_features)))$r.squared)
    cat("\n")
    
    cat(paste0(species, tissue_dot, " R2 of aa + codon features: "))
    cat(summary(lm(get(paste0(species, tissue_dot, ".log10.TR")) ~ as.matrix(cbind(aa_features, codon_features))))$r.squared)
    cat("\n\n")
  }
}
```

# Generate tri-nucleotide frequencies in 5'ofTICE, uTICE, dTICE, and 3'ofTICE regions for all genes in each tissue of each species
```{r, eval = FALSE}
for (species in names(species_tissue_list)) {
	for (species_tissue in species_tissue_list[[species]]) {
		if (species_tissue!="") {
			species.UTR_seqs <- get(paste(species, species_tissue, "UTR_seqs", sep="."))
			species.CDS_seqs <- get(paste(species, species_tissue, "CDS_seqs", sep="."))
			connector <- paste0(".", species_tissue)
		} else {
			species.UTR_seqs <- get(paste(species, "UTR_seqs", sep="."))
			species.CDS_seqs <- get(paste(species, "CDS_seqs", sep="."))
			connector <- ""
		}
	  
	  if (species != "hs") {
			species.uTICE_len <- species_uTICE_len_list[[species]]
			species.dTICE_len <- species_dTICE_len_list[[species]]
		} else {
			species.uTICE_len <- species_uTICE_len_list[[species]][[species_tissue]]
			species.dTICE_len <- species_dTICE_len_list[[species]][[species_tissue]]
		}
	  
	  # 5' of TICE
	  species.UTR_seqs_5ofTICE <- substr( x=species.UTR_seqs, start=1, stop=nchar(species.UTR_seqs)-species.uTICE_len )
	  species.UTR_tri_counts_5ofTICE <- sapply(tri_nts, FUN=function(nuc) {
	    stri_count_regex(species.UTR_seqs_5ofTICE, paste0('(?=',nuc,')'))
	  })
	  rownames(species.UTR_tri_counts_5ofTICE) <- names(species.UTR_seqs_5ofTICE)
	  species.UTR_tri_freqs_5ofTICE <- species.UTR_tri_counts_5ofTICE / rowSums(species.UTR_tri_counts_5ofTICE)
	  species.UTR_tri_freqs_5ofTICE[is.na(species.UTR_tri_freqs_5ofTICE)] <- 0

	  saveRDS(species.UTR_tri_freqs_5ofTICE, file=paste0("processed_data_w_polyA/tri_freqs/", species, connector, ".UTR_tri_freqs_5ofTICE.rds"))
	  
	  # uTICE
	  species.UTR_seqs_uTICE <- substr( x=species.UTR_seqs, start=nchar(species.UTR_seqs)-species.uTICE_len+1, stop=nchar(species.UTR_seqs) )
	  species.UTR_tri_counts_uTICE <- sapply(tri_nts, FUN=function(nuc) {
	    stri_count_regex(species.UTR_seqs_uTICE, paste0('(?=',nuc,')'))
	  })
	  rownames(species.UTR_tri_counts_uTICE) <- names(species.UTR_seqs_uTICE)
	  species.UTR_tri_freqs_uTICE <- species.UTR_tri_counts_uTICE / rowSums(species.UTR_tri_counts_uTICE)
	  species.UTR_tri_freqs_uTICE[is.na(species.UTR_tri_freqs_uTICE)] <- 0
	  
	  saveRDS(species.UTR_tri_freqs_uTICE, file=paste0("processed_data_w_polyA/tri_freqs/", species, connector, ".UTR_tri_freqs_uTICE.rds"))
	  
	  # dTICE
	  species.CDS_seqs_dTICE <- substr( x=species.CDS_seqs, start=4, stop=species.dTICE_len )
	  species.CDS_tri_counts_dTICE <- sapply(tri_nts, FUN=function(nuc) {
	    stri_count_regex(species.CDS_seqs_dTICE, paste0('(?=',nuc,')'))
	  })
	  rownames(species.CDS_tri_counts_dTICE) <- names(species.CDS_seqs_dTICE)
	  species.CDS_tri_freqs_dTICE <- species.CDS_tri_counts_dTICE / rowSums(species.CDS_tri_counts_dTICE)
	  species.CDS_tri_freqs_dTICE[is.na(species.CDS_tri_freqs_dTICE)] <- 0
	  
	  saveRDS(species.CDS_tri_freqs_dTICE, file=paste0("processed_data_w_polyA/tri_freqs/", species, connector, ".CDS_tri_freqs_dTICE.rds"))
	  
	  # 3' of TICE
	  species.CDS_seqs_3ofTICE <- substr( x=species.CDS_seqs, start=species.dTICE_len+1, stop=nchar(species.CDS_seqs) )
	  species.CDS_tri_counts_3ofTICE <- sapply(tri_nts, FUN=function(nuc) {
	    stri_count_regex(species.CDS_seqs_3ofTICE, paste0('(?=',nuc,')'))
	  })
	  rownames(species.CDS_tri_counts_3ofTICE) <- names(species.CDS_seqs_3ofTICE)
	  species.CDS_tri_freqs_3ofTICE <- species.CDS_tri_counts_3ofTICE / rowSums(species.CDS_tri_counts_3ofTICE)
	  species.CDS_tri_freqs_3ofTICE[is.na(species.CDS_tri_freqs_3ofTICE)] <- 0
	  
	  saveRDS(species.CDS_tri_freqs_3ofTICE, file=paste0("processed_data_w_polyA/tri_freqs/", species, connector, ".CDS_tri_freqs_3ofTICE.rds"))	  
	}
}
```
# information needed to plot a trinucleotide ratio heatmap
```{r}
## trinucleotide table
tri_nts_table <- rbind(
  matrix(tri_nts[1:16], nrow=4),
  matrix(tri_nts[17:32], nrow=4),
  matrix(tri_nts[33:48], nrow=4),
  matrix(tri_nts[49:64], nrow=4)
)

## creates a own color palette from blue to red
my_palette <- colorRampPalette(c(rgb(103, 169, 207, maxColorValue = 255), rgb(247, 247, 247, maxColorValue = 255), rgb(239, 138, 98, maxColorValue = 255)))(n = 299)
```

# Find the minimum, the 2nd minimum, and the 3rd minimum folding energy windows in the 5'UTR for every gene; make the windows non-overlapping
```{r, eval = FALSE}
# find the optimal window length based on the R2 of the minimum folding energy and log10(TR)
for (species in names(species_tissue_list)) {
	for (species_tissue in species_tissue_list[[species]]) {
		if (species_tissue!="") {
			tissue_dot <- paste0(".", species_tissue)
			tissue_us <- paste0("_", species_tissue)
		} else {
			tissue_dot <- ""
			tissue_us <- ""
		}
		if (species == "sp") {
			fold_energy_features_all <- readRDS(paste0("processed_data_w_polyA/folding_energy_features_selected/", species, tissue_us, "_UTR_folding_energy_features_selected_trun.rds"))[[4]]
		} else {
			fold_energy_features_all <- readRDS(paste0("processed_data_w_polyA/folding_energy_features_selected/", species, tissue_us, "_UTR_folding_energy_features_selected.rds"))[[4]]
		}
		
		R2s_min_fold_energy_features <- sapply(window_lens, FUN=function(window_len) {
			min_fold_energy_features <- fold_energy_features_all[, c(which(grepl(paste0("^min", "_", window_len, "$"), colnames(fold_energy_features_all))), which(grepl(paste0("^min", "_", window_len, "_"), colnames(fold_energy_features_all))), which(grepl(paste0("^Intercept", "_", window_len), colnames(fold_energy_features_all))))]
			summary(lm(get(paste0(species, tissue_dot, ".log10.TR")) ~ as.matrix(min_fold_energy_features)))$r.squared
		})
		
		min_fold_energy_window_len <- as.numeric(window_lens[which.max(R2s_min_fold_energy_features)])
		
		min_fold_energy_features <- fold_energy_features_all[, c(which(grepl(paste0("^min", "_", min_fold_energy_window_len, "$"), colnames(fold_energy_features_all))), which(grepl(paste0("^min", "_", min_fold_energy_window_len, "_"), colnames(fold_energy_features_all))), which(grepl(paste0("^Intercept", "_", min_fold_energy_window_len), colnames(fold_energy_features_all))))]
		
		UTR_folding_energy_all <- readRDS(file=paste0("processed_data_w_polyA/var_win_fold_UTR5_CDSshort_all/", species, tissue_us, "_5UTR_CDS_folding_energy_", min_fold_energy_window_len, ".rds"))
		species_genes <- get(paste0(species, tissue_dot, ".genes"))
		UTR_folding_energy_all <- UTR_folding_energy_all$UTR_folding_energy_all[species_genes]
		
		if (species != "sp") {
			min_2ndmin_3rdmin_fold_energy_features <- t( sapply(UTR_folding_energy_all, FUN=function(x) {
				if (is.null(x)) {
					return(rep(NA, 3))
				} else {
					n <- length(x)
					x1 <- min(x)
					x1_loc <- which.min(x)
					rm_idx <- max(1, x1_loc - min_fold_energy_window_len + 1):min(n, x1_loc + min_fold_energy_window_len)
					x_kept <- x[-rm_idx]
					if (length(x_kept) > 0) {
						x2 <- min(x_kept)
						x2_loc <- which(x==x2)
						rm_idx <- c(rm_idx, max(1, x2_loc - min_fold_energy_window_len + 1):min(n, x2_loc + min_fold_energy_window_len))
						x_kept <- x[-rm_idx]
						if (length(x_kept) > 0) {
							x3 <- min(x_kept)
						} else {
							x3 <- NA
						}
					} else {
						x2 <- NA
						x3 <- NA
					}
				}
				return(c(x1, x2, x3))
			}) )
		} else { # sp, needs truncation to remove long windows that extend beyond +30
			trun_pos <- as.numeric(window_len) - 30 + 1 # the position (-trun_pos) relative to AUG; windows whose starting positions to the 3' of trun_pos should be removed
			UTR_folding_energy_all_trun <- lapply(UTR_folding_energy_all, FUN=function(x) {
				if (trun_pos > length(x)) {
					x <- NULL
				} else if (trun_pos > 0) {
					x <- rev(rev(x)[trun_pos:length(x)])
				}
				return(x)
			})
			min_2ndmin_3rdmin_fold_energy_features <- t( sapply(UTR_folding_energy_all_trun, FUN=function(x) {
				if (is.null(x)) {
					return(rep(NA, 3))
				} else {
					n <- length(x)
					x1 <- min(x)
					x1_loc <- which.min(x)
					rm_idx <- max(1, x1_loc - min_fold_energy_window_len + 1):min(n, x1_loc + min_fold_energy_window_len)
					x_kept <- x[-rm_idx]
					if (length(x_kept) > 0) {
						x2 <- min(x_kept)
						x2_loc <- which(x==x2)
						rm_idx <- c(rm_idx, max(1, x2_loc - min_fold_energy_window_len + 1):min(n, x2_loc + min_fold_energy_window_len))
						x_kept <- x[-rm_idx]
						if (length(x_kept) > 0) {
							x3 <- min(x_kept)
						} else {
							x3 <- NA
						}
					} else {
						x2 <- NA
						x3 <- NA
					}
				}
				return(c(x1, x2, x3))
			}) )
		}

		if (species != "hs") {
		  min_fold_energy_features.div <- div_original_features(min_2ndmin_3rdmin_fold_energy_features[,1], paste("min_fold_energy", min_fold_energy_window_len, sep="_"), get(paste0(species, tissue_dot, ".log10.UTR_lens")), get(paste0(species, ".len_breaks")))
		
		  second_min_fold_energy_features.div <- div_original_features(min_2ndmin_3rdmin_fold_energy_features[,2], paste("2ndmin_fold_energy", min_fold_energy_window_len, sep="_"), get(paste0(species, tissue_dot, ".log10.UTR_lens")), get(paste0(species, ".len_breaks")))
		
		  third_min_fold_energy_features.div <- div_original_features(min_2ndmin_3rdmin_fold_energy_features[,3], paste("3rdmin_fold_energy", min_fold_energy_window_len, sep="_"), get(paste0(species, tissue_dot, ".log10.UTR_lens")), get(paste0(species, ".len_breaks")))
		} else {
		  min_fold_energy_features.div <- div_original_features(min_2ndmin_3rdmin_fold_energy_features[,1], paste("min_fold_energy", min_fold_energy_window_len, sep="_"), get(paste0(species, tissue_dot, ".log10.UTR_lens")), get(paste0(species, tissue_dot, ".len_breaks")))
		
		  second_min_fold_energy_features.div <- div_original_features(min_2ndmin_3rdmin_fold_energy_features[,2], paste("2ndmin_fold_energy", min_fold_energy_window_len, sep="_"), get(paste0(species, tissue_dot, ".log10.UTR_lens")), get(paste0(species, tissue_dot, ".len_breaks")))
		
		  third_min_fold_energy_features.div <- div_original_features(min_2ndmin_3rdmin_fold_energy_features[,3], paste("3rdmin_fold_energy", min_fold_energy_window_len, sep="_"), get(paste0(species, tissue_dot, ".log10.UTR_lens")), get(paste0(species, tissue_dot, ".len_breaks")))
		}
		
		saveRDS(min_fold_energy_features.div, file=paste0("processed_data_w_polyA/min_2ndmin_3rdmin_fold_energy_features/", species, tissue_us, "_min_fold_energy_features.div.rds"))
		
		saveRDS(second_min_fold_energy_features.div, file=paste0("processed_data_w_polyA/min_2ndmin_3rdmin_fold_energy_features/", species, tissue_us, "_second_min_fold_energy_features.div.rds"))
		
		saveRDS(third_min_fold_energy_features.div, file=paste0("processed_data_w_polyA/min_2ndmin_3rdmin_fold_energy_features/", species, tissue_us, "_third_min_fold_energy_features.div.rds"))
		
		saveRDS(min_2ndmin_3rdmin_fold_energy_features, file=paste0("processed_data_w_polyA/min_2ndmin_3rdmin_fold_energy_features/", species, tissue_us, "_min_2ndmin_3rdmin_fold_energy_features"))
	}
}
```

# Figure 1. Distributions of log10(TR), log10(5'UTR length), and log10(CDS length) 
```{r Figure1, fig.width=5, fig.height=5, fig.path='figures/', dev=c('pdf'), out.width="50%"}
# a list of species and colors
species_colors <- list(
  sc="#EB9D41",
  sp="#94241D",
  at="#22549C",
  mm="#2F6E42",
  hs="#CEC448"
)

# number of species
m <- length(species_colors)

# number of genes in each exemplar tissue of each species
num_genes <- NULL
# distributions of the log10(TR), log10(UTR_lens) and log10(CDS_lens) values of each exemplar tissue in each species after aligning their median
center.log10.TR <- NULL
for (species in names(species_tissue_list)) {
	for (i in 1:length(species_exemplar_tissue_list[[species]])) {
		species_tissue <- species_exemplar_tissue_list[[species]][i]
		if (species_tissue!="") {
			tissue_dot <- paste0(".", species_tissue)
		} else {
			tissue_dot <- ""
		}
		num_genes <- c(num_genes, sum(!is.na(get(paste0(species, tissue_dot, ".log10.TR")))))
		center.log10.TR <- c(center.log10.TR, median(get(paste0(species, tissue_dot, ".log10.TR"))))
	}
}
center.log10.TR <- mean(center.log10.TR)
den.log10.TR <- vector("list", 0)
ymax.log10.TR <- NULL
xmin.log10.TR <- NULL
xmax.log10.TR <- NULL
den.log10.UTR_lens <- vector("list", 0)
ymax.log10.UTR_lens <- NULL
xmin.log10.UTR_lens <- NULL
xmax.log10.UTR_lens <- NULL
den.log10.CDS_lens <- vector("list", 0)
ymax.log10.CDS_lens <- NULL
xmin.log10.CDS_lens <- NULL
xmax.log10.CDS_lens <- NULL
for (species in names(species_tissue_list)) {
	for (i in 1:length(species_exemplar_tissue_list[[species]])) {
		species_tissue <- species_exemplar_tissue_list[[species]][i]
		if (species_tissue!="") {
			tissue_dot <- paste0(".", species_tissue)
		} else {
			tissue_dot <- ""
		}
		x <- get(paste0(species, tissue_dot, ".log10.TR"))
		den.log10.TR[[species]] <- density(x-median(x)+center.log10.TR)
		ymax.log10.TR <- c(ymax.log10.TR, max(den.log10.TR[[species]]$y))
		xmin.log10.TR <- c(xmin.log10.TR, min(den.log10.TR[[species]]$x))
		xmax.log10.TR <- c(xmax.log10.TR, max(den.log10.TR[[species]]$x))
		x <- log(nchar(get(paste0(species, tissue_dot, ".UTR_seqs")))+1, 10)
		den.log10.UTR_lens[[species]] <- density(x)
		ymax.log10.UTR_lens <- c(ymax.log10.UTR_lens, max(den.log10.UTR_lens[[species]]$y))
		xmin.log10.UTR_lens <- c(xmin.log10.UTR_lens, min(den.log10.UTR_lens[[species]]$x))
		xmax.log10.UTR_lens <- c(xmax.log10.UTR_lens, max(den.log10.UTR_lens[[species]]$x))
		x <- log(nchar(get(paste0(species, tissue_dot, ".CDS_seqs")))+1, 10)
		den.log10.CDS_lens[[species]] <- density(x)
		ymax.log10.CDS_lens <- c(ymax.log10.CDS_lens, max(den.log10.CDS_lens[[species]]$y))
		xmin.log10.CDS_lens <- c(xmin.log10.CDS_lens, min(den.log10.CDS_lens[[species]]$x))
		xmax.log10.CDS_lens <- c(xmax.log10.CDS_lens, max(den.log10.CDS_lens[[species]]$x))
	}
}
max_idx.log10.TR <- which(ymax.log10.TR==max(ymax.log10.TR))
max_idx.log10.UTR_lens <- which(ymax.log10.UTR_lens==max(ymax.log10.UTR_lens))
max_idx.log10.CDS_lens <- which(ymax.log10.CDS_lens==max(ymax.log10.CDS_lens))

# plot the distributions
plot(den.log10.TR[[max_idx.log10.TR]], lwd=2, col=species_colors[[max_idx.log10.TR]], xlim=c(min(xmin.log10.TR), max(xmax.log10.TR)), ylim=c(0,max(ymax.log10.TR)), main="Distributions of log10(TR) values")
for (i in 1:m) {
	if (i != max_idx.log10.TR) {
		lines(den.log10.TR[[i]], lwd=2, col=species_colors[[i]])
	} 
}
legend("topleft", paste(paste(names(species_exemplar_tissue_list), unlist(species_exemplar_tissue_list)), paste(num_genes, "genes"), sep=": "), lwd=2, col=unlist(species_colors), bty="n")

plot(den.log10.UTR_lens[[max_idx.log10.UTR_lens]], lwd=2, col=species_colors[[max_idx.log10.UTR_lens]], xlim=c(min(c(xmin.log10.UTR_lens,xmin.log10.CDS_lens)), max(c(xmax.log10.UTR_lens,xmax.log10.CDS_lens))), ylim=c(0,max(ymax.log10.UTR_lens)), main="Distributions of log10(UTR len + 1) values", axes=F, xlab="length (bp)")
axis(1, log(c(0,10,25,50,100,250,500,1000,2500,5000,10000)+1,10), paste(c(0,10,25,50,100,250,500,1000,2500,5000,10000)), las=2)
axis(2)
box()
for (i in 1:m) {
	if (i != max_idx.log10.UTR_lens) {
		lines(den.log10.UTR_lens[[i]], lwd=2, col=species_colors[[i]])
	} 
}
legend("topleft", paste(paste(names(species_exemplar_tissue_list), unlist(species_exemplar_tissue_list)), paste(num_genes, "genes"), sep=": "), lwd=2, col=unlist(species_colors), bty="n")

plot(den.log10.CDS_lens[[max_idx.log10.CDS_lens]], lwd=2, col=species_colors[[max_idx.log10.CDS_lens]], xlim=c(min(c(xmin.log10.UTR_lens,xmin.log10.CDS_lens)), max(c(xmax.log10.UTR_lens,xmax.log10.CDS_lens))), ylim=c(0,max(ymax.log10.CDS_lens)), main="Distributions of log10(CDS len + 1) values", axes=F, xlab="length (bp)")
axis(1, log(c(0,10,25,50,100,250,500,1000,2500,5000,10000)+1,10), paste(c(0,10,25,50,100,250,500,1000,2500,5000,10000)), las=2)
axis(2)
box()
for (i in 1:m) {
	if (i != max_idx.log10.CDS_lens) {
		lines(den.log10.CDS_lens[[i]], lwd=2, col=species_colors[[i]])
	} 
}
legend("topleft", paste(paste(names(species_exemplar_tissue_list), unlist(species_exemplar_tissue_list)), paste(num_genes, "genes"), sep=": "), lwd=2, col=unlist(species_colors), bty="n")
```

# Figure 1 - figure supplement 1. Distributions of log10(TR) in all the tissues
```{r FigureS1, fig.width=5, fig.height=5, fig.path='figures/', dev=c('pdf'), out.width="50%"}
# a list of species and colors
species_tissue_colors <- list(
  sc="#EB9D41",
  sp=list("#94241D", "alt"="#DA3A32"),
  at=list("leaf"="#22549C", "root"="#3881B4", "shoot"="#4CAAE7"),
  mm=list("nih3t3"="#2F6E42", "liver"="#5C9345", "kidney"="#76B958"),
  hs="#CEC448"
)

# number of species
m <- length(unlist(species_tissue_colors))

# number of genes in each tissue of each species
num_genes <- NULL
# distributions of the log10(TR) of each tissue in each species after aligning their median
center.log10.TR <- NULL
for (species in names(species_tissue_list)) {
	for (i in 1:length(species_tissue_list[[species]])) {
		species_tissue <- species_tissue_list[[species]][i]
		if (species_tissue!="") {
			tissue_dot <- paste0(".", species_tissue)
		} else {
			tissue_dot <- ""
		}
		num_genes <- c(num_genes, sum(!is.na(get(paste0(species, tissue_dot, ".log10.TR")))))
		center.log10.TR <- c(center.log10.TR, median(get(paste0(species, tissue_dot, ".log10.TR"))))
	}
}
center.log10.TR <- mean(center.log10.TR)
den.log10.TR <- vector("list", 0)
ymax.log10.TR <- NULL
xmin.log10.TR <- NULL
xmax.log10.TR <- NULL
for (species in names(species_tissue_list)) {
	for (i in 1:length(species_tissue_list[[species]])) {
		species_tissue <- species_tissue_list[[species]][i]
		if (species_tissue!="") {
			tissue_dot <- paste0(".", species_tissue)
		} else {
			tissue_dot <- ""
		}
		name <- paste0(species, tissue_dot)
		x <- get(paste0(species, tissue_dot, ".log10.TR"))
		den.log10.TR[[name]] <- density(x-median(x)+center.log10.TR)
		ymax.log10.TR <- c(ymax.log10.TR, max(den.log10.TR[[name]]$y))
		xmin.log10.TR <- c(xmin.log10.TR, min(den.log10.TR[[name]]$x))
		xmax.log10.TR <- c(xmax.log10.TR, max(den.log10.TR[[name]]$x))
	}
}
max_idx.log10.TR <- which(ymax.log10.TR==max(ymax.log10.TR))

# plot the distributions
plot(den.log10.TR[[max_idx.log10.TR]], lwd=2, col=unlist(species_tissue_colors)[max_idx.log10.TR], xlim=c(min(xmin.log10.TR), max(xmax.log10.TR)), ylim=c(0,max(ymax.log10.TR)), main="Distributions of log10(TR) values")
for (i in 1:m) {
	if (i != max_idx.log10.TR) {
		lines(den.log10.TR[[i]], lwd=2, col=unlist(species_tissue_colors)[i])
	} 
}
legend("topleft", paste(names(unlist(species_tissue_colors)), paste(num_genes, "genes"), sep=": "), lwd=2, col=unlist(species_tissue_colors), bty="n")
```

# Figure 2 - figure supplement 1. Folding energy of the top and bottom 20 translated genes
## the following code only needs to be run once
```{r, eval = FALSE}
# a function to generate the top and bottom genes' folding energy values aligned at AUG
top_bottom_TR_genes_fold_engy_AUG_aligned <- function(n_left, n_right, top_genes, bottom_genes, UTR_folding_energy_all, CDS_folding_energy_all) {
	# n_left: the number of positions on the left of AUG
	# n_right: the number of positions on the right of AUG
	top_results <- sapply(top_genes, FUN=function(x) {
	    c( rev(rev(UTR_folding_energy_all[[x]])[1:n_left]), CDS_folding_energy_all[[x]][1:n_right] )
	})

	bottom_results <- sapply(bottom_genes, FUN=function(x) {
	    c( rev(rev(UTR_folding_energy_all[[x]])[1:n_left]), CDS_folding_energy_all[[x]][1:n_right] )
	})
	
	return(list(top=top_results, bottom=bottom_results))
}

n_left <- 250
n_right <- 50
for (species in names(species_tissue_list)) {
  for (species_tissue in species_tissue_list[[species]]) {
    if (species_tissue!="") {
			tissue_dot <- paste0(".", species_tissue)
			tissue_us <- paste0("_", species_tissue)
		} else {
			tissue_dot <- ""
			tissue_us <- ""
		}
    UTR_seqs <- get(paste0(species, tissue_dot, ".UTR_seqs"))
		log10.UTR_len <- log(nchar(UTR_seqs), 10)

    folding_energy_all <- readRDS(paste0("processed_data_w_polyA/35_win_fold_UTR5_CDSfull_all/", species, tissue_us, "_UTR5_full_CDS_window_35.rds"))
    UTR_folding_energy_all <- folding_energy_all$UTR_folding_energy_all
    CDS_folding_energy_all <- folding_energy_all$CDS_folding_energy_all
    rm(folding_energy_all)
    gc()
    
    log10.TR <- get(paste0(species, tissue_dot, ".log10.TR"))
    log10.tmp <- sort(log10.TR[log10.UTR_len<=log(n_left,10)])
    TR_top_20_genes <- names(rev(log10.tmp)[1:20])
    TR_bottom_20_genes <- names(log10.tmp[1:20])
    
    fold_engy_AUG_aligned <- top_bottom_TR_genes_fold_engy_AUG_aligned(n_left=n_left, n_right=n_right, top_genes=TR_top_20_genes, bottom_genes=TR_bottom_20_genes, UTR_folding_energy_all, CDS_folding_energy_all)
    TR_top_20_heatmap <- fold_engy_AUG_aligned$top
    TR_bottom_20_heatmap <- fold_engy_AUG_aligned$bottom
    rm(fold_engy_AUG_aligned)
    
    labels <- as.character( c((-n_left):(-1), 1:n_right) )
    breakpoints <- c( seq(1, n_left, by=10), n_left, n_left+1, rev(seq(n_left+n_right, n_left+1, by=-10)) )
    labels[-breakpoints] <- ""
    
    rownames(TR_top_20_heatmap) <- labels
    rownames(TR_bottom_20_heatmap) <- labels
    
    saveRDS(TR_top_20_heatmap, file=paste0("processed_data_w_polyA/TR_top_bottom_20_heatmaps/", species, tissue_dot, ".TR_top_20_heatmap.rds"))
    saveRDS(TR_bottom_20_heatmap, file=paste0("processed_data_w_polyA/TR_top_bottom_20_heatmaps/", species, tissue_dot, ".TR_bottom_20_heatmap.rds"))
  }
}
```
## plot
```{r FigureS2, fig.width=7, fig.height=5, fig.path='figures/', dev=c('pdf'), out.width="50%"}
n_left <- 250
n_right <- 50
breakpoints <- c( seq(1, n_left, by=10), n_left, n_left+1, rev(seq(n_left+n_right, n_left+1, by=-10)) )

for (species in names(species_tissue_list)) {
  for (species_tissue in species_tissue_list[[species]]) {
    if (species_tissue!="") {
			tissue_dot <- paste0(".", species_tissue)
			tissue_us <- paste0("_", species_tissue)
		} else {
			tissue_dot <- ""
			tissue_us <- ""
		}
    TR_top_20_heatmap <- readRDS(file=paste0("processed_data_w_polyA/TR_top_bottom_20_heatmaps/", species, tissue_dot, ".TR_top_20_heatmap.rds"))
    TR_bottom_20_heatmap <- readRDS(file=paste0("processed_data_w_polyA/TR_top_bottom_20_heatmaps/", species, tissue_dot, ".TR_bottom_20_heatmap.rds"))
    heatmap.range <- range(c(TR_top_20_heatmap, TR_bottom_20_heatmap), na.rm=T)
    print(
      levelplot(TR_top_20_heatmap, scales=list(x=list(at=breakpoints, rot=90, cex=.3)), at=seq(heatmap.range[1], heatmap.range[2], length.out=31), main=paste0(species, tissue_dot, " TR top 20 genes"), col.regions=heat.colors(120))
    )
    print(
      levelplot(TR_bottom_20_heatmap, scales=list(x=list(at=breakpoints, rot=90, cex=.3)), at=seq(heatmap.range[1], heatmap.range[2], length.out=31), main=paste0(species, tissue_dot, " TR bottom 20 genes"), col.regions=heat.colors(120))
    )
  }
}
```

# Figure 3. R2s of RNA folding energy features
## the following code only needs to be run once
```{r, eval = FALSE}
for (species in names(species_tissue_list)) {
	for (species_tissue in species_tissue_list[[species]]) {
		if (species_tissue!="") {
			tissue_dot <- paste0(".", species_tissue)
			tissue_us <- paste0("_", species_tissue)
		} else {
			tissue_dot <- ""
			tissue_us <- ""
		}
		
		load(file=paste0("processed_data_w_polyA/", species, tissue_dot, ".Rdata"))
		fold_energy_features_selected <- readRDS(file=paste0("processed_data_w_polyA/folding_energy_features_selected/", species, tissue_us, "_UTR_folding_energy_features_selected.rds"))
		
		all_features <- fold_energy_features_selected$all_features
		if (species != "sp") {
		  selected_feature_R2 <- fold_energy_features_selected$selected_feature_R2
		} else {
		  selected_feature_R2 <- readRDS(file=paste0("processed_data_w_polyA/folding_energy_features_selected/", species, tissue_us, "_UTR_folding_energy_features_selected_trun.rds"))$selected_feature_R2
		}
		
		
		feature_names <- c("5'most", "5'mostTICE", "-1window", "min", "10%", "25%", "75%", "90%", "max", "mean", "%<=20%", "sum<=5%", "sum<=10%", "sum<=20%", "%>90%", "%>80%", "sum>90%", "sum>80%", "model all-1", "whole", "model all", "RNA fold")
		names(feature_names) <- c("5' cap", "5'mostTICE", "-1", "min", "10%", "25%", "75%", "90%", "max", "mean", "%<=20%", "sum<=5%", "sum<=10%", "sum<=20%", "%>90%", "%>80%", "sum>90%", "sum>80%", "model all-1", "whole", "model all", "RNA fold")
		
		feature_window_R2s <- matrix(NA, nrow=length(window_lens), ncol=length(feature_names)) # a matrix to store the results
		
		rownames(feature_window_R2s) <- window_lens
		colnames(feature_window_R2s) <- names(feature_names)
		
		for (j in 1:length(feature_names)) {
		  feature_name <- feature_names[j]
		  if (!(feature_name %in% c("whole", "RNA fold"))) { # these two features do not vary with window lengths
		    for (i in 1:length(window_lens)) {
		      window_len <- window_lens[i]
		      if (!(feature_name %in% c("model all-1", "model all"))) { # single features
		        tmp1 <- which(colnames(all_features)==paste0(feature_name, "_", window_len))
		        tmp2 <- which(grepl(paste0("^", feature_name, "_", window_len, "_"), colnames(all_features)))
		        if (length(tmp1)>0 & length(tmp2)==0) {
		          idx <- c( tmp1, which(grepl("Intercept", colnames(all_features))) )
		        } else if (length(tmp1)==0 & length(tmp2)>0) {
		          idx <- c( tmp2, which(grepl("Intercept", colnames(all_features))) )
		        } else {
		          idx <- NULL
		        }
		      } else if (feature_name == "model all-1") {
		        if (length(tmp1)>0 & length(tmp2)==0) {
		          idx <- c( which(grepl(paste0("_", window_len, "$"), colnames(all_features))), which(grepl("Intercept", colnames(all_features))) )
		        } else if (length(tmp1)==0 & length(tmp2)>0) {
		          idx <- c( which(grepl(paste0("_", window_len, "_"), colnames(all_features))), which(grepl("Intercept", colnames(all_features))) )
		        } else {
		          idx <- NULL
		        }
		      } else if (feature_name == "model all") {
		        if (length(tmp1)>0 & length(tmp2)==0) {
		          idx <- c( which(grepl(paste0("_", window_len, "$"), colnames(all_features))), which(grepl("Intercept", colnames(all_features))), which(grepl("whole", colnames(all_features))) )
		        } else if (length(tmp1)==0 & length(tmp2)>0) {
		          idx <- c( which(grepl(paste0("_", window_len, "_"), colnames(all_features))), which(grepl("Intercept", colnames(all_features))), which(grepl("whole", colnames(all_features))) )
		        } else {
		          idx <- NULL
		        }
		      }
		      features <- all_features[,idx]
		      if (length(idx) > 0) {
		        feature_window_R2s[i,j] <- summary(lm(get(paste0(species, tissue_dot, ".log10.TR"))~as.matrix(features)))$r.squared
		      } else {
		        feature_window_R2s[i,j] <- 0
		      }
		    }
		  } else if (feature_name == "whole") {
		    idx <- c( which(grepl("whole", colnames(all_features))), which(grepl("Intercept", colnames(all_features))) )
		    features <- all_features[,idx]
		    feature_window_R2s[,j] <- summary(lm(get(paste0(species, tissue_dot, ".log10.TR"))~as.matrix(features)))$r.squared
		  } else { # feature == "RNA fold" the BIC selected ones
		    feature_window_R2s[,j] <- selected_feature_R2
		  }
		}
		write.csv(t(feature_window_R2s), file=paste0("processed_data_w_polyA/folding_energy_feature_R2s/", species, ".", species_tissue, ".csv"), quote=F)
		saveRDS(feature_window_R2s, file=paste0("processed_data_w_polyA/folding_energy_feature_R2s/", species, ".", species_tissue, ".rds"))
	}
}
```
## plot
```{r Figure3, fig.width=7, fig.height=3, fig.path='figures/', dev=c('pdf')}
# correlation between each RNA folding energy and log10(TR)
for (species in names(species_tissue_list)) {
	for (species_tissue in species_tissue_list[[species]]) {
	  feature_window_R2s <- readRDS(file=paste0("processed_data_w_polyA/folding_energy_feature_R2s/", species, ".", species_tissue, ".rds"))
		# make a histogram
		window_cols <- c("#4F71BE", "#DF8344", "#A5A5A5", "#F6C243", "#6A9AD0", "#7EAB55", "#2D4474", "#944D20", "#636363", "#937524", "#355D8D", "#4B6733", "#718DCB", "#E59B65", "#B7B7B7", "#F7CF55", "#86ADD9", "#97C072", "#3C599C", "#C4672C", "#848484", "#000000")
		names(window_cols) <- c(window_lens, "NA")
		feature_window_R2s_for_plot <- setNames(melt(feature_window_R2s), c("Window_length", "Feature", 'R2'))
		feature_window_R2s_for_plot[,1] <- as.factor(feature_window_R2s_for_plot[,1])
		
		theme_set(theme_bw())
		print( ggplot(feature_window_R2s_for_plot, aes(x=Feature, y=R2, fill=Window_length)) + geom_bar(stat="identity", position=position_dodge()) + scale_fill_manual(values=window_cols) + theme(axis.text.x = element_text(angle = 45, vjust = 1, hjust=1)) + ggtitle(paste0(species, ".", species_tissue)) )
	}
}
```

# Figure 5. Pearson correlation(s) between min fold energy window feature(s) and log(TR) 
## the following code only needs to be run once
```{r, eval = FALSE}
min_fold_energy_window_feature_cors <- NULL
rownames_min_fold_energy_window_feature_cors <- NULL
colnames_min_fold_energy_window_feature_cors <- c("contig stem", "max stem", "all")

for (species in names(species_tissue_list)) {
	for (species_tissue in species_tissue_list[[species]]) {
		if (species_tissue!="") {
			tissue_dot <- paste0(".", species_tissue)
			tissue_us <- paste0("_", species_tissue)
		} else {
			tissue_dot <- ""
			tissue_us <- ""
		}
	  min_fold_energy_window_features <- read.table(paste0("data/min_window_seq_parameters/", species, tissue_us, ".tsv"), header=T, sep="\t", stringsAsFactors=FALSE)
	  
	  # feature 1
	  max_continuous_stem <- min_fold_energy_window_features$max_continuous_stem
	  names(max_continuous_stem) <- min_fold_energy_window_features$id
	  cor_max_continuous_stem <- cor(get(paste0(species, tissue_dot, ".log10.TR")), max_continuous_stem[get(paste0(species, tissue_dot, ".genes"))], use="complete.obs")
	  
	  # feature 2
	  max_stem_paired <- min_fold_energy_window_features$max_stem_paired
	  names(max_stem_paired) <- min_fold_energy_window_features$id
	  cor_max_stem_paired <- cor(get(paste0(species, tissue_dot, ".log10.TR")), max_stem_paired[get(paste0(species, tissue_dot, ".genes"))], use="complete.obs")
	  
	  # feature 3
	  paired_nts <- min_fold_energy_window_features$paired_nts
	  names(paired_nts) <- min_fold_energy_window_features$id
	  cor_paired_nts <- cor(get(paste0(species, tissue_dot, ".log10.TR")), paired_nts[get(paste0(species, tissue_dot, ".genes"))], use="complete.obs")
	  
	  min_fold_energy_window_feature_cors <- rbind(min_fold_energy_window_feature_cors,
	                                               c(cor_max_continuous_stem, 
	                                                 cor_max_stem_paired, 
	                                                 cor_paired_nts)
	                                               )
	  rownames_min_fold_energy_window_feature_cors <- c(
	    rownames_min_fold_energy_window_feature_cors,
	    paste0(species, tissue_dot)
	  )
	}
}
rownames(min_fold_energy_window_feature_cors) <- rownames_min_fold_energy_window_feature_cors
colnames(min_fold_energy_window_feature_cors) <- colnames_min_fold_energy_window_feature_cors

write.csv(min_fold_energy_window_feature_cors, file="tables/min_fold_energy_window_feature_cors.csv", quote=F)
```
## plot
```{r Figure5, fig.width=7, fig.height=3, fig.path='figures/', dev=c('pdf')}
min_fold_energy_window_feature_cors <- read.csv(file="tables/min_fold_energy_window_feature_cors.csv")
colnames(min_fold_energy_window_feature_cors)[1] <- "species.tissue"

min_fold_energy_window_feature_cols <- c("#AECDE1", "#447DB2", "#386895")
min_fold_energy_window_feature_cors_for_plot <- setNames(melt(min_fold_energy_window_feature_cors), c("Species_Tissue", "Feature", 'Pearson_Cor'))

theme_set(theme_bw())
print( ggplot(min_fold_energy_window_feature_cors_for_plot, aes(x=Species_Tissue, y=Pearson_Cor, fill=Feature)) + geom_bar(stat="identity", position=position_dodge()) + scale_fill_manual(values=min_fold_energy_window_feature_cols) + theme(axis.text.x = element_text(angle = 45, vjust = 1, hjust=1)) + ggtitle("min fold energy window features") )		
```

# Figure 6. Difference(s) of min fold energy window feature(s) between top TR and bottom TR genes 
## the following code only needs to be run once
```{r, eval = FALSE}
min_fold_energy_window_feature_top_bottom_diffs <- NULL
rownames_min_fold_energy_window_feature_top_bottom_diffs <- NULL
colnames_min_fold_energy_window_feature_top_bottom_diffs <- c("MFE", "TR", "loop", "contig stem", "max stem", "all")

for (species in names(species_tissue_list)) {
	for (species_tissue in species_tissue_list[[species]]) {
		if (species_tissue!="") {
			tissue_dot <- paste0(".", species_tissue)
			tissue_us <- paste0("_", species_tissue)
		} else {
			tissue_dot <- ""
			tissue_us <- ""
		}
	  min_fold_energy_window_features <- read.table(paste0("data/min_window_seq_parameters/", species, tissue_us, ".tsv"), header=T, sep="\t", stringsAsFactors=FALSE)
	  
	  # feature 1
	  mfe <- min_fold_energy_window_features$mfe
	  names(mfe) <- min_fold_energy_window_features$id
	  diff_mfe <- mean(mfe[get(paste0(species, tissue_dot, ".TR_bottom_genes"))], na.rm=T) - mean(mfe[get(paste0(species, tissue_dot, ".TR_top_genes"))], na.rm=T)
	  
	  # feature 2
	  log10.TR <- get(paste0(species, tissue_dot, ".log10.TR"))
	  diff_log10.TR <- mean(log10.TR[get(paste0(species, tissue_dot, ".TR_bottom_genes"))], na.rm=T) - mean(log10.TR[get(paste0(species, tissue_dot, ".TR_top_genes"))], na.rm=T)
	  
	  # feature 3
	  max_loop <- min_fold_energy_window_features$max_loop
	  names(max_loop) <- min_fold_energy_window_features$id
	  diff_max_loop <- mean(max_loop[get(paste0(species, tissue_dot, ".TR_bottom_genes"))], na.rm=T) - mean(max_loop[get(paste0(species, tissue_dot, ".TR_top_genes"))], na.rm=T)
	  
	  # feature 4
	  max_continuous_stem <- min_fold_energy_window_features$max_continuous_stem
	  names(max_continuous_stem) <- min_fold_energy_window_features$id
	  diff_max_continuous_stem <- mean(max_continuous_stem[get(paste0(species, tissue_dot, ".TR_bottom_genes"))], na.rm=T) - mean(max_continuous_stem[get(paste0(species, tissue_dot, ".TR_top_genes"))], na.rm=T)
	  
	  # feature 5
	  max_stem_paired <- min_fold_energy_window_features$max_stem_paired
	  names(max_stem_paired) <- min_fold_energy_window_features$id
	  diff_max_stem_paired <- mean(max_stem_paired[get(paste0(species, tissue_dot, ".TR_bottom_genes"))], na.rm=T) - mean(max_stem_paired[get(paste0(species, tissue_dot, ".TR_top_genes"))], na.rm=T)
	  
	  # feature 6
	  paired_nts <- min_fold_energy_window_features$paired_nts
	  names(paired_nts) <- min_fold_energy_window_features$id
	  diff_paired_nts <- mean(paired_nts[get(paste0(species, tissue_dot, ".TR_bottom_genes"))], na.rm=T) - mean(paired_nts[get(paste0(species, tissue_dot, ".TR_top_genes"))], na.rm=T)
	  
	  min_fold_energy_window_feature_top_bottom_diffs <- rbind(min_fold_energy_window_feature_top_bottom_diffs,
	                                               c(diff_mfe,
	                                                 diff_log10.TR,
	                                                 diff_max_loop,
	                                                 diff_max_continuous_stem, 
	                                                 diff_max_stem_paired, 
	                                                 diff_paired_nts)
	                                               )
	  rownames_min_fold_energy_window_feature_top_bottom_diffs <- c(
	    rownames_min_fold_energy_window_feature_top_bottom_diffs,
	    paste0(species, tissue_dot)
	  )
	}
}
rownames(min_fold_energy_window_feature_top_bottom_diffs) <- rownames_min_fold_energy_window_feature_top_bottom_diffs
colnames(min_fold_energy_window_feature_top_bottom_diffs) <- colnames_min_fold_energy_window_feature_top_bottom_diffs

write.csv(min_fold_energy_window_feature_top_bottom_diffs, file="tables/min_fold_energy_window_feature_top_bottom_diffs.csv", quote=F)
```
## plot
```{r Figure6, fig.width=7, fig.height=3, fig.path='figures/', dev=c('pdf')}
min_fold_energy_window_feature_top_bottom_diffs <- read.csv(file="tables/min_fold_energy_window_feature_top_bottom_diffs.csv")
colnames(min_fold_energy_window_feature_top_bottom_diffs)[1] <- "species.tissue"

# only keep the exemplar tissues
min_fold_energy_window_feature_top_bottom_diffs_for_plot <- setNames(melt(min_fold_energy_window_feature_top_bottom_diffs[min_fold_energy_window_feature_top_bottom_diffs$species.tissue %in% species_exemplar_tissue_names,]), c("Species_Tissue", "Feature", 'Bottom_minus_Top_Diff'))

min_fold_energy_window_feature_top_bottom_diffs_for_plot$Species_Tissue <- factor(min_fold_energy_window_feature_top_bottom_diffs_for_plot$Species_Tissue, levels=c("sc", "sp.alt", "at.leaf", "mm.nih3t3", "hs.hela2"))

theme_set(theme_bw())
print( ggplot(min_fold_energy_window_feature_top_bottom_diffs_for_plot, aes(x=Feature, y=Bottom_minus_Top_Diff, fill=Species_Tissue)) + geom_bar(stat="identity", position=position_dodge()) + scale_fill_manual(values=species_cols[c("sc", "sp.alt", "at.leaf", "mm.nih3t3", "hs.hela2")]) + theme(axis.text.x = element_text(angle = 45, vjust = 1, hjust=1)) + ggtitle("min fold energy window features (TR bottom - TR top)") )		
```

# Figure 6 - figure supplement 1. Distribution(s) of the number(s) of paired nucleotides in high/low MFE genes or high/low TR genes
```{r FigureS6, fig.width=5, fig.height=5, fig.path='figures/', dev=c('pdf'), out.width="50%"}
#my_plot_hook <- function(x, options)
#  paste("\n", knitr::hook_plot_tex(x, options), "\n")
#knitr::knit_hooks$set(plot = my_plot_hook)

myred <- "#88342D"

for (species in names(species_tissue_list)) {
	for (species_tissue in species_tissue_list[[species]]) {
		if (species_tissue!="") {
			tissue_dot <- paste0(".", species_tissue)
			tissue_us <- paste0("_", species_tissue)
		} else {
			tissue_dot <- ""
			tissue_us <- ""
		}
	  min_fold_energy_window_features <- read.table(paste0("data/min_window_seq_parameters/", species, tissue_us, ".tsv"), header=T, sep="\t", stringsAsFactors=FALSE)
	  
	  mfe_quantiles <- quantile(min_fold_energy_window_features$mfe, prob=c(0.1, 0.9))
	  
	  for (feature_name in c("paired_nts")) {
		  feature <- min_fold_energy_window_features[,feature_name]
		  names(feature) <- min_fold_energy_window_features$id
		  
		  density1 <- density(feature[min_fold_energy_window_features$mfe>=mfe_quantiles[2]], bw=1, na.rm=TRUE)
		  density2 <- density(feature[min_fold_energy_window_features$mfe<=mfe_quantiles[1]], bw=1, na.rm=TRUE)
		  density3 <- density(feature[get(paste0(species, tissue_dot, ".TR_top_genes"))], bw=1, na.rm=TRUE)
		  density4 <- density(feature[get(paste0(species, tissue_dot, ".TR_bottom_genes"))], bw=1, na.rm=TRUE)
		  
		  mean1 <- mean(feature[min_fold_energy_window_features$mfe>=mfe_quantiles[2]], na.rm=TRUE)
		  mean2 <- mean(feature[min_fold_energy_window_features$mfe<=mfe_quantiles[1]], na.rm=TRUE)
		  mean3 <- mean(feature[get(paste0(species, tissue_dot, ".TR_top_genes"))], na.rm=TRUE)
		  mean4 <- mean(feature[get(paste0(species, tissue_dot, ".TR_bottom_genes"))], na.rm=TRUE)
		  
		  plot(0,0,type="n",xlim=range(c(density1$x,density2$x,density3$x,density4$x)), ylim=range(c(density1$y,density2$y,density3$y,density4$y)), xaxt='n', xlab=feature, ylab="Density", main=paste0(species, tissue_dot))
		  axis(1)
		  lines(density1, col=1, lwd=3)
		  abline(v=mean1, lty=2, col=1, lwd=3)
		  lines(density2, col=myred, lwd=3)
		  abline(v=mean2, lty=2, col=myred, lwd=3)
		  legend("topright", c("Top MFE", "Bottom MFE"), col=c(1, myred), lty=c(1,1), bty="n")
		  arrows(x0=mean1, y0=0, x1=mean2, y1=0, col=1, length=0.05, code=3)
		  text(x=(mean1+mean2)/2, y=0.01, labels=round(mean2-mean1, 1))
		  text(x=mean1-1, y=0.02, labels=round(mean1, 1))
		  text(x=mean2+1, y=0.02, labels=round(mean2, 1), col=myred)
	
		  plot(0,0,type="n",xlim=range(c(density1$x,density2$x,density3$x,density4$x)), ylim=range(c(density1$y,density2$y,density3$y,density4$y)), xaxt='n', xlab=feature, ylab="Density", main=paste0(species, tissue_dot))
		  axis(1)
		  lines(density3, col=1, lwd=3)
		  abline(v=mean3, lty=2, col=1, lwd=3)
		  lines(density4, col=myred, lwd=3)
		  abline(v=mean4, lty=2, col=myred, lwd=3)
		  legend("topright", c("Top TR", "Bottom TR"), col=c(1, myred), lty=c(1,1), bty="n")
		  arrows(x0=mean3, y0=0, x1=mean4, y1=0, col=1, length=0.05, code=3)
		  text(x=(mean3+mean4)/2, y=0.01, labels=round(mean4-mean3, 1))
		  text(x=mean3-1, y=0.02, labels=round(mean3, 1))
		  text(x=mean4+1, y=0.02, labels=round(mean4, 1), col=myred)
	  }
	}
}
```

# Figure 7. The 5' and 3' boundaries of TICE.
```{r, eval = FALSE}
for (species in names(species_exemplar_tissue_list)) {
	for (species_tissue in species_exemplar_tissue_list[[species]]) {
		if (species_tissue!="") {
			species.genes <- get(paste(species, species_tissue, "genes", sep="."))
			species.UTR_seqs <- get(paste(species, species_tissue, "UTR_seqs", sep="."))
			species.CDS_seqs <- get(paste(species, species_tissue, "CDS_seqs", sep="."))
			species.log10.TR <- get(paste(species, species_tissue, "log10.TR", sep="."))
			species.TR_top_genes <- get(paste(species, species_tissue, "TR_top_genes", sep="."))
			species.TR_bottom_genes <- get(paste(species, species_tissue, "TR_bottom_genes", sep="."))
			connector <- paste0(species_tissue, ".")
		} else {
			species.genes <- get(paste(species, "genes", sep="."))
			species.UTR_seqs <- get(paste(species, "UTR_seqs", sep="."))
			species.CDS_seqs <- get(paste(species, "CDS_seqs", sep="."))
			species.log10.TR <- get(paste(species, "log10.TR", sep="."))
			species.TR_top_genes <- get(paste(species, "TR_top_genes", sep="."))
			species.TR_bottom_genes <- get(paste(species, "TR_bottom_genes", sep="."))
			connector <- ""
		}
		species.log10.UTR_len <- log(nchar(species.UTR_seqs), 10)
		if (species != "hs") {
			species.uTICE_len <- species_uTICE_len_list[[species]]
		} else {
			species.uTICE_len <- species_uTICE_len_list[[species]][[species_tissue]]
		}
		
		## for the genes in the top and bottom cohort, find the 5' UTR sequences and CDS sequences
		species.TR_top_UTR_seqs <- species.UTR_seqs[species.TR_top_genes]
		species.TR_top_CDS_seqs <- species.CDS_seqs[species.TR_top_genes]
				
		species.TR_bottom_UTR_seqs <- species.UTR_seqs[species.TR_bottom_genes]
		species.TR_bottom_CDS_seqs <- species.CDS_seqs[species.TR_bottom_genes]
		
		# construct Position Weight Matrices (PWMs) based on the top 10% TR genes with different 5' UTR lengths and ORF lengths
		max_UTR_len <- 100
		len_breaks <- seq(5, max_UTR_len, by=5) # 5' UTR lengths vary from 100 bp to 5 bp
		
		## top TR
		species.TR_top_PWM_max_UTR_len <- UTR_pwm(UTR_seqs=species.TR_top_UTR_seqs, n=max_UTR_len) # no CDS length
		
		species.TR_top_PWM_max_UTR_len_CDS_13 <- UTR_CDS_pwm(UTR_seqs=species.TR_top_UTR_seqs, CDS_seqs=species.TR_top_CDS_seqs, n_UTR=max_UTR_len, n_CDS=13) # 13 bp CDS
		colnames(species.TR_top_PWM_max_UTR_len_CDS_13) <- c("A", "C", "G", "T")

		species.TR_top_PWM_max_UTR_len_CDS_23 <- UTR_CDS_pwm(UTR_seqs=species.TR_top_UTR_seqs, CDS_seqs=species.TR_top_CDS_seqs, n_UTR=max_UTR_len, n_CDS=23) # 23 bp CDS
		colnames(species.TR_top_PWM_max_UTR_len_CDS_23) <- c("A", "C", "G", "T")

		species.TR_top_PWM_max_UTR_len_CDS_28 <- UTR_CDS_pwm(UTR_seqs=species.TR_top_UTR_seqs, CDS_seqs=species.TR_top_CDS_seqs, n_UTR=max_UTR_len, n_CDS=28) # 28 bp CDS
		colnames(species.TR_top_PWM_max_UTR_len_CDS_28) <- c("A", "C", "G", "T")

		species.TR_top_PWM_max_UTR_len_CDS_33 <- UTR_CDS_pwm(UTR_seqs=species.TR_top_UTR_seqs, CDS_seqs=species.TR_top_CDS_seqs, n_UTR=max_UTR_len, n_CDS=33) # 33 bp CDS
		colnames(species.TR_top_PWM_max_UTR_len_CDS_33) <- c("A", "C", "G", "T")
		
		saveRDS(species.TR_top_PWM_max_UTR_len_CDS_33, file=paste0("processed_data_w_polyA/AUGPWM_R2s/", species, ".", connector, "TR_top_PWM_AUG_aligned.rds"))

		## bottom TR
		species.TR_bottom_PWM_max_UTR_len <- UTR_pwm(UTR_seqs=species.TR_bottom_UTR_seqs, n=max_UTR_len) # no CDS length
		
		species.TR_bottom_PWM_max_UTR_len_CDS_13 <- UTR_CDS_pwm(UTR_seqs=species.TR_bottom_UTR_seqs, CDS_seqs=species.TR_bottom_CDS_seqs, n_UTR=max_UTR_len, n_CDS=13) # 13 bp CDS
		colnames(species.TR_bottom_PWM_max_UTR_len_CDS_13) <- c("A", "C", "G", "T")

		species.TR_bottom_PWM_max_UTR_len_CDS_23 <- UTR_CDS_pwm(UTR_seqs=species.TR_bottom_UTR_seqs, CDS_seqs=species.TR_bottom_CDS_seqs, n_UTR=max_UTR_len, n_CDS=23) # 23 bp CDS
		colnames(species.TR_bottom_PWM_max_UTR_len_CDS_23) <- c("A", "C", "G", "T")

		species.TR_bottom_PWM_max_UTR_len_CDS_28 <- UTR_CDS_pwm(UTR_seqs=species.TR_bottom_UTR_seqs, CDS_seqs=species.TR_bottom_CDS_seqs, n_UTR=max_UTR_len, n_CDS=28) # 28 bp CDS
		colnames(species.TR_bottom_PWM_max_UTR_len_CDS_28) <- c("A", "C", "G", "T")

		species.TR_bottom_PWM_max_UTR_len_CDS_33 <- UTR_CDS_pwm(UTR_seqs=species.TR_bottom_UTR_seqs, CDS_seqs=species.TR_bottom_CDS_seqs, n_UTR=max_UTR_len, n_CDS=33) # 33 bp CDS
		colnames(species.TR_bottom_PWM_max_UTR_len_CDS_33) <- c("A", "C", "G", "T")
		
		saveRDS(species.TR_bottom_PWM_max_UTR_len_CDS_33, file=paste0("processed_data_w_polyA/AUGPWM_R2s/", species, ".", connector, "TR_bottom_PWM_AUG_aligned.rds"))
		
	
		# PWM scores (aligned at AUG)

		## top TR
		### varying 5' UTR lengths, 0 bp CDS length
		filename <- paste0("processed_data_w_polyA/AUGPWM_R2s/", species, ".", connector, "TR_top_PWM_max_UTR_len_score.rds")
		if (!file.exists(filename)) {
		  species.TR_top_PWM_max_UTR_len_score <- sapply(len_breaks, FUN=function(i){
		    UTR_pwm_score( species.TR_top_PWM_max_UTR_len[(i-5+1):nrow(species.TR_top_PWM_max_UTR_len),], species.UTR_seqs )
		  })
		  saveRDS(species.TR_top_PWM_max_UTR_len_score, file=filename)
		}

		### varying 5' UTR lengths, 13 bp CDS length
		filename <- paste0("processed_data_w_polyA/AUGPWM_R2s/", species, ".", connector, "TR_top_PWM_max_UTR_len_CDS_13_score.rds")
		if (!file.exists(filename)) {
		  species.UTR_CDS_13_seqs <- sapply(species.genes, FUN=function(x) {
		    paste( species.UTR_seqs[x], paste(strsplit(species.CDS_seqs[x], "")[[1]][4:13], collapse=""), sep="" )
		  })
		  species.TR_top_PWM_max_UTR_len_CDS_13_score <- sapply(len_breaks, FUN=function(i){
		    UTR_pwm_score( species.TR_top_PWM_max_UTR_len_CDS_13[(i-5+1):nrow(species.TR_top_PWM_max_UTR_len_CDS_13),], species.UTR_CDS_13_seqs )
		  })
		  saveRDS(species.TR_top_PWM_max_UTR_len_CDS_13_score, file=filename)
		}

		### varying 5' UTR lengths, 23 bp CDS length
		filename <- paste0("processed_data_w_polyA/AUGPWM_R2s/", species, ".", connector, "TR_top_PWM_max_UTR_len_CDS_23_score.rds")
		if (!file.exists(filename)) {
		  species.UTR_CDS_23_seqs <- sapply(species.genes, FUN=function(x) {
		    paste( species.UTR_seqs[x], paste(strsplit(species.CDS_seqs[x], "")[[1]][4:23], collapse=""), sep="" )
		  })
		  species.TR_top_PWM_max_UTR_len_CDS_23_score <- sapply(len_breaks, FUN=function(i){
		    UTR_pwm_score( species.TR_top_PWM_max_UTR_len_CDS_23[(i-5+1):nrow(species.TR_top_PWM_max_UTR_len_CDS_23),], species.UTR_CDS_23_seqs )
		  })
		  saveRDS(species.TR_top_PWM_max_UTR_len_CDS_23_score, file=filename)
		}

		### varying 5' UTR lengths, 28 bp CDS length
		filename <- paste0("processed_data_w_polyA/AUGPWM_R2s/", species, ".", connector, "TR_top_PWM_max_UTR_len_CDS_28_score.rds")
		if (!file.exists(filename)) {
		  species.UTR_CDS_28_seqs <- sapply(species.genes, FUN=function(x) {
		    paste( species.UTR_seqs[x], paste(strsplit(species.CDS_seqs[x], "")[[1]][4:28], collapse=""), sep="" )
		  })
		  species.TR_top_PWM_max_UTR_len_CDS_28_score <- sapply(len_breaks, FUN=function(i){
		    UTR_pwm_score( species.TR_top_PWM_max_UTR_len_CDS_28[(i-5+1):nrow(species.TR_top_PWM_max_UTR_len_CDS_28),], species.UTR_CDS_28_seqs )
		  })
		  saveRDS(species.TR_top_PWM_max_UTR_len_CDS_28_score, file=filename)
		}

		### varying 5' UTR lengths, 33 bp CDS length
		filename <- paste0("processed_data_w_polyA/AUGPWM_R2s/", species, ".", connector, "TR_top_PWM_max_UTR_len_CDS_33_score.rds")
		if (!file.exists(filename)) {
		  species.UTR_CDS_33_seqs <- sapply(species.genes, FUN=function(x) {
		    paste( species.UTR_seqs[x], paste(strsplit(species.CDS_seqs[x], "")[[1]][4:33], collapse=""), sep="" )
		  })
		  species.TR_top_PWM_max_UTR_len_CDS_33_score <- sapply(len_breaks, FUN=function(i){
		    UTR_pwm_score( species.TR_top_PWM_max_UTR_len_CDS_33[(i-5+1):nrow(species.TR_top_PWM_max_UTR_len_CDS_33),], species.UTR_CDS_33_seqs )
		  })
		  saveRDS(species.TR_top_PWM_max_UTR_len_CDS_33_score, file=filename)
		} 
	}
}
```

```{r Figure7, fig.width=5, fig.height=5, fig.path='figures/', dev=c('pdf'), out.width="50%"}
# a function to plot a UTR-CDS PWM
plot_UTR_CDS_pwm <- function(PWM, n_UTR, n_CDS, title) {
	PWM_plot <- rbind(
	  PWM[1:n_UTR,],
	  c(1,0,0,0),
	  c(0,0,0,1),
	  c(0,0,1,0),
	  PWM[(n_UTR+1):nrow(PWM),]
	)
	colnames(PWM_plot) <- c("A", "C", "G", "U")
	PWM_motif <- new("pcm", mat=t(PWM_plot), name=title)
	plot(PWM_motif, ic.scale=F)
}

max_UTR_len <- 100
len_breaks <- seq(5, max_UTR_len, by=5) # 5' UTR lengths vary from 100 bp to 5 bp

for (species in names(species_exemplar_tissue_list)) {
	for (species_tissue in species_exemplar_tissue_list[[species]]) {
	    if (species_tissue!="") {
			  species.log10.TR <- get(paste(species, species_tissue, "log10.TR", sep="."))
			  species.UTR_seqs <- get(paste(species, species_tissue, "UTR_seqs", sep="."))
			  connector <- paste0(species_tissue, ".")
		  } else {
			  species.log10.TR <- get(paste(species, "log10.TR", sep="."))
			  species.UTR_seqs <- get(paste(species, "UTR_seqs", sep="."))
			  connector <- ""
		  }
	    species.log10.UTR_len <- log(nchar(species.UTR_seqs), 10)
	    
	  	species.TR_top_PWM_AUG_aligned <- readRDS(file=paste0("processed_data_w_polyA/AUGPWM_R2s/", species, ".", connector, "TR_top_PWM_AUG_aligned.rds"))
	  	species.TR_bottom_PWM_AUG_aligned <- readRDS(file=paste0("processed_data_w_polyA/AUGPWM_R2s/", species, ".", connector, "TR_bottom_PWM_AUG_aligned.rds"))
	 	  ## plot the PWM matrices
	 	  plot_UTR_CDS_pwm(species.TR_top_PWM_AUG_aligned[21:nrow(species.TR_top_PWM_AUG_aligned),], n_UTR=80, n_CDS=33, title=paste0(species, ".", connector, "TR_top_PWM"))
	 	  plot_UTR_CDS_pwm(species.TR_bottom_PWM_AUG_aligned[21:nrow(species.TR_bottom_PWM_AUG_aligned),], n_UTR=80, n_CDS=33, title=paste0(species, ".", connector, "TR_bottom_PWM"))
	  
	  	# PWM scores (aligned at AUG)
	  	species.TR_top_PWM_max_UTR_len_score <- readRDS(paste0("processed_data_w_polyA/AUGPWM_R2s/", species, ".", connector, "TR_top_PWM_max_UTR_len_score.rds"))
	  
	  	species.TR_top_PWM_max_UTR_len_CDS_13_score <- readRDS(paste0("processed_data_w_polyA/AUGPWM_R2s/", species, ".", connector, "TR_top_PWM_max_UTR_len_CDS_13_score.rds"))
	  
	  	species.TR_top_PWM_max_UTR_len_CDS_23_score <- readRDS(paste0("processed_data_w_polyA/AUGPWM_R2s/", species, ".", connector, "TR_top_PWM_max_UTR_len_CDS_23_score.rds"))
	  
	  	species.TR_top_PWM_max_UTR_len_CDS_28_score <- readRDS(paste0("processed_data_w_polyA/AUGPWM_R2s/", species, ".", connector, "TR_top_PWM_max_UTR_len_CDS_28_score.rds"))
	  
	  	species.TR_top_PWM_max_UTR_len_CDS_33_score <- readRDS(paste0("processed_data_w_polyA/AUGPWM_R2s/", species, ".", connector, "TR_top_PWM_max_UTR_len_CDS_33_score.rds"))
	  
	  	# calculate the R2s for genes with different minimum 5' UTR lengths
	  	## top TR
		species.TR_top_R2s_w_TR <- sapply(1:length(len_breaks), FUN=function(idx) {
	    	long_idx <- which(species.log10.UTR_len >= log(100-len_breaks[idx]+5,10))
	    	c( cor(species.log10.TR[long_idx], species.TR_top_PWM_max_UTR_len_score[,idx][long_idx])^2,
	    	cor(species.log10.TR[long_idx], species.TR_top_PWM_max_UTR_len_CDS_13_score[,idx][long_idx])^2,
	    	cor(species.log10.TR[long_idx], species.TR_top_PWM_max_UTR_len_CDS_23_score[,idx][long_idx])^2,
	    	cor(species.log10.TR[long_idx], species.TR_top_PWM_max_UTR_len_CDS_28_score[,idx][long_idx])^2,
	    	cor(species.log10.TR[long_idx], species.TR_top_PWM_max_UTR_len_CDS_33_score[,idx][long_idx])^2
	    	)
		})
		# plot the R2's
		
		## top TR
		plot(x=seq(from=-100, to=-5, by=5), y=species.TR_top_R2s_w_TR[1,], ylim=range(species.TR_top_R2s_w_TR), main=paste0(species, ".", connector), xlab="length 5' UTR TR top PWM", ylab="R2 logTR vs. TR top PWM score", pch=20, axes=T, col=gray(.9))
		points(x=seq(from=-100, to=-5, by=5), y=species.TR_top_R2s_w_TR[2,], pch=20, col=gray(0.7))
		points(x=seq(from=-100, to=-5, by=5), y=species.TR_top_R2s_w_TR[3,], pch=20, col=gray(0.5))
		points(x=seq(from=-100, to=-5, by=5), y=species.TR_top_R2s_w_TR[4,], pch=20, col=gray(0.3))
		points(x=seq(from=-100, to=-5, by=5), y=species.TR_top_R2s_w_TR[5,], pch=20, col=gray(0.1))
		max_idx <- which(species.TR_top_R2s_w_TR==max(species.TR_top_R2s_w_TR), arr.ind=T)
		abline(v=seq(from=-100, to=-5, by=5)[which.max(species.TR_top_R2s_w_TR[max_idx[1],])], lty=2)
		legend("bottomleft", c("5' UTR only", "5' UTR & +4 to +13", "5' UTR & +4 to +23", "5' UTR & +4 to +28", "5' UTR & +4 to +33"), pch=c(20,20,20,20,20), col=c(gray(0.9),gray(0.7),gray(0.5),gray(0.3),gray(0.1)), bty="n")
	}
}
```

# Figure 7 - figure supplement 2. R2s between uTICE (PWM only or PWM + selected di/tri nucleotides), dTICE (PWM only or PWM + selected di/tri nucleotides), or TICE (PWM only or PWM + selected di/tri nucleotides)
## the following code only needs to be run once
```{r, eval = FALSE}
R2s_uTICE_dTICE_TICE <- NULL
rownames_R2s_uTICE_dTICE_TICE <- NULL
colnames_R2s_uTICE_dTICE_TICE <- c("uTICE_PWM", "uTICE_PWM+di/tri", "dTICE_PWM", "dTICE_PWM+di/tri", "TICE_PWM", "TICE_PWM+di/tri")

for (species in names(species_tissue_list)) {
	for (species_tissue in species_tissue_list[[species]]) {
		if (species_tissue!="") {
			tissue_dot <- paste0(".", species_tissue)
			tissue_us <- paste0("_", species_tissue)
		} else {
			tissue_dot <- ""
			tissue_us <- ""
		}
	  
	  uTICE_features <- get(paste0(species, tissue_dot, ".uTICE_features"))[[1]]
	  dTICE_features <- get(paste0(species, tissue_dot, ".dTICE_features"))[[1]]
	  
	  uTICE_PWM_features <- uTICE_features[, c(which(grepl("^PWM", colnames(uTICE_features))), which(grepl("^Intercept", colnames(uTICE_features))))]
	  dTICE_PWM_features <- dTICE_features[, c(which(grepl("^PWM", colnames(dTICE_features))), which(grepl("^Intercept", colnames(dTICE_features))))]
	  
	  TICE_features <- cbind(uTICE_features, dTICE_features)
	  TICE_PWM_features <- cbind(uTICE_PWM_features, dTICE_PWM_features)
	    
	  R2s_uTICE_dTICE_TICE <- rbind(R2s_uTICE_dTICE_TICE,
	                                               c(
	                                                 summary(lm(get(paste0(species, tissue_dot, ".log10.TR")) ~ as.matrix(uTICE_PWM_features)))$r.squared,
	                                                 summary(lm(get(paste0(species, tissue_dot, ".log10.TR")) ~ as.matrix(uTICE_features)))$r.squared,
	                                                 summary(lm(get(paste0(species, tissue_dot, ".log10.TR")) ~ as.matrix(dTICE_PWM_features)))$r.squared,
	                                                 summary(lm(get(paste0(species, tissue_dot, ".log10.TR")) ~ as.matrix(dTICE_features)))$r.squared,
	                                                 summary(lm(get(paste0(species, tissue_dot, ".log10.TR")) ~ as.matrix(TICE_PWM_features)))$r.squared,
	                                                 summary(lm(get(paste0(species, tissue_dot, ".log10.TR")) ~ as.matrix(TICE_features)))$r.squared
	                                                 )
	                                               )
	  rownames_R2s_uTICE_dTICE_TICE <- c(
	    rownames_R2s_uTICE_dTICE_TICE,
	    paste0(species, tissue_dot)
	  )
	}
}
rownames(R2s_uTICE_dTICE_TICE) <- rownames_R2s_uTICE_dTICE_TICE
colnames(R2s_uTICE_dTICE_TICE) <- colnames_R2s_uTICE_dTICE_TICE

write.csv(R2s_uTICE_dTICE_TICE, file="tables/R2s_uTICE_dTICE_TICE.csv", quote=F)
```
## plot
```{r FigureS7.2, fig.width=5, fig.height=5, fig.path='figures/', dev=c('pdf'), out.width="50%"}
R2s_uTICE_dTICE_TICE <- read.csv(file="tables/R2s_uTICE_dTICE_TICE.csv")
colnames(R2s_uTICE_dTICE_TICE)[1] <- "species.tissue"

# only keep the exemplar tissues
for (species in names(species_tissue_list)) {
	for (i in 1:length(species_exemplar_tissue_list[[species]])) {
		species_tissue <- species_exemplar_tissue_list[[species]][i]
		if (species_tissue!="") {
			tissue_dot <- paste0(".", species_tissue)
			tissue_us <- paste0("_", species_tissue)
		} else {
			tissue_dot <- ""
			tissue_us <- ""
		}
		tmp_for_plot <- as.data.frame( matrix(R2s_uTICE_dTICE_TICE[R2s_uTICE_dTICE_TICE$species.tissue==paste0(species, tissue_dot),2:ncol(R2s_uTICE_dTICE_TICE)], ncol=1) )
		colnames(tmp_for_plot) <- "R2"
		tmp_for_plot$Feature <- factor( rep(c("uTICE", "dTICE", "TICE"), each=2), levels=c("uTICE", "dTICE", "TICE") )
		tmp_for_plot$ifPWMonly <- factor( rep(c("PWM", "PWM+di/tri"), 3), levels=c("PWM", "PWM+di/tri") )
		tmp_for_plot$R2 <- as.numeric(tmp_for_plot$R2)
		
		theme_set(theme_bw())
print( ggplot(tmp_for_plot, aes(x=Feature, y=R2, fill=ifPWMonly)) + geom_bar(stat="identity", position=position_dodge()) + scale_fill_manual(values=c("#AECDE1", "#447DB2")) + theme(axis.text.x = element_text(angle = 45, vjust = 1, hjust=1)) + ggtitle(paste0(species, tissue_dot)) )
	}
}
```

# Figure 8. R2 of 5'ofTICE vs. R2 of TICE vs. R2 of 5'motifs (5'ofTICE + TICE)
## the following code only needs to be run once
```{r, eval = FALSE}
R2s_5ofTICE_TICE_5motifs <- NULL
rownames_R2s_5ofTICE_TICE_5motifs <- NULL
colnames_R2s_5ofTICE_TICE_5motifs <- c("5'ofTICE", "TICE", "5'motifs")

for (species in names(species_tissue_list)) {
	for (species_tissue in species_tissue_list[[species]]) {
		if (species_tissue!="") {
			tissue_dot <- paste0(".", species_tissue)
			tissue_us <- paste0("_", species_tissue)
		} else {
			tissue_dot <- ""
			tissue_us <- ""
		}
	  
	  R2s_5ofTICE_TICE_5motifs <- rbind(R2s_5ofTICE_TICE_5motifs,
	                                               c(
	                                                 summary(lm(get(paste0(species, tissue_dot, ".log10.TR")) ~ as.matrix(get(paste0(species, tissue_dot, ".5ofTICE_features"))[[1]])))$r.squared,
	                                                 summary(lm(get(paste0(species, tissue_dot, ".log10.TR")) ~ as.matrix(cbind(get(paste0(species, tissue_dot, ".uTICE_features"))[[1]], get(paste0(species, tissue_dot, ".dTICE_features"))[[1]]))))$r.squared,
	                                                 summary(lm(get(paste0(species, tissue_dot, ".log10.TR")) ~ as.matrix(cbind(get(paste0(species, tissue_dot, ".uTICE_features"))[[1]], get(paste0(species, tissue_dot, ".dTICE_features"))[[1]], get(paste0(species, tissue_dot, ".5ofTICE_features"))[[1]]))))$r.squared
	                                                 )
	                                               )
	  rownames_R2s_5ofTICE_TICE_5motifs <- c(
	    rownames_R2s_5ofTICE_TICE_5motifs,
	    paste0(species, tissue_dot)
	  )
	}
}
rownames(R2s_5ofTICE_TICE_5motifs) <- rownames_R2s_5ofTICE_TICE_5motifs
colnames(R2s_5ofTICE_TICE_5motifs) <- colnames_R2s_5ofTICE_TICE_5motifs

write.csv(R2s_5ofTICE_TICE_5motifs, file="tables/R2s_5ofTICE_TICE_5motifs.csv", quote=F)
```
## plot
```{r Figure8, fig.width=5, fig.height=5, fig.path='figures/', dev=c('pdf'), out.width="50%"}
R2s_5ofTICE_TICE_5motifs <- read.csv(file="tables/R2s_5ofTICE_TICE_5motifs.csv")
colnames(R2s_5ofTICE_TICE_5motifs)[1] <- "species.tissue"

# only keep the exemplar tissues
R2s_5ofTICE_TICE_5motifs_for_plot <- setNames(melt(R2s_5ofTICE_TICE_5motifs[R2s_5ofTICE_TICE_5motifs$species.tissue %in% species_exemplar_tissue_names,]), c("Species_Tissue", "Feature", 'R2'))

for (species in names(species_tissue_list)) {
	for (i in 1:length(species_exemplar_tissue_list[[species]])) {
		species_tissue <- species_exemplar_tissue_list[[species]][i]
		if (species_tissue!="") {
			tissue_dot <- paste0(".", species_tissue)
			tissue_us <- paste0("_", species_tissue)
		} else {
			tissue_dot <- ""
			tissue_us <- ""
		}
		tmp_for_plot <- R2s_5ofTICE_TICE_5motifs_for_plot[R2s_5ofTICE_TICE_5motifs_for_plot$Species_Tissue==paste0(species, tissue_dot),]
		
		tmp_for_plot$Feature <- factor(tmp_for_plot$Feature, levels=c("X5.ofTICE", "TICE", "X5.motifs"))
		
		theme_set(theme_bw())
print( ggplot(tmp_for_plot, aes(x=Feature, y=R2, fill=Species_Tissue)) + geom_bar(stat="identity", position=position_dodge()) + scale_fill_manual(values="#447DB2") + theme(axis.text.x = element_text(angle = 45, vjust = 1, hjust=1)) + ggtitle(paste0(species, tissue_dot)) )
	}
}
```

# Figure 8 - figure supplement 1. 5' cap PWMs and R2s of various PWM boundaries
## the following code only needs to be run once
```{r, eval = FALSE}
for (species in names(species_exemplar_tissue_list)) {
	for (species_tissue in species_exemplar_tissue_list[[species]]) {
		if (species_tissue!="") {
			species.genes <- get(paste(species, species_tissue, "genes", sep="."))
			species.UTR_seqs <- get(paste(species, species_tissue, "UTR_seqs", sep="."))
			species.log10.TR <- get(paste(species, species_tissue, "log10.TR", sep="."))
			species.TR_top_genes <- get(paste(species, species_tissue, "TR_top_genes", sep="."))
			species.TR_bottom_genes <- get(paste(species, species_tissue, "TR_bottom_genes", sep="."))
			connector <- paste0(species_tissue, ".")
		} else {
			species.genes <- get(paste(species, "genes", sep="."))
			species.UTR_seqs <- get(paste(species, "UTR_seqs", sep="."))
			species.log10.TR <- get(paste(species, "log10.TR", sep="."))
			species.TR_top_genes <- get(paste(species, "TR_top_genes", sep="."))
			species.TR_bottom_genes <- get(paste(species, "TR_bottom_genes", sep="."))
			connector <- ""
		}
		species.log10.UTR_len <- log(nchar(species.UTR_seqs), 10)
		if (species != "hs") {
			species.uTICE_len <- species_uTICE_len_list[[species]]
		} else {
			species.uTICE_len <- species_uTICE_len_list[[species]][[species_tissue]]
		}
		
		## for the genes in the top and bottom cohort, find the 5' UTR sequences and CDS sequences
		species.TR_top_UTR_seqs <- species.UTR_seqs[species.TR_top_genes]		
		species.TR_bottom_UTR_seqs <- species.UTR_seqs[species.TR_bottom_genes]
		
		# construct Position Weight Matrices (PWMs) based on the top or bottom 10% TR genes from the 5' end with different lengths, excluding the TICE sequence
		len_breaks <- c(1:10, seq(from=15, to=100, by=5)) # varying UTR from the 5' end; do not include CDS sequences or the uTICE region
		
		## top TR
		species.TR_top_PWM_5_aligned  <- UTR_pwm_5_aligned(UTR_seqs=substr( x=species.TR_top_UTR_seqs, start=1, stop=nchar(species.TR_top_UTR_seqs)-species.uTICE_len ), n=max(len_breaks))
		colnames(species.TR_top_PWM_5_aligned) <- c("A", "T", "C", "G")
		saveRDS(species.TR_top_PWM_5_aligned, file=paste0("processed_data_w_polyA/5'PWM_R2s/", species, ".", connector, "TR_top_PWM_5_aligned.rds"))

		## bottom TR
		species.TR_bottom_PWM_5_aligned  <- UTR_pwm_5_aligned(UTR_seqs=substr( x=species.TR_bottom_UTR_seqs, start=1, stop=nchar(species.TR_bottom_UTR_seqs)-species.uTICE_len ), n=max(len_breaks))
		colnames(species.TR_bottom_PWM_5_aligned) <- c("A", "T", "C", "G")
		saveRDS(species.TR_bottom_PWM_5_aligned, file=paste0("processed_data_w_polyA/5'PWM_R2s/", species, ".", connector, "TR_bottom_PWM_5_aligned.rds"))
	
		# PWM scores (aligned at the 5' end)

		## top TR
		species.TR_top_PWM_5_aligned_score <- sapply(len_breaks, FUN=function(i){
	    	if (i==1) {
	      	  PWM_tmp <- matrix(species.TR_top_PWM_5_aligned[1:i,], nrow=1)
	      	  colnames(PWM_tmp) <- c("A", "T", "C", "G")
	    	} else {
	      	  PWM_tmp <- species.TR_top_PWM_5_aligned[1:i,]
	    	}
	    	UTR_pwm_5_aligned_score( PWM_tmp, species.UTR_seqs )
		})
		saveRDS(species.TR_top_PWM_5_aligned_score, file=paste0("processed_data_w_polyA/5'PWM_R2s/", species, ".", connector, "TR_top_PWM_5_aligned_score.rds"))
	}
}
```
## plot
```{r FigureS8, fig.width=5, fig.height=5, fig.path='figures/', dev=c('pdf'), out.width="50%"}
# a function to plot a 5' cap PWM
plot_pwm_5_aligned <- function(PWM, title) {
	colnames(PWM) <- c("A", "U", "C", "G")
	PWM_motif <- new("pcm", mat=t(PWM), name=title)
	plot(PWM_motif, ic.scale=F)
}

len_breaks <- c(1:10, seq(from=15, to=100, by=5))

for (species in names(species_exemplar_tissue_list)) {
	for (species_tissue in species_exemplar_tissue_list[[species]]) {
	  if (species_tissue!="") {
			species.log10.TR <- get(paste(species, species_tissue, "log10.TR", sep="."))
			connector <- paste0(species_tissue, ".")
		} else {
			species.log10.TR <- get(paste(species, "log10.TR", sep="."))
			connector <- ""
		}
	  
	  species.TR_top_PWM_5_aligned <- readRDS(file=paste0("processed_data_w_polyA/5'PWM_R2s/", species, ".", connector, "TR_top_PWM_5_aligned.rds"))
	  species.TR_bottom_PWM_5_aligned <- readRDS(file=paste0("processed_data_w_polyA/5'PWM_R2s/", species, ".", connector, "TR_bottom_PWM_5_aligned.rds"))
	  ## plot the PWM matrices
	  plot_pwm_5_aligned(species.TR_top_PWM_5_aligned[1:50,], title=paste0(species, ".", connector, "TR_top_PWM_5_aligned"))
	  plot_pwm_5_aligned(species.TR_bottom_PWM_5_aligned[1:50,], title=paste0(species, ".", connector, "TR_bottom_PWM_5_aligned"))
	  
	  # PWM scores (aligned at the 5' end)
	  species.TR_top_PWM_5_aligned_score <- readRDS(file=paste0("processed_data_w_polyA/5'PWM_R2s/", species, ".", connector, "TR_top_PWM_5_aligned_score.rds"))
	  
	  # calculate the R2s for genes with different minimum 5' UTR lengths
		species.TR_top_R2s_w_TR <- sapply(1:length(len_breaks), FUN=function(idx) {
		    R2 <- cor(species.log10.TR, species.TR_top_PWM_5_aligned_score[,idx])^2
		    return(R2)
		})
		
		# plot the R2's
		plot(x=len_breaks, y=species.TR_top_R2s_w_TR, main=paste0(species, ".", connector), xlab="length of 5' UTR TR top PWM (5' aligned)", ylab="R2 logTR vs. TR top PWM score", pch=20, axes=T, col=1)
	}
}
```

# Figure 9 - figure supplement 1. Ratios of trinucleotide frequencies between the top and bottom TR genes in the 5' of TICE region, the uTICE region, the dTICE region, and the 3' of TICE region
## the following code only needs to be run once
```{r, eval = FALSE}
col_max <- NULL # record the maximum value to plot in next part

# Remove genes with short 5' UTRs. Keep genes with at least 3 bp in each region of interest
for (species in names(species_exemplar_tissue_list)) {
	for (species_tissue in species_exemplar_tissue_list[[species]]) {
		if (species_tissue!="") {
			species.genes <- get(paste(species, species_tissue, "genes", sep="."))
			species.UTR_seqs <- get(paste(species, species_tissue, "UTR_seqs", sep="."))
			species.CDS_seqs <- get(paste(species, species_tissue, "CDS_seqs", sep="."))
			species.log10.TR <- get(paste(species, species_tissue, "log10.TR", sep="."))
			species.TR_top_genes <- get(paste(species, species_tissue, "TR_top_genes", sep="."))
			species.TR_bottom_genes <- get(paste(species, species_tissue, "TR_bottom_genes", sep="."))
			connector <- paste0(".", species_tissue)
		} else {
			species.genes <- get(paste(species, "genes", sep="."))
			species.UTR_seqs <- get(paste(species, "UTR_seqs", sep="."))
			species.CDS_seqs <- get(paste(species, "CDS_seqs", sep="."))
			species.log10.TR <- get(paste(species, "log10.TR", sep="."))
			species.TR_top_genes <- get(paste(species, "TR_top_genes", sep="."))
			species.TR_bottom_genes <- get(paste(species, "TR_bottom_genes", sep="."))
			connector <- ""
		}
	  
	  if (species != "hs") {
			species.uTICE_len <- species_uTICE_len_list[[species]]
			species.dTICE_len <- species_dTICE_len_list[[species]]
		} else {
			species.uTICE_len <- species_uTICE_len_list[[species]][[species_tissue]]
			species.dTICE_len <- species_dTICE_len_list[[species]][[species_tissue]]
		}
	  
	  species.UTR_tri_freqs_5ofTICE <- readRDS(file=paste0("processed_data_w_polyA/tri_freqs/", species, connector, ".UTR_tri_freqs_5ofTICE.rds"))
	  species.UTR_tri_freqs_uTICE <- readRDS(file=paste0("processed_data_w_polyA/tri_freqs/", species, connector, ".UTR_tri_freqs_uTICE.rds"))
	  species.CDS_tri_freqs_dTICE <- readRDS(file=paste0("processed_data_w_polyA/tri_freqs/", species, connector, ".CDS_tri_freqs_dTICE.rds"))
	  species.CDS_tri_freqs_3ofTICE <- readRDS(file=paste0("processed_data_w_polyA/tri_freqs/", species, connector, ".CDS_tri_freqs_3ofTICE.rds"))
	  
	  species.TR_top_UTR_seqs <- species.UTR_seqs[species.TR_top_genes]
	  species.TR_bottom_UTR_seqs <- species.UTR_seqs[species.TR_bottom_genes]
	  species.TR_top_CDS_seqs <- species.CDS_seqs[species.TR_top_genes]
	  species.TR_bottom_CDS_seqs <- species.CDS_seqs[species.TR_bottom_genes]
	  
	  # 5' of TICE
	  species.UTR_min_len <- species.uTICE_len + 3
	  
	  species.UTR_tri_freq_top_bottom_ratios_5ofTICE <- colMeans(species.UTR_tri_freqs_5ofTICE[species.TR_top_genes[nchar(species.TR_top_UTR_seqs)>=species.UTR_min_len],]) / colMeans(species.UTR_tri_freqs_5ofTICE[species.TR_bottom_genes[nchar(species.TR_bottom_UTR_seqs)>=species.UTR_min_len],])
	  species.UTR_tri_freq_top_bottom_ratios_5ofTICE_table <- rbind(
	    matrix(species.UTR_tri_freq_top_bottom_ratios_5ofTICE[1:16], nrow=4),
	    matrix(species.UTR_tri_freq_top_bottom_ratios_5ofTICE[17:32], nrow=4),
	    matrix(species.UTR_tri_freq_top_bottom_ratios_5ofTICE[33:48], nrow=4),
	    matrix(species.UTR_tri_freq_top_bottom_ratios_5ofTICE[49:64], nrow=4)
	  )
	  
	  write.csv(species.UTR_tri_freq_top_bottom_ratios_5ofTICE_table, file=paste0("tables/tri_freq_tables/", species, connector, ".UTR_tri_freq_top_bottom_ratios_5ofTICE_table.csv"), quote=F)
	            
	  # uTICE
	  species.UTR_tri_freq_top_bottom_ratios_uTICE <- colMeans(species.UTR_tri_freqs_uTICE[species.TR_top_genes[nchar(species.TR_top_UTR_seqs)>=3],]) / colMeans(species.UTR_tri_freqs_uTICE[species.TR_bottom_genes[nchar(species.TR_bottom_UTR_seqs)>=3],])
	  species.UTR_tri_freq_top_bottom_ratios_uTICE_table <- rbind(
	    matrix(species.UTR_tri_freq_top_bottom_ratios_uTICE[1:16], nrow=4),
	    matrix(species.UTR_tri_freq_top_bottom_ratios_uTICE[17:32], nrow=4),
	    matrix(species.UTR_tri_freq_top_bottom_ratios_uTICE[33:48], nrow=4),
	    matrix(species.UTR_tri_freq_top_bottom_ratios_uTICE[49:64], nrow=4)
	  )
	  
	  write.csv(species.UTR_tri_freq_top_bottom_ratios_uTICE_table, file=paste0("tables/tri_freq_tables/", species, connector, ".UTR_tri_freq_top_bottom_ratios_uTICE_table.csv"), quote=F)
	            
	  # dTICE
	  species.CDS_tri_freq_top_bottom_ratios_dTICE <- colMeans(species.CDS_tri_freqs_dTICE[species.TR_top_genes[nchar(species.TR_top_CDS_seqs)>=3],]) / colMeans(species.CDS_tri_freqs_dTICE[species.TR_bottom_genes[nchar(species.TR_bottom_CDS_seqs)>=3],])
	  species.CDS_tri_freq_top_bottom_ratios_dTICE_table <- rbind(
	    matrix(species.CDS_tri_freq_top_bottom_ratios_dTICE[1:16], nrow=4),
	    matrix(species.CDS_tri_freq_top_bottom_ratios_dTICE[17:32], nrow=4),
	    matrix(species.CDS_tri_freq_top_bottom_ratios_dTICE[33:48], nrow=4),
	    matrix(species.CDS_tri_freq_top_bottom_ratios_dTICE[49:64], nrow=4)
	  )
	  write.csv(species.CDS_tri_freq_top_bottom_ratios_dTICE_table, file=paste0("tables/tri_freq_tables/", species, connector, ".CDS_tri_freq_top_bottom_ratios_dTICE_table.csv"), quote=F)
	            
	  # 3' of TICE
	  species.CDS_min_len <- species.dTICE_len + 3
	  
	  species.CDS_tri_freq_top_bottom_ratios_3ofTICE <- colMeans(species.CDS_tri_freqs_3ofTICE[species.TR_top_genes[nchar(species.TR_top_CDS_seqs)>=species.CDS_min_len],]) / colMeans(species.CDS_tri_freqs_3ofTICE[species.TR_bottom_genes[nchar(species.TR_bottom_CDS_seqs)>=species.CDS_min_len],])
	  species.CDS_tri_freq_top_bottom_ratios_3ofTICE_table <- rbind(
	    matrix(species.CDS_tri_freq_top_bottom_ratios_3ofTICE[1:16], nrow=4),
	    matrix(species.CDS_tri_freq_top_bottom_ratios_3ofTICE[17:32], nrow=4),
	    matrix(species.CDS_tri_freq_top_bottom_ratios_3ofTICE[33:48], nrow=4),
	    matrix(species.CDS_tri_freq_top_bottom_ratios_3ofTICE[49:64], nrow=4)
	  )
	  write.csv(species.CDS_tri_freq_top_bottom_ratios_3ofTICE_table, file=paste0("tables/tri_freq_tables/", species, connector, ".CDS_tri_freq_top_bottom_ratios_3ofTICE_table.csv"), quote=F)
	  
	  col_max <- max(col_max, max(species.UTR_tri_freq_top_bottom_ratios_5ofTICE_table), max(species.UTR_tri_freq_top_bottom_ratios_uTICE_table), max(species.CDS_tri_freq_top_bottom_ratios_dTICE_table), max(species.CDS_tri_freq_top_bottom_ratios_3ofTICE_table))
	}
}

saveRDS(col_max, file="processed_data_w_polyA/tri_freq_tables/col_max.rds")
```
## plot
```{r FigureS9, fig.width=5, fig.height=5, fig.path='figures/', dev=c('pdf'), out.width="50%"}
col_max <- readRDS(file="processed_data_w_polyA/tri_freq_tables/col_max.rds")

col_breaks <- c(seq(0, 0.8, length=100),  # for blue
  seq(0.81, 1.25, length=100),           # for white
  seq(1.26, col_max, length=100))       # for red

for (species in names(species_exemplar_tissue_list)) {
	for (species_tissue in species_exemplar_tissue_list[[species]]) {
		if (species_tissue!="") {
			connector <- paste0(".", species_tissue)
		} else {
			connector <- ""
		}
	  
	  # 5' of TICE
	  species.UTR_tri_freq_top_bottom_ratios_5ofTICE_table <- as.matrix(read.csv(file=paste0("tables/tri_freq_tables/", species, connector, ".UTR_tri_freq_top_bottom_ratios_5ofTICE_table.csv"), header=T)[,-1])
	  
	  heatmap.2(species.UTR_tri_freq_top_bottom_ratios_5ofTICE_table,
	            cellnote = tri_nts_table,  # same data set for cell labels
	            main = paste0(species, connector, " top/bottom ratios\nin the 5ofTICE region"), # heat map title
	            notecol="black",      # change font color of cell labels to black
	            density.info="none",  # turns off density plot inside color legend
	            trace="none",         # turns off trace lines inside the heat map
	            #margins =c(12,9),     # widens margins around plot
	            col=my_palette,       # use on color palette defined earlier
	            breaks=col_breaks,    # enable color transition at specified limits
	            dendrogram="none",    # only draw a row dendrogram
	            Rowv="NA",            # turn off row clustering
	            Colv="NA",            # turn off column clustering
	            labRow="",            # turn off row labeling
	            labCol=""             # turn off column labeling
	  )
	            
	  # uTICE
	  species.UTR_tri_freq_top_bottom_ratios_uTICE_table <- as.matrix(read.csv(file=paste0("tables/tri_freq_tables/", species, connector, ".UTR_tri_freq_top_bottom_ratios_uTICE_table.csv"), header=T)[,-1])

	  heatmap.2(species.UTR_tri_freq_top_bottom_ratios_uTICE_table,
	            cellnote = tri_nts_table,  # same data set for cell labels
	            main = paste0(species, connector, " top/bottom ratios\nin the uTICE region"), # heat map title
	            notecol="black",      # change font color of cell labels to black
	            density.info="none",  # turns off density plot inside color legend
	            trace="none",         # turns off trace lines inside the heat map
	            #margins =c(12,9),     # widens margins around plot
	            col=my_palette,       # use on color palette defined earlier
	            breaks=col_breaks,    # enable color transition at specified limits
	            dendrogram="none",    # only draw a row dendrogram
	            Rowv="NA",            # turn off row clustering
	            Colv="NA",            # turn off column clustering
	            labRow="",            # turn off row labeling
	            labCol=""             # turn off column labeling
	  )
	  
	  # dTICE
	  species.CDS_tri_freq_top_bottom_ratios_dTICE_table <- as.matrix(read.csv(file=paste0("tables/tri_freq_tables/", species, connector, ".CDS_tri_freq_top_bottom_ratios_dTICE_table.csv"), header=T)[,-1])

	  heatmap.2(species.CDS_tri_freq_top_bottom_ratios_dTICE_table,
	            cellnote = tri_nts_table,  # same data set for cell labels
	            main = paste0(species, connector, " top/bottom ratios\nin the dTICE region"), # heat map title
	            notecol="black",      # change font color of cell labels to black
	            density.info="none",  # turns off density plot inside color legend
	            trace="none",         # turns off trace lines inside the heat map
	            #margins =c(12,9),     # widens margins around plot
	            col=my_palette,       # use on color palette defined earlier
	            breaks=col_breaks,    # enable color transition at specified limits
	            dendrogram="none",    # only draw a row dendrogram
	            Rowv="NA",            # turn off row clustering
	            Colv="NA",            # turn off column clustering
	            labRow="",            # turn off row labeling
	            labCol=""             # turn off column labeling
	  )
	            
	  # 3' of TICE
	  species.CDS_tri_freq_top_bottom_ratios_3ofTICE_table <- as.matrix(read.csv(file=paste0("tables/tri_freq_tables/", species, connector, ".CDS_tri_freq_top_bottom_ratios_3ofTICE_table.csv"), header=T)[,-1])
	  
	  heatmap.2(species.CDS_tri_freq_top_bottom_ratios_3ofTICE_table,
	            cellnote = tri_nts_table,  # same data set for cell labels
	            main = paste0(species, connector, " top/bottom ratios\nin the uTICE region"), # heat map title
	            notecol="black",      # change font color of cell labels to black
	            density.info="none",  # turns off density plot inside color legend
	            trace="none",         # turns off trace lines inside the heat map
	            #margins =c(12,9),     # widens margins around plot
	            col=my_palette,       # use on color palette defined earlier
	            breaks=col_breaks,    # enable color transition at specified limits
	            dendrogram="none",    # only draw a row dendrogram
	            Rowv="NA",            # turn off row clustering
	            Colv="NA",            # turn off column clustering
	            labRow="",            # turn off row labeling
	            labCol=""             # turn off column labeling
	  )
	}
}

```

# Figure 10. Pearson correlations between the top/bottom trinucleotide ratios of two regions within a species
```{r Figure10, fig.width=7, fig.height=7, fig.path='figures/', dev=c('pdf'), out.width="50%"}
for (species in names(species_exemplar_tissue_list)) {
	for (species_tissue in species_exemplar_tissue_list[[species]]) {
		if (species_tissue!="") {
			connector <- paste0(".", species_tissue)
		} else {
			connector <- ""
		}
	  
	  # 5' of TICE
	  species.UTR_tri_freq_top_bottom_ratios_5ofTICE_table <- as.numeric(unlist(read.csv(file=paste0("tables/tri_freq_tables/", species, connector, ".UTR_tri_freq_top_bottom_ratios_5ofTICE_table.csv"), header=T)[,-1]))
	  
	  # uTICE
	  species.UTR_tri_freq_top_bottom_ratios_uTICE_table <- as.numeric(unlist(read.csv(file=paste0("tables/tri_freq_tables/", species, connector, ".UTR_tri_freq_top_bottom_ratios_uTICE_table.csv"), header=T)[,-1]))

	  # dTICE
	  species.CDS_tri_freq_top_bottom_ratios_dTICE_table <- as.numeric(unlist(read.csv(file=paste0("tables/tri_freq_tables/", species, connector, ".CDS_tri_freq_top_bottom_ratios_dTICE_table.csv"), header=T)[,-1]))

	  # 3' of TICE
	  species.CDS_tri_freq_top_bottom_ratios_3ofTICE_table <- as.numeric(unlist(read.csv(file=paste0("tables/tri_freq_tables/", species, connector, ".CDS_tri_freq_top_bottom_ratios_3ofTICE_table.csv"), header=T)[,-1]))
	  
	  # the correlation table
	  species.cors <- matrix(NA, nrow=4, ncol=4)
	  rownames(species.cors) <- c("5ofTICE", "uTICE", "dTICE", "3ofTICE")
	  colnames(species.cors) <- c("5ofTICE", "uTICE", "dTICE", "3ofTICE")
	  species.cors["uTICE", "5ofTICE"] <- cor(species.UTR_tri_freq_top_bottom_ratios_uTICE_table, species.UTR_tri_freq_top_bottom_ratios_5ofTICE_table)
	  species.cors["dTICE", "5ofTICE"] <- cor(species.CDS_tri_freq_top_bottom_ratios_dTICE_table, species.UTR_tri_freq_top_bottom_ratios_5ofTICE_table)
	  species.cors["dTICE", "uTICE"] <- cor(species.CDS_tri_freq_top_bottom_ratios_dTICE_table, species.UTR_tri_freq_top_bottom_ratios_uTICE_table)
	  species.cors["3ofTICE", "5ofTICE"] <- cor(species.CDS_tri_freq_top_bottom_ratios_3ofTICE_table, species.UTR_tri_freq_top_bottom_ratios_5ofTICE_table)
	  species.cors["3ofTICE", "uTICE"] <- cor(species.CDS_tri_freq_top_bottom_ratios_3ofTICE_table, species.UTR_tri_freq_top_bottom_ratios_uTICE_table)
	  species.cors["3ofTICE", "dTICE"] <- cor(species.CDS_tri_freq_top_bottom_ratios_3ofTICE_table, species.CDS_tri_freq_top_bottom_ratios_dTICE_table)
	  
	  col_breaks <- c(seq(-1,-0.1,length=100),  # for blue
	                  seq(-0.09,0.09,length=100),  # for white
	                  seq(0.1,1,length=100)  # for red
	                  )
	  
	  # make the within-species correlation heatmap
	  heatmap.2(species.cors,
	            main = paste0(species, connector, " cor of\ntrinucleotide ratios"), # heat map title
	            cellnote = round(species.cors, 2),  # same data set for cell labels
	            notecol="black",      # change font color of cell labels to black
	            density.info="none",  # turns off density plot inside color legend
	            trace="none",         # turns off trace lines inside the heat map
	            margins =c(15,15),     # widens margins around plot
	            col=my_palette,       # use on color palette defined earlier
	            breaks=col_breaks,    # enable color transition at specified limits
	            dendrogram="none",    # only draw a row dendrogram
	            Rowv="NA",            # turn off row clustering
	            Colv="NA"            # turn off column clustering
	            )
	}
}
```

# Figure 11. Pearson correlations between the top/bottom trinucleotide ratios of the region of two species
```{r Figure11, fig.width=7, fig.height=7, fig.path='figures/', dev=c('pdf'), out.width="50%"}
btw_species_cors <- NULL
rownames_btw_species_cors <- NULL
colnames_btw_species_cors <- c("5ofTICE vs 5ofTICE", "uTICE vs uTICE", "dTICE vs dTICE", "3ofTICE vs 3ofTICE")
for (i in 1:(length(species_exemplar_tissue_list)-1)) {
  for (j in (i+1):length(species_exemplar_tissue_list)) {
    species1 <- names(species_exemplar_tissue_list)[i]
    species2 <- names(species_exemplar_tissue_list)[j]
    species1_tissue <- species_exemplar_tissue_list[[species1]]
    species2_tissue <- species_exemplar_tissue_list[[species2]]
    if (species1_tissue!="") {
			connector1 <- paste0(".", species1_tissue)
		} else {
			connector1 <- ""
		}
    if (species2_tissue!="") {
			connector2 <- paste0(".", species2_tissue)
		} else {
			connector2 <- ""
		}
    
    # 5' of TICE
	  species1.UTR_tri_freq_top_bottom_ratios_5ofTICE_table <- as.numeric(unlist(read.csv(file=paste0("tables/tri_freq_tables/", species1, connector1, ".UTR_tri_freq_top_bottom_ratios_5ofTICE_table.csv"), header=T)[,-1]))
	  species2.UTR_tri_freq_top_bottom_ratios_5ofTICE_table <- as.numeric(unlist(read.csv(file=paste0("tables/tri_freq_tables/", species2, connector2, ".UTR_tri_freq_top_bottom_ratios_5ofTICE_table.csv"), header=T)[,-1]))
	  
	  # uTICE
	  species1.UTR_tri_freq_top_bottom_ratios_uTICE_table <- as.numeric(unlist(read.csv(file=paste0("tables/tri_freq_tables/", species1, connector1, ".UTR_tri_freq_top_bottom_ratios_uTICE_table.csv"), header=T)[,-1]))
	  species2.UTR_tri_freq_top_bottom_ratios_uTICE_table <- as.numeric(unlist(read.csv(file=paste0("tables/tri_freq_tables/", species2, connector2, ".UTR_tri_freq_top_bottom_ratios_uTICE_table.csv"), header=T)[,-1]))

	  # dTICE
	  species1.CDS_tri_freq_top_bottom_ratios_dTICE_table <- as.numeric(unlist(read.csv(file=paste0("tables/tri_freq_tables/", species1, connector1, ".CDS_tri_freq_top_bottom_ratios_dTICE_table.csv"), header=T)[,-1]))
	  species2.CDS_tri_freq_top_bottom_ratios_dTICE_table <- as.numeric(unlist(read.csv(file=paste0("tables/tri_freq_tables/", species2, connector2, ".CDS_tri_freq_top_bottom_ratios_dTICE_table.csv"), header=T)[,-1]))

	  # 3' of TICE
	  species1.CDS_tri_freq_top_bottom_ratios_3ofTICE_table <- as.numeric(unlist(read.csv(file=paste0("tables/tri_freq_tables/", species1, connector1, ".CDS_tri_freq_top_bottom_ratios_3ofTICE_table.csv"), header=T)[,-1]))
	  species2.CDS_tri_freq_top_bottom_ratios_3ofTICE_table <- as.numeric(unlist(read.csv(file=paste0("tables/tri_freq_tables/", species2, connector2, ".CDS_tri_freq_top_bottom_ratios_3ofTICE_table.csv"), header=T)[,-1]))
	  
	  rownames_btw_species_cors <- c(
	    rownames_btw_species_cors,
	    paste0(species1, connector1, " vs. ", species2, connector2)
	  )
	  
	  btw_species_cors <- rbind(
	    btw_species_cors,
	    c(
	      cor(species1.UTR_tri_freq_top_bottom_ratios_5ofTICE_table, species2.UTR_tri_freq_top_bottom_ratios_5ofTICE_table),
	      cor(species1.UTR_tri_freq_top_bottom_ratios_uTICE_table, species2.UTR_tri_freq_top_bottom_ratios_uTICE_table),
	      cor(species1.CDS_tri_freq_top_bottom_ratios_dTICE_table, species2.CDS_tri_freq_top_bottom_ratios_dTICE_table),
	      cor(species1.CDS_tri_freq_top_bottom_ratios_3ofTICE_table, species2.CDS_tri_freq_top_bottom_ratios_3ofTICE_table)
	    )
	  )
  }
}
rownames(btw_species_cors) <- rownames_btw_species_cors
colnames(btw_species_cors) <- colnames_btw_species_cors

# make the between-species correlation heatmap
heatmap.2(btw_species_cors,
  main = "Cor of trinucleotide\nratios between species", # heat map title
  cellnote = round(btw_species_cors, 2),  # same data set for cell labels
  notecol="black",      # change font color of cell labels to black
  density.info="none",  # turns off density plot inside color legend
  trace="none",         # turns off trace lines inside the heat map
  margins =c(15,15),     # widens margins around plot
  col=my_palette,       # use on color palette defined earlier
  breaks=col_breaks,    # enable color transition at specified limits
  dendrogram="none",    # only draw a row dendrogram
  Rowv="NA",            # turn off row clustering
  Colv="NA"            # turn off column clustering
)
```

# Figure 12. Collinearity analysis: 5'biochem (i.e. uAUG and RNAfold) and 5'motif (i.e. 5'ofTICE+uTICE+dTICE)
```{r}
for (species in names(species_tissue_list)) {
  for (species_tissue in species_tissue_list[[species]]) {
    if (species_tissue!="") {
			tissue_dot <- paste0(".", species_tissue)
			tissue_us <- paste0("_", species_tissue)
		} else {
			tissue_dot <- ""
			tissue_us <- ""
		}
    
    features_uAUG <- get(paste0(species, tissue_dot, ".uAUG_counts.div"))
    
    features_RNAfold <- get(paste0(species, tissue_dot, ".fold_energy_features_selected"))
    
    features_5biochem <- cbind(
      features_uAUG,
      features_RNAfold
    )
    
    features_5motif <- cbind(
      get(paste0(species, tissue_dot, ".5ofTICE_features"))[[1]],
      get(paste0(species, tissue_dot, ".uTICE_features"))[[1]], 
      get(paste0(species, tissue_dot, ".dTICE_features"))[[1]]
    )
    
    # uAUG vs. 5motif
    R2_uAUG <- summary(lm(get(paste0(species, tissue_dot, ".log10.TR")) ~ as.matrix(features_uAUG)))$r.squared
    R2_5motif <- summary(lm(get(paste0(species, tissue_dot, ".log10.TR")) ~ as.matrix(features_5motif)))$r.squared
    R2_uAUG_5motif <- summary(lm(get(paste0(species, tissue_dot, ".log10.TR")) ~ as.matrix(cbind(features_uAUG, features_5motif))))$r.squared
    
    cat(paste0(species, tissue_dot, "\nR2 of uAUG+5motif: "))
    cat(R2_uAUG_5motif)
    cat("\n")
    cat("% of uAUG: ")
    cat(round(R2_uAUG/R2_uAUG_5motif * 100, 3))
    cat("%\n")
    cat("% of 5motif: ")
    cat(round(R2_5motif/R2_uAUG_5motif * 100, 3))
    cat("%\n")
    cat("% of collinearity: ")
    cat(round((R2_uAUG+R2_5motif-R2_uAUG_5motif)/R2_uAUG_5motif * 100, 3))
    cat("%\n\n")
    
    # RNAfold vs. 5motif
    R2_RNAfold <- summary(lm(get(paste0(species, tissue_dot, ".log10.TR")) ~ as.matrix(features_RNAfold)))$r.squared
    R2_RNAfold_5motif <- summary(lm(get(paste0(species, tissue_dot, ".log10.TR")) ~ as.matrix(cbind(features_RNAfold, features_5motif))))$r.squared
    
    cat("R2 of RNAfold+5motif: ")
    cat(R2_RNAfold_5motif)
    cat("\n")
    cat("% of RNAfold: ")
    cat(round(R2_RNAfold/R2_RNAfold_5motif * 100, 3))
    cat("%\n")
    cat("% of 5motif: ")
    cat(round(R2_5motif/R2_RNAfold_5motif * 100, 3))
    cat("%\n")
    cat("% of collinearity: ")
    cat(round((R2_RNAfold+R2_5motif-R2_RNAfold_5motif)/R2_RNAfold_5motif * 100, 3))
    cat("%\n\n")
    
    # 5biochem vs. 5motif
    R2_5biochem <- summary(lm(get(paste0(species, tissue_dot, ".log10.TR")) ~ as.matrix(features_5biochem)))$r.squared
    R2_5biochem_5motif <- summary(lm(get(paste0(species, tissue_dot, ".log10.TR")) ~ as.matrix(cbind(features_5biochem, features_5motif))))$r.squared
    
    cat("R2 of 5biochem+5motif: ")
    cat(R2_5biochem_5motif)
    cat("\n")
    cat("% of 5biochem: ")
    cat(round(R2_5biochem/R2_5biochem_5motif * 100, 3))
    cat("%\n")
    cat("% of 5motif: ")
    cat(round(R2_5motif/R2_5biochem_5motif * 100, 3))
    cat("%\n")
    cat("% of collinearity: ")
    cat(round((R2_5biochem+R2_5motif-R2_5biochem_5motif)/R2_5biochem_5motif * 100, 3))
    cat("%\n\n")
  }
}
```

# A comment in the text - Collinearity analysis: 5'UTR length and 5'biochem+5'motif
```{r}
for (species in names(species_tissue_list)) {
  for (species_tissue in species_tissue_list[[species]]) {
    if (species_tissue!="") {
			tissue_dot <- paste0(".", species_tissue)
			tissue_us <- paste0("_", species_tissue)
		} else {
			tissue_dot <- ""
			tissue_us <- ""
		}
    
    features_UTRlens <- get(paste0(species, tissue_dot, ".log10.UTR_lens.div"))

    features_uAUG <- get(paste0(species, tissue_dot, ".uAUG_counts.div"))
    
    features_RNAfold <- get(paste0(species, tissue_dot, ".fold_energy_features_selected"))
    
    features_5biochem <- cbind(
      features_uAUG,
      features_RNAfold
    )
    
    features_5motif <- cbind(
      get(paste0(species, tissue_dot, ".5ofTICE_features"))[[1]],
      get(paste0(species, tissue_dot, ".uTICE_features"))[[1]], 
      get(paste0(species, tissue_dot, ".dTICE_features"))[[1]]
    )
    
    features_5biochem_5motif <- cbind(
      features_5biochem,
      features_5motif
    )
    
    # UTR_lens vs. 5biochem+5motif
    R2_UTRlens <- summary(lm(get(paste0(species, tissue_dot, ".log10.TR")) ~ as.matrix(features_UTRlens)))$r.squared
    R2_5biochem_5motif <- summary(lm(get(paste0(species, tissue_dot, ".log10.TR")) ~ as.matrix(features_5biochem_5motif)))$r.squared
    R2_UTRlens_5biochem_5motif <- summary(lm(get(paste0(species, tissue_dot, ".log10.TR")) ~ as.matrix(cbind(features_UTRlens, features_5biochem_5motif))))$r.squared
    
    cat(paste0(species, tissue_dot, "\nR2 of UTRlens+5biochem+5motif: "))
    cat(R2_UTRlens_5biochem_5motif)
    cat("\n")
    cat("% of UTRlens: ")
    cat(round(R2_UTRlens/R2_UTRlens_5biochem_5motif * 100, 3))
    cat("%\n")
    cat("% of 5motif: ")
    cat(round(R2_5biochem_5motif/R2_UTRlens_5biochem_5motif * 100, 3))
    cat("%\n")
    cat("% of collinearity: ")
    cat(round((R2_UTRlens+R2_5biochem_5motif-R2_UTRlens_5biochem_5motif)/R2_UTRlens_5biochem_5motif * 100, 3))
    cat("%\n\n")
  }
}
```

# Figure 13. R2s of six cis-control elements (an excel file to store the R2 values first)
## functions
```{r}
# a function to calculate the R2 for a full model with 8 features (five mega features + three single features)
calc_8feature_full_model_R2 <- function(species.log10.TR, species.uTICE_features, species.dTICE_features, species.5ofTICE_features, species.61codon_features, species.log10.UTR_lens, species.log10.CDS_lens, species.uAUG_counts, species.fold_energy_features_selected, species.polyA_lens, len_breaks, ifplot=F, title="") {
	n <- length(species.log10.UTR_lens)
	UTR_lens <- 10^species.log10.UTR_lens
	
	features <- cbind(species.uTICE_features, species.dTICE_features, species.5ofTICE_features, species.61codon_features, species.log10.CDS_lens, species.uAUG_counts, species.fold_energy_features_selected, species.polyA_lens)
	
	lm_selected <- lm(species.log10.TR ~ as.matrix(features), y=TRUE)
	
	if (ifplot) {
		plot(x=lm_selected$fitted, y=lm_selected$y, xlab="log10 TR (predicted)", ylab="log10 TR", main=paste(title, "R2 =", round(summary(lm_selected)$r.squared, 2)))
	}
	return(summary(lm_selected)$r.squared)
}

# a function to calculate the R2 for a full model with five mega features + two single features (uORFs, CDS length)
calc_5_2_feature_full_model_R2 <- function(species.log10.TR, species.uTICE_features, species.dTICE_features, species.5ofTICE_features, species.61codon_features, species.log10.UTR_lens, species.log10.CDS_lens, species.uAUG_counts, species.fold_energy_features_selected, len_breaks, ifplot=F, title="") {
	n <- length(species.log10.UTR_lens)
	UTR_lens <- 10^species.log10.UTR_lens
	
	features <- cbind(species.uTICE_features, species.dTICE_features, species.5ofTICE_features, species.61codon_features, species.log10.CDS_lens, species.uAUG_counts, species.fold_energy_features_selected)
	
	lm_selected <- lm(species.log10.TR ~ as.matrix(features), y=TRUE)
	
	if (ifplot) {
		plot(x=lm_selected$fitted, y=lm_selected$y, xlab="log10 TR (predicted)", ylab="log10 TR", main=paste(title, "R2 =", round(summary(lm_selected)$r.squared, 2)))
	}
	return(summary(lm_selected)$r.squared)
}
```
## the following code only needs to be run once
```{r, eval = FALSE}
six_element_R2s <- NULL
rownames_six_element_R2s <- NULL
colnames_six_element_R2s <- c("uAUGs",	"RNA fold",	"5' motifs",	"CDS length",	"codon usage",	"CDS RNA fold",	"poly A", "full model 8 features",	"full model 5 features")
for (species in names(species_tissue_list)) {
  for (species_tissue in species_tissue_list[[species]]) {
    if (species_tissue!="") {
			tissue_dot <- paste0(".", species_tissue)
			tissue_us <- paste0("_", species_tissue)
		} else {
			tissue_dot <- ""
			tissue_us <- ""
		}
    rownames_six_element_R2s <- c(rownames_six_element_R2s, paste0(species, tissue_dot))
    tmp_R2s <- NULL
    ## uAUG_counts
    tmp_R2s <- c(tmp_R2s,
                 summary(lm(get(paste0(species, tissue_dot, ".log10.TR")) ~ get(paste0(species, tissue_dot, ".uAUG_counts.div"))))$r.squared
                 )
    ## selected UTR folding energy features
    tmp_R2s <- c(tmp_R2s,
                 summary(lm(get(paste0(species, tissue_dot, ".log10.TR")) ~ as.matrix(get(paste0(species, tissue_dot, ".fold_energy_features_selected")))))$r.squared
                 )
    ## 5' motifs (uTICE + dTICE + 5' of TICE features)
    TICE_features <- cbind(get(paste0(species, tissue_dot, ".uTICE_features"))[[1]], get(paste0(species, tissue_dot, ".dTICE_features"))[[1]], get(paste0(species, tissue_dot, ".5ofTICE_features"))[[1]])
    tmp_R2s <- c(tmp_R2s,
                 summary(lm(get(paste0(species, tissue_dot, ".log10.TR")) ~ as.matrix(TICE_features)))$r.squared
                 )
    ## log10.CDS_lens
    tmp_R2s <- c(tmp_R2s,
                 summary(lm(get(paste0(species, tissue_dot, ".log10.TR")) ~ get(paste0(species, tissue_dot, ".log10.CDS_lens.div"))))$r.squared
                 )
    ## codon features
    tmp_R2s <- c(tmp_R2s,
                 summary(lm(get(paste0(species, tissue_dot, ".log10.TR")) ~ as.matrix(get(paste0(species, tissue_dot, ".codon_features"))[[1]])))$r.squared
                 )
    ## CDS fold energy
    tmp_R2s <- c(tmp_R2s,
                 summary(lm(get(paste0(species, tissue_dot, ".log10.TR")) ~ get(paste0(species, tissue_dot, ".CDS_fold_energy.div"))))$r.squared
                 )
    ## polyA_lens
    tmp_R2s <- c(tmp_R2s,
                 summary(lm(get(paste0(species, tissue_dot, ".log10.TR")) ~ get(paste0(species, tissue_dot, ".polyA_lens.div"))))$r.squared
                 )
    
    ## 8-feature model
    tmp_R2s <- c(tmp_R2s,
                 calc_8feature_full_model_R2(get(paste0(species, tissue_dot, ".log10.TR")), get(paste0(species, tissue_dot, ".uTICE_features"))[[1]], get(paste0(species, tissue_dot, ".dTICE_features"))[[1]], get(paste0(species, tissue_dot, ".5ofTICE_features"))[[1]], get(paste0(species, tissue_dot, ".codon_features"))[[1]], get(paste0(species, tissue_dot, ".log10.UTR_lens.div")), get(paste0(species, tissue_dot, ".log10.CDS_lens.div")), get(paste0(species, tissue_dot, ".uAUG_counts.div")), get(paste0(species, tissue_dot, ".fold_energy_features_selected")), get(paste0(species, tissue_dot, ".polyA_lens.div")), get(paste0(species, tissue_dot, ".len_breaks")), ifplot=F, title=paste(species, species_tissue))
                 )
    
    ## 5-feature model
    tmp_R2s <- c(tmp_R2s,
                 calc_5_2_feature_full_model_R2(get(paste0(species, tissue_dot, ".log10.TR")), get(paste0(species, tissue_dot, ".uTICE_features"))[[1]], get(paste0(species, tissue_dot, ".dTICE_features"))[[1]], get(paste0(species, tissue_dot, ".5ofTICE_features"))[[1]], get(paste0(species, tissue_dot, ".codon_features"))[[1]], get(paste0(species, tissue_dot, ".log10.UTR_lens.div")), get(paste0(species, tissue_dot, ".log10.CDS_lens.div")), get(paste0(species, tissue_dot, ".uAUG_counts.div")), get(paste0(species, tissue_dot, ".fold_energy_features_selected")), get(paste0(species, tissue_dot, ".len_breaks")), ifplot=F, title=paste(species, species_tissue))
                 )
    
    six_element_R2s <- rbind(six_element_R2s, tmp_R2s)
  }
}
rownames(six_element_R2s) <- rownames_six_element_R2s
colnames(six_element_R2s) <- colnames_six_element_R2s

write.csv(six_element_R2s, file="tables/six_element_R2s.csv", quote=F)
```
## plot
```{r Figure13, fig.width=3.5, fig.height=3.5, fig.path='figures/', dev=c('png'), dpi=300, out.width="50%"}
for (species in names(species_tissue_list)) {
	for (i in 1:length(species_tissue_list[[species]])) {
		species_tissue <- species_tissue_list[[species]][i]
		if (species_tissue!="") {
			tissue_dot <- paste0(".", species_tissue)
			tissue_us <- paste0("_", species_tissue)
		} else {
			tissue_dot <- ""
			tissue_us <- ""
		}
		calc_5_2_feature_full_model_R2(get(paste0(species, tissue_dot, ".log10.TR")), get(paste0(species, tissue_dot, ".uTICE_features"))[[1]], get(paste0(species, tissue_dot, ".dTICE_features"))[[1]], get(paste0(species, tissue_dot, ".5ofTICE_features"))[[1]], get(paste0(species, tissue_dot, ".codon_features"))[[1]], get(paste0(species, tissue_dot, ".log10.UTR_lens.div")), get(paste0(species, tissue_dot, ".log10.CDS_lens.div")), get(paste0(species, tissue_dot, ".uAUG_counts.div")), get(paste0(species, tissue_dot, ".fold_energy_features_selected")), get(paste0(species, tissue_dot, ".len_breaks")), ifplot=T, title=paste(species, species_tissue))
	}
}
```

# Figure 14. Collinearity analysis of four features: uAUGs, 5'RNA fold, CDS length, and codon usage
```{r Figure14, fig.width=7, fig.height=7, fig.path='figures/', dev=c('pdf'), out.width="50%"}
## A function to calculate pairwise collinearity of every two features
calc_pariwise_collinearity <- function(features_list, y) {
	p <- length(features_list)
	feature_names <- names(features_list)
	
	uni_feature_R2s <- sapply(1:p, FUN=function(i) {
		tmp_features_all <- cbind(y, features_list[[i]])
		colnames(tmp_features_all)[1] <- "y"
		## remove duplicated column names
		tmp_features_all <- tmp_features_all[, !duplicated(colnames(tmp_features_all))]
		tmp_features_all <- as.data.frame(tmp_features_all)
		## model
		tmp_features_model <- lm(y~., data=tmp_features_all)
		summary(tmp_features_model)$r.squared
	})
	
	feature_idx_combn <- combn(p, 2)
	bi_feature_R2s <- apply(feature_idx_combn, 2, FUN=function(feature_idx) {
		tmp_features_all <- y
		for (i in feature_idx) {
			tmp_features_all <- cbind(tmp_features_all, features_list[[i]])
		}
		colnames(tmp_features_all)[1] <- "y"
		## remove duplicated column names
		tmp_features_all <- tmp_features_all[, !duplicated(colnames(tmp_features_all))]
		tmp_features_all <- as.data.frame(tmp_features_all)
		## model
		tmp_features_model <- lm(y~., data=tmp_features_all)
		summary(tmp_features_model)$r.squared
	})
	
	result <- matrix(NA, p, p)
	colnames(result) <- feature_names
	rownames(result) <- feature_names
	# the order of bi_feature_R2s should correspond to the columns of feature_idx_combn
	for (i in 1:length(bi_feature_R2s)) {
		idx <- feature_idx_combn[,i]
		redundancy <- sum(uni_feature_R2s[idx]) - bi_feature_R2s[i]
		redundancy <- max(0, redundancy)
		result[idx[2], idx[1]] <- max(
		  round(redundancy/uni_feature_R2s[idx[1]] * 100, 3),
		  round(redundancy/uni_feature_R2s[idx[2]] * 100, 3)
		)
  }
  result
}

## A function to make levelplot showing values
myPanel <- function(x, y, z, ...) {
    panel.levelplot(x,y,z,...)
    panel.text(x, y, round(z,2))
}

for (species in names(species_tissue_list)) {
  for (species_tissue in species_tissue_list[[species]]) {
    if (species_tissue!="") {
			tissue_dot <- paste0(".", species_tissue)
			tissue_us <- paste0("_", species_tissue)
		} else {
			tissue_dot <- ""
			tissue_us <- ""
		}
    
		features_list <- list(
			uAUG=get(paste0(species, tissue_dot, ".uAUG_counts.div")),
			RNAfold=get(paste0(species, tissue_dot, ".fold_energy_features_selected")),
			CDSlen=get(paste0(species, tissue_dot, ".log10.CDS_lens.div")),
			CodonUsage=get(paste0(species, tissue_dot, ".codon_features"))[[1]]
		)
		tmp <- calc_pariwise_collinearity(features_list, get(paste0(species, tissue_dot, ".log10.TR")))
		tmp[tmp > 100] <- 100 # truncate the value at 100
		assign(x=paste0(species, tissue_dot, ".pariwise_collinearity"),
			value=tmp[-1, -4]/100
		)
		grid <- setNames(melt(get(paste0(species, tissue_dot, ".pariwise_collinearity"))), c("Feature_1", "Feature_2", "Collinearity"))
		print(
		  levelplot(Collinearity ~ Feature_1*Feature_2, grid, panel = myPanel, scales=list(x=list(rot=90)), main=paste(species, species_tissue), col.regions=rev(heat.colors(120)), cuts=100, at=seq(0,1,0.01))
		)  

		filename <- paste0("tables/pairwise_collinearity/", species, tissue_dot, ".pariwise_collinearity.csv")
		if (!file.exists(filename)) {
		  write.csv(get(paste0(species, tissue_dot, ".pariwise_collinearity"))/100, file=filename, quote=F)
		}
  }
}
```

# Collinearity analysis: (codon frequencies in the 1st half of CDS) and (codon frequencies in the 2nd half of CDS)
```{r}
for (species in names(species_tissue_list)) {
  for (species_tissue in species_tissue_list[[species]]) {
    if (species_tissue!="") {
			tissue_dot <- paste0(".", species_tissue)
			tissue_us <- paste0("_", species_tissue)
		} else {
			tissue_dot <- ""
			tissue_us <- ""
		}
    
    codon_features_1 <- readRDS(paste0("processed_data_w_polyA/codon_features/", species, tissue_dot, ".codon_features_1.rds"))[[1]]
    
    codon_features_2 <- readRDS(paste0("processed_data_w_polyA/codon_features/", species, tissue_dot, ".codon_features_2.rds"))[[1]]
    
    codon_features_1_2 <- cbind(
      codon_features_1,
      codon_features_2
    )
    
    # codon_features_1 vs. codon_features_2
    R2_features_1 <- summary(lm(get(paste0(species, tissue_dot, ".log10.TR")) ~ as.matrix(codon_features_1)))$r.squared
    R2_features_2 <- summary(lm(get(paste0(species, tissue_dot, ".log10.TR")) ~ as.matrix(codon_features_2)))$r.squared
    R2_features_1_2 <- summary(lm(get(paste0(species, tissue_dot, ".log10.TR")) ~ as.matrix(codon_features_1_2)))$r.squared
    
    cat(paste0(species, tissue_dot, "\nR2 of codon freq 1st + 2nd halves: "))
    cat(R2_features_1_2)
    cat("\n")
    cat("% of codon freq 1st half: ")
    cat(round(R2_features_1/R2_features_1_2 * 100, 3))
    cat("%\n")
    cat("% of codon freq 2nd half: ")
    cat(round(R2_features_2/R2_features_1_2 * 100, 3))
    cat("%\n")
    cat("% of collinearity: ")
    cat(round((R2_features_1+R2_features_2-R2_features_1_2)/R2_features_1_2 * 100, 3))
    cat("%\n\n")
  }
}
```

# Figure 15. Collinearity analysis of min fold energy, 2nd min fold energy, and 3rd min fold energy
```{r}
for (species in names(species_tissue_list)) {
  for (species_tissue in species_tissue_list[[species]]) {
    if (species_tissue!="") {
			tissue_dot <- paste0(".", species_tissue)
			tissue_us <- paste0("_", species_tissue)
		} else {
			tissue_dot <- ""
			tissue_us <- ""
		}
    
    # min fold energy vs. 2nd min fold energy
    features_1 <- readRDS(file=paste0("processed_data_w_polyA/min_2ndmin_3rdmin_fold_energy_features/", species, tissue_us, "_min_fold_energy_features.div.rds"))
    
    features_2 <- readRDS(file=paste0("processed_data_w_polyA/min_2ndmin_3rdmin_fold_energy_features/", species, tissue_us, "_second_min_fold_energy_features.div.rds"))
    
    features_1_2 <- cbind(
      features_1,
      features_2
    )    
    
    nonNA_idx <- which(complete.cases(features_1_2))
    
    R2_features_1 <- summary(lm(get(paste0(species, tissue_dot, ".log10.TR"))[nonNA_idx] ~ as.matrix(features_1[nonNA_idx,])))$r.squared
    R2_features_2 <- summary(lm(get(paste0(species, tissue_dot, ".log10.TR"))[nonNA_idx] ~ as.matrix(features_2[nonNA_idx,])))$r.squared
    R2_features_1_2 <- summary(lm(get(paste0(species, tissue_dot, ".log10.TR"))[nonNA_idx] ~ as.matrix(features_1_2[nonNA_idx,])))$r.squared
    
    cat(paste0(species, tissue_dot, "\nR2 of min fold energy + 2nd min fold energy: "))
    cat(R2_features_1_2)
    cat("\n")
    cat("% of min fold energy: ")
    cat(round(R2_features_1/R2_features_1_2 * 100, 3))
    cat("%\n")
    cat("% of 2nd min fold energy: ")
    cat(round(R2_features_2/R2_features_1_2 * 100, 3))
    cat("%\n")
    cat("% of collinearity: ")
    cat(round((R2_features_1+R2_features_2-R2_features_1_2)/R2_features_1_2 * 100, 3))
    cat("%\n")
    cat("# of genes: ")
    cat(length(nonNA_idx))
    cat("\n\n")
	
    # min fold energy vs. 3rd min fold energy
    features_1 <- readRDS(file=paste0("processed_data_w_polyA/min_2ndmin_3rdmin_fold_energy_features/", species, tissue_us, "_min_fold_energy_features.div.rds"))
    
    features_2 <- readRDS(file=paste0("processed_data_w_polyA/min_2ndmin_3rdmin_fold_energy_features/", species, tissue_us, "_third_min_fold_energy_features.div.rds"))
    
    features_1_2 <- cbind(
      features_1,
      features_2
    )
    
    nonNA_idx <- which(complete.cases(features_1_2))
    
    R2_features_1 <- summary(lm(get(paste0(species, tissue_dot, ".log10.TR"))[nonNA_idx] ~ as.matrix(features_1[nonNA_idx,])))$r.squared
    R2_features_2 <- summary(lm(get(paste0(species, tissue_dot, ".log10.TR"))[nonNA_idx] ~ as.matrix(features_2[nonNA_idx,])))$r.squared
    R2_features_1_2 <- summary(lm(get(paste0(species, tissue_dot, ".log10.TR"))[nonNA_idx] ~ as.matrix(features_1_2[nonNA_idx,])))$r.squared
    
    cat(paste0("R2 of min fold energy + 3rd min fold energy: "))
    cat(R2_features_1_2)
    cat("\n")
    cat("% of min fold energy: ")
    cat(round(R2_features_1/R2_features_1_2 * 100, 3))
    cat("%\n")
    cat("% of 3rd min fold energy: ")
    cat(round(R2_features_2/R2_features_1_2 * 100, 3))
    cat("%\n")
    cat("% of collinearity: ")
    cat(round((R2_features_1+R2_features_2-R2_features_1_2)/R2_features_1_2 * 100, 3))
    cat("%\n")
    cat("# of genes: ")
    cat(length(nonNA_idx))
    cat("\n\n")
    
    # Pairwise Pearson correlations of min fold energy vs. 2nd min fold energy vs. 3rd min fold energy
    min_2ndmin_3rdmin_fold_energy_features <- readRDS(file=paste0("processed_data_w_polyA/min_2ndmin_3rdmin_fold_energy_features/", species, tissue_us, "_min_2ndmin_3rdmin_fold_energy_features"))[nonNA_idx,]
    cat(paste0("Pearson cor of min fold energy vs. 2nd min fold energy: "))
    cat(cor(min_2ndmin_3rdmin_fold_energy_features[,1], min_2ndmin_3rdmin_fold_energy_features[,2]))
    cat("\n")
    cat(paste0("Pearson cor of min fold energy vs. 3rd min fold energy: "))
    cat(cor(min_2ndmin_3rdmin_fold_energy_features[,1], min_2ndmin_3rdmin_fold_energy_features[,3]))
    cat("\n")
    cat(paste0("Pearson cor of 2nd min fold energy vs. 3rd min fold energy: "))
    cat(cor(min_2ndmin_3rdmin_fold_energy_features[,2], min_2ndmin_3rdmin_fold_energy_features[,3]))
    cat("\n\n")
  }
}
```

# Figure 16A. Collinearity analysis of aa freqs, codon freqs, and synonymous codon ratios
```{r Figure16A, fig.width=7, fig.height=7, fig.path='figures/', dev=c('pdf'), out.width="50%"}
# collinearity between codon freqs, aa freqs, and syn codon ratios
for (species in names(species_tissue_list)) {
	for (i in 1:length(species_tissue_list[[species]])) {
		species_tissue <- species_tissue_list[[species]][i]
		if (species_tissue!="") {
			tissue_dot <- paste0(".", species_tissue)
			tissue_us <- paste0("_", species_tissue)
		} else {
			tissue_dot <- ""
			tissue_us <- ""
		}
		
		aa_features <- readRDS(paste0("processed_data_w_polyA/aa_features/", species, tissue_dot, ".aa_features.rds"))[[1]]
		codon_features <- readRDS(paste0("processed_data_w_polyA/codon_features/", species, tissue_dot, ".codon_features.rds"))[[1]]
		syncodon_features <- readRDS(paste0("processed_data_w_polyA/syncodon_features/", species, tissue_dot, ".syncodon_features.rds"))[[1]]
		
		keep_idx <- which(apply(syncodon_features, 1, FUN=function(x) !(any(x==Inf | is.na(x)))))
		
		log10.CDS_lens <- get(paste0(species, tissue_dot, ".log10.CDS_lens"))
		
		cat(paste0(species, tissue_dot, "\n"))
		cat(paste0("# genes with non-Inf syn codon ratios: ", length(keep_idx), "\n"))
		cat(paste0("median CDS length of included genes: ", median(10^log10.CDS_lens[keep_idx]), "\n"))
		cat(paste0("median CDS length of excluded genes: ", median(10^log10.CDS_lens[-keep_idx]), "\n"))
		
		aa_syncodon_features <- cbind(aa_features, syncodon_features)
		codon_syncodon_features <- cbind(codon_features, syncodon_features)
		
		R2_aa_features <- summary(lm(get(paste0(species, tissue_dot, ".log10.TR"))[keep_idx] ~ as.matrix(aa_features[keep_idx,])))$r.squared
		R2_syncodon_features <- summary(lm(get(paste0(species, tissue_dot, ".log10.TR"))[keep_idx] ~ as.matrix(syncodon_features[keep_idx,])))$r.squared
		R2_aa_syncodon_features <- summary(lm(get(paste0(species, tissue_dot, ".log10.TR"))[keep_idx] ~ as.matrix(aa_syncodon_features[keep_idx,])))$r.squared
		
		cat(paste0("R2 of aa_features + syncodon_features: "))
		cat(R2_aa_syncodon_features)
		cat("\n")
		cat("% of aa_features: ")
		cat(round(R2_aa_features/R2_aa_syncodon_features * 100, 3))
		cat("%\n")
		cat("% of syncodon_features: ")
		cat(round(R2_syncodon_features/R2_aa_syncodon_features * 100, 3))
		cat("%\n")
		cat("% of collinearity: ")
		cat(round((R2_aa_features+R2_syncodon_features-R2_aa_syncodon_features)/R2_aa_syncodon_features * 100, 3))
		cat("%\n\n")
		
		R2_codon_features <- summary(lm(get(paste0(species, tissue_dot, ".log10.TR"))[keep_idx] ~ as.matrix(codon_features[keep_idx,])))$r.squared
		R2_codon_syncodon_features <- summary(lm(get(paste0(species, tissue_dot, ".log10.TR"))[keep_idx] ~ as.matrix(codon_syncodon_features[keep_idx,])))$r.squared
		
		cat(paste0("R2 of codon_features + syncodon_features: "))
		cat(R2_codon_syncodon_features)
		cat("\n")
		cat("% of codon_features: ")
		cat(round(R2_codon_features/R2_codon_syncodon_features * 100, 3))
		cat("%\n")
		cat("% of syncodon_features: ")
		cat(round(R2_syncodon_features/R2_codon_syncodon_features * 100, 3))
		cat("%\n")
		cat("% of collinearity: ")
		cat(round((R2_codon_features+R2_syncodon_features-R2_codon_syncodon_features)/R2_codon_syncodon_features * 100, 3))
		cat("%\n\n")		
	}
}
```

```{r, eval=FALSE}
# calculate the correlation between each syn codon's ratio and its corresponding AA's freq
for (species in names(species_tissue_list)) {
	for (i in 1:length(species_tissue_list[[species]])) {
		species_tissue <- species_tissue_list[[species]][i]
		if (species_tissue!="") {
			tissue_dot <- paste0(".", species_tissue)
			tissue_us <- paste0("_", species_tissue)
		} else {
			tissue_dot <- ""
			tissue_us <- ""
		}
		## count the frequency of 64 codons (including 3 stop codons) in every gene
		CDS_seqs <- get(paste0(species, tissue_dot, ".CDS_seqs"))
		codon_counts <- t( sapply(CDS_seqs, FUN=function(seq) {
		    sst <- strsplit(seq, "")[[1]]
		    n <- length(sst)
		    out <- paste0(sst[seq(from=1, to=n, by=3)], sst[seq(from=2, to=n, by=3)], sst[seq(from=3, to=n, by=3)])
		    table(out)[tri_nts]
		}) )
		codon_counts[is.na(codon_counts)] <- 0
		codon_freqs <- codon_counts / rowSums(codon_counts)
		rownames(codon_freqs) <- names(CDS_seqs)
		colnames(codon_freqs) <- tri_nts
		
		## count the frequency of 20 amino acids (AAs) in every gene
		aa_counts <- sapply(aas, FUN=function(aa) {
		    idx <- which(tri_nts_to_aas==aa)
		    if (length(idx)==1) {
		        codon_counts[,idx]
		    } else {
		        rowSums(codon_counts[,idx])
		    }   
		})
		aa_freqs <- aa_counts / rowSums(aa_counts)
		rownames(aa_freqs) <- names(CDS_seqs)
		colnames(aa_freqs) <- aas
		
		## normalize the codon counts by their corresponding AA count
		syncodon_ratios <- codon_freqs[,nonstop_codons]
		for (aa in aas) {
			tmp_codons <- names(tri_nts_to_aas[which(tri_nts_to_aas==aa)])
			syncodon_ratios[,tmp_codons] <- codon_freqs[,tmp_codons] / rowSums(codon_freqs[,tmp_codons, drop=F])
		}		
		
		cor_codon_aa <- data.frame(
			codon = nonstop_codons,
			aa = tri_nts_to_aas[nonstop_codons],
			cor = sapply(nonstop_codons, FUN=function(codon) {
			  x <- syncodon_ratios[,codon]
			  y <- aa_freqs[,tri_nts_to_aas[codon]]
			  keep_idx <- which(x!=Inf & !is.na(x))
				cor(x[keep_idx], y[keep_idx])
			})
		)
		
		cat(paste0("\n", species, tissue_dot, "\n"))
		cat(paste0("# genes with non-Inf syn codon ratios: ", sum(apply(syncodon_ratios, 1, FUN=function(z) !any(z==Inf | is.na(z)))), "\n"))
		print(cor_codon_aa)	
	}
}
```

# Figure 16B. Collinearity analysis of four features: aa freqs (or codon freqs, or synonymous codon ratios), CDS length, 5'RNA fold, and uAUG
```{r Figure16B, fig.width=7, fig.height=7, fig.path='figures/', dev=c('pdf'), out.width="50%"}
## A function to calculate pairwise collinearity of every two features
calc_syncodon_vs_features_collinearity <- function(features_list, syncodon_features, imp_feature_name, y) {
	p <- length(features_list)
	feature_names <- names(features_list)
	
	syncodon_R2 <- summary(lm(y ~ as.matrix(syncodon_features)))$r.squared
	  
	uni_feature_R2s <- sapply(1:p, FUN=function(i) {
		tmp_features_all <- cbind(y, features_list[[i]])
		colnames(tmp_features_all)[1] <- "y"
		## remove duplicated column names
		tmp_features_all <- tmp_features_all[, !duplicated(colnames(tmp_features_all))]
		tmp_features_all <- as.data.frame(tmp_features_all)
		## model
		tmp_features_model <- lm(y~., data=tmp_features_all)
		summary(tmp_features_model)$r.squared
	})
	names(uni_feature_R2s) <- feature_names
	
	bi_feature_R2s <- sapply(1:p, FUN=function(i) {
		tmp_features_all <- cbind(y, features_list[[i]], syncodon_features)
		colnames(tmp_features_all)[1] <- "y"
		## remove duplicated column names
		tmp_features_all <- tmp_features_all[, !duplicated(colnames(tmp_features_all))]
		tmp_features_all <- as.data.frame(tmp_features_all)
		## model
		tmp_features_model <- lm(y~., data=tmp_features_all)
		summary(tmp_features_model)$r.squared
	})
	names(bi_feature_R2s) <- paste(imp_feature_name, feature_names, sep="-")
	
	result <- matrix(NA, p, 1)
	rownames(result) <- feature_names
	# the order of bi_feature_R2s should correspond to the columns of feature_idx_combn
	for (i in 1:p) {
		redundancy <- syncodon_R2 + uni_feature_R2s[i] - bi_feature_R2s[i]
		redundancy <- max(0, redundancy)
		result[i, 1] <- max(
		  round(redundancy/syncodon_R2 * 100, 3),
		  round(redundancy/uni_feature_R2s[i] * 100, 3)
		)
	}
	
	cat(paste0("R2 of ", imp_feature_name, ": "))
	cat(syncodon_R2)
	cat("\n")
	for (i in 1:p) {
	  feature_name <- feature_names[i]
	  cat(paste0("R2 of ", feature_name, ": "))
	  cat(uni_feature_R2s[i])
	  cat("\n")
	  cat(paste0("R2 of ", imp_feature_name, " + ", feature_name, ": "))
	  cat(bi_feature_R2s[i])
	  cat("\n")
	}
	cat("\n")
  result
}

## A function to make levelplot showing values
myPanel <- function(x, y, z, ...) {
    panel.levelplot(x,y,z,...)
    panel.text(x, y, round(z,2))
}

for (species in names(species_tissue_list)) {
  for (species_tissue in species_tissue_list[[species]]) {
    if (species_tissue!="") {
			tissue_dot <- paste0(".", species_tissue)
			tissue_us <- paste0("_", species_tissue)
		} else {
			tissue_dot <- ""
			tissue_us <- ""
		}
    
    aa_features <- readRDS(paste0("processed_data_w_polyA/aa_features/", species, tissue_dot, ".aa_features.rds"))[[1]]
    codon_features <- readRDS(paste0("processed_data_w_polyA/codon_features/", species, tissue_dot, ".codon_features.rds"))[[1]]
		syncodon_features <- readRDS(paste0("processed_data_w_polyA/syncodon_features/", species, tissue_dot, ".syncodon_features.rds"))[[1]]
		
		# only keep the genes whose syncodon features are all non-Inf and non-NA
		keep_idx <- which(apply(syncodon_features, 1, FUN=function(x) !(any(x==Inf | is.na(x)))))
		# keep_idx <- 1:nrow(syncodon_features) # all genes
		# keep_idx <- which(apply(syncodon_features, 1, FUN=function(x) any(x==Inf | is.na(x)))) # removed genes
		
		aa_features <- aa_features[keep_idx,]
		codon_features <- codon_features[keep_idx,]
		syncodon_features <- syncodon_features[keep_idx,]
		features_list <- list(
			uAUG=get(paste0(species, tissue_dot, ".uAUG_counts.div"))[keep_idx,],
			RNAfold=get(paste0(species, tissue_dot, ".fold_energy_features_selected"))[keep_idx,],
			CDSlen=get(paste0(species, tissue_dot, ".log10.CDS_lens.div"))[keep_idx,]
		)
		
		cat(paste0(species, tissue_dot, "\n"))
		
		aa_tmp <- calc_syncodon_vs_features_collinearity(features_list, aa_features, "aa freqs", get(paste0(species, tissue_dot, ".log10.TR"))[keep_idx])
		
		colnames(aa_tmp) <- "AA freqs"
		aa_tmp[aa_tmp > 100] <- 100 # truncate the value at 100
		assign(x=paste0(species, tissue_dot, ".aa_collinearity"),
			value=aa_tmp/100
		)
		
		codon_tmp <- calc_syncodon_vs_features_collinearity(features_list, codon_features, "codon freqs", get(paste0(species, tissue_dot, ".log10.TR"))[keep_idx])
		colnames(codon_tmp) <- "codon ratios"
		codon_tmp[codon_tmp > 100] <- 100 # truncate the value at 100
		assign(x=paste0(species, tissue_dot, ".codon_collinearity"),
			value=codon_tmp/100
		)
		
		syncodon_tmp <- calc_syncodon_vs_features_collinearity(features_list, syncodon_features, "syncodon ratios", get(paste0(species, tissue_dot, ".log10.TR"))[keep_idx])
		colnames(syncodon_tmp) <- "syncodon ratios"
		syncodon_tmp[syncodon_tmp > 100] <- 100 # truncate the value at 100
		assign(x=paste0(species, tissue_dot, ".syncodon_collinearity"),
			value=syncodon_tmp/100
		)
		
		aa_grid <- setNames(melt(get(paste0(species, tissue_dot, ".aa_collinearity"))), c("Feature_1", "Feature_2", "Collinearity"))
		print(
		  levelplot(Collinearity ~ Feature_1*Feature_2, aa_grid, panel = myPanel, scales=list(x=list(rot=90)), main=paste(species, species_tissue, "aa freqs"), col.regions=rev(heat.colors(120)), cuts=100, at=seq(0,1,0.01))
		)  

		codon_grid <- setNames(melt(get(paste0(species, tissue_dot, ".codon_collinearity"))), c("Feature_1", "Feature_2", "Collinearity"))
		print(
		  levelplot(Collinearity ~ Feature_1*Feature_2, codon_grid, panel = myPanel, scales=list(x=list(rot=90)), main=paste(species, species_tissue, "codon ratios"), col.regions=rev(heat.colors(120)), cuts=100, at=seq(0,1,0.01))
		)
		
		syncodon_grid <- setNames(melt(get(paste0(species, tissue_dot, ".syncodon_collinearity"))), c("Feature_1", "Feature_2", "Collinearity"))
		print(
		  levelplot(Collinearity ~ Feature_1*Feature_2, syncodon_grid, panel = myPanel, scales=list(x=list(rot=90)), main=paste(species, species_tissue, "syncodon ratios"), col.regions=rev(heat.colors(120)), cuts=100, at=seq(0,1,0.01))
		)

		filename <- paste0("tables/pairwise_collinearity/", species, tissue_dot, ".aa_collinearity.csv")
		if (!file.exists(filename)) {
		  write.csv(get(paste0(species, tissue_dot, ".aa_collinearity"))/100, file=filename, quote=F)
		}
		
		filename <- paste0("tables/pairwise_collinearity/", species, tissue_dot, ".codon_collinearity.csv")
		if (!file.exists(filename)) {
		  write.csv(get(paste0(species, tissue_dot, ".codon_collinearity"))/100, file=filename, quote=F)
		}
		
		filename <- paste0("tables/pairwise_collinearity/", species, tissue_dot, ".syncodon_collinearity.csv")
		if (!file.exists(filename)) {
		  write.csv(get(paste0(species, tissue_dot, ".syncodon_collinearity"))/100, file=filename, quote=F)
		}
  }
}
```

# Figure 17. Collinearity analysis of three features: uAUGs, 5'RNA fold, and CDS length, with AA usage and synonymous codon usage
```{r}
## A function to calculate collinearity between aa usage (or synonymous codon usage) and another feature
calc_aa_syncodon_collinearity <- function(features_list, aa_features, codon_features, y) {
	p <- length(features_list)
	feature_names <- names(features_list)
	
	uni_feature_R2s <- sapply(1:p, FUN=function(i) {
		tmp_features_all <- cbind(y, features_list[[i]])
		colnames(tmp_features_all)[1] <- "y"
		## remove duplicated column names
		tmp_features_all <- tmp_features_all[, !duplicated(colnames(tmp_features_all))]
		tmp_features_all <- as.data.frame(tmp_features_all)
		## model
		tmp_features_model <- lm(y~., data=tmp_features_all)
		summary(tmp_features_model)$r.squared
	})
	
	aa_features_R2 <- summary(lm(y~as.matrix(aa_features)))$r.squared
	codon_features_R2 <- summary(lm(y~as.matrix(codon_features)))$r.squared
	syncodon_features_R2 <- codon_features_R2 - aa_features_R2
	
	bi_feature_R2s <- sapply(1:p, FUN=function(i) {
		tmp_features_all <- cbind(y, features_list[[i]], aa_features)
		colnames(tmp_features_all)[1] <- "y"
		## remove duplicated column names
		tmp_features_all <- tmp_features_all[, !duplicated(colnames(tmp_features_all))]
		tmp_features_all <- as.data.frame(tmp_features_all)
		## model
		tmp_features_model <- lm(y~., data=tmp_features_all)
		aa_bi_feature_R2 <- summary(tmp_features_model)$r.squared
		
		tmp_features_all <- cbind(y, features_list[[i]], codon_features)
		colnames(tmp_features_all)[1] <- "y"
		## remove duplicated column names
		tmp_features_all <- tmp_features_all[, !duplicated(colnames(tmp_features_all))]
		tmp_features_all <- as.data.frame(tmp_features_all)
		## model
		tmp_features_model <- lm(y~., data=tmp_features_all)
		codon_bi_feature_R2 <- summary(tmp_features_model)$r.squared
		
		return(c(aa_bi_feature_R2, codon_bi_feature_R2-aa_bi_feature_R2))
	})
	
	aa_result <- rep(NA, p)
	names(aa_result) <- feature_names
	
	syncodon_result <- rep(NA, p)
	names(syncodon_result) <- feature_names
	
	# the order of bi_feature_R2s should correspond to feature_names
	for (i in 1:p) {
		aa_redundancy <- (uni_feature_R2s[i] + aa_features_R2) - bi_feature_R2s[1,i]
		aa_result[i] <- max(
		  round(aa_redundancy/uni_feature_R2s[i] * 100, 3),
		  round(aa_redundancy/aa_features_R2 * 100, 3)
		)
		
		syncodon_redundancy <- (uni_feature_R2s[i] + syncodon_features_R2) - bi_feature_R2s[2,i]
		syncodon_result[i] <- max(
		  round(syncodon_redundancy/uni_feature_R2s[i] * 100, 3),
		  round(syncodon_redundancy/syncodon_features_R2 * 100, 3)
		)
	}
	return(list(aa_result, syncodon_result))
}

for (species in names(species_tissue_list)) {
  for (species_tissue in species_tissue_list[[species]]) {
		if (species_tissue!="") {
			tissue_dot <- paste0(".", species_tissue)
			tissue_us <- paste0("_", species_tissue)
		} else {
			tissue_dot <- ""
			tissue_us <- ""
		}
    
		features_list <- list(
			uAUG=get(paste0(species, tissue_dot, ".uAUG_counts.div")),
			RNAfold=get(paste0(species, tissue_dot, ".fold_energy_features_selected")),
			CDSlen=get(paste0(species, tissue_dot, ".log10.CDS_lens.div"))
		)
		aa_features <- get(paste0(species, tissue_dot, ".aa_features"))[[1]]
		codon_features <- get(paste0(species, tissue_dot, ".codon_features"))[[1]]
		tmp <- calc_aa_syncodon_collinearity(features_list, aa_features, codon_features, get(paste0(species, tissue_dot, ".log10.TR")))
		aa_result <- tmp[[1]]
		syncodon_result <- tmp[[2]]
		aa_result[aa_result > 100] <- 100 # truncate the value at 100
		syncodon_result[syncodon_result > 100] <- 100 # truncate the value at 100
		
		aa_result <- aa_result/100
		names(aa_result) <- names(features_list)
		syncodon_result <- syncodon_result/100
		names(syncodon_result) <- names(features_list)
		
		cat(paste0(species, tissue_dot, "\ncollinearity with AA usage: \n"))
		for (i in 1:length(features_list)) {
		  cat(names(features_list)[i])
		  cat("\n")
		  cat(aa_result[i])
		  cat("\n")
		}
		cat("\n")
		
		cat(paste0(species, tissue_dot, "\ncollinearity with synonymous codon usage: \n"))
		for (i in 1:length(features_list)) {
		  cat(names(features_list)[i])
		  cat("\n")
		  cat(syncodon_result[i])
		  cat("\n")
		}
		cat("\n")
  }
}
```

# Figure 18. Analysis of aa freqs / codon freqs / synonymous codon ratios vs. tRNA abundance for sc
```{r Figure18, fig.width=7, fig.height=7, fig.path='figures/', dev=c('pdf'), out.width="50%"}
for (species in "sc") {
	for (i in 1:length(species_tissue_list[[species]])) {
		species_tissue <- species_tissue_list[[species]][i]
		if (species_tissue!="") {
			tissue_dot <- paste0(".", species_tissue)
			tissue_us <- paste0("_", species_tissue)
		} else {
			tissue_dot <- ""
			tissue_us <- ""
		}
		
		aa_freqs <- readRDS(paste0("processed_data_w_polyA/aa_features/", species, tissue_dot, ".aa_freqs.rds"))
		codon_freqs <- readRDS(paste0("processed_data_w_polyA/codon_features/", species, tissue_dot, ".codon_freqs.rds"))
		syncodon_ratios <- readRDS(paste0("processed_data_w_polyA/syncodon_features/", species, tissue_dot, ".syncodon_ratios.rds"))
		
		genes <- get(paste0(species, tissue_dot, ".genes"))
		
		TR_top_genes <- get(paste0(species, tissue_dot, ".TR_top_genes"))
		TR_bottom_genes <- get(paste0(species, tissue_dot, ".TR_bottom_genes"))
		
		keep_idx <- which(apply(syncodon_ratios, 1, FUN=function(x) !(any(x==Inf | is.na(x)))))
		
		aa_freqs <- aa_freqs[keep_idx,]
		codon_freqs <- codon_freqs[keep_idx,]
		syncodon_ratios <- syncodon_ratios[keep_idx,]
		genes <- genes[keep_idx]
		TR_top_genes <- TR_top_genes[TR_top_genes %in% genes]
		TR_bottom_genes <- TR_bottom_genes[TR_bottom_genes %in% genes]
		
		aa_tRNA_abun <- readRDS(paste0("processed_data_w_polyA/tRNA_abun/", species, tissue_dot, ".aa_tRNA_abun.rds"))
		codon_tRNA_abun <- readRDS(paste0("processed_data_w_polyA/tRNA_abun/", species, tissue_dot, ".codon_tRNA_abun.rds"))
		
		TR_top_avg_aa_freqs <- colMeans(aa_freqs[TR_top_genes,])
		TR_bottom_avg_aa_freqs <- colMeans(aa_freqs[TR_bottom_genes,])
		
		TR_top_avg_codon_freqs <- colMeans(codon_freqs[TR_top_genes,])
		TR_bottom_avg_codon_freqs <- colMeans(codon_freqs[TR_bottom_genes,])
		
		TR_top_avg_syncodon_ratios <- colMeans(syncodon_ratios[TR_top_genes,])
		TR_bottom_avg_syncodon_ratios <- colMeans(syncodon_ratios[TR_bottom_genes,])
		
		# output the values
		
		## R2s of tRNA abun ~ AA freqs (TR top vs. bottom)
		cat(paste0(species, tissue_dot, "\n"))
		cat("Top TR AA freqs. vs tRNAs abundaces: ")
		cat(
		  cor(
		    TR_top_avg_aa_freqs,
		    aa_tRNA_abun
		  )^2
		)
		cat("\n")
		cat("Bottom TR AA freqs. vs tRNAs abundaces: ")
		cat(
		  cor(
		    TR_bottom_avg_aa_freqs,
		    aa_tRNA_abun
		  )^2
		)
		cat("\n")
		cat("Test if TR top > TR bottom:\n")
		print(
		  cocor.dep.groups.overlap(
		    r.jk=cor(aa_tRNA_abun, TR_top_avg_aa_freqs), 
		    r.jh=cor(aa_tRNA_abun, TR_bottom_avg_aa_freqs),  
		    r.kh=cor(TR_top_avg_aa_freqs, TR_bottom_avg_aa_freqs), 
		    n=20, 
		    alternative = "greater",
		    var.labels = c("tRNA_abun", "TR_top_aa_freq", "TR_bottom_aa_freq")
		  )
		)
		
		## R2s of tRNA abun ~ codon freqs (TR top vs. bottom)
		cat(paste0(species, tissue_dot, "\n"))
		cat("Top TR codon freqs. vs tRNAs abundaces: ")
		cat(
		  cor(
		    TR_top_avg_codon_freqs,
		    codon_tRNA_abun
		  )^2
		)
		cat("\n")
		cat("Bottom TR codon freqs. vs tRNAs abundaces: ")
		cat(
		  cor(
		    TR_bottom_avg_codon_freqs,
		    codon_tRNA_abun
		  )^2
		)
		cat("\n")
		cat("Test if TR top > TR bottom:\n")
		print(
		  cocor.dep.groups.overlap(
		    r.jk=cor(codon_tRNA_abun, TR_top_avg_codon_freqs), 
		    r.jh=cor(codon_tRNA_abun, TR_bottom_avg_codon_freqs),  
		    r.kh=cor(TR_top_avg_codon_freqs, TR_bottom_avg_codon_freqs), 
		    n=20, 
		    alternative = "greater",
		    var.labels = c("tRNA_abun", "TR_top_aa_freq", "TR_bottom_aa_freq")
		  )
		)
		
		## R2s of tRNA abun ~ syncodon ratios (TR top vs. bottom)
		cat(paste0(species, tissue_dot, "\n"))
		cat("Top TR syncodon ratios. vs tRNAs abundaces: ")
		cat(
		  cor(
		    TR_top_avg_syncodon_ratios,
		    codon_tRNA_abun
		  )^2
		)
		cat("\n")
		cat("Bottom TR syncodon ratios. vs tRNAs abundaces: ")
		cat(
		  cor(
		    TR_bottom_avg_syncodon_ratios,
		    codon_tRNA_abun
		  )^2
		)
		cat("\n")
		cat("Test if TR top > TR bottom:\n")
		print(
		  cocor.dep.groups.overlap(
		    r.jk=cor(codon_tRNA_abun, TR_top_avg_syncodon_ratios), 
		    r.jh=cor(codon_tRNA_abun, TR_bottom_avg_syncodon_ratios),  
		    r.kh=cor(TR_top_avg_syncodon_ratios, TR_bottom_avg_syncodon_ratios), 
		    n=20, 
		    alternative = "greater",
		    var.labels = c("tRNA_abun", "TR_top_aa_freq", "TR_bottom_aa_freq")
		  )
		)
		
		# scatterplots of tRNA abundance vs. TR top/bottom ratios
		TR_top_vs_bottom_avg_aa_freqs_ratios <- TR_top_avg_aa_freqs / TR_bottom_avg_aa_freqs
		TR_top_vs_bottom_avg_codon_freqs_ratios <- TR_top_avg_codon_freqs / TR_bottom_avg_codon_freqs
		TR_top_vs_bottom_avg_syncodon_ratios_ratios <- TR_top_avg_syncodon_ratios / TR_bottom_avg_syncodon_ratios
		
		plot(x=TR_top_vs_bottom_avg_aa_freqs_ratios, 
		     y=aa_tRNA_abun, 
		     xlab="TR top/bottom AA freq. ratios", 
		     ylab="tRNA abundance"
		)
		title(paste("cor =",
            round(cor(x=TR_top_vs_bottom_avg_aa_freqs_ratios, y=aa_tRNA_abun), 2), 
            "p =",
            round(cor.test(TR_top_vs_bottom_avg_aa_freqs_ratios, aa_tRNA_abun, alternative="greater")$p.value*3, 3))
		)
		abline(v=1, lty=2)
		
		plot(x=TR_top_vs_bottom_avg_codon_freqs_ratios, 
		     y=codon_tRNA_abun, 
		     xlab="TR top/bottom codon freq. ratios", 
		     ylab="tRNA abundance"
		)
		title(paste("cor =",
            round(cor(x=TR_top_vs_bottom_avg_codon_freqs_ratios, y=codon_tRNA_abun), 2), 
            "p =",
            round(cor.test(TR_top_vs_bottom_avg_codon_freqs_ratios, codon_tRNA_abun, alternative="greater")$p.value*3, 3))
		)
		abline(v=1, lty=2)
		
		plot(x=TR_top_vs_bottom_avg_syncodon_ratios_ratios, 
		     y=codon_tRNA_abun, 
		     xlab="TR top/bottom syncodon ratio ratios", 
		     ylab="tRNA abundance"
		)
		title(paste("cor =",
            round(cor(x=TR_top_vs_bottom_avg_syncodon_ratios_ratios, y=codon_tRNA_abun), 2), 
            "p =",
            round(cor.test(TR_top_vs_bottom_avg_syncodon_ratios_ratios, codon_tRNA_abun, alternative="greater")$p.value*3, 3))
		)
		abline(v=1, lty=2)
	}
}
```

# Supplementary file 2. Output the processed information to an Excel file, one tab per species & tissue
```{r, eval=FALSE}
# columns
## Gene
## 5' UTR seq
## CDS seq
## log10 mRNA (RPKM)
## log10 TR (RPKM)
## 5' motif score
## 5' UTR RNA fold engy
## 5'UTR #uAUGs
## log10 CDS length
## codon freq.
## AA freq.
## syn.codon preference
## poly A length
## log10 5' UTR length

for (species in names(species_tissue_list)) {
	for (species_tissue in species_tissue_list[[species]]) {
		if (species_tissue!="") {
			tissue_dot <- paste0(".", species_tissue)
			tissue_us <- paste0("_", species_tissue)
		} else {
			tissue_dot <- ""
			tissue_us <- ""
		}
		supp2 <- data.frame(
			Gene = get(paste0(species, tissue_dot, ".genes"))
		)
		supp2$"5' UTR seq" <- get(paste0(species, tissue_dot, ".UTR_seqs"))
		supp2$"CDS seq" <- get(paste0(species, tissue_dot, ".CDS_seqs"))
		supp2$"log10 mRNA (RPKM)" <- get(paste0(species, tissue_dot, ".log10.mRNA"))
		supp2$"log10 TR (RPKM)" <- get(paste0(species, tissue_dot, ".log10.TR"))
		supp2$"5' motif score" <- get(paste0(species, tissue_dot, ".log10.TR"))
		
	    features_5motif <- cbind(
	      get(paste0(species, tissue_dot, ".5ofTICE_features"))[[1]],
	      get(paste0(species, tissue_dot, ".uTICE_features"))[[1]], 
	      get(paste0(species, tissue_dot, ".dTICE_features"))[[1]]
	    )
		supp2$"5' motif score" <- fitted(lm(get(paste0(species, tissue_dot, ".log10.TR")) ~ as.matrix(features_5motif), na.action="na.exclude"))
		
		features_RNAfold <- get(paste0(species, tissue_dot, ".fold_energy_features_selected"))
		supp2$"5' UTR RNA fold engy" <- fitted(lm(get(paste0(species, tissue_dot, ".log10.TR")) ~ as.matrix(features_RNAfold), na.action="na.exclude"))
		
		supp2$"5'UTR #uAUGs" <- get(paste0(species, tissue_dot, ".uAUG_counts"))
		
		features_CDSlen <- get(paste0(species, tissue_dot, ".log10.CDS_lens.div"))
		supp2$"log10 CDS length" <- fitted(lm(get(paste0(species, tissue_dot, ".log10.TR")) ~ as.matrix(features_CDSlen), na.action="na.exclude"))
		
		features_CodonUsage <- get(paste0(species, tissue_dot, ".codon_features"))[[1]]
		supp2$"codon freq." <- fitted(lm(get(paste0(species, tissue_dot, ".log10.TR")) ~ as.matrix(features_CodonUsage), na.action="na.exclude"))
		
		features_AAUsage <- get(paste0(species, tissue_dot, ".aa_features"))[[1]]
		supp2$"AA freq." <- fitted(lm(get(paste0(species, tissue_dot, ".log10.TR")) ~ as.matrix(features_AAUsage), na.action="na.exclude"))
		
		features_SynCodonUsage <- readRDS(paste0("processed_data_w_polyA/syncodon_features/", species, tissue_dot, ".syncodon_features.rds"))[[1]]
		supp2$"syn.codon preference" <- fitted(lm(get(paste0(species, tissue_dot, ".log10.TR")) ~ as.matrix(features_SynCodonUsage), na.action="na.exclude"))
		
		supp2$"poly A length" <- get(paste0(species, tissue_dot, ".polyA_lens"))		
		supp2$"log10 5' UTR length" <- get(paste0(species, tissue_dot, ".log10.UTR_lens"))
		
		write.csv(supp2, file=paste0("supp/SuppFile2.", species, tissue_dot, ".csv"), quote=FALSE, row.names=FALSE)
		rm(supp2)
		gc()
	}
}
```

# Supplementary file 4. Output the RNA secondary structures of the minimum folding energy windows (the most folded windows)
```{r, eval=FALSE}
# columns
## id
## seq
## mfe
## mfe_struct
## max_loop
## max_continuous_stem
## max_stem
## all paired nucleotides
## log TR

for (species in names(species_tissue_list)) {
	for (species_tissue in species_tissue_list[[species]]) {
		if (species_tissue!="") {
			tissue_dot <- paste0(".", species_tissue)
			tissue_us <- paste0("_", species_tissue)
		} else {
			tissue_dot <- ""
			tissue_us <- ""
		}
		
		min_fold_energy_window_features <- read.table(paste0("data/min_window_seq_parameters/", species, tissue_us, ".tsv"), header=T, sep="\t", stringsAsFactors=FALSE)
		rownames(min_fold_energy_window_features) <- min_fold_energy_window_features$id
		
		supp4 <- data.frame(
			id = get(paste0(species, tissue_dot, ".genes"))
		)
		supp4$"seq" <- min_fold_energy_window_features$seq[supp4$id]
		supp4$"mfe" <- min_fold_energy_window_features$mfe[supp4$id]
		supp4$"mfe_struct" <- min_fold_energy_window_features$mfe_struct[supp4$id]
		supp4$"max_loop" <- min_fold_energy_window_features$max_loop[supp4$id]
		supp4$"max_continuous_stem" <- min_fold_energy_window_features$max_continuous_stem[supp4$id]
		supp4$"max_stem" <- min_fold_energy_window_features$max_stem_paired[supp4$id]
		supp4$"all paired nucleotides" <- min_fold_energy_window_features$paired_nts[supp4$id]
		supp4$"log TR" <- get(paste0(species, tissue_dot, ".log10.TR"))
		
		write.csv(supp4, file=paste0("supp/SuppFile4.", species, tissue_dot, ".csv"), quote=FALSE, row.names=FALSE)
	}
}
```

# Supplementary file 5. The frequencies of nucleotides at each nucleotide position of PWMs. i.	The AUG aligned PWMs for the top and bottom cohorts shown in Figure 7. ie from -80 to +35. PWMs for all 10 datasets. ii.	The 5' cap aligned PWMs for the 5' most 50 nucleotides for the top and bottom cohorts shown in Figure 8—figure supplement 1.
```{r, eval=FALSE}
for (species in names(species_exemplar_tissue_list)) {
	for (species_tissue in species_exemplar_tissue_list[[species]]) {
	  if (species_tissue!="") {
			species.log10.TR <- get(paste(species, species_tissue, "log10.TR", sep="."))
			connector <- paste0(species_tissue, ".")
		} else {
			species.log10.TR <- get(paste(species, "log10.TR", sep="."))
			connector <- ""
		}
	  
	  species.TR_top_PWM_AUG_aligned <- readRDS(file=paste0("processed_data_w_polyA/AUGPWM_R2s/", species, ".", connector, "TR_top_PWM_AUG_aligned.rds"))
	  write.csv(species.TR_top_PWM_AUG_aligned[21:nrow(species.TR_top_PWM_AUG_aligned),], file=paste0("supp/SuppFile5.", species, tissue_dot, ".TR_top_PWM_AUG_aligned.csv"), quote=FALSE, row.names=TRUE)
	  
	  species.TR_bottom_PWM_AUG_aligned <- readRDS(file=paste0("processed_data_w_polyA/AUGPWM_R2s/", species, ".", connector, "TR_bottom_PWM_AUG_aligned.rds"))
	  write.csv(species.TR_bottom_PWM_AUG_aligned[21:nrow(species.TR_bottom_PWM_AUG_aligned),], file=paste0("supp/SuppFile5.", species, tissue_dot, ".TR_bottom_PWM_AUG_aligned.csv"), quote=FALSE, row.names=TRUE)
	  
	  species.TR_top_PWM_5_aligned <- readRDS(file=paste0("processed_data_w_polyA/5'PWM_R2s/", species, ".", connector, "TR_top_PWM_5_aligned.rds"))
	  write.csv(species.TR_top_PWM_5_aligned[1:50,], file=paste0("supp/SuppFile5.", species, tissue_dot, ".TR_top_PWM_5_aligned.csv"), quote=FALSE, row.names=TRUE)
	  
	  species.TR_bottom_PWM_5_aligned <- readRDS(file=paste0("processed_data_w_polyA/5'PWM_R2s/", species, ".", connector, "TR_bottom_PWM_5_aligned.rds"))
	  write.csv(species.TR_bottom_PWM_5_aligned[1:50,], file=paste0("supp/SuppFile5.", species, tissue_dot, ".TR_bottom_PWM_5_aligned.csv"), quote=FALSE, row.names=TRUE)
	}
}
```

# Supplementary file 6. Compare the Sc. TR data from Weinberg et al to that of Cuperus et al.
```{r}
# columns
## gene name
## cupress 5' UTR seq
## weinberg 5' UTR seq
## cupress log 10 growth rate
## weinberg log 10 IE

sc.cuperus.data <- read_xlsx("data/Supplementary file 6 Cuperus vs weinberg.xlsx", sheet=1, range="A2:N2259")
sc.cuperus.data <- as.data.frame(sc.cuperus.data)

sc.cuperus.genes <- sc.cuperus.data[,1]
sc.cuperus.log10.TR <- sc.cuperus.data[,2]
names(sc.cuperus.log10.TR) <- sc.cuperus.genes
sc.cuperus.UTR_lens <- sc.cuperus.data$"5' UTR length"
names(sc.cuperus.UTR_lens) <- sc.cuperus.genes
sc.cuperus.UTR_seqs <- sc.cuperus.data$"UTR sequence"
names(sc.cuperus.UTR_seqs) <- sc.cuperus.genes

rm(sc.cuperus.data)

sc.common.genes <- intersect(sc.genes, sc.cuperus.genes)
# check the UTR lengths
sc.UTR_lens <- 10^sc.log10.UTR_lens
idx <- which(abs(sc.UTR_lens[sc.common.genes] - sc.cuperus.UTR_lens[sc.common.genes]) < 2)

cor(sc.log10.TR[sc.common.genes][idx], sc.cuperus.log10.TR[sc.common.genes][idx])

cor(sc.log10.TR[sc.common.genes][idx], sc.cuperus.log10.TR[sc.common.genes][idx])^2

supp6 <- data.frame(
	gene = sc.common.genes[idx]
)
supp6$"cupress 5' UTR seq" <- sc.cuperus.UTR_seqs[sc.common.genes][idx]
supp6$"weinberg 5' UTR seq" <- sc.UTR_seqs[sc.common.genes][idx]
supp6$"cupress log 10 growth rate" <- sc.cuperus.log10.TR[sc.common.genes][idx]
supp6$"weinberg log 10 IE" <- sc.log10.TR[sc.common.genes][idx]

write.csv(supp6, file="supp/SuppFile6.csv", quote=FALSE, row.names=FALSE)
```
````
